# Dozens of genetic variants sustain adaptation to urban spatial heterogeneity in *Arabidopsis thaliana*

**DOI:** 10.64898/2026.06.27.734931

**Authors:** Justine Floret, Anja Linstädter, Huiyao Zhang, Vera Hesen, Swan Portalier, Lee Weinand, Gaelle Bustarret, Kirsten Bell, Fabrice Roux, Tahir Ali, Margarita Takou, Gregor Schmitz, Stanislav Kopriva, Juliette de Meaux

## Abstract

Urbanization creates mosaics of microhabitats that sustain diverse plant communities and offer untapped opportunities to understand contemporary ecological and evolutionary processes. Using a city-wide colonization experiment in Cologne (Germany), we identified the environmental factors limiting establishment and persistence of the ruderal species *Arabidopsis thaliana* and the genetic variants enabling adaptation. We show that standing genetic variation is essential for realizing the species’ urban niche, enabling rapid adaptation to fine-scale gradients in disturbance, vegetation, and soil conditions. Despite originating from only two parental genotypes, populations evolved substantial adaptive differentiation within three generations, revealing a highly polygenic basis of local adaptation. More broadly, our approach establishes cities as powerful open-air laboratories for uncovering and fostering ecological and evolutionary processes that generate and sustain biodiversity in human-dominated landscapes.

## Introduction

Global biodiversity is declining at an unprecedented rate, as a growing human population drives the destruction, fragmentation, and degradation of natural habitats (Cowie et al. 2022). At the same time, urbanization is accelerating worldwide, with cities projected to house more than ten billion people by 2050 (UN-Habitat 2024). Cities are not homogeneous but instead form fine-scale mosaics of habitat sites that differ in microclimate, soil conditions, and disturbance regimes, including parks, roadside verges, and built-up surfaces (Chatzidimitriou and Yannas 2015; Tresch et al. 2018). This spatial heterogeneity creates steep, replicated environmental gradients that shape the structure of species communities, allowing cities to sustain diverse plant communities, despite intense anthropogenic pressure (Aronson et al. 2014; Hahs et al. 2023). If, in addition, cities can host active evolutionary dynamics, they could even become beacons of biodiversity (Szulkin et al. 2020; Boehnke et al. 2022). Studies comparing rural and urban populations of plants and animals often reveal that urban species have adapted to the unique challenges of city life (Santangelo et al. 2022; Thompson et al. 2025). For example, research has identified a correlation between leaf color and intra-urban temperature variations in *Oxalis corniculata* (Fukano et al. 2023). Yet, we do not know how much natural variation is available within cities that can be mobilized to enable rapid adaptation to environmental stressors at the scale of a single city (Ibañez et al. 2017; Wu et al. 2026).

Here, we used an urban landscape as an open-air laboratory to simultaneously identify the environmental variables that set evolutionary pressures on populations of *Arabidopsis thaliana* and the genetic variants that support their evolution after three generations (Fig. 1). Urban populations of *A. thaliana* harbor genetic diversity representative of broad regional gene pools (Bomblies et al. 2010; Schmitz et al. 2024). Although colonization appears largely stochastic, genetic variation is not randomly distributed across microhabitats. In particular, allelic variation associated with germination and flowering time tracks urban disturbance gradients (Schmitz et al. 2024), suggesting that cities may select for different ecotypes across urban environments.

**Figure 1:**
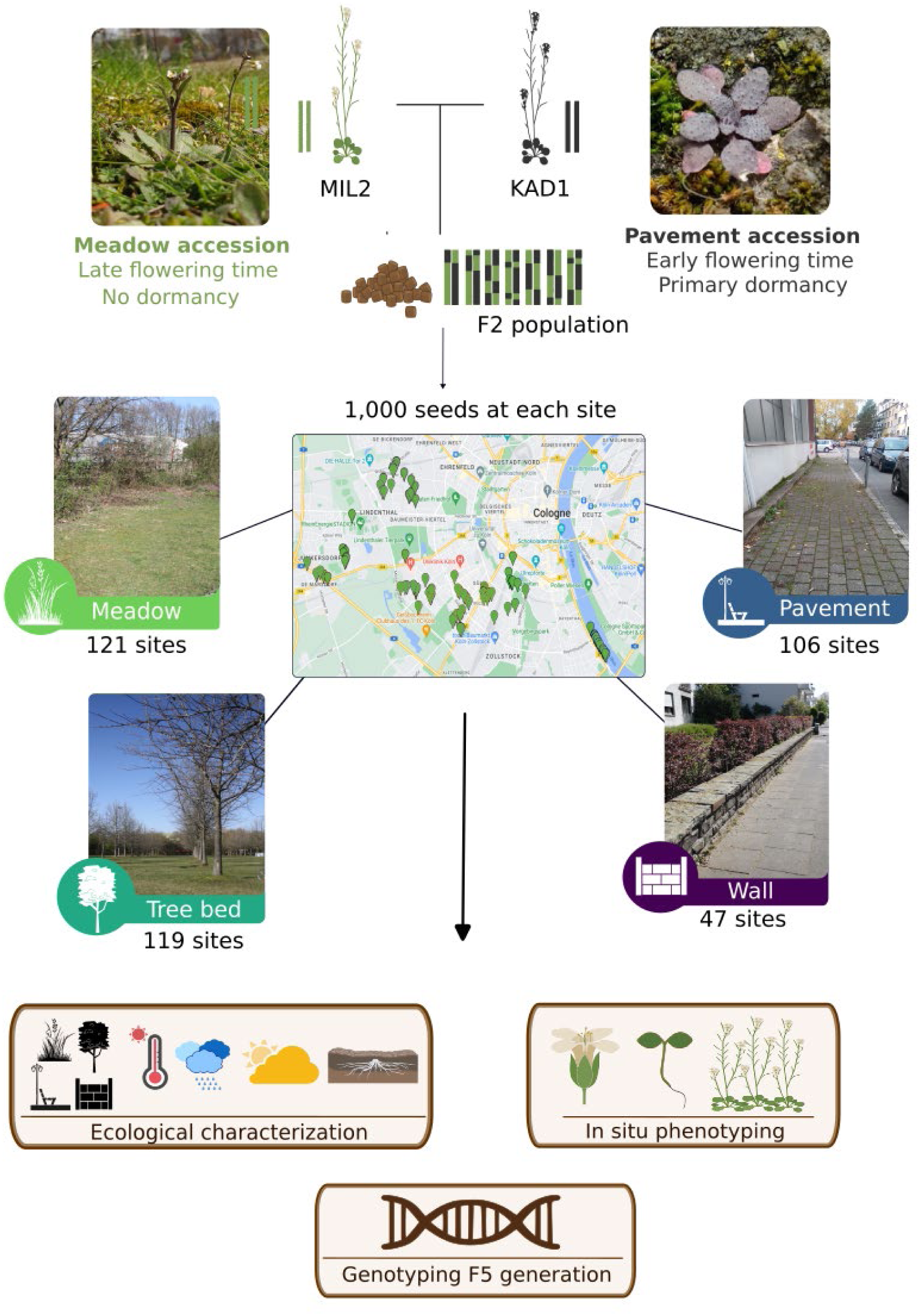
Two genetically different local accessions of *A. thaliana* (MIL2 and KAD1) collected at two contrasting urban microhabitats in Cologne have been crossed to produce an F2 population. In 2021, in each of 393 natural urban sites corresponding to 121 meadows, 119 pavements, 106 tree beds, and 47 walls, 1,000 seeds of the F2 population were sown.

### Urban experimental evolution in a heterogeneous city

To identify the environmental parameters that exert selective pressures limiting population establishment and fostering adaptation, we conducted a city-wide colonization experiment with *A. thaliana* across Cologne, Germany, treating the urban landscape as a set of spatially replicated environmental gradients. Urban habitats were classified *a priori* into four types representing common urban plant habitats that differ in substrate availability, vegetation cover, and disturbance regime: meadows, tree beds, pavements and walls (Fig. 2A). These habitat types span a gradient of increasing human impact (Giuliani et al. 2024a), are themselves variable and harbor spontaneous *A. thaliana* populations (Schmitz et al. 2024). We crossed two previously characterized ecotypes (MIL2 and KAD1), originating from contrasting urban microhabitats in this city – a managed urban meadow and a wall, respectively (Schmitz et al. 2024) – and generated a segregating F2 population. These parental genotypes differed in the genetic architecture of key growth, life-history and resource acquisition traits and capture substantial phenotypic divergence among 12 *A. thaliana* genotypes from Cologne as quantified in common-garden experiments (Schmitz et al. 2024; Fig. S1). We identified 393 sites in Cologne previously unoccupied habitat (121 meadows, 106 tree beds, 119 pavements, and 47 walls; Fig. 1). In fall 2021, we distributed 1,000 F2 seeds per site, forming a population of recombined genomes of the two local urban ecotypes. Establishment was assessed in spring 2022, sites were characterized for abiotic environmental conditions, vegetation characteristics, and disturbance regimes (Fig. S2-S3, Table 1). A second campaign of establishment was started in summer 2022, and population persistence was monitored. In 2025, a total of 68 populations persisted with at least five individuals. These populations were transplanted to the greenhouse, amplified, individually phenotyped in common garden for their flowering time and dormancy levels, and ultimately pool-sequenced (Fig 1, Suppl. Methods section).

**Figure 2:**
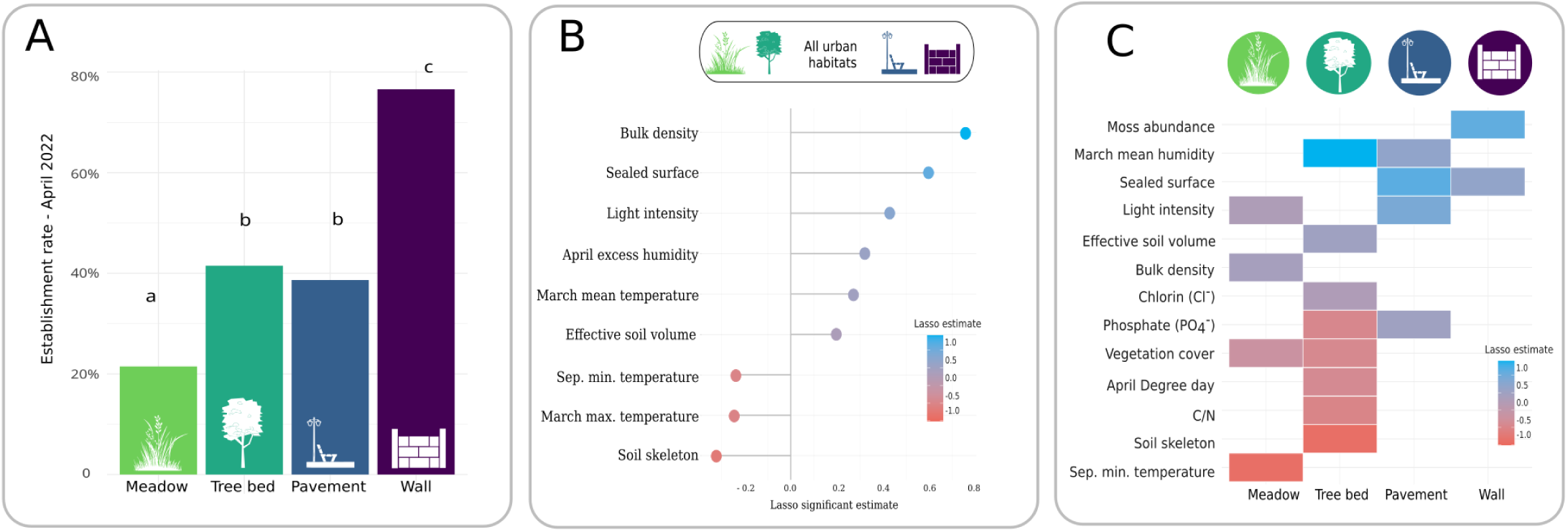
The role of the ecological factors on the establishment of *A. thaliana*’s in natural urban environments. Between April 2021 and June 2023, 33 ecological variables were measured at each site (Table 1). **A.** Frequency of *A. thaliana* population established per habitat type in April 2022. The letters indicate a significant pairwise difference (p <0.05) after Tukey’s comparisons of means. B. Across all urban environments: ecological factors positively (blue) or negatively (red) associated with the establishment of *A. thaliana* in April 2022 (first generation). **C.** Per urban habitat: Ecological factors positively (blue) or negatively (red) associated with the establishment of *A. thaliana* in April 2022 (first generation).

**TABLE 1:**
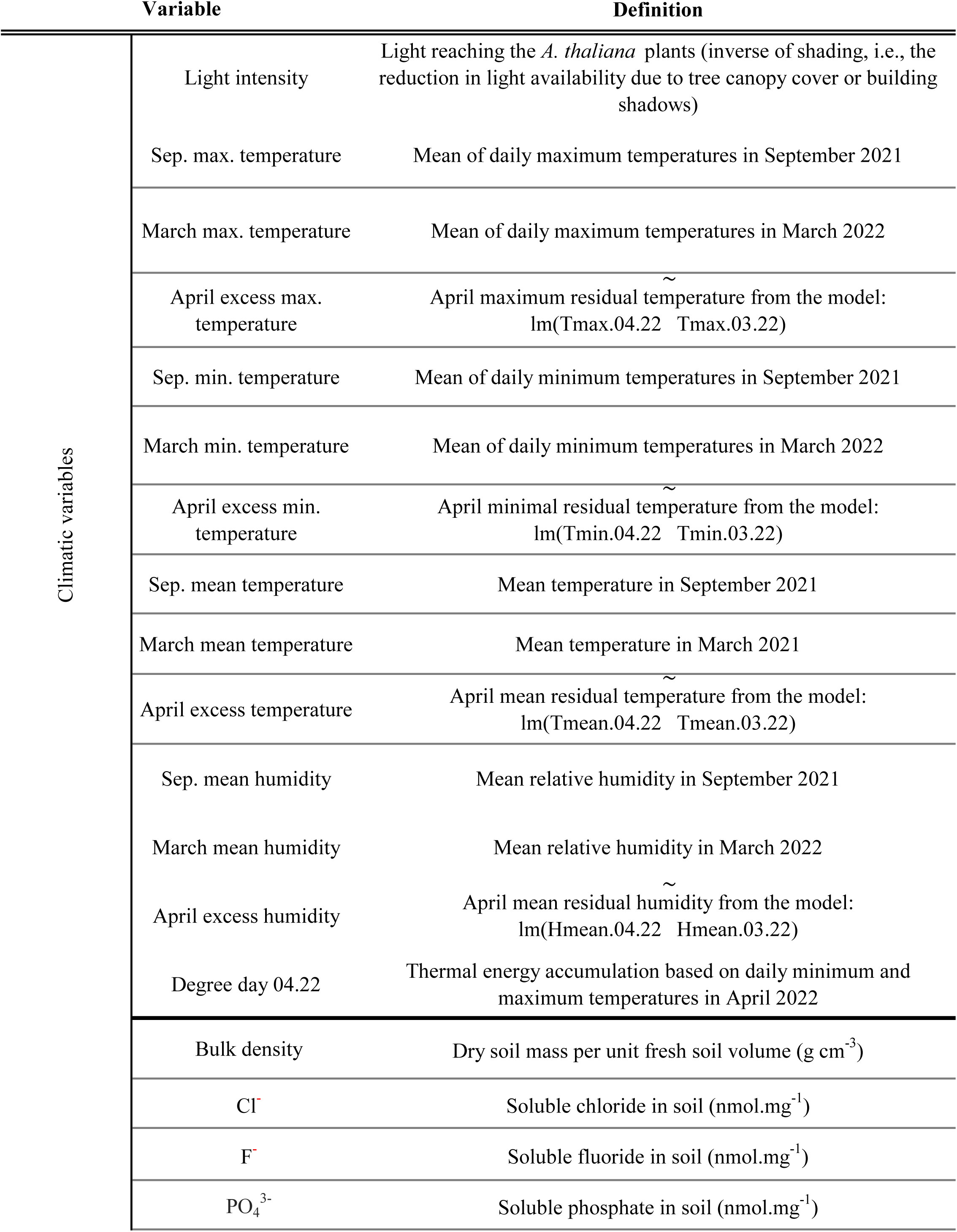
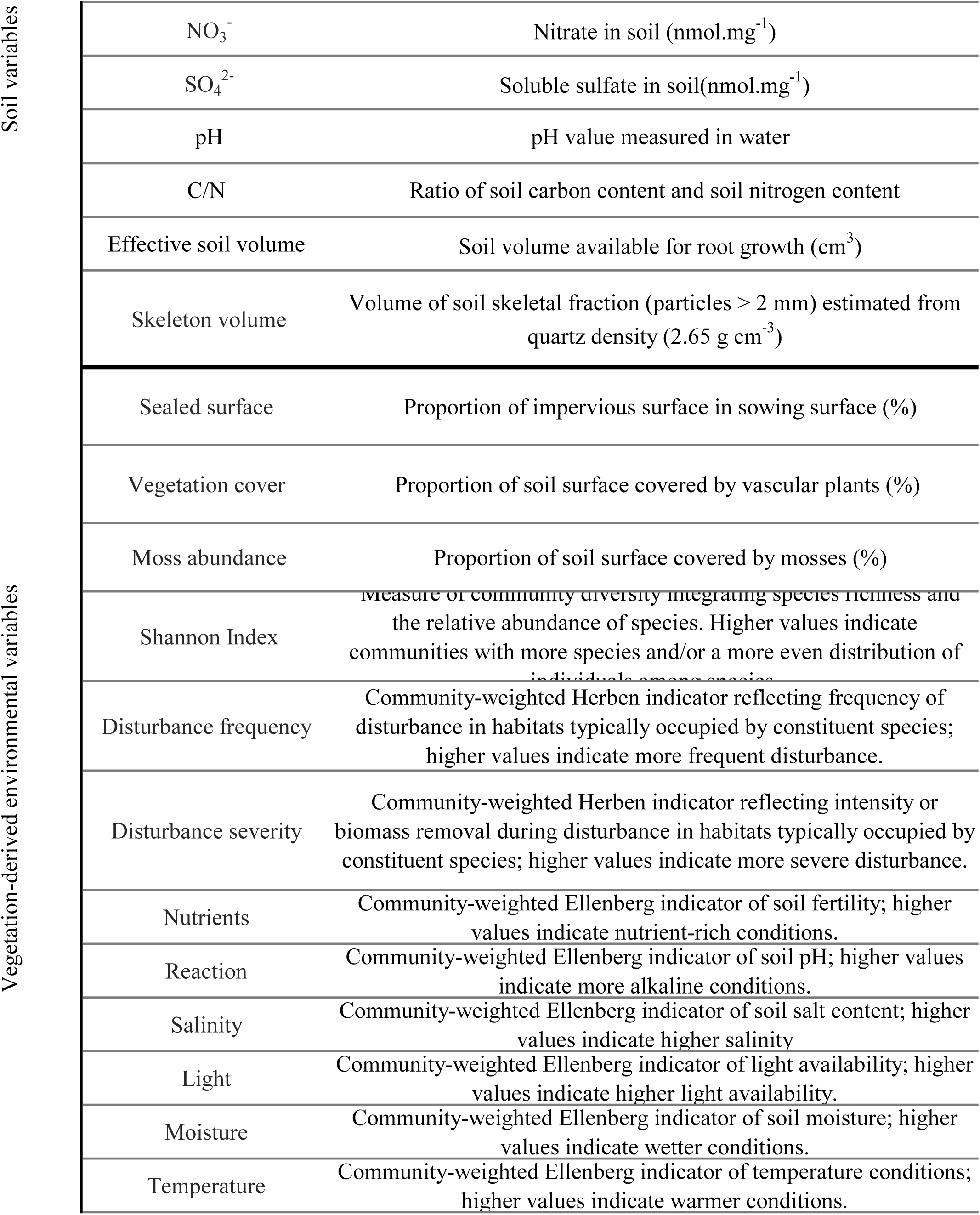
Environmental factors characterizing urban habitats in Cologne (Germany). Variables were recorded across key phenological stages of Arabidopsis thaliana: at sowing (September 2021), end of vegetative growth (February 2022), and fruit ripening (May 2022), across 393 sites.

|  | Variable | Definition |
| --- | --- | --- |
| Climatic variables | Light intensity | Light reaching the *A. thaliana* plants (inverse of shading, i.e., the reduction in light availability due to tree canopy cover or building shadows) |
|  | Sep. max. temperature | Mean of daily maximum temperatures in September 2021 |
|  | March max. temperature | Mean of daily maximum temperatures in March 2022 |
| | April excess max. temperature | April maximum residual temperature from the model:<br>$\text{lm}(\text{Tmax.04.22} \sim \text{Tmax.03.22})$ |
|  | Sep. min. temperature | Mean of daily minimum temperatures in September 2021 |
|  | March min. temperature | Mean of daily minimum temperatures in March 2022 |
| | April excess min. temperature | April minimal residual temperature from the model:<br>$\text{lm}(\text{Tmin.04.22} \sim \text{Tmin.03.22})$ |
|  | Sep. mean temperature | Mean temperature in September 2021 |
|  | March mean temperature | Mean temperature in March 2021 |
| | April excess temperature | April mean residual temperature from the model:<br>$\text{lm}(\text{Tmean.04.22} \sim \text{Tmean.03.22})$ |
|  | Sep. mean humidity | Mean relative humidity in September 2021 |
|  | March mean humidity | Mean relative humidity in March 2022 |
| | April excess humidity | April mean residual humidity from the model:<br>$\text{lm}(\text{Hmean.04.22} \sim \text{Hmean.03.22})$ |
|  | Degree day 04.22 | Thermal energy accumulation based on daily minimum and maximum temperatures in April 2022 |
|  | Bulk density | Dry soil mass per unit fresh soil volume (g cm<sup>-3</sup>) |
|  | Cl<sup>-</sup> | Soluble chloride in soil (nmol.mg<sup>-1</sup>) |
|  | F<sup>-</sup> | Soluble fluoride in soil (nmol.mg<sup>-1</sup>) |
|  | PO<sub>4</sub><sup>3-</sup> | Soluble phosphate in soil (nmol.mg<sup>-1</sup>) |

Soil variables
|  |  |
| --- | --- |
| $\text{NO}_3^-$ | Nitrate in soil ($\text{nmol.mg}^{-1}$) |
| $\text{SO}_4^{2-}$ | Soluble sulfate in soil($\text{nmol.mg}^{-1}$) |
| pH | pH value measured in water |
| C/N | Ratio of soil carbon content and soil nitrogen content |
| Effective soil volume | Soil volume available for root growth ($\text{cm}^3$) |
| Skeleton volume | Volume of soil skeletal fraction (particles > 2 mm) estimated from quartz density ($2.65 \text{ g cm}^{-3}$) |

Vegetation-derived environmental variables
|  |  |
| --- | --- |
| Sealed surface | Proportion of impervious surface in sowing surface (%) |
| Vegetation cover | Proportion of soil surface covered by vascular plants (%) |
| Moss abundance | Proportion of soil surface covered by mosses (%) |
| Shannon Index | Measure of community diversity integrating species richness and the relative abundance of species. Higher values indicate communities with more species and/or a more even distribution of individuals among species. |
| Disturbance frequency | Community-weighted Herben indicator reflecting frequency of disturbance in habitats typically occupied by constituent species; higher values indicate more frequent disturbance. |
| Disturbance severity | Community-weighted Herben indicator reflecting intensity or biomass removal during disturbance in habitats typically occupied by constituent species; higher values indicate more severe disturbance. |
| Nutrients | Community-weighted Ellenberg indicator of soil fertility; higher values indicate nutrient-rich conditions. |
| Reaction | Community-weighted Ellenberg indicator of soil pH; higher values indicate more alkaline conditions. |
| Salinity | Community-weighted Ellenberg indicator of soil salt content; higher values indicate higher salinity |
| Light | Community-weighted Ellenberg indicator of light availability; higher values indicate higher light availability. |
| Moisture | Community-weighted Ellenberg indicator of soil moisture; higher values indicate wetter conditions. |
| Temperature | Community-weighted Ellenberg indicator of temperature conditions; higher values indicate warmer conditions. |

### The urban niche of *A. thaliana*

The experimental distribution of the same F2 seed set across 393 unmanipulated habitat sites enabled an unbiased identification of environmental parameters that delimit the environmental niche allowing *A. thaliana* population recruitment and persistence within the city of Cologne. Because *A. thaliana* is a ruderal species, whose vegetative biomass and seed production decline strongly in the presence of companion plants (Palacio-Lopez et al. 2020; Libourel et al. 2021; Bastias et al. 2024; Baron et al. 2025), we expected establishment to be favored in disturbed and sparsely vegetated habitats. Consistent with this expectation, habitat type significantly influenced establishment (Fig. 2A). Walls exhibited the highest establishment rate (77%), whereas meadows showed the lowest (21%; F_3,389_ = 14.76, *p* = 1.05e-09). In contrast, population persistence did not differ between habitat types (χ^2^ *=* 1.68, F = 6, p = 0.95; Fig S4).

To identify environmental drivers of establishment, we first analyzed indicators derived from the resident plant community (Ellenberg et al. 1991; Midolo et al. 2023). Because plant communities integrate environmental conditions over comparatively long-time scales, they provide reliable proxies for resource availability and disturbance regimes (Diekmann 2003). Establishment was strongly favored by higher disturbance severity (503-fold increase per unit, p = 2.3e-02), but declined with nutrient availability (1.20-fold decrease per unit, p = 2.7e-02).

We next analyzed a large set of directly measured abiotic and biotic environmental variables (Fig. 2B; Table 2). Across habitats, establishment was associated with environmental conditions characteristic of urban grey spaces: *A. thaliana* established preferentially in shallow, physically constrained soils with high bulk density (12.84-fold increase, p = 1.71e-04) and low skeletal content (1.05-fold decrease per unit, p = 2.19e-02), as well as in sites with greater surface sealing and higher light availability (both p < 0.01). These conditions are typically associated with low soil development and reduced resource availability, which limit the growth of more resource-demanding competitors and thereby create opportunities for this ruderal species (Mathur et al. 2014; Taylor et al. 2017; Correa et al. 2019; Libourel et al. 2021; Giuliani et al. 2024b). Microclimatic conditions further contributed to the establishment, with higher temperatures in September and March and lower air humidity in March and April associated with reduced establishment (all p < 0.05). These results were consistent with analyses based on plant-community indicators, which suggests that *A. thaliana* establishment is favored in harsh urban habitats characterized by frequent disturbance and low resource availability.

**TABLE 2:** Environmental factors influencing the establishment of *Arabidopsis thaliana* in urban habitats. LASSO regression was used to identify environmental predictors of establishment (recorded April 2022). Variables selected by the LASSO model were subsequently analysed using a generalized linear model (GLM) to estimate their effects.

|  |  | Df | glm estimate | P-value | F values | P-value(F) |
| --- | --- | --- | --- | --- | --- | --- |
| All urban environments | Sep. min. temperature | 1 | -0.18 | 4.25e-02 | 4.14 | 4.27e-02 |
|  | March max. temperature | 1 | -0.24 | 2.23e-03 | 9.65 | 2.05e-03 |
|  | March mean humidity | 1 | 0.01 | 3.25e-02 | 4.71 | 3.07e-02 |
|  | Effective soil volume | 1 | 2.35 | 1.85e-02 | 5.28 | 2.22e-02 |
|  | Skeleton volume | 1 | -0.05 | 2.19e-02 | 5.60 | 1.85e-02 |
|  | Bulk density | 1 | 9.46 | 1.71e-04 | 19.01 | 1.71e-05 |
|  | Moss abundance | 1 | 2.36 | 1.58e-01 | 2.48 | 1.16e-01 |
|  | Sealed surface | 1 | 2.72 | 7.58e-04 | 11.12 | 9.44e-04 |
|  | April mean humidity | 1 | 0.03 | 4.28e-03 | 8.50 | 3.77e-03 |
|  | Light intensity | 1 | 0.00 | 6.51e-05 | 16.49 | 6.03e-05 |
|  | Residual | 356 |  |  |  |  |
| Meadow | Sep. min. temperature | 1 | -0.95 | 2.86e-04 | 17.94 | 4.66e-05 |
|  | Bulk density | 1 | 12.19 | 2.13e-03 | 12.86 | 4.97e-04 |
|  | Vegetation cover | 1 | -3.32 | 1.45e-02 | 8.15 | 5.11e-03 |
|  | Light intensity | 1 | 1.27e-03 | 3.68e-02 | 5.94 | 1.63e-02 |
|  | Residuals | 113 |  |  |  |  |
| Tree bed | March mean humidity | 1 | 0.15 | 5.99e-05 | 21.10 | 1.43e-05 |
|  | Skeleton volume | 1 | -0.17 | 5.76e-03 | 9.99 | 2.15e-03 |
|  | C/N ratio | 1 | -0.40 | 2.54e-02 | 6.55 | 1.22e-02 |
|  | Effective soil volume | 1 | 8.49 | 2.50e-03 | 11.97 | 8.31e-04 |
|  | Phosphate (PO<sub>4</sub><sup>3-</sup>) | 1 | -3.52e-03 | 8.29e-04 | 14.64 | 2.41e-04 |
|  | Vegetation cover | 1 | -2.40 | 3.28e-02 | 5.97 | 1.65e-02 |
|  | April Degree days | 1 | -0.02 | 4.87e-02 | 5.09 | 2.65e-02 |
|  | Chloride (Cl<sup>-</sup>) | 1 | 1.44e-03 | 4.35e-03 | 10.65 | 1.56e-03 |
|  | Residuals | 89 |  |  |  |  |
| Pavement | Sealed surface | 1 | 6.85 | 5.08e-05 | 29.50 | 4.36e-07 |
|  | Light intensity | 1 | 3.56e-03 | 9.02e-05 | 26.76 | 1.30e-06 |
|  | March mean humidity | 1 | 0.29 | 1.06e-03 | 18.33 | 4.48e-05 |
|  | Phosphate (PO<sub>4</sub><sup>3-</sup>) | 1 | 4.06e-03 | 3.20e-02 | 8.92 | 3.59e-03 |
|  | Residuals | 94 |  |  |  |  |
| Wall | Moss abundance | 1 | 9.01 | 2.47e-04 | 24.80 | 1.08e-05 |
|  | Sealed surface | 1 | 7.80 | 1.18e-02 | 11.70 | 1.38e-03 |
|  | Residuals | 43 |  |  |  |  |

Beyond these general patterns, habitat-specific models revealed additional context-dependent environmental filters (Fig. 2C; Table 2). In meadows, establishment declined with decreasing minimal temperature in September (0.39-fold decrease per unit, p = 2.86e-04). In tree beds, establishment decreased with increasing skeletal content (1.19-fold decease, p = 5.76e-03), but increased with higher humidity in March (1.16-fold increase, p = 5.99e-05). On pavements, establishment was promoted by greater surface sealing (943-fold increase per unit, p = 5.08e-05) and higher light availability (1.001-fold increase per unit, p = 9.02e-05), consistent with the general association with harsh urban habitats. Walls represented the extreme of the human impact gradient, corresponding to the harshest urban habitats with minimal and highly discontinuous soil availability. Here, establishment increased with both surface sealing (2.44e03-fold increase per unit, p = 1.18e-02) and moss abundance (8180-fold increase per unit, p = 2.47e-04). The latter suggests either microsite amelioration or direct facilitation via mosses, for example through enhanced moisture retention and temperature buffering in otherwise harsh environments (Anthelme et al. 2014; Lett et al. 2020). In harsh environments, bryophytes consistently facilitate seedling emergence and survival, often outweighing competitive effects (Soudzilovskaia et al. 2011).

In pavements, another highly modified urban habitat, higher resource availability (in particular PO₄³⁻ concentration and humidity) had a weak but significant positive effect on *A. thaliana* establishment, a pattern not observed in other habitat types (Fig. 2D, Table 2). Similar mechanisms are known from drylands, where establishment is frequently restricted to protected microsites that accumulate water and fine soil (Loayza et al. 2017; Shemesh 2025). Most ecological filters limiting the urban recruitment niche of *A. thaliana* likely operate broadly across less human-modified environments. In contrast, the significant effect of chloride concentration (1.004-fold increase per unit, p = 1.56e-03) suggests that a pollutant specifically associated to urban habitats can locally promote *A. thaliana* establishment, presumably by reducing competition (Fig. 2D, Table 2).

### Dozens of genetic variants contribute to local adaptation in the city

Our experimental design allowed us to directly link establishment to the reservoir of genetic variation that was mobilized for rapid adaptation. Indeed, we could interrogate what environmental, phenotypic and phenological variation was associated with genome-wide allele frequency changes after three generations of population differentiation. By design, in the initial F2 seed population, parental alleles are present at frequency 0.5. In the absence of selection, allele frequencies are expected to diverge across populations due to drift, and thus to be independent of environmental variation. We used simulations that accounted for population size and recombination along the genomes of individuals in each population to calibrate false discovery rates (Fig. S5), and to account for missing environmental data (with some sets of predictors only available for 32 of the 68 sequenced populations). With this approach, we identified 18 genomic regions in which allele frequencies were associated with environmental variables at a false discovery rate of 0.25, implying that about 13 of these variants are true positive (Fig. 3). We thus conclude that of the 36 least correlated environmental parameters, approximately 12-14 ecological factors select for alleles differentiating the genomes of the meadow parent (MIL2) and the wall parent (KAD1) (Fig. 3, Table S1). The strong statistical support of some variant associations and/or their co-localization with large effect mutations in known candidate genes confirm that many of these loci are very likely to be true positives.

**Figure 3:**
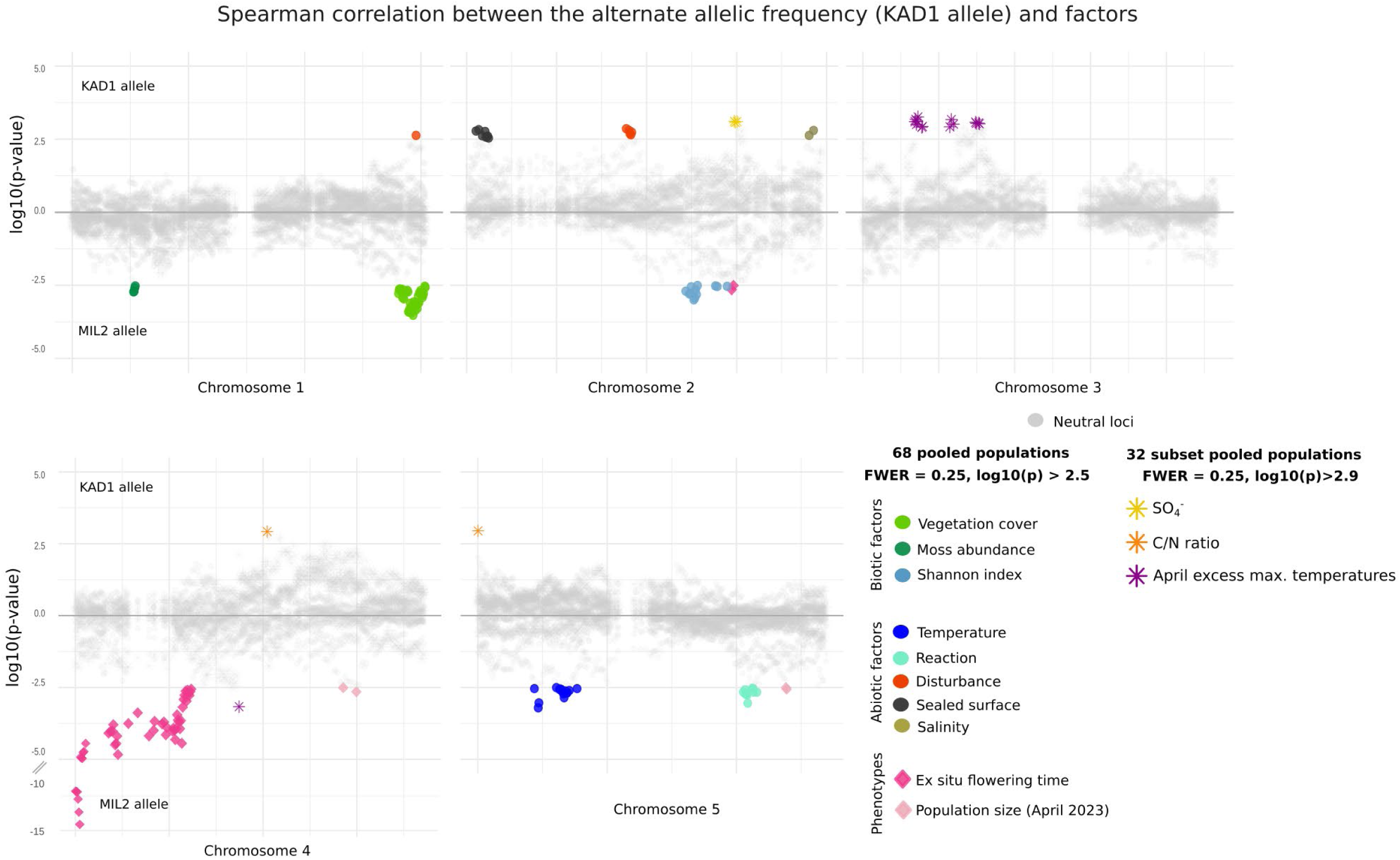
Genomic regions associated with phenotypes and environmental factors. While positive p-values indicate a positive correlation between the median KAD1 allele frequency and environmental factors, negative p-values indicate a positive correlation between the median MIL2 allele frequency and environmental factors. Non-significant correlations between allele frequencies and environmental factors are in grey.

For example, disturbance, local population size, and elevated April temperature were associated with genomic regions on two different chromosomes, a pattern that is highly unlikely to arise by chance (family-wise error rate FWER<0.03, Fig. S5). In addition, a MIL2 allele at the end of chromosome 1 was associated with increased vegetation cover (FWER < 0.05). This region overlaps the gene *PERK13*. The parent KAD1 carries a *PERK13* allele previously shown to reduce the ability to adjust plant size in response to competition with grasses, a trait known to be under strong selection in competitive environments (Frachon et al. 2018; Libourel et al. 2026). Finally, the positive association of the KAD1 allele on chromosome 2 correlates with soil sulfate levels and maps to a chromosomal segment containing a loss-of-function allele of the phosphate transporter PHT1:4 due to a deletion having removed the C-terminal part (amino acids 493-535). This transporter belongs to the regulatory network underlying sulfur content variation in *A. thaliana* (de Jager et al. 2024).

Adaptive differences across the genome are strikingly aligned with ecology at the site of origin of the parental genotypes. The MIL2 allele carried alleles whose frequency increased with vegetation cover, moss abundance, and plant community diversity (Spearman ρ=0.43, 0.37 and 0.40, respectively), three factors indicative of more competition-prone and stressful conditions. The MIL2 allele was also more frequent in sites hosting larger populations (on chromosome 4, R=-0.37, p = 2.24e-03; and on chromosome 5, R= -0.36, p= 2.74e-03; Table S1). In contrast, the KAD1 allele increased in frequency in low pH plant communities (Ellenberg Index Reaction), with increasing soil sulfate levels, higher salinity, higher disturbance, higher sealed surface and lower soil N content (Spearman ρ= -0.40, 0.56, 0.38, 0.37-0.39, 0.38, 0.54, respectively; Table S1). Despite their geographic proximity, the two genotypes thus show signatures consistent with local adaptation to meadow and wall, their urban environments of origin.

The number of associated genomic regions shows that their local adaptation is polygenic and most likely mediated by multiple traits providing advantages to diverse environmental factors. Interestingly, we did not find significant associations between genomic variation and any of the four habitat types, but with many of the continuous environmental factors that together differentiate habitat types (Fig. S6). This indicates that variants tend to have factor-specific effects, and it is the assemblage of multiple adaptive alleles that allows specialization to different kinds of environments.

### Competition, disturbance and genetics shape the habitat carrying capacity of *A. thaliana*

Our experiment demonstrates that genetic variation influences not only the establishment niche but also a second key component of the species’ niche: local carrying capacity. Indeed, population size varied across habitat types, but was stable over the years, indicating that established sites differed in their maximum population size, i.e., their carrying capacity. The most suitable urban environment in terms of establishment (i.e., walls) exhibited the lowest carrying capacity (population size) across habitat types (Fig. 4A, F_3,148_ = 13.68, p = 1.6e-02). Thus, contrary to expectation, habitat suitability and carrying capacity were not aligned (Yim et al. 2024; Holian et al. 2026).

**Figure 4:**
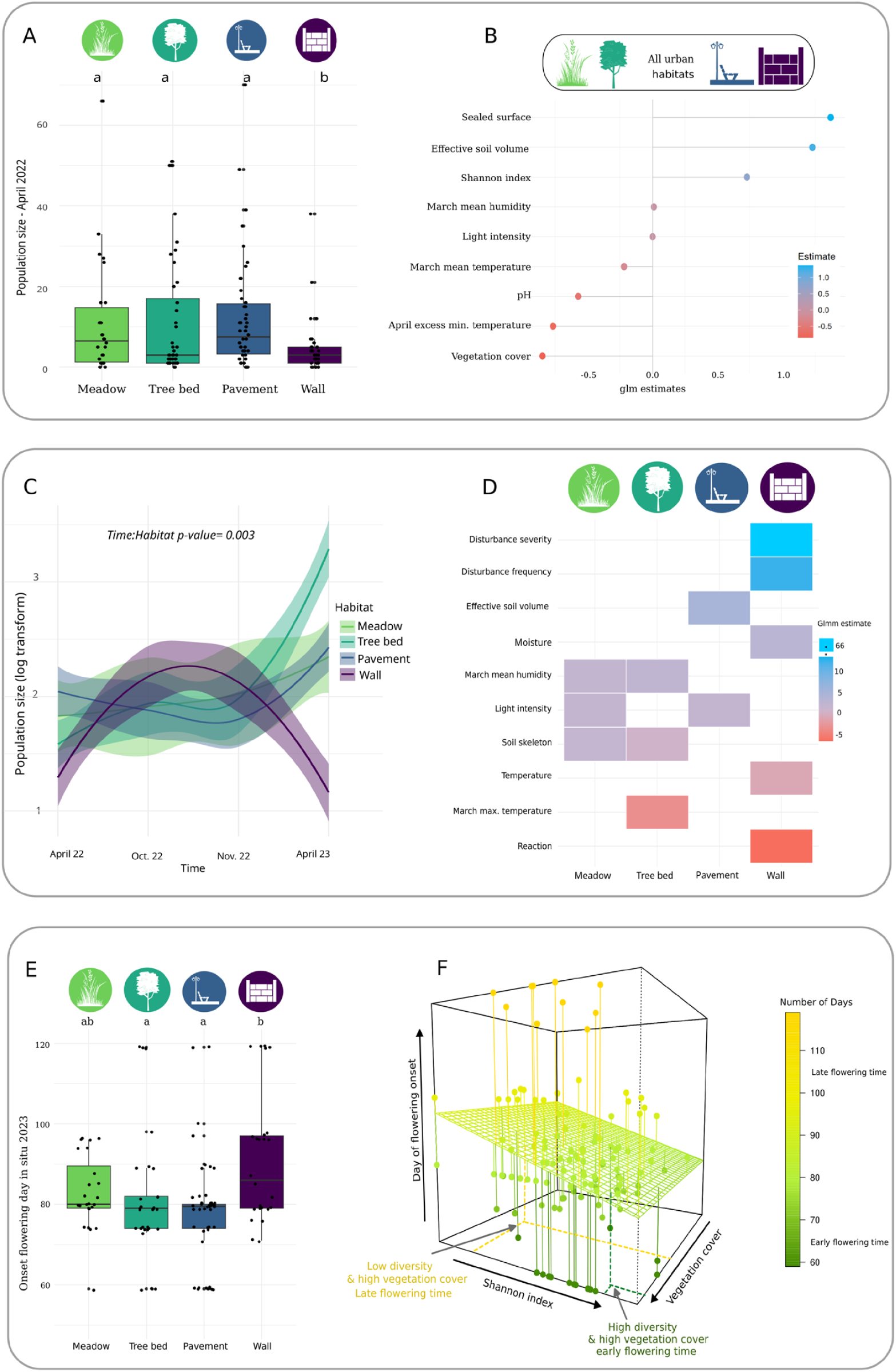
The role of the ecological factors on *A. thaliana’s* population biology in natural urban environments. Across 393 urban sites monitored between April 2021 and June 2023, 37 ecological factors were measured (Table 1). **A**. Population size of established population in April 2022. **B.** Across all urban environments: Ecological factors positively (blue) or negatively (red) associated with the population size in April 2022, April 2023 and April 2024. **C.** Population size of established population over a generation, from April 2022 to April 2023. **D.** Ecological factors positively or negatively associated with the population size of the second generation in October and November 2022, and April 2023. **E.** Onset flowering, the first flowering plant in the population, per established population in Julian days in April 2023. **F.** Onset flowering time per established population along gradient of biodiversity (Shannon index) and plant density (Vegetation cover).

Across habitats, variation in population size depended mainly on factors associated with competitive pressure (Table S2). Populations tended to be larger in sites with reduced dominance of single competitive species (Fig. 4B). Population size also increased with higher Shannon diversity (with a 1.36-fold increase; χ^2^ = 32.45, p = 1.22e-08), greater effective soil volume (with a 1.23-fold increase; χ^2^ = 8.85, p = 2.94e-03), lower vegetation cover (1.19-fold decrease per unit; χ^2^ = 18.43, p *=* 1.76e-05), and surfaces were sealed (1.37- fold increase, χ^2^ = 27.14, p = 1.89e-07, Fig. 4B, Table S2). These findings align with the ruderal life-history strategy of *A. thaliana*, with high levels of competition limiting the carrying capacity (Takou et al. 2019). The carrying capacity varies over time and this differs between the habitats (Fig. 4C). This is likely due to the complex effects of the biotic and abiotic drivers of carrying capacity that changed across habitat types (Fig. 4C-D). Some factors even had antagonistic effects. The severity and frequency of environmental disturbance were the main positive predictors of population size on walls (50.49-fold increase per unit, χ^2^ = 4.83, p = 2.80e-02, and, 5.52-fold increase per unit, χ^2^ = 5.16, p = 2.31e-02 respectively). Meanwhile in meadows, disturbance frequency decreased population size (1.95-fold decrease per unit, χ^2^ = 4.75, p *=* 2.94e-02).

Our genetic analysis shows that genetic variation further explains variation in population size. Indeed, on chromosome 4, we detected a genomic region associated with variation in population size, and so carrying capacity. This region overlapped a region weakly associated with variation in soil phosphate content (Fig. S6), which contains *PHR1*, a gene needed to respond to phosphate starvation. The KAD1 allele of *PHR1*, which presents a leucine instead of the conserved proline residue at position 203 in the protein, is identical to a known QTL allele defective in root growth response to starvation (Reymond et al. 2006). The direction of the association suggests that populations with the MIL2 allele can be larger in sites with low soil phosphate content, but its significance does not pass acceptable FWER thresholds (Fig. S5). Further investigations are needed to validate the effect of *PHR1*. While our data show that genetic variation at specific loci can contribute to broadening the establishment niche of *A. thaliana* urban populations, we suggest that it can also enhance their carrying capacity in specific environments.

### Genetic variation broadens the species’ urban niche

This experiment is unique in that it allows an experimental characterization of the species niche for the city of Cologne. In particular, it allows us to isolate separately parameters of the recruitment niche (establishment) and the parameters of the persistence niche (carrying capacity), and identify the dimensions of the niche that are supported by natural genetic variation differentiating two local urban genotypes. Our results show that the ecological dimensions that drive establishment or persistence overlap with the ecological dimensions that associate with differentiation at the genomic level. For example, we found genetic variants which widen the gradients of vegetation cover, soil phosphate content, sealed surface, moss abundance or disturbance that are suitable for population establishment (Fig. 3, Table S3). Yet, some environmental variables, such as soil C:N ratio and sulfate concentration, do not measurably influence establishment probability but have selected specific alleles (Fig. 2B-C). We conclude that C:N ratio and soil sulfate would also impose limits to establishment in the absence of suitable genetic variation. The genetic variation present in spontaneous populations of *A. thaliana* thus broadens the urban niche of the species, allowing growth beyond just the disturbed environments typical of ruderal species, as well as across a range of resource levels. In addition, alleles carried by the meadow parent, MIL2, promote population establishment in habitats with high moss abundance, which in our city occurs on walls and not on meadows. This suggests that recombination between urban genotypes could even extend the niche of the species. Our study thus documents the underlying complexity of species niches, beyond the presence/absence data that are classically used to define them (Ikeda et al. 2017). With 68 sequenced populations, our power to interrogate associations between genetic variation and interacting environmental factors has some limits, but it is tempting to speculate that the interaction between variants will further modulate the impact of secondary environmental filters on population diversification.

### Habitat diversity influences the diversity of phenology in situ, but not its adaptive relevance

Genetic variation controlling phenology has been intensively studied and is believed to provide essential adaptation to novel environments (Hancock et al. 2025). Flowering changed along disturbance gradients in spontaneous *A. thaliana* populations of the city of Cologne and the parental lines differed in the regulation of both flowering time and dormancy (Schmitz et al. 2024). Our experiment thus offers a framework to test the relevance of these alleles for defining phenology in the urban environment and test their relevance for adaptation.

First, both population dynamics of seedling recruitment and seed dormancy varied between the types of habitat (Fig. 4C, Time:Habitat F_6,412_ = 3.30, p = 3.15e-14). On walls, census population size peaked in autumn and subsequently declined in spring. This observation stands in contrast with tree beds and pavements, where population census size increased throughout the year, indicating that, contrary to walls, the germination rate was higher than the death rate in spring in these environments. The severity and frequency of environmental disturbance were significantly associated with census size dynamics on walls (66.42-fold increase in population size, χ² = 5.07, p = 2.43e-02, and 9.14-fold increase, χ² = 7.53, p = 6.08e-03, respectively; Fig. 4D). Populations differed significantly in the germination rate (interaction population * time after harvest, χ² = 4076.4, p < 2.2 e-16, Fig. S7), yet it did not associate with census size dynamics nor with any of the environmental parameters measured. We thus could not establish the adaptive relevance of genetic variation in seed dormancy in the city of Cologne.

As previously described (Schmitz et al. 2024), the timing of flowering of plants growing in situ in unmanipulated habitat sites was also habitat-specific and associated with environmental variables. On walls, onset of flowering occurred later than in the other three habitats (Fig. 4E, Table S3, F_3,126_ = 0.73, p= 1.1e-02). Overall, *A. thaliana* populations with relatively higher moss abundance (23.5-fold increase per unit, χ^2^ =9.87, p = 1.66e-03), greater light intensity (1-fold increase per unit, χ^2^ = 301.81, p = 1.33e-67), and higher humidity in March (1.02-fold increase per unit, χ^2^ = 4.36, p = 3.67e-02) flowered later. However, the interaction between light intensity and mean March humidity (χ^2^ = 7193.55, p < 2e-16) associated with accelerated flowering. The interaction between Shannon index and vegetation cover (χ² = 17.87, p = 2.36e−05) further indicated that populations growing in places where vegetation cover was high, flowered earlier in an environment with greater biodiversity than in an environment with lower biodiversity (Fig. 4F, Table S3). *A. thaliana* thus appears to be under pressure to reproduce before later-growing species in the seasonal succession (F. Cahill Jr et al. 2005; Bittebiere and Mony 2015).

Our experiment provided a specific test of the adaptive relevance of genetic variation in a well-known regulator of flowering, the gene *FRIGIDA* (Johanson et al. 2000). Indeed, KAD1 presented a non-functional *FRIGIDA* allele and genetic differences in average flowering time of the 68 sequenced populations associated strongly with allelic variation at the edge of chromosome 4 (Spearman ρ =0.78, p < 10e-07, FDR < 0.001), where the gene is located (Fig. 3, Table S1-S4). The correlation between ex situ flowering time, which only reflects genetic differences in the regulation of this trait, and in situ flowering time (population phenology) were positively correlated, especially in sites exposed to high levels of light (χ² = 10.1, p = 1.45e-03). We thus conclude that part of the phenotypic variation observed under natural conditions has a genetic basis (χ² = 10.1, p = 1.45e-3). Yet, genomic regions associated with flowering time, including major QTL such as *FRIGIDA* on chromosomes 4 but also a second weakly associated allele on chromosome 2 (Reeves and Coupland 2001), were not located in genomic regions associating with environmental variables (Fig. 3, Table S4). Rather, it was flowering time in situ (population phenology) that associated strongly with environmental variables. In contrast to expectations, we thus found no evidence for an adaptive role of flowering regulation by *FRIGIDA* under long-day conditions in the city of Cologne (Schmitz et al. 2024). Together, these results show that germination and flowering time of populations growing in unmanipulated urban sites are primarily shaped by phenotypic plasticity in response to local abiotic and biotic environmental conditions.

## Conclusion

We present a genetically informed, experimentally resolved decomposition of the realized urban niche into recruitment and persistence components, integrating natural environmental heterogeneity with standardized propagule input. This framework directly links ecological filtering with genome-wide differentiation under real-world conditions, without the biases of fully controlled greenhouse or climate chamber experiments. Our results show that *Arabidopsis thaliana* occupies urban environments through two complementary ecological strategies. In disturbance-prone urban green habitats, it behaves as a typical ruderal species, avoiding competition through temporal niche escape and rapid life-cycle completion. In contrast, harsh urban grey habitats support exceptionally high recruitment but low population densities, indicating that conditions favoring establishment differ from those supporting long-term persistence. Together, these findings demonstrate that recruitment and persistence niches can be partially decoupled within a single species, extending the classical view of *A. thaliana* as a predominantly disturbance-bound ruderal species.

Our experiment further shows that this niche cannot be realized without a contribution of local genetic diversity. The genomic analysis of persisting populations identifies a large reservoir of genetic variants that can be rapidly mobilized to promote population establishment along fine-scale environmental gradients, including disturbance, vegetation structure, soil chemistry, and habitat carrying capacity. This polygenic basis reinforces the central role of standing genetic variation in enabling rapid evolutionary processes under spatially structured selection, even at the scale of a single city. More broadly, our findings show that substantial adaptive differentiation can emerge over remarkably short spatial and temporal scales from only two parental genotypes. The experimental framework introduced here – combining field-realistic environments, controlled seed input, and genome-wide inference – provides a powerful and transferable tool for dissecting recruitment and persistence as adaptive processes in heterogeneous landscapes, and for understanding how realized niches are assembled in nature. Finally, our study highlights the value of cities as replicated open-air laboratories for ecology and evolution. Urban environmental mosaics not only enable analysis of the mechanisms generating ecological and genetic diversity, but also offer opportunities to identify processes that may help maintain and promote biodiversity in human-dominated landscapes. Thus, while urbanization remains a major challenge for biodiversity conservation, cities may also provide unexpected opportunities to understand, and potentially foster, the processes that sustain biodiversity.

## Acknowledgements

We thank Thibault Leroy for helpful discussions; Johanna Lucia Schmid, Anna Hakimzadeh, Esi Annobil, Anna Dölz, Thorben Kunisch, Melvin Brodbeck and Clara Hoffmann for their help in population characterization; Volker Kummer for support in plant identification, as well as Simone Brockmann and Gabi Gehrmann for their help with soil analyses. Lina Abdelwahed, Leen Abraham, Lea Hördemann, Neda Rahnamae, Renan Grenado Chavez and Annalisa Falzone-Tegtmeier are acknowledged for occasional field support.

## Author contributions

Conceptualization: JM, AL, JF

Methodology: JM, AL, JF, SP, FB, TA, GS, SK

Investigation: JF, HZ, LW, GB, KB, GS

Visualization: JF, HZ, VH

Funding acquisition: JM, AL

Project administration: JM Supervision: JM, AL

Writing – original draft: JF, JM, AL

Writing – review & editing: SK, FR, MT, TA, GB

## Data and materials availability

The data and R scripts are available at the github: https://github.com/jfloret23/Urban_niche_Ath.

