## Supplementary Material for "Dozens of genetic variants sustain adaptation to urban spatial heterogeneity in *Arabidopsis thaliana*"

**The PDF file includes:**

Materials and Methods
Figs. S1 to S10
Tables S1 to S8
References

### Materials and Methods

#### Study design

To understand how environmental heterogeneity within a city shapes population establishment and adaptation, we conducted a city-wide colonization experiment with *Arabidopsis thaliana* across Cologne, Germany, treating the urban landscape as a spatially replicated environmental gradient. Urban habitats were classified *a priori* into four habitat types that represent common plant habitats within cities and differ in substrate availability, vegetation cover, and disturbance regime (Bonthoux et al. 2019): meadows and tree beds (green spaces) and pavements and walls (grey spaces). Meadows and tree beds represent vegetated urban habitats, whereas pavements and walls represent highly impervious substrates, together spanning a gradient of increasing human impact (Giuliani et al. 2024). These four habitat types have been previously found to be harbor spontaneous *A. thaliana* populations (Schmitz et al. 2024). We aimed to sample 100 suitable, unoccupied sites per habitat type that were representative of each category for experimental colonization.

To identify suitable sites, 616 candidate sites distributed across Cologne and the four habitat types were surveyed twice monthly between March and May 2021 and once monthly in July and August 2021. Sites were only retained if repeated surveys confirmed the absence of spontaneous *A. thaliana* populations. In Spring 2022 and 2023, 393 sites were selected for experimental colonization, comprising 121 meadows, 106 tree beds, 119 pavements, and 47 walls (Fig 1 and 2A). The number of suitable wall habitats remained limited, due to the destruction of the targeted sites by human activity, resulting in lower replication for this habitat type. Biotic and abiotic environmental conditions were subsequently quantified at all sites during 2022 and 2023.

#### Characterization of biotic and abiotic conditions of each site

##### Surface characteristics and vegetation-derived environmental variables

Surface characteristics (including vegetation cover) was quantified at three ecologically relevant time points during the *A. thaliana* life cycle: prior to germination (at sowing; September 2021), at the end of the vegetative phase (February 2022), and at fruit ripening (May 2022; Table S5). To this end, we established standardized experimental plots (1 m$^2$) at each site, where *A. thaliana* seeds were sown (see below). Plot shape was mostly quadratic but was adjusted in some cases to account for narrow patch shapes. At each sampling event, the ground cover of sealed (impervious) surface, vascular plant cover (vegetation cover), moss cover, and bare soil cover was recorded.
Plant community composition was recorded in April 2022, at the time of peak standing biomass as the ground cover of all vascular plant species. Species identification was conducted on-site using plant identification tools, i.e., the Flora Incognita application (Mäder et al. 2021) and the identification guide "Guide des fleurs de France et d'Europe" by Delachaux (2017). In total, 2,431 plant species records were assembled across 393 sites (6.2 species on average; Table S6). From

these data, we calculated standard diversity metrics, i.e. Shannon diversity (Shannon 1948) and Pielou's evenness (Pielou 1966).

We harnessed plant species' indicator function to characterize sites' abiotic environmental conditions and disturbance regimes. Specifically, we calculated Ellenberg indicator values (EIV) (Ellenberg et al. 1991) which provide species-specific optima along key environmental gradients in light availability (L), temperature (T), soil moisture (F), soil reaction (R), and nutrient availability (N). EIVs are widely used as community-derived proxies for abiotic environmental conditions in both natural and urban ecosystems (Diekmann 2003; Schmitz et al. 2024). In addition, we calculated disturbance frequency and severity (Herben et al. 2016; Midolo et al. 2023) as disturbance regime descriptors. Community-weighted means of EIVs and disturbance indices were calculated for each site using species-specific values (Table S7), excluding *A. thaliana*. In total, 28 surface characteristics and vegetation-derived variables were compiled (Table S5).

Soil variables

Soil sampling was conducted in April 2022. At each site, three soil cores were collected at the outer corners of the 1 $m^2$ experimental plots using a cylindrical corer (4.5 cm height, 5 cm diameter; Table S2). Where coring was not feasible (e.g., pavements), samples were collected using a shovel. Soil cores were pooled per site to obtain a composite sample and stored at 4°C until processing. Samples were homogenized and approximately two-thirds were oven-dried at 55°C for ≥48 h to preserve organic matter, then sieved (2 mm mesh) to separate fine soil from coarse fragments (skeletal material). Skeleton mass was recorded, and skeleton volume was estimated from mass assuming a quartz density of 2.65 g $cm^{-3}$. Dried subsamples were used for (i) soil pH and CN analysis, and for (ii) determination of bulk density and ion content after additional drying at 105° C for ≥48 h.

Soil pH was measured following (Mclean 1982). Ten grams of soil were suspended in 25 mL deionised water, equilibrated for ≥30 min, and measured using a calibrated glass electrode (pH 7 and 4 standards). Soil carbon content (C) and soil nitrogen content (N) were measured by high-temperature combustion on milled samples using an EA3000 Euro Vector CNS analyser (EuroVector, Pavia, Italy). Bulk density was calculated as dry soil mass per unit bulk soil volume of fine earth, with correction for coarse fragments. Bulk volume was derived from core volume minus the estimated volume of skeletal material. Soluble anions ($F^-$, $Cl^-$, $NO_3^-$, $PO_4^{3-}$, and $SO_4^{2-}$) were quantified by ion chromatography (Dionex ICS-1100; Thermo Scientific, Darmstadt, Germany) following aqueous extraction (0.400 ±0.005 g soil in 1 mL distilled water, 6 h incubation at 30°C). Separation was performed using a Dionex IonPacAS22 RFIC 4x 250 mm analytic column. The running buffer included 4.5 mM $Na_2CO_3$ /1.4 mM $NaHCO_3$ and the anions were quantified using standard curves generated using external standard calibration.

Soil depth was measured at each site in July 2023 using metal probes (maximum depth 50 cm; three replicates per site). Effective soil volume ($m^3$) was calculated as the product of fresh soil volume and mean soil depth by the bare soil surface. This metric represents effective rooting soil depth after accounting for surface sealing. When soil sampling was not feasible (e.g., vertical walls), soil properties were approximated using Ellenberg indicator values (nutrient, soil moisture and reaction) derived from local plant communities (community-weighted means based on plant species' cover values). Overall, 24 soil-related variables were measured (Table S7).

Climate variables

DS1923-F5 Hygrochron data loggers (iButton, Whitewater, USA) were used to record soil temperature and relative humidity over one year in the vicinity of experimental plots. A total of 33 loggers were deployed in September 2021, coinciding with seed sowing (with 20 loggers installed in meadow sites, 9 in tree beds, and 3 in pavement sites; and only one could be installed on walls due to the lack of suitable fixing points). Loggers were buried at approximately 4 cm soil depth at one corner of each plot. Where sites were spatially close (e.g., within the same parking lot or street segment), a single data logger was used. Devices recorded soil temperature and relative humidity six times per day (8:00 am, 12:00 pm, 4:00 pm, 8:00 pm, 12:00 am, and 4:00 am). After one year, data were extracted and processed in R using the *pollen* package (Nowosad 2019). We derived mean monthly temperature and humidity, mean daily maximum and minimum temperatures, and growing degree days.

Light intensity was measured indirectly using the Photone Grow Light Meter application (Lightray Innovation GmbH, Zurich, Switzerland). Accuracy was validated against a SpectroSense 2 quantum sensor (Skye Instruments Ltd., Llandrindod Wells, UK) across seven sites with three replicate measurements per site ($R^2 = 0.99$, $p = 1.66e\text{-}08$). At each site, light intensity was first measured first in an unshaded reference area (representing ambient light at the time of day) and immediately afterwards at plant level. Shading intensity was calculated as the difference between ambient and plant-level light. Altogether, we derived 10 climatic factors from the 67 climatic variables measured (Table S8).

Imputation of missing data

Due to environmental constraints, some ecological parameters could not be measured at several sites. These included temperature and humidity recorded with iButtons, as well as soil collection at locations with limited soil volume, such as walls or pavements. To complete missing data, we tested two imputation methods in R (version 4.4.1): imputePCA from the missMDA package (version 1.19 (Josse and Husson 2016)) and missForest from the missForest package (version 1.5 (Stekhoven and Bühlmann 2012)). To assess which method more accurately inferred the missing values, a subset of the dataset with no missing data was used, and observed values were randomly and non-randomly replaced by NA. Both imputation methods were then applied to this modified dataset.

Statistical analysis

The statistical analysis and data visualization were performed in R (version 4.5.2 (R core team, 2022)) using the packages lsmeans (version 2.30-2), emmeans (version 1.11.1), glmmLasso (version 1.6.3), and ggplot2 (version 3.5.2).

- Imputing the missing data

The resulting imputed datasets were compared to the original observed data using Pearson's Chi-square test, mean squared error, F-test, and correlation coefficient, and were also compared to each other using the following linear mixed model:

lmer ( Data_Imputed ~ Variables + Method + ( Method|Variables))

- Ecological variables

The 393 sown sites were characterized in spring 2022 and July 2023 for a total of 120 measured ecological variables, which we subsequently slimmed down to 37 uncorrelated variables (Fig S6, Table 1). Correlations between the ecological variables were computed using the function rcorr from the package Hmisc (version 5.2-3), the Spearman method, and visualized with the package corrplot (version 0.95). When two ecologically important variables were correlated, one was retained in the dataset, and the other was included as residual of the linear model:

lm(Variable2 ~ Variable1)

The characterization of urban habitats was performed using penalized logistic regression (Lasso), suitable for models with a high number of variables. A 10-fold cross-validation was used to calibrate the lambda coefficient. The regression was carried out on standardized variables with the following model:

glmnet(ecological_variable, urban_habitat, al pha = 1, lambda = lambda_min , standardize = TRUE, family = multinomial, type.multinomial = grouped)

A general linear model (GLM) was then fitted with the 10 most important Lasso variables to estimate their significance and effects on urban habitats:

glm(Urban_habitat ~ Variable1 + . . . + Variable + residuals, family = negativebinomial )

To give a practical sense of how much a variable impacts the dependent variable, we extracted the exponential of the estimate and present it in % of increase per unit of the explanatory variable.

**In situ population biology associating with different ecological factors**

*Arabidopsis thaliana* F2 population

Among previously characterized ecotypes collected in spontaneous *A. thaliana* stands in Cologne, the lines MIL2 and KAD1 differ in the regulation of germination and flowering time, as well as in fruit number (Schmitz et al. 2024). These lines were selected as parents to generate F1 individuals.

F1 seeds were then amplified to obtain the F2 population. The amplification was conducted under controlled conditions in a growth chamber (Percival) to prevent contamination with laboratory lines. Plants were grown in 6 cm diameter pots filled with VM type soil (Einheitserde, Sinntal-Altengronau, Germany) for four weeks in short days (8 hours of light at 18◦C/16 hours of night at 16◦C) followed by 6 weeks in short days at 4◦C for vernalization. Plants were then grown in long-day conditions (20◦ C the day and 18◦C during the night with 16 hours of light and 15 minutes of infrared light per day) until fruits ripened and seeds were harvested.

<u>Sowing in 393 urban sites</u>

In total, 393 sites were sown in September 2021: 121 meadows, 119 pavements, 106 tree beds, and 47 walls. At each site, 1,000 seeds of the MIL2 × KAD1 F2 population were sown using a calibrated 2ml Eppendorf "salt shaker" device with perforated lid. Sowing was performed within a delimited 1 $m^2$ quadrat in each site, except on walls, where seeds were sown along a 2 m transect. To increase the number of established populations across the urban gradient, a second sowing campaign was conducted in July 2022 across 77 additional sites in Cologne (10 meadows, 1 tree bed, 20 pavements, and 46 walls). The same F2 population and sowing protocol were used, but seed density was increased to 10,000 seeds per site to ensure sufficient propagule input under highly heterogeneous urban conditions.

<u>Population biology and phenotypic scoring</u>

Each year, sites were monitored at several critical stages of the *A. thaliana* life cycle: germination (September, October), flowering (March, April), and fruiting (May). Sites where no population was established were further monitored at least twice per year (October and April) to ensure we did not miss a population that would have germinated late. These observations enabled quantification of establishment and persistence rates, population size, and onset of flowering:

- Establishment rate was defined as the proportion of sown sites in which *A. thaliana* was present (as observed in spring 2022).

- Persistence score quantified the population's ability to survive across years. A score of 1 indicated that the population germinated and flowered only in April 2022 (first generation) before extinction; a score of 2 indicated persistence for two generations (until April 2023); and a score of 3 indicated continued presence in April 2024 (third generation).

- Population size was determined as the number of individuals per established population. Counts were performed each spring, as well as at multiple time points from autumn 2022 to spring 2023 (second generation): October and November 2022 (after autumn germination, when rosettes were sufficiently developed for identification), February 2023 (end of winter), and April 2023 (spring).

• Onset of flowering was scored in 2023 and recorded as the number of days from 1st January to the first flowering event within each population, measured over a seven-week period in spring 2023. Individuals that flowered in November 2022 were assigned a flowering day of −61 (with day 0 as 1 January 2023). Plants that did not flower during the observation period were given a flowering date of 30 April (onset day 120). Populations sown in September 2021 underwent selection over the whole life cycle, whereas populations sown in July 2022 only had selection for survival during the winter season 2022-2023.

Statistical analysis

The statistical analysis and data visualization were performed in R (version 4.5.2) using the packages lsmeans (version 2.30-2), emmeans (version 1.11.1), car (version 3.1-3), glmmLasso (version 1.6.3), and ggplot2 (version 3.5.2).

- Association between urban habitat types and population biology.

The effect of habitat types on population establishment was determined using a GLM model:

glm(Phenotype ∼ Habitat_type + residuals)

The model family depended on the phenotype considered: establishment was modeled with a binomial distribution, persistence score and onset of flowering time with Poisson distributions, and population size with a quasi-binomial distribution. The significance of habitat type was assessed using the Anova function from the car package (version 3.1-3) in combination with emmeans for post hoc comparisons.

- Association between ecological variables and population biology.

As in the characterization of urban habitats, the association between ecological variables and the established populations was performed using Lasso regression with a 10-fold cross-validation on standardized variables with the following model:

glmnet(Phenotype, ecological_variables, al pha = 1, lambda = lambda_min, standardize = TRUE)

A general linear model was then fitted with the 10 most important Lasso variables to estimate their significance and effects on urban habitats:

glm(Phenotype ∼ Variable1 + . . . + Variable + residuals)

For each phenotype, models were first performed across all habitats and then separately for each habitat type, except for onset of flowering, for which habitat-specific analyses were not performed due to insufficient statistical power.

### Ex situ population biology and the genetic basis of phenotypic variation

Bulk-segregant analysis of flowering time variation in the F2 population

A total of 1,000 seeds of the F2 population MIL2 x KAD1 were planted in 6 cm diameter pots filled with VM soil and grown in a growth chamber (Percival) in long-day conditions (20◦ C during the day and 18◦ C during the night with 16 hours of light) and 60% humidity (as described in (Schmitz et al. 2024)). Flowering time was defined as the number of days between sowing and the first flower for each replicate. Leaf material of the 25% earliest and the 25% latest flowering individuals were bulked and DNA from the two pools was isolated.

The two bulks were sequenced using Illumina 1.3+ Whole Genome Sequencing (WGS), with 60x coverage and 2x150 bp reads. The adapters and poor quality regions were trimmed using Fastp, the mapping to reference genome (TAIR10.1) was done with bwa-mem2 (v2.2.1), and the filtering with samtools (v1.13) to include only the reads that had two good pairs with both pairs well mapped (-f 3). Quality was checked with FastQC for fast files (FastQC) and qualimap for bam files. The Single Nucleotide Polymorphism (SNP) calling was realized with bcftools v1.18 (Danecek et al., 2021), using the MIL2 DNA consensus sequence as a reference, without insertion-deletion (indel) and with p-value of 0.01. The QTL mapping was realized with the R package QTLseqr.

Transplanting F5 generations for DNA sequencing and genetic analysis of phenotypic variation across populations

In February and March 2025, 1,766 plants from 168 populations, 23 meadows, 58 pavements, 38 tree beds, and 49 walls, were transferred to the greenhouse. Per population, up to 20 individuals were randomly sampled. Plants were extracted at the rosette stage, stored in sealed plastic bags with humid paper at 4◦ C overnight and transplanted the next morning in 6 cm diameter pots filled with VM soil. In April 2025, populations where plants were not observed or were too small for transfer were visited again and additional plants from these populations were transplanted. In May 2025, F6 seeds were collected from all transplanted individuals for further analyses.

Primary dormancy measurement.

At least four weeks after seed harvest, a random subsample of up to ten individuals (with at least 50 seeds per individuals) was taken for each transplanted population, excluding populations with less than four individuals. In total, 350 individuals were selected from 87 transplanted populations: 13 meadows, 30 pavements, 21 tree beds, and 23 walls. Germination tests were conducted five, eight, eleven, and thirteen weeks after fruit ripening. The proportion of germination was determined on 30 to 50 seeds per transplanted F5 individual, placed on moist filter paper in 12 well micro plates (approximately 600 µL water). Seeds were incubated under long-day conditions for seven days, after which the number of germinated and non-germinated seeds was counted. A seed was considered germinated when cotyledons were visible.

Ex situ flowering time

The same F5 individuals as for the dormancy experiments were used to determine the time to flowering in common garden conditions. The experiment was performed on 958 individuals from 17 meadow habitats, 38 pavements, 23 tree beds, and 34 walls with 4 to 33 replicates per population. Five seeds per amplified F5 plant were planted in 6 cm diameter pots filled with VM soil, weeded down to one after germination and grown in a growth chamber (Percival) in long-day conditions (20◦ C during the day and 18◦ C during the night with 16 hours of light) and 60% humidity (as described in (Schmitz et al. 2024)).

Statistical analysis

The statistical analysis and data visualization were performed in R (version 4.5.2) using the packages lsmeans (version2.30-2), emmeans (version 1.11.1), car (version 3.1-3), glmmLasso (version 1.6.3), and ggplot2 (version 3.5.2).

- Ex situ flowering time

To account for the experimental design, varying numbers of replicates, and both individual and population-level effects, flowering time estimates were derived using a generalized linear mixed-effects model (GLMM):

$$\text{glm}(\text{Floweringtime} \sim \text{Population/Individual} + \text{Tray}, f\,amily = \text{negativebinomial})$$

These estimates were then used to assess associations with environmental indices and habitat types via a Gaussian GLM:

$$\text{glm}(\text{Exsitu\_onsetFT} \sim \text{Variable1} + \ldots + \text{Variable} + \text{residuals})$$

Ex situ onset of flowering time was defined as the flowering of the 25% earliest individuals per population. The relationship between in situ and ex situ onsets of flowering time was modelled using a linear model (LM):

$$\text{lm}(\text{Insitu\_onsetFT} \sim \text{Exsitu\_onsetFT} \cdot \text{Habitat})$$

- Primary dormancy

Germination ratios (germinated seeds / total seeds) were calculated for each individual across the three experimental runs. Germination speed slopes were estimated using a GLM:

$$\text{glmer}\ ((\text{germinated\_seeds}, \text{non\_germinated\_seeds}) \sim \text{Week} + (\text{Week}|\ \text{Population}) + (\text{Week}|\ \text{Population} : \text{Individual}\ ), \text{family} = \text{binomial}\ )$$

The germination slope was extracted as the estimates of the (Week| Population) effect. Associations between these germination slopes and ecological variables were subsequently tested using a GLM, as described for ex situ flowering time.

- Correlation between the ex situ flowering time and the germination slope.

For each phenotype, models were first performed across all habitats and then separately for each habitat type, except for onset of flowering, for which habitat-specific analyses were not performed due to insufficient statistical power. As previously described, correlations between ex situ flowering time estimates and germination ratios estimates from the final experimental run were evaluated using Spearman's rank correlation test.

### Association between the environment and the genome

#### DNA extraction

Among the 168 transplanted populations, 74 populations containing more than 10 individuals were selected for pool sequencing. Between 10 and 20 individuals were randomly chosen from each population, and equivalent amounts of leaf material were sampled and pooled. In addition to the 74 F5 populations, three F2 populations consisting of 300 individuals each were also pooled as control. Genomic DNA was extracted from approximately 100 mg of frozen plant tissue using a cetyltrimethylammonium bromide (CTAB)-based protocol. Tissue was ground to a fine powder in liquid nitrogen and transferred to 2 mL microcentrifuge tubes. Samples were mixed with 700 µL of pre-warmed (65 °C) CTAB extraction buffer (100 mM Tris-HCl pH 8.0, 20 mM EDTA pH 8.0, 1.4 M NaCl, 2% w/v CTAB, 2% w/v PVP-40, and 0.5% β-mercaptoethanol added fresh) and 12 µL RNase A (10 mg/mL). The mixture was incubated at 65 °C for 30 minutes with occasional mixing, followed by centrifugation at maximum speed for 5 minutes at room temperature. The supernatant was transferred to a new tube.

An equal volume of chloroform:isoamyl alcohol (24:1) was added, mixed by vortexing for 10 seconds, and centrifuged at 12,000 × g for 10 minutes at room temperature. The aqueous phase was carefully transferred to a fresh tube. DNA was precipitated by adding 0.1 volumes of 3 M sodium acetate (pH 5.2) and 2.5 volumes of cold absolute ethanol, followed by gentle inversion. Samples were incubated at −20 °C for 2 hours to overnight and centrifuged at 13,000 rpm for 10 minutes at 4 °C. The resulting DNA pellet was washed with 500 µL of 70% ethanol, centrifuged for 5 minutes, air-dried, and resuspended in 50 µL TE buffer (10 mM Tris-HCl, 1 mM EDTA, pH 8.0).

Whole-genome sequencing (WGS) was conducted using Illumina short reads of 150 bp, with a coverage of 100x per pool.

#### Bioinformatic pipeline

We maintained reads with quality score (Q) above 30 (MultiQC v1.22.3). The PCR duplicates were removed using the assigned unique molecular identifier (UMI) and UMI-tools v1.1.6. Processed reads were then mapped to the *A. thaliana* TAIR10 reference genome using bwa-mem2 v2.2.1 and filtered with SAMtools v1.18. SNP calling was performed with BCFtools v1.19. Whole-genome sequencing data from both parental lines, MIL2 (ERR4592348 from the project: PRJEB40091 in the ENA base) and KAD1 (ERR4585755), were included in the SNP calling (Schmitz et al., 2024). Variants were filtered by a minimum mapping quality of 30, site quality of

40, and genotype quality (GQ) of 25. Indels and multi-allelic variants were removed to retain only biallelic SNPs.

Selecting segregating loci in the F2 populations

To concentrate on the most reliable SNP candidates, only exonic sites were used, which differed between the parents by at least one SNP, showed an allele frequency in the F2 pools between 0.4 and 0.6. and did not show missing data in the parental whole-genome sequences (Fig. S8). A total of 159,098 sites differentiating KAD1 and MIL2 were kept for the next analysis: 41,537 at chromosome 1, 20,487 at chromosome 2, 28,365 at chromosome 3, 29,109 at chromosome 4, and 39,600 at chromosome 5. We note that there was no sign of segregation distorsion or biased allelic transmission in this population.

Polarizing the MIL2 parental allele as reference allele

The sequencing reads were mapped to the *Arabidopsis thaliana* Col-0 reference genome. Because the reference allele did not consistently correspond to a single parental genotype, reference bias affected the orientation of alleles across loci. As part of the sites matched the MIL2 genotype and others the KAD1 genotype, the allelic polarization was inconsistent, leading to artificial blocks and random fluctuations in allele frequencies within short genomic regions in the F5 generation (Fig S9). The reference allele was polarized, such that the MIL2 allele was designated as reference and KAD1 as alternate (Fig S9).

Median allelic frequency per 50 kb windows

In order to identify genomic regions associated with the phenotypes and environmental factors, allele frequencies per pool were calculated as the alternate allele sequencing depth divided by total depth (AD/DP). Median alternate allele frequencies were then computed in 50 kb windows across chromosomes, taking into account the strong linkage. For each of the 68 populations retained, 1,971 non-overlapping 50 kb windows (each containing more than 10 SNPs) were computed: 502 on chromosome 1, 314 on chromosome 2, 376 on chromosome 3, 317 on chromosome 4, and 462 on chromosome 5. Pools containing plants that were not the offspring of the original MIL2xKAD1 F2 population could be easily discared, because median frequencies alternated across neighbouring 50kb windows (Fig. S10). In total, eight populations were discarded, and 68 populations were retained.

Statistical analysis

The statistical analysis and data visualization were performed in R (version 4.5.2) using the packages ggplot2 (version 3.5.2) and the function cor.test from the package stat.

Associations between ex situ phenotypes (flowering time, germination slope), environmental variables, and median allelic frequencies across 50 kb windows were evaluated using the following pipeline:

• 1. Only the windows with more than 10 SNP were retained.

• 2. For each window, Spearman's rank correlation was computed between population median allelic frequencies and the phenotype or environmental variable.

• 3. A randomization test was performed by shuffling the variable 1,000 times and recomputing Spearman's rank correlation each time.

Windows were deemed significant if the observed correlation coefficient (R) fell within the top 5% of the randomized R distribution and the p-value exceeded 0.01. QTL positions were mapped along the genome using −log10(p-value) when R was positive or log10(p-value) when R was negative; non-significant windows were annotated as neutral.

For each identified QTL region, the median allelic frequency from the most significant window across the 68 populations was correlated with and plotted against the corresponding variable.

Estimation of the FWER (Family Wise Error Rate)

To quantify our false positive rate, we simulated the experimental design, with a forward in time Wright Fisher individual-based model, for a set of 32 or 68 populations recombining independently for three generations under neutrality. The simulation starts from a F1 heterozygous individual. For each study site, we generated 1000 genotypes resulting from the F1 individual undergoing selfing, as the F2 starting seed pool. For each population, we sample Ns "seeds" without replacement from the 1000 genotypes, to simulate the bottleneck which followed the sowing throughout the city, Ns being the average population size measured at site s. We then sequentially generate F3, F4 and F5 generations. To generate seeds of generation Fi+1, we randomly choose with replacement Ns plants at generation Fi and we let them undergo selfing. We modelled a realistic recombination landscape, using the genetic distances from A. thaliana's recombination map taken from https://github.com/LohmuellerLab/arabidopsis_recomb_maps. All of the simulated genomes contain all the SNPs included in the VCF of the F5. The SNPs are grouped into windows of 50 kb, and a Spearman's rank correlation test is performed between the median frequency of each window and the environmental variable measured at each site (we chose Shannon index as test variable). This simulation framework under neutrality allows us to obtain the distribution of the minimum p-values for the Spearman rank correlation test across the genome under the null hypothesis. We then computed the p-value thresholds that yield a family-wise error rate (FWER) of 0.05, 0.1 and 0.25. These were determined as the corresponding quantile of the distribution of the 499 smallest p-value obtained from each of our 499 simulation replicates. We find the p-value thresholds to be 1.46e-4, 3.14e-4 and 1.17e-3 for the set of 32 populations, and 6.45e-4, 9.56e-4 and 3.19e-3 for the set of 66 populations, respectively for each target FWER.

Data availability

The data and R scripts are available at the github: https://github.com/jfloret23/Urban_niche_Ath

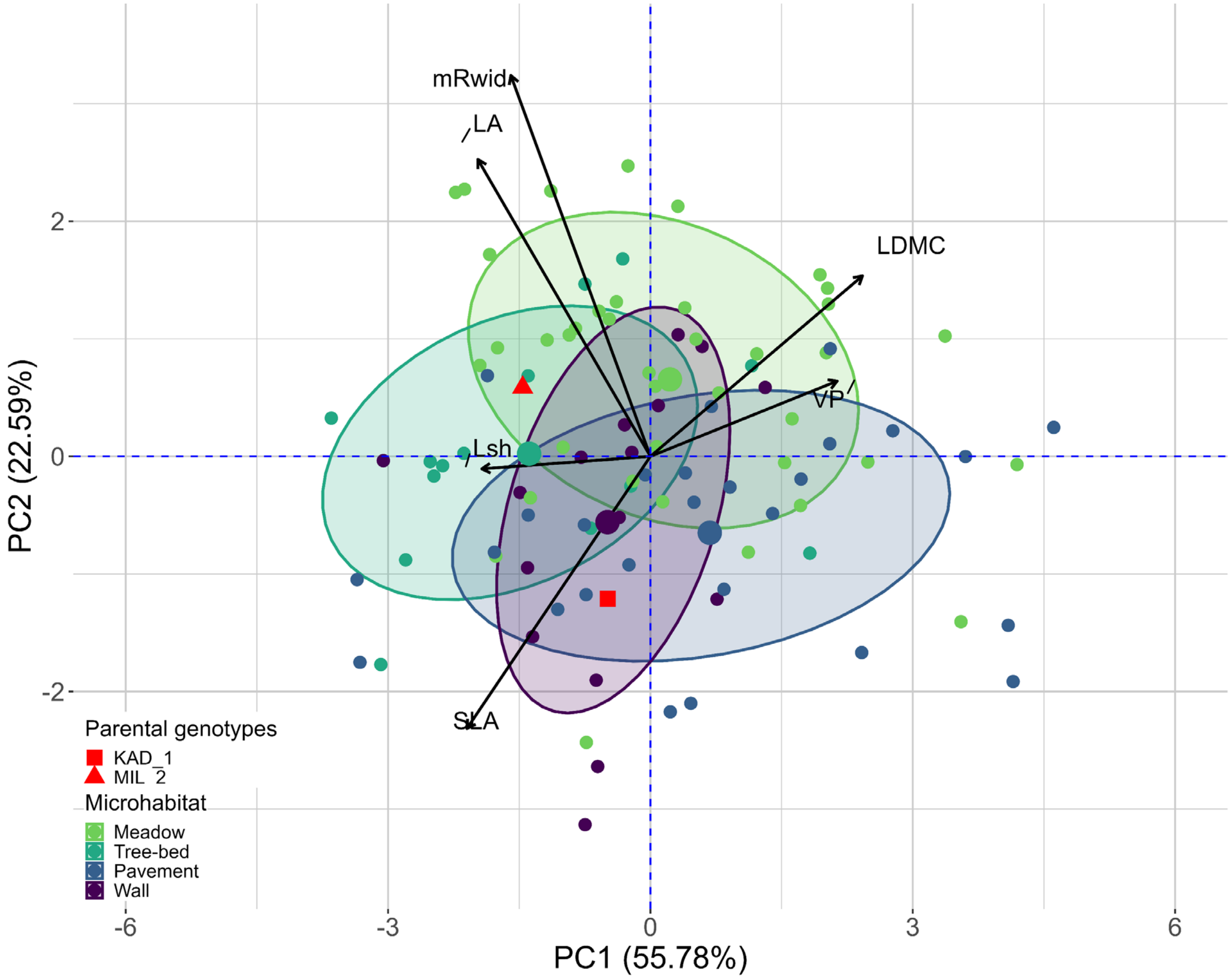

**Figure S1.** Divergent trait syndromes of the parental genotypes used in the city-wide colonization experiment relative to phenotypic variation among *Arabidopsis thaliana* genotypes from Cologne. Trait values were measured in a greenhouse common-garden experiment on 3–8 individuals per genotype representing 18 genotypes from four urban habitat types (meadow, tree bed, pavement, and wall). Mean values for the pavement-derived genotype KAD1 and the meadow-derived genotype MIL2, which served as parental genotypes in the colonization experiment, are marked in red. Traits include growth traits (mean rosette width, mRwid, cm; length of the longest shoot at seed maturation, Lsh, cm), the life-history trait vegetative period (VP, days; corresponds to flowering time), and resource-acquisition traits (leaf area, LA, mm$^2$; specific leaf area, SLA, mm$^2$ mg$^{-1}$; and leaf dry matter content, LDMC, %).

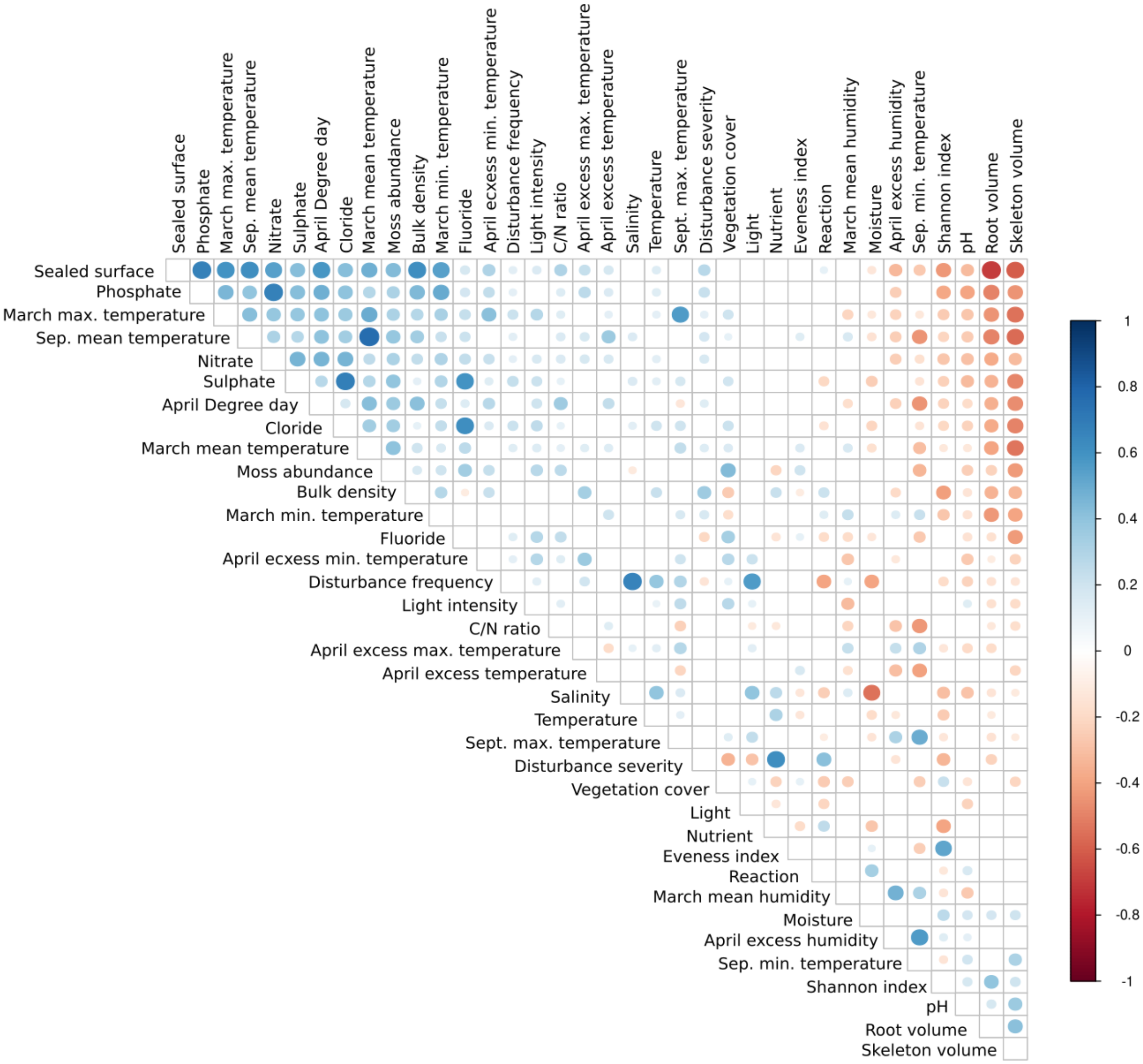

**Fig. S2.** Correlation coefficients between environmental factors used in the analysis. The ecological factors were measured at different key phenological stages of *A. thaliana*: between September 2021 and May 2022 across 393 sites in Cologne. Spearman's rank correlation coefficients were computed among these factors.

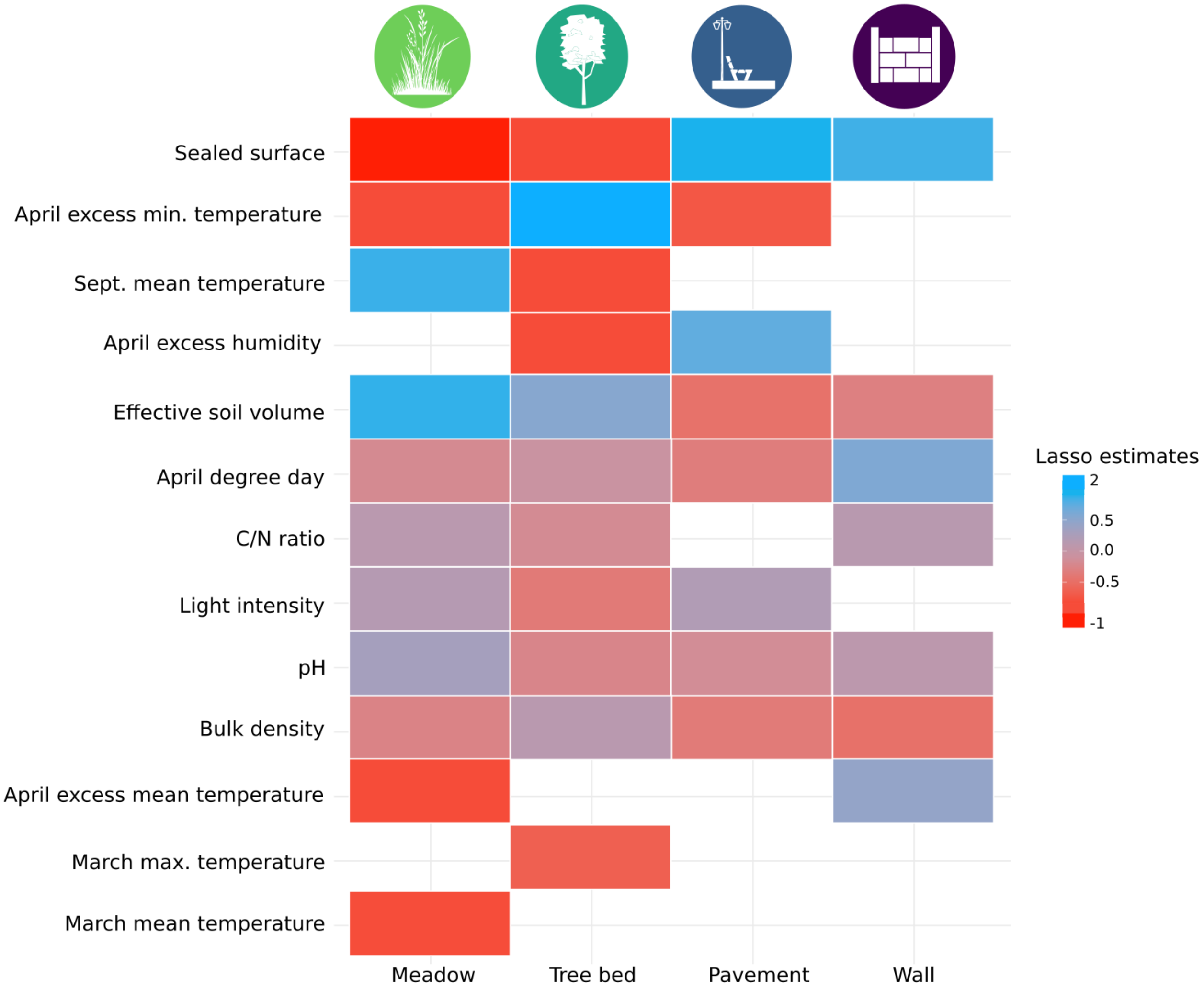

**Fig. S3.** Environmental factors characterizing the urban habitats. The ecological factors were measured at different key phenological stages of *A. thaliana* between September 2021 and May 2022 across 393 sites in Cologne. Lasso regression was first performed on the complete dataset, and then a GLM was fitted using the 10 most important variables selected by the Lasso. Only the significant variables from the GLM were retained.

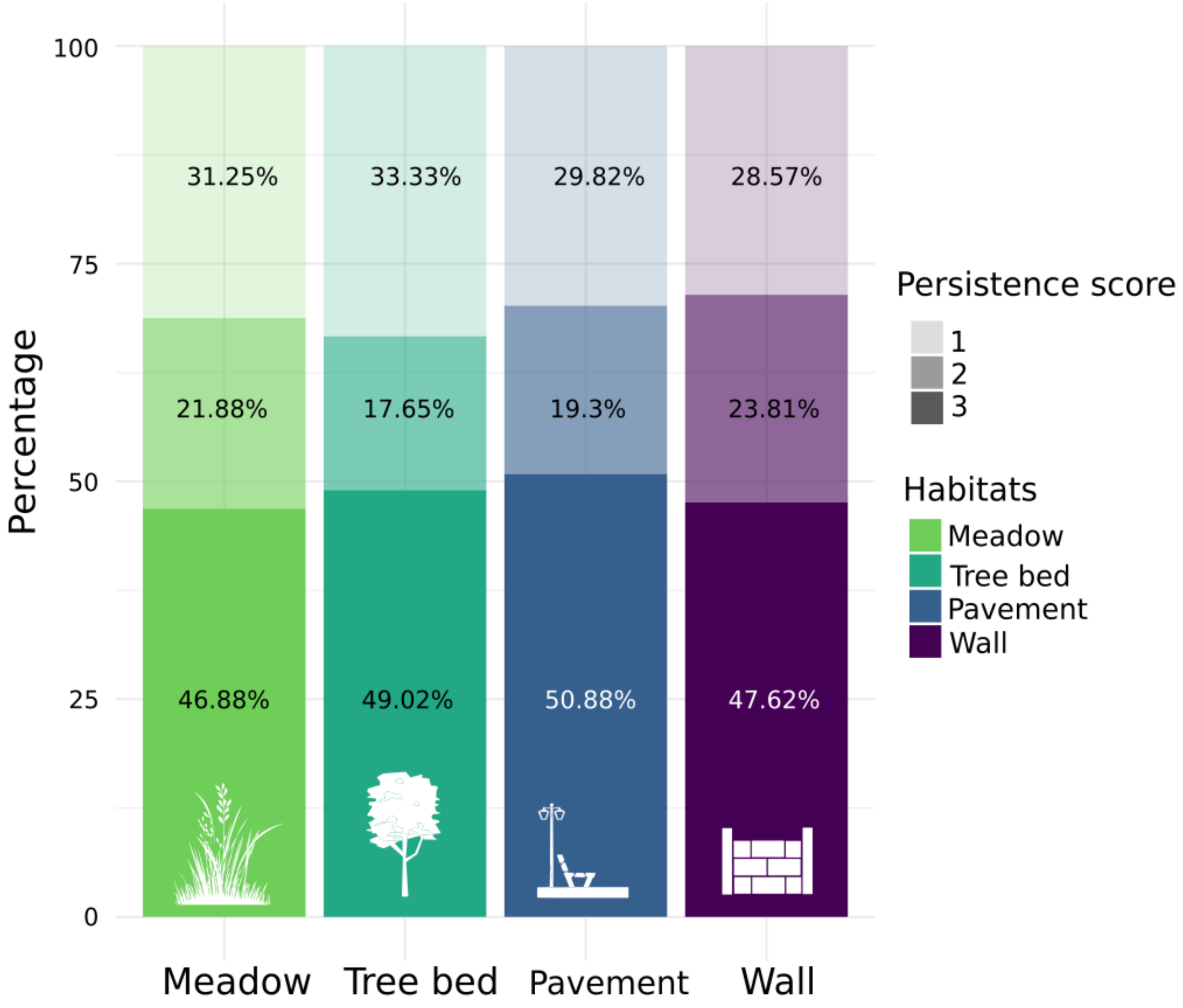

**Fig. S4.** Persistence score: quantifies the population's ability to survive across years. 1: the population germinated and flowered only in April 2022 (first generation) before extinction; 2: persistence for two generations (until April 2023); and 3: continued presence in April 2024 (third generation).

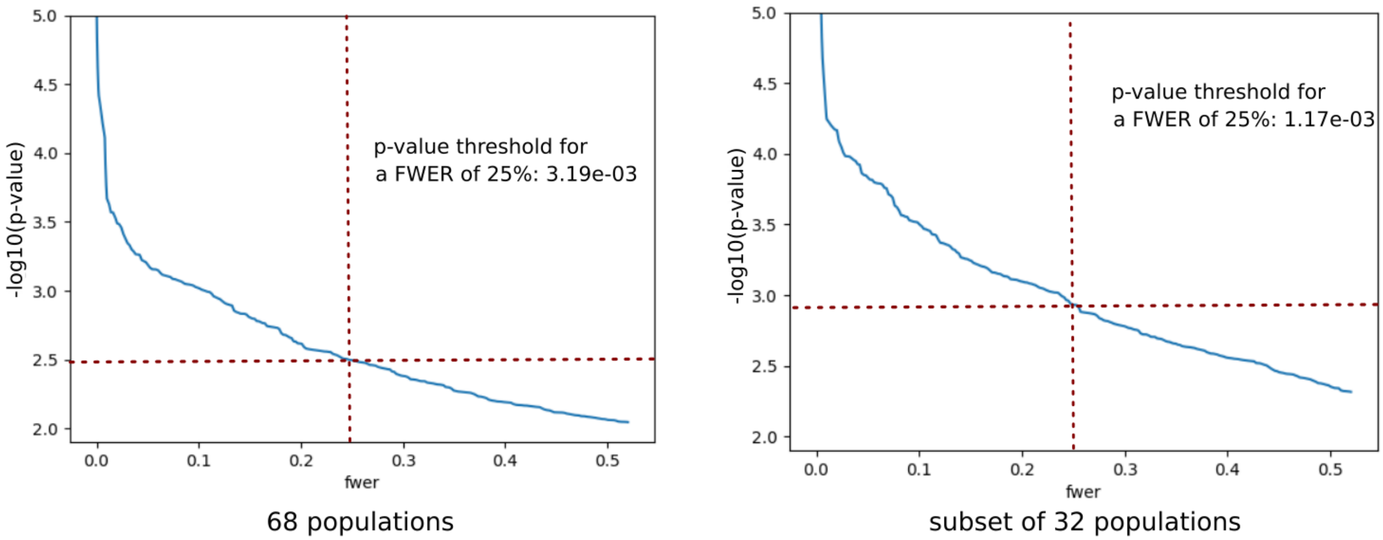

**Fig. S5.** Simulation of the experiment under neutrality, the FWER was computed from the output as the proportion of simulated replicates yielding one or more positive tests, among the Spearman's ranks correlation test performed. The model was realized with Shannon index as environmental variable, the p-value threshold was computed from 499 simulation replicates under neutralityas a function of FWER.

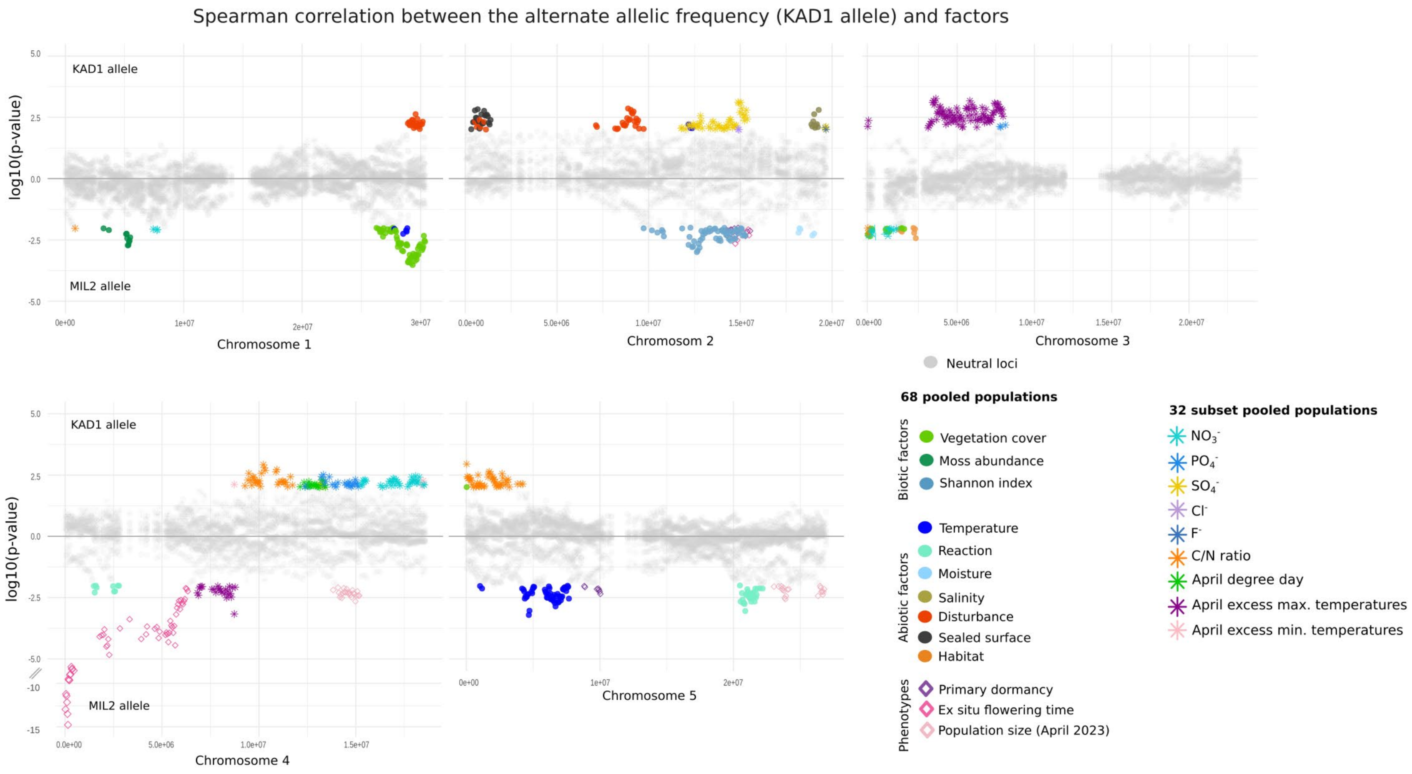

**Figure S6.** Genomic regions associated with phenotypes and environmental factors. Positive p-value indicate a positive correlation between the median KAD1 allele frequency and the factor, a negative p-value a correlation with the median MIL2 allele frequency. The color grey indicate a non-significant correlation

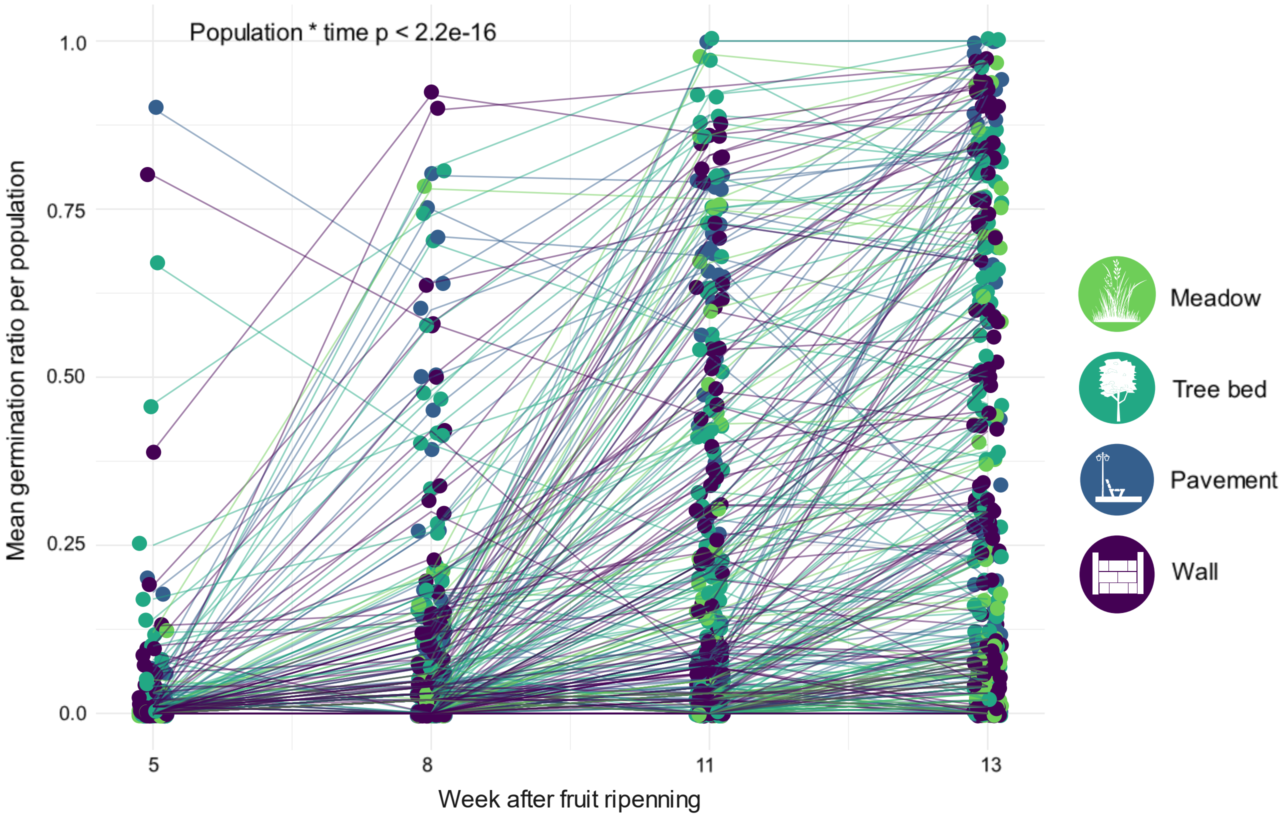

**Figure S7:** Release of primary dormancy across time in transplanted *A. thaliana* populations. Germination rate (number of germinated seeds relative to total seeds) was measured on the progeny of 359 transplanted individuals from 87 populations at different time points after fruit ripening (number of weeks).

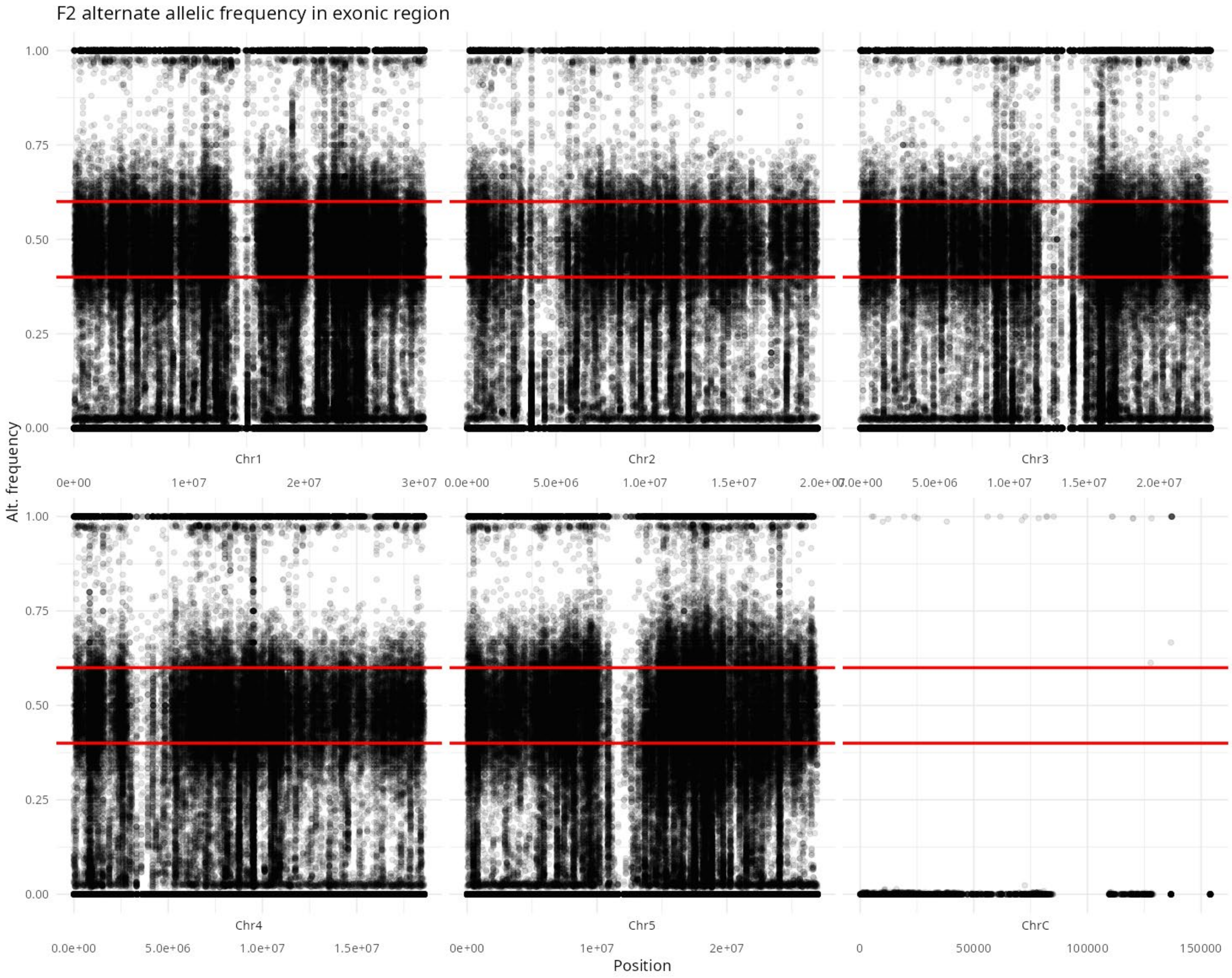

**Fig. S8.** Alternative allelic frequency of the F2 pools. The F2 pools (at least 300 individuals per pool) were mapped on the Col0 reference genome.

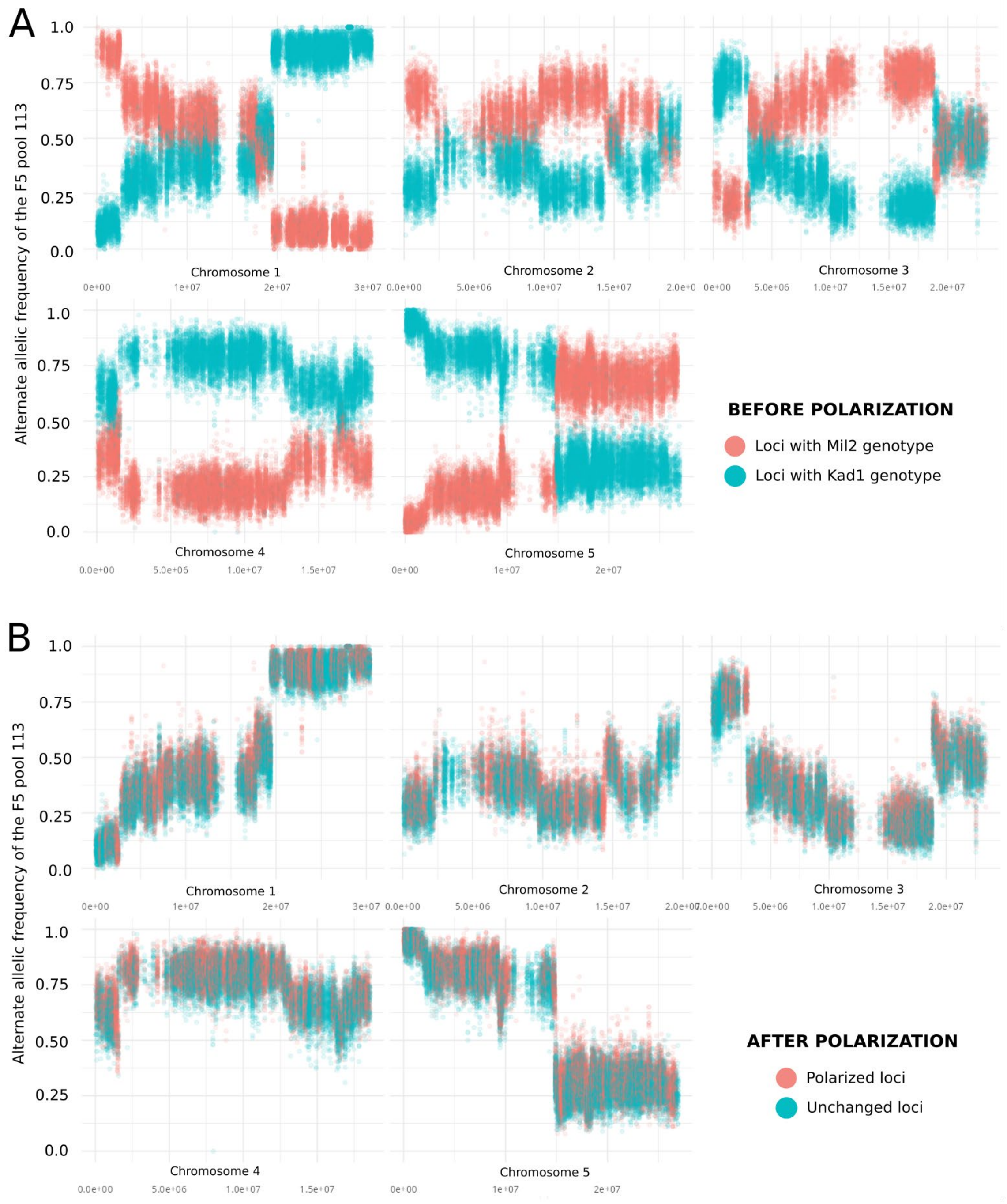

**Fig. S9.** Polarization of the reference allele in the F5 pools. Sequencing reads were mapped to the *Arabidopsis thaliana* Col0 reference genome. **A.** Example of alternate allele frequencies in an F5 pool before polarization; some sites matched the MIL2 genotype (pink) and the others KAD1 genotype (blue). **B.** The same F5 pool after polarization, with the genotype MIL2 considered as the reference.

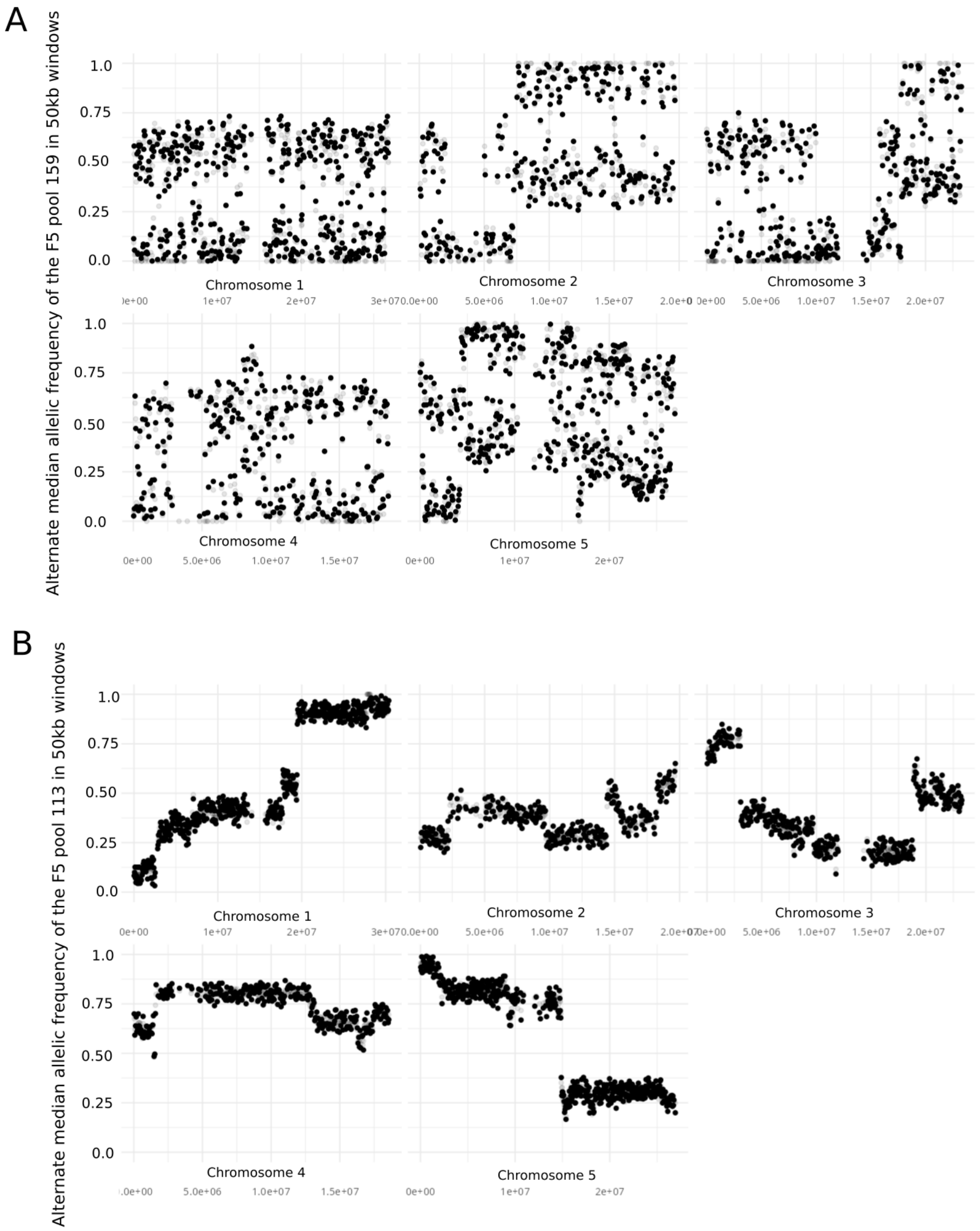

**Fig. S10**. Median allele frequencies in two exemplary F5 pools 113 and 159, calculated in 50 kb windows across the genome. Sequencing reads were mapped to the *A. thaliana* Col0 reference genome, and alleles were polarized so that the MIL2 genotype was considered as the reference. A. Example of a contaminated pool containing reads from other *A. thaliana* accessions, which was removed from the analysis. B. Example of an expected allelic frequency distribution in an F5 pool.

**Table S1: Association between genomic regions and population biology and environmental factors.**

**Table S2: Environmental factors influencing the size of *Arabidopsis thaliana* populations across urban habitats.** In September 2021, 1,000 seeds of an *A. thaliana* F2 population were sown at 393 sites across four urban habitat types in Cologne (Germany). Between April 2021 and June 2023, a suite of environmental variables were recorded at each site (Tables S1-3) or in selected sites (climatic variables; Table S4). Effects were analysed using a negative binomial generalized linear mixed model (glmm: Number_Individuals ~ Environmental_parameters + (1| Census_Years), family = negative.binomial), followed by Type II analysis of deviance tests.

**Table S3: Environmental factors influencing onset flowering time in *Arabidopsis thaliana* across urban habitats.** LASSO regression was used to identify environmental predictors of flowering onset in established populations (April 2023). Variables selected by the LASSO model were subsequently analysed using a generalized linear model (GLM) to estimate their effects.

**Table S4: Ex situ flowering time and environmental predictors of in situ flowering onset in Arabidopsis thaliana urban populations.** In situ flowering onset was recorded in April 2023 across 168 populations. Ex situ flowering time (FT) was measured under controlled conditions for 112 transplanted populations.

**Table S5: Surface characteristics and vegetation-derived environment variables measured across 393 experimental sites in the city of Cologne.** Surface characteristics were recorded at three key phenological stages of *A. thaliana*: seed sowing (September 2021), the end of the vegetative stage (February 2022), and fruit ripening (May 2022).

**Table S6: Plant species recorded at each site together with their corresponding Ellenberg indicator values (EIVs) and disturbance indices.** Only vascular plant species, excluding *Arabidopsis thaliana*, were considered. Community-weighted means of EIVs and disturbance indices were calculated from the relative cover of non-indifferent after standardizing their total cover to 100% (see Table S1).

**Table S7: Soil variables measured from topsoil samples (0-4.5 cm) collected in April 2022.** Soil depth was measured separately in July 2023.

**Table S8: Climatic variables measured at urban habitat patches.** For temperature and humidity, hygrochron data loggers (iButton, Whitewater, USA) were buried in soil (ca. 4 cm depth) at selected sites (n = 33).

Bonthoux, Sébastien, Lolita Voisin, Sabine Bouché-Pillon, and Simon Chollet. 2019. "More than Weeds: Spontaneous Vegetation in Streets as a Neglected Element of Urban Biodiversity." *Landscape and Urban Planning* 185 (May): 163–72. https://doi.org/10.1016/j.landurbplan.2019.02.009.

Diekmann, Martin. 2003. "Species Indicator Values as an Important Tool in Applied Plant Ecology – a Review." *Basic and Applied Ecology* 4 (6): 493–506. https://doi.org/10.1078/1439-1791-00185.

Ellenberg, H., H. Weber, R. Düll, Volkmar Wirth, Willy Werner, and D. Paulissen. 1991. "Zeigwerte von Pflanzen in MittelEuropa." *Scripta Geobotanica* 18 (January): 248.

Giuliani, Licida M., Paul D. Hallett, and Kenneth W. Loades. 2024. "Effects of Soil Structure Complexity to Root Growth of Plants with Contrasting Root Architecture." *Soil and Tillage Research* 238 (May): 106023. https://doi.org/10.1016/j.still.2024.106023.

Herben, Tomáš, Milan Chytrý, and Jitka Klimešová. 2016. "A Quest for Species-Level Indicator Values for Disturbance." *Journal of Vegetation Science* 27 (3): 628–36. https://doi.org/10.1111/jvs.12384.

Josse, Julie, and François Husson. 2016. "missMDA: A Package for Handling Missing Values in Multivariate Data Analysis." *Journal of Statistical Software* 70 (April): 1–31. https://doi.org/10.18637/jss.v070.i01.

Mäder, Patrick, David Boho, Michael Rzanny, et al. 2021. "The Flora Incognita App – Interactive Plant Species Identification." *Methods in Ecology and Evolution* 12 (7): 1335–42. https://doi.org/10.1111/2041-210X.13611.

Mclean, E. o. 1982. "Soil pH and Lime Requirement." In *Methods of Soil Analysis*. John Wiley & Sons, Ltd. https://doi.org/10.2134/agronmonogr9.2.2ed.c12.

Midolo, Gabriele, Tomáš Herben, Irena Axmanová, et al. 2023. "Disturbance Indicator Values for European Plants." *Global Ecology and Biogeography* 32 (1): 24–34. https://doi.org/10.1111/geb.13603.

Nowosad, J. 2019. "Pollen: Analysis of Aerobiological Data." *R Package Version 0.71*.

Pielou, E. C. 1966. "The Measurement of Diversity in Different Types of Biological Collections." *Journal of Theoretical Biology* 13 (December): 131–44. https://doi.org/10.1016/0022-5193(66)90013-0.

Schmitz, Gregor, Anja Linstädter, Anke S. K. Frank, et al. 2024. "Environmental Filtering of Life-History Trait Diversity in Urban Populations of Arabidopsis Thaliana." *Journal of Ecology* 112 (1): 14–27. https://doi.org/10.1111/1365-2745.14211.

Shannon, C. E. 1948. *A Mathematical Theory of Communication*.

Stekhoven, Daniel J., and Peter Bühlmann. 2012. “MissForest--Non-Parametric Missing Value Imputation for Mixed-Type Data.” *Bioinformatics (Oxford, England)* 28 (1): 112–18. https://doi.org/10.1093/bioinformatics/btr597.

### Dozens of genetic variants sustain adaptation to urban spatial heterogeneity in *Arabidopsis thaliana*

Authors: Justine Floret, Anja Linstädter, Huiyao Zhang, Vera Hesen, Swan Portalier, Lee Weinand, Gaelle Bustarret, Kirsten Bell, Fabrice Roux, Tahir Ali, Margarita Takou, Gregor Schmitz, Stanislav Kopriva, and Juliette de Meaux

**Table S1: Association between genomic regions, population biology and environmental factors.**

**Table S2: Environmental factors influencing the size of *Arabidopsis thaliana* populations across urban habitats.** In September 2021, 1,000 seeds of an *A. thaliana* F2 population were sown at 393 sites across four urban habitat types in Cologne (Germany). Between April 2021 and June 2023, a suite of environmental variables were recorded at each site (Tables S1-3) or in selected sites (climatic variables; Table S4). Effects were analysed using a negative binomial generalized linear mixed model (glmm: Number_Individuals ~ Environmental_parameters + (1| Census_Years), family = negative.binomial), followed by Type II analysis of deviance tests.

**Table S3: Environmental factors influencing in situ onset flowering time in *Arabidopsis thaliana* across urban habitats.** LASSO regression was used to identify environmental predictors of flowering onset in established populations (April 2023). Variables selected by the LASSO model were subsequently analysed using a generalized linear model (GLM) to estimate their effects.

**Table S4: Ex situ flowering time and environmental predictors of in situ flowering onset in Arabidopsis thaliana urban populations.** In situ flowering onset was recorded in April 2023 across 168 populations. Ex situ flowering time (FT) was measured under controlled conditions for 112 transplanted populations.

**TABLE S5: Surface characteristics and vegetation-derived environment variables measured across 393 experimental sites in the city of Cologne.** Surface characteristics were recorded at three key phenological stages of *A. thaliana*: seed sowing (September 2021), the end of the vegetative stage (February 2022), and fruit ripening (May 2022).

**Table S6: Plant species recorded at each site together with their corresponding Ellenberg indicator values (EIVs) and disturbance indices.** Only vascular plant species, excluding *Arabidopsis thaliana*, were considered. Community-weighted means of EIVs and disturbance indices were calculated from the relative cover of non-indifferent after standardizing their total cover to 100% (see Table S1).

**Table S7: Soil variables measured from topsoil samples (0-4.5 cm) collected in April 2022.** Soil depth was measured separately in July 2023.

**Table S8: Climatic variables measured at urban habitat patches.** For temperature and humidity, hygrochron data loggers (iButton, Whitewater, USA) were buried in soil (ca. 4 cm depth) at selected sites (n = 33).

**Table S1:** Association between genomic regions, population biology and environmental factors

| Chrm | Pos | Factor | pvalue | R | logpvalue | Category | Effect |
|---|---|---|---|---|---|---|---|
| Chr1 | 5275000 | Moss abundance | 1.89E-03 | -3.75E-01 | -2.72E+00 | Indexes | Negative |
| Chr1 | 5325000 | Moss abundance | 1.96E-03 | -3.74E-01 | -2.71E+00 | Indexes | Negative |
| Chr1 | 5375000 | Moss abundance | 2.58E-03 | -3.65E-01 | -2.59E+00 | Indexes | Negative |
| Chr1 | 5425000 | Moss abundance | 2.99E-03 | -3.60E-01 | -2.52E+00 | Indexes | Negative |
| Chr1 | 28075000 | Vegetation cover | 1.66E-03 | -3.80E-01 | -2.78E+00 | Indexes | Negative |
| Chr1 | 28125000 | Vegetation cover | 2.26E-03 | -3.69E-01 | -2.65E+00 | Indexes | Negative |
| Chr1 | 28175000 | Vegetation cover | 2.37E-03 | -3.68E-01 | -2.62E+00 | Indexes | Negative |
| Chr1 | 28225000 | Vegetation cover | 2.22E-03 | -3.70E-01 | -2.65E+00 | Indexes | Negative |
| Chr1 | 28275000 | Vegetation cover | 1.94E-03 | -3.75E-01 | -2.71E+00 | Indexes | Negative |
| Chr1 | 28325000 | Vegetation cover | 2.17E-03 | -3.71E-01 | -2.66E+00 | Indexes | Negative |
| Chr1 | 28375000 | Vegetation cover | 2.32E-03 | -3.69E-01 | -2.63E+00 | Indexes | Negative |
| Chr1 | 28425000 | Vegetation cover | 1.18E-03 | -3.91E-01 | -2.93E+00 | Indexes | Negative |
| Chr1 | 28525000 | Vegetation cover | 1.09E-03 | -3.93E-01 | -2.96E+00 | Indexes | Negative |
| Chr1 | 28775000 | Vegetation cover | 1.97E-03 | -3.74E-01 | -2.71E+00 | Indexes | Negative |
| Chr1 | 28825000 | Vegetation cover | 2.14E-03 | -3.71E-01 | -2.67E+00 | Indexes | Negative |
| Chr1 | 28875000 | Vegetation cover | 1.88E-03 | -3.76E-01 | -2.73E+00 | Indexes | Negative |
| Chr1 | 28925000 | Vegetation cover | 3.94E-04 | -4.24E-01 | -3.40E+00 | Indexes | Negative |
| Chr1 | 28975000 | Vegetation cover | 3.84E-04 | -4.24E-01 | -3.42E+00 | Indexes | Negative |
| Chr1 | 29025000 | Vegetation cover | 4.86E-04 | -4.18E-01 | -3.31E+00 | Indexes | Negative |
| Chr1 | 29075000 | Vegetation cover | 7.76E-04 | -4.04E-01 | -3.11E+00 | Indexes | Negative |
| Chr1 | 29125000 | Vegetation cover | 5.39E-04 | -4.15E-01 | -3.27E+00 | Indexes | Negative |
| Chr1 | 29175000 | Vegetation cover | 3.60E-04 | -4.26E-01 | -3.44E+00 | Indexes | Negative |
| Chr1 | 29225000 | Vegetation cover | 6.14E-04 | -4.11E-01 | -3.21E+00 | Indexes | Negative |
| Chr1 | 29275000 | Vegetation cover | 6.67E-04 | -4.08E-01 | -3.18E+00 | Indexes | Negative |
| Chr1 | 29325000 | Vegetation cover | 3.00E-04 | -4.31E-01 | -3.52E+00 | Indexes | Negative |
| Chr1 | 29375000 | Vegetation cover | 8.60E-04 | -4.01E-01 | -3.07E+00 | Indexes | Negative |
| Chr1 | 29425000 | Vegetation cover | 6.84E-04 | -4.07E-01 | -3.17E+00 | Indexes | Negative |
| Chr1 | 29475000 | Vegetation cover | 6.53E-04 | -4.09E-01 | -3.18E+00 | Indexes | Negative |
| Chr1 | 29525000 | Vegetation cover | 4.38E-04 | -4.21E-01 | -3.36E+00 | Indexes | Negative |
| Chr1 | 29575000 | Vegetation cover | 8.16E-04 | -4.02E-01 | -3.09E+00 | Indexes | Negative |
| Chr1 | 29575000 | Disturbance | 2.35E-03 | 3.74E-01 | 2.63E+00 | Indexes | Positive |
| Chr1 | 29625000 | Vegetation cover | 6.71E-04 | -4.08E-01 | -3.17E+00 | Indexes | Negative |
| Chr1 | 29675000 | Vegetation cover | 5.09E-04 | -4.16E-01 | -3.29E+00 | Indexes | Negative |
| Chr1 | 29725000 | Vegetation cover | 7.66E-04 | -4.04E-01 | -3.12E+00 | Indexes | Negative |
| Chr1 | 29775000 | Vegetation cover | 1.37E-03 | -3.86E-01 | -2.86E+00 | Indexes | Negative |
| Chr1 | 29825000 | Vegetation cover | 8.03E-04 | -4.03E-01 | -3.10E+00 | Indexes | Negative |
| Chr1 | 29875000 | Vegetation cover | 1.77E-03 | -3.78E-01 | -2.75E+00 | Indexes | Negative |
| Chr1 | 29925000 | Vegetation cover | 1.76E-03 | -3.78E-01 | -2.75E+00 | Indexes | Negative |
| Chr1 | 29975000 | Vegetation cover | 2.11E-03 | -3.72E-01 | -2.68E+00 | Indexes | Negative |
| Chr1 | 30025000 | Vegetation cover | 1.62E-03 | -3.81E-01 | -2.79E+00 | Indexes | Negative |
| Chr1 | 30125000 | Vegetation cover | 1.22E-03 | -3.90E-01 | -2.92E+00 | Indexes | Negative |
| Chr1 | 30175000 | Vegetation cover | 1.42E-03 | -3.85E-01 | -2.85E+00 | Indexes | Negative |
| Chr1 | 30225000 | Vegetation cover | 1.66E-03 | -3.80E-01 | -2.78E+00 | Indexes | Negative |
| Chr1 | 30325000 | Vegetation cover | 2.94E-03 | -3.61E-01 | -2.53E+00 | Indexes | Negative |
| Chr1 | 30375000 | Vegetation cover | 2.69E-03 | -3.64E-01 | -2.57E+00 | Indexes | Negative |
| Chr2 | 525000 | Sealed surface | 1.66E-03 | 3.80E-01 | 2.78E+00 | Indexes | Positive |
| Chr2 | 675000 | Sealed surface | 1.48E-03 | 3.83E-01 | 2.83E+00 | Indexes | Positive |
| Chr2 | 875000 | Sealed surface | 2.42E-03 | 3.67E-01 | 2.62E+00 | Indexes | Positive |
| Chr2 | 1025000 | Sealed surface | 1.72E-03 | 3.79E-01 | 2.76E+00 | Indexes | Positive |
| Chr2 | 1075000 | Sealed surface | 2.68E-03 | 3.64E-01 | 2.57E+00 | Indexes | Positive |
| Chr2 | 1125000 | Sealed surface | 2.62E-03 | 3.65E-01 | 2.58E+00 | Indexes | Positive |
| Chr2 | 1175000 | Sealed surface | 2.46E-03 | 3.67E-01 | 2.61E+00 | Indexes | Positive |
| Chr2 | 1225000 | Sealed surface | 2.88E-03 | 3.61E-01 | 2.54E+00 | Indexes | Positive |
| Chr2 | 8875000 | Disturbance | 1.38E-03 | 3.91E-01 | 2.86E+00 | Indexes | Positive |
| Chr2 | 9025000 | Disturbance | 1.57E-03 | 3.87E-01 | 2.80E+00 | Indexes | Positive |
| Chr2 | 9075000 | Disturbance | 2.04E-03 | 3.79E-01 | 2.69E+00 | Indexes | Positive |
| Chr2 | 9125000 | Disturbance | 2.24E-03 | 3.75E-01 | 2.65E+00 | Indexes | Positive |
| Chr2 | 9175000 | Disturbance | 1.83E-03 | 3.82E-01 | 2.74E+00 | Indexes | Positive |
| Chr2 | 12175000 | Shannon index | 1.99E-03 | -3.74E-01 | -2.70E+00 | Indexes | Negative |
| Chr2 | 12375000 | Shannon index | 1.62E-03 | -3.81E-01 | -2.79E+00 | Indexes | Negative |
| Chr2 | 12425000 | Shannon index | 1.70E-03 | -3.79E-01 | -2.77E+00 | Indexes | Negative |
| Chr2 | 12475000 | Shannon index | 2.80E-03 | -3.62E-01 | -2.55E+00 | Indexes | Negative |
| Chr2 | 12625000 | Shannon index | 1.01E-03 | -3.96E-01 | -3.00E+00 | Indexes | Negative |
| Chr2 | 12675000 | Shannon index | 1.15E-03 | -3.92E-01 | -2.94E+00 | Indexes | Negative |
| Chr2 | 12725000 | Shannon index | 2.21E-03 | -3.70E-01 | -2.66E+00 | Indexes | Negative |
| Chr2 | 12775000 | Shannon index | 1.53E-03 | -3.83E-01 | -2.82E+00 | Indexes | Negative |

| | | | | | | | |
|---|---|---|---|---|---|---|---|
| Chr2 | 12825000 | Shannon index | 3.10E-03 | -3.59E-01 | -2.51E+00 | Indexes | Negative |
| Chr2 | 13825000 | Shannon index | 3.01E-03 | -3.60E-01 | -2.52E+00 | Indexes | Negative |
| Chr2 | 13925000 | Shannon index | 2.88E-03 | -3.61E-01 | -2.54E+00 | Indexes | Negative |
| Chr2 | 14475000 | Shannon index | 2.91E-03 | -3.61E-01 | -2.54E+00 | Indexes | Negative |
| Chr2 | 14725000 | ex situ Flowering time | 2.24E-03 | -3.65E-01 | -2.65E+00 | Phenotype | Negative |
| Chr2 | 14825000 | ex situ Flowering time | 3.12E-03 | -3.53E-01 | -2.51E+00 | Phenotype | Negative |
| Chr2 | 14875000 | SO4 | 7.70E-04 | 5.64E-01 | 3.11E+00 | Factors | Positive |
| Chr2 | 14925000 | SO4 | 8.40E-04 | 5.61E-01 | 3.08E+00 | Factors | Positive |
| Chr2 | 15025000 | SO4 | 7.85E-04 | 5.63E-01 | 3.11E+00 | Factors | Positive |
| Chr2 | 19025000 | Salinity | 2.35E-03 | 3.74E-01 | 2.63E+00 | Indexes | Positive |
| Chr2 | 19275000 | Salinity | 1.58E-03 | 3.87E-01 | 2.80E+00 | Indexes | Positive |
| Chr3 | 3425000 | April excess max. temperature | 7.91E-04 | 5.63E-01 | 3.10E+00 | Factors | Positive |
| Chr3 | 3475000 | April excess max. temperature | 6.61E-04 | 5.70E-01 | 3.18E+00 | Factors | Positive |
| Chr3 | 3525000 | April excess max. temperature | 9.83E-04 | 5.55E-01 | 3.01E+00 | Factors | Positive |
| Chr3 | 3575000 | April excess max. temperature | 7.71E-04 | 5.64E-01 | 3.11E+00 | Factors | Positive |
| Chr3 | 3625000 | April excess max. temperature | 5.47E-04 | 5.77E-01 | 3.26E+00 | Factors | Positive |
| Chr3 | 3875000 | April excess max. temperature | 1.23E-03 | 5.46E-01 | 2.91E+00 | Factors | Positive |
| Chr3 | 3925000 | April excess max. temperature | 1.16E-03 | 5.48E-01 | 2.94E+00 | Factors | Positive |
| Chr3 | 5725000 | April excess max. temperature | 1.19E-03 | 5.47E-01 | 2.92E+00 | Factors | Positive |
| Chr3 | 5825000 | April excess max. temperature | 6.73E-04 | 5.69E-01 | 3.17E+00 | Factors | Positive |
| Chr3 | 5975000 | April excess max. temperature | 9.64E-04 | 5.56E-01 | 3.02E+00 | Factors | Positive |
| Chr3 | 7425000 | April excess max. temperature | 8.32E-04 | 5.61E-01 | 3.08E+00 | Factors | Positive |
| Chr3 | 7475000 | April excess max. temperature | 9.08E-04 | 5.58E-01 | 3.04E+00 | Factors | Positive |
| Chr3 | 7625000 | April excess max. temperature | 8.86E-04 | 5.59E-01 | 3.05E+00 | Factors | Positive |
| Chr3 | 7675000 | April excess max. temperature | 9.39E-04 | 5.57E-01 | 3.03E+00 | Factors | Positive |
| Chr4 | 25000 | ex situ Flowering time | 7.17E-13 | -7.38E-01 | -1.21E+01 | Phenotype | Negative |
| Chr4 | 75000 | ex situ Flowering time | 6.80E-13 | -7.38E-01 | -1.22E+01 | Phenotype | Negative |
| Chr4 | 125000 | ex situ Flowering time | 6.41E-13 | -7.39E-01 | -1.22E+01 | Phenotype | Negative |
| Chr4 | 175000 | ex situ Flowering time | 1.90E-13 | -7.50E-01 | -1.27E+01 | Phenotype | Negative |
| Chr4 | 225000 | ex situ Flowering time | 2.14E-14 | -7.68E-01 | -1.37E+01 | Phenotype | Negative |
| Chr4 | 275000 | ex situ Flowering time | 2.71E-15 | -7.84E-01 | -1.46E+01 | Phenotype | Negative |
| Chr4 | 325000 | ex situ Flowering time | 2.14E-10 | -6.78E-01 | -9.67E+00 | Phenotype | Negative |
| Chr4 | 375000 | ex situ Flowering time | 1.89E-10 | -6.79E-01 | -9.72E+00 | Phenotype | Negative |
| Chr4 | 425000 | ex situ Flowering time | 1.75E-10 | -6.80E-01 | -9.76E+00 | Phenotype | Negative |
| Chr4 | 475000 | ex situ Flowering time | 4.65E-10 | -6.69E-01 | -9.33E+00 | Phenotype | Negative |
| Chr4 | 525000 | ex situ Flowering time | 5.47E-10 | -6.67E-01 | -9.26E+00 | Phenotype | Negative |
| Chr4 | 625000 | ex situ Flowering time | 2.17E-09 | -6.49E-01 | -8.66E+00 | Phenotype | Negative |
| Chr4 | 675000 | ex situ Flowering time | 1.46E-09 | -6.54E-01 | -8.84E+00 | Phenotype | Negative |
| Chr4 | 725000 | ex situ Flowering time | 1.40E-09 | -6.55E-01 | -8.85E+00 | Phenotype | Negative |
| Chr4 | 775000 | ex situ Flowering time | 8.30E-09 | -6.31E-01 | -8.08E+00 | Phenotype | Negative |
| Chr4 | 825000 | ex situ Flowering time | 7.04E-09 | -6.33E-01 | -8.15E+00 | Phenotype | Negative |
| Chr4 | 875000 | ex situ Flowering time | 1.06E-09 | -6.58E-01 | -8.98E+00 | Phenotype | Negative |
| Chr4 | 925000 | ex situ Flowering time | 1.17E-08 | -6.26E-01 | -7.93E+00 | Phenotype | Negative |
| Chr4 | 975000 | ex situ Flowering time | 2.08E-08 | -6.17E-01 | -7.68E+00 | Phenotype | Negative |
| Chr4 | 1025000 | ex situ Flowering time | 2.73E-08 | -6.13E-01 | -7.56E+00 | Phenotype | Negative |
| Chr4 | 1075000 | ex situ Flowering time | 2.98E-08 | -6.12E-01 | -7.53E+00 | Phenotype | Negative |
| Chr4 | 1125000 | ex situ Flowering time | 1.46E-08 | -6.22E-01 | -7.84E+00 | Phenotype | Negative |
| Chr4 | 1175000 | ex situ Flowering time | 1.24E-08 | -6.25E-01 | -7.91E+00 | Phenotype | Negative |
| Chr4 | 1225000 | ex situ Flowering time | 1.44E-08 | -6.23E-01 | -7.84E+00 | Phenotype | Negative |
| Chr4 | 1275000 | ex situ Flowering time | 1.53E-08 | -6.22E-01 | -7.82E+00 | Phenotype | Negative |
| Chr4 | 1325000 | ex situ Flowering time | 7.14E-08 | -5.98E-01 | -7.15E+00 | Phenotype | Negative |
| Chr4 | 1375000 | ex situ Flowering time | 1.47E-07 | -5.87E-01 | -6.83E+00 | Phenotype | Negative |
| Chr4 | 1425000 | ex situ Flowering time | 2.20E-07 | -5.80E-01 | -6.66E+00 | Phenotype | Negative |
| Chr4 | 1475000 | ex situ Flowering time | 1.88E-07 | -5.82E-01 | -6.73E+00 | Phenotype | Negative |
| Chr4 | 1525000 | ex situ Flowering time | 4.21E-07 | -5.69E-01 | -6.38E+00 | Phenotype | Negative |
| Chr4 | 1575000 | ex situ Flowering time | 1.35E-07 | -5.88E-01 | -6.87E+00 | Phenotype | Negative |
| Chr4 | 1625000 | ex situ Flowering time | 1.75E-07 | -5.84E-01 | -6.76E+00 | Phenotype | Negative |
| Chr4 | 1775000 | ex situ Flowering time | 8.11E-05 | -4.59E-01 | -4.09E+00 | Phenotype | Negative |
| Chr4 | 1875000 | ex situ Flowering time | 9.25E-05 | -4.56E-01 | -4.03E+00 | Phenotype | Negative |
| Chr4 | 1975000 | ex situ Flowering time | 9.62E-05 | -4.55E-01 | -4.02E+00 | Phenotype | Negative |
| Chr4 | 2025000 | ex situ Flowering time | 1.60E-04 | -4.42E-01 | -3.80E+00 | Phenotype | Negative |

| | | | | | | | |
|---|---|---|---|---|---|---|---|
| Chr4 | 2125000 | ex situ Flowering time | 3.18E-05 | -4.82E-01 | -4.50E+00 | Phenotype | Negative |
| Chr4 | 2175000 | ex situ Flowering time | 3.57E-05 | -4.79E-01 | -4.45E+00 | Phenotype | Negative |
| Chr4 | 2225000 | ex situ Flowering time | 6.42E-05 | -4.65E-01 | -4.19E+00 | Phenotype | Negative |
| Chr4 | 2275000 | ex situ Flowering time | 1.46E-05 | -4.99E-01 | -4.83E+00 | Phenotype | Negative |
| Chr4 | 2325000 | ex situ Flowering time | 4.66E-06 | -5.23E-01 | -5.33E+00 | Phenotype | Negative |
| Chr4 | 2375000 | ex situ Flowering time | 3.65E-06 | -5.28E-01 | -5.44E+00 | Phenotype | Negative |
| Chr4 | 2425000 | ex situ Flowering time | 6.24E-07 | -5.62E-01 | -6.20E+00 | Phenotype | Negative |
| Chr4 | 2475000 | ex situ Flowering time | 7.67E-07 | -5.58E-01 | -6.12E+00 | Phenotype | Negative |
| Chr4 | 2525000 | ex situ Flowering time | 9.60E-07 | -5.54E-01 | -6.02E+00 | Phenotype | Negative |
| Chr4 | 2575000 | ex situ Flowering time | 1.06E-06 | -5.52E-01 | -5.98E+00 | Phenotype | Negative |
| Chr4 | 2625000 | ex situ Flowering time | 9.69E-08 | -5.93E-01 | -7.01E+00 | Phenotype | Negative |
| Chr4 | 2675000 | ex situ Flowering time | 3.71E-07 | -5.71E-01 | -6.43E+00 | Phenotype | Negative |
| Chr4 | 2725000 | ex situ Flowering time | 1.79E-07 | -5.83E-01 | -6.75E+00 | Phenotype | Negative |
| Chr4 | 2775000 | ex situ Flowering time | 6.23E-08 | -6.00E-01 | -7.21E+00 | Phenotype | Negative |
| Chr4 | 2825000 | ex situ Flowering time | 1.76E-04 | -4.40E-01 | -3.76E+00 | Phenotype | Negative |
| Chr4 | 3325000 | ex situ Flowering time | 4.15E-04 | -4.16E-01 | -3.38E+00 | Phenotype | Negative |
| Chr4 | 3925000 | ex situ Flowering time | 6.50E-05 | -4.65E-01 | -4.19E+00 | Phenotype | Negative |
| Chr4 | 4175000 | ex situ Flowering time | 1.01E-04 | -4.54E-01 | -4.00E+00 | Phenotype | Negative |
| Chr4 | 4225000 | ex situ Flowering time | 2.10E-04 | -4.35E-01 | -3.68E+00 | Phenotype | Negative |
| Chr4 | 4625000 | ex situ Flowering time | 1.70E-04 | -4.41E-01 | -3.77E+00 | Phenotype | Negative |
| Chr4 | 4725000 | ex situ Flowering time | 1.99E-04 | -4.36E-01 | -3.70E+00 | Phenotype | Negative |
| Chr4 | 4825000 | ex situ Flowering time | 7.11E-05 | -4.63E-01 | -4.15E+00 | Phenotype | Negative |
| Chr4 | 4875000 | ex situ Flowering time | 1.30E-04 | -4.48E-01 | -3.89E+00 | Phenotype | Negative |
| Chr4 | 5175000 | ex situ Flowering time | 9.54E-05 | -4.55E-01 | -4.02E+00 | Phenotype | Negative |
| Chr4 | 5225000 | ex situ Flowering time | 1.20E-04 | -4.49E-01 | -3.92E+00 | Phenotype | Negative |
| Chr4 | 5275000 | ex situ Flowering time | 1.03E-04 | -4.54E-01 | -3.99E+00 | Phenotype | Negative |
| Chr4 | 5325000 | ex situ Flowering time | 4.86E-05 | -4.72E-01 | -4.31E+00 | Phenotype | Negative |
| Chr4 | 5375000 | ex situ Flowering time | 1.17E-04 | -4.50E-01 | -3.93E+00 | Phenotype | Negative |
| Chr4 | 5425000 | ex situ Flowering time | 3.53E-04 | -4.21E-01 | -3.45E+00 | Phenotype | Negative |
| Chr4 | 5475000 | ex situ Flowering time | 2.31E-04 | -4.32E-01 | -3.64E+00 | Phenotype | Negative |
| Chr4 | 5525000 | ex situ Flowering time | 1.98E-04 | -4.36E-01 | -3.70E+00 | Phenotype | Negative |
| Chr4 | 5575000 | ex situ Flowering time | 1.16E-04 | -4.50E-01 | -3.94E+00 | Phenotype | Negative |
| Chr4 | 5625000 | ex situ Flowering time | 2.22E-04 | -4.33E-01 | -3.65E+00 | Phenotype | Negative |
| Chr4 | 5675000 | ex situ Flowering time | 3.59E-05 | -4.79E-01 | -4.44E+00 | Phenotype | Negative |
| Chr4 | 5725000 | ex situ Flowering time | 6.51E-04 | -4.03E-01 | -3.19E+00 | Phenotype | Negative |
| Chr4 | 5775000 | ex situ Flowering time | 1.21E-03 | -3.84E-01 | -2.92E+00 | Phenotype | Negative |
| Chr4 | 5825000 | ex situ Flowering time | 1.70E-03 | -3.74E-01 | -2.77E+00 | Phenotype | Negative |
| Chr4 | 5875000 | ex situ Flowering time | 2.35E-03 | -3.63E-01 | -2.63E+00 | Phenotype | Negative |
| Chr4 | 5925000 | ex situ Flowering time | 1.06E-03 | -3.88E-01 | -2.97E+00 | Phenotype | Negative |
| Chr4 | 5975000 | ex situ Flowering time | 2.54E-03 | -3.60E-01 | -2.59E+00 | Phenotype | Negative |
| Chr4 | 6025000 | ex situ Flowering time | 1.88E-03 | -3.70E-01 | -2.73E+00 | Phenotype | Negative |
| Chr4 | 6075000 | ex situ Flowering time | 1.67E-03 | -3.74E-01 | -2.78E+00 | Phenotype | Negative |
| Chr4 | 6125000 | ex situ Flowering time | 2.28E-03 | -3.64E-01 | -2.64E+00 | Phenotype | Negative |
| Chr4 | 6175000 | ex situ Flowering time | 2.77E-03 | -3.57E-01 | -2.56E+00 | Phenotype | Negative |
| Chr4 | 8725000 | April excess max. temperature | 6.79E-04 | -5.69E-01 | -3.17E+00 | Factors | Negative |
| Chr4 | 10225000 | C/N ratio | 1.19E-03 | 5.47E-01 | 2.92E+00 | Factors | Positive |
| Chr4 | 14275000 | Population size 2023 | 3.10E-03 | -3.59E-01 | -2.51E+00 | Phenotype | Negative |
| Chr4 | 14975000 | Population size 2023 | 2.24E-03 | -3.70E-01 | -2.65E+00 | Phenotype | Negative |
| Chr5 | 25000 | C/N ratio | 1.13E-03 | 5.49E-01 | 2.95E+00 | Factors | Positive |
| Chr5 | 4375000 | Temperature | 2.88E-03 | -3.67E-01 | -2.54E+00 | Indexes | Negative |
| Chr5 | 4675000 | Temperature | 6.20E-04 | -4.16E-01 | -3.21E+00 | Indexes | Negative |
| Chr5 | 4725000 | Temperature | 9.23E-04 | -4.04E-01 | -3.03E+00 | Indexes | Negative |
| Chr5 | 6075000 | Temperature | 3.13E-03 | -3.64E-01 | -2.50E+00 | Indexes | Negative |
| Chr5 | 6325000 | Temperature | 2.69E-03 | -3.69E-01 | -2.57E+00 | Indexes | Negative |
| Chr5 | 6375000 | Temperature | 2.80E-03 | -3.68E-01 | -2.55E+00 | Indexes | Negative |
| Chr5 | 6475000 | Temperature | 2.64E-03 | -3.70E-01 | -2.58E+00 | Indexes | Negative |
| Chr5 | 6525000 | Temperature | 2.30E-03 | -3.74E-01 | -2.64E+00 | Indexes | Negative |
| Chr5 | 6625000 | Temperature | 1.91E-03 | -3.81E-01 | -2.72E+00 | Indexes | Negative |
| Chr5 | 6675000 | Temperature | 1.39E-03 | -3.91E-01 | -2.86E+00 | Indexes | Negative |
| Chr5 | 6775000 | Temperature | 2.38E-03 | -3.73E-01 | -2.62E+00 | Indexes | Negative |
| Chr5 | 6875000 | Temperature | 1.99E-03 | -3.79E-01 | -2.70E+00 | Indexes | Negative |
| Chr5 | 6925000 | Temperature | 2.11E-03 | -3.77E-01 | -2.68E+00 | Indexes | Negative |
| Chr5 | 7075000 | Temperature | 2.47E-03 | -3.72E-01 | -2.61E+00 | Indexes | Negative |
| Chr5 | 7675000 | Temperature | 2.88E-03 | -3.67E-01 | -2.54E+00 | Indexes | Negative |
| Chr5 | 20525000 | Reaction | 2.20E-03 | -3.76E-01 | -2.66E+00 | Indexes | Negative |
| Chr5 | 20675000 | Reaction | 2.62E-03 | -3.70E-01 | -2.58E+00 | Indexes | Negative |
| Chr5 | 20725000 | Reaction | 1.84E-03 | -3.82E-01 | -2.74E+00 | Indexes | Negative |
| Chr5 | 20775000 | Reaction | 1.97E-03 | -3.80E-01 | -2.71E+00 | Indexes | Negative |
| Chr5 | 20825000 | Reaction | 1.74E-03 | -3.84E-01 | -2.76E+00 | Indexes | Negative |
| Chr5 | 20875000 | Reaction | 8.99E-04 | -4.05E-01 | -3.05E+00 | Indexes | Negative |
| Chr5 | 21225000 | Reaction | 2.94E-03 | -3.66E-01 | -2.53E+00 | Indexes | Negative |
| Chr5 | 21275000 | Reaction | 2.91E-03 | -3.66E-01 | -2.54E+00 | Indexes | Negative |
| Chr5 | 21325000 | Reaction | 2.16E-03 | -3.77E-01 | -2.66E+00 | Indexes | Negative |

| | | | | | | | |
|---|---|---|---|---|---|---|---|
| Chr5 | 21575000 | Reaction | 2.17E-03 | -3.76E-01 | -2.66E+00 | Indexes | Negative |
| Chr5 | 23825000 | Population size 2023 | 3.13E-03 | -3.58E-01 | -2.50E+00 | Phenotype | Negative |
| Chr5 | 23875000 | Population size 2023 | 2.74E-03 | -3.63E-01 | -2.56E+00 | Phenotype | Negative |

**Table S2: Environmental factors influencing the size of *Arabidopsis thaliana* populations across urban habitats.** In September 2021, 1,000 seeds of an *A. thaliana* F2 population were sown at 393 sites across four urban habitat types in Cologne (Germany). Between April 2021 and June 2023, a suite of environmental variables were recorded at each site (Tables S1-3) or in selected sites (climatic variables; Table S4). Effects were analysed using a negative binomial generalized linear mixed model (glmm: Number_Individuals ~ Environmental_parameters + (1| Census_Years), family = negative.binomial), followed by Type II analysis of deviance tests.

| Habitat | Variable | Df | glm estimate | p-value | χ2 | p-value(χ2) | |
|---|---|---|---|---|---|---|---|
| Meadow | April min. temperature | 1 | 13.54 | 8.96E-03 | 6.83 | 8.96E-03 | ** |
| | Sealed surface | 1 | 6.35 | 3.80E-03 | 8.38 | 3.80E-03 | ** |
| | March mean temperature | 1 | 2.42 | 2.01E-02 | 5.4 | 2.01E-02 | * |
| | March mean humidity | 1 | 0.3 | 1.64E-03 | 9.91 | 1.64E-03 | ** |
| | Light intensity | 1 | 1.20E-03 | 5.76E-04 | 11.85 | 5.76E-04 | *** |
| | Disturbance frequency | 1 | -1.95 | 2.94E-02 | 4.75 | 2.94E-02 | * |
| | March max. temperature | 1 | -2.66 | 1.31E-02 | 6.16 | 1.31E-02 | * |
| | Residuals | 152 | | | | | |
| Tree bed | Light intensity | 1 | -1.41E-03 | 7.94E-03 | 7.05 | 7.94E-03 | ** |
| | Residuals | 104 | | | | | |
| Pavement | Sealed surface | 1 | 6.27 | 3.04E-02 | 4.68 | 3.04E-02 | * |
| | Light intensity | 1 | 1.17E-03 | 2.49E-02 | 5.03 | 2.49E-02 | * |
| | March max. temperature | 1 | -0.48 | 3.13E-02 | 4.64 | 3.13E-02 | * |
| | Residuals | 137 | | | | | |
| Wall | Disturbance severity | 1 | 50.49 | 2.80E-02 | 4.83 | 2.80E-02 | * |
| | Disturbance frequency | 1 | 5.52 | 2.31E-02 | 5.16 | 2.31E-02 | * |
| | mean Reaction | 1 | -7.51 | 3.00E-03 | 8.8 | 3.00E-03 | ** |
| | Residuals | 107 | | | | | |

**Table S3: Environmental factors influencing onset flowering time in *Arabidopsis thaliana* across urban habitats.** LASSO regression was used to identify environmental predictors of flowering onset in established populations (April 2023). Variables selected by the LASSO model were subsequently analysed using a generalized linear model (GLM) to estimate their effects.

| Variables | Df | glm estimate | p-value | χ2 | p-value(χ2) | |
|---|---|---|---|---|---|---|
| Moss abundance | 1 | 2.35E-01 | 1.66E-03 | 9.89E+00 | 1.66E-03 | ** |
| March mean temperature | 1 | 2.09E-02 | 3.68E-02 | 4.36E+00 | 3.68E-02 | * |
| Skeleton volume | 1 | 5.36E-03 | 1.94E-02 | 5.46E+00 | 1.94E-02 | * |
| Light intensity | 1 | 4.81E-04 | 2.00E-16 | 3.02E+02 | 1.33E-67 | *** |
| Chloride (Cl-) | 1 | 2.39E-05 | 2.97E-01 | 1.09E+00 | 2.97E-01 | |
| Light intensity * March mean humidity | 1 | -6.36E-06 | 2.00E-16 | 7.19E+03 | 2.00E-16 | *** |
| March mean humidity | 1 | -4.27E-03 | 1.52E-09 | 2.32E+02 | 2.03E-52 | *** |
| Vegetation cover * Shannon index | 1 | -1.01E-01 | 2.36E-05 | 1.79E+01 | 2.36E-05 | *** |
| Residual | 115 | | | | | |

**Table S4: Ex situ flowering time and environmental predictors of in situ flowering onset in *Arabidopsis thaliana* urban populations.** In situ flowering onset was recorded in April 2023 across 168 populations. Ex situ flowering time (FT) was measured under controlled conditions for 112 transplanted populations.

| **Variable** | **Df** | **χ2** | **p-value (χ2)** | |
|---|---|---|---|---|
| *Ex situ* onset FT | 1 | 6.79E-01 | 4.10E-01 | |
| Light intensity | 1 | 1.46E+00 | 2.27E-01 | |
| Vegetation cover | 1 | 9.78E-01 | 3.23E-01 | |
| Sealed surface | 1 | 1.53E+00 | 2.16E-01 | |
| Moss abundance | 1 | 4.89E-01 | 4.84E-01 | |
| Shannon index | 1 | 2.19E+00 | 1.39E-01 | |
| Habitat type[1] | 3 | 3.94E+00 | 2.68E-01 | |
| Temperature | 1 | 3.46E+00 | 6.27E-02 | . |
| Moisture | 1 | 4.1742 | 0.041044 | * |
| Salinity | 1 | 5.4277 | 0.01982 | * |
| Reaction | 1 | 7.48E-01 | 3.87E-01 | |
| Nutrients | 1 | 1.59E-01 | 6.90E-01 | |
| Light | 1 | 1.64E-01 | 6.86E-01 | |
| Disturbance severity | 1 | 4.49E+00 | 3.41E-02 | * |
| *Ex situ* onset FT : Light | 1 | 1.01E+01 | 1.45E-03 | ** |

[1] Meadow, tree bed, pavement, wall

**TABLE S5: Surface characteristics and vegetation-derived environment variables measured across 393 experimental sites in the city of Cologne.** Surface characteristics were recorded at three key phenological stages of *A. thaliana*: seed sowing (September 2021), the end of the vegetative stage (February 2022), and fruit ripening (May 2022).

| Variable | Definition | Variables | Measured on |
|---|---|---|---|
| Sowing surface | Area of the surface where the F2 seeds were sown (cm²) | 3 | Sep. 2021, Feb. 2022, May 2022 |
| Sealed surface | Proportion of impervious surface in sowing surface (%) | 3 | Sep. 2021, Feb. 2022, May 2022 |
| Soil surface | Proportion of soil in sowing surface (%), indicating the habitable area for *Arabidopsis thaliana* establishment | 3 | Sep. 2021, Feb. 2022, May 2022 |
| Bare soil | Proportion of soil surface not covered by vascular plants or mosses | 3 | Sep. 2021, Feb. 2022, May 2022 |
| Vegetation cover | Proportion of soil surface covered by vascular plants (%) | 3 | Sep. 2021, Feb. 2022, May 2022 |
| Moss abundance | Proportion of soil surface covered by mosses (%) | 3 | Sep. 2021, Feb. 2022, May 2022 |
| Shannon Index | Measure of community diversity integrating species richness and the relative abundance of species. Higher values indicate communities with more species and/or a more even distribution of individuals among species. | 1 | May 2022 |
| Evenness | Measure of diversity (Pielou's evenness index) expressing how evenly abundance is distributed among the species present in a community. Values range from 0 to 1, with 1 indicating that all species are equally abundant. | 1 | May 2022 |
| Disturbance frequency | Community-weighted Herben indicator reflecting frequency of disturbance in habitats typically occupied by constituent species; higher values indicate more frequent disturbance. | 1 | May 2022 |
| Disturbance severity | Community-weighted Herben indicator reflecting intensity or biomass removal during disturbance in habitats typically occupied by constituent species; higher values indicate more severe disturbance. | 1 | May 2022 |
| Nutrients | Community-weighted Ellenberg indicator of soil fertility; higher values indicate nutrient-rich conditions. | 1 | May 2022 |
| Reaction | Community-weighted Ellenberg indicator of soil pH; higher values indicate more alkaline conditions. | 1 | May 2022 |
| Salinity | Community-weighted Ellenberg indicator of soil salt content; higher values indicate higher salinity. | 1 | May 2022 |
| Light | Community-weighted Ellenberg indicator of light availability; higher values indicate higher light availability. | 1 | May 2022 |
| Moisture | Community-weighted Ellenberg indicator of soil moisture; higher values indicate wetter conditions. | 1 | May 2022 |
| Temperature | Community-weighted Ellenberg indicator of temperature conditions; higher values indicate warmer conditions. | 1 | May 2022 |

**TABLE S6: Plant species recorded at each site together with their corresponding Ellenberg indicator values (EIVs) and disturbance indices.** Only vascular plant species, excluding *Arabidopsis thaliana*, were considered. Community-weighted means of EIVs and disturbance indices were calculated from the relative cover of non-indifferent after standardizing their total cover to 100% (see Table S1 ).

| Site_ID | WFO_name | Ellenberg_name | Abundance | Total_abundance | Adjust_Abundance | Light | Temperature | Moisture | Reaction | Nutrients | Salinity | Disturbance Severity | Disturbance Severity herblayer | Disturbance Frequency | Disturbance Frequency herblayer | Mowing Frequency | Grazing Pressure | Soil Disturbance |
|---|---|---|---|---|---|---|---|---|---|---|---|---|---|---|---|---|---|---|
| 1 | Taraxacum officinale F.H.Wigg. | Taraxacum officinale agg. | 0.28 | 0.73 | 0.384 | NA | NA | NA | NA | NA | NA | NA | NA | NA | NA | NA | NA | NA |
| 1 | Poa annua L. | Poa annua | 0.25 | 0.73 | 0.273 | 7.4 | 5.3 | NA | 6 | 7.7 | 0.8 | 0.65201 | 0.63791 | 1.54898 | 2.03286 | 0.77966 | 0.37419 | 0.45721 |
| 1 | Veronica arvensis L. | Veronica arvensis | 0.15 | 0.73 | 0.400 | 6.5 | NA | NA | 5.4 | NA | 0 | 0.59769 | 0.45571 | 0.64594 | 1.51584 | 0.28825 | 0.22901 | 0.32316 |
| 1 | Stellaria media (L.) Vill. | Stellaria media agg. | 0.05 | 0.73 | 0.600 | 7.4 | 5.3 | NA | 6 | 7.7 | 0.8 | 0.65201 | 0.63791 | 1.54898 | 2.03286 | 0.77966 | 0.37419 | 0.45721 |
| 2 | Poa annua L. | Poa annua | 0.15 | 0.55 | 0.161 | 7.4 | 5.3 | NA | 6 | 7.7 | 0.8 | 0.65201 | 0.63791 | 1.54898 | 2.03286 | 0.77966 | 0.37419 | 0.45721 |
| 2 | Hordeum murinum L. | Hordeum murinum agg. | 0.15 | 0.55 | 0.455 | 7.4 | 5.3 | NA | 6 | 7.7 | 0.8 | 0.65201 | 0.63791 | 1.54898 | 2.03286 | 0.77966 | 0.37419 | 0.45721 |
| 2 | Viola odorata L. | Viola odorata | 0.25 | 0.55 | 0.500 | 7.4 | 5.3 | NA | 6 | 7.7 | 0.8 | 0.65201 | 0.63791 | 1.54898 | 2.03286 | 0.77966 | 0.37419 | 0.45721 |
| 3 | Taraxacum officinale F.H.Wigg. | Taraxacum officinale agg. | 0.25 | 0.5 | 0.556 | 7.4 | 5.3 | NA | 6 | 7.7 | 0.8 | 0.65201 | 0.63791 | 1.54898 | 2.03286 | 0.77966 | 0.37419 | 0.45721 |
| 3 | Poa annua L. | Poa annua | 0.23 | 0.5 | 0.333 | 7.4 | 5.3 | NA | 6 | 7.7 | 0.8 | 0.65201 | 0.63791 | 1.54898 | 2.03286 | 0.77966 | 0.37419 | 0.45721 |
| 3 | Sagina procumbens L. | Sagina procumbens | 0.02 | 0.5 | 0.056 | 7.4 | 5.3 | NA | 6 | 7.7 | 0.8 | 0.65201 | 0.63791 | 1.54898 | 2.03286 | 0.77966 | 0.37419 | 0.45721 |
| 4 | Cardamine hirsuta L. | Cardamine hirsuta | 0.1 | 0.25 | 0.400 | 7.4 | 5.3 | NA | 6 | 7.7 | 0.8 | 0.65201 | 0.63791 | 1.54898 | 2.03286 | 0.77966 | 0.37419 | 0.45721 |
| 4 | Poa annua L. | Poa annua | 0.15 | 0.25 | 0.023 | 6.9 | 5.9 | NA | 6.8 | 7 | 0 | 0.84157 | 0.83772 | 1.85474 | 2.04042 | 0.4856 | 0.14178 | 0.80173 |
| 5 | Poa annua L. | Poa annua | 0.3 | 0.5 | 0.235 | 8.1 | NA | 3.5 | 7.5 | 5 | 0.3 | 0.75184 | 0.72654 | 1.3537 | 1.85024 | 0.717 | 0.18024 | 0.48387 |
| 5 | Veronica arvensis L. | Veronica arvensis | 0.02 | 0.5 | 0.150 | 6.3 | NA | 5.9 | 6.3 | 6.9 | 0.1 | 0.6037 | 0.3946 | 0.44368 | 1.50139 | 0.30164 | 0.21315 | 0.21494 |
| 5 | Sagina procumbens L. | Sagina procumbens | 0.02 | 0.5 | 0.300 | 7.1 | 5.3 | 4.5 | 5.9 | 4.8 | 0.3 | 0.52518 | 0.5113 | 1.61045 | 2.05509 | 0.96495 | 0.27897 | 0.32911 |
| 5 | Taraxacum officinale F.H.Wigg. | Taraxacum officinale agg. | 0.16 | 0.5 | 0.200 | NA | NA | NA | NA | NA | NA | NA | NA | NA | NA | NA | NA | NA |
| 6 | Poa annua L. | Poa annua | 0.1 | 0.62 | 0.154 | NA | NA | NA | NA | NA | NA | NA | NA | NA | NA | NA | NA | NA |
| 6 | Taraxacum officinale F.H.Wigg. | Taraxacum officinale agg. | 0.25 | 0.62 | 0.111 | NA | NA | NA | NA | NA | NA | NA | NA | NA | NA | NA | NA | NA |
| 6 | Veronica arvensis L. | Veronica arvensis | 0.02 | 0.62 | 0.214 | 7.2 | 5.7 | 4.3 | 6.8 | NA | 0.1 | 0.54405 | 0.41742 | 0.70503 | 1.63482 | 0.49706 | 0.18636 | 0.24515 |
| 6 | Viola odorata L. | Viola odorata | 0.2 | 0.62 | 0.071 | 6.3 | NA | 5.9 | 6.3 | 6.9 | 0.1 | 0.6037 | 0.3946 | 0.44368 | 1.50139 | 0.30164 | 0.21315 | 0.21494 |
| 6 | Senecio vulgaris L. | Senecio vulgaris | 0.05 | 0.62 | 0.067 | NA | NA | NA | NA | NA | NA | NA | NA | NA | NA | NA | NA | NA |
| 7 | Poa annua L. | Poa annua | 0.25 | 0.55 | 0.200 | NA | NA | NA | NA | NA | NA | NA | NA | NA | NA | NA | NA | NA |
| 7 | Taraxacum officinale F.H.Wigg. | Taraxacum officinale agg. | 0.25 | 0.55 | 0.278 | NA | NA | NA | NA | NA | NA | NA | NA | NA | NA | NA | NA | NA |
| 7 | Veronica arvensis L. | Veronica arvensis | 0.05 | 0.55 | 0.250 | NA | NA | NA | NA | NA | NA | NA | NA | NA | NA | NA | NA | NA |
| 8 | Poa annua L. | Poa annua | 0.25 | 0.5 | 0.688 | 7.4 | 5.3 | NA | 6 | 7.7 | 0.8 | 0.65201 | 0.63791 | 1.54898 | 2.03286 | 0.77966 | 0.37419 | 0.45721 |
| 8 | Taraxacum officinale F.H.Wigg. | Taraxacum officinale agg. | 0.15 | 0.5 | 0.111 | 6.3 | NA | 5.9 | 6.3 | 6.9 | 0.1 | 0.6037 | 0.3946 | 0.44368 | 1.50139 | 0.30164 | 0.21315 | 0.21494 |
| 8 | Veronica arvensis L. | Veronica arvensis | 0.1 | 0.5 | 0.263 | 6.3 | NA | 5.9 | 6.3 | 6.9 | 0.1 | 0.6037 | 0.3946 | 0.44368 | 1.50139 | 0.30164 | 0.21315 | 0.21494 |
| 9 | Poa annua L. | Poa annua | 0.25 | 0.45 | 0.063 | 6.8 | 6.2 | 4.4 | NA | NA | 0.3 | 0.7871 | 0.75023 | 1.37944 | 1.92919 | 0.40631 | 0.19607 | 0.63681 |
| 9 | Taraxacum officinale F.H.Wigg. | Taraxacum officinale agg. | 0.2 | 0.45 | 0.056 | 7.3 | 6.1 | 3.8 | 6 | 5.1 | 0 | 0.62987 | 0.59281 | 1.27995 | 1.90491 | 0.6275 | 0.3036 | 0.42608 |
| 10 | Taraxacum officinale F.H.Wigg. | Taraxacum officinale agg. | 0.2 | 0.3 | 0.083 | 7.4 | 5.8 | 4.2 | 6.7 | 5.3 | 0 | 0.62702 | 0.59749 | 1.35516 | 1.96644 | 0.57704 | 0.23154 | 0.49642 |
| 10 | Sagina procumbens L. | Sagina procumbens | 0.05 | 0.3 | 0.789 | 7.4 | 5.3 | NA | 6 | 7.7 | 0.8 | 0.65201 | 0.63791 | 1.54898 | 2.03286 | 0.77966 | 0.37419 | 0.45721 |
| 10 | Cardamine hirsuta L. | Cardamine hirsuta | 0.05 | 0.3 | 0.765 | 7.4 | 5.3 | NA | 6 | 7.7 | 0.8 | 0.65201 | 0.63791 | 1.54898 | 2.03286 | 0.77966 | 0.37419 | 0.45721 |
| 12 | Poa annua L. | Poa annua | 0.05 | 0.15 | 0.056 | 6.2 | 5.5 | NA | 6.9 | 8.5 | 0.1 | 0.60658 | 0.39025 | 0.44214 | 1.39281 | 0.24736 | 0.18526 | 0.18179 |
| 12 | Taraxacum officinale F.H.Wigg. | Taraxacum officinale agg. | 0.1 | 0.15 | 0.786 | 7.4 | 5.3 | NA | 6 | 7.7 | 0.8 | 0.65201 | 0.63791 | 1.54898 | 2.03286 | 0.77966 | 0.37419 | 0.45721 |
| 13 | Draba verna L. | Erophila verna | 0.05 | 0.25 | 0.200 | NA | NA | NA | NA | NA | NA | NA | NA | NA | NA | NA | NA | NA |
| 13 | Cardamine hirsuta L. | Cardamine hirsuta | 0.05 | 0.25 | 0.550 | 7.4 | 5.3 | NA | 6 | 7.7 | 0.8 | 0.65201 | 0.63791 | 1.54898 | 2.03286 | 0.77966 | 0.37419 | 0.45721 |
| 13 | Veronica arvensis L. | Veronica arvensis | 0.05 | 0.25 | 0.200 | 8.1 | NA | 3.9 | 6.9 | 6.4 | 0.4 | 0.80098 | 0.76713 | 1.29016 | 1.92909 | 0.75604 | 0.19458 | 0.52486 |
| 13 | Poa annua L. | Poa annua | 0.1 | 0.25 | 0.150 | 8.1 | NA | NA | 5.2 | NA | 0.1 | 0.51276 | 0.49581 | 1.51667 | 2.20989 | 0.98959 | 0.46997 | 0.3594 |
| 14 | Taraxacum officinale F.H.Wigg. | Taraxacum officinale agg. | 0.15 | 0.25 | 0.050 | 8.1 | NA | NA | 5.2 | NA | 0.1 | 0.51276 | 0.49581 | 1.51667 | 2.20989 | 0.98959 | 0.46997 | 0.3594 |
| 14 | Veronica arvensis L. | Veronica arvensis | 0.05 | 0.25 | 0.100 | 8.1 | NA | NA | 5.2 | NA | 0.1 | 0.51276 | 0.49581 | 1.51667 | 2.20989 | 0.98959 | 0.46997 | 0.3594 |
| 14 | Cardamine hirsuta L. | Cardamine hirsuta | 0.05 | 0.25 | 0.083 | 8.1 | NA | NA | 5.2 | NA | 0.1 | 0.51276 | 0.49581 | 1.51667 | 2.20989 | 0.98959 | 0.46997 | 0.3594 |
| 15 | Taraxacum officinale F.H.Wigg. | Taraxacum officinale agg. | 0.25 | 0.5 | 0.167 | 8.1 | NA | NA | 5.2 | NA | 0.1 | 0.51276 | 0.49581 | 1.51667 | 2.20989 | 0.98959 | 0.46997 | 0.3594 |
| 15 | Poa annua L. | Poa annua | 0.2 | 0.5 | 0.108 | 8.1 | NA | NA | 5.2 | NA | 0.1 | 0.51276 | 0.49581 | 1.51667 | 2.20989 | 0.98959 | 0.46997 | 0.3594 |
| 15 | Veronica arvensis L. | Veronica arvensis | 0.05 | 0.5 | 0.094 | 8.1 | NA | NA | 5.2 | NA | 0.1 | 0.51276 | 0.49581 | 1.51667 | 2.20989 | 0.98959 | 0.46997 | 0.3594 |
| 16 | Taraxacum officinale F.H.Wigg. | Taraxacum officinale agg. | 0.15 | 0.2 | 0.315 | 8.1 | NA | NA | 5.2 | NA | 0.1 | 0.51276 | 0.49581 | 1.51667 | 2.20989 | 0.98959 | 0.46997 | 0.3594 |
| 16 | Poa annua L. | Poa annua | 0.04 | 0.2 | 0.172 | 8.1 | NA | NA | 5.2 | NA | 0.1 | 0.51276 | 0.49581 | 1.51667 | 2.20989 | 0.98959 | 0.46997 | 0.3594 |
| 16 | Veronica arvensis L. | Veronica arvensis | 0.01 | 0.2 | 0.109 | 8.1 | NA | NA | 5.2 | NA | 0.1 | 0.51276 | 0.49581 | 1.51667 | 2.20989 | 0.98959 | 0.46997 | 0.3594 |
| 17 | Taraxacum officinale F.H.Wigg. | Taraxacum officinale agg. | 0.1 | 0.25 | 0.149 | 8.1 | NA | NA | 5.2 | NA | 0.1 | 0.51276 | 0.49581 | 1.51667 | 2.20989 | 0.98959 | 0.46997 | 0.3594 |
| 17 | Poa annua L. | Poa annua | 0.1 | 0.25 | 0.245 | 8.1 | NA | NA | 5.2 | NA | 0.1 | 0.51276 | 0.49581 | 1.51667 | 2.20989 | 0.98959 | 0.46997 | 0.3594 |
| 17 | Sagina procumbens L. | Sagina procumbens | 0.05 | 0.25 | 0.400 | 8.1 | NA | NA | 5.2 | NA | 0.1 | 0.51276 | 0.49581 | 1.51667 | 2.20989 | 0.98959 | 0.46997 | 0.3594 |
| 18 | Sagina procumbens L. | Sagina procumbens | 0.01 | 0.02 | 0.150 | 6.3 | NA | 5.9 | 6.3 | 6.9 | 0.1 | 0.6037 | 0.3946 | 0.44368 | 1.50139 | 0.30164 | 0.21315 | 0.21494 |
| 18 | Veronica arvensis L. | Veronica arvensis | 0.01 | 0.02 | 0.556 | 6 | NA | 6.4 | 7.5 | 6.1 | 0.1 | 0.67416 | 0.35728 | 0.31529 | 1.53539 | 0.05981 | 0.19852 | 0.1369 |
| 19 | Poa annua L. | Poa annua | 0.1 | 0.25 | 0.500 | 6 | NA | 6.4 | 7.5 | 6.1 | 0.1 | 0.67416 | 0.35728 | 0.31529 | 1.53539 | 0.05981 | 0.19852 | 0.1369 |
| 19 | Veronica arvensis L. | Veronica arvensis | 0.05 | 0.25 | 0.800 | 6 | NA | 6.4 | 7.5 | 6.1 | 0.1 | 0.67416 | 0.35728 | 0.31529 | 1.53539 | 0.05981 | 0.19852 | 0.1369 |
| 19 | Cardamine hirsuta L. | Cardamine hirsuta | 0.05 | 0.25 | 0.706 | 6 | NA | 6.4 | 7.5 | 6.1 | 0.1 | 0.67416 | 0.35728 | 0.31529 | 1.53539 | 0.05981 | 0.19852 | 0.1369 |
| 19 | Sagina procumbens L. | Sagina procumbens | 0.05 | 0.25 | 1.000 | 6 | NA | 6.4 | 7.5 | 6.1 | 0.1 | 0.67416 | 0.35728 | 0.31529 | 1.53539 | 0.05981 | 0.19852 | 0.1369 |
| 20 | Poa annua L. | Poa annua | 0.1 | 0.82 | 0.949 | 6 | NA | 6.4 | 7.5 | 6.1 | 0.1 | 0.67416 | 0.35728 | 0.31529 | 1.53539 | 0.05981 | 0.19852 | 0.1369 |
| 20 | Veronica arvensis L. | Veronica arvensis | 0.22 | 0.82 | 0.770 | 6 | NA | 6.4 | 7.5 | 6.1 | 0.1 | 0.67416 | 0.35728 | 0.31529 | 1.53539 | 0.05981 | 0.19852 | 0.1369 |
| 20 | Taraxacum officinale F.H.Wigg. | Taraxacum officinale agg. | 0.5 | 0.82 | 0.700 | 6 | NA | 6.4 | 7.5 | 6.1 | 0.1 | 0.67416 | 0.35728 | 0.31529 | 1.53539 | 0.05981 | 0.19852 | 0.1369 |
| 21 | Poa annua L. | Poa annua | 0.25 | 0.85 | 0.368 | 6 | NA | 6.4 | 7.5 | 6.1 | 0.1 | 0.67416 | 0.35728 | 0.31529 | 1.53539 | 0.05981 | 0.19852 | 0.1369 |
| 21 | Veronica arvensis L. | Veronica arvensis | 0.05 | 0.85 | 0.050 | 7.3 | 5.4 | 4.1 | 7.6 | 4.2 | 0.4 | 0.58728 | 0.56055 | 1.34601 | 1.94534 | 0.82357 | 0.23743 | 0.34603 |
| 21 | Taraxacum officinale F.H.Wigg. | Taraxacum officinale agg. | 0.55 | 0.85 | 0.050 | 7.3 | 5.4 | 4.1 | 7.6 | 4.2 | 0.4 | 0.58728 | 0.56055 | 1.34601 | 1.94534 | 0.82357 | 0.23743 | 0.34603 |
| 22 | Poa annua L. | Poa annua | 0.05 | 0.6 | 0.020 | 7.3 | 5.4 | 4.1 | 7.6 | 4.2 | 0.4 | 0.58728 | 0.56055 | 1.34601 | 1.94534 | 0.82357 | 0.23743 | 0.34603 |
| 22 | Veronica arvensis L. | Veronica arvensis | 0.05 | 0.6 | 0.080 | 7.3 | 5.4 | 4.1 | 7.6 | 4.2 | 0.4 | 0.58728 | 0.56055 | 1.34601 | 1.94534 | 0.82357 | 0.23743 | 0.34603 |
| 22 | Taraxacum officinale F.H.Wigg. | Taraxacum officinale agg. | 0.3 | 0.6 | 0.100 | 7.3 | 5.4 | 4.1 | 7.6 | 4.2 | 0.4 | 0.58728 | 0.56055 | 1.34601 | 1.94534 | 0.82357 | 0.23743 | 0.34603 |
| 22 | Cardamine hirsuta L. | Cardamine hirsuta | 0.05 | 0.6 | 0.084 | 7.3 | 5.4 | 4.1 | 7.6 | 4.2 | 0.4 | 0.58728 | 0.56055 | 1.34601 | 1.94534 | 0.82357 | 0.23743 | 0.34603 |
| 22 | Viola odorata L. | Viola odorata | 0.15 | 0.6 | 0.100 | 8.1 | NA | 3.5 | 6.1 | NA | 0.2 | 0.68175 | 0.6716 | 1.70824 | 1.99767 | 0.41848 | 0.25411 | 0.57318 |
| 23 | Poa annua L. | Poa annua | 0.05 | 0.9 | 0.100 | NA | NA | NA | NA | NA | NA | NA | NA | NA | NA | NA | NA | NA |
| 23 | Veronica arvensis L. | Veronica arvensis | 0.5 | 0.9 | 0.050 | 8.1 | NA | 3.5 | 6.1 | NA | 0.2 | 0.68175 | 0.6716 | 1.70824 | 1.99767 | 0.41848 | 0.25411 | 0.57318 |
| 23 | Taraxacum officinale F.H.Wigg. | Taraxacum officinale agg. | 0.25 | 0.9 | 0.080 | 7.3 | 5.4 | 4.1 | 7.6 | 4.2 | 0.4 | 0.58728 | 0.56055 | 1.34601 | 1.94534 | 0.82357 | 0.23743 | 0.34603 |
| 23 | Parietaria judaica L. | Parietaria judaica (diffusa) | 0.1 | 0.9 | 0.100 | 7.3 | 5.4 | 4.1 | 7.6 | 4.2 | 0.4 | 0.58728 | 0.56055 | 1.34601 | 1.94534 | 0.82357 | 0.23743 | 0.34603 |
| 24 | Poa annua L. | Poa annua | 0.1 | 0.25 | 0.080 | 7.3 | 5.4 | 4.1 | 7.6 | 4.2 | 0.4 | 0.58728 | 0.56055 | 1.34601 | 1.94534 | 0.82357 | 0.23743 | 0.34603 |
| 24 | Veronica arvensis L. | Veronica arvensis | 0.05 | 0.25 | 0.100 | 7.3 | 5.4 | 4.1 | 7.6 | 4.2 | 0.4 | 0.58728 | 0.56055 | 1.34601 | 1.94534 | 0.82357 | 0.23743 | 0.34603 |
| 24 | Taraxacum officinale F.H.Wigg. | Taraxacum officinale agg. | 0.1 | 0.25 | 0.080 | 7.3 | 5.4 | 4.1 | 7.6 | 4.2 | 0.4 | 0.58728 | 0.56055 | 1.34601 | 1.94534 | 0.82357 | 0.23743 | 0.34603 |
| 25 | Cardamine hirsuta L. | Cardamine hirsuta | 0.05 | 0.05 | 0.100 | 8.1 | NA | 3.5 | 6.1 | NA | 0.2 | 0.68175 | 0.6716 | 1.70824 | 1.99767 | 0.41848 | 0.25411 | 0.57318 |
| 27 | Taraxacum officinale F.H.Wigg. | Taraxacum officinale agg. | 0.08 | 0.08 | 0.105 | 8.1 | NA | 3.5 | 6.1 | NA | 0.2 | 0.68175 | 0.6716 | 1.70824 | 1.99767 | 0.41848 | 0.25411 | 0.57318 |
| 29 | Taraxacum officinale F.H.Wigg. | Taraxacum officinale agg. | 0.05 | 0.15 | 0.100 | 8.1 | NA | 3.5 | 6.1 | NA | 0.2 | 0.68175 | 0.6716 | 1.70824 | 1.99767 | 0.41848 | 0.25411 | 0.57318 |
| 29 | Sagina procumbens L. | Sagina procumbens | 0.1 | 0.15 | 0.080 | 8.1 | NA | 3.5 | 6.1 | NA | 0.2 | 0.68175 | 0.6716 | 1.70824 | 1.99767 | 0.41848 | 0.25411 | 0.57318 |
| 30 | Veronica persica Poir. | Veronica persica (tournefortii) | 0.02 | 0.88 | 0.160 | 7.3 | 5.4 | 4.1 | 7.6 | 4.2 | 0.4 | 0.58728 | 0.56055 | 1.34601 | 1.94534 | 0.82357 | 0.23743 | 0.34603 |
| 30 | Plantago lanceolata L. | Plantago lanceolata | 0.1 | 0.88 | 0.222 | 7.3 | 5.4 | 4.1 | 7.6 | 4.2 | 0.4 | 0.58728 | 0.56055 | 1.34601 | 1.94534 | 0.82357 | 0.23743 | 0.34603 |
| 30 | Valeriana locusta L. | Valerianella locusta (olitoria) | 0.42 | 0.88 | 0.308 | 7.3 | 5.4 | 4.1 | 7.6 | 4.2 | 0.4 | 0.58728 | 0.56055 | 1.34601 | 1.94534 | 0.82357 | 0.23743 | 0.34603 |
| 30 | Cotoneaster dammeri C.K.Schneid. | NA | 0.05 | 0.88 | 0.133 | 7.3 | 5.4 | 4.1 | 7.6 | 4.2 | 0.4 | 0.58728 | 0.56055 | 1.34601 | 1.94534 | 0.82357 | 0.23743 | 0.34603 |
| 30 | Lepidium draba L. | Cardaria draba | 0.05 | 0.88 | 0.286 | 7.3 | 5.4 | 4.1 | 7.6 | 4.2 | 0.4 | 0.58728 | 0.56055 | 1.34601 | 1.94534 | 0.82357 | 0.23743 | 0.34603 |
| 30 | Cerastium glomeratum Thuill. | Cerastium glomeratum | 0.02 | 0.88 | 0.178 | 7.3 | 5.4 | 4.1 | 7.6 | 4.2 | 0.4 | 0.58728 | 0.56055 | 1.34601 | 1.94534 | 0.82357 | 0.23743 | 0.34603 |
| 30 | Taraxacum officinale F.H.Wigg. | Taraxacum officinale agg. | 0.05 | 0.88 | 0.100 | 7.3 | 5.4 | 4.1 | 7.6 | 4.2 | 0.4 | 0.58728 | 0.56055 | 1.34601 | 1.94534 | 0.82357 | 0.23743 | 0.34603 |
| 30 | Arenaria serpyllifolia L. | Arenaria serpyllifolia agg. | 0.02 | 0.88 | 0.083 | 7.3 | 5.4 | 4.1 | 7.6 | 4.2 | 0.4 | 0.58728 | 0.56055 | 1.34601 | 1.94534 | 0.82357 | 0.23743 | 0.34603 |
| 30 | Daucus carota L. | Daucus carota | 0.05 | 0.88 | 0.172 | 7.3 | 5.4 | 4.1 | 7.6 | 4.2 | 0.4 | 0.58728 | 0.56055 | 1.34601 | 1.94534 | 0.82357 | 0.23743 | 0.34603 |
| 30 | Bromus hordeaceus L. | Bromus hordeaceus (mollis) | 0.1 | 0.88 | 0.190 | 7.3 | 5.4 | 4.1 | 7.6 | 4.2 | 0.4 | 0.58728 | 0.56055 | 1.34601 | 1.94534 | 0.82357 | 0.23743 | 0.34603 |
| 31 | Lepidium draba L. | Cardaria draba | 0.2 | 0.85 | 0.100 | 7.3 | 5.4 | 4.1 | 7.6 | 4.2 | 0.4 | 0.58728 | 0.56055 | 1.34601 | 1.94534 | 0.82357 | 0.23743 | 0.34603 |
| 31 | Plantago lanceolata L. | Plantago lanceolata | 0.05 | 0.85 | 0.150 | NA | NA | NA | NA | NA | NA | NA | NA | NA | NA | NA | NA | NA |
| 31 | Taraxacum officinale F.H.Wigg. | Taraxacum officinale agg. | 0.1 | 0.85 | 0.150 | 7.9 | NA | 4.1 | 5.8 | NA | 0.9 | 0.54139 | 0.51128 | 1.31363 | 1.93633 | 0.8112 | 0.25085 | 0.32953 |
| 31 | Holcus lanatus L. | Holcus lanatus | 0.2 | 0.85 | 0.227 | 7.9 | NA | 4.1 | 5.8 | NA | 0.9 | 0.54139 | 0.51128 | 1.31363 | 1.93633 | 0.8112 | 0.25085 | 0.32953 |
| 31 | Lolium perenne L. | Lolium perenne | 0.3 | 0.85 | 0.263 | 7.9 | NA | 4.1 | 5.8 | NA | 0.9 | 0.54139 | 0.51128 | 1.31363 | 1.93633 | 0.8112 | 0.25085 | 0.32953 |
| 32 | Glechoma hederacea L. | Glechoma hederacea | 0.15 | 1 | 0.250 | 7.9 | NA | 4.1 | 5.8 | NA | 0.9 | 0.54139 | 0.51128 | 1.31363 | 1.93633 | 0.8112 | 0.25085 | 0.32953 |
| 32 | Urtica dioica L. | Urtica dioica | 0.1 | 1 | 0.200 | 7.9 | NA | 4.1 | 5.8 | NA | 0.9 | 0.54139 | 0.51128 | 1.31363 | 1.93633 | 0.8112 | 0.25085 | 0.32953 |
| 32 | Holcus lanatus L. | Holcus lanatus | 0.15 | 1 | 0.400 | 7.9 | NA | 4.1 | 5.8 | NA | 0.9 | 0.54139 | 0.51128 | 1.31363 | 1.93633 | 0.8112 | 0.25085 | 0.32953 |
| 32 | Lolium perenne L. | Lolium perenne | 0.35 | 1 | 0.255 | 7.9 | NA | 4.1 | 5.8 | NA | 0.9 | 0.54139 | 0.51128 | 1.31363 | 1.93633 | 0.8112 | 0.25085 | 0.32953 |
| 32 | Dipsacus fullonum L. | Dipsacus fullonum (sylvestris) | 0.25 | 1 | 0.250 | 7.9 | NA | 4.1 | 5.8 | NA | 0.9 | 0.54139 | 0.51128 | 1.31363 | 1.93633 | 0.8112 | 0.25085 | 0.32953 |
| 33 | Plantago lanceolata L. | Plantago lanceolata | 0.3 | 1 | 0.051 | 7.9 | NA | 4.1 | 5.8 | NA | 0.9 | 0.54139 | 0.51128 | 1.31363 | 1.93633 | 0.8112 | 0.25085 | 0.32953 |
| 33 | Valeriana locusta L. | Valerianella locusta (olitoria) | 0.1 | 1 | 0.100 | 7.9 | NA | 4.1 | 5.8 | NA | 0.9 | 0.54139 | 0.51128 | 1.31363 | 1.93633 | 0.8112 | 0.25085 | 0.32953 |
| 33 | Medicago lupulina L. | Medicago lupulina | 0.1 | 1 | 0.250 | 7.9 | NA | 4.1 | 5.8 | NA | 0.9 | 0.54139 | 0.51128 | 1.31363 | 1.93633 | 0.8112 | 0.25085 | 0.32953 |

| | | | | | | | | | | | | | | | | | | |
|---|---|---|---|---|---|---|---|---|---|---|---|---|---|---|---|---|---|---|
| 33 | Vicia sativa L. | Vicia sativa agg. | 0.05 | 1 | 0.110 | 7.9 | NA | 4.1 | 5.8 | NA | 0.9 | 0.54139 | 0.51128 | 1.31363 | 1.93633 | 0.8112 | 0.25085 | 0.32953 |
| 33 | Veronica persica Poir. | Veronica persica (tournefortii) | 0.05 | 1 | 0.100 | 7.9 | NA | 4.1 | 5.8 | NA | 0.9 | 0.54139 | 0.51128 | 1.31363 | 1.93633 | 0.8112 | 0.25085 | 0.32953 |
| 33 | Linaria vulgaris Mill. | Linaria vulgaris | 0.05 | 1 | 0.050 | 7.9 | NA | 4.1 | 5.8 | NA | 0.9 | 0.54139 | 0.51128 | 1.31363 | 1.93633 | 0.8112 | 0.25085 | 0.32953 |
| 33 | Lolium perenne L. | Lolium perenne | 0.3 | 1 | 0.256 | 7.9 | NA | 4.1 | 5.8 | NA | 0.9 | 0.54139 | 0.51128 | 1.31363 | 1.93633 | 0.8112 | 0.25085 | 0.32953 |
| 33 | Ranunculus repens L. | Ranunculus repens | 0.05 | 1 | 0.200 | 7.9 | NA | 4.1 | 5.8 | NA | 0.9 | 0.54139 | 0.51128 | 1.31363 | 1.93633 | 0.8112 | 0.25085 | 0.32953 |
| 34 | Taraxacum officinale F.H.Wigg. | Taraxacum officinale agg. | 0.05 | 0.25 | 0.050 | 7.9 | NA | 4.1 | 5.8 | NA | 0.9 | 0.54139 | 0.51128 | 1.31363 | 1.93633 | 0.8112 | 0.25085 | 0.32953 |
| 34 | Galium mollugo L. | Galium mollugo mollugo (elatum) | 0.1 | 0.25 | 0.130 | NA | NA | NA | NA | NA | NA | NA | NA | NA | NA | NA | NA | NA |
| 34 | Rubus pseudojaponicus Koidz. | Rubus caesius | 0.05 | 0.25 | 0.230 | NA | NA | NA | NA | NA | NA | NA | NA | NA | NA | NA | NA | NA |
| 34 | Urtica dioica L. | Urtica dioica | 0.05 | 0.25 | 0.172 | NA | NA | NA | NA | NA | NA | NA | NA | NA | NA | NA | NA | NA |
| 35 | Taraxacum officinale F.H.Wigg. | Taraxacum officinale agg. | 0.1 | 0.65 | 0.105 | NA | NA | NA | NA | NA | NA | NA | NA | NA | NA | NA | NA | NA |
| 35 | Glechoma hederacea L. | Glechoma hederacea | 0.05 | 0.65 | 0.100 | NA | NA | NA | NA | NA | NA | NA | NA | NA | NA | NA | NA | NA |
| 35 | Sanguisorba minor Scop. | anguisorba minor (Poterium sanguisorb | 0.05 | 0.65 | 0.100 | NA | NA | NA | NA | NA | NA | NA | NA | NA | NA | NA | NA | NA |
| 35 | Galium verum L. | Galium verum agg. (Cruciata glabra) | 0.1 | 0.65 | 0.050 | NA | NA | NA | NA | NA | NA | NA | NA | NA | NA | NA | NA | NA |
| 35 | Poa annua L. | Poa annua | 0.35 | 0.65 | 0.080 | NA | NA | NA | NA | NA | NA | NA | NA | NA | NA | NA | NA | NA |
| 36 | Taraxacum officinale F.H.Wigg. | Taraxacum officinale agg. | 0.1 | 0.9 | 0.080 | NA | NA | NA | NA | NA | NA | NA | NA | NA | NA | NA | NA | NA |
| 36 | Poa annua L. | Poa annua | 0.4 | 0.9 | 0.056 | 8.1 | NA | NA | 5.2 | NA | 0.1 | 0.51276 | 0.49581 | 1.51667 | 2.20989 | 0.98959 | 0.46997 | 0.3594 |
| 36 | Galium verum L. | Galium verum agg. (Cruciata glabra) | 0.15 | 0.9 | 0.077 | 8.1 | NA | NA | 5.2 | NA | 0.1 | 0.51276 | 0.49581 | 1.51667 | 2.20989 | 0.98959 | 0.46997 | 0.3594 |
| 36 | Cirsium vulgare (Savi) Ten. | Cirsium vulgare (lanceolatum) | 0.1 | 0.9 | 0.064 | 8.1 | NA | NA | 5.2 | NA | 0.1 | 0.51276 | 0.49581 | 1.51667 | 2.20989 | 0.98959 | 0.46997 | 0.3594 |
| 36 | Geum urbanum L. | Geum urbanum | 0.1 | 0.9 | 0.050 | NA | NA | NA | 7 | NA | 0 | 0.66957 | 0.20824 | 0.12967 | 1.34554 | 0.07292 | 0.20428 | 0.13248 |
| 36 | Valeriana locusta L. | Valerianella locusta (olitoria) | 0.05 | 0.9 | 0.026 | NA | NA | NA | NA | NA | NA | NA | NA | NA | NA | NA | NA | NA |
| 37 | Galium mollugo L. | Galium mollugo mollugo (elatum) | 0.15 | 0.7 | 0.235 | 7.1 | 5.3 | 4.5 | 5.9 | 4.8 | 0.3 | 0.52518 | 0.5113 | 1.61045 | 2.05509 | 0.96495 | 0.27897 | 0.32911 |
| 37 | Poa annua L. | Poa annua | 0.4 | 0.7 | 0.300 | 7.1 | 5.3 | 4.5 | 5.9 | 4.8 | 0.3 | 0.52518 | 0.5113 | 1.61045 | 2.05509 | 0.96495 | 0.27897 | 0.32911 |
| 37 | Dipsacus fullonum L. | Dipsacus fullonum (sylvestris) | 0.15 | 0.7 | 0.100 | 7.1 | 5.3 | 4.5 | 5.9 | 4.8 | 0.3 | 0.52518 | 0.5113 | 1.61045 | 2.05509 | 0.96495 | 0.27897 | 0.32911 |
| 38 | Glechoma hederacea L. | Glechoma hederacea | 0.05 | 0.7 | 0.200 | 7.1 | 5.3 | 4.5 | 5.9 | 4.8 | 0.3 | 0.52518 | 0.5113 | 1.61045 | 2.05509 | 0.96495 | 0.27897 | 0.32911 |
| 38 | Valeriana locusta L. | Valerianella locusta (olitoria) | 0.05 | 0.7 | 0.100 | 7.3 | 5.4 | 4.1 | 7.6 | 4.2 | 0.4 | 0.58728 | 0.56055 | 1.34601 | 1.94534 | 0.82357 | 0.23743 | 0.34603 |
| 38 | Poa annua L. | Poa annua | 0.55 | 0.7 | 0.100 | 7.3 | 5.4 | 4.1 | 7.6 | 4.2 | 0.4 | 0.58728 | 0.56055 | 1.34601 | 1.94534 | 0.82357 | 0.23743 | 0.34603 |
| 38 | Geranium dissectum L. | Geranium dissectum | 0.05 | 0.7 | 0.057 | 7.3 | 5.4 | 4.1 | 7.6 | 4.2 | 0.4 | 0.58728 | 0.56055 | 1.34601 | 1.94534 | 0.82357 | 0.23743 | 0.34603 |
| 39 | Taraxacum officinale F.H.Wigg. | Taraxacum officinale agg. | 0.05 | 0.75 | 0.010 | 8.5 | NA | 2.5 | NA | NA | 0 | 0.506 | 0.44835 | 1.02121 | 1.67002 | 0.34316 | 0.25824 | 0.34742 |
| 39 | Poa annua L. | Poa annua | 0.35 | 0.75 | 0.042 | 7.3 | 5.4 | 4.1 | 7.6 | 4.2 | 0.4 | 0.58728 | 0.56055 | 1.34601 | 1.94534 | 0.82357 | 0.23743 | 0.34603 |
| 39 | Dipsacus fullonum L. | Dipsacus fullonum (sylvestris) | 0.15 | 0.75 | 0.080 | 7.3 | 5.4 | 4.1 | 7.6 | 4.2 | 0.4 | 0.58728 | 0.56055 | 1.34601 | 1.94534 | 0.82357 | 0.23743 | 0.34603 |
| 39 | Jacobaea analoga (DC.) Veldkamp | Senecio jacobaea | 0.15 | 0.75 | 0.050 | 7.3 | NA | 3.9 | 6.1 | 3.9 | 0.1 | 0.55903 | 0.41538 | 0.65189 | 1.65181 | 0.43719 | 0.24279 | 0.21709 |
| 39 | Sisymbrium officinale (L.) Scop. | Sisymbrium officinale | 0.05 | 0.75 | 0.080 | 7.3 | 5.4 | 4.1 | 7.6 | 4.2 | 0.4 | 0.58728 | 0.56055 | 1.34601 | 1.94534 | 0.82357 | 0.23743 | 0.34603 |
| 40 | Taraxacum officinale F.H.Wigg. | Taraxacum officinale agg. | 0.2 | 1 | 0.040 | 7.3 | 5.4 | 4.1 | 7.6 | 4.2 | 0.4 | 0.58728 | 0.56055 | 1.34601 | 1.94534 | 0.82357 | 0.23743 | 0.34603 |
| 40 | Poa annua L. | Poa annua | 0.55 | 1 | 0.025 | 7.3 | 5.4 | 4.1 | 7.6 | 4.2 | 0.4 | 0.58728 | 0.56055 | 1.34601 | 1.94534 | 0.82357 | 0.23743 | 0.34603 |
| 40 | Dipsacus fullonum L. | Dipsacus fullonum (sylvestris) | 0.2 | 1 | 0.200 | 8.6 | NA | 2.7 | 4.4 | 1.6 | 0 | 0.53557 | 0.50485 | 1.29532 | 1.90211 | 0.72298 | 0.28495 | 0.2778 |
| 40 | Galium mollugo L. | Galium mollugo mollugo (elatum) | 0.05 | 1 | 0.125 | 8.6 | NA | 2.7 | 4.4 | 1.6 | 0 | 0.53557 | 0.50485 | 1.29532 | 1.90211 | 0.72298 | 0.28495 | 0.2778 |
| 41 | Taraxacum officinale F.H.Wigg. | Taraxacum officinale agg. | 0.25 | 0.9 | 0.100 | 7.3 | 5.4 | 4.1 | 7.6 | 4.2 | 0.4 | 0.58728 | 0.56055 | 1.34601 | 1.94534 | 0.82357 | 0.23743 | 0.34603 |
| 41 | Poa annua L. | Poa annua | 0.35 | 0.9 | 0.100 | 8.6 | NA | 2.7 | 4.4 | 1.6 | 0 | 0.53557 | 0.50485 | 1.29532 | 1.90211 | 0.72298 | 0.28495 | 0.2778 |
| 41 | Glechoma hederacea L. | Glechoma hederacea | 0.05 | 0.9 | 0.066 | 7.3 | 5.8 | 4.8 | 5.3 | 5.6 | 0 | 0.67083 | 0.63295 | 1.24583 | 1.99939 | 0.56795 | 0.28417 | 0.48569 |
| 41 | Veronica serpyllifolia L. | Veronica serpyllifolia | 0.05 | 0.9 | 0.050 | 8.1 | NA | NA | 5.2 | NA | 0.1 | 0.51276 | 0.49581 | 1.51667 | 2.20989 | 0.98959 | 0.46997 | 0.3594 |
| 41 | Myosotis ramosissima Rochel | Myosotis ramosissima (coll. Hispida) | 0.05 | 0.9 | 0.105 | 7.1 | 5.3 | 4.5 | 5.9 | 4.8 | 0.3 | 0.52518 | 0.5113 | 1.61045 | 2.05509 | 0.96495 | 0.27897 | 0.32911 |
| 41 | Prunella vulgaris L. | Prunella vulgaris | 0.15 | 0.9 | 0.050 | 7.3 | NA | 3.9 | 6.1 | 3.9 | 0.1 | 0.55903 | 0.41538 | 0.65189 | 1.65181 | 0.43719 | 0.24279 | 0.21709 |
| 42 | Taraxacum officinale F.H.Wigg. | Taraxacum officinale agg. | 0.25 | 1 | 0.250 | 7.1 | 5.3 | 4.5 | 5.9 | 4.8 | 0.3 | 0.52518 | 0.5113 | 1.61045 | 2.05509 | 0.96495 | 0.27897 | 0.32911 |
| 42 | Poa annua L. | Poa annua | 0.33 | 1 | 0.187 | 7.1 | 5.3 | 4.5 | 5.9 | 4.8 | 0.3 | 0.52518 | 0.5113 | 1.61045 | 2.05509 | 0.96495 | 0.27897 | 0.32911 |
| 42 | Myosotis arvensis Hill | Myosotis arvensis (intermedia) | 0.05 | 1 | 0.156 | 7.1 | 5.3 | 4.5 | 5.9 | 4.8 | 0.3 | 0.52518 | 0.5113 | 1.61045 | 2.05509 | 0.96495 | 0.27897 | 0.32911 |
| 42 | Sagina procumbens L. | Sagina procumbens | 0.02 | 1 | 0.242 | 7.1 | 5.3 | 4.5 | 5.9 | 4.8 | 0.3 | 0.52518 | 0.5113 | 1.61045 | 2.05509 | 0.96495 | 0.27897 | 0.32911 |
| 42 | Jacobaea vulgaris Gaertn. | Senecio jacobaea | 0.05 | 1 | 0.170 | 7.1 | 5.3 | 4.5 | 5.9 | 4.8 | 0.3 | 0.52518 | 0.5113 | 1.61045 | 2.05509 | 0.96495 | 0.27897 | 0.32911 |
| 42 | Galium mollugo L. | Galium mollugo mollugo (elatum) | 0.1 | 1 | 0.131 | 7.1 | 5.3 | 4.5 | 5.9 | 4.8 | 0.3 | 0.52518 | 0.5113 | 1.61045 | 2.05509 | 0.96495 | 0.27897 | 0.32911 |
| 42 | Prunella vulgaris L. | Prunella vulgaris | 0.15 | 1 | 0.080 | 7.1 | 5.3 | 4.5 | 5.9 | 4.8 | 0.3 | 0.52518 | 0.5113 | 1.61045 | 2.05509 | 0.96495 | 0.27897 | 0.32911 |
| 42 | Geranium pusillum L. | Geranium pusillum | 0.02 | 1 | 0.050 | 7.3 | 5.4 | 4.1 | 7.6 | 4.2 | 0.4 | 0.58728 | 0.56055 | 1.34601 | 1.94534 | 0.82357 | 0.23743 | 0.34603 |
| 42 | Glechoma hederacea L. | Glechoma hederacea | 0.03 | 1 | 0.080 | 7.9 | NA | 4.1 | 5.8 | NA | 0.9 | 0.54139 | 0.51128 | 1.31363 | 1.93633 | 0.8112 | 0.25085 | 0.32953 |
| 43 | Poa annua L. | Poa annua | 0.55 | 0.8 | 0.021 | 7.3 | 5.4 | 4.1 | 7.6 | 4.2 | 0.4 | 0.58728 | 0.56055 | 1.34601 | 1.94534 | 0.82357 | 0.23743 | 0.34603 |
| 43 | Dipsacus fullonum L. | Dipsacus fullonum (sylvestris) | 0.1 | 0.8 | 0.020 | 7.3 | 5.4 | 4.1 | 7.6 | 4.2 | 0.4 | 0.58728 | 0.56055 | 1.34601 | 1.94534 | 0.82357 | 0.23743 | 0.34603 |
| 43 | Rubus pseudojaponicus Koidz. | Rubus caesius | 0.1 | 0.8 | 0.020 | 7.3 | 5.4 | 4.1 | 7.6 | 4.2 | 0.4 | 0.58728 | 0.56055 | 1.34601 | 1.94534 | 0.82357 | 0.23743 | 0.34603 |
| 43 | Taraxacum officinale F.H.Wigg. | Taraxacum officinale agg. | 0.05 | 0.8 | 0.100 | 7.3 | 5.4 | 4.1 | 7.6 | 4.2 | 0.4 | 0.58728 | 0.56055 | 1.34601 | 1.94534 | 0.82357 | 0.23743 | 0.34603 |
| 45 | Glechoma hederacea L. | Glechoma hederacea | 0.1 | 0.9 | 0.120 | 7.3 | 5.4 | 4.1 | 7.6 | 4.2 | 0.4 | 0.58728 | 0.56055 | 1.34601 | 1.94534 | 0.82357 | 0.23743 | 0.34603 |
| 45 | Sherardia arvensis L. | Sherardia arvensis | 0.15 | 0.9 | 0.052 | 7.3 | 5.4 | 4.1 | 7.6 | 4.2 | 0.4 | 0.58728 | 0.56055 | 1.34601 | 1.94534 | 0.82357 | 0.23743 | 0.34603 |
| 45 | Veronica persica Poir. | Veronica persica (tournefortii) | 0.1 | 0.9 | 0.020 | 7.3 | 5.4 | 4.1 | 7.6 | 4.2 | 0.4 | 0.58728 | 0.56055 | 1.34601 | 1.94534 | 0.82357 | 0.23743 | 0.34603 |
| 45 | Bellis perennis L. | Bellis perennis | 0.05 | 0.9 | 0.080 | 7.3 | 5.4 | 4.1 | 7.6 | 4.2 | 0.4 | 0.58728 | 0.56055 | 1.34601 | 1.94534 | 0.82357 | 0.23743 | 0.34603 |
| 45 | Lysimachia arvensis (L.) U.Manns & Anderb. | Anagallis arvensis | 0.1 | 0.9 | 0.063 | 7.3 | 5.4 | 4.1 | 7.6 | 4.2 | 0.4 | 0.58728 | 0.56055 | 1.34601 | 1.94534 | 0.82357 | 0.23743 | 0.34603 |
| 45 | Parthenocissus quinquefolia Planch. | NA | 0.05 | 0.9 | 0.053 | 7.3 | 5.4 | 4.1 | 7.6 | 4.2 | 0.4 | 0.58728 | 0.56055 | 1.34601 | 1.94534 | 0.82357 | 0.23743 | 0.34603 |
| 45 | Prunella vulgaris L. | Prunella vulgaris | 0.15 | 0.9 | 0.102 | 7.3 | 5.4 | 4.1 | 7.6 | 4.2 | 0.4 | 0.58728 | 0.56055 | 1.34601 | 1.94534 | 0.82357 | 0.23743 | 0.34603 |
| 45 | Sonchus oleraceus L. | Sonchus oleraceus | 0.05 | 0.9 | 0.050 | 7.3 | 5.4 | 4.1 | 7.6 | 4.2 | 0.4 | 0.58728 | 0.56055 | 1.34601 | 1.94534 | 0.82357 | 0.23743 | 0.34603 |
| 45 | Herniaria glabra L. | Herniaria glabra | 0.15 | 0.9 | 0.050 | 7.3 | 5.4 | 4.1 | 7.6 | 4.2 | 0.4 | 0.58728 | 0.56055 | 1.34601 | 1.94534 | 0.82357 | 0.23743 | 0.34603 |
| 47 | Glechoma hederacea L. | Glechoma hederacea | 0.25 | 0.95 | 0.050 | 7.3 | 5.4 | 4.1 | 7.6 | 4.2 | 0.4 | 0.58728 | 0.56055 | 1.34601 | 1.94534 | 0.82357 | 0.23743 | 0.34603 |
| 47 | Hypericum maculatum Crantz | Hypericum maculatum | 0.15 | 0.95 | 0.020 | 7.3 | 5.4 | 4.1 | 7.6 | 4.2 | 0.4 | 0.58728 | 0.56055 | 1.34601 | 1.94534 | 0.82357 | 0.23743 | 0.34603 |
| 47 | Taraxacum officinale F.H.Wigg. | Taraxacum officinale agg. | 0.45 | 0.95 | 0.080 | 7.3 | 5.4 | 4.1 | 7.6 | 4.2 | 0.4 | 0.58728 | 0.56055 | 1.34601 | 1.94534 | 0.82357 | 0.23743 | 0.34603 |
| 47 | Veronica persica Poir. | Veronica persica (tournefortii) | 0.05 | 0.95 | 0.050 | 7.9 | NA | 4.1 | 5.8 | NA | 0.9 | 0.54139 | 0.51128 | 1.31363 | 1.93633 | 0.8112 | 0.25085 | 0.32953 |
| 47 | Parthenocissus quinquefolia Planch. | NA | 0.05 | 0.95 | 0.080 | 7.9 | NA | 4.1 | 5.8 | NA | 0.9 | 0.54139 | 0.51128 | 1.31363 | 1.93633 | 0.8112 | 0.25085 | 0.32953 |
| 53 | Lysimachia arvensis (L.) U.Manns & Anderb. | Anagallis arvensis | 0.05 | 0.8 | 0.040 | 7.9 | NA | 4.1 | 5.8 | NA | 0.9 | 0.54139 | 0.51128 | 1.31363 | 1.93633 | 0.8112 | 0.25085 | 0.32953 |
| 53 | Taraxacum officinale F.H.Wigg. | Taraxacum officinale agg. | 0.12 | 0.8 | 0.429 | NA | NA | NA | NA | NA | NA | NA | NA | NA | NA | NA | NA | NA |
| 53 | Geranium molle L. | Geranium molle | 0.05 | 0.8 | 0.400 | 6.7 | NA | NA | 6.5 | 7.7 | 0.1 | 0.70945 | 0.61373 | 0.80583 | 1.7717 | 0.43958 | 0.26577 | 0.51423 |
| 53 | Valeriana locusta L. | Valerianella locusta (olitoria) | 0.05 | 0.8 | 0.500 | 7.7 | NA | NA | 5.6 | NA | 0.7 | 0.53223 | 0.52325 | 1.73719 | 2.14421 | 0.94278 | 0.39374 | 0.35054 |
| 53 | Galium aparine L. | Galium aparine aparine | 0.02 | 0.8 | 0.306 | 6.3 | NA | 5.9 | 6.3 | 6.9 | 0.1 | 0.6037 | 0.3946 | 0.44368 | 1.50139 | 0.30164 | 0.21315 | 0.21494 |
| 53 | Silene noctiflora L. | Silene noctiflora (Melandrium noct.) | 0.15 | 0.8 | 0.100 | NA | NA | NA | NA | NA | NA | NA | NA | NA | NA | NA | NA | NA |
| 53 | Sonchus oleraceus L. | Sonchus oleraceus | 0.05 | 0.8 | 0.150 | 7.4 | 5.3 | NA | 6 | 7.7 | 0.8 | 0.65201 | 0.63791 | 1.54898 | 2.03286 | 0.77966 | 0.37419 | 0.45721 |
| 53 | Poa annua L. | Poa annua | 0.31 | 0.8 | 0.455 | 5.2 | 5.9 | 5.4 | 7.3 | 8 | 0 | 0.67947 | 0.25214 | 0.1495 | 1.33661 | 0.07498 | 0.19025 | 0.12819 |
| 54 | Geranium molle L. | Geranium molle | 0.05 | 0.9 | 0.020 | NA | NA | NA | NA | NA | NA | NA | NA | NA | NA | NA | NA | NA |
| 54 | Lysimachia arvensis (L.) U.Manns & Anderb. | Anagallis arvensis | 0.05 | 0.9 | 1.000 | NA | NA | NA | NA | NA | NA | NA | NA | NA | NA | NA | NA | NA |
| 54 | Cirsium arvense (L.) Scop. | Cirsium arvense | 0.1 | 0.9 | 0.636 | 8.4 | NA | 4 | 6.3 | NA | 0.1 | 0.76525 | 0.73026 | 1.22636 | 1.78874 | 0.7034 | 0.16135 | 0.51807 |
| 54 | Poa annua L. | Poa annua | 0.7 | 0.9 | 0.133 | NA | NA | NA | NA | NA | NA | NA | NA | NA | NA | NA | NA | NA |
| 55 | Valeriana locusta L. | Valerianella locusta (olitoria) | 0.05 | 0.6 | 0.145 | NA | NA | NA | NA | NA | NA | NA | NA | NA | NA | NA | NA | NA |
| 55 | Geranium molle L. | Geranium molle | 0.02 | 0.6 | 0.659 | 7.2 | 5 | 5.1 | 6.1 | NA | 0.1 | 0.52316 | 0.5171 | 1.84298 | 2.10722 | 1.43318 | 0.13855 | 0.37874 |
| 55 | Veronica persica Poir. | Veronica persica (tournefortii) | 0.02 | 0.6 | 0.101 | 8.1 | NA | 3.5 | 7.5 | 5 | 0.3 | 0.75184 | 0.72654 | 1.3537 | 1.85024 | 0.717 | 0.18024 | 0.48387 |
| 55 | Poa annua L. | Poa annua | 0.49 | 0.6 | 0.481 | 7.2 | 5 | 5.1 | 6.1 | NA | 0.1 | 0.52316 | 0.5171 | 1.84298 | 2.10722 | 1.43318 | 0.13855 | 0.37874 |
| 55 | Galium aparine L. | Galium aparine aparine | 0.02 | 0.6 | 0.067 | 8.1 | 6.2 | NA | 7.1 | 7.6 | 0.1 | 0.8111 | 0.79007 | 1.4109 | 1.89172 | 0.89447 | 0.17462 | 0.49967 |
| 56 | Poa annua L. | Poa annua | 0.75 | 0.95 | 0.269 | 7.2 | 5 | 5.1 | 6.1 | NA | 0.1 | 0.52316 | 0.5171 | 1.84298 | 2.10722 | 1.43318 | 0.13855 | 0.37874 |
| 56 | Veronica hederifolia L. | Veronica hederifolia agg. | 0.05 | 0.95 | 1.000 | 7.4 | 5 | NA | NA | NA | 0.1 | 0.52506 | 0.51076 | 1.56059 | 2.06148 | 1.07287 | 0.38107 | 0.34662 |
| 56 | Bromus tectorum L. | Bromus tectorum | 0.15 | 0.95 | 0.250 | 7.4 | 5 | NA | NA | NA | 0.1 | 0.52506 | 0.51076 | 1.56059 | 2.06148 | 1.07287 | 0.38107 | 0.34662 |
| 57 | Poa annua L. | Poa annua | 0.65 | 0.85 | 0.280 | 6.7 | NA | NA | 6.5 | 7.7 | 0.1 | 0.70945 | 0.61373 | 0.80583 | 1.7717 | 0.43958 | 0.26577 | 0.51423 |
| 57 | Taraxacum officinale F.H.Wigg. | Taraxacum officinale agg. | 0.1 | 0.85 | 0.333 | 6.9 | 6.3 | 3.9 | 6.3 | 7.1 | 0.1 | 0.77916 | 0.67616 | 0.95204 | 1.78267 | 0.43197 | 0.25047 | 0.50688 |
| 57 | Sedum album L. | Sedum album | 0.1 | 0.85 | 0.640 | 7.4 | 5 | NA | NA | NA | 0.1 | 0.52506 | 0.51076 | 1.56059 | 2.06148 | 1.07287 | 0.38107 | 0.34662 |
| 58 | Urtica dioica L. | Urtica dioica | 0.05 | 0.9 | 0.650 | NA | NA | NA | NA | NA | NA | NA | NA | NA | NA | NA | NA | NA |
| 58 | Poa annua L. | Poa annua | 0.7 | 0.9 | 0.444 | NA | NA | NA | NA | NA | NA | NA | NA | NA | NA | NA | NA | NA |
| 58 | Ranunculus bulbosus L. | Ranunculus bulbosus | 0.05 | 0.9 | 0.316 | 7.4 | 5.3 | NA | 6 | 7.7 | 0.8 | 0.65201 | 0.63791 | 1.54898 | 2.03286 | 0.77966 | 0.37419 | 0.45721 |
| 58 | Pastinaca sativa L. | Pastinaca sativa | 0.1 | 0.9 | 0.300 | 7.4 | 5.3 | NA | 6 | 7.7 | 0.8 | 0.65201 | 0.63791 | 1.54898 | 2.03286 | 0.77966 | 0.37419 | 0.45721 |
| 59 | Poa annua L. | Poa annua | 0.85 | 0.95 | 0.200 | 7.4 | 5.3 | NA | 6 | 7.7 | 0.8 | 0.65201 | 0.63791 | 1.54898 | 2.03286 | 0.77966 | 0.37419 | 0.45721 |
| 59 | Bromus tectorum L. | Bromus tectorum | 0.1 | 0.95 | 1.000 | 7.4 | 5.3 | NA | 6 | 7.7 | 0.8 | 0.65201 | 0.63791 | 1.54898 | 2.03286 | 0.77966 | 0.37419 | 0.45721 |
| 60 | Poa annua L. | Poa annua | 0.8 | 0.85 | 0.222 | 7.4 | 5.3 | NA | 6 | 7.7 | 0.8 | 0.65201 | 0.63791 | 1.54898 | 2.03286 | 0.77966 | 0.37419 | 0.45721 |
| 60 | Cerastium fontanum Baumg. | Cerastium fontanum fontanum | 0.05 | 0.85 | 0.300 | 7.4 | 5.3 | NA | 6 | 7.7 | 0.8 | 0.65201 | 0.63791 | 1.54898 | 2.03286 | 0.77966 | 0.37419 | 0.45721 |
| 61 | Poa annua L. | Poa annua | 0.75 | 0.95 | 1.000 | 7.4 | 5.3 | NA | 6 | 7.7 | 0.8 | 0.65201 | 0.63791 | 1.54898 | 2.03286 | 0.77966 | 0.37419 | 0.45721 |
| 61 | Potentilla reptans L. | Potentilla reptans | 0.2 | 0.95 | 0.500 | 7.4 | 5.3 | NA | 6 | 7.7 | 0.8 | 0.65201 | 0.63791 | 1.54898 | 2.03286 | 0.77966 | 0.37419 | 0.45721 |
| 62 | Poa annua L. | Poa annua | 0.85 | 0.9 | 0.300 | 7.4 | 5.3 | NA | 6 | 7.7 | 0.8 | 0.65201 | 0.63791 | 1.54898 | 2.03286 | 0.77966 | 0.37419 | 0.45721 |
| 62 | Geum urbanum L. | Geum urbanum | 0.05 | 0.9 | 0.444 | 7.4 | 5.3 | NA | 6 | 7.7 | 0.8 | 0.65201 | 0.63791 | 1.54898 | 2.03286 | 0.77966 | 0.37419 | 0.45721 |
| 63 | Poa annua L. | Poa annua | 0.85 | 0.9 | 0.500 | 7.4 | 5.3 | NA | 6 | 7.7 | 0.8 | 0.65201 | 0.63791 | 1.54898 | 2.03286 | 0.77966 | 0.37419 | 0.45721 |
| 63 | Geum urbanum L. | Geum urbanum | 0.05 | 0.9 | 0.600 | 7.4 | 5.3 | NA | 6 | 7.7 | 0.8 | 0.65201 | 0.63791 | 1.54898 | 2.03286 | 0.77966 | 0.37419 | 0.45721 |

| | | | | | | | | | | | | | | | | | | |
|---|---|---|---|---|---|---|---|---|---|---|---|---|---|---|---|---|---|---|
| 64 | Poa annua L. | Poa annua | 0.8 | 0.85 | 0.316 | NA | NA | NA | NA | NA | NA | NA | NA | NA | NA | NA | NA | NA |
| 64 | Lepidium draba L. | Cardaria draba | 0.05 | 0.85 | 0.250 | NA | NA | NA | NA | NA | NA | NA | NA | NA | NA | NA | NA | NA |
| 65 | Poa annua L. | Poa annua | 0.55 | 0.7 | 0.090 | NA | NA | NA | NA | NA | NA | NA | NA | NA | NA | NA | NA | NA |
| 65 | Taraxacum officinale F.H.Wigg. | Taraxacum officinale agg. | 0.05 | 0.7 | 0.800 | 7.4 | 5.3 | NA | 6 | 7.7 | 0.8 | 0.65201 | 0.63791 | 1.54898 | 2.03286 | 0.77966 | 0.37419 | 0.45721 |
| 65 | Euphorbia peplus L. | Euphorbia peplus | 0.05 | 0.7 | 0.800 | NA | NA | NA | NA | NA | NA | NA | NA | NA | NA | NA | NA | NA |
| 65 | Lepidium draba L. | Cardaria draba | 0.05 | 0.7 | 0.200 | 7.2 | NA | NA | 6.5 | 8.4 | 0.1 | 0.5933 | 0.54722 | 1.04949 | 1.60983 | 0.64032 | 0.25162 | 0.31795 |
| 66 | Taraxacum officinale F.H.Wigg. | Taraxacum officinale agg. | 0.2 | 1 | 0.337 | NA | NA | NA | NA | NA | NA | NA | NA | NA | NA | NA | NA | NA |
| 66 | Poa annua L. | Poa annua | 0.2 | 1 | 0.295 | NA | NA | NA | NA | NA | NA | NA | NA | NA | NA | NA | NA | NA |
| 66 | Veronica hederifolia L. | Veronica hederifolia agg. | 0.3 | 1 | 0.350 | 7.4 | 5.3 | NA | 6 | 7.7 | 0.8 | 0.65201 | 0.63791 | 1.54898 | 2.03286 | 0.77966 | 0.37419 | 0.45721 |
| 66 | Geum urbanum L. | Geum urbanum | 0.05 | 1 | 0.308 | 7.4 | 5.3 | NA | 6 | 7.7 | 0.8 | 0.65201 | 0.63791 | 1.54898 | 2.03286 | 0.77966 | 0.37419 | 0.45721 |
| 66 | Cirsium vulgare (Savi) Ten. | Cirsium vulgare (lanceolatum) | 0.1 | 1 | 0.250 | 7.1 | 7.8 | 4.1 | 6.2 | 6 | 0.1 | 0.76637 | 0.65623 | 0.78647 | 1.82229 | 0.56568 | 0.18972 | 0.47454 |
| 66 | Arctium minus (Hill) Bernh. | Arctium minus | 0.1 | 1 | 1.000 | 7.4 | 5.3 | NA | 6 | 7.7 | 0.8 | 0.65201 | 0.63791 | 1.54898 | 2.03286 | 0.77966 | 0.37419 | 0.45721 |
| 66 | Lepidium draba L. | Cardaria draba | 0.05 | 1 | 0.100 | NA | NA | NA | NA | NA | NA | NA | NA | NA | NA | NA | NA | NA |
| 67 | Taraxacum officinale F.H.Wigg. | Taraxacum officinale agg. | 0.1 | 0.95 | 0.100 | 7.4 | 5.3 | NA | 6 | 7.7 | 0.8 | 0.65201 | 0.63791 | 1.54898 | 2.03286 | 0.77966 | 0.37419 | 0.45721 |
| 67 | Bellis perennis L. | Bellis perennis | 0.1 | 0.95 | 0.526 | 7.4 | 5.3 | NA | 6 | 7.7 | 0.8 | 0.65201 | 0.63791 | 1.54898 | 2.03286 | 0.77966 | 0.37419 | 0.45721 |
| 67 | Lysimachia arvensis (L.) U.Manns & Anderb. | Anagallis arvensis | 0.05 | 0.95 | 0.111 | NA | NA | NA | NA | NA | NA | NA | NA | NA | NA | NA | NA | NA |
| 67 | Veronica hederifolia L. | Veronica hederifolia agg. | 0.35 | 0.95 | 0.111 | 7.1 | 4.9 | NA | 6 | NA | 1 | 0.59517 | 0.59064 | 1.89772 | 2.12267 | 0.65091 | 0.41213 | 0.37185 |
| 67 | Poa annua L. | Poa annua | 0.35 | 0.95 | 0.053 | 7.1 | 4.9 | NA | 6 | NA | 1 | 0.59517 | 0.59064 | 1.89772 | 2.12267 | 0.65091 | 0.41213 | 0.37185 |
| 68 | Poa annua L. | Poa annua | 0.55 | 1 | 0.235 | 8.1 | NA | 3.9 | 6.9 | 6.4 | 0.4 | 0.80098 | 0.76713 | 1.29016 | 1.92909 | 0.75604 | 0.19458 | 0.52486 |
| 68 | Valeriana locusta L. | Valerianella locusta (olitoria) | 0.35 | 1 | 0.900 | 7.4 | 5.2 | 4.7 | 6.1 | NA | 0 | 0.77406 | 0.76726 | 1.79113 | 2.07279 | 0.64778 | 0.2119 | 0.65925 |
| 68 | Veronica hederifolia L. | Veronica hederifolia agg. | 0.1 | 1 | 0.350 | 7.4 | 5.3 | NA | 6 | 7.7 | 0.8 | 0.65201 | 0.63791 | 1.54898 | 2.03286 | 0.77966 | 0.37419 | 0.45721 |
| 69 | Poa annua L. | Poa annua | 0.2 | 0.8 | 0.900 | 7.4 | 5.3 | NA | 6 | 7.7 | 0.8 | 0.65201 | 0.63791 | 1.54898 | 2.03286 | 0.77966 | 0.37419 | 0.45721 |
| 69 | Veronica hederifolia L. | Veronica hederifolia agg. | 0.4 | 0.8 | 0.400 | 7.8 | 5.3 | 3.8 | 6.6 | 5 | 0.2 | 0.49579 | 0.46867 | 1.36306 | 1.85033 | 0.62791 | 0.29201 | 0.3217 |
| 69 | Valeriana locusta L. | Valerianella locusta (olitoria) | 0.2 | 0.8 | 0.750 | 6.7 | NA | NA | 6.5 | 7.7 | 0.1 | 0.70945 | 0.61373 | 0.80583 | 1.7717 | 0.43958 | 0.26577 | 0.51423 |
| 70 | Hordeum murinum L. | Hordeum murinum agg. | 0.2 | 1 | 0.200 | NA | NA | NA | NA | NA | NA | NA | NA | NA | NA | NA | NA | NA |
| 70 | Alliaria petiolata (M.Bieb.) | Alliaria petiolata | 0.2 | 1 | 0.600 | 7.7 | NA | NA | 5.6 | NA | 0.7 | 0.53223 | 0.52325 | 1.73719 | 2.14421 | 0.94278 | 0.39374 | 0.35054 |
| 70 | Geum urbanum L. | Geum urbanum | 0.2 | 1 | 0.500 | 7.4 | 5.3 | NA | 6 | 7.7 | 0.8 | 0.65201 | 0.63791 | 1.54898 | 2.03286 | 0.77966 | 0.37419 | 0.45721 |
| 70 | Sonchus asper (L.) Hill | Sonchus asper | 0.2 | 1 | 0.100 | 6.3 | NA | 5.9 | 6.3 | 6.9 | 0.1 | 0.6037 | 0.3946 | 0.44368 | 1.50139 | 0.30164 | 0.21315 | 0.21494 |
| 70 | Hypochaeris radicata L. | Hypochoeris radicata | 0.2 | 1 | 0.950 | 7.4 | 5.3 | NA | 6 | 7.7 | 0.8 | 0.65201 | 0.63791 | 1.54898 | 2.03286 | 0.77966 | 0.37419 | 0.45721 |
| 71 | Bellis perennis L. | Bellis perennis | 0.15 | 1 | 0.950 | 8.1 | NA | 3.9 | 6.9 | 6.4 | 0.4 | 0.80098 | 0.76713 | 1.29016 | 1.92909 | 0.75604 | 0.19458 | 0.52486 |
| 71 | Taraxacum officinale F.H.Wigg. | Taraxacum officinale agg. | 0.1 | 1 | 1.000 | 7.4 | 5.3 | NA | 6 | 7.7 | 0.8 | 0.65201 | 0.63791 | 1.54898 | 2.03286 | 0.77966 | 0.37419 | 0.45721 |
| 71 | Erodium cicutarium (L.) | Erodium cicutarium cicutarium | 0.05 | 1 | 0.944 | NA | NA | NA | NA | NA | NA | NA | NA | NA | NA | NA | NA | NA |
| 71 | Cerastium arvense L. | Cerastium arvense | 0.05 | 1 | 0.750 | 7.4 | 5.3 | NA | 6 | 7.7 | 0.8 | 0.65201 | 0.63791 | 1.54898 | 2.03286 | 0.77966 | 0.37419 | 0.45721 |
| 71 | Medicago lupulina L. | Medicago lupulina | 0.1 | 1 | 0.500 | NA | NA | NA | NA | NA | NA | NA | NA | NA | NA | NA | NA | NA |
| 71 | Poa annua L. | Poa annua | 0.45 | 1 | 0.050 | 6.7 | NA | NA | 6.5 | 7.7 | 0.1 | 0.70945 | 0.61373 | 0.80583 | 1.7717 | 0.43958 | 0.26577 | 0.51423 |
| 71 | Hypochaeris radicata L. | Hypochoeris radicata | 0.1 | 1 | 0.300 | 6.3 | 5 | 7.3 | 5.8 | 6.5 | 0.7 | 0.53692 | 0.48641 | 1.02734 | 1.80642 | 0.6097 | 0.33182 | 0.26598 |
| 72 | Bellis perennis L. | Bellis perennis | 0.05 | 1 | 0.125 | NA | NA | NA | NA | NA | NA | NA | NA | NA | NA | NA | NA | NA |
| 72 | Taraxacum officinale F.H.Wigg. | Taraxacum officinale agg. | 0.05 | 1 | 0.100 | 6.3 | NA | 5.9 | 6.3 | 6.9 | 0.1 | 0.6037 | 0.3946 | 0.44368 | 1.50139 | 0.30164 | 0.21315 | 0.21494 |
| 72 | Geranium molle L. | Geranium molle | 0.29 | 1 | 0.050 | NA | NA | 4.5 | 6.6 | 6.6 | 0 | 0.78059 | 0.60963 | 0.54418 | 1.63701 | 0.26733 | 0.16232 | 0.54354 |
| 72 | Cerastium arvense L. | Cerastium arvense | 0.24 | 1 | 0.050 | 7.3 | 6.1 | 3.8 | 6 | 5.1 | 0 | 0.62987 | 0.59281 | 1.27995 | 1.90491 | 0.6275 | 0.3036 | 0.42608 |
| 72 | Geranium molle L. | Geranium molle | 0.05 | 1 | 0.027 | 5.2 | 5.9 | 5.4 | 7.3 | 8 | 0 | 0.67947 | 0.25214 | 0.1495 | 1.33661 | 0.07498 | 0.19025 | 0.12819 |
| 72 | Medicago lupulina L. | Medicago lupulina | 0.1 | 1 | 0.250 | 7.4 | 5.3 | NA | 6 | 7.7 | 0.8 | 0.65201 | 0.63791 | 1.54898 | 2.03286 | 0.77966 | 0.37419 | 0.45721 |
| 72 | Cirsium vulgare (Savi) Ten. | Cirsium vulgare (lanceolatum) | 0.05 | 1 | 0.050 | NA | NA | 4.5 | 6.6 | 6.6 | 0 | 0.78059 | 0.60963 | 0.54418 | 1.63701 | 0.26733 | 0.16232 | 0.54354 |
| 72 | Poa annua L. | Poa annua | 0.15 | 1 | 0.147 | NA | NA | 4.5 | 6.6 | 6.6 | 0 | 0.78059 | 0.60963 | 0.54418 | 1.63701 | 0.26733 | 0.16232 | 0.54354 |
| 72 | Potentilla reptans L. | Potentilla reptans | 0.02 | 1 | 0.190 | 5.2 | 5.9 | 5.4 | 7.3 | 8 | 0 | 0.67947 | 0.25214 | 0.1495 | 1.33661 | 0.07498 | 0.19025 | 0.12819 |
| 73 | Bellis perennis L. | Bellis perennis | 0.1 | 1 | 0.303 | 5.2 | 5.9 | 5.4 | 7.3 | 8 | 0 | 0.67947 | 0.25214 | 0.1495 | 1.33661 | 0.07498 | 0.19025 | 0.12819 |
| 73 | Taraxacum officinale F.H.Wigg. | Taraxacum officinale agg. | 0.05 | 1 | 0.560 | 5.2 | 5.9 | 5.4 | 7.3 | 8 | 0 | 0.67947 | 0.25214 | 0.1495 | 1.33661 | 0.07498 | 0.19025 | 0.12819 |
| 73 | Sherardia arvensis L. | Sherardia arvensis | 0.05 | 1 | 0.190 | 5.2 | 5.9 | 5.4 | 7.3 | 8 | 0 | 0.67947 | 0.25214 | 0.1495 | 1.33661 | 0.07498 | 0.19025 | 0.12819 |
| 73 | Erodium cicutarium (L.) | Erodium cicutarium cicutarium | 0.02 | 1 | 0.800 | 7.4 | 5.3 | NA | 6 | 7.7 | 0.8 | 0.65201 | 0.63791 | 1.54898 | 2.03286 | 0.77966 | 0.37419 | 0.45721 |
| 73 | Medicago lupulina L. | Medicago lupulina | 0.1 | 1 | 0.300 | 5.2 | 5.9 | 5.4 | 7.3 | 8 | 0 | 0.67947 | 0.25214 | 0.1495 | 1.33661 | 0.07498 | 0.19025 | 0.12819 |
| 73 | Dipsacus fullonum L. | Dipsacus fullonum (sylvestris) | 0.05 | 1 | 0.100 | 5.2 | 5.9 | 5.4 | 7.3 | 8 | 0 | 0.67947 | 0.25214 | 0.1495 | 1.33661 | 0.07498 | 0.19025 | 0.12819 |
| 73 | Lolium perenne L. | Lolium perenne | 0.15 | 1 | 0.333 | 7 | 5.6 | 4.4 | 5.9 | 5.4 | 0.1 | 0.64871 | 0.63945 | 1.70759 | 2.05577 | 0.56937 | 0.21649 | 0.54803 |
| 73 | Cirsium vulgare (Savi) Ten. | Cirsium vulgare (lanceolatum) | 0.05 | 1 | 0.100 | 6.3 | NA | 5.9 | 6.3 | 6.9 | 0.1 | 0.6037 | 0.3946 | 0.44368 | 1.50139 | 0.30164 | 0.21315 | 0.21494 |
| 73 | Poa annua L. | Poa annua | 0.35 | 1 | 0.156 | 7.4 | 5.9 | 3.8 | 6 | NA | 0 | 0.7907 | 0.77966 | 1.64404 | 2.03844 | 0.57393 | 0.17512 | 0.69169 |
| 73 | Fragaria vesca L. | Fragaria vesca | 0.05 | 1 | 1.000 | 5.2 | 5.9 | 5.4 | 7.3 | 8 | 0 | 0.67947 | 0.25214 | 0.1495 | 1.33661 | 0.07498 | 0.19025 | 0.12819 |
| 73 | Potentilla reptans L. | Potentilla reptans | 0.03 | 1 | 0.153 | 6.3 | NA | 5.9 | 6.3 | 6.9 | 0.1 | 0.6037 | 0.3946 | 0.44368 | 1.50139 | 0.30164 | 0.21315 | 0.21494 |
| 74 | Bellis perennis L. | Bellis perennis | 0.05 | 0.6 | 0.250 | 7.4 | 5.3 | NA | 6 | 7.7 | 0.8 | 0.65201 | 0.63791 | 1.54898 | 2.03286 | 0.77966 | 0.37419 | 0.45721 |
| 74 | Hypochaeris radicata L. | Hypochoeris radicata | 0.1 | 0.6 | 0.110 | 6.3 | NA | 5.9 | 6.3 | 6.9 | 0.1 | 0.6037 | 0.3946 | 0.44368 | 1.50139 | 0.30164 | 0.21315 | 0.21494 |
| 74 | Poa annua L. | Poa annua | 0.13 | 0.6 | 0.150 | 5.2 | 5.9 | 5.4 | 7.3 | 8 | 0 | 0.67947 | 0.25214 | 0.1495 | 1.33661 | 0.07498 | 0.19025 | 0.12819 |
| 74 | Medicago lupulina L. | Medicago lupulina | 0.2 | 0.6 | 0.100 | 6.3 | NA | 5.9 | 6.3 | 6.9 | 0.1 | 0.6037 | 0.3946 | 0.44368 | 1.50139 | 0.30164 | 0.21315 | 0.21494 |
| 74 | Cerastium arvense L. | Cerastium arvense | 0.05 | 0.6 | 0.210 | 6.3 | NA | 5.9 | 6.3 | 6.9 | 0.1 | 0.6037 | 0.3946 | 0.44368 | 1.50139 | 0.30164 | 0.21315 | 0.21494 |
| 74 | Geranium molle L. | Geranium molle | 0.05 | 0.6 | 0.620 | 7.2 | NA | NA | 6.5 | 8.4 | 0.1 | 0.5933 | 0.54722 | 1.04949 | 1.60983 | 0.64032 | 0.25162 | 0.31795 |
| 74 | Geranium pusillum L. | Geranium pusillum | 0.02 | 0.6 | 0.286 | 7.4 | 5.3 | NA | 6 | 7.7 | 0.8 | 0.65201 | 0.63791 | 1.54898 | 2.03286 | 0.77966 | 0.37419 | 0.45721 |
| 75 | Bellis perennis L. | Bellis perennis | 0.15 | 0.9 | 0.111 | 6.9 | 5.9 | NA | 6.8 | 7 | 0 | 0.84157 | 0.83772 | 1.85474 | 2.04042 | 0.4856 | 0.14178 | 0.80173 |
| 75 | Taraxacum officinale F.H.Wigg. | Taraxacum officinale agg. | 0.08 | 0.9 | 0.300 | 7.4 | 5.3 | NA | 6 | 7.7 | 0.8 | 0.65201 | 0.63791 | 1.54898 | 2.03286 | 0.77966 | 0.37419 | 0.45721 |
| 75 | Poa annua L. | Poa annua | 0.24 | 0.9 | 0.200 | 6.3 | NA | 5.9 | 6.3 | 6.9 | 0.1 | 0.6037 | 0.3946 | 0.44368 | 1.50139 | 0.30164 | 0.21315 | 0.21494 |
| 75 | Geranium molle L. | Geranium molle | 0.05 | 0.9 | 0.211 | 6.3 | NA | 5.9 | 6.3 | 6.9 | 0.1 | 0.6037 | 0.3946 | 0.44368 | 1.50139 | 0.30164 | 0.21315 | 0.21494 |
| 75 | Potentilla grandiflora L. | Potentilla grandiflora | 0.2 | 0.9 | 0.118 | NA | NA | NA | NA | NA | NA | NA | NA | NA | NA | NA | NA | NA |
| 75 | Medicago lupulina L. | Medicago lupulina | 0.1 | 0.9 | 0.200 | NA | NA | NA | NA | NA | NA | NA | NA | NA | NA | NA | NA | NA |
| 75 | Cerastium arvense L. | Cerastium arvense | 0.08 | 0.9 | 0.100 | 7.3 | 6.1 | 3.8 | 6 | 5.1 | 0 | 0.62987 | 0.59281 | 1.27995 | 1.90491 | 0.6275 | 0.3036 | 0.42608 |
| 76 | Bellis perennis L. | Bellis perennis | 0.1 | 0.93 | 0.200 | 8.4 | NA | 3 | 6 | NA | 0 | 0.46249 | 0.43915 | 1.39189 | 1.83687 | 0.38437 | 0.30571 | 0.38883 |
| 76 | Taraxacum officinale F.H.Wigg. | Taraxacum officinale agg. | 0.05 | 0.93 | 0.050 | 7.7 | NA | NA | 5.6 | NA | 0.7 | 0.53223 | 0.52325 | 1.73719 | 2.14421 | 0.94278 | 0.39374 | 0.35054 |
| 76 | Poa annua L. | Poa annua | 0.19 | 0.93 | 0.091 | 7.7 | NA | NA | 5.6 | NA | 0.7 | 0.53223 | 0.52325 | 1.73719 | 2.14421 | 0.94278 | 0.39374 | 0.35054 |
| 76 | Geranium molle L. | Geranium molle | 0.2 | 0.93 | 0.136 | 8.1 | NA | NA | 5.2 | NA | 0.1 | 0.51276 | 0.49581 | 1.51667 | 2.20989 | 0.98959 | 0.46997 | 0.3594 |
| 76 | Cerastium arvense L. | Cerastium arvense | 0.05 | 0.93 | 0.050 | 8.1 | NA | NA | 5.2 | NA | 0.1 | 0.51276 | 0.49581 | 1.51667 | 2.20989 | 0.98959 | 0.46997 | 0.3594 |
| 76 | Medicago lupulina L. | Medicago lupulina | 0.2 | 0.93 | 0.650 | 7 | 7 | 5 | 7 | 7 | 0 | NA | NA | NA | NA | NA | NA | NA |
| 76 | Achillea millefolium L. | Achillea millefolium millefolium | 0.05 | 0.93 | 0.824 | 7.4 | 5.3 | NA | 6 | 7.7 | 0.8 | 0.65201 | 0.63791 | 1.54898 | 2.03286 | 0.77966 | 0.37419 | 0.45721 |
| 76 | Geranium pusillum L. | Geranium pusillum | 0.02 | 0.93 | 0.300 | NA | NA | 5.3 | 6.2 | NA | 0.1 | 0.68682 | 0.14816 | 0.05813 | 1.31254 | 0.01433 | 0.20126 | 0.10505 |
| 76 | Capsella bursa-pastoris Medik. | Capsella bursa-pastoris | 0.02 | 0.93 | 0.200 | 7.3 | NA | NA | 5.8 | 5.9 | 0.5 | 0.63207 | 0.61738 | 1.52973 | 2.00883 | 0.70322 | 0.36696 | 0.39725 |
| 76 | Hypochaeris radicata L. | Hypochoeris radicata | 0.05 | 0.93 | 0.600 | 5.2 | 5.6 | 4.6 | 7.5 | 3.8 | 0 | 0.65844 | 0.17774 | 0.10594 | 1.36663 | 0.08038 | 0.20249 | 0.12191 |
| 77 | Bellis perennis L. | Bellis perennis | 0.08 | 0.85 | 0.278 | 7.7 | NA | NA | 5.6 | NA | 0.7 | 0.53223 | 0.52325 | 1.73719 | 2.14421 | 0.94278 | 0.39374 | 0.35054 |
| 77 | Taraxacum officinale F.H.Wigg. | Taraxacum officinale agg. | 0.08 | 0.85 | 0.450 | NA | NA | NA | NA | NA | NA | NA | NA | NA | NA | NA | NA | NA |
| 77 | Lolium perenne L. | Lolium perenne | 0.1 | 0.85 | 0.600 | 7.4 | 5.3 | NA | 6 | 7.7 | 0.8 | 0.65201 | 0.63791 | 1.54898 | 2.03286 | 0.77966 | 0.37419 | 0.45721 |
| 77 | Medicago lupulina L. | Medicago lupulina | 0.1 | 0.85 | 0.600 | 7.4 | 5.3 | NA | 6 | 7.7 | 0.8 | 0.65201 | 0.63791 | 1.54898 | 2.03286 | 0.77966 | 0.37419 | 0.45721 |
| 77 | Myosotis ramosissima Rochel | Myosotis ramosissima (coll. Hispida) | 0.05 | 0.85 | 0.150 | 7.7 | NA | NA | 5.6 | NA | 0.7 | 0.53223 | 0.52325 | 1.73719 | 2.14421 | 0.94278 | 0.39374 | 0.35054 |
| 77 | Veronica arvensis L. | Veronica arvensis | 0.02 | 0.85 | 0.200 | 7.7 | NA | NA | 5.6 | NA | 0.7 | 0.53223 | 0.52325 | 1.73719 | 2.14421 | 0.94278 | 0.39374 | 0.35054 |
| 77 | Cerastium semidecandrum L. | Cerastium semidecandrum | 0.04 | 0.85 | 0.250 | 7.7 | NA | NA | 5.6 | NA | 0.7 | 0.53223 | 0.52325 | 1.73719 | 2.14421 | 0.94278 | 0.39374 | 0.35054 |
| 77 | Geranium molle L. | Geranium molle | 0.05 | 0.85 | 0.300 | 7.7 | NA | NA | 5.6 | NA | 0.7 | 0.53223 | 0.52325 | 1.73719 | 2.14421 | 0.94278 | 0.39374 | 0.35054 |
| 77 | Vicia sativa L. | Vicia sativa agg. | 0.02 | 0.85 | 0.100 | 7.4 | 5.3 | NA | 6 | 7.7 | 0.8 | 0.65201 | 0.63791 | 1.54898 | 2.03286 | 0.77966 | 0.37419 | 0.45721 |
| 77 | Hypochaeris radicata L. | Hypochoeris radicata | 0.1 | 0.85 | 0.158 | NA | NA | NA | NA | NA | NA | NA | NA | NA | NA | NA | NA | NA |
| 77 | Trifolium dubium Sibth. | Trifolium dubium (minus) | 0.08 | 0.85 | 0.444 | NA | NA | NA | NA | NA | NA | NA | NA | NA | NA | NA | NA | NA |
| 77 | Prunella vulgaris L. | Prunella vulgaris | 0.09 | 0.85 | 0.333 | NA | NA | NA | NA | NA | NA | NA | NA | NA | NA | NA | NA | NA |
| 77 | Cerastium fontanum Baumg. | Cerastium fontanum fontanum | 0.04 | 0.85 | 0.118 | NA | NA | NA | NA | NA | NA | NA | NA | NA | NA | NA | NA | NA |
| 78 | Bellis perennis L. | Bellis perennis | 0.29 | 0.92 | 0.056 | NA | NA | NA | NA | NA | NA | NA | NA | NA | NA | NA | NA | NA |
| 78 | Taraxacum officinale F.H.Wigg. | Taraxacum officinale agg. | 0.05 | 0.92 | 0.600 | 7.4 | 5.3 | NA | 6 | 7.7 | 0.8 | 0.65201 | 0.63791 | 1.54898 | 2.03286 | 0.77966 | 0.37419 | 0.45721 |
| 78 | Lolium perenne L. | Lolium perenne | 0.1 | 0.92 | 0.100 | 7.4 | 5.3 | NA | 6 | 7.7 | 0.8 | 0.65201 | 0.63791 | 1.54898 | 2.03286 | 0.77966 | 0.37419 | 0.45721 |
| 78 | Geum urbanum L. | Geum urbanum | 0.05 | 0.92 | 0.300 | 7.4 | 5.3 | NA | 6 | 7.7 | 0.8 | 0.65201 | 0.63791 | 1.54898 | 2.03286 | 0.77966 | 0.37419 | 0.45721 |
| 78 | Myosotis ramosissima Rochel | Myosotis ramosissima (coll. Hispida) | 0.02 | 0.92 | 0.579 | NA | NA | NA | NA | NA | NA | NA | NA | NA | NA | NA | NA | NA |
| 78 | Trifolium repens L. | Trifolium repens | 0.05 | 0.92 | 0.970 | 7.3 | NA | 4 | 6.7 | 5.6 | 0.1 | 0.69239 | 0.67894 | 1.55406 | 1.88846 | 0.70954 | 0.18169 | 0.49207 |
| 78 | Prunella vulgaris L. | Prunella vulgaris | 0.23 | 0.92 | 1.000 | NA | NA | NA | NA | NA | NA | NA | NA | NA | NA | NA | NA | NA |
| 78 | Veronica arvensis L. | Veronica arvensis | 0.02 | 0.92 | 1.000 | NA | NA | NA | NA | NA | NA | NA | NA | NA | NA | NA | NA | NA |
| 78 | Ranunculus repens L. | Ranunculus repens | 0.04 | 0.92 | 0.694 | NA | NA | NA | NA | NA | NA | NA | NA | NA | NA | NA | NA | NA |
| 78 | Fragaria vesca L. | Fragaria vesca | 0.05 | 0.92 | 0.230 | NA | NA | NA | NA | NA | NA | NA | NA | NA | NA | NA | NA | NA |
| 78 | Myosotis arvensis Hill | Myosotis arvensis (intermedia) | 0.02 | 0.92 | 0.038 | NA | NA | NA | NA | NA | NA | NA | NA | NA | NA | NA | NA | NA |
| 79 | Bellis perennis L. | Bellis perennis | 0.15 | 0.87 | 0.118 | 8.4 | NA | 3 | 6 | NA | 0 | 0.46249 | 0.43915 | 1.39189 | 1.83687 | 0.38437 | 0.30571 | 0.38883 |
| 79 | Taraxacum officinale F.H.Wigg. | Taraxacum officinale agg. | 0.09 | 0.87 | 0.268 | 8.1 | NA | NA | 5.2 | NA | 0.1 | 0.51276 | 0.49581 | 1.51667 | 2.20989 | 0.98959 | 0.46997 | 0.3594 |

| | | | | | | | | | | | | | | | | | | |
|---|---|---|---|---|---|---|---|---|---|---|---|---|---|---|---|---|---|---|
| 79 | Lolium perenne L. | Lolium perenne | 0.09 | 0.87 | 0.186 | 8.1 | NA | NA | 5.2 | NA | 0.1 | 0.51276 | 0.49581 | 1.51667 | 2.20989 | 0.98959 | 0.46997 | 0.3594 |
| 79 | Cerastium arvense L. | Cerastium arvense | 0.04 | 0.87 | 0.083 | 8.1 | NA | NA | 5.2 | NA | 0.1 | 0.51276 | 0.49581 | 1.51667 | 2.20989 | 0.98959 | 0.46997 | 0.3594 |
| 79 | Veronica serpyllifolia L. | Veronica serpyllifolia | 0.05 | 0.87 | 0.020 | 8.5 | NA | 2.5 | NA | NA | 0 | 0.506 | 0.44835 | 1.02121 | 1.67002 | 0.34316 | 0.25824 | 0.34742 |
| 79 | Medicago lupulina L. | Medicago lupulina | 0.1 | 0.87 | 0.042 | 8.5 | NA | 2.5 | NA | NA | 0 | 0.506 | 0.44835 | 1.02121 | 1.67002 | 0.34316 | 0.25824 | 0.34742 |
| 79 | Veronica arvensis L. | Veronica arvensis | 0.02 | 0.87 | 0.270 | 7.4 | 5.3 | NA | 6 | 7.7 | 0.8 | 0.65201 | 0.63791 | 1.54898 | 2.03286 | 0.77966 | 0.37419 | 0.45721 |
| 79 | Fragaria vesca L. | Fragaria vesca | 0.04 | 0.87 | 0.050 | 6.5 | NA | NA | 5.4 | NA | 0 | 0.59769 | 0.45571 | 0.64594 | 1.51584 | 0.28825 | 0.22901 | 0.32316 |
| 79 | Cerastium fontanum Baumg. | Cerastium fontanum fontanum | 0.04 | 0.87 | 0.050 | 6.5 | NA | NA | 5.4 | NA | 0 | 0.59769 | 0.45571 | 0.64594 | 1.51584 | 0.28825 | 0.22901 | 0.32316 |
| 79 | Achillea millefolium L. | Achillea millefolium millefolium | 0.01 | 0.87 | 0.053 | 6.5 | NA | NA | 5.4 | NA | 0 | 0.59769 | 0.45571 | 0.64594 | 1.51584 | 0.28825 | 0.22901 | 0.32316 |
| 79 | Cerastium semidecandrum L. | Cerastium semidecandrum | 0.04 | 0.87 | 0.200 | NA | NA | NA | NA | NA | NA | NA | NA | NA | NA | NA | NA | NA |
| 79 | Prunella vulgaris L. | Prunella vulgaris | 0.1 | 0.87 | 1.000 | 7.1 | 4.9 | NA | 6 | NA | 1 | 0.59517 | 0.59064 | 1.89772 | 2.12267 | 0.65091 | 0.41213 | 0.37185 |
| 79 | Hypochaeris radicata L. | Hypochoeris radicata | 0.1 | 0.87 | 0.556 | NA | NA | NA | NA | NA | NA | NA | NA | NA | NA | NA | NA | NA |
| 80 | Bellis perennis L. | Bellis perennis | 0.1 | 0.92 | 0.320 | NA | NA | NA | NA | NA | NA | NA | NA | NA | NA | NA | NA | NA |
| 80 | Taraxacum officinale F.H.Wigg. | Taraxacum officinale agg. | 0.09 | 0.92 | 0.611 | NA | NA | NA | NA | NA | NA | NA | NA | NA | NA | NA | NA | NA |
| 80 | Lolium perenne L. | Lolium perenne | 0.1 | 0.92 | 0.600 | NA | NA | NA | NA | NA | NA | NA | NA | NA | NA | NA | NA | NA |
| 80 | Cerastium arvense L. | Cerastium arvense | 0.05 | 0.92 | 0.500 | NA | NA | NA | NA | NA | NA | NA | NA | NA | NA | NA | NA | NA |
| 80 | Veronica serpyllifolia L. | Veronica serpyllifolia | 0.08 | 0.92 | 0.200 | NA | NA | NA | NA | NA | NA | NA | NA | NA | NA | NA | NA | NA |
| 80 | Myosotis ramosissima Rochel | Myosotis ramosissima (coll. Hispida) | 0.01 | 0.92 | 0.204 | NA | NA | NA | NA | NA | NA | NA | NA | NA | NA | NA | NA | NA |
| 80 | Geranium molle L. | Geranium molle | 0.08 | 0.92 | 0.106 | NA | NA | NA | NA | NA | NA | NA | NA | NA | NA | NA | NA | NA |
| 80 | Medicago lupulina L. | Medicago lupulina | 0.05 | 0.92 | 0.300 | 7.4 | 5.3 | NA | 6 | 7.7 | 0.8 | 0.65201 | 0.63791 | 1.54898 | 2.03286 | 0.77966 | 0.37419 | 0.45721 |
| 80 | Cerastium fontanum Baumg. | Cerastium fontanum fontanum | 0.05 | 0.92 | 0.220 | 7 | NA | NA | NA | NA | 0.3 | 0.56899 | 0.4288 | 0.64817 | 1.63031 | 0.49671 | 0.22558 | 0.25973 |
| 80 | Trifolium repens L. | Trifolium repens | 0.05 | 0.92 | 0.100 | NA | NA | NA | NA | NA | NA | NA | NA | NA | NA | NA | NA | NA |
| 80 | Cerastium semidecandrum L. | Cerastium semidecandrum | 0.04 | 0.92 | 0.193 | 7 | NA | NA | NA | NA | 0.3 | 0.56899 | 0.4288 | 0.64817 | 1.63031 | 0.49671 | 0.22558 | 0.25973 |
| 80 | Achillea millefolium L. | Achillea millefolium millefolium | 0.01 | 0.92 | 0.700 | NA | NA | NA | NA | NA | NA | NA | NA | NA | NA | NA | NA | NA |
| 80 | Fragaria vesca L. | Fragaria vesca | 0.04 | 0.92 | 0.684 | NA | NA | NA | NA | NA | NA | NA | NA | NA | NA | NA | NA | NA |
| 80 | Veronica arvensis L. | Veronica arvensis | 0.01 | 0.92 | 0.150 | NA | NA | NA | NA | NA | NA | NA | NA | NA | NA | NA | NA | NA |
| 80 | Poa annua L. | Poa annua | 0.08 | 0.92 | 0.100 | NA | NA | NA | NA | NA | NA | NA | NA | NA | NA | NA | NA | NA |
| 80 | Prunella vulgaris L. | Prunella vulgaris | 0.08 | 0.92 | 0.050 | NA | NA | NA | NA | NA | NA | NA | NA | NA | NA | NA | NA | NA |
| 81 | Bellis perennis L. | Bellis perennis | 0.1 | 0.67 | 0.050 | NA | NA | NA | NA | NA | NA | NA | NA | NA | NA | NA | NA | NA |
| 81 | Taraxacum officinale F.H.Wigg. | Taraxacum officinale agg. | 0.1 | 0.67 | 0.060 | NA | NA | NA | NA | NA | NA | NA | NA | NA | NA | NA | NA | NA |
| 81 | Lolium perenne L. | Lolium perenne | 0.08 | 0.67 | 0.062 | NA | NA | NA | NA | NA | NA | NA | NA | NA | NA | NA | NA | NA |
| 81 | Cerastium arvense L. | Cerastium arvense | 0.05 | 0.67 | 0.250 | NA | NA | NA | NA | NA | NA | NA | NA | NA | NA | NA | NA | NA |
| 81 | Veronica serpyllifolia L. | Veronica serpyllifolia | 0.05 | 0.67 | 0.200 | NA | NA | NA | NA | NA | NA | NA | NA | NA | NA | NA | NA | NA |
| 81 | Medicago lupulina L. | Medicago lupulina | 0.04 | 0.67 | 0.800 | NA | NA | NA | NA | NA | NA | NA | NA | NA | NA | NA | NA | NA |
| 81 | Ranunculus repens L. | Ranunculus repens | 0.04 | 0.67 | 0.340 | 7.3 | NA | NA | 7.6 | 7.7 | 0.4 | 0.75479 | 0.72524 | 1.28692 | 1.77374 | 0.61324 | 0.18283 | 0.56773 |
| 81 | Cerastium semidecandrum L. | Cerastium semidecandrum | 0.02 | 0.67 | 0.333 | NA | NA | NA | NA | NA | NA | NA | NA | NA | NA | NA | NA | NA |
| 81 | Veronica arvensis L. | Veronica arvensis | 0.02 | 0.67 | 0.556 | NA | NA | NA | NA | NA | NA | NA | NA | NA | NA | NA | NA | NA |
| 81 | Geranium molle L. | Geranium molle | 0.04 | 0.67 | 0.200 | NA | NA | NA | NA | NA | NA | NA | NA | NA | NA | NA | NA | NA |
| 81 | Hypochaeris radicata L. | Hypochoeris radicata | 0.08 | 0.67 | 0.342 | 7.4 | 5.3 | NA | 6 | 7.7 | 0.8 | 0.65201 | 0.63791 | 1.54898 | 2.03286 | 0.77966 | 0.37419 | 0.45721 |
| 81 | Poa annua L. | Poa annua | 0.05 | 0.67 | 0.273 | 8.1 | NA | 3.9 | 6.9 | 6.4 | 0.4 | 0.80098 | 0.76713 | 1.29016 | 1.92909 | 0.75604 | 0.19458 | 0.52486 |
| 82 | Bellis perennis L. | Bellis perennis | 0.25 | 1.02 | 0.600 | 7.4 | 5.3 | NA | 6 | 7.7 | 0.8 | 0.65201 | 0.63791 | 1.54898 | 2.03286 | 0.77966 | 0.37419 | 0.45721 |
| 82 | Taraxacum officinale F.H.Wigg. | Taraxacum officinale agg. | 0.2 | 1.02 | 0.667 | NA | NA | NA | NA | NA | NA | NA | NA | NA | NA | NA | NA | NA |
| 82 | Poa annua L. | Poa annua | 0.16 | 1.02 | 0.200 | 8.1 | NA | 3.1 | 6.6 | NA | 0 | 0.56066 | 0.54465 | 1.58041 | 1.97627 | 0.33838 | 0.29416 | 0.45216 |
| 82 | Trifolium repens L. | Trifolium repens | 0.3 | 1.02 | 0.600 | NA | NA | NA | NA | NA | NA | NA | NA | NA | NA | NA | NA | NA |
| 82 | Geranium molle L. | Geranium molle | 0.05 | 1.02 | 0.500 | NA | NA | NA | NA | NA | NA | NA | NA | NA | NA | NA | NA | NA |
| 82 | Medicago lupulina L. | Medicago lupulina | 0.02 | 1.02 | 0.750 | NA | NA | NA | NA | NA | NA | NA | NA | NA | NA | NA | NA | NA |
| 82 | Sherardia arvensis L. | Sherardia arvensis | 0.04 | 1.02 | 0.400 | NA | NA | NA | NA | NA | NA | NA | NA | NA | NA | NA | NA | NA |
| 83 | Bellis perennis L. | Bellis perennis | 0.4 | 1 | 0.400 | 7.4 | 5.3 | NA | 6 | 7.7 | 0.8 | 0.65201 | 0.63791 | 1.54898 | 2.03286 | 0.77966 | 0.37419 | 0.45721 |
| 83 | Taraxacum officinale F.H.Wigg. | Taraxacum officinale agg. | 0.08 | 1 | 0.122 | 7.4 | 5.3 | NA | 6 | 7.7 | 0.8 | 0.65201 | 0.63791 | 1.54898 | 2.03286 | 0.77966 | 0.37419 | 0.45721 |
| 83 | Veronica arvensis L. | Veronica arvensis | 0.07 | 1 | 0.294 | 7.4 | 5.3 | NA | 6 | 7.7 | 0.8 | 0.65201 | 0.63791 | 1.54898 | 2.03286 | 0.77966 | 0.37419 | 0.45721 |
| 83 | Cerastium arvense L. | Cerastium arvense | 0.05 | 1 | 0.083 | 7.4 | 5.3 | NA | 6 | 7.7 | 0.8 | 0.65201 | 0.63791 | 1.54898 | 2.03286 | 0.77966 | 0.37419 | 0.45721 |
| 83 | Cerastium fontanum Baumg. | Cerastium fontanum fontanum | 0.25 | 1 | 0.556 | 7 | 5.6 | 4.4 | 5.9 | 5.4 | 0.1 | 0.64871 | 0.63945 | 1.70759 | 2.05577 | 0.56937 | 0.21649 | 0.54803 |
| 83 | Geranium molle L. | Geranium molle | 0.15 | 1 | 0.200 | 7 | 5.6 | 4.4 | 5.9 | 5.4 | 0.1 | 0.64871 | 0.63945 | 1.70759 | 2.05577 | 0.56937 | 0.21649 | 0.54803 |
| 84 | Glechoma hederacea L. | Glechoma hederacea | 0.15 | 1 | 1.000 | 6.5 | NA | NA | 5.4 | NA | 0 | 0.59769 | 0.45571 | 0.64594 | 1.51584 | 0.28825 | 0.22901 | 0.32316 |
| 84 | Geum urbanum L. | Geum urbanum | 0.2 | 1 | 0.114 | 7.1 | 5.3 | 4.5 | 5.9 | 4.8 | 0.3 | 0.52518 | 0.5113 | 1.61045 | 2.05509 | 0.96495 | 0.27897 | 0.32911 |
| 84 | Populus alba L. | Populus alba | 0.25 | 1 | 0.059 | 7.1 | 5.3 | 4.5 | 5.9 | 4.8 | 0.3 | 0.52518 | 0.5113 | 1.61045 | 2.05509 | 0.96495 | 0.27897 | 0.32911 |
| 84 | Bromus diandrus Roth | NA | 0.05 | 1 | 0.100 | 6.2 | 5.5 | NA | 6.9 | 8.5 | 0.1 | 0.60658 | 0.39025 | 0.44214 | 1.39281 | 0.24736 | 0.18526 | 0.18179 |
| 84 | Lapsana communis L. | Lapsana communis | 0.2 | 1 | 0.100 | 7.4 | 5.8 | 4.2 | 6.7 | 5.3 | 0 | 0.62702 | 0.59749 | 1.35516 | 1.96644 | 0.57704 | 0.23154 | 0.49642 |
| 84 | Carduus nutant L. | Carduus nutans agg. | 0.05 | 1 | 0.400 | 7.2 | 5.7 | 4.3 | 6.8 | NA | 0.1 | 0.54405 | 0.41742 | 0.70503 | 1.63482 | 0.49706 | 0.18636 | 0.24515 |
| 84 | Taraxacum officinale F.H.Wigg. | Taraxacum officinale agg. | 0.1 | 1 | 0.444 | 7.4 | 5.3 | NA | 6 | 7.7 | 0.8 | 0.65201 | 0.63791 | 1.54898 | 2.03286 | 0.77966 | 0.37419 | 0.45721 |
| 85 | Populus alba L. | Populus alba | 0.25 | 0.45 | 0.571 | 7.4 | 5.3 | NA | 6 | 7.7 | 0.8 | 0.65201 | 0.63791 | 1.54898 | 2.03286 | 0.77966 | 0.37419 | 0.45721 |
| 85 | Achillea millefolium L. | Achillea millefolium millefolium | 0.1 | 0.45 | 0.071 | 7.4 | 5.8 | 4.2 | 6.7 | 5.3 | 0 | 0.62702 | 0.59749 | 1.35516 | 1.96644 | 0.57704 | 0.23154 | 0.49642 |
| 85 | Veronica arvensis L. | Veronica arvensis | 0.05 | 0.45 | 0.467 | 7.4 | 5.3 | NA | 6 | 7.7 | 0.8 | 0.65201 | 0.63791 | 1.54898 | 2.03286 | 0.77966 | 0.37419 | 0.45721 |
| 85 | Glechoma hederacea L. | Glechoma hederacea | 0.05 | 0.45 | 0.550 | 7.4 | 5.3 | NA | 6 | 7.7 | 0.8 | 0.65201 | 0.63791 | 1.54898 | 2.03286 | 0.77966 | 0.37419 | 0.45721 |
| 86 | Populus alba L. | Populus alba | 0.35 | 0.7 | 0.389 | 7.4 | 5.3 | NA | 6 | 7.7 | 0.8 | 0.65201 | 0.63791 | 1.54898 | 2.03286 | 0.77966 | 0.37419 | 0.45721 |
| 86 | Achillea millefolium L. | Achillea millefolium millefolium | 0.05 | 0.7 | 0.330 | 7.4 | 5.3 | NA | 6 | 7.7 | 0.8 | 0.65201 | 0.63791 | 1.54898 | 2.03286 | 0.77966 | 0.37419 | 0.45721 |
| 86 | Medicago lupulina L. | Medicago lupulina | 0.13 | 0.7 | 0.125 | 7.9 | 6.2 | NA | 6.7 | 6.4 | 0.2 | 0.62749 | 0.58201 | 1.11218 | 1.53794 | 0.69579 | 0.21134 | 0.22838 |
| 86 | Vicia sativa L. | Vicia sativa agg. | 0.08 | 0.7 | 0.167 | 7.2 | 6 | 4.1 | 6.8 | 4.9 | 0 | 0.72892 | 0.658 | 1.07912 | 1.91928 | 0.406 | 0.27624 | 0.47339 |
| 86 | Bromus diandrus Roth | NA | 0.05 | 0.7 | 0.158 | 7.3 | 4.1 | 5.7 | 3.7 | 2.9 | 0 | 0.50708 | 0.41392 | 0.80188 | 1.46639 | 0.47762 | 0.18385 | 0.2417 |
| 86 | Veronica arvensis L. | Veronica arvensis | 0.02 | 0.7 | 0.150 | NA | NA | NA | NA | NA | NA | NA | NA | NA | NA | NA | NA | NA |
| 86 | Myosotis arvensis Hill | Myosotis arvensis (intermedia) | 0.02 | 0.7 | 0.056 | 6.8 | 6.2 | 4.4 | NA | NA | 0.3 | 0.7871 | 0.75023 | 1.37944 | 1.92919 | 0.40631 | 0.19607 | 0.63681 |
| 87 | Populus alba L. | Populus alba | 0.6 | 0.75 | 0.895 | 7.4 | 5.3 | NA | 6 | 7.7 | 0.8 | 0.65201 | 0.63791 | 1.54898 | 2.03286 | 0.77966 | 0.37419 | 0.45721 |
| 87 | Achillea millefolium L. | Achillea millefolium millefolium | 0.05 | 0.75 | 0.941 | 7.4 | 5.3 | NA | 6 | 7.7 | 0.8 | 0.65201 | 0.63791 | 1.54898 | 2.03286 | 0.77966 | 0.37419 | 0.45721 |
| 87 | Medicago lupulina L. | Medicago lupulina | 0.1 | 0.75 | 0.789 | 7.4 | 5.3 | NA | 6 | 7.7 | 0.8 | 0.65201 | 0.63791 | 1.54898 | 2.03286 | 0.77966 | 0.37419 | 0.45721 |
| 88 | Populus alba L. | Populus alba | 0.6 | 0.85 | 0.944 | 7.4 | 5.3 | NA | 6 | 7.7 | 0.8 | 0.65201 | 0.63791 | 1.54898 | 2.03286 | 0.77966 | 0.37419 | 0.45721 |
| 88 | Achillea millefolium L. | Achillea millefolium millefolium | 0.2 | 0.85 | 0.944 | 7.4 | 5.3 | NA | 6 | 7.7 | 0.8 | 0.65201 | 0.63791 | 1.54898 | 2.03286 | 0.77966 | 0.37419 | 0.45721 |
| 88 | Medicago lupulina L. | Medicago lupulina | 0.05 | 0.85 | 0.941 | 7.4 | 5.3 | NA | 6 | 7.7 | 0.8 | 0.65201 | 0.63791 | 1.54898 | 2.03286 | 0.77966 | 0.37419 | 0.45721 |
| 89 | Populus alba L. | Populus alba | 0.93 | 0.93 | 0.071 | NA | NA | NA | NA | NA | NA | NA | NA | NA | NA | NA | NA | NA |
| 90 | Populus alba L. | Populus alba | 0.93 | 0.98 | 0.200 | 7.4 | 5.3 | NA | 6 | 7.7 | 0.8 | 0.65201 | 0.63791 | 1.54898 | 2.03286 | 0.77966 | 0.37419 | 0.45721 |
| 90 | Senecio inaequidens DC. | Senecio inaequidens | 0.05 | 0.98 | 0.105 | NA | NA | NA | NA | NA | NA | NA | NA | NA | NA | NA | NA | NA |
| 91 | Populus alba L. | Populus alba | 0.77 | 1 | 0.350 | 7.4 | 5.8 | 4.2 | 6.7 | 5.3 | 0 | 0.62702 | 0.59749 | 1.35516 | 1.96644 | 0.57704 | 0.23154 | 0.49642 |
| 91 | Achillea millefolium L. | Achillea millefolium millefolium | 0.1 | 1 | 0.250 | 7.4 | 5.3 | NA | 6 | 7.7 | 0.8 | 0.65201 | 0.63791 | 1.54898 | 2.03286 | 0.77966 | 0.37419 | 0.45721 |
| 91 | Medicago lupulina L. | Medicago lupulina | 0.08 | 1 | 0.200 | 5.2 | 5.9 | 5.4 | 7.3 | 8 | 0 | 0.67947 | 0.25214 | 0.1495 | 1.33661 | 0.07498 | 0.19025 | 0.12819 |
| 91 | Poa annua L. | Poa annua | 0.05 | 1 | 0.100 | NA | NA | NA | NA | NA | NA | NA | NA | NA | NA | NA | NA | NA |
| 92 | Populus alba L. | Populus alba | 0.63 | 0.9 | 0.050 | NA | NA | NA | NA | NA | NA | NA | NA | NA | NA | NA | NA | NA |
| 92 | Achillea millefolium L. | Achillea millefolium millefolium | 0.1 | 0.9 | 0.050 | NA | NA | NA | NA | NA | NA | NA | NA | NA | NA | NA | NA | NA |
| 92 | Erodium cicutarium (L.) | Erodium cicutarium cicutarium | 0.15 | 0.9 | 0.167 | 8.1 | NA | NA | 4.4 | 2.8 | 0.3 | 0.48868 | 0.46508 | 1.4426 | 2.04965 | 0.7278 | 0.33931 | 0.35628 |
| 92 | Medicago lupulina L. | Medicago lupulina | 0.02 | 0.9 | 0.089 | NA | NA | NA | NA | NA | NA | NA | NA | NA | NA | NA | NA | NA |
| 93 | Populus alba L. | Populus alba | 0.35 | 0.95 | 0.054 | NA | NA | NA | NA | NA | NA | NA | NA | NA | NA | NA | NA | NA |
| 93 | Achillea millefolium L. | Achillea millefolium millefolium | 0.1 | 0.95 | 0.094 | NA | NA | NA | NA | NA | NA | NA | NA | NA | NA | NA | NA | NA |
| 93 | Erodium cicutarium (L.) | Erodium cicutarium cicutarium | 0.1 | 0.95 | 0.054 | NA | NA | NA | NA | NA | NA | NA | NA | NA | NA | NA | NA | NA |
| 93 | Poa annua L. | Poa annua | 0.38 | 0.95 | 0.103 | NA | NA | NA | NA | NA | NA | NA | NA | NA | NA | NA | NA | NA |
| 93 | Medicago lupulina L. | Medicago lupulina | 0.02 | 0.95 | 0.098 | NA | NA | NA | NA | NA | NA | NA | NA | NA | NA | NA | NA | NA |
| 94 | Medicago lupulina L. | Medicago lupulina | 0.05 | 1 | 0.149 | NA | NA | NA | NA | NA | NA | NA | NA | NA | NA | NA | NA | NA |
| 94 | Taraxacum officinale F.H.Wigg. | Taraxacum officinale agg. | 0.1 | 1 | 0.196 | NA | NA | NA | NA | NA | NA | NA | NA | NA | NA | NA | NA | NA |
| 94 | Erodium cicutarium (L.) | Erodium cicutarium cicutarium | 0.1 | 1 | 0.080 | NA | NA | NA | NA | NA | NA | NA | NA | NA | NA | NA | NA | NA |
| 94 | Potentilla argentea L. | Potentilla argentea agg. | 0.15 | 1 | 0.200 | NA | 5.4 | 5.7 | 6.8 | 7 | 0 | 0.67742 | 0.24559 | 0.14546 | 1.36309 | 0.06547 | 0.19208 | 0.12644 |
| 94 | Herniaria glabra L. | Herniaria glabra | 0.05 | 1 | 0.071 | 7.9 | NA | 4.1 | 5.8 | NA | 0.9 | 0.54139 | 0.51128 | 1.31363 | 1.93633 | 0.8112 | 0.25085 | 0.32953 |
| 94 | Poa annua L. | Poa annua | 0.2 | 1 | 0.067 | 7.9 | NA | 4.1 | 5.8 | NA | 0.9 | 0.54139 | 0.51128 | 1.31363 | 1.93633 | 0.8112 | 0.25085 | 0.32953 |
| 94 | Bromus sterilis L. | Bromus sterilis | 0.25 | 1 | 0.235 | 7.9 | NA | 4.1 | 5.8 | NA | 0.9 | 0.54139 | 0.51128 | 1.31363 | 1.93633 | 0.8112 | 0.25085 | 0.32953 |
| 94 | Sedum acre L. | Sedum acre | 0.1 | 1 | 0.100 | 7.9 | NA | 4.1 | 5.8 | NA | 0.9 | 0.54139 | 0.51128 | 1.31363 | 1.93633 | 0.8112 | 0.25085 | 0.32953 |
| 95 | Medicago lupulina L. | Medicago lupulina | 0.05 | 1 | 0.111 | 7.9 | NA | 4.1 | 5.8 | NA | 0.9 | 0.54139 | 0.51128 | 1.31363 | 1.93633 | 0.8112 | 0.25085 | 0.32953 |
| 95 | Taraxacum officinale F.H.Wigg. | Taraxacum officinale agg. | 0.15 | 1 | 0.105 | 7.9 | NA | 4.1 | 5.8 | NA | 0.9 | 0.54139 | 0.51128 | 1.31363 | 1.93633 | 0.8112 | 0.25085 | 0.32953 |
| 95 | Achillea millefolium L. | Achillea millefolium millefolium | 0.08 | 1 | 0.100 | NA | NA | NA | NA | NA | NA | NA | NA | NA | NA | NA | NA | NA |
| 95 | Achillea millefolium L. | Achillea millefolium millefolium | 0.05 | 1 | 0.150 | NA | NA | NA | NA | NA | NA | NA | NA | NA | NA | NA | NA | NA |
| 95 | Potentilla argentea L. | Potentilla argentea agg. | 0.08 | 1 | 0.150 | NA | NA | NA | NA | NA | NA | NA | NA | NA | NA | NA | NA | NA |
| 95 | Poa annua L. | Poa annua | 0.05 | 1 | 0.120 | NA | NA | NA | NA | NA | NA | NA | NA | NA | NA | NA | NA | NA |

| | | | | | | | | | | | | | | | | | | |
|---|---|---|---|---|---|---|---|---|---|---|---|---|---|---|---|---|---|---|
| 95 | Bromus sterilis L. | Bromus sterilis | 0.2 | 1 | 0.100 | NA | NA | NA | NA | NA | NA | NA | NA | NA | NA | NA | NA | NA |
| 95 | Bromus hordeaceus L. | Bromus hordeaceus (mollis) | 0.15 | 1 | 0.053 | NA | NA | NA | NA | NA | NA | NA | NA | NA | NA | NA | NA | NA |
| 95 | Cerastium semidecandrum L. | Cerastium semidecandrum | 0.05 | 1 | 0.050 | 8.6 | NA | 2.7 | 4.4 | 1.6 | 0 | 0.53557 | 0.50485 | 1.29532 | 1.90211 | 0.72298 | 0.28495 | 0.2778 |
| 95 | Sedum acre L. | Sedum acre | 0.1 | 1 | 0.100 | 6.7 | 5.3 | 4.5 | 5.3 | 5.8 | 0 | 0.75464 | 0.73051 | 1.36974 | 1.95508 | 0.45974 | 0.15567 | 0.6659 |
| 95 | Veronica arvensis L. | Veronica arvensis | 0.04 | 1 | 0.050 | 8.6 | NA | 2.7 | 4.4 | 1.6 | 0 | 0.53557 | 0.50485 | 1.29532 | 1.90211 | 0.72298 | 0.28495 | 0.2778 |
| 96 | Medicago lupulina L. | Medicago lupulina | 0.02 | 1 | 0.050 | NA | NA | NA | NA | NA | NA | NA | NA | NA | NA | NA | NA | NA |
| 96 | Taraxacum officinale F.H.Wigg. | Taraxacum officinale agg. | 0.15 | 1 | 0.100 | 8.6 | NA | 2.7 | 4.4 | 1.6 | 0 | 0.53557 | 0.50485 | 1.29532 | 1.90211 | 0.72298 | 0.28495 | 0.2778 |
| 96 | Veronica arvensis L. | Veronica arvensis | 0.02 | 1 | 0.080 | 8.6 | NA | 2.7 | 4.4 | 1.6 | 0 | 0.53557 | 0.50485 | 1.29532 | 1.90211 | 0.72298 | 0.28495 | 0.2778 |
| 96 | Achillea millefolium L. | Achillea millefolium millefolium | 0.1 | 1 | 0.050 | 8.6 | NA | 2.7 | 4.4 | 1.6 | 0 | 0.53557 | 0.50485 | 1.29532 | 1.90211 | 0.72298 | 0.28495 | 0.2778 |
| 96 | Potentilla argentea L. | Potentilla argentea agg. | 0.1 | 1 | 0.100 | 8.6 | NA | 2.7 | 4.4 | 1.6 | 0 | 0.53557 | 0.50485 | 1.29532 | 1.90211 | 0.72298 | 0.28495 | 0.2778 |
| 96 | Poa annua L. | Poa annua | 0.1 | 1 | 0.060 | 8.6 | NA | 2.7 | 4.4 | 1.6 | 0 | 0.53557 | 0.50485 | 1.29532 | 1.90211 | 0.72298 | 0.28495 | 0.2778 |
| 96 | Bromus sterilis L. | Bromus sterilis | 0.51 | 1 | 0.105 | NA | NA | NA | NA | NA | NA | NA | NA | NA | NA | NA | NA | NA |
| 97 | Medicago lupulina L. | Medicago lupulina | 0.08 | 1 | 0.200 | NA | NA | NA | NA | NA | NA | NA | NA | NA | NA | NA | NA | NA |
| 97 | Taraxacum officinale F.H.Wigg. | Taraxacum officinale agg. | 0.12 | 1 | 0.100 | 8.6 | NA | 2.7 | 4.4 | 1.6 | 0 | 0.53557 | 0.50485 | 1.29532 | 1.90211 | 0.72298 | 0.28495 | 0.2778 |
| 97 | Veronica arvensis L. | Veronica arvensis | 0.05 | 1 | 0.100 | 8.1 | NA | 3.5 | 6.1 | NA | 0.2 | 0.68175 | 0.6716 | 1.70824 | 1.99767 | 0.41848 | 0.25411 | 0.57318 |
| 97 | Erodium cicutarium (L.) | Erodium cicutarium cicutarium | 0.08 | 1 | 0.111 | 8.1 | NA | 3.5 | 6.1 | NA | 0.2 | 0.68175 | 0.6716 | 1.70824 | 1.99767 | 0.41848 | 0.25411 | 0.57318 |
| 97 | Achillea millefolium L. | Achillea millefolium millefolium | 0.05 | 1 | 0.154 | 8.1 | NA | 3.5 | 6.1 | NA | 0.2 | 0.68175 | 0.6716 | 1.70824 | 1.99767 | 0.41848 | 0.25411 | 0.57318 |
| 97 | Potentilla argentea L. | Potentilla argentea agg. | 0.05 | 1 | 0.167 | 8.1 | NA | 3.5 | 6.1 | NA | 0.2 | 0.68175 | 0.6716 | 1.70824 | 1.99767 | 0.41848 | 0.25411 | 0.57318 |
| 97 | Cerastium semidecandrum L. | Cerastium semidecandrum | 0.05 | 1 | 0.286 | 8.1 | NA | 3.5 | 6.1 | NA | 0.2 | 0.68175 | 0.6716 | 1.70824 | 1.99767 | 0.41848 | 0.25411 | 0.57318 |
| 97 | Myosotis arvensis Hill | Myosotis arvensis (intermedia) | 0.05 | 1 | 0.178 | 8.1 | NA | 3.5 | 6.1 | NA | 0.2 | 0.68175 | 0.6716 | 1.70824 | 1.99767 | 0.41848 | 0.25411 | 0.57318 |
| 97 | Bellis perennis L. | Bellis perennis | 0.1 | 1 | 0.080 | 8.1 | NA | 3.5 | 6.1 | NA | 0.2 | 0.68175 | 0.6716 | 1.70824 | 1.99767 | 0.41848 | 0.25411 | 0.57318 |
| 97 | Poa annua L. | Poa annua | 0.24 | 1 | 0.083 | 8.5 | NA | 3.5 | 6.3 | 4.1 | 0.2 | 0.57633 | 0.55499 | 1.44489 | 1.81436 | 0.579 | 0.25616 | 0.27657 |
| 97 | Bromus sterilis L. | Bromus sterilis | 0.13 | 1 | 0.345 | 7.9 | NA | 4.1 | 5.8 | NA | 0.9 | 0.54139 | 0.51128 | 1.31363 | 1.93633 | 0.8112 | 0.25085 | 0.32953 |
| 98 | Medicago lupulina L. | Medicago lupulina | 0.1 | 1 | 0.379 | 7.9 | NA | 4.1 | 5.8 | NA | 0.9 | 0.54139 | 0.51128 | 1.31363 | 1.93633 | 0.8112 | 0.25085 | 0.32953 |
| 98 | Taraxacum officinale F.H.Wigg. | Taraxacum officinale agg. | 0.1 | 1 | 0.250 | 8.1 | NA | 3.5 | 6.1 | NA | 0.2 | 0.68175 | 0.6716 | 1.70824 | 1.99767 | 0.41848 | 0.25411 | 0.57318 |
| 98 | Potentilla argentea L. | Potentilla argentea agg. | 0.05 | 1 | 0.100 | 8.5 | NA | 3.5 | 6.3 | 4.1 | 0.2 | 0.57633 | 0.55499 | 1.44489 | 1.81436 | 0.579 | 0.25616 | 0.27657 |
| 98 | Plantago lanceolata L. | Plantago lanceolata | 0.1 | 1 | 0.050 | 7.3 | 6.1 | 3.8 | 6 | 5.1 | 0 | 0.62987 | 0.59281 | 1.27995 | 1.90491 | 0.6275 | 0.3036 | 0.42608 |
| 98 | Myosotis arvensis Hill | Myosotis arvensis (intermedia) | 0.08 | 1 | 0.227 | 7.3 | 6.1 | 3.8 | 6 | 5.1 | 0 | 0.62987 | 0.59281 | 1.27995 | 1.90491 | 0.6275 | 0.3036 | 0.42608 |
| 98 | Cerastium semidecandrum L. | Cerastium semidecandrum | 0.05 | 1 | 0.158 | NA | NA | NA | NA | NA | NA | NA | NA | NA | NA | NA | NA | NA |
| 98 | Lactuca virosa Hablitz | Lactuca virosa | 0.1 | 1 | 0.080 | NA | NA | NA | NA | NA | NA | NA | NA | NA | NA | NA | NA | NA |
| 98 | Poa annua L. | Poa annua | 0.2 | 1 | 0.250 | 8.6 | NA | 2.7 | 4.4 | 1.6 | 0 | 0.53557 | 0.50485 | 1.29532 | 1.90211 | 0.72298 | 0.28495 | 0.2778 |
| 98 | Bromus sterilis L. | Bromus sterilis | 0.2 | 1 | 0.050 | NA | NA | NA | NA | NA | NA | NA | NA | NA | NA | NA | NA | NA |
| 98 | Lapsana communis L. | Lapsana communis | 0.02 | 1 | 0.102 | NA | NA | NA | NA | NA | NA | NA | NA | NA | NA | NA | NA | NA |
| 99 | Medicago lupulina L. | Medicago lupulina | 0.08 | 0.95 | 0.150 | NA | NA | NA | NA | NA | NA | NA | NA | NA | NA | NA | NA | NA |
| 99 | Taraxacum officinale F.H.Wigg. | Taraxacum officinale agg. | 0.05 | 0.95 | 0.459 | NA | NA | NA | NA | NA | NA | NA | NA | NA | NA | NA | NA | NA |
| 99 | Erodium cicutarium (L.) | Erodium cicutarium cicutarium | 0.08 | 0.95 | 0.100 | NA | NA | NA | NA | NA | NA | NA | NA | NA | NA | NA | NA | NA |
| 99 | Achillea millefolium L. | Achillea millefolium millefolium | 0.04 | 0.95 | 0.150 | NA | NA | NA | NA | NA | NA | NA | NA | NA | NA | NA | NA | NA |
| 99 | Potentilla argentea L. | Potentilla argentea agg. | 0.05 | 0.95 | 0.050 | NA | NA | NA | NA | NA | NA | NA | NA | NA | NA | NA | NA | NA |
| 99 | Poa annua L. | Poa annua | 0.23 | 0.95 | 0.200 | NA | NA | NA | NA | NA | NA | NA | NA | NA | NA | NA | NA | NA |
| 99 | Bromus sterilis L. | Bromus sterilis | 0.15 | 0.95 | 0.250 | NA | NA | NA | NA | NA | NA | NA | NA | NA | NA | NA | NA | NA |
| 99 | Cerastium semidecandrum L. | Cerastium semidecandrum | 0.1 | 0.95 | 0.144 | 7.1 | 5.3 | 4.5 | 5.9 | 4.8 | 0.3 | 0.52518 | 0.5113 | 1.61045 | 2.05509 | 0.96495 | 0.27897 | 0.32911 |
| 99 | Rumex acetosella L. | Rumex acetosella acetosella | 0.05 | 0.95 | 0.050 | NA | NA | NA | NA | NA | NA | NA | NA | NA | NA | NA | NA | NA |
| 99 | Myosotis arvensis Hill | Myosotis arvensis (intermedia) | 0.02 | 0.95 | 0.200 | NA | NA | NA | NA | NA | NA | NA | NA | NA | NA | NA | NA | NA |
| 99 | Plantago lanceolata L. | Plantago lanceolata | 0.1 | 0.95 | 0.170 | 7.3 | NA | NA | 5.8 | 5.9 | 0.5 | 0.63207 | 0.61738 | 1.52973 | 2.00883 | 0.70322 | 0.36696 | 0.39725 |
| 100 | Erodium cicutarium (L.) | Erodium cicutarium cicutarium | 0.1 | 1 | 0.050 | 7 | 5.6 | 4.4 | 5.9 | 5.4 | 0.1 | 0.64871 | 0.63945 | 1.70759 | 2.05577 | 0.56937 | 0.21649 | 0.54803 |
| 100 | Potentilla argentea L. | Potentilla argentea agg. | 0.05 | 1 | 0.172 | 8.1 | NA | NA | 5.2 | NA | 0.1 | 0.51276 | 0.49581 | 1.51667 | 2.20989 | 0.98959 | 0.46997 | 0.3594 |
| 100 | Poa annua L. | Poa annua | 0.35 | 1 | 0.211 | 6.7 | NA | NA | 6.5 | 7.7 | 0.1 | 0.70945 | 0.61373 | 0.80583 | 1.7717 | 0.43958 | 0.26577 | 0.51423 |
| 100 | Bromus sterilis L. | Bromus sterilis | 0.3 | 1 | 0.350 | 6.7 | NA | 4.9 | 6.8 | 5 | 0 | 0.66057 | 0.18432 | 0.11528 | 1.34075 | 0.06671 | 0.20032 | 0.12881 |
| 100 | Medicago lupulina L. | Medicago lupulina | 0.06 | 1 | 0.350 | 6.7 | NA | 4.9 | 6.8 | 5 | 0 | 0.66057 | 0.18432 | 0.11528 | 1.34075 | 0.06671 | 0.20032 | 0.12881 |
| 100 | Myosotis arvensis Hill | Myosotis arvensis (intermedia) | 0.02 | 1 | 0.040 | 6.7 | NA | 4.9 | 6.8 | 5 | 0 | 0.66057 | 0.18432 | 0.11528 | 1.34075 | 0.06671 | 0.20032 | 0.12881 |
| 100 | Sonchus asper (L.) Hill | Sonchus asper | 0.05 | 1 | 0.130 | 6.3 | NA | 5.9 | 6.3 | 6.9 | 0.1 | 0.6037 | 0.3946 | 0.44368 | 1.50139 | 0.30164 | 0.21315 | 0.21494 |
| 100 | Echium vulgare L. | Echium vulgare | 0.02 | 1 | 0.060 | 6.3 | NA | 5.9 | 6.3 | 6.9 | 0.1 | 0.6037 | 0.3946 | 0.44368 | 1.50139 | 0.30164 | 0.21315 | 0.21494 |
| 100 | Plantago coronopus L. | Plantago coronopus | 0.05 | 1 | 0.097 | 6.7 | NA | 4.9 | 6.8 | 5 | 0 | 0.66057 | 0.18432 | 0.11528 | 1.34075 | 0.06671 | 0.20032 | 0.12881 |
| 101 | Taraxacum officinale F.H.Wigg. | Taraxacum officinale agg. | 0.1 | 1 | 0.077 | NA | NA | NA | 7 | NA | 0 | 0.66957 | 0.20824 | 0.12967 | 1.34554 | 0.07292 | 0.20428 | 0.13248 |
| 101 | Myosotis arvensis Hill | Myosotis arvensis (intermedia) | 0.1 | 1 | 0.103 | NA | NA | NA | 7 | NA | 0 | 0.66957 | 0.20824 | 0.12967 | 1.34554 | 0.07292 | 0.20428 | 0.13248 |
| 101 | Veronica arvensis L. | Veronica arvensis | 0.02 | 1 | 0.100 | 7.7 | NA | NA | 5.6 | NA | 0.7 | 0.53223 | 0.52325 | 1.73719 | 2.14421 | 0.94278 | 0.39374 | 0.35054 |
| 101 | Echium vulgare L. | Echium vulgare | 0.05 | 1 | 0.263 | 7.1 | 5.3 | 4.5 | 5.9 | 4.8 | 0.3 | 0.52518 | 0.5113 | 1.61045 | 2.05509 | 0.96495 | 0.27897 | 0.32911 |
| 101 | Erodium cicutarium (L.) | Erodium cicutarium cicutarium | 0.05 | 1 | 0.235 | 7.3 | 5.4 | 4.1 | 7.6 | 4.2 | 0.4 | 0.58728 | 0.56055 | 1.34601 | 1.94534 | 0.82357 | 0.23743 | 0.34603 |
| 101 | Potentilla argentea L. | Potentilla argentea agg. | 0.05 | 1 | 0.200 | 7.3 | 5.4 | 4.1 | 7.6 | 4.2 | 0.4 | 0.58728 | 0.56055 | 1.34601 | 1.94534 | 0.82357 | 0.23743 | 0.34603 |
| 101 | Poa annua L. | Poa annua | 0.37 | 1 | 0.200 | 7.3 | 5.4 | 4.1 | 7.6 | 4.2 | 0.4 | 0.58728 | 0.56055 | 1.34601 | 1.94534 | 0.82357 | 0.23743 | 0.34603 |
| 101 | Bromus sterilis L. | Bromus sterilis | 0.21 | 1 | 0.250 | 7.3 | 5.4 | 4.1 | 7.6 | 4.2 | 0.4 | 0.58728 | 0.56055 | 1.34601 | 1.94534 | 0.82357 | 0.23743 | 0.34603 |
| 101 | Hypochaeris radicata L. | Hypochoeris radicata | 0.05 | 1 | 0.050 | 8.4 | NA | 3 | 6 | NA | 0 | 0.46249 | 0.43915 | 1.39189 | 1.83687 | 0.38437 | 0.30571 | 0.38883 |
| 102 | Erodium cicutarium (L.) | Erodium cicutarium cicutarium | 0.05 | 1 | 0.100 | 7.9 | NA | 4.1 | 5.8 | NA | 0.9 | 0.54139 | 0.51128 | 1.31363 | 1.93633 | 0.8112 | 0.25085 | 0.32953 |
| 102 | Potentilla argentea L. | Potentilla argentea agg. | 0.05 | 1 | 0.045 | 7.9 | NA | 4.1 | 5.8 | NA | 0.9 | 0.54139 | 0.51128 | 1.31363 | 1.93633 | 0.8112 | 0.25085 | 0.32953 |
| 102 | Myosotis arvensis Hill | Myosotis arvensis (intermedia) | 0.05 | 1 | 0.200 | 8.6 | NA | 2.7 | 4.4 | 1.6 | 0 | 0.53557 | 0.50485 | 1.29532 | 1.90211 | 0.72298 | 0.28495 | 0.2778 |
| 102 | Veronica arvensis L. | Veronica arvensis | 0.02 | 1 | 0.260 | 7.1 | 5.3 | 4.5 | 5.9 | 4.8 | 0.3 | 0.52518 | 0.5113 | 1.61045 | 2.05509 | 0.96495 | 0.27897 | 0.32911 |
| 102 | Cerastium semidecandrum L. | Cerastium semidecandrum | 0.05 | 1 | 0.050 | 8.4 | NA | 3 | 6 | NA | 0 | 0.46249 | 0.43915 | 1.39189 | 1.83687 | 0.38437 | 0.30571 | 0.38883 |
| 102 | Poa annua L. | Poa annua | 0.33 | 1 | 0.100 | 7.3 | 5.4 | 4.1 | 7.6 | 4.2 | 0.4 | 0.58728 | 0.56055 | 1.34601 | 1.94534 | 0.82357 | 0.23743 | 0.34603 |
| 102 | Bromus sterilis L. | Bromus sterilis | 0.45 | 1 | 0.100 | 7.1 | 5.3 | 4.5 | 5.9 | 4.8 | 0.3 | 0.52518 | 0.5113 | 1.61045 | 2.05509 | 0.96495 | 0.27897 | 0.32911 |
| 103 | Medicago lupulina L. | Medicago lupulina | 0.08 | 1 | 0.150 | 7.9 | NA | 4.1 | 5.8 | NA | 0.9 | 0.54139 | 0.51128 | 1.31363 | 1.93633 | 0.8112 | 0.25085 | 0.32953 |
| 103 | Taraxacum officinale F.H.Wigg. | Taraxacum officinale agg. | 0.05 | 1 | 0.123 | 8.1 | NA | 3.5 | 6.1 | NA | 0.2 | 0.68175 | 0.6716 | 1.70824 | 1.99767 | 0.41848 | 0.25411 | 0.57318 |
| 103 | Veronica arvensis L. | Veronica arvensis | 0.04 | 1 | 0.047 | 7.3 | 5.4 | 4.1 | 7.6 | 4.2 | 0.4 | 0.58728 | 0.56055 | 1.34601 | 1.94534 | 0.82357 | 0.23743 | 0.34603 |
| 103 | Myosotis arvensis Hill | Myosotis arvensis (intermedia) | 0.04 | 1 | 0.125 | 7.1 | 5.3 | 4.5 | 5.9 | 4.8 | 0.3 | 0.52518 | 0.5113 | 1.61045 | 2.05509 | 0.96495 | 0.27897 | 0.32911 |
| 103 | Hypochaeris radicata L. | Hypochoeris radicata | 0.1 | 1 | 0.080 | 7.1 | 5.3 | 4.5 | 5.9 | 4.8 | 0.3 | 0.52518 | 0.5113 | 1.61045 | 2.05509 | 0.96495 | 0.27897 | 0.32911 |
| 103 | Cerastium semidecandrum L. | Cerastium semidecandrum | 0.1 | 1 | 0.050 | 7.3 | 5.4 | 4.1 | 7.6 | 4.2 | 0.4 | 0.58728 | 0.56055 | 1.34601 | 1.94534 | 0.82357 | 0.23743 | 0.34603 |
| 103 | Poa annua L. | Poa annua | 0.35 | 1 | 0.066 | 8.1 | NA | NA | 5.2 | NA | 0.1 | 0.51276 | 0.49581 | 1.51667 | 2.20989 | 0.98959 | 0.46997 | 0.3594 |
| 103 | Bromus sterilis L. | Bromus sterilis | 0.19 | 1 | 0.108 | 7.3 | 5.4 | 4.1 | 7.6 | 4.2 | 0.4 | 0.58728 | 0.56055 | 1.34601 | 1.94534 | 0.82357 | 0.23743 | 0.34603 |
| 103 | Rosa fujisanensis Makino | NA | 0.05 | 1 | 0.090 | 7.1 | 5.3 | 4.5 | 5.9 | 4.8 | 0.3 | 0.52518 | 0.5113 | 1.61045 | 2.05509 | 0.96495 | 0.27897 | 0.32911 |
| 104 | Medicago lupulina L. | Medicago lupulina | 0.1 | 1 | 0.158 | 7.3 | 5.4 | 4.1 | 7.6 | 4.2 | 0.4 | 0.58728 | 0.56055 | 1.34601 | 1.94534 | 0.82357 | 0.23743 | 0.34603 |
| 104 | Potentilla argentea L. | Potentilla argentea agg. | 0.1 | 1 | 0.150 | 7.1 | NA | NA | NA | 4.6 | 0 | 0.69216 | 0.66495 | 1.32487 | 1.98453 | 0.49303 | 0.16764 | 0.56484 |
| 104 | Potentilla argentea L. | Potentilla argentea agg. | 0.05 | 1 | 0.167 | 7.3 | 5.4 | 4.1 | 7.6 | 4.2 | 0.4 | 0.58728 | 0.56055 | 1.34601 | 1.94534 | 0.82357 | 0.23743 | 0.34603 |
| 104 | Plantago lanceolata L. | Plantago lanceolata | 0.05 | 1 | 0.165 | 7.3 | 5.4 | 4.1 | 7.6 | 4.2 | 0.4 | 0.58728 | 0.56055 | 1.34601 | 1.94534 | 0.82357 | 0.23743 | 0.34603 |
| 104 | Hypochaeris radicata L. | Hypochoeris radicata | 0.05 | 1 | 0.156 | 7.3 | 5.4 | 4.1 | 7.6 | 4.2 | 0.4 | 0.58728 | 0.56055 | 1.34601 | 1.94534 | 0.82357 | 0.23743 | 0.34603 |
| 104 | Poa annua L. | Poa annua | 0.5 | 1 | 0.303 | 7.3 | 5.4 | 4.1 | 7.6 | 4.2 | 0.4 | 0.58728 | 0.56055 | 1.34601 | 1.94534 | 0.82357 | 0.23743 | 0.34603 |
| 104 | Bromus sterilis L. | Bromus sterilis | 0.15 | 1 | 0.100 | 7.3 | 5.4 | 4.1 | 7.6 | 4.2 | 0.4 | 0.58728 | 0.56055 | 1.34601 | 1.94534 | 0.82357 | 0.23743 | 0.34603 |
| 105 | Medicago lupulina L. | Medicago lupulina | 0.08 | 1 | 0.081 | 7.3 | 5.4 | 4.1 | 7.6 | 4.2 | 0.4 | 0.58728 | 0.56055 | 1.34601 | 1.94534 | 0.82357 | 0.23743 | 0.34603 |
| 105 | Potentilla argentea L. | Potentilla argentea agg. | 0.08 | 1 | 0.100 | 7.3 | 5.4 | 4.1 | 7.6 | 4.2 | 0.4 | 0.58728 | 0.56055 | 1.34601 | 1.94534 | 0.82357 | 0.23743 | 0.34603 |
| 105 | Taraxacum officinale F.H.Wigg. | Taraxacum officinale agg. | 0.05 | 1 | 0.100 | 7.9 | NA | 4.1 | 5.8 | NA | 0.9 | 0.54139 | 0.51128 | 1.31363 | 1.93633 | 0.8112 | 0.25085 | 0.32953 |
| 105 | Veronica arvensis L. | Veronica arvensis | 0.1 | 1 | 0.100 | 7.1 | 5.3 | 4.5 | 5.9 | 4.8 | 0.3 | 0.52518 | 0.5113 | 1.61045 | 2.05509 | 0.96495 | 0.27897 | 0.32911 |
| 105 | Erodium cicutarium (L.) | Erodium cicutarium cicutarium | 0.08 | 1 | 0.084 | 7.9 | NA | 4.1 | 5.8 | NA | 0.9 | 0.54139 | 0.51128 | 1.31363 | 1.93633 | 0.8112 | 0.25085 | 0.32953 |
| 105 | Alliaria petiolata (M.Bieb.) | Alliaria petiolata | 0.02 | 1 | 0.082 | 7.9 | NA | 4.1 | 5.8 | NA | 0.9 | 0.54139 | 0.51128 | 1.31363 | 1.93633 | 0.8112 | 0.25085 | 0.32953 |
| 105 | Crepis capillaris (L.) Wallr. | Crepis capillaris | 0.05 | 1 | 0.100 | 7.9 | NA | 4.1 | 5.8 | NA | 0.9 | 0.54139 | 0.51128 | 1.31363 | 1.93633 | 0.8112 | 0.25085 | 0.32953 |
| 105 | Geum urbanum L. | Geum urbanum | 0.08 | 1 | 0.080 | 7.9 | NA | 4.1 | 5.8 | NA | 0.9 | 0.54139 | 0.51128 | 1.31363 | 1.93633 | 0.8112 | 0.25085 | 0.32953 |
| 105 | Poa annua L. | Poa annua | 0.3 | 1 | 0.050 | 7.9 | NA | 4.1 | 5.8 | NA | 0.9 | 0.54139 | 0.51128 | 1.31363 | 1.93633 | 0.8112 | 0.25085 | 0.32953 |
| 105 | Bromus sterilis L. | Bromus sterilis | 0.16 | 1 | 0.083 | 7.9 | NA | 4.1 | 5.8 | NA | 0.9 | 0.54139 | 0.51128 | 1.31363 | 1.93633 | 0.8112 | 0.25085 | 0.32953 |
| 106 | Medicago lupulina L. | Medicago lupulina | 0.1 | 1 | 0.080 | 7.9 | NA | 4.1 | 5.8 | NA | 0.9 | 0.54139 | 0.51128 | 1.31363 | 1.93633 | 0.8112 | 0.25085 | 0.32953 |
| 106 | Potentilla argentea L. | Potentilla argentea agg. | 0.05 | 1 | 0.100 | 7.9 | NA | 4.1 | 5.8 | NA | 0.9 | 0.54139 | 0.51128 | 1.31363 | 1.93633 | 0.8112 | 0.25085 | 0.32953 |
| 106 | Erodium cicutarium (L.) | Erodium cicutarium cicutarium | 0.09 | 1 | 0.063 | 7.9 | NA | 4.1 | 5.8 | NA | 0.9 | 0.54139 | 0.51128 | 1.31363 | 1.93633 | 0.8112 | 0.25085 | 0.32953 |
| 106 | Sedum acre L. | Sedum acre | 0.1 | 1 | 0.084 | 7.9 | NA | 4.1 | 5.8 | NA | 0.9 | 0.54139 | 0.51128 | 1.31363 | 1.93633 | 0.8112 | 0.25085 | 0.32953 |
| 106 | Taraxacum officinale F.H.Wigg. | Taraxacum officinale agg. | 0.05 | 1 | 0.082 | 7.9 | NA | 4.1 | 5.8 | NA | 0.9 | 0.54139 | 0.51128 | 1.31363 | 1.93633 | 0.8112 | 0.25085 | 0.32953 |
| 106 | Poa annua L. | Poa annua | 0.34 | 1 | 0.120 | 7.9 | NA | 4.1 | 5.8 | NA | 0.9 | 0.54139 | 0.51128 | 1.31363 | 1.93633 | 0.8112 | 0.25085 | 0.32953 |
| 106 | Bromus sterilis L. | Bromus sterilis | 0.27 | 1 | 0.080 | 7.9 | NA | 4.1 | 5.8 | NA | 0.9 | 0.54139 | 0.51128 | 1.31363 | 1.93633 | 0.8112 | 0.25085 | 0.32953 |
| 107 | Medicago lupulina L. | Medicago lupulina | 0.08 | 1 | 0.100 | 7.9 | NA | 4.1 | 5.8 | NA | 0.9 | 0.54139 | 0.51128 | 1.31363 | 1.93633 | 0.8112 | 0.25085 | 0.32953 |
| 107 | Potentilla argentea L. | Potentilla argentea agg. | 0.1 | 1 | 0.100 | 7.9 | NA | 4.1 | 5.8 | NA | 0.9 | 0.54139 | 0.51128 | 1.31363 | 1.93633 | 0.8112 | 0.25085 | 0.32953 |
| 107 | Taraxacum officinale F.H.Wigg. | Taraxacum officinale agg. | 0.05 | 1 | 0.050 | 7.9 | NA | 4.1 | 5.8 | NA | 0.9 | 0.54139 | 0.51128 | 1.31363 | 1.93633 | 0.8112 | 0.25085 | 0.32953 |
| 107 | Veronica arvensis L. | Veronica arvensis | 0.05 | 1 | 0.030 | 8.6 | NA | 2.7 | 4.4 | 1.6 | 0 | 0.53557 | 0.50485 | 1.29532 | 1.90211 | 0.72298 | 0.28495 | 0.2778 |

| | | | | | | | | | | | | | | | | | | |
|---|---|---|---|---|---|---|---|---|---|---|---|---|---|---|---|---|---|---|
| 107 | Rumex acetosella L. | Rumex acetosella acetosella | 0.1 | 1 | 0.020 | 7 | 5.6 | 4.4 | 5.9 | 5.4 | 0.1 | 0.64871 | 0.63945 | 1.70759 | 2.05577 | 0.56937 | 0.21649 | 0.54803 |
| 107 | Myosotis arvensis Hill | Myosotis arvensis (intermedia) | 0.05 | 1 | 0.080 | 8.6 | NA | 2.7 | 4.4 | 1.6 | 0 | 0.53557 | 0.50485 | 1.29532 | 1.90211 | 0.72298 | 0.28495 | 0.2778 |
| 107 | Sisymbrium officinale (L.) Scop. | Sisymbrium officinale | 0.05 | 1 | 0.429 | 7.4 | 5.3 | NA | 6 | 7.7 | 0.8 | 0.65201 | 0.63791 | 1.54898 | 2.03286 | 0.77966 | 0.37419 | 0.45721 |
| 107 | Poa annua L. | Poa annua | 0.3 | 1 | 0.250 | NA | NA | NA | NA | NA | NA | NA | NA | NA | NA | NA | NA | NA |
| 107 | Bromus sterilis L. | Bromus sterilis | 0.22 | 1 | 0.010 | 6.3 | NA | 5.9 | 6.3 | 6.9 | 0.1 | 0.6037 | 0.3946 | 0.44368 | 1.50139 | 0.30164 | 0.21315 | 0.21494 |
| 108 | Erodium cicutarium (L.) | Erodium cicutarium cicutarium | 0.1 | 1 | 0.010 | NA | 5.4 | 5.7 | 6.8 | 7 | 0 | 0.67742 | 0.24559 | 0.14546 | 1.36309 | 0.06547 | 0.19208 | 0.12644 |
| 108 | Potentilla argentea L. | Potentilla argentea agg. | 0.06 | 1 | 0.200 | 6.7 | NA | NA | 6.5 | 7.7 | 0.1 | 0.70945 | 0.61373 | 0.80583 | 1.7717 | 0.43958 | 0.26577 | 0.51423 |
| 108 | Echium vulgare L. | Echium vulgare | 0.02 | 1 | 0.460 | 5.2 | 5.9 | 5.4 | 7.3 | 8 | 0 | 0.67947 | 0.25214 | 0.1495 | 1.33661 | 0.07498 | 0.19025 | 0.12819 |
| 108 | Polygonum aviculare L. | Polygonum aviculare agg. | 0.02 | 1 | 0.202 | NA | 5.4 | 5.7 | 6.8 | 7 | 0 | 0.67742 | 0.24559 | 0.14546 | 1.36309 | 0.06547 | 0.19208 | 0.12644 |
| 108 | Poa annua L. | Poa annua | 0.3 | 1 | 0.010 | 7.2 | 5.5 | 4.5 | 6 | 7.6 | 0.3 | 0.72905 | 0.70188 | 1.35981 | 1.84158 | 0.57912 | 0.20081 | 0.60725 |
| 108 | Bromus sterilis L. | Bromus sterilis | 0.5 | 1 | 0.157 | 7.9 | NA | 4.1 | 5.8 | NA | 0.9 | 0.54139 | 0.51128 | 1.31363 | 1.93633 | 0.8112 | 0.25085 | 0.32953 |
| 109 | Erodium cicutarium (L.) | Erodium cicutarium cicutarium | 0.1 | 0.95 | 0.083 | NA | NA | NA | NA | NA | NA | NA | NA | NA | NA | NA | NA | NA |
| 109 | Taraxacum officinale F.H.Wigg. | Taraxacum officinale agg. | 0.1 | 0.95 | 0.204 | NA | NA | NA | NA | NA | NA | NA | NA | NA | NA | NA | NA | NA |
| 109 | Plantago major L. | Plantago major ssp. major | 0.05 | 0.95 | 0.435 | 7.4 | 5.3 | NA | 6 | 7.7 | 0.8 | 0.65201 | 0.63791 | 1.54898 | 2.03286 | 0.77966 | 0.37419 | 0.45721 |
| 109 | Myosotis arvensis Hill | Myosotis arvensis (intermedia) | 0.05 | 0.95 | 0.111 | 8.4 | NA | 3 | 7.7 | 3.3 | 0 | 0.62073 | 0.55773 | 1.05738 | 1.76031 | 0.36863 | 0.23711 | 0.33892 |
| 109 | Geum urbanum L. | Geum urbanum | 0.05 | 0.95 | 0.151 | 8 | 7.1 | 4.3 | 6.5 | 6.5 | 0 | 0.77 | 0.73636 | 1.30212 | 1.89005 | 0.65758 | 0.2 | 0.54 |
| 109 | Alliaria petiolata (M.Bieb.) | Alliaria petiolata | 0.05 | 0.95 | 0.050 | NA | NA | 4.5 | 6.6 | 6.6 | 0 | 0.78059 | 0.60963 | 0.54418 | 1.63701 | 0.26733 | 0.16232 | 0.54354 |
| 109 | Poa annua L. | Poa annua | 0.45 | 0.95 | 0.050 | NA | NA | 4.5 | 6.6 | 6.6 | 0 | 0.78059 | 0.60963 | 0.54418 | 1.63701 | 0.26733 | 0.16232 | 0.54354 |
| 109 | Bromus sterilis L. | Bromus sterilis | 0.1 | 0.95 | 0.111 | 7.9 | 4.8 | 6 | 7.4 | 6 | 0 | 0.63414 | 0.54614 | 0.83673 | 1.6192 | 0.40196 | 0.19592 | 0.32047 |
| 110 | Erodium cicutarium (L.) | Erodium cicutarium cicutarium | 0.1 | 1 | 0.240 | 7.9 | 5.7 | 4.2 | 7 | 7.2 | 0.1 | 0.76998 | 0.7507 | 1.46055 | 1.90983 | 0.91556 | 0.1857 | 0.55131 |
| 110 | Taraxacum officinale F.H.Wigg. | Taraxacum officinale agg. | 0.2 | 1 | 0.150 | 7.4 | 5.3 | NA | 6 | 7.7 | 0.8 | 0.65201 | 0.63791 | 1.54898 | 2.03286 | 0.77966 | 0.37419 | 0.45721 |
| 110 | Potentilla argentea L. | Potentilla argentea agg. | 0.15 | 1 | 0.333 | 7.4 | 5.3 | NA | 6 | 7.7 | 0.8 | 0.65201 | 0.63791 | 1.54898 | 2.03286 | 0.77966 | 0.37419 | 0.45721 |
| 110 | Myosotis arvensis Hill | Myosotis arvensis (intermedia) | 0.05 | 1 | 0.158 | NA | NA | NA | NA | NA | NA | NA | NA | NA | NA | NA | NA | NA |
| 110 | Polygonum aviculare L. | Polygonum aviculare agg. | 0.05 | 1 | 0.700 | 8.1 | NA | 3.9 | 6.9 | 6.4 | 0.4 | 0.80098 | 0.76713 | 1.29016 | 1.92909 | 0.75604 | 0.19458 | 0.52486 |
| 110 | Poa annua L. | Poa annua | 0.2 | 1 | 0.800 | 8.1 | NA | 3.9 | 6.9 | 6.4 | 0.4 | 0.80098 | 0.76713 | 1.29016 | 1.92909 | 0.75604 | 0.19458 | 0.52486 |
| 110 | Bromus sterilis L. | Bromus sterilis | 0.25 | 1 | 0.111 | 6.7 | NA | NA | 6.5 | 7.7 | 0.1 | 0.70945 | 0.61373 | 0.80583 | 1.7717 | 0.43958 | 0.26577 | 0.51423 |
| 111 | Erodium cicutarium (L.) | Erodium cicutarium cicutarium | 0.08 | 1 | 0.400 | 8.1 | NA | 3.9 | 6.9 | 6.4 | 0.4 | 0.80098 | 0.76713 | 1.29016 | 1.92909 | 0.75604 | 0.19458 | 0.52486 |
| 111 | Potentilla argentea L. | Potentilla argentea agg. | 0.1 | 1 | 0.300 | 8.1 | NA | 3.9 | 6.9 | 6.4 | 0.4 | 0.80098 | 0.76713 | 1.29016 | 1.92909 | 0.75604 | 0.19458 | 0.52486 |
| 111 | Alliaria petiolata (M.Bieb.) | Alliaria petiolata | 0.04 | 1 | 0.111 | 8.1 | NA | 3.9 | 6.9 | 6.4 | 0.4 | 0.80098 | 0.76713 | 1.29016 | 1.92909 | 0.75604 | 0.19458 | 0.52486 |
| 111 | Veronica arvensis L. | Veronica arvensis | 0.04 | 1 | 0.400 | 8.1 | NA | 3.9 | 6.9 | 6.4 | 0.4 | 0.80098 | 0.76713 | 1.29016 | 1.92909 | 0.75604 | 0.19458 | 0.52486 |
| 111 | Poa annua L. | Poa annua | 0.54 | 1 | 0.400 | 6.7 | NA | NA | 6.5 | 7.7 | 0.1 | 0.70945 | 0.61373 | 0.80583 | 1.7717 | 0.43958 | 0.26577 | 0.51423 |
| 111 | Bromus sterilis L. | Bromus sterilis | 0.2 | 1 | 0.526 | 7.4 | 5.3 | NA | 6 | 7.7 | 0.8 | 0.65201 | 0.63791 | 1.54898 | 2.03286 | 0.77966 | 0.37419 | 0.45721 |
| 112 | Medicago lupulina L. | Medicago lupulina | 0.08 | 0.5 | 0.050 | 7.2 | 5.5 | 4.5 | 6 | 7.6 | 0.3 | 0.72905 | 0.70188 | 1.35981 | 1.84158 | 0.57912 | 0.20081 | 0.60725 |
| 112 | Erodium cicutarium (L.) | Erodium cicutarium cicutarium | 0.05 | 0.5 | 0.150 | 8.1 | NA | NA | 4.4 | 2.8 | 0.3 | 0.48868 | 0.46508 | 1.4426 | 2.04965 | 0.7278 | 0.33931 | 0.35628 |
| 112 | Geranium molle L. | Geranium molle | 0.02 | 0.5 | 0.200 | 6.7 | NA | NA | 6.5 | 7.7 | 0.1 | 0.70945 | 0.61373 | 0.80583 | 1.7717 | 0.43958 | 0.26577 | 0.51423 |
| 112 | Achillea millefolium L. | Achillea millefolium millefolium | 0.05 | 0.5 | 0.200 | 7.4 | 5.3 | NA | 6 | 7.7 | 0.8 | 0.65201 | 0.63791 | 1.54898 | 2.03286 | 0.77966 | 0.37419 | 0.45721 |
| 112 | Bromus diandrus Roth | NA | 0.2 | 0.5 | 0.200 | 7 | NA | NA | 7.5 | 4.8 | 0 | 0.66959 | 0.18576 | 0.11569 | 1.33503 | 0.03304 | 0.20329 | 0.10324 |
| 112 | Bromus sterilis L. | Bromus sterilis | 0.1 | 0.5 | 0.510 | 7.4 | 5.3 | NA | 6 | 7.7 | 0.8 | 0.65201 | 0.63791 | 1.54898 | 2.03286 | 0.77966 | 0.37419 | 0.45721 |
| 113 | Medicago lupulina L. | Medicago lupulina | 0.1 | 0.45 | 0.421 | 7.4 | 5.3 | NA | 6 | 7.7 | 0.8 | 0.65201 | 0.63791 | 1.54898 | 2.03286 | 0.77966 | 0.37419 | 0.45721 |
| 113 | Erodium cicutarium (L.) | Erodium cicutarium cicutarium | 0.05 | 0.45 | 0.400 | 6.7 | NA | NA | 6.5 | 7.7 | 0.1 | 0.70945 | 0.61373 | 0.80583 | 1.7717 | 0.43958 | 0.26577 | 0.51423 |
| 113 | Achillea millefolium L. | Achillea millefolium millefolium | 0.04 | 0.45 | 0.615 | 6.7 | NA | NA | 6.5 | 7.7 | 0.1 | 0.70945 | 0.61373 | 0.80583 | 1.7717 | 0.43958 | 0.26577 | 0.51423 |
| 113 | Geranium molle L. | Geranium molle | 0.02 | 0.45 | 0.150 | NA | NA | NA | NA | NA | NA | NA | NA | NA | NA | NA | NA | NA |
| 113 | Bromus diandrus Roth | NA | 0.2 | 0.45 | 0.700 | 7.4 | 5.3 | NA | 6 | 7.7 | 0.8 | 0.65201 | 0.63791 | 1.54898 | 2.03286 | 0.77966 | 0.37419 | 0.45721 |
| 113 | Cerastium semidecandrum L. | Cerastium semidecandrum | 0.02 | 0.45 | 0.300 | 7.4 | 5.2 | 4.7 | 6.1 | NA | 0 | 0.77406 | 0.76726 | 1.79113 | 2.07279 | 0.64778 | 0.2119 | 0.65925 |
| 113 | Veronica arvensis L. | Veronica arvensis | 0.02 | 0.45 | 0.105 | NA | NA | NA | NA | NA | NA | NA | NA | NA | NA | NA | NA | NA |
| 114 | Medicago lupulina L. | Medicago lupulina | 0.08 | 0.26 | 0.889 | NA | NA | NA | NA | NA | NA | NA | NA | NA | NA | NA | NA | NA |
| 114 | Erodium cicutarium (L.) | Erodium cicutarium cicutarium | 0.04 | 0.26 | 0.111 | NA | NA | NA | NA | NA | NA | NA | NA | NA | NA | NA | NA | NA |
| 114 | Achillea millefolium L. | Achillea millefolium millefolium | 0.08 | 0.26 | 0.053 | 6.7 | NA | NA | 6.5 | 7.7 | 0.1 | 0.70945 | 0.61373 | 0.80583 | 1.7717 | 0.43958 | 0.26577 | 0.51423 |
| 114 | Stellaria media (L.) Vill. | Stellaria media agg. | 0.02 | 0.26 | 0.588 | 7.4 | 5.3 | NA | 6 | 7.7 | 0.8 | 0.65201 | 0.63791 | 1.54898 | 2.03286 | 0.77966 | 0.37419 | 0.45721 |
| 114 | Bromus hordeaceus L. | Bromus hordeaceus (mollis) | 0.04 | 0.26 | 0.050 | 7.4 | 5.3 | NA | 6 | 7.7 | 0.8 | 0.65201 | 0.63791 | 1.54898 | 2.03286 | 0.77966 | 0.37419 | 0.45721 |
| 115 | Medicago lupulina L. | Medicago lupulina | 0.04 | 0.3 | 0.350 | NA | NA | NA | NA | NA | NA | NA | NA | NA | NA | NA | NA | NA |
| 115 | Erodium cicutarium (L.) | Erodium cicutarium cicutarium | 0.05 | 0.3 | 0.100 | 7.8 | 5.3 | 3.8 | 6.6 | 5 | 0.2 | 0.49579 | 0.46867 | 1.36306 | 1.85033 | 0.62791 | 0.29201 | 0.3217 |
| 115 | Achillea millefolium L. | Achillea millefolium millefolium | 0.1 | 0.3 | 0.400 | NA | NA | NA | NA | NA | NA | NA | NA | NA | NA | NA | NA | NA |
| 115 | Taraxacum officinale F.H.Wigg. | Taraxacum officinale agg. | 0.05 | 0.3 | 0.400 | 7.8 | NA | 5.3 | 6.3 | 7.3 | 0 | NA | NA | NA | NA | NA | NA | NA |
| 115 | Geranium molle L. | Geranium molle | 0.02 | 0.3 | 0.250 | NA | NA | 5.3 | 6.2 | NA | 0.1 | 0.68682 | 0.14816 | 0.05813 | 1.31254 | 0.01433 | 0.20126 | 0.10505 |
| 115 | Stellaria media (L.) Vill. | Stellaria media agg. | 0.04 | 0.3 | 0.500 | 7.4 | 5.3 | NA | 6 | 7.7 | 0.8 | 0.65201 | 0.63791 | 1.54898 | 2.03286 | 0.77966 | 0.37419 | 0.45721 |
| 116 | Medicago lupulina L. | Medicago lupulina | 0.1 | 0.35 | 0.400 | 7.4 | 5.3 | NA | 6 | 7.7 | 0.8 | 0.65201 | 0.63791 | 1.54898 | 2.03286 | 0.77966 | 0.37419 | 0.45721 |
| 116 | Erodium cicutarium (L.) | Erodium cicutarium cicutarium | 0.1 | 0.35 | 0.200 | 7.7 | NA | NA | 5.6 | NA | 0.7 | 0.53223 | 0.52325 | 1.73719 | 2.14421 | 0.94278 | 0.39374 | 0.35054 |
| 116 | Achillea millefolium L. | Achillea millefolium millefolium | 0.1 | 0.35 | 0.450 | 7.7 | NA | NA | 5.6 | NA | 0.7 | 0.53223 | 0.52325 | 1.73719 | 2.14421 | 0.94278 | 0.39374 | 0.35054 |
| 116 | Erigeron canadensis L. | Conyza canadensis | 0.05 | 0.35 | 0.050 | 7.7 | NA | NA | 5.6 | NA | 0.7 | 0.53223 | 0.52325 | 1.73719 | 2.14421 | 0.94278 | 0.39374 | 0.35054 |
| 117 | Medicago lupulina L. | Medicago lupulina | 0.08 | 0.45 | 0.050 | NA | NA | NA | NA | NA | NA | NA | NA | NA | NA | NA | NA | NA |
| 117 | Erodium cicutarium (L.) | Erodium cicutarium cicutarium | 0.08 | 0.45 | 0.056 | 7.4 | 5.3 | NA | 6 | 7.7 | 0.8 | 0.65201 | 0.63791 | 1.54898 | 2.03286 | 0.77966 | 0.37419 | 0.45721 |
| 117 | Achillea millefolium L. | Achillea millefolium millefolium | 0.12 | 0.45 | 0.250 | 6.7 | NA | NA | 6.5 | 7.7 | 0.1 | 0.70945 | 0.61373 | 0.80583 | 1.7717 | 0.43958 | 0.26577 | 0.51423 |
| 117 | Stellaria media (L.) Vill. | Stellaria media agg. | 0.02 | 0.45 | 0.050 | NA | NA | 4.5 | 6.6 | 6.6 | 0 | 0.78059 | 0.60963 | 0.54418 | 1.63701 | 0.26733 | 0.16232 | 0.54354 |
| 117 | Cerastium semidecandrum L. | Cerastium semidecandrum | 0.05 | 0.45 | 0.100 | NA | NA | NA | NA | NA | NA | NA | NA | NA | NA | NA | NA | NA |
| 117 | Geranium molle L. | Geranium molle | 0.1 | 0.45 | 0.070 | 7.3 | 4.3 | NA | 5.9 | NA | 0.1 | 0.51602 | 0.49187 | 1.37487 | 2.0182 | 0.89491 | 0.26179 | 0.32262 |
| 118 | Medicago lupulina L. | Medicago lupulina | 0.05 | 0.5 | 0.125 | 7.3 | 5.4 | 4.1 | 7.6 | 4.2 | 0.4 | 0.58728 | 0.56055 | 1.34601 | 1.94534 | 0.82357 | 0.23743 | 0.34603 |
| 118 | Erodium cicutarium (L.) | Erodium cicutarium cicutarium | 0.04 | 0.5 | 0.050 | NA | NA | 4.5 | 6.6 | 6.6 | 0 | 0.78059 | 0.60963 | 0.54418 | 1.63701 | 0.26733 | 0.16232 | 0.54354 |
| 118 | Achillea millefolium L. | Achillea millefolium millefolium | 0.15 | 0.5 | 0.050 | 6.7 | NA | NA | 6.5 | 7.7 | 0.1 | 0.70945 | 0.61373 | 0.80583 | 1.7717 | 0.43958 | 0.26577 | 0.51423 |
| 118 | Taraxacum officinale F.H.Wigg. | Taraxacum officinale agg. | 0.1 | 0.5 | 0.150 | 7.9 | 5.7 | 4.2 | 7 | 7.2 | 0.1 | 0.76998 | 0.7507 | 1.46055 | 1.90983 | 0.91556 | 0.1857 | 0.55131 |
| 118 | Cerastium semidecandrum L. | Cerastium semidecandrum | 0.05 | 0.5 | 0.027 | NA | NA | 4.5 | 6.6 | 6.6 | 0 | 0.78059 | 0.60963 | 0.54418 | 1.63701 | 0.26733 | 0.16232 | 0.54354 |
| 118 | Senecio inaequidens DC. | Senecio inaequidens | 0.11 | 0.5 | 0.150 | 7.9 | 5.7 | 4.2 | 7 | 7.2 | 0.1 | 0.76998 | 0.7507 | 1.46055 | 1.90983 | 0.91556 | 0.1857 | 0.55131 |
| 119 | Medicago lupulina L. | Medicago lupulina | 0.05 | 0.6 | 0.100 | 6.7 | NA | NA | 6.5 | 7.7 | 0.1 | 0.70945 | 0.61373 | 0.80583 | 1.7717 | 0.43958 | 0.26577 | 0.51423 |
| 119 | Echium vulgare L. | Echium vulgare | 0.05 | 0.6 | 0.338 | 6.7 | NA | NA | 6.5 | 7.7 | 0.1 | 0.70945 | 0.61373 | 0.80583 | 1.7717 | 0.43958 | 0.26577 | 0.51423 |
| 119 | Achillea millefolium L. | Achillea millefolium millefolium | 0.3 | 0.6 | 0.190 | NA | NA | 4.5 | 6.6 | 6.6 | 0 | 0.78059 | 0.60963 | 0.54418 | 1.63701 | 0.26733 | 0.16232 | 0.54354 |
| 119 | Poa annua L. | Poa annua | 0.2 | 0.6 | 0.232 | NA | NA | 4.5 | 6.6 | 6.6 | 0 | 0.78059 | 0.60963 | 0.54418 | 1.63701 | 0.26733 | 0.16232 | 0.54354 |
| 120 | Medicago lupulina L. | Medicago lupulina | 0.1 | 0.58 | 0.100 | NA | NA | 4.5 | 6.6 | 6.6 | 0 | 0.78059 | 0.60963 | 0.54418 | 1.63701 | 0.26733 | 0.16232 | 0.54354 |
| 120 | Achillea millefolium L. | Achillea millefolium millefolium | 0.2 | 0.58 | 0.050 | NA | NA | 4.5 | 6.6 | 6.6 | 0 | 0.78059 | 0.60963 | 0.54418 | 1.63701 | 0.26733 | 0.16232 | 0.54354 |
| 120 | Cerastium semidecandrum L. | Cerastium semidecandrum | 0.05 | 0.58 | 0.050 | 6.3 | 5 | 7.3 | 5.8 | 6.5 | 0.7 | 0.53692 | 0.48641 | 1.02734 | 1.80642 | 0.6097 | 0.33182 | 0.26598 |
| 120 | Poa annua L. | Poa annua | 0.23 | 0.58 | 0.400 | 6.3 | NA | 5.9 | 6.3 | 6.9 | 0.1 | 0.6037 | 0.3946 | 0.44368 | 1.50139 | 0.30164 | 0.21315 | 0.21494 |
| 121 | Medicago lupulina L. | Medicago lupulina | 0.11 | 0.58 | 0.400 | 6.3 | NA | 5.9 | 6.3 | 6.9 | 0.1 | 0.6037 | 0.3946 | 0.44368 | 1.50139 | 0.30164 | 0.21315 | 0.21494 |
| 121 | Achillea millefolium L. | Achillea millefolium millefolium | 0.22 | 0.58 | 0.333 | 6.7 | NA | NA | 6.5 | 7.7 | 0.1 | 0.70945 | 0.61373 | 0.80583 | 1.7717 | 0.43958 | 0.26577 | 0.51423 |
| 121 | Poa annua L. | Poa annua | 0.25 | 0.58 | 0.050 | 7.7 | NA | NA | 5.6 | NA | 0.7 | 0.53223 | 0.52325 | 1.73719 | 2.14421 | 0.94278 | 0.39374 | 0.35054 |
| 122 | Medicago lupulina L. | Medicago lupulina | 0.04 | 0.4 | 0.052 | 7.4 | 5.3 | NA | 6 | 7.7 | 0.8 | 0.65201 | 0.63791 | 1.54898 | 2.03286 | 0.77966 | 0.37419 | 0.45721 |
| 122 | Erodium cicutarium (L.) | Erodium cicutarium cicutarium | 0.1 | 0.4 | 0.306 | NA | NA | NA | 7 | NA | 0 | 0.66957 | 0.20824 | 0.12967 | 1.34554 | 0.07292 | 0.20428 | 0.13248 |
| 122 | Achillea millefolium L. | Achillea millefolium millefolium | 0.05 | 0.4 | 0.250 | 6.9 | 5.9 | NA | 6.8 | 7 | 0 | 0.84157 | 0.83772 | 1.85474 | 2.04042 | 0.4856 | 0.14178 | 0.80173 |
| 122 | Stellaria media (L.) Vill. | Stellaria media agg. | 0.02 | 0.4 | 0.170 | NA | NA | NA | 7 | NA | 0 | 0.66957 | 0.20824 | 0.12967 | 1.34554 | 0.07292 | 0.20428 | 0.13248 |
| 122 | Cerastium semidecandrum L. | Cerastium semidecandrum | 0.04 | 0.4 | 0.300 | 6.3 | NA | 5.9 | 6.3 | 6.9 | 0.1 | 0.6037 | 0.3946 | 0.44368 | 1.50139 | 0.30164 | 0.21315 | 0.21494 |
| 122 | Senecio inaequidens DC. | Senecio inaequidens | 0.15 | 0.4 | 0.050 | 6.3 | 5 | 7.3 | 5.8 | 6.5 | 0.7 | 0.53692 | 0.48641 | 1.02734 | 1.80642 | 0.6097 | 0.33182 | 0.26598 |
| 123 | Bromus sterilis L. | Bromus sterilis | 0.15 | 1 | 0.110 | NA | NA | 4.5 | 6.6 | 6.6 | 0 | 0.78059 | 0.60963 | 0.54418 | 1.63701 | 0.26733 | 0.16232 | 0.54354 |
| 123 | Echium vulgare L. | Echium vulgare | 0.1 | 1 | 0.130 | 6.3 | 5 | 7.3 | 5.8 | 6.5 | 0.7 | 0.53692 | 0.48641 | 1.02734 | 1.80642 | 0.6097 | 0.33182 | 0.26598 |
| 123 | Polygonum aviculare L. | Polygonum aviculare agg. | 0.65 | 1 | 0.612 | 7.7 | NA | NA | 6.7 | 7.2 | 0.1 | 0.5681 | 0.51951 | 1.12401 | 1.73868 | 0.64408 | 0.31575 | 0.32626 |
| 123 | Taraxacum officinale F.H.Wigg. | Taraxacum officinale agg. | 0.1 | 1 | 0.333 | 7.4 | 5.3 | NA | 6 | 7.7 | 0.8 | 0.65201 | 0.63791 | 1.54898 | 2.03286 | 0.77966 | 0.37419 | 0.45721 |
| 124 | Achillea millefolium L. | Achillea millefolium millefolium | 0.15 | 1 | 0.100 | 6.5 | NA | NA | 5.4 | NA | 0 | 0.59769 | 0.45571 | 0.64594 | 1.51584 | 0.28825 | 0.22901 | 0.32316 |
| 124 | Geranium molle L. | Geranium molle | 0.05 | 1 | 0.350 | 7.4 | 5.3 | NA | 6 | 7.7 | 0.8 | 0.65201 | 0.63791 | 1.54898 | 2.03286 | 0.77966 | 0.37419 | 0.45721 |
| 124 | Taraxacum officinale F.H.Wigg. | Taraxacum officinale agg. | 0.05 | 1 | 0.316 | 7.4 | 5.3 | NA | 6 | 7.7 | 0.8 | 0.65201 | 0.63791 | 1.54898 | 2.03286 | 0.77966 | 0.37419 | 0.45721 |
| 124 | Bromus sterilis L. | Bromus sterilis | 0.75 | 1 | 0.471 | 8.4 | NA | 3 | 6 | NA | 0 | 0.46249 | 0.43915 | 1.39189 | 1.83687 | 0.38437 | 0.30571 | 0.38883 |
| 125 | Achillea millefolium L. | Achillea millefolium millefolium | 0.25 | 1.1 | 0.100 | 7 | 5.6 | 4.4 | 5.9 | 5.4 | 0.1 | 0.64871 | 0.63945 | 1.70759 | 2.05577 | 0.56937 | 0.21649 | 0.54803 |
| 125 | Geranium molle L. | Geranium molle | 0.25 | 1.1 | 0.100 | 7 | 5.6 | 4.4 | 5.9 | 5.4 | 0.1 | 0.64871 | 0.63945 | 1.70759 | 2.05577 | 0.56937 | 0.21649 | 0.54803 |
| 125 | Bromus sterilis L. | Bromus sterilis | 0.15 | 1.1 | 0.100 | NA | NA | NA | NA | NA | NA | NA | NA | NA | NA | NA | NA | NA |
| 125 | Poa annua L. | Poa annua | 0.45 | 1.1 | 0.100 | 7 | 5.6 | 4.4 | 5.9 | 5.4 | 0.1 | 0.64871 | 0.63945 | 1.70759 | 2.05577 | 0.56937 | 0.21649 | 0.54803 |
| 126 | Achillea millefolium L. | Achillea millefolium millefolium | 0.25 | 0.95 | 0.091 | 8.1 | NA | NA | 5.2 | NA | 0.1 | 0.51276 | 0.49581 | 1.51667 | 2.20989 | 0.98959 | 0.46997 | 0.3594 |
| 126 | Bromus sterilis L. | Bromus sterilis | 0.15 | 0.95 | 0.136 | 8.4 | NA | 3 | 6 | NA | 0 | 0.46249 | 0.43915 | 1.39189 | 1.83687 | 0.38437 | 0.30571 | 0.38883 |
| 126 | Poa annua L. | Poa annua | 0.5 | 0.95 | 0.200 | 6.8 | NA | 5.6 | 5.8 | NA | 0.1 | 0.52169 | 0.45097 | 0.93044 | 1.87015 | 0.67593 | 0.2975 | 0.27054 |
| 126 | Senecio inaequidens DC. | Senecio inaequidens | 0.05 | 0.95 | 0.100 | 7.3 | 5.4 | 4.1 | 7.6 | 4.2 | 0.4 | 0.58728 | 0.56055 | 1.34601 | 1.94534 | 0.82357 | 0.23743 | 0.34603 |
| 127 | Achillea millefolium L. | Achillea millefolium millefolium | 0.25 | 1 | 0.059 | 6.5 | NA | NA | 5.4 | NA | 0 | 0.59769 | 0.45571 | 0.64594 | 1.51584 | 0.28825 | 0.22901 | 0.32316 |

| | | | | | | | | | | | | | | | | | | |
|---|---|---|---|---|---|---|---|---|---|---|---|---|---|---|---|---|---|---|
| 127 | Taraxacum officinale F.H.Wigg. | Taraxacum officinale agg. | 0.08 | 1 | 0.300 | NA | NA | 4.5 | 6.6 | 6.6 | 0 | 0.78059 | 0.60963 | 0.54418 | 1.63701 | 0.26733 | 0.16232 | 0.54354 |
| 127 | Bromus sterilis L. | Bromus sterilis | 0.15 | 1 | 0.050 | 7 | 5.6 | 4.4 | 5.9 | 5.4 | 0.1 | 0.64871 | 0.63945 | 1.70759 | 2.05577 | 0.56937 | 0.21649 | 0.54803 |
| 127 | Poa annua L. | Poa annua | 0.38 | 1 | 0.200 | NA | NA | NA | NA | NA | NA | NA | NA | NA | NA | NA | NA | NA |
| 127 | Geranium molle L. | Geranium molle | 0.1 | 1 | 0.222 | NA | NA | NA | NA | NA | NA | NA | NA | NA | NA | NA | NA | NA |
| 127 | Senecio inaequidens DC. | Senecio inaequidens | 0.04 | 1 | 0.100 | NA | NA | NA | NA | NA | NA | NA | NA | NA | NA | NA | NA | NA |
| 128 | Achillea millefolium L. | Achillea millefolium millefolium | 0.2 | 1 | 0.050 | 7 | 5.6 | 4.4 | 5.9 | 5.4 | 0.1 | 0.64871 | 0.63945 | 1.70759 | 2.05577 | 0.56937 | 0.21649 | 0.54803 |
| 128 | Potentilla argentea L. | Potentilla argentea agg. | 0.25 | 1 | 0.050 | 7 | 5.6 | 4.4 | 5.9 | 5.4 | 0.1 | 0.64871 | 0.63945 | 1.70759 | 2.05577 | 0.56937 | 0.21649 | 0.54803 |
| 128 | Medicago lupulina L. | Medicago lupulina | 0.05 | 1 | 0.450 | 7.4 | 5.3 | NA | 6 | 7.7 | 0.8 | 0.65201 | 0.63791 | 1.54898 | 2.03286 | 0.77966 | 0.37419 | 0.45721 |
| 128 | Taraxacum officinale F.H.Wigg. | Taraxacum officinale agg. | 0.05 | 1 | 0.500 | 7.4 | 5.3 | NA | 6 | 7.7 | 0.8 | 0.65201 | 0.63791 | 1.54898 | 2.03286 | 0.77966 | 0.37419 | 0.45721 |
| 128 | Poa annua L. | Poa annua | 0.45 | 1 | 0.500 | 7.4 | 5.3 | NA | 6 | 7.7 | 0.8 | 0.65201 | 0.63791 | 1.54898 | 2.03286 | 0.77966 | 0.37419 | 0.45721 |
| 129 | Achillea millefolium L. | Achillea millefolium millefolium | 0.4 | 1 | 0.700 | 7.4 | 5.3 | NA | 6 | 7.7 | 0.8 | 0.65201 | 0.63791 | 1.54898 | 2.03286 | 0.77966 | 0.37419 | 0.45721 |
| 129 | Taraxacum officinale F.H.Wigg. | Taraxacum officinale agg. | 0.05 | 1 | 0.300 | 7.6 | NA | NA | 6.5 | 6.3 | 0.7 | 0.75548 | 0.75048 | 1.8409 | 2.08419 | 0.63746 | 0.23177 | 0.58814 |
| 129 | Geranium molle L. | Geranium molle | 0.1 | 1 | 0.421 | NA | NA | NA | NA | NA | NA | NA | NA | NA | NA | NA | NA | NA |
| 129 | Poa annua L. | Poa annua | 0.43 | 1 | 0.278 | NA | NA | NA | NA | NA | NA | NA | NA | NA | NA | NA | NA | NA |
| 129 | Medicago lupulina L. | Medicago lupulina | 0.02 | 1 | 0.278 | NA | NA | NA | NA | NA | NA | NA | NA | NA | NA | NA | NA | NA |
| 130 | Achillea millefolium L. | Achillea millefolium millefolium | 0.25 | 0.98 | 0.235 | 7.3 | 5.4 | 4.1 | 7.6 | 4.2 | 0.4 | 0.58728 | 0.56055 | 1.34601 | 1.94534 | 0.82357 | 0.23743 | 0.34603 |
| 130 | Taraxacum officinale F.H.Wigg. | Taraxacum officinale agg. | 0.1 | 0.98 | 0.222 | 7.3 | 5.4 | 4.1 | 7.6 | 4.2 | 0.4 | 0.58728 | 0.56055 | 1.34601 | 1.94534 | 0.82357 | 0.23743 | 0.34603 |
| 130 | Geranium molle L. | Geranium molle | 0.05 | 0.98 | 0.050 | 8.1 | NA | NA | 5.2 | NA | 0.1 | 0.51276 | 0.49581 | 1.51667 | 2.20989 | 0.98959 | 0.46997 | 0.3594 |
| 130 | Poa annua L. | Poa annua | 0.4 | 0.98 | 0.350 | 8 | NA | NA | 6.1 | 6.3 | 0.1 | 0.56333 | 0.55394 | 1.72195 | 2.16326 | 1.07172 | 0.43309 | 0.38415 |
| 130 | Polygonum aviculare L. | Polygonum aviculare agg. | 0.05 | 0.98 | 0.700 | 8.1 | NA | 3.9 | 6.9 | 6.4 | 0.4 | 0.80098 | 0.76713 | 1.29016 | 1.92909 | 0.75604 | 0.19458 | 0.52486 |
| 130 | Medicago lupulina L. | Medicago lupulina | 0.13 | 0.98 | 0.421 | 6.6 | 5.6 | 5 | 5.8 | 8.3 | 0 | 0.6044 | 0.52396 | 0.8458 | 1.44474 | 0.64843 | 0.16995 | 0.28076 |
| 131 | Achillea millefolium L. | Achillea millefolium millefolium | 0.25 | 1 | 0.030 | NA | NA | NA | NA | NA | NA | NA | NA | NA | NA | NA | NA | NA |
| 131 | Taraxacum officinale F.H.Wigg. | Taraxacum officinale agg. | 0.15 | 1 | 0.153 | 7.4 | 5.3 | NA | 6 | 7.7 | 0.8 | 0.65201 | 0.63791 | 1.54898 | 2.03286 | 0.77966 | 0.37419 | 0.45721 |
| 131 | Geranium molle L. | Geranium molle | 0.25 | 1 | 0.100 | 8.1 | NA | NA | 5.2 | NA | 0.1 | 0.51276 | 0.49581 | 1.51667 | 2.20989 | 0.98959 | 0.46997 | 0.3594 |
| 131 | Poa annua L. | Poa annua | 0.2 | 1 | 0.064 | 7.7 | NA | NA | 5.6 | NA | 0.7 | 0.53223 | 0.52325 | 1.73719 | 2.14421 | 0.94278 | 0.39374 | 0.35054 |
| 131 | Polygonum aviculare L. | Polygonum aviculare agg. | 0.15 | 1 | 0.074 | NA | NA | 5.3 | 6.2 | NA | 0.1 | 0.68682 | 0.14816 | 0.05813 | 1.31254 | 0.01433 | 0.20126 | 0.10505 |
| 132 | Achillea millefolium L. | Achillea millefolium millefolium | 0.05 | 0.98 | 0.143 | 7.3 | 6.1 | 3.8 | 6 | 5.1 | 0 | 0.62987 | 0.59281 | 1.27995 | 1.90491 | 0.6275 | 0.3036 | 0.42608 |
| 132 | Taraxacum officinale F.H.Wigg. | Taraxacum officinale agg. | 0.45 | 0.98 | 0.140 | 7 | 5.6 | 4.4 | 5.9 | 5.4 | 0.1 | 0.64871 | 0.63945 | 1.70759 | 2.05577 | 0.56937 | 0.21649 | 0.54803 |
| 132 | Potentilla argentea L. | Potentilla argentea agg. | 0.1 | 0.98 | 0.133 | NA | NA | NA | NA | NA | NA | NA | NA | NA | NA | NA | NA | NA |
| 132 | Poa annua L. | Poa annua | 0.3 | 0.98 | 0.020 | 6.5 | NA | NA | 5.4 | NA | 0 | 0.59769 | 0.45571 | 0.64594 | 1.51584 | 0.28825 | 0.22901 | 0.32316 |
| 132 | Bromus sterilis L. | Bromus sterilis | 0.05 | 0.98 | 0.032 | NA | NA | NA | NA | NA | NA | NA | NA | NA | NA | NA | NA | NA |
| 132 | Geranium molle L. | Geranium molle | 0.03 | 0.98 | 0.150 | 7.7 | NA | NA | 6.7 | 7.2 | 0.1 | 0.5681 | 0.51951 | 1.12401 | 1.73868 | 0.64408 | 0.31575 | 0.32626 |
| 133 | Achillea millefolium L. | Achillea millefolium millefolium | 0.1 | 1 | 0.900 | 7.4 | 5.3 | NA | 6 | 7.7 | 0.8 | 0.65201 | 0.63791 | 1.54898 | 2.03286 | 0.77966 | 0.37419 | 0.45721 |
| 133 | Taraxacum officinale F.H.Wigg. | Taraxacum officinale agg. | 0.1 | 1 | 0.200 | 6.6 | 5.7 | NA | 6.3 | 7.6 | 0.1 | 0.67317 | 0.45023 | 0.43732 | 1.4267 | 0.22701 | 0.17304 | 0.27906 |
| 133 | Plantago lanceolata L. | Plantago lanceolata | 0.25 | 1 | 0.421 | 6.6 | 5.7 | NA | 6.3 | 7.6 | 0.1 | 0.67317 | 0.45023 | 0.43732 | 1.4267 | 0.22701 | 0.17304 | 0.27906 |
| 133 | Poa annua L. | Poa annua | 0.38 | 1 | 0.800 | 8.1 | NA | 3.9 | 6.9 | 6.4 | 0.4 | 0.80098 | 0.76713 | 1.29016 | 1.92909 | 0.75604 | 0.19458 | 0.52486 |
| 133 | Bromus sterilis L. | Bromus sterilis | 0.15 | 1 | 0.222 | 7.1 | 4.9 | NA | 6 | NA | 1 | 0.59517 | 0.59064 | 1.89772 | 2.12267 | 0.65091 | 0.41213 | 0.37185 |
| 133 | Medicago lupulina L. | Medicago lupulina | 0.02 | 1 | 0.330 | 8.1 | NA | NA | 4.4 | 2.8 | 0.3 | 0.48868 | 0.46508 | 1.4426 | 2.04965 | 0.7278 | 0.33931 | 0.35628 |
| 134 | Achillea millefolium L. | Achillea millefolium millefolium | 0.25 | 1 | 0.278 | 7.1 | 4.9 | NA | 6 | NA | 1 | 0.59517 | 0.59064 | 1.89772 | 2.12267 | 0.65091 | 0.41213 | 0.37185 |
| 134 | Taraxacum officinale F.H.Wigg. | Taraxacum officinale agg. | 0.15 | 1 | 0.100 | 6.7 | NA | NA | 6.5 | 7.7 | 0.1 | 0.70945 | 0.61373 | 0.80583 | 1.7717 | 0.43958 | 0.26577 | 0.51423 |
| 134 | Medicago lupulina L. | Medicago lupulina | 0.35 | 1 | 0.050 | 7.1 | 4.9 | NA | 6 | NA | 1 | 0.59517 | 0.59064 | 1.89772 | 2.12267 | 0.65091 | 0.41213 | 0.37185 |
| 134 | Poa annua L. | Poa annua | 0.15 | 1 | 0.740 | NA | 6 | NA | NA | NA | NA | NA | NA | NA | NA | NA | NA | NA |
| 134 | Bromus sterilis L. | Bromus sterilis | 0.1 | 1 | 0.153 | NA | 6 | NA | NA | NA | NA | NA | NA | NA | NA | NA | NA | NA |
| 135 | Achillea millefolium L. | Achillea millefolium millefolium | 0.11 | 1 | 0.426 | NA | 6 | NA | NA | NA | NA | NA | NA | NA | NA | NA | NA | NA |
| 135 | Taraxacum officinale F.H.Wigg. | Taraxacum officinale agg. | 0.05 | 1 | 0.300 | NA | NA | NA | NA | NA | NA | NA | NA | NA | NA | NA | NA | NA |
| 135 | Medicago lupulina L. | Medicago lupulina | 0.3 | 1 | 0.220 | NA | NA | NA | NA | NA | NA | NA | NA | NA | NA | NA | NA | NA |
| 135 | Poa annua L. | Poa annua | 0.35 | 1 | 0.300 | 7.7 | NA | NA | 5.6 | NA | 0.7 | 0.53223 | 0.52325 | 1.73719 | 2.14421 | 0.94278 | 0.39374 | 0.35054 |
| 135 | Bromus sterilis L. | Bromus sterilis | 0.1 | 1 | 0.193 | 8.1 | NA | 3.5 | 7.5 | 5 | 0.3 | 0.75184 | 0.72654 | 1.3537 | 1.85024 | 0.717 | 0.18024 | 0.48387 |
| 135 | Bellis perennis L. | Bellis perennis | 0.02 | 1 | 0.200 | 7.4 | 5.3 | NA | 6 | 7.7 | 0.8 | 0.65201 | 0.63791 | 1.54898 | 2.03286 | 0.77966 | 0.37419 | 0.45721 |
| 135 | Stellaria media (L.) Vill. | Stellaria media agg. | 0.02 | 1 | 0.105 | 7 | NA | NA | NA | NA | 0.3 | 0.56899 | 0.4288 | 0.64817 | 1.63031 | 0.49671 | 0.22558 | 0.25973 |
| 135 | Plantago lanceolata L. | Plantago lanceolata | 0.05 | 1 | 0.700 | 7.4 | 5.3 | NA | 6 | 7.7 | 0.8 | 0.65201 | 0.63791 | 1.54898 | 2.03286 | 0.77966 | 0.37419 | 0.45721 |
| 136 | Achillea millefolium L. | Achillea millefolium millefolium | 0.1 | 1 | 0.100 | 7.1 | 5.3 | 4.5 | 5.9 | 4.8 | 0.3 | 0.52518 | 0.5113 | 1.61045 | 2.05509 | 0.96495 | 0.27897 | 0.32911 |
| 136 | Taraxacum officinale F.H.Wigg. | Taraxacum officinale agg. | 0.2 | 1 | 0.040 | 7 | 5.6 | 4.4 | 5.9 | 5.4 | 0.1 | 0.64871 | 0.63945 | 1.70759 | 2.05577 | 0.56937 | 0.21649 | 0.54803 |
| 136 | Medicago lupulina L. | Medicago lupulina | 0.05 | 1 | 0.300 | 7.1 | 5.3 | 4.5 | 5.9 | 4.8 | 0.3 | 0.52518 | 0.5113 | 1.61045 | 2.05509 | 0.96495 | 0.27897 | 0.32911 |
| 136 | Poa annua L. | Poa annua | 0.45 | 1 | 0.210 | 7.1 | 5.3 | 4.5 | 5.9 | 4.8 | 0.3 | 0.52518 | 0.5113 | 1.61045 | 2.05509 | 0.96495 | 0.27897 | 0.32911 |
| 136 | Hypochaeris radicata L. | Hypochoeris radicata | 0.2 | 1 | 0.186 | 7.1 | 5.3 | 4.5 | 5.9 | 4.8 | 0.3 | 0.52518 | 0.5113 | 1.61045 | 2.05509 | 0.96495 | 0.27897 | 0.32911 |
| 137 | Achillea millefolium L. | Achillea millefolium millefolium | 0.05 | 1 | 0.350 | 8 | NA | NA | 6.1 | 6.3 | 0.1 | 0.56333 | 0.55394 | 1.72195 | 2.16326 | 1.07172 | 0.43309 | 0.38415 |
| 137 | Taraxacum officinale F.H.Wigg. | Taraxacum officinale agg. | 0.25 | 1 | 0.250 | 8 | NA | NA | 6.1 | 6.3 | 0.1 | 0.56333 | 0.55394 | 1.72195 | 2.16326 | 1.07172 | 0.43309 | 0.38415 |
| 137 | Medicago lupulina L. | Medicago lupulina | 0.05 | 1 | 0.200 | 7.2 | NA | NA | 6.7 | 7.8 | 0.1 | 0.79972 | 0.7689 | 1.29856 | 1.89634 | 0.65193 | 0.17352 | 0.67176 |
| 137 | Poa annua L. | Poa annua | 0.2 | 1 | 0.330 | 6.9 | 6.5 | NA | 7.3 | 6.3 | 0 | 0.67274 | 0.19896 | 0.13039 | 1.32576 | 0.04466 | 0.20186 | 0.11616 |
| 137 | Geranium molle L. | Geranium molle | 0.05 | 1 | 0.667 | 7.1 | 7.8 | 4.1 | 6.2 | 6 | 0.1 | 0.76637 | 0.65623 | 0.78647 | 1.82229 | 0.56568 | 0.18972 | 0.47454 |
| 137 | Polygonum aviculare L. | Polygonum aviculare agg. | 0.05 | 1 | 0.333 | 7.8 | NA | 4.6 | 6.2 | 5.8 | 0.1 | 0.71321 | 0.68276 | 1.29546 | 1.8275 | 0.62188 | 0.18605 | 0.48742 |
| 137 | Plantago lanceolata L. | Plantago lanceolata | 0.35 | 1 | 0.089 | 7.3 | 5.4 | 4.1 | 7.6 | 4.2 | 0.4 | 0.58728 | 0.56055 | 1.34601 | 1.94534 | 0.82357 | 0.23743 | 0.34603 |
| 138 | Achillea millefolium L. | Achillea millefolium millefolium | 0.23 | 0.9 | 0.500 | NA | NA | NA | NA | NA | NA | NA | NA | NA | NA | NA | NA | NA |
| 138 | Plantago lanceolata L. | Plantago lanceolata | 0.13 | 0.9 | 0.040 | 7 | 5.6 | 4.4 | 5.9 | 5.4 | 0.1 | 0.64871 | 0.63945 | 1.70759 | 2.05577 | 0.56937 | 0.21649 | 0.54803 |
| 138 | Erodium cicutarium (L.) | Erodium cicutarium cicutarium | 0.08 | 0.9 | 0.403 | NA | NA | NA | NA | NA | NA | NA | NA | NA | NA | NA | NA | NA |
| 138 | Hypochaeris radicata L. | Hypochoeris radicata | 0.18 | 0.9 | 0.455 | NA | NA | NA | NA | NA | NA | NA | NA | NA | NA | NA | NA | NA |
| 138 | Potentilla argentea L. | Potentilla argentea agg. | 0.05 | 0.9 | 0.300 | NA | NA | NA | NA | NA | NA | NA | NA | NA | NA | NA | NA | NA |
| 138 | Poa annua L. | Poa annua | 0.18 | 0.9 | 0.444 | NA | NA | NA | NA | NA | NA | NA | NA | NA | NA | NA | NA | NA |
| 138 | Bromus sterilis L. | Bromus sterilis | 0.05 | 0.9 | 0.167 | 7.1 | 4.9 | NA | 6 | NA | 1 | 0.59517 | 0.59064 | 1.89772 | 2.12267 | 0.65091 | 0.41213 | 0.37185 |
| 139 | Achillea millefolium L. | Achillea millefolium millefolium | 0.2 | 1 | 0.667 | NA | NA | NA | NA | NA | NA | NA | NA | NA | NA | NA | NA | NA |
| 139 | Taraxacum officinale F.H.Wigg. | Taraxacum officinale agg. | 0.05 | 1 | 0.200 | 6.5 | NA | NA | 5.4 | NA | 0 | 0.59769 | 0.45571 | 0.64594 | 1.51584 | 0.28825 | 0.22901 | 0.32316 |
| 139 | Medicago lupulina L. | Medicago lupulina | 0.2 | 1 | 0.200 | 7 | 5.6 | 4.4 | 5.9 | 5.4 | 0.1 | 0.64871 | 0.63945 | 1.70759 | 2.05577 | 0.56937 | 0.21649 | 0.54803 |
| 139 | Poa annua L. | Poa annua | 0.55 | 1 | 0.500 | 7.1 | 4.9 | NA | 6 | NA | 1 | 0.59517 | 0.59064 | 1.89772 | 2.12267 | 0.65091 | 0.41213 | 0.37185 |
| 140 | Achillea millefolium L. | Achillea millefolium millefolium | 0.05 | 1 | 0.200 | 7 | 5.6 | 4.4 | 5.9 | 5.4 | 0.1 | 0.64871 | 0.63945 | 1.70759 | 2.05577 | 0.56937 | 0.21649 | 0.54803 |
| 140 | Taraxacum officinale F.H.Wigg. | Taraxacum officinale agg. | 0.2 | 1 | 0.268 | 7 | 5.6 | 4.4 | 5.9 | 5.4 | 0.1 | 0.64871 | 0.63945 | 1.70759 | 2.05577 | 0.56937 | 0.21649 | 0.54803 |
| 140 | Medicago lupulina L. | Medicago lupulina | 0.4 | 1 | 0.059 | 7 | 5.6 | 4.4 | 5.9 | 5.4 | 0.1 | 0.64871 | 0.63945 | 1.70759 | 2.05577 | 0.56937 | 0.21649 | 0.54803 |
| 140 | Poa annua L. | Poa annua | 0.35 | 1 | 0.083 | 7 | 5.6 | 4.4 | 5.9 | 5.4 | 0.1 | 0.64871 | 0.63945 | 1.70759 | 2.05577 | 0.56937 | 0.21649 | 0.54803 |
| 141 | Taraxacum officinale F.H.Wigg. | Taraxacum officinale agg. | 0.13 | 1 | 0.278 | NA | NA | NA | NA | NA | NA | NA | NA | NA | NA | NA | NA | NA |
| 141 | Plantago major L. | Plantago major ssp. major | 0.17 | 1 | 0.400 | NA | NA | NA | NA | NA | NA | NA | NA | NA | NA | NA | NA | NA |
| 141 | Fragaria vesca L. | Fragaria vesca | 0.25 | 1 | 1.000 | NA | NA | NA | NA | NA | NA | NA | NA | NA | NA | NA | NA | NA |
| 141 | Prunella vulgaris L. | Prunella vulgaris | 0.17 | 1 | 0.333 | NA | NA | NA | NA | NA | NA | NA | NA | NA | NA | NA | NA | NA |
| 141 | Stellaria media (L.) Vill. | Stellaria media agg. | 0.02 | 1 | 0.477 | 7.4 | 5.8 | 4.2 | 6.7 | 5.3 | 0 | 0.62702 | 0.59749 | 1.35516 | 1.96644 | 0.57704 | 0.23154 | 0.49642 |
| 141 | Dactylis glomerata L. | Dactylis glomerata | 0.15 | 1 | 0.118 | NA | NA | NA | NA | NA | NA | NA | NA | NA | NA | NA | NA | NA |
| 141 | Poa annua L. | Poa annua | 0.11 | 1 | 0.150 | 7.2 | NA | NA | 5.5 | 4.7 | 0.5 | 0.51067 | 0.4744 | 1.23473 | 1.90522 | 0.81709 | 0.28699 | 0.29055 |
| 142 | Taraxacum officinale F.H.Wigg. | Taraxacum officinale agg. | 0.23 | 1 | 0.100 | 7.3 | 5.4 | 4.1 | 7.6 | 4.2 | 0.4 | 0.58728 | 0.56055 | 1.34601 | 1.94534 | 0.82357 | 0.23743 | 0.34603 |
| 142 | Veronica arvensis L. | Veronica arvensis | 0.05 | 1 | 0.200 | 6.9 | 5.3 | 6.4 | 7.3 | 6.7 | 1 | 0.60499 | 0.3452 | 0.36505 | 1.40223 | 0.13192 | 0.19582 | 0.16678 |
| 142 | Cerastium semidecandrum L. | Cerastium semidecandrum | 0.08 | 1 | 0.077 | 6.3 | NA | 5.9 | 6.3 | 6.9 | 0.1 | 0.6037 | 0.3946 | 0.44368 | 1.50139 | 0.30164 | 0.21315 | 0.21494 |
| 142 | Viola odorata L. | Viola odorata | 0.05 | 1 | 0.167 | 7.3 | NA | NA | 6.5 | 2.7 | 0.3 | 0.50155 | 0.46101 | 1.19593 | 1.8416 | 0.70249 | 0.27259 | 0.24586 |
| 142 | Sonchus oleraceus L. | Sonchus oleraceus | 0.05 | 1 | 0.786 | 7.4 | 5.3 | NA | 6 | 7.7 | 0.8 | 0.65201 | 0.63791 | 1.54898 | 2.03286 | 0.77966 | 0.37419 | 0.45721 |
| 142 | Bromus sterilis L. | Bromus sterilis | 0.1 | 1 | 0.200 | 7.9 | 6.2 | NA | 6.7 | 6.4 | 0.2 | 0.62749 | 0.58201 | 1.11218 | 1.53794 | 0.69579 | 0.21134 | 0.22838 |
| 142 | Poa annua L. | Poa annua | 0.36 | 1 | 0.200 | 7.9 | 6.2 | NA | 6.7 | 6.4 | 0.2 | 0.62749 | 0.58201 | 1.11218 | 1.53794 | 0.69579 | 0.21134 | 0.22838 |
| 142 | Stellaria media (L.) Vill. | Stellaria media agg. | 0.08 | 1 | 0.056 | 6.3 | NA | 5.9 | 6.3 | 6.9 | 0.1 | 0.6037 | 0.3946 | 0.44368 | 1.50139 | 0.30164 | 0.21315 | 0.21494 |
| 143 | Taraxacum officinale F.H.Wigg. | Taraxacum officinale agg. | 0.1 | 0.58 | 0.050 | 6.7 | 5.3 | 4.5 | 5.3 | 5.8 | 0 | 0.75464 | 0.73051 | 1.36974 | 1.95508 | 0.45974 | 0.15567 | 0.6659 |
| 143 | Bellis perennis L. | Bellis perennis | 0.1 | 0.58 | 0.111 | 6.9 | 5.9 | NA | 6.8 | 7 | 0 | 0.84157 | 0.83772 | 1.85474 | 2.04042 | 0.4856 | 0.14178 | 0.80173 |
| 143 | Viola odorata L. | Viola odorata | 0.1 | 0.58 | 0.474 | NA | NA | NA | NA | NA | NA | NA | NA | NA | NA | NA | NA | NA |
| 143 | Stellaria media (L.) Vill. | Stellaria media agg. | 0.05 | 0.58 | 0.063 | 7.3 | 6.1 | 3.8 | 6 | 5.1 | 0 | 0.62987 | 0.59281 | 1.27995 | 1.90491 | 0.6275 | 0.3036 | 0.42608 |
| 143 | Prunella vulgaris L. | Prunella vulgaris | 0.08 | 0.58 | 0.111 | 8 | 5 | NA | 6.1 | 6.6 | 0.4 | 0.62393 | 0.60267 | 1.39471 | 1.77442 | 0.6433 | 0.24014 | 0.43141 |
| 143 | Bromus sterilis L. | Bromus sterilis | 0.05 | 0.58 | 0.033 | 7.3 | 6.1 | 3.8 | 6 | 5.1 | 0 | 0.62987 | 0.59281 | 1.27995 | 1.90491 | 0.6275 | 0.3036 | 0.42608 |
| 143 | Poa annua L. | Poa annua | 0.1 | 0.58 | 0.053 | NA | NA | 4.5 | 6.6 | 6.6 | 0 | 0.78059 | 0.60963 | 0.54418 | 1.63701 | 0.26733 | 0.16232 | 0.54354 |
| 144 | Taraxacum officinale F.H.Wigg. | Taraxacum officinale agg. | 0.1 | 0.95 | 0.118 | NA | NA | NA | NA | NA | NA | NA | NA | NA | NA | NA | NA | NA |
| 144 | Stellaria media (L.) Vill. | Stellaria media agg. | 0.2 | 0.95 | 0.778 | 7.4 | 5.3 | NA | 6 | 7.7 | 0.8 | 0.65201 | 0.63791 | 1.54898 | 2.03286 | 0.77966 | 0.37419 | 0.45721 |
| 144 | Fragaria vesca L. | Fragaria vesca | 0.1 | 0.95 | 0.105 | NA | NA | NA | NA | NA | NA | NA | NA | NA | NA | NA | NA | NA |
| 144 | Prunella vulgaris L. | Prunella vulgaris | 0.2 | 0.95 | 0.059 | 6.7 | NA | NA | 5.2 | NA | 0.4 | 0.59706 | 0.49429 | 0.69366 | 1.8006 | 0.482 | 0.19483 | 0.19138 |
| 144 | Viola odorata L. | Viola odorata | 0.05 | 0.95 | 0.211 | 6.6 | NA | NA | 6.9 | 5.2 | 0.4 | 0.54634 | 0.5169 | 1.30759 | 1.85566 | 0.79671 | 0.26499 | 0.28358 |
| 144 | Bromus sterilis L. | Bromus sterilis | 0.1 | 0.95 | 0.056 | NA | 5.4 | 5.7 | 6.8 | 7 | 0 | 0.67742 | 0.24559 | 0.14546 | 1.36309 | 0.06547 | 0.19208 | 0.12644 |

| | | | | | | | | | | | | | | | | | | |
|---|---|---|---|---|---|---|---|---|---|---|---|---|---|---|---|---|---|---|
| 144 | Poa annua L. | Poa annua | 0.12 | 0.95 | 0.056 | NA | 5.4 | 5.7 | 6.8 | 7 | 0 | 0.67742 | 0.24559 | 0.14546 | 1.36309 | 0.06547 | 0.19208 | 0.12644 |
| 144 | Glechoma hederacea L. | Glechoma hederacea | 0.08 | 0.95 | 0.059 | 8.1 | NA | 3.5 | 7.5 | 5 | 0.3 | 0.75184 | 0.72654 | 1.3537 | 1.85024 | 0.717 | 0.18024 | 0.48387 |
| 145 | Taraxacum officinale F.H.Wigg. | Taraxacum officinale agg. | 0.1 | 1 | 0.071 | 6.9 | 6.3 | 3.9 | 6.3 | 7.1 | 0.1 | 0.77916 | 0.67616 | 0.95204 | 1.78267 | 0.43197 | 0.25047 | 0.50688 |
| 145 | Fragaria vesca L. | Fragaria vesca | 0.35 | 1 | 0.300 | NA | NA | 4.5 | 6.6 | 6.6 | 0 | 0.78059 | 0.60963 | 0.54418 | 1.63701 | 0.26733 | 0.16232 | 0.54354 |
| 145 | Glechoma hederacea L. | Glechoma hederacea | 0.1 | 1 | 0.105 | 8.1 | NA | NA | 5.2 | NA | 0.1 | 0.51276 | 0.49581 | 1.51667 | 2.20989 | 0.98959 | 0.46997 | 0.3594 |
| 145 | Viola odorata L. | Viola odorata | 0.05 | 1 | 0.100 | NA | NA | 4.5 | 6.6 | 6.6 | 0 | 0.78059 | 0.60963 | 0.54418 | 1.63701 | 0.26733 | 0.16232 | 0.54354 |
| 145 | Prunella vulgaris L. | Prunella vulgaris | 0.05 | 1 | 0.500 | NA | NA | 4.5 | 6.6 | 6.6 | 0 | 0.78059 | 0.60963 | 0.54418 | 1.63701 | 0.26733 | 0.16232 | 0.54354 |
| 145 | Bromus sterilis L. | Bromus sterilis | 0.1 | 1 | 0.200 | NA | 5.4 | 5.7 | 6.8 | 7 | 0 | 0.67742 | 0.24559 | 0.14546 | 1.36309 | 0.06547 | 0.19208 | 0.12644 |
| 145 | Poa annua L. | Poa annua | 0.25 | 1 | 0.050 | 8.1 | NA | 3.5 | 6.1 | NA | 0.2 | 0.68175 | 0.6716 | 1.70824 | 1.99767 | 0.41848 | 0.25411 | 0.57318 |
| 146 | Taraxacum officinale F.H.Wigg. | Taraxacum officinale agg. | 0.1 | 1 | 0.290 | 7.3 | 6.1 | 3.8 | 6 | 5.1 | 0 | 0.62987 | 0.59281 | 1.27995 | 1.90491 | 0.6275 | 0.3036 | 0.42608 |
| 146 | Fragaria vesca L. | Fragaria vesca | 0.35 | 1 | 0.050 | 7.2 | 6 | 4.1 | 6.8 | 4.9 | 0 | 0.72892 | 0.658 | 1.07912 | 1.91928 | 0.406 | 0.27624 | 0.47339 |
| 146 | Glechoma hederacea L. | Glechoma hederacea | 0.1 | 1 | 0.217 | 7.4 | 5.3 | NA | 6 | 7.7 | 0.8 | 0.65201 | 0.63791 | 1.54898 | 2.03286 | 0.77966 | 0.37419 | 0.45721 |
| 146 | Viola odorata L. | Viola odorata | 0.05 | 1 | 0.267 | 7.4 | 5.3 | NA | 6 | 7.7 | 0.8 | 0.65201 | 0.63791 | 1.54898 | 2.03286 | 0.77966 | 0.37419 | 0.45721 |
| 146 | Prunella vulgaris L. | Prunella vulgaris | 0.04 | 1 | 0.204 | 7.4 | 5.3 | NA | 6 | 7.7 | 0.8 | 0.65201 | 0.63791 | 1.54898 | 2.03286 | 0.77966 | 0.37419 | 0.45721 |
| 146 | Hypericum perforatum L. | Hypericum perforatum | 0.08 | 1 | 0.118 | 8 | NA | NA | 6.1 | 6.3 | 0.1 | 0.56333 | 0.55394 | 1.72195 | 2.16326 | 1.07172 | 0.43309 | 0.38415 |
| 146 | Cerastium semidecandrum L. | Cerastium semidecandrum | 0.04 | 1 | 0.109 | 8 | NA | NA | 6.1 | 6.3 | 0.1 | 0.56333 | 0.55394 | 1.72195 | 2.16326 | 1.07172 | 0.43309 | 0.38415 |
| 146 | Bromus sterilis L. | Bromus sterilis | 0.05 | 1 | 0.103 | 8 | NA | NA | 6.1 | 6.3 | 0.1 | 0.56333 | 0.55394 | 1.72195 | 2.16326 | 1.07172 | 0.43309 | 0.38415 |
| 146 | Poa annua L. | Poa annua | 0.19 | 1 | 0.109 | 8 | NA | NA | 6.1 | 6.3 | 0.1 | 0.56333 | 0.55394 | 1.72195 | 2.16326 | 1.07172 | 0.43309 | 0.38415 |
| 147 | Taraxacum officinale F.H.Wigg. | Taraxacum officinale agg. | 0.05 | 1 | 0.119 | 8 | NA | NA | 6.1 | 6.3 | 0.1 | 0.56333 | 0.55394 | 1.72195 | 2.16326 | 1.07172 | 0.43309 | 0.38415 |
| 147 | Fragaria vesca L. | Fragaria vesca | 0.04 | 1 | 0.157 | 7.4 | 5.3 | NA | 6 | 7.7 | 0.8 | 0.65201 | 0.63791 | 1.54898 | 2.03286 | 0.77966 | 0.37419 | 0.45721 |
| 147 | Glechoma hederacea L. | Glechoma hederacea | 0.05 | 1 | 0.070 | 7 | 5.6 | 4.4 | 5.9 | 5.4 | 0.1 | 0.64871 | 0.63945 | 1.70759 | 2.05577 | 0.56937 | 0.21649 | 0.54803 |
| 147 | Viola odorata L. | Viola odorata | 0.09 | 1 | 0.250 | 6 | NA | 6.4 | 7.5 | 6.1 | 0.1 | 0.67416 | 0.35728 | 0.31529 | 1.53539 | 0.05981 | 0.19852 | 0.1369 |
| 147 | Prunella vulgaris L. | Prunella vulgaris | 0.09 | 1 | 0.186 | 7.3 | 5.4 | 4.1 | 7.6 | 4.2 | 0.4 | 0.58728 | 0.56055 | 1.34601 | 1.94534 | 0.82357 | 0.23743 | 0.34603 |
| 147 | Trifolium repens L. | Trifolium repens | 0.15 | 1 | 0.133 | 7.3 | 5.4 | 4.1 | 7.6 | 4.2 | 0.4 | 0.58728 | 0.56055 | 1.34601 | 1.94534 | 0.82357 | 0.23743 | 0.34603 |
| 147 | Cerastium semidecandrum L. | Cerastium semidecandrum | 0.04 | 1 | 0.059 | 7.3 | 5.4 | 4.1 | 7.6 | 4.2 | 0.4 | 0.58728 | 0.56055 | 1.34601 | 1.94534 | 0.82357 | 0.23743 | 0.34603 |
| 147 | Hypericum perforatum L. | Hypericum perforatum | 0.05 | 1 | 0.051 | 8.4 | 7.2 | 2.6 | 6.8 | 3.6 | 0.2 | 0.58203 | 0.49726 | 0.89718 | 1.5951 | 0.3257 | 0.17939 | 0.36267 |
| 147 | Cerastium fontanum Baumg. | Cerastium fontanum fontanum | 0.04 | 1 | 0.080 | 7.3 | 5.4 | 4.1 | 7.6 | 4.2 | 0.4 | 0.58728 | 0.56055 | 1.34601 | 1.94534 | 0.82357 | 0.23743 | 0.34603 |
| 147 | Lolium perenne L. | Lolium perenne | 0.05 | 1 | 0.167 | 8.1 | NA | 3.5 | 6.1 | NA | 0.2 | 0.68175 | 0.6716 | 1.70824 | 1.99767 | 0.41848 | 0.25411 | 0.57318 |
| 147 | Stellaria media (L.) Vill. | Stellaria media agg. | 0.1 | 1 | 0.105 | 8.1 | NA | 3.5 | 6.1 | NA | 0.2 | 0.68175 | 0.6716 | 1.70824 | 1.99767 | 0.41848 | 0.25411 | 0.57318 |
| 147 | Bromus sterilis L. | Bromus sterilis | 0.1 | 1 | 0.100 | 8.1 | NA | 3.5 | 6.1 | NA | 0.2 | 0.68175 | 0.6716 | 1.70824 | 1.99767 | 0.41848 | 0.25411 | 0.57318 |
| 147 | Poa annua L. | Poa annua | 0.15 | 1 | 0.080 | 7.9 | NA | 4.1 | 5.8 | NA | 0.9 | 0.54139 | 0.51128 | 1.31363 | 1.93633 | 0.8112 | 0.25085 | 0.32953 |
| 148 | Taraxacum officinale F.H.Wigg. | Taraxacum officinale agg. | 0.08 | 1 | 0.020 | 7 | 5.6 | 4.4 | 5.9 | 5.4 | 0.1 | 0.64871 | 0.63945 | 1.70759 | 2.05577 | 0.56937 | 0.21649 | 0.54803 |
| 148 | Glechoma hederacea L. | Glechoma hederacea | 0.13 | 1 | 0.050 | 7 | 5.6 | 4.4 | 5.9 | 5.4 | 0.1 | 0.64871 | 0.63945 | 1.70759 | 2.05577 | 0.56937 | 0.21649 | 0.54803 |
| 148 | Viola odorata L. | Viola odorata | 0.13 | 1 | 0.050 | 8.6 | NA | 2.7 | 4.4 | 1.6 | 0 | 0.53557 | 0.50485 | 1.29532 | 1.90211 | 0.72298 | 0.28495 | 0.2778 |
| 148 | Hypericum perforatum L. | Hypericum perforatum | 0.04 | 1 | 0.084 | 8.1 | NA | 3.5 | 6.1 | NA | 0.2 | 0.68175 | 0.6716 | 1.70824 | 1.99767 | 0.41848 | 0.25411 | 0.57318 |
| 148 | Trifolium repens L. | Trifolium repens | 0.05 | 1 | 0.350 | 7.4 | 5.3 | NA | 6 | 7.7 | 0.8 | 0.65201 | 0.63791 | 1.54898 | 2.03286 | 0.77966 | 0.37419 | 0.45721 |
| 148 | Chaerophyllum temulum L. | Chaerophyllum temulum | 0.04 | 1 | 0.020 | 7 | 5.6 | 4.4 | 5.9 | 5.4 | 0.1 | 0.64871 | 0.63945 | 1.70759 | 2.05577 | 0.56937 | 0.21649 | 0.54803 |
| 148 | Cerastium fontanum Baumg. | Cerastium fontanum fontanum | 0.08 | 1 | 0.050 | 6.7 | 5.3 | 4.5 | 5.3 | 5.8 | 0 | 0.75464 | 0.73051 | 1.36974 | 1.95508 | 0.45974 | 0.15567 | 0.6659 |
| 148 | Lolium perenne L. | Lolium perenne | 0.05 | 1 | 0.040 | 7 | 5.6 | 4.4 | 5.9 | 5.4 | 0.1 | 0.64871 | 0.63945 | 1.70759 | 2.05577 | 0.56937 | 0.21649 | 0.54803 |
| 148 | Poa annua L. | Poa annua | 0.2 | 1 | 0.050 | 8.6 | NA | 2.7 | 4.4 | 1.6 | 0 | 0.53557 | 0.50485 | 1.29532 | 1.90211 | 0.72298 | 0.28495 | 0.2778 |
| 148 | Prunella vulgaris L. | Prunella vulgaris | 0.13 | 1 | 0.050 | NA | NA | NA | NA | NA | NA | NA | NA | NA | NA | NA | NA | NA |
| 148 | Plantago major L. | Plantago major ssp. major | 0.05 | 1 | 0.090 | 8.1 | NA | 3.5 | 6.1 | NA | 0.2 | 0.68175 | 0.6716 | 1.70824 | 1.99767 | 0.41848 | 0.25411 | 0.57318 |
| 148 | Bellis perennis L. | Bellis perennis | 0.02 | 1 | 0.050 | NA | NA | NA | NA | NA | NA | NA | NA | NA | NA | NA | NA | NA |
| 149 | Taraxacum officinale F.H.Wigg. | Taraxacum officinale agg. | 0.08 | 1 | 0.020 | 8.5 | NA | 3.5 | 6.3 | 4.1 | 0.2 | 0.57633 | 0.55499 | 1.44489 | 1.81436 | 0.579 | 0.25616 | 0.27657 |
| 149 | Glechoma hederacea L. | Glechoma hederacea | 0.06 | 1 | 0.053 | 7.3 | NA | NA | 5.8 | 5.9 | 0.5 | 0.63207 | 0.61738 | 1.52973 | 2.00883 | 0.70322 | 0.36696 | 0.39725 |
| 149 | Viola odorata L. | Viola odorata | 0.04 | 1 | 0.150 | 8.6 | NA | 2.7 | 4.4 | 1.6 | 0 | 0.53557 | 0.50485 | 1.29532 | 1.90211 | 0.72298 | 0.28495 | 0.2778 |
| 149 | Fragaria vesca L. | Fragaria vesca | 0.23 | 1 | 0.040 | 5.2 | 5.9 | 5.4 | 7.3 | 8 | 0 | 0.67947 | 0.25214 | 0.1495 | 1.33661 | 0.07498 | 0.19025 | 0.12819 |
| 149 | Prunella vulgaris L. | Prunella vulgaris | 0.23 | 1 | 0.040 | 7.3 | 6.1 | 3.8 | 6 | 5.1 | 0 | 0.62987 | 0.59281 | 1.27995 | 1.90491 | 0.6275 | 0.3036 | 0.42608 |
| 149 | Chaerophyllum temulum L. | Chaerophyllum temulum | 0.05 | 1 | 0.089 | 7.9 | NA | 4.1 | 5.8 | NA | 0.9 | 0.54139 | 0.51128 | 1.31363 | 1.93633 | 0.8112 | 0.25085 | 0.32953 |
| 149 | Trifolium repens L. | Trifolium repens | 0.05 | 1 | 0.308 | 7.9 | NA | 4.1 | 5.8 | NA | 0.9 | 0.54139 | 0.51128 | 1.31363 | 1.93633 | 0.8112 | 0.25085 | 0.32953 |
| 149 | Lolium perenne L. | Lolium perenne | 0.12 | 1 | 0.333 | 7.9 | NA | 4.1 | 5.8 | NA | 0.9 | 0.54139 | 0.51128 | 1.31363 | 1.93633 | 0.8112 | 0.25085 | 0.32953 |
| 149 | Poa annua L. | Poa annua | 0.1 | 1 | 0.286 | 7.9 | NA | 4.1 | 5.8 | NA | 0.9 | 0.54139 | 0.51128 | 1.31363 | 1.93633 | 0.8112 | 0.25085 | 0.32953 |
| 149 | Plantago major L. | Plantago major ssp. major | 0.04 | 1 | 0.267 | 7.9 | NA | 4.1 | 5.8 | NA | 0.9 | 0.54139 | 0.51128 | 1.31363 | 1.93633 | 0.8112 | 0.25085 | 0.32953 |
| 150 | Bellis perennis L. | Bellis perennis | 0.04 | 0.72 | 0.300 | 7.9 | NA | 4.1 | 5.8 | NA | 0.9 | 0.54139 | 0.51128 | 1.31363 | 1.93633 | 0.8112 | 0.25085 | 0.32953 |
| 150 | Fragaria vesca L. | Fragaria vesca | 0.07 | 0.72 | 0.500 | 7.9 | NA | 4.1 | 5.8 | NA | 0.9 | 0.54139 | 0.51128 | 1.31363 | 1.93633 | 0.8112 | 0.25085 | 0.32953 |
| 150 | Glechoma hederacea L. | Glechoma hederacea | 0.03 | 0.72 | 0.086 | 8.4 | NA | 3 | 6 | NA | 0 | 0.46249 | 0.43915 | 1.39189 | 1.83687 | 0.38437 | 0.30571 | 0.38883 |
| 150 | Viola odorata L. | Viola odorata | 0.05 | 0.72 | 0.125 | 7.9 | NA | 4.1 | 5.8 | NA | 0.9 | 0.54139 | 0.51128 | 1.31363 | 1.93633 | 0.8112 | 0.25085 | 0.32953 |
| 150 | Trifolium repens L. | Trifolium repens | 0.07 | 0.72 | 0.650 | 7.6 | NA | NA | 6.5 | 6.3 | 0.7 | 0.75548 | 0.75048 | 1.8409 | 2.08419 | 0.63746 | 0.23177 | 0.58814 |
| 150 | Cerastium fontanum Baumg. | Cerastium fontanum fontanum | 0.07 | 0.72 | 0.050 | NA | NA | NA | NA | NA | NA | NA | NA | NA | NA | NA | NA | NA |
| 150 | Medicago lupulina L. | Medicago lupulina | 0.04 | 0.72 | 0.136 | NA | NA | NA | NA | NA | NA | NA | NA | NA | NA | NA | NA | NA |
| 150 | Prunella vulgaris L. | Prunella vulgaris | 0.07 | 0.72 | 0.526 | 7.4 | 5.3 | NA | 6 | 7.7 | 0.8 | 0.65201 | 0.63791 | 1.54898 | 2.03286 | 0.77966 | 0.37419 | 0.45721 |
| 150 | Taraxacum officinale F.H.Wigg. | Taraxacum officinale agg. | 0.13 | 0.72 | 0.150 | NA | NA | NA | NA | NA | NA | NA | NA | NA | NA | NA | NA | NA |
| 150 | Hypericum perforatum L. | Hypericum perforatum | 0.02 | 0.72 | 0.050 | 7.3 | 5.4 | 4.1 | 7.6 | 4.2 | 0.4 | 0.58728 | 0.56055 | 1.34601 | 1.94534 | 0.82357 | 0.23743 | 0.34603 |
| 150 | Poa annua L. | Poa annua | 0.13 | 0.72 | 0.100 | 7.3 | 6.1 | 3.8 | 6 | 5.1 | 0 | 0.62987 | 0.59281 | 1.27995 | 1.90491 | 0.6275 | 0.3036 | 0.42608 |
| 151 | Bellis perennis L. | Bellis perennis | 0.05 | 0.65 | 0.051 | 7.3 | 6.1 | 3.8 | 6 | 5.1 | 0 | 0.62987 | 0.59281 | 1.27995 | 1.90491 | 0.6275 | 0.3036 | 0.42608 |
| 151 | Viola odorata L. | Viola odorata | 0.05 | 0.65 | 0.250 | 7.3 | 6.1 | 3.8 | 6 | 5.1 | 0 | 0.62987 | 0.59281 | 1.27995 | 1.90491 | 0.6275 | 0.3036 | 0.42608 |
| 151 | Trifolium repens L. | Trifolium repens | 0.04 | 0.65 | 0.102 | 8.6 | NA | 2.7 | 4.4 | 1.6 | 0 | 0.53557 | 0.50485 | 1.29532 | 1.90211 | 0.72298 | 0.28495 | 0.2778 |
| 151 | Medicago lupulina L. | Medicago lupulina | 0.04 | 0.65 | 0.250 | 7.1 | 5.3 | 4.5 | 5.9 | 4.8 | 0.3 | 0.52518 | 0.5113 | 1.61045 | 2.05509 | 0.96495 | 0.27897 | 0.32911 |
| 151 | Chaerophyllum temulum L. | Chaerophyllum temulum | 0.04 | 0.65 | 0.350 | 7.3 | 5.4 | 4.1 | 7.6 | 4.2 | 0.4 | 0.58728 | 0.56055 | 1.34601 | 1.94534 | 0.82357 | 0.23743 | 0.34603 |
| 151 | Symphoricarpos albus (L.) S.F.Blake | NA | 0.05 | 0.65 | 0.300 | 7.3 | 5.4 | 4.1 | 7.6 | 4.2 | 0.4 | 0.58728 | 0.56055 | 1.34601 | 1.94534 | 0.82357 | 0.23743 | 0.34603 |
| 151 | Prunella vulgaris L. | Prunella vulgaris | 0.1 | 0.65 | 0.050 | 7.3 | 5.4 | 4.1 | 7.6 | 4.2 | 0.4 | 0.58728 | 0.56055 | 1.34601 | 1.94534 | 0.82357 | 0.23743 | 0.34603 |
| 151 | Cerastium fontanum Baumg. | Cerastium fontanum fontanum | 0.04 | 0.65 | 0.050 | 7.3 | 5.4 | 4.1 | 7.6 | 4.2 | 0.4 | 0.58728 | 0.56055 | 1.34601 | 1.94534 | 0.82357 | 0.23743 | 0.34603 |
| 151 | Glechoma hederacea L. | Glechoma hederacea | 0.04 | 0.65 | 0.089 | 8.1 | NA | 3.5 | 6.1 | NA | 0.2 | 0.68175 | 0.6716 | 1.70824 | 1.99767 | 0.41848 | 0.25411 | 0.57318 |
| 151 | Hypericum perforatum L. | Hypericum perforatum | 0.04 | 0.65 | 0.200 | 7.3 | 5.4 | 4.1 | 7.6 | 4.2 | 0.4 | 0.58728 | 0.56055 | 1.34601 | 1.94534 | 0.82357 | 0.23743 | 0.34603 |
| 151 | Bromus sterilis L. | Bromus sterilis | 0.05 | 0.65 | 0.400 | 7.3 | 5.4 | 4.1 | 7.6 | 4.2 | 0.4 | 0.58728 | 0.56055 | 1.34601 | 1.94534 | 0.82357 | 0.23743 | 0.34603 |
| 151 | Poa annua L. | Poa annua | 0.11 | 0.65 | 0.250 | 6.7 | NA | 4.9 | 6.8 | 5 | 0 | 0.66057 | 0.18432 | 0.11528 | 1.34075 | 0.06671 | 0.20032 | 0.12881 |
| 152 | Bellis perennis L. | Bellis perennis | 0.05 | 0.78 | 0.080 | 8.4 | NA | 3 | 6 | NA | 0 | 0.46249 | 0.43915 | 1.39189 | 1.83687 | 0.38437 | 0.30571 | 0.38883 |
| 152 | Viola odorata L. | Viola odorata | 0.08 | 0.78 | 0.105 | 6.7 | NA | 4.9 | 6.8 | 5 | 0 | 0.66057 | 0.18432 | 0.11528 | 1.34075 | 0.06671 | 0.20032 | 0.12881 |
| 152 | Trifolium repens L. | Trifolium repens | 0.09 | 0.78 | 0.100 | 6.3 | NA | 5.9 | 6.3 | 6.9 | 0.1 | 0.6037 | 0.3946 | 0.44368 | 1.50139 | 0.30164 | 0.21315 | 0.21494 |
| 152 | Prunella vulgaris L. | Prunella vulgaris | 0.09 | 0.78 | 0.100 | 6.3 | NA | 5.9 | 6.3 | 6.9 | 0.1 | 0.6037 | 0.3946 | 0.44368 | 1.50139 | 0.30164 | 0.21315 | 0.21494 |
| 152 | Taraxacum officinale F.H.Wigg. | Taraxacum officinale agg. | 0.09 | 0.78 | 0.050 | 6.3 | NA | 5.9 | 6.3 | 6.9 | 0.1 | 0.6037 | 0.3946 | 0.44368 | 1.50139 | 0.30164 | 0.21315 | 0.21494 |
| 152 | Fragaria vesca L. | Fragaria vesca | 0.03 | 0.78 | 0.130 | NA | NA | NA | 7 | NA | 0 | 0.66957 | 0.20824 | 0.12967 | 1.34554 | 0.07292 | 0.20428 | 0.13248 |
| 152 | Veronica arvensis L. | Veronica arvensis | 0.01 | 0.78 | 0.040 | NA | NA | NA | 7 | NA | 0 | 0.66957 | 0.20824 | 0.12967 | 1.34554 | 0.07292 | 0.20428 | 0.13248 |
| 152 | Bromus sterilis L. | Bromus sterilis | 0.09 | 0.78 | 0.042 | 6.3 | NA | 5.9 | 6.3 | 6.9 | 0.1 | 0.6037 | 0.3946 | 0.44368 | 1.50139 | 0.30164 | 0.21315 | 0.21494 |
| 152 | Poa annua L. | Poa annua | 0.25 | 0.78 | 0.062 | 7.7 | NA | NA | 5.6 | NA | 0.7 | 0.53223 | 0.52325 | 1.73719 | 2.14421 | 0.94278 | 0.39374 | 0.35054 |
| 153 | Viola odorata L. | Viola odorata | 0.05 | 1 | 0.115 | 7.7 | NA | NA | 5.6 | NA | 0.7 | 0.53223 | 0.52325 | 1.73719 | 2.14421 | 0.94278 | 0.39374 | 0.35054 |
| 153 | Trifolium repens L. | Trifolium repens | 0.1 | 1 | 0.100 | NA | NA | NA | NA | NA | NA | NA | NA | NA | NA | NA | NA | NA |
| 153 | Taraxacum officinale F.H.Wigg. | Taraxacum officinale agg. | 0.1 | 1 | 0.263 | 7.3 | 5.4 | 4.1 | 7.6 | 4.2 | 0.4 | 0.58728 | 0.56055 | 1.34601 | 1.94534 | 0.82357 | 0.23743 | 0.34603 |
| 153 | Bellis perennis L. | Bellis perennis | 0.05 | 1 | 0.176 | 8.6 | NA | 2.7 | 4.4 | 1.6 | 0 | 0.53557 | 0.50485 | 1.29532 | 1.90211 | 0.72298 | 0.28495 | 0.2778 |
| 153 | Veronica arvensis L. | Veronica arvensis | 0.02 | 1 | 0.050 | 8.6 | NA | 2.7 | 4.4 | 1.6 | 0 | 0.53557 | 0.50485 | 1.29532 | 1.90211 | 0.72298 | 0.28495 | 0.2778 |
| 153 | Prunella vulgaris L. | Prunella vulgaris | 0.1 | 1 | 0.050 | 8.1 | NA | NA | 5.2 | NA | 0.1 | 0.51276 | 0.49581 | 1.51667 | 2.20989 | 0.98959 | 0.46997 | 0.3594 |
| 153 | Medicago lupulina L. | Medicago lupulina | 0.15 | 1 | 0.250 | 7.4 | 5.3 | NA | 6 | 7.7 | 0.8 | 0.65201 | 0.63791 | 1.54898 | 2.03286 | 0.77966 | 0.37419 | 0.45721 |
| 153 | NA | NA | 0.04 | 1 | 0.040 | 7.9 | NA | 4.1 | 5.8 | NA | 0.9 | 0.54139 | 0.51128 | 1.31363 | 1.93633 | 0.8112 | 0.25085 | 0.32953 |
| 153 | Cerastium fontanum Baumg. | Cerastium fontanum fontanum | 0.04 | 1 | 0.050 | 8.4 | NA | 3 | 6 | NA | 0 | 0.46249 | 0.43915 | 1.39189 | 1.83687 | 0.38437 | 0.30571 | 0.38883 |
| 153 | Bromus sterilis L. | Bromus sterilis | 0.05 | 1 | 0.045 | 8.4 | NA | 3 | 6 | NA | 0 | 0.46249 | 0.43915 | 1.39189 | 1.83687 | 0.38437 | 0.30571 | 0.38883 |
| 153 | Poa annua L. | Poa annua | 0.3 | 1 | 0.050 | 7.9 | NA | 4.1 | 5.8 | NA | 0.9 | 0.54139 | 0.51128 | 1.31363 | 1.93633 | 0.8112 | 0.25085 | 0.32953 |
| 154 | Taraxacum officinale F.H.Wigg. | Taraxacum officinale agg. | 0.02 | 0.76 | 0.156 | 8.6 | NA | 2.7 | 4.4 | 1.6 | 0 | 0.53557 | 0.50485 | 1.29532 | 1.90211 | 0.72298 | 0.28495 | 0.2778 |
| 154 | Plantago lanceolata L. | Plantago lanceolata | 0.2 | 0.76 | 0.050 | NA | NA | NA | NA | NA | NA | NA | NA | NA | NA | NA | NA | NA |
| 154 | Medicago lupulina L. | Medicago lupulina | 0.2 | 0.76 | 0.100 | 7.1 | 5.3 | 4.5 | 5.9 | 4.8 | 0.3 | 0.52518 | 0.5113 | 1.61045 | 2.05509 | 0.96495 | 0.27897 | 0.32911 |
| 154 | Potentilla argentea L. | Potentilla argentea agg. | 0.04 | 0.76 | 0.040 | 8.1 | NA | NA | 5.2 | NA | 0.1 | 0.51276 | 0.49581 | 1.51667 | 2.20989 | 0.98959 | 0.46997 | 0.3594 |
| 154 | Poa annua L. | Poa annua | 0.3 | 0.76 | 0.570 | NA | NA | NA | NA | NA | NA | NA | NA | NA | NA | NA | NA | NA |
| 155 | Plantago lanceolata L. | Plantago lanceolata | 0.2 | 0.85 | 0.062 | 8.1 | NA | NA | 5.2 | NA | 0.1 | 0.51276 | 0.49581 | 1.51667 | 2.20989 | 0.98959 | 0.46997 | 0.3594 |
| 155 | Medicago lupulina L. | Medicago lupulina | 0.2 | 0.85 | 0.047 | 8.4 | 7.2 | 2.6 | 6.8 | 3.6 | 0.2 | 0.58203 | 0.49726 | 0.89718 | 1.5951 | 0.3257 | 0.17939 | 0.36267 |
| 155 | Potentilla argentea L. | Potentilla argentea agg. | 0.15 | 0.85 | 0.063 | 7.3 | NA | 3.9 | 6.1 | 3.9 | 0.1 | 0.55903 | 0.41538 | 0.65189 | 1.65181 | 0.43719 | 0.24279 | 0.21709 |
| 155 | Poa annua L. | Poa annua | 0.2 | 0.85 | 0.100 | 8.6 | NA | 2.7 | 4.4 | 1.6 | 0 | 0.53557 | 0.50485 | 1.29532 | 1.90211 | 0.72298 | 0.28495 | 0.2778 |
| 155 | Herniaria glabra L. | Herniaria glabra | 0.1 | 0.85 | 0.060 | 7.3 | NA | 3.9 | 6.1 | 3.9 | 0.1 | 0.55903 | 0.41538 | 0.65189 | 1.65181 | 0.43719 | 0.24279 | 0.21709 |

| | | | | | | | | | | | | | | | | | | |
|---|---|---|---|---|---|---|---|---|---|---|---|---|---|---|---|---|---|---|
| 156 | Plantago lanceolata L. | Plantago lanceolata | 0.3 | 1 | 0.053 | 8.6 | NA | 2.7 | 4.4 | 1.6 | 0 | 0.53557 | 0.50485 | 1.29532 | 1.90211 | 0.72298 | 0.28495 | 0.2778 |
| 156 | Medicago lupulina L. | Medicago lupulina | 0.2 | 1 | 0.108 | 8.6 | NA | 2.7 | 4.4 | 1.6 | 0 | 0.53557 | 0.50485 | 1.29532 | 1.90211 | 0.72298 | 0.28495 | 0.2778 |
| 156 | Potentilla argentea L. | Potentilla argentea agg. | 0.05 | 1 | 0.180 | 6.6 | NA | NA | 6.9 | 5.2 | 0.4 | 0.54634 | 0.5169 | 1.30759 | 1.85566 | 0.79671 | 0.26499 | 0.28358 |
| 156 | Poa annua L. | Poa annua | 0.45 | 1 | 0.053 | 8.6 | NA | 2.7 | 4.4 | 1.6 | 0 | 0.53557 | 0.50485 | 1.29532 | 1.90211 | 0.72298 | 0.28495 | 0.2778 |
| 157 | Plantago lanceolata L. | Plantago lanceolata | 0.1 | 1 | 0.200 | 7.3 | 5.4 | 4.1 | 7.6 | 4.2 | 0.4 | 0.58728 | 0.56055 | 1.34601 | 1.94534 | 0.82357 | 0.23743 | 0.34603 |
| 157 | Medicago lupulina L. | Medicago lupulina | 0.2 | 1 | 0.083 | 7.1 | NA | NA | NA | 4.6 | 0 | 0.69216 | 0.66495 | 1.32487 | 1.98453 | 0.49303 | 0.16764 | 0.56484 |
| 157 | Bellis perennis L. | Bellis perennis | 0.05 | 1 | 0.187 | 8.6 | NA | 2.7 | 4.4 | 1.6 | 0 | 0.53557 | 0.50485 | 1.29532 | 1.90211 | 0.72298 | 0.28495 | 0.2778 |
| 157 | Herniaria glabra L. | Herniaria glabra | 0.1 | 1 | 0.104 | 8.6 | NA | 2.7 | 4.4 | 1.6 | 0 | 0.53557 | 0.50485 | 1.29532 | 1.90211 | 0.72298 | 0.28495 | 0.2778 |
| 157 | Matricaria discoidea DC. | Matricaria discoidea | 0.1 | 1 | 0.303 | 7.9 | NA | 4.1 | 5.8 | NA | 0.9 | 0.54139 | 0.51128 | 1.31363 | 1.93633 | 0.8112 | 0.25085 | 0.32953 |
| 157 | Spergularia rubra J.Presl & C.Presl | Spergularia rubra | 0.02 | 1 | 0.600 | 7.4 | 5.3 | NA | 6 | 7.7 | 0.8 | 0.65201 | 0.63791 | 1.54898 | 2.03286 | 0.77966 | 0.37419 | 0.45721 |
| 157 | Poa annua L. | Poa annua | 0.3 | 1 | 0.101 | 8.6 | NA | 2.7 | 4.4 | 1.6 | 0 | 0.53557 | 0.50485 | 1.29532 | 1.90211 | 0.72298 | 0.28495 | 0.2778 |
| 157 | Sagina procumbens L. | Sagina procumbens | 0.13 | 1 | 0.050 | 8.1 | NA | NA | 4.4 | 2.8 | 0.3 | 0.48868 | 0.46508 | 1.4426 | 2.04965 | 0.7278 | 0.33931 | 0.35628 |
| 158 | Plantago lanceolata L. | Plantago lanceolata | 0.2 | 1 | 0.150 | NA | NA | NA | NA | NA | NA | NA | NA | NA | NA | NA | NA | NA |
| 158 | Medicago lupulina L. | Medicago lupulina | 0.25 | 1 | 0.200 | NA | NA | NA | NA | NA | NA | NA | NA | NA | NA | NA | NA | NA |
| 158 | Poa annua L. | Poa annua | 0.25 | 1 | 0.011 | 8.1 | NA | 3.5 | 6.1 | NA | 0.2 | 0.68175 | 0.6716 | 1.70824 | 1.99767 | 0.41848 | 0.25411 | 0.57318 |
| 158 | Taraxacum officinale F.H.Wigg. | Taraxacum officinale agg. | 0.2 | 1 | 0.153 | NA | NA | NA | NA | NA | NA | NA | NA | NA | NA | NA | NA | NA |
| 158 | Sagina procumbens L. | Sagina procumbens | 0.1 | 1 | 0.020 | 8.1 | NA | 3.5 | 6.1 | NA | 0.2 | 0.68175 | 0.6716 | 1.70824 | 1.99767 | 0.41848 | 0.25411 | 0.57318 |
| 159 | Medicago lupulina L. | Medicago lupulina | 0.1 | 1 | 0.100 | NA | NA | NA | NA | NA | NA | NA | NA | NA | NA | NA | NA | NA |
| 159 | Cerastium semidecandrum L. | Cerastium semidecandrum | 0.05 | 1 | 0.020 | 8.1 | NA | 3.5 | 6.1 | NA | 0.2 | 0.68175 | 0.6716 | 1.70824 | 1.99767 | 0.41848 | 0.25411 | 0.57318 |
| 159 | Achillea millefolium L. | Achillea millefolium millefolium | 0.04 | 1 | 0.208 | 8.1 | NA | NA | 4.4 | 2.8 | 0.3 | 0.48868 | 0.46508 | 1.4426 | 2.04965 | 0.7278 | 0.33931 | 0.35628 |
| 159 | Vicia hirsuta (L.) Gray | Vicia hirsuta | 0.05 | 1 | 0.050 | 8.1 | NA | 3.5 | 6.1 | NA | 0.2 | 0.68175 | 0.6716 | 1.70824 | 1.99767 | 0.41848 | 0.25411 | 0.57318 |
| 159 | Senecio inaequidens DC. | Senecio inaequidens | 0.05 | 1 | 0.050 | 8.1 | NA | 3.5 | 6.1 | NA | 0.2 | 0.68175 | 0.6716 | 1.70824 | 1.99767 | 0.41848 | 0.25411 | 0.57318 |
| 159 | Poa annua L. | Poa annua | 0.61 | 1 | 0.105 | NA | NA | NA | NA | NA | NA | NA | NA | NA | NA | NA | NA | NA |
| 159 | Lysimachia arvensis (L.) U.Manns & Anderb. | Anagallis arvensis | 0.02 | 1 | 0.051 | 8.5 | NA | 3.5 | 6.3 | 4.1 | 0.2 | 0.57633 | 0.55499 | 1.44489 | 1.81436 | 0.579 | 0.25616 | 0.27657 |
| 159 | Potentilla argentea L. | Potentilla argentea agg. | 0.02 | 1 | 0.050 | 8.4 | 7.2 | 2.6 | 6.8 | 3.6 | 0.2 | 0.58203 | 0.49726 | 0.89718 | 1.5951 | 0.3257 | 0.17939 | 0.36267 |
| 159 | Myosotis ramosissima Rochel | Myosotis ramosissima (coll. Hispida) | 0.04 | 1 | 0.100 | 8.1 | NA | NA | 4.4 | 2.8 | 0.3 | 0.48868 | 0.46508 | 1.4426 | 2.04965 | 0.7278 | 0.33931 | 0.35628 |
| 159 | Hypericum perforatum L. | Hypericum perforatum | 0.02 | 1 | 0.100 | NA | NA | NA | NA | NA | NA | NA | NA | NA | NA | NA | NA | NA |
| 160 | Medicago lupulina L. | Medicago lupulina | 0.1 | 1 | 0.100 | 8.5 | NA | 3.5 | 6.3 | 4.1 | 0.2 | 0.57633 | 0.55499 | 1.44489 | 1.81436 | 0.579 | 0.25616 | 0.27657 |
| 160 | Achillea millefolium L. | Achillea millefolium millefolium | 0.1 | 1 | 0.130 | NA | NA | NA | NA | NA | NA | NA | NA | NA | NA | NA | NA | NA |
| 160 | Cerastium semidecandrum L. | Cerastium semidecandrum | 0.05 | 1 | 0.130 | 8.1 | NA | NA | 4.4 | 2.8 | 0.3 | 0.48868 | 0.46508 | 1.4426 | 2.04965 | 0.7278 | 0.33931 | 0.35628 |
| 160 | Plantago lanceolata L. | Plantago lanceolata | 0.08 | 1 | 0.100 | 8.1 | NA | NA | 4.4 | 2.8 | 0.3 | 0.48868 | 0.46508 | 1.4426 | 2.04965 | 0.7278 | 0.33931 | 0.35628 |
| 160 | Poa annua L. | Poa annua | 0.6 | 1 | 0.150 | 8.1 | NA | NA | 4.4 | 2.8 | 0.3 | 0.48868 | 0.46508 | 1.4426 | 2.04965 | 0.7278 | 0.33931 | 0.35628 |
| 160 | Vicia hirsuta (L.) Gray | Vicia hirsuta | 0.02 | 1 | 0.143 | 6.7 | NA | NA | 6.5 | 7.7 | 0.1 | 0.70945 | 0.61373 | 0.80583 | 1.7717 | 0.43958 | 0.26577 | 0.51423 |
| 160 | Hypericum perforatum L. | Hypericum perforatum | 0.05 | 1 | 0.100 | 7.4 | 5.3 | NA | 6 | 7.7 | 0.8 | 0.65201 | 0.63791 | 1.54898 | 2.03286 | 0.77966 | 0.37419 | 0.45721 |
| 161 | Medicago lupulina L. | Medicago lupulina | 0.05 | 0.88 | 0.400 | 7.4 | 5.3 | NA | 6 | 7.7 | 0.8 | 0.65201 | 0.63791 | 1.54898 | 2.03286 | 0.77966 | 0.37419 | 0.45721 |
| 161 | Achillea millefolium L. | Achillea millefolium millefolium | 0.04 | 0.88 | 0.245 | 7.7 | NA | NA | 5.6 | NA | 0.7 | 0.53223 | 0.52325 | 1.73719 | 2.14421 | 0.94278 | 0.39374 | 0.35054 |
| 161 | Cerastium semidecandrum L. | Cerastium semidecandrum | 0.04 | 0.88 | 0.600 | 7.4 | 5.3 | NA | 6 | 7.7 | 0.8 | 0.65201 | 0.63791 | 1.54898 | 2.03286 | 0.77966 | 0.37419 | 0.45721 |
| 161 | Potentilla argentea L. | Potentilla argentea agg. | 0.05 | 0.88 | 0.150 | NA | 5.4 | 5.7 | 6.8 | 7 | 0 | 0.67742 | 0.24559 | 0.14546 | 1.36309 | 0.06547 | 0.19208 | 0.12644 |
| 161 | Hypericum perforatum L. | Hypericum perforatum | 0.04 | 0.88 | 0.202 | NA | NA | NA | NA | NA | NA | NA | NA | NA | NA | NA | NA | NA |
| 161 | Vicia hirsuta (L.) Gray | Vicia hirsuta | 0.02 | 0.88 | 0.408 | 7.1 | NA | 4.4 | 7 | 5.9 | 0.1 | 0.82845 | 0.8207 | 1.75144 | 1.9998 | 0.52148 | 0.15007 | 0.73628 |
| 161 | Euphorbia cyparissias L. | Euphorbia cyparissias | 0.15 | 0.88 | 0.084 | 8.1 | 6.2 | NA | 7.1 | 7.6 | 0.1 | 0.8111 | 0.79007 | 1.4109 | 1.89172 | 0.89447 | 0.17462 | 0.49967 |
| 161 | Plantago lanceolata L. | Plantago lanceolata | 0.04 | 0.88 | 0.333 | 7.2 | 5 | 5.1 | 6.1 | NA | 0.1 | 0.52316 | 0.5171 | 1.84298 | 2.10722 | 1.43318 | 0.13855 | 0.37874 |
| 161 | Poa annua L. | Poa annua | 0.2 | 0.88 | 0.204 | 7.2 | 5 | 5.1 | 6.1 | NA | 0.1 | 0.52316 | 0.5171 | 1.84298 | 2.10722 | 1.43318 | 0.13855 | 0.37874 |
| 161 | Silene latifolia Poir. | NA | 0.23 | 0.88 | 0.072 | 6.3 | 5 | 7.3 | 5.8 | 6.5 | 0.7 | 0.53692 | 0.48641 | 1.02734 | 1.80642 | 0.6097 | 0.33182 | 0.26598 |
| 161 | Lysimachia arvensis (L.) U.Manns & Anderb. | Anagallis arvensis | 0.02 | 0.88 | 0.290 | NA | NA | NA | NA | NA | NA | NA | NA | NA | NA | NA | NA | NA |
| 162 | Myosotis ramosissima Rochel | Myosotis ramosissima (coll. Hispida) | 0.01 | 1 | 0.038 | 7.9 | 5.1 | 5.1 | 6.7 | 7.7 | 0.2 | 0.75642 | 0.75467 | 2.08644 | 2.23861 | 0.83282 | 0.22413 | 0.56443 |
| 162 | Potentilla argentea L. | Potentilla argentea agg. | 0.2 | 1 | 0.333 | 7.2 | 5 | 5.1 | 6.1 | NA | 0.1 | 0.52316 | 0.5171 | 1.84298 | 2.10722 | 1.43318 | 0.13855 | 0.37874 |
| 162 | Achillea millefolium L. | Achillea millefolium millefolium | 0.05 | 1 | 0.054 | 7.7 | NA | NA | 6.4 | NA | 0 | 0.51083 | 0.48089 | 1.29849 | 1.93695 | 0.96262 | 0.19429 | 0.30238 |
| 162 | Plantago lanceolata L. | Plantago lanceolata | 0.1 | 1 | 0.250 | 7.4 | 5.3 | NA | 6 | 7.7 | 0.8 | 0.65201 | 0.63791 | 1.54898 | 2.03286 | 0.77966 | 0.37419 | 0.45721 |
| 162 | Bellis perennis L. | Bellis perennis | 0.05 | 1 | 0.050 | 7.4 | 5.3 | NA | 6 | 7.7 | 0.8 | 0.65201 | 0.63791 | 1.54898 | 2.03286 | 0.77966 | 0.37419 | 0.45721 |
| 162 | Poa annua L. | Poa annua | 0.24 | 1 | 0.556 | NA | NA | NA | NA | NA | NA | NA | NA | NA | NA | NA | NA | NA |
| 162 | Liquidambar styraciflua L. | NA | 0.3 | 1 | 0.050 | 6.7 | NA | NA | 6.5 | 7.7 | 0.1 | 0.70945 | 0.61373 | 0.80583 | 1.7717 | 0.43958 | 0.26577 | 0.51423 |
| 162 | Medicago lupulina L. | Medicago lupulina | 0.05 | 1 | 0.050 | 5.7 | NA | NA | 6.4 | 7.4 | 0 | 0.68859 | 0.26809 | 0.17316 | 1.34857 | 0.09545 | 0.19552 | 0.14349 |
| 163 | Medicago lupulina L. | Medicago lupulina | 0.04 | 0.96 | 0.105 | 6.7 | NA | NA | 6.5 | 7.7 | 0.1 | 0.70945 | 0.61373 | 0.80583 | 1.7717 | 0.43958 | 0.26577 | 0.51423 |
| 163 | Plantago lanceolata L. | Plantago lanceolata | 0.25 | 0.96 | 0.200 | 6.7 | NA | NA | 6.5 | 7.7 | 0.1 | 0.70945 | 0.61373 | 0.80583 | 1.7717 | 0.43958 | 0.26577 | 0.51423 |
| 163 | Potentilla argentea L. | Potentilla argentea agg. | 0.15 | 0.96 | 0.100 | NA | NA | NA | NA | NA | NA | NA | NA | NA | NA | NA | NA | NA |
| 163 | Rumex crispus L. | Rumex crispus | 0.15 | 0.96 | 0.200 | NA | NA | NA | NA | NA | NA | NA | NA | NA | NA | NA | NA | NA |
| 163 | Taraxacum officinale F.H.Wigg. | Taraxacum officinale agg. | 0.05 | 0.96 | 0.111 | NA | NA | NA | NA | NA | NA | NA | NA | NA | NA | NA | NA | NA |
| 163 | Veronica arvensis L. | Veronica arvensis | 0.02 | 0.96 | 0.100 | NA | NA | NA | NA | NA | NA | NA | NA | NA | NA | NA | NA | NA |
| 163 | Poa annua L. | Poa annua | 0.3 | 0.96 | 0.158 | 7 | NA | NA | 7.5 | 4.8 | 0 | 0.66959 | 0.18576 | 0.11569 | 1.33503 | 0.03304 | 0.20329 | 0.10324 |
| 164 | Medicago lupulina L. | Medicago lupulina | 0.08 | 1 | 0.100 | 7.3 | NA | NA | 5.8 | 5.9 | 0.5 | 0.63207 | 0.61738 | 1.52973 | 2.00883 | 0.70322 | 0.36696 | 0.39725 |
| 164 | Cerastium semidecandrum L. | Cerastium semidecandrum | 0.05 | 1 | 0.200 | 7.1 | 7.8 | 4.1 | 6.2 | 6 | 0.1 | 0.76637 | 0.65623 | 0.78647 | 1.82229 | 0.56568 | 0.18972 | 0.47454 |
| 164 | Taraxacum officinale F.H.Wigg. | Taraxacum officinale agg. | 0.05 | 1 | 0.600 | NA | NA | 5.3 | 6.2 | NA | 0.1 | 0.68682 | 0.14816 | 0.05813 | 1.31254 | 0.01433 | 0.20126 | 0.10505 |
| 164 | ene latifolia subsp. alba (Mill.) Greuter & Burd | Silene pratensis (alba) | 0.5 | 1 | 0.071 | 7.3 | 6.1 | 3.8 | 6 | 5.1 | 0 | 0.62987 | 0.59281 | 1.27995 | 1.90491 | 0.6275 | 0.3036 | 0.42608 |
| 164 | Poa annua L. | Poa annua | 0.32 | 1 | 0.105 | 7.3 | 6.1 | 3.8 | 6 | 5.1 | 0 | 0.62987 | 0.59281 | 1.27995 | 1.90491 | 0.6275 | 0.3036 | 0.42608 |
| 165 | Hypericum perforatum L. | Hypericum perforatum | 0.05 | 1 | 0.130 | 6.7 | NA | NA | 5.2 | NA | 0.4 | 0.59706 | 0.49429 | 0.69366 | 1.8006 | 0.482 | 0.19483 | 0.19138 |
| 165 | Medicago lupulina L. | Medicago lupulina | 0.1 | 1 | 0.077 | 7.1 | 4.9 | NA | 6 | NA | 1 | 0.59517 | 0.59064 | 1.89772 | 2.12267 | 0.65091 | 0.41213 | 0.37185 |
| 165 | Plantago lanceolata L. | Plantago lanceolata | 0.1 | 1 | 0.200 | 7.4 | 5.3 | NA | 6 | 7.7 | 0.8 | 0.65201 | 0.63791 | 1.54898 | 2.03286 | 0.77966 | 0.37419 | 0.45721 |
| 165 | Potentilla argentea L. | Potentilla argentea agg. | 0.05 | 1 | 0.200 | 7.1 | 4.9 | NA | 6 | NA | 1 | 0.59517 | 0.59064 | 1.89772 | 2.12267 | 0.65091 | 0.41213 | 0.37185 |
| 165 | Poa annua L. | Poa annua | 0.35 | 1 | 0.400 | 6.7 | NA | NA | 6.5 | 7.7 | 0.1 | 0.70945 | 0.61373 | 0.80583 | 1.7717 | 0.43958 | 0.26577 | 0.51423 |
| 165 | Bromus sterilis L. | Bromus sterilis | 0.35 | 1 | 0.211 | 7.3 | 5.4 | 4.1 | 7.6 | 4.2 | 0.4 | 0.58728 | 0.56055 | 1.34601 | 1.94534 | 0.82357 | 0.23743 | 0.34603 |
| 166 | Medicago lupulina L. | Medicago lupulina | 0.08 | 1 | 0.778 | 8.1 | NA | 3.9 | 6.9 | 6.4 | 0.4 | 0.80098 | 0.76713 | 1.29016 | 1.92909 | 0.75604 | 0.19458 | 0.52486 |
| 166 | Plantago lanceolata L. | Plantago lanceolata | 0.1 | 1 | 0.895 | 8.1 | NA | 3.9 | 6.9 | 6.4 | 0.4 | 0.80098 | 0.76713 | 1.29016 | 1.92909 | 0.75604 | 0.19458 | 0.52486 |
| 166 | Bellis perennis L. | Bellis perennis | 0.04 | 1 | 0.059 | 6.7 | NA | NA | 6.5 | 7.7 | 0.1 | 0.70945 | 0.61373 | 0.80583 | 1.7717 | 0.43958 | 0.26577 | 0.51423 |
| 166 | Poa annua L. | Poa annua | 0.68 | 1 | 0.050 | 6.7 | NA | NA | 6.5 | 7.7 | 0.1 | 0.70945 | 0.61373 | 0.80583 | 1.7717 | 0.43958 | 0.26577 | 0.51423 |
| 166 | Hypericum perforatum L. | Hypericum perforatum | 0.05 | 1 | 0.300 | 6.7 | NA | NA | 6.5 | 7.7 | 0.1 | 0.70945 | 0.61373 | 0.80583 | 1.7717 | 0.43958 | 0.26577 | 0.51423 |
| 166 | Lysimachia arvensis (L.) U.Manns & Anderb. | Anagallis arvensis | 0.02 | 1 | 0.200 | 7.4 | 5.3 | NA | 6 | 7.7 | 0.8 | 0.65201 | 0.63791 | 1.54898 | 2.03286 | 0.77966 | 0.37419 | 0.45721 |
| 166 | Veronica arvensis L. | Veronica arvensis | 0.03 | 1 | 0.267 | NA | NA | NA | NA | NA | NA | NA | NA | NA | NA | NA | NA | NA |
| 167 | Medicago lupulina L. | Medicago lupulina | 0.04 | 1 | 0.100 | NA | NA | 4.5 | 6.6 | 6.6 | 0 | 0.78059 | 0.60963 | 0.54418 | 1.63701 | 0.26733 | 0.16232 | 0.54354 |
| 167 | Achillea millefolium L. | Achillea millefolium millefolium | 0.15 | 1 | 0.100 | 6.8 | NA | 5.6 | 5.8 | NA | 0.1 | 0.52169 | 0.45097 | 0.93044 | 1.87015 | 0.67593 | 0.2975 | 0.27054 |
| 167 | Bromus sterilis L. | Bromus sterilis | 0.57 | 1 | 0.400 | 7.4 | 5.3 | NA | 6 | 7.7 | 0.8 | 0.65201 | 0.63791 | 1.54898 | 2.03286 | 0.77966 | 0.37419 | 0.45721 |
| 167 | Cerastium semidecandrum L. | Cerastium semidecandrum | 0.04 | 1 | 0.200 | 7.9 | 5.7 | 4.2 | 7 | 7.2 | 0.1 | 0.76998 | 0.7507 | 1.46055 | 1.90983 | 0.91556 | 0.1857 | 0.55131 |
| 167 | Potentilla reptans L. | Potentilla reptans | 0.05 | 1 | 0.200 | 7.7 | NA | NA | 5.6 | NA | 0.7 | 0.53223 | 0.52325 | 1.73719 | 2.14421 | 0.94278 | 0.39374 | 0.35054 |
| 167 | Plantago lanceolata L. | Plantago lanceolata | 0.05 | 1 | 0.050 | 6.3 | NA | 5.9 | 6.3 | 6.9 | 0.1 | 0.6037 | 0.3946 | 0.44368 | 1.50139 | 0.30164 | 0.21315 | 0.21494 |
| 167 | Liquidambar styraciflua L. | NA | 0.1 | 1 | 0.063 | 7.9 | 5.7 | 4.2 | 7 | 7.2 | 0.1 | 0.76998 | 0.7507 | 1.46055 | 1.90983 | 0.91556 | 0.1857 | 0.55131 |
| 168 | Medicago lupulina L. | Medicago lupulina | 0.02 | 0.81 | 0.150 | 7.9 | 5.7 | 4.2 | 7 | 7.2 | 0.1 | 0.76998 | 0.7507 | 1.46055 | 1.90983 | 0.91556 | 0.1857 | 0.55131 |
| 168 | Erodium cicutarium (L.) | Erodium cicutarium cicutarium | 0.1 | 0.81 | 0.120 | NA | NA | NA | NA | NA | NA | NA | NA | NA | NA | NA | NA | NA |
| 168 | Bellis perennis L. | Bellis perennis | 0.05 | 0.81 | 0.100 | 6.7 | NA | NA | 6.5 | 7.7 | 0.1 | 0.70945 | 0.61373 | 0.80583 | 1.7717 | 0.43958 | 0.26577 | 0.51423 |
| 168 | Potentilla argentea L. | Potentilla argentea agg. | 0.15 | 0.81 | 0.200 | NA | NA | NA | NA | NA | NA | NA | NA | NA | NA | NA | NA | NA |
| 168 | Plantago lanceolata L. | Plantago lanceolata | 0.1 | 0.81 | 0.050 | NA | NA | 4.5 | 6.6 | 6.6 | 0 | 0.78059 | 0.60963 | 0.54418 | 1.63701 | 0.26733 | 0.16232 | 0.54354 |
| 168 | Liquidambar styraciflua L. | NA | 0.05 | 0.81 | 0.400 | NA | NA | NA | NA | NA | NA | NA | NA | NA | NA | NA | NA | NA |
| 168 | Cerastium semidecandrum L. | Cerastium semidecandrum | 0.05 | 0.81 | 0.368 | 7.4 | 5.3 | NA | 6 | 7.7 | 0.8 | 0.65201 | 0.63791 | 1.54898 | 2.03286 | 0.77966 | 0.37419 | 0.45721 |
| 168 | Veronica arvensis L. | Veronica arvensis | 0.02 | 0.81 | 0.190 | 6.3 | NA | 5.9 | 6.3 | 6.9 | 0.1 | 0.6037 | 0.3946 | 0.44368 | 1.50139 | 0.30164 | 0.21315 | 0.21494 |
| 168 | Hypericum perforatum L. | Hypericum perforatum | 0.1 | 0.81 | 0.253 | 6.3 | NA | 5.9 | 6.3 | 6.9 | 0.1 | 0.6037 | 0.3946 | 0.44368 | 1.50139 | 0.30164 | 0.21315 | 0.21494 |
| 168 | Poa annua L. | Poa annua | 0.17 | 0.81 | 0.100 | 6.3 | NA | 5.9 | 6.3 | 6.9 | 0.1 | 0.6037 | 0.3946 | 0.44368 | 1.50139 | 0.30164 | 0.21315 | 0.21494 |
| 169 | Potentilla argentea L. | Potentilla argentea agg. | 0.17 | 0.85 | 0.200 | 6.3 | NA | 5.9 | 6.3 | 6.9 | 0.1 | 0.6037 | 0.3946 | 0.44368 | 1.50139 | 0.30164 | 0.21315 | 0.21494 |
| 169 | Medicago lupulina L. | Medicago lupulina | 0.04 | 0.85 | 0.150 | 7.2 | NA | NA | 6.5 | 8.4 | 0.1 | 0.5933 | 0.54722 | 1.04949 | 1.60983 | 0.64032 | 0.25162 | 0.31795 |
| 169 | Senecio inaequidens DC. | Senecio inaequidens | 0.04 | 0.85 | 0.300 | 7.4 | 5.3 | NA | 6 | 7.7 | 0.8 | 0.65201 | 0.63791 | 1.54898 | 2.03286 | 0.77966 | 0.37419 | 0.45721 |
| 169 | Plantago lanceolata L. | Plantago lanceolata | 0.1 | 0.85 | 0.100 | 7.3 | 5.4 | 4.1 | 7.6 | 4.2 | 0.4 | 0.58728 | 0.56055 | 1.34601 | 1.94534 | 0.82357 | 0.23743 | 0.34603 |
| 169 | Poa annua L. | Poa annua | 0.5 | 0.85 | 0.222 | 5.7 | NA | NA | 7.3 | 5.4 | 0 | 0.6918 | 0.13881 | 0.03984 | 1.31841 | 0.01173 | 0.19643 | 0.10363 |
| 170 | Potentilla argentea L. | Potentilla argentea agg. | 0.1 | 0.8 | 0.200 | 7.4 | 5.3 | NA | 6 | 7.7 | 0.8 | 0.65201 | 0.63791 | 1.54898 | 2.03286 | 0.77966 | 0.37419 | 0.45721 |
| 170 | Plantago lanceolata L. | Plantago lanceolata | 0.1 | 0.8 | 0.104 | 5.2 | 5.9 | 5.4 | 7.3 | 8 | 0 | 0.67947 | 0.25214 | 0.1495 | 1.33661 | 0.07498 | 0.19025 | 0.12819 |
| 170 | Hypericum perforatum L. | Hypericum perforatum | 0.05 | 0.8 | 0.204 | 7.4 | 5.3 | NA | 6 | 7.7 | 0.8 | 0.65201 | 0.63791 | 1.54898 | 2.03286 | 0.77966 | 0.37419 | 0.45721 |
| 170 | Medicago lupulina L. | Medicago lupulina | 0.04 | 0.8 | 0.380 | 8.3 | 5 | NA | 7.2 | 7.8 | 0.1 | 0.67936 | 0.56623 | 0.73548 | 1.48508 | 0.50389 | 0.1867 | 0.27262 |
| 170 | Senecio inaequidens DC. | Senecio inaequidens | 0.04 | 0.8 | 0.170 | 7.4 | 5.3 | NA | 6 | 7.7 | 0.8 | 0.65201 | 0.63791 | 1.54898 | 2.03286 | 0.77966 | 0.37419 | 0.45721 |
| 170 | Sedum acre L. | Sedum acre | 0.04 | 0.8 | 0.050 | 8.1 | NA | NA | 5.2 | NA | 0.1 | 0.51276 | 0.49581 | 1.51667 | 2.20989 | 0.98959 | 0.46997 | 0.3594 |

| | | | | | | | | | | | | | | | | | | |
|---|---|---|---|---|---|---|---|---|---|---|---|---|---|---|---|---|---|---|
| 170 | Lysimachia arvensis (L.) U.Manns & Anderb. | Anagallis arvensis | 0.04 | 0.8 | 0.100 | NA | NA | 4.5 | 6.6 | 6.6 | 0 | 0.78059 | 0.60963 | 0.54418 | 1.63701 | 0.26733 | 0.16232 | 0.54354 |
| 170 | Poa annua L. | Poa annua | 0.39 | 0.8 | 0.050 | 6.2 | 5.5 | NA | 6.9 | 8.5 | 0.1 | 0.60658 | 0.39025 | 0.44214 | 1.39281 | 0.24736 | 0.18526 | 0.18179 |
| 171 | Medicago lupulina L. | Medicago lupulina | 0.1 | 1 | 0.130 | 6.3 | NA | 5.9 | 6.3 | 6.9 | 0.1 | 0.6037 | 0.3946 | 0.44368 | 1.50139 | 0.30164 | 0.21315 | 0.21494 |
| 171 | Plantago lanceolata L. | Plantago lanceolata | 0.08 | 1 | 0.051 | 7 | 5.6 | 4.4 | 5.9 | 5.4 | 0.1 | 0.64871 | 0.63945 | 1.70759 | 2.05577 | 0.56937 | 0.21649 | 0.54803 |
| 171 | Potentilla argentea L. | Potentilla argentea agg. | 0.1 | 1 | 0.111 | 7 | 5.6 | 4.4 | 5.9 | 5.4 | 0.1 | 0.64871 | 0.63945 | 1.70759 | 2.05577 | 0.56937 | 0.21649 | 0.54803 |
| 171 | Hypericum perforatum L. | Hypericum perforatum | 0.08 | 1 | 0.050 | 6.9 | 5.9 | NA | 6.8 | 7 | 0 | 0.84157 | 0.83772 | 1.85474 | 2.04042 | 0.4856 | 0.14178 | 0.80173 |
| 171 | Euphorbia cyparissias L. | Euphorbia cyparissias | 0.25 | 1 | 0.300 | 6.3 | 5 | 7.3 | 5.8 | 6.5 | 0.7 | 0.53692 | 0.48641 | 1.02734 | 1.80642 | 0.6097 | 0.33182 | 0.26598 |
| 171 | Poa annua L. | Poa annua | 0.39 | 1 | 0.263 | 6.3 | 5 | 7.3 | 5.8 | 6.5 | 0.7 | 0.53692 | 0.48641 | 1.02734 | 1.80642 | 0.6097 | 0.33182 | 0.26598 |
| 172 | Potentilla argentea L. | Potentilla argentea agg. | 0.1 | 1 | 0.235 | NA | NA | NA | NA | NA | NA | NA | NA | NA | NA | NA | NA | NA |
| 172 | Medicago lupulina L. | Medicago lupulina | 0.05 | 1 | 0.250 | 7.3 | 5.4 | 4.1 | 7.6 | 4.2 | 0.4 | 0.58728 | 0.56055 | 1.34601 | 1.94534 | 0.82357 | 0.23743 | 0.34603 |
| 172 | Hypericum perforatum L. | Hypericum perforatum | 0.06 | 1 | 0.050 | 7.9 | 5.7 | 4.2 | 7 | 7.2 | 0.1 | 0.76998 | 0.7507 | 1.46055 | 1.90983 | 0.91556 | 0.1857 | 0.55131 |
| 172 | Senecio inaequidens DC. | Senecio inaequidens | 0.1 | 1 | 0.100 | 7.1 | 4.9 | NA | 6 | NA | 1 | 0.59517 | 0.59064 | 1.89772 | 2.12267 | 0.65091 | 0.41213 | 0.37185 |
| 172 | Poa annua L. | Poa annua | 0.55 | 1 | 0.050 | 7.9 | NA | 4.1 | 5.8 | NA | 0.9 | 0.54139 | 0.51128 | 1.31363 | 1.93633 | 0.8112 | 0.25085 | 0.32953 |
| 172 | Plantago lanceolata L. | Plantago lanceolata | 0.04 | 1 | 0.045 | 7 | 5.6 | 4.4 | 5.9 | 5.4 | 0.1 | 0.64871 | 0.63945 | 1.70759 | 2.05577 | 0.56937 | 0.21649 | 0.54803 |
| 172 | Liquidambar styraciflua L. | NA | 0.1 | 1 | 0.091 | 7.3 | 5.4 | 4.1 | 7.6 | 4.2 | 0.4 | 0.58728 | 0.56055 | 1.34601 | 1.94534 | 0.82357 | 0.23743 | 0.34603 |
| 173 | Cerastium glomeratum Thuill. | Cerastium glomeratum | 0.05 | 0.76 | 0.100 | 7.3 | 5.4 | 4.1 | 7.6 | 4.2 | 0.4 | 0.58728 | 0.56055 | 1.34601 | 1.94534 | 0.82357 | 0.23743 | 0.34603 |
| 173 | Bellis perennis L. | Bellis perennis | 0.05 | 0.76 | 0.200 | 7.4 | 5.3 | NA | 6 | 7.7 | 0.8 | 0.65201 | 0.63791 | 1.54898 | 2.03286 | 0.77966 | 0.37419 | 0.45721 |
| 173 | Potentilla argentea L. | Potentilla argentea agg. | 0.04 | 0.76 | 0.400 | 7.4 | 5.3 | NA | 6 | 7.7 | 0.8 | 0.65201 | 0.63791 | 1.54898 | 2.03286 | 0.77966 | 0.37419 | 0.45721 |
| 173 | Medicago lupulina L. | Medicago lupulina | 0.1 | 0.76 | 0.400 | 7.4 | 5.3 | NA | 6 | 7.7 | 0.8 | 0.65201 | 0.63791 | 1.54898 | 2.03286 | 0.77966 | 0.37419 | 0.45721 |
| 173 | Hypochaeris radicata L. | Hypochoeris radicata | 0.04 | 0.76 | 0.100 | 7.4 | 5.3 | NA | 6 | 7.7 | 0.8 | 0.65201 | 0.63791 | 1.54898 | 2.03286 | 0.77966 | 0.37419 | 0.45721 |
| 173 | Hypericum perforatum L. | Hypericum perforatum | 0.04 | 0.76 | 0.111 | 7.3 | 5.4 | 4.1 | 7.6 | 4.2 | 0.4 | 0.58728 | 0.56055 | 1.34601 | 1.94534 | 0.82357 | 0.23743 | 0.34603 |
| 173 | Plantago lanceolata L. | Plantago lanceolata | 0.05 | 0.76 | 0.100 | 7.3 | NA | NA | 7.6 | 7.7 | 0.4 | 0.75479 | 0.72524 | 1.28692 | 1.77374 | 0.61324 | 0.18283 | 0.56773 |
| 173 | Dactylis glomerata L. | Dactylis glomerata | 0.15 | 0.76 | 0.300 | 7.6 | NA | NA | 5.8 | NA | 0.5 | 0.55139 | 0.54271 | 1.75426 | 2.13166 | 0.95232 | 0.27891 | 0.39876 |
| 173 | Lysimachia arvensis (L.) U.Manns & Anderb. | Anagallis arvensis | 0.04 | 0.76 | 0.100 | 7.6 | NA | NA | 5.8 | NA | 0.5 | 0.55139 | 0.54271 | 1.75426 | 2.13166 | 0.95232 | 0.27891 | 0.39876 |
| 173 | Sedum acre L. | Sedum acre | 0.2 | 0.76 | 0.100 | 6.3 | NA | 5.9 | 6.3 | 6.9 | 0.1 | 0.6037 | 0.3946 | 0.44368 | 1.50139 | 0.30164 | 0.21315 | 0.21494 |
| 174 | Medicago lupulina L. | Medicago lupulina | 0.04 | 0.37 | 0.050 | 6.3 | NA | 5.9 | 6.3 | 6.9 | 0.1 | 0.6037 | 0.3946 | 0.44368 | 1.50139 | 0.30164 | 0.21315 | 0.21494 |
| 174 | Potentilla argentea L. | Potentilla argentea agg. | 0.04 | 0.37 | 0.100 | 7.3 | NA | NA | 5.8 | 5.9 | 0.5 | 0.63207 | 0.61738 | 1.52973 | 2.00883 | 0.70322 | 0.36696 | 0.39725 |
| 174 | Plantago lanceolata L. | Plantago lanceolata | 0.05 | 0.37 | 0.600 | 8.1 | NA | 3.9 | 6.9 | 6.4 | 0.4 | 0.80098 | 0.76713 | 1.29016 | 1.92909 | 0.75604 | 0.19458 | 0.52486 |
| 174 | Potentilla reptans L. | Potentilla reptans | 0.06 | 0.37 | 0.105 | 6.6 | 5.7 | NA | 6.3 | 7.6 | 0.1 | 0.67317 | 0.45023 | 0.43732 | 1.4267 | 0.22701 | 0.17304 | 0.27906 |
| 174 | Hypericum perforatum L. | Hypericum perforatum | 0.04 | 0.37 | 0.222 | 7.8 | NA | 4.6 | 6.2 | 5.8 | 0.1 | 0.71321 | 0.68276 | 1.29546 | 1.8275 | 0.62188 | 0.18605 | 0.48742 |
| 174 | Veronica arvensis L. | Veronica arvensis | 0.04 | 0.37 | 0.389 | 7 | 5.6 | 4.4 | 5.9 | 5.4 | 0.1 | 0.64871 | 0.63945 | 1.70759 | 2.05577 | 0.56937 | 0.21649 | 0.54803 |
| 174 | Poa annua L. | Poa annua | 0.1 | 0.37 | 0.294 | 7 | 5.6 | 4.4 | 5.9 | 5.4 | 0.1 | 0.64871 | 0.63945 | 1.70759 | 2.05577 | 0.56937 | 0.21649 | 0.54803 |
| 175 | Bellis perennis L. | Bellis perennis | 0.05 | 1 | 0.278 | 7 | 5.6 | 4.4 | 5.9 | 5.4 | 0.1 | 0.64871 | 0.63945 | 1.70759 | 2.05577 | 0.56937 | 0.21649 | 0.54803 |
| 175 | Plantago lanceolata L. | Plantago lanceolata | 0.09 | 1 | 0.010 | 7.4 | 5.2 | 4.7 | 6.1 | NA | 0 | 0.77406 | 0.76726 | 1.79113 | 2.07279 | 0.64778 | 0.2119 | 0.65925 |
| 175 | Potentilla reptans L. | Potentilla reptans | 0.18 | 1 | 0.550 | 8.1 | NA | 3.9 | 6.9 | 6.4 | 0.4 | 0.80098 | 0.76713 | 1.29016 | 1.92909 | 0.75604 | 0.19458 | 0.52486 |
| 175 | Hypochaeris radicata L. | Hypochoeris radicata | 0.09 | 1 | 0.153 | 7.1 | 4.9 | NA | 6 | NA | 1 | 0.59517 | 0.59064 | 1.89772 | 2.12267 | 0.65091 | 0.41213 | 0.37185 |
| 175 | Medicago lupulina L. | Medicago lupulina | 0.11 | 1 | 0.050 | NA | 5.4 | 5.7 | 6.8 | 7 | 0 | 0.67742 | 0.24559 | 0.14546 | 1.36309 | 0.06547 | 0.19208 | 0.12644 |
| 175 | Rumex acetosella L. | Rumex acetosella acetosella | 0.15 | 1 | 0.231 | 6.3 | 5 | 7.3 | 5.8 | 6.5 | 0.7 | 0.53692 | 0.48641 | 1.02734 | 1.80642 | 0.6097 | 0.33182 | 0.26598 |
| 175 | Hypericum perforatum L. | Hypericum perforatum | 0.19 | 1 | 0.074 | NA | NA | NA | 7 | NA | 0 | 0.66957 | 0.20824 | 0.12967 | 1.34554 | 0.07292 | 0.20428 | 0.13248 |
| 175 | Poa annua L. | Poa annua | 0.14 | 1 | 0.071 | NA | NA | NA | NA | NA | NA | NA | NA | NA | NA | NA | NA | NA |
| 176 | Plantago lanceolata L. | Plantago lanceolata | 0.1 | 0.95 | 0.349 | 8.4 | NA | 3 | 6 | NA | 0 | 0.46249 | 0.43915 | 1.39189 | 1.83687 | 0.38437 | 0.30571 | 0.38883 |
| 176 | Medicago lupulina L. | Medicago lupulina | 0.15 | 0.95 | 0.067 | 7.3 | 6.1 | 3.8 | 6 | 5.1 | 0 | 0.62987 | 0.59281 | 1.27995 | 1.90491 | 0.6275 | 0.3036 | 0.42608 |
| 176 | Potentilla argentea L. | Potentilla argentea agg. | 0.05 | 0.95 | 0.357 | 6.6 | NA | NA | 6.9 | 5.2 | 0.4 | 0.54634 | 0.5169 | 1.30759 | 1.85566 | 0.79671 | 0.26499 | 0.28358 |
| 176 | Bellis perennis L. | Bellis perennis | 0.02 | 0.95 | 0.042 | NA | NA | NA | NA | NA | NA | NA | NA | NA | NA | NA | NA | NA |
| 176 | Dactylis glomerata L. | Dactylis glomerata | 0.2 | 0.95 | 0.050 | 7.1 | 4.3 | NA | 5.1 | NA | 0.1 | 0.5318 | 0.50357 | 1.29074 | 2.04581 | 0.808 | 0.3645 | 0.36613 |
| 176 | Poa annua L. | Poa annua | 0.43 | 0.95 | 0.050 | 6.5 | 6.7 | 5.3 | 6.7 | 6.7 | 0 | 0.6884 | 0.39438 | 0.41506 | 1.39788 | 0.17506 | 0.19382 | 0.19382 |
| 177 | Hypericum perforatum L. | Hypericum perforatum | 0.05 | 1 | 0.080 | 7.1 | 4.9 | NA | 6 | NA | 1 | 0.59517 | 0.59064 | 1.89772 | 2.12267 | 0.65091 | 0.41213 | 0.37185 |
| 177 | Vicia hirsuta (L.) Gray | Vicia hirsuta | 0.15 | 1 | 0.263 | NA | NA | NA | NA | NA | NA | NA | NA | NA | NA | NA | NA | NA |
| 177 | Medicago lupulina L. | Medicago lupulina | 0.2 | 1 | 0.167 | 6.7 | NA | NA | 6.5 | 7.7 | 0.1 | 0.70945 | 0.61373 | 0.80583 | 1.7717 | 0.43958 | 0.26577 | 0.51423 |
| 177 | Senecio inaequidens DC. | Senecio inaequidens | 0.1 | 1 | 0.350 | 7.1 | 4.9 | NA | 6 | NA | 1 | 0.59517 | 0.59064 | 1.89772 | 2.12267 | 0.65091 | 0.41213 | 0.37185 |
| 177 | Bromus hordeaceus L. | Bromus hordeaceus (mollis) | 0.1 | 1 | 0.111 | 8.4 | NA | 4 | 6.3 | NA | 0.1 | 0.76525 | 0.73026 | 1.22636 | 1.78874 | 0.7034 | 0.16135 | 0.51807 |
| 177 | Poa annua L. | Poa annua | 0.4 | 1 | 0.100 | 7.1 | 4.9 | NA | 6 | NA | 1 | 0.59517 | 0.59064 | 1.89772 | 2.12267 | 0.65091 | 0.41213 | 0.37185 |
| 178 | Plantago lanceolata L. | Plantago lanceolata | 0.15 | 0.6 | 0.450 | 8.9 | 5.8 | 3 | 7.6 | 2.6 | 0.2 | 0.5898 | 0.52038 | 0.9391 | 1.66393 | 0.49914 | 0.25017 | 0.26404 |
| 178 | Medicago lupulina L. | Medicago lupulina | 0.1 | 0.6 | 0.020 | 7.3 | 5.4 | 4.1 | 7.6 | 4.2 | 0.4 | 0.58728 | 0.56055 | 1.34601 | 1.94534 | 0.82357 | 0.23743 | 0.34603 |
| 178 | Vicia hirsuta (L.) Gray | Vicia hirsuta | 0.05 | 0.6 | 0.051 | 7.3 | 5.4 | 4.1 | 7.6 | 4.2 | 0.4 | 0.58728 | 0.56055 | 1.34601 | 1.94534 | 0.82357 | 0.23743 | 0.34603 |
| 178 | Hypochaeris radicata L. | Hypochoeris radicata | 0.1 | 0.6 | 0.213 | 6.8 | 6.2 | 4.4 | NA | NA | 0.3 | 0.7871 | 0.75023 | 1.37944 | 1.92919 | 0.40631 | 0.19607 | 0.63681 |
| 178 | Potentilla reptans L. | Potentilla reptans | 0.05 | 0.6 | 0.300 | 6.7 | NA | NA | 6.5 | 7.7 | 0.1 | 0.70945 | 0.61373 | 0.80583 | 1.7717 | 0.43958 | 0.26577 | 0.51423 |
| 178 | Hypericum perforatum L. | Hypericum perforatum | 0.02 | 0.6 | 0.220 | 8.1 | NA | 3.9 | 6.9 | 6.4 | 0.4 | 0.80098 | 0.76713 | 1.29016 | 1.92909 | 0.75604 | 0.19458 | 0.52486 |
| 178 | Achillea millefolium L. | Achillea millefolium millefolium | 0.04 | 0.6 | 0.050 | 7.6 | NA | NA | 6.5 | 6.3 | 0.7 | 0.75548 | 0.75048 | 1.8409 | 2.08419 | 0.63746 | 0.23177 | 0.58814 |
| 178 | Poa annua L. | Poa annua | 0.09 | 0.6 | 0.193 | 7.1 | 5.3 | 4.5 | 5.9 | 4.8 | 0.3 | 0.52518 | 0.5113 | 1.61045 | 2.05509 | 0.96495 | 0.27897 | 0.32911 |
| 179 | Plantago lanceolata L. | Plantago lanceolata | 0.17 | 0.91 | 0.100 | 7.3 | 5.4 | 4.1 | 7.6 | 4.2 | 0.4 | 0.58728 | 0.56055 | 1.34601 | 1.94534 | 0.82357 | 0.23743 | 0.34603 |
| 179 | Medicago lupulina L. | Medicago lupulina | 0.15 | 0.91 | 0.150 | 6.9 | 6.5 | NA | 7.3 | 6.3 | 0 | 0.67274 | 0.19896 | 0.13039 | 1.32576 | 0.04466 | 0.20186 | 0.11616 |
| 179 | Potentilla argentea L. | Potentilla argentea agg. | 0.17 | 0.91 | 0.100 | 7.3 | 5.4 | 4.1 | 7.6 | 4.2 | 0.4 | 0.58728 | 0.56055 | 1.34601 | 1.94534 | 0.82357 | 0.23743 | 0.34603 |
| 179 | Poa annua L. | Poa annua | 0.4 | 0.91 | 0.170 | 6.6 | NA | NA | 6.9 | 5.2 | 0.4 | 0.54634 | 0.5169 | 1.30759 | 1.85566 | 0.79671 | 0.26499 | 0.28358 |
| 179 | Bellis perennis L. | Bellis perennis | 0.02 | 0.91 | 0.250 | 7.3 | 6.1 | 3.8 | 6 | 5.1 | 0 | 0.62987 | 0.59281 | 1.27995 | 1.90491 | 0.6275 | 0.3036 | 0.42608 |
| 180 | Plantago lanceolata L. | Plantago lanceolata | 0.15 | 0.96 | 0.150 | 6.6 | NA | NA | 6.9 | 5.2 | 0.4 | 0.54634 | 0.5169 | 1.30759 | 1.85566 | 0.79671 | 0.26499 | 0.28358 |
| 180 | Medicago lupulina L. | Medicago lupulina | 0.15 | 0.96 | 0.186 | 6.6 | NA | NA | 6.9 | 5.2 | 0.4 | 0.54634 | 0.5169 | 1.30759 | 1.85566 | 0.79671 | 0.26499 | 0.28358 |
| 180 | Potentilla argentea L. | Potentilla argentea agg. | 0.1 | 0.96 | 0.220 | 7.4 | 5.3 | NA | 6 | 7.7 | 0.8 | 0.65201 | 0.63791 | 1.54898 | 2.03286 | 0.77966 | 0.37419 | 0.45721 |
| 180 | Hypericum perforatum L. | Hypericum perforatum | 0.02 | 0.96 | 0.200 | 7.4 | 5.3 | NA | 6 | 7.7 | 0.8 | 0.65201 | 0.63791 | 1.54898 | 2.03286 | 0.77966 | 0.37419 | 0.45721 |
| 180 | Liquidambar styraciflua L. | NA | 0.05 | 0.96 | 0.330 | 7.2 | NA | NA | 6.7 | 7.8 | 0.1 | 0.79972 | 0.7689 | 1.29856 | 1.89634 | 0.65193 | 0.17352 | 0.67176 |
| 180 | Poa annua L. | Poa annua | 0.49 | 0.96 | 0.111 | 7.4 | 5.2 | 4.7 | 6.1 | NA | 0 | 0.77406 | 0.76726 | 1.79113 | 2.07279 | 0.64778 | 0.2119 | 0.65925 |
| 181 | Plantago lanceolata L. | Plantago lanceolata | 0.08 | 0.33 | 0.278 | 7.4 | 5.3 | NA | 6 | 7.7 | 0.8 | 0.65201 | 0.63791 | 1.54898 | 2.03286 | 0.77966 | 0.37419 | 0.45721 |
| 181 | Medicago lupulina L. | Medicago lupulina | 0.1 | 0.33 | 0.205 | 7 | 5.6 | 4.4 | 5.9 | 5.4 | 0.1 | 0.64871 | 0.63945 | 1.70759 | 2.05577 | 0.56937 | 0.21649 | 0.54803 |
| 181 | Achillea millefolium L. | Achillea millefolium millefolium | 0.1 | 0.33 | 0.460 | 7.4 | 5.3 | NA | 6 | 7.7 | 0.8 | 0.65201 | 0.63791 | 1.54898 | 2.03286 | 0.77966 | 0.37419 | 0.45721 |
| 181 | Bellis perennis L. | Bellis perennis | 0.01 | 0.33 | 0.040 | 7.1 | 4.9 | NA | 6 | NA | 1 | 0.59517 | 0.59064 | 1.89772 | 2.12267 | 0.65091 | 0.41213 | 0.37185 |
| 181 | Hypericum perforatum L. | Hypericum perforatum | 0.04 | 0.33 | 0.032 | 7 | 5.6 | 4.4 | 5.9 | 5.4 | 0.1 | 0.64871 | 0.63945 | 1.70759 | 2.05577 | 0.56937 | 0.21649 | 0.54803 |
| 182 | Plantago lanceolata L. | Plantago lanceolata | 0.17 | 1 | 0.091 | 7 | 5.6 | 4.4 | 5.9 | 5.4 | 0.1 | 0.64871 | 0.63945 | 1.70759 | 2.05577 | 0.56937 | 0.21649 | 0.54803 |
| 182 | Medicago lupulina L. | Medicago lupulina | 0.1 | 1 | 0.200 | 7 | 5.6 | 4.4 | 5.9 | 5.4 | 0.1 | 0.64871 | 0.63945 | 1.70759 | 2.05577 | 0.56937 | 0.21649 | 0.54803 |
| 182 | Poa annua L. | Poa annua | 0.6 | 1 | 0.200 | 6.5 | NA | NA | 5.4 | NA | 0 | 0.59769 | 0.45571 | 0.64594 | 1.51584 | 0.28825 | 0.22901 | 0.32316 |
| 182 | Bromus hordeaceus L. | Bromus hordeaceus (mollis) | 0.11 | 1 | 0.400 | 7.4 | 5.3 | NA | 6 | 7.7 | 0.8 | 0.65201 | 0.63791 | 1.54898 | 2.03286 | 0.77966 | 0.37419 | 0.45721 |
| 182 | Hypericum perforatum L. | Hypericum perforatum | 0.02 | 1 | 0.200 | 7.4 | 5.3 | NA | 6 | 7.7 | 0.8 | 0.65201 | 0.63791 | 1.54898 | 2.03286 | 0.77966 | 0.37419 | 0.45721 |
| 183 | Plantago lanceolata L. | Plantago lanceolata | 0.13 | 0.99 | 0.400 | 7.4 | 5.3 | NA | 6 | 7.7 | 0.8 | 0.65201 | 0.63791 | 1.54898 | 2.03286 | 0.77966 | 0.37419 | 0.45721 |
| 183 | Medicago lupulina L. | Medicago lupulina | 0.08 | 0.99 | 0.500 | 7 | 5.6 | 4.4 | 5.9 | 5.4 | 0.1 | 0.64871 | 0.63945 | 1.70759 | 2.05577 | 0.56937 | 0.21649 | 0.54803 |
| 183 | Potentilla argentea L. | Potentilla argentea agg. | 0.1 | 0.99 | 0.200 | 6.5 | NA | NA | 5.4 | NA | 0 | 0.59769 | 0.45571 | 0.64594 | 1.51584 | 0.28825 | 0.22901 | 0.32316 |
| 183 | Hypericum perforatum L. | Hypericum perforatum | 0.04 | 0.99 | 0.610 | NA | NA | NA | NA | NA | NA | NA | NA | NA | NA | NA | NA | NA |
| 183 | Veronica arvensis L. | Veronica arvensis | 0.02 | 0.99 | 0.647 | NA | NA | NA | NA | NA | NA | NA | NA | NA | NA | NA | NA | NA |
| 183 | Poa annua L. | Poa annua | 0.4 | 0.99 | 0.500 | NA | NA | NA | NA | NA | NA | NA | NA | NA | NA | NA | NA | NA |
| 183 | Bromus hordeaceus L. | Bromus hordeaceus (mollis) | 0.1 | 0.99 | 0.111 | 6.8 | NA | NA | 7.6 | 6.3 | 0.5 | 0.69759 | 0.6081 | 0.78594 | 1.30011 | 0.36881 | 0.19765 | 0.32063 |
| 183 | Dactylis glomerata L. | Dactylis glomerata | 0.08 | 0.99 | 0.667 | 7.1 | 4.9 | NA | 6 | NA | 1 | 0.59517 | 0.59064 | 1.89772 | 2.12267 | 0.65091 | 0.41213 | 0.37185 |
| 183 | Lolium perenne L. | Lolium perenne | 0.04 | 0.99 | 0.057 | NA | NA | NA | NA | NA | NA | NA | NA | NA | NA | NA | NA | NA |
| 184 | Plantago lanceolata L. | Plantago lanceolata | 0.08 | 1 | 0.350 | 8 | NA | NA | 6.1 | 6.3 | 0.1 | 0.56333 | 0.55394 | 1.72195 | 2.16326 | 1.07172 | 0.43309 | 0.38415 |
| 184 | Medicago lupulina L. | Medicago lupulina | 0.1 | 1 | 0.050 | NA | NA | NA | 6.6 | NA | 0 | 0.65326 | 0.6245 | 1.32492 | 1.96985 | 0.60133 | 0.19748 | 0.49592 |
| 184 | Hypochaeris radicata L. | Hypochoeris radicata | 0.05 | 1 | 0.200 | 6.2 | 5.5 | NA | 6.9 | 8.5 | 0.1 | 0.60658 | 0.39025 | 0.44214 | 1.39281 | 0.24736 | 0.18526 | 0.18179 |
| 184 | Potentilla argentea L. | Potentilla argentea agg. | 0.05 | 1 | 0.077 | 7.4 | 6.1 | 3.4 | 7.4 | 2.5 | 0.1 | 0.51564 | 0.45185 | 1.01243 | 1.75065 | 0.57143 | 0.28256 | 0.18911 |
| 184 | Hypericum perforatum L. | Hypericum perforatum | 0.04 | 1 | 0.111 | 7.7 | NA | NA | 6.7 | 7.2 | 0.1 | 0.5681 | 0.51951 | 1.12401 | 1.73868 | 0.64408 | 0.31575 | 0.32626 |
| 184 | Cerastium semidecandrum L. | Cerastium semidecandrum | 0.05 | 1 | 0.214 | 7.9 | 6.2 | NA | 6.7 | 6.4 | 0.2 | 0.62749 | 0.58201 | 1.11218 | 1.53794 | 0.69579 | 0.21134 | 0.22838 |
| 184 | Euphorbia cyparissias L. | Euphorbia cyparissias | 0.3 | 1 | 0.200 | 7.8 | 5.3 | 3.8 | 6.6 | 5 | 0.2 | 0.49579 | 0.46867 | 1.36306 | 1.85033 | 0.62791 | 0.29201 | 0.3217 |
| 184 | Poa annua L. | Poa annua | 0.29 | 1 | 0.050 | 7.2 | 5.7 | 4.3 | 6.8 | NA | 0.1 | 0.54405 | 0.41742 | 0.70503 | 1.63482 | 0.49706 | 0.18636 | 0.24515 |
| 184 | Taraxacum officinale F.H.Wigg. | Taraxacum officinale agg. | 0.04 | 1 | 0.056 | 7.1 | 4.3 | NA | 5.1 | NA | 0.1 | 0.5318 | 0.50357 | 1.29074 | 2.04581 | 0.808 | 0.3645 | 0.36613 |
| 185 | Medicago lupulina L. | Medicago lupulina | 0.05 | 1 | 0.020 | 7.1 | 4.9 | NA | 6 | NA | 1 | 0.59517 | 0.59064 | 1.89772 | 2.12267 | 0.65091 | 0.41213 | 0.37185 |
| 185 | Achillea millefolium L. | Achillea millefolium millefolium | 0.1 | 1 | 0.125 | 6.9 | 5.3 | 6.4 | 7.3 | 6.7 | 1 | 0.60499 | 0.3452 | 0.36505 | 1.40223 | 0.13192 | 0.19582 | 0.16678 |
| 185 | Taraxacum officinale F.H.Wigg. | Taraxacum officinale agg. | 0.15 | 1 | 0.056 | 8.1 | NA | NA | 5.2 | NA | 0.1 | 0.51276 | 0.49581 | 1.51667 | 2.20989 | 0.98959 | 0.46997 | 0.3594 |
| 185 | Potentilla argentea L. | Potentilla argentea agg. | 0.05 | 1 | 0.053 | 6.9 | 5.9 | NA | 6.8 | 7 | 0 | 0.84157 | 0.83772 | 1.85474 | 2.04042 | 0.4856 | 0.14178 | 0.80173 |
| 185 | Plantago lanceolata L. | Plantago lanceolata | 0.06 | 1 | 0.063 | 7.4 | 5.8 | 4.2 | 6.7 | 5.3 | 0 | 0.62702 | 0.59749 | 1.35516 | 1.96644 | 0.57704 | 0.23154 | 0.49642 |
| 185 | Echium vulgare L. | Echium vulgare | 0.05 | 1 | 0.778 | 7.4 | 5.3 | NA | 6 | 7.7 | 0.8 | 0.65201 | 0.63791 | 1.54898 | 2.03286 | 0.77966 | 0.37419 | 0.45721 |
| 185 | Poa annua L. | Poa annua | 0.54 | 1 | 0.033 | 6.9 | 5.9 | NA | 6.8 | 7 | 0 | 0.84157 | 0.83772 | 1.85474 | 2.04042 | 0.4856 | 0.14178 | 0.80173 |

| | | | | | | | | | | | | | | | | | | |
|---|---|---|---|---|---|---|---|---|---|---|---|---|---|---|---|---|---|---|
| 186 | Achillea millefolium L. | Achillea millefolium millefolium | 0.08 | 1 | 0.158 | NA | NA | NA | NA | NA | NA | NA | NA | NA | NA | NA | NA | NA |
| 186 | Plantago lanceolata L. | Plantago lanceolata | 0.1 | 1 | 0.118 | 8.6 | 6.3 | 2.6 | 7 | 1.9 | 0.1 | 0.4586 | 0.38628 | 0.91458 | 1.54485 | 0.13464 | 0.30586 | 0.2467 |
| 186 | Taraxacum officinale F.H.Wigg. | Taraxacum officinale agg. | 0.2 | 1 | 0.056 | NA | NA | 3.9 | 6.5 | 2.9 | 0 | 0.52109 | 0.46149 | 1.00552 | 1.92327 | 0.79082 | 0.28293 | 0.2827 |
| 186 | Potentilla argentea L. | Potentilla argentea agg. | 0.04 | 1 | 0.071 | 8.1 | NA | 3.5 | 7.5 | 5 | 0.3 | 0.75184 | 0.72654 | 1.3537 | 1.85024 | 0.717 | 0.18024 | 0.48387 |
| 186 | Bromus hordeaceus L. | Bromus hordeaceus (mollis) | 0.09 | 1 | 0.050 | NA | 5.4 | 5.7 | 6.8 | 7 | 0 | 0.67742 | 0.24559 | 0.14546 | 1.36309 | 0.06547 | 0.19208 | 0.12644 |
| 186 | Poa annua L. | Poa annua | 0.26 | 1 | 0.053 | 6.8 | 6.2 | 4.4 | NA | NA | 0.3 | 0.7871 | 0.75023 | 1.37944 | 1.92919 | 0.40631 | 0.19607 | 0.63681 |
| 186 | Bromus sterilis L. | Bromus sterilis | 0.23 | 1 | 0.250 | 7.4 | 5.8 | 4.2 | 6.7 | 5.3 | 0 | 0.62702 | 0.59749 | 1.35516 | 1.96644 | 0.57704 | 0.23154 | 0.49642 |
| 187 | Medicago lupulina L. | Medicago lupulina | 0.02 | 0.95 | 0.200 | 7.3 | NA | NA | 6.7 | 6.6 | 1 | 0.7393 | 0.71983 | 1.39902 | 1.79157 | 0.57605 | 0.19228 | 0.59493 |
| 187 | Achillea millefolium L. | Achillea millefolium millefolium | 0.08 | 0.95 | 0.050 | 8.2 | NA | 3.5 | 5.7 | 3.2 | 0 | 0.46142 | 0.41736 | 1.14878 | 1.78546 | 0.38405 | 0.27764 | 0.246 |
| 187 | Erodium cicutarium (L.) | Erodium cicutarium cicutarium | 0.01 | 0.95 | 0.240 | 8.2 | NA | 3.5 | 5.7 | 3.2 | 0 | 0.46142 | 0.41736 | 1.14878 | 1.78546 | 0.38405 | 0.27764 | 0.246 |
| 187 | Taraxacum officinale F.H.Wigg. | Taraxacum officinale agg. | 0.1 | 0.95 | 0.020 | 8.1 | NA | 3.5 | 6.1 | NA | 0.2 | 0.68175 | 0.6716 | 1.70824 | 1.99767 | 0.41848 | 0.25411 | 0.57318 |
| 187 | Potentilla argentea L. | Potentilla argentea agg. | 0.05 | 0.95 | 0.333 | 7.3 | 5.4 | 4.1 | 7.6 | 4.2 | 0.4 | 0.58728 | 0.56055 | 1.34601 | 1.94534 | 0.82357 | 0.23743 | 0.34603 |
| 187 | Bromus hordeaceus L. | Bromus hordeaceus (mollis) | 0.09 | 0.95 | 0.056 | 7.3 | 6.1 | 3.8 | 6 | 5.1 | 0 | 0.62987 | 0.59281 | 1.27995 | 1.90491 | 0.6275 | 0.3036 | 0.42608 |
| 187 | Poa annua L. | Poa annua | 0.3 | 0.95 | 0.215 | 7.3 | 6.1 | 3.8 | 6 | 5.1 | 0 | 0.62987 | 0.59281 | 1.27995 | 1.90491 | 0.6275 | 0.3036 | 0.42608 |
| 187 | Bromus sterilis L. | Bromus sterilis | 0.25 | 0.95 | 0.118 | 7.3 | 5.4 | 4.1 | 7.6 | 4.2 | 0.4 | 0.58728 | 0.56055 | 1.34601 | 1.94534 | 0.82357 | 0.23743 | 0.34603 |
| 187 | Echium vulgare L. | Echium vulgare | 0.05 | 0.95 | 0.054 | NA | 5.4 | 5.7 | 6.8 | 7 | 0 | 0.67742 | 0.24559 | 0.14546 | 1.36309 | 0.06547 | 0.19208 | 0.12644 |
| 188 | Medicago lupulina L. | Medicago lupulina | 0.02 | 0.98 | 0.046 | 8.2 | NA | 3.5 | 5.7 | 3.2 | 0 | 0.46142 | 0.41736 | 1.14878 | 1.78546 | 0.38405 | 0.27764 | 0.246 |
| 188 | Achillea millefolium L. | Achillea millefolium millefolium | 0.08 | 0.98 | 0.054 | 8.2 | NA | 3.5 | 5.7 | 3.2 | 0 | 0.46142 | 0.41736 | 1.14878 | 1.78546 | 0.38405 | 0.27764 | 0.246 |
| 188 | Taraxacum officinale F.H.Wigg. | Taraxacum officinale agg. | 0.15 | 0.98 | 0.075 | 8.2 | NA | 3.5 | 5.7 | 3.2 | 0 | 0.46142 | 0.41736 | 1.14878 | 1.78546 | 0.38405 | 0.27764 | 0.246 |
| 188 | Potentilla argentea L. | Potentilla argentea agg. | 0.02 | 0.98 | 0.294 | 7.7 | NA | NA | 5.6 | NA | 0.7 | 0.53223 | 0.52325 | 1.73719 | 2.14421 | 0.94278 | 0.39374 | 0.35054 |
| 188 | Plantago lanceolata L. | Plantago lanceolata | 0.05 | 0.98 | 0.050 | 8.2 | NA | 3.5 | 5.7 | 3.2 | 0 | 0.46142 | 0.41736 | 1.14878 | 1.78546 | 0.38405 | 0.27764 | 0.246 |
| 188 | Bellis perennis L. | Bellis perennis | 0.08 | 0.98 | 0.050 | NA | NA | NA | NA | NA | NA | NA | NA | NA | NA | NA | NA | NA |
| 188 | Veronica arvensis L. | Veronica arvensis | 0.02 | 0.98 | 0.222 | 7.9 | NA | 4.1 | 5.8 | NA | 0.9 | 0.54139 | 0.51128 | 1.31363 | 1.93633 | 0.8112 | 0.25085 | 0.32953 |
| 188 | Hypochaeris radicata L. | Hypochoeris radicata | 0.09 | 0.98 | 0.114 | NA | NA | NA | 6.6 | NA | 0 | 0.65326 | 0.6245 | 1.32492 | 1.96985 | 0.60133 | 0.19748 | 0.49592 |
| 188 | Poa annua L. | Poa annua | 0.3 | 0.98 | 0.050 | 7.4 | 5.3 | NA | 6 | 7.7 | 0.8 | 0.65201 | 0.63791 | 1.54898 | 2.03286 | 0.77966 | 0.37419 | 0.45721 |
| 188 | Bromus sterilis L. | Bromus sterilis | 0.13 | 0.98 | 0.022 | 7.3 | 5.4 | 4.1 | 7.6 | 4.2 | 0.4 | 0.58728 | 0.56055 | 1.34601 | 1.94534 | 0.82357 | 0.23743 | 0.34603 |
| 188 | Echium vulgare L. | Echium vulgare | 0.04 | 0.98 | 0.150 | 8.6 | NA | 2.7 | 4.4 | 1.6 | 0 | 0.53557 | 0.50485 | 1.29532 | 1.90211 | 0.72298 | 0.28495 | 0.2778 |
| 189 | Medicago lupulina L. | Medicago lupulina | 0.02 | 1 | 0.050 | 7.9 | NA | 4.1 | 5.8 | NA | 0.9 | 0.54139 | 0.51128 | 1.31363 | 1.93633 | 0.8112 | 0.25085 | 0.32953 |
| 189 | Achillea millefolium L. | Achillea millefolium millefolium | 0.1 | 1 | 0.100 | 7.9 | NA | 4.1 | 5.8 | NA | 0.9 | 0.54139 | 0.51128 | 1.31363 | 1.93633 | 0.8112 | 0.25085 | 0.32953 |
| 189 | Erodium cicutarium (L.) | Erodium cicutarium cicutarium | 0.02 | 1 | 0.080 | 8.1 | NA | 3.5 | 6.1 | NA | 0.2 | 0.68175 | 0.6716 | 1.70824 | 1.99767 | 0.41848 | 0.25411 | 0.57318 |
| 189 | Taraxacum officinale F.H.Wigg. | Taraxacum officinale agg. | 0.1 | 1 | 0.100 | 7.1 | 5.3 | 4.5 | 5.9 | 4.8 | 0.3 | 0.52518 | 0.5113 | 1.61045 | 2.05509 | 0.96495 | 0.27897 | 0.32911 |
| 189 | Potentilla argentea L. | Potentilla argentea agg. | 0.02 | 1 | 0.042 | 7.9 | NA | 4.1 | 5.8 | NA | 0.9 | 0.54139 | 0.51128 | 1.31363 | 1.93633 | 0.8112 | 0.25085 | 0.32953 |
| 189 | Hypochaeris radicata L. | Hypochoeris radicata | 0.05 | 1 | 0.300 | NA | NA | NA | NA | NA | NA | NA | NA | NA | NA | NA | NA | NA |
| 189 | Lactuca serriola L. | Lactuca serriola (scariola) | 0.05 | 1 | 0.050 | 8.5 | NA | 3.5 | 6.3 | 4.1 | 0.2 | 0.57633 | 0.55499 | 1.44489 | 1.81436 | 0.579 | 0.25616 | 0.27657 |
| 189 | Poa annua L. | Poa annua | 0.39 | 1 | 0.020 | 7 | 5.6 | 4.4 | 5.9 | 5.4 | 0.1 | 0.64871 | 0.63945 | 1.70759 | 2.05577 | 0.56937 | 0.21649 | 0.54803 |
| 189 | Bromus sterilis L. | Bromus sterilis | 0.15 | 1 | 0.040 | 6.7 | 5.3 | 4.5 | 5.3 | 5.8 | 0 | 0.75464 | 0.73051 | 1.36974 | 1.95508 | 0.45974 | 0.15567 | 0.6659 |
| 189 | Echium vulgare L. | Echium vulgare | 0.1 | 1 | 0.050 | 7.1 | 5.3 | 4.5 | 5.9 | 4.8 | 0.3 | 0.52518 | 0.5113 | 1.61045 | 2.05509 | 0.96495 | 0.27897 | 0.32911 |
| 190 | Medicago lupulina L. | Medicago lupulina | 0.1 | 1 | 0.100 | 7 | 5.6 | 4.4 | 5.9 | 5.4 | 0.1 | 0.64871 | 0.63945 | 1.70759 | 2.05577 | 0.56937 | 0.21649 | 0.54803 |
| 190 | Achillea millefolium L. | Achillea millefolium millefolium | 0.08 | 1 | 0.100 | 8.2 | NA | 2.1 | 6.6 | 1.7 | 0.4 | 0.4561 | 0.42853 | 1.29025 | 1.77233 | 0.32666 | 0.30641 | 0.32553 |
| 190 | Taraxacum officinale F.H.Wigg. | Taraxacum officinale agg. | 0.1 | 1 | 0.050 | 7 | 5.6 | 4.4 | 5.9 | 5.4 | 0.1 | 0.64871 | 0.63945 | 1.70759 | 2.05577 | 0.56937 | 0.21649 | 0.54803 |
| 190 | Plantago lanceolata L. | Plantago lanceolata | 0.04 | 1 | 0.020 | 7.6 | NA | NA | 6.5 | 6.3 | 0.7 | 0.75548 | 0.75048 | 1.8409 | 2.08419 | 0.63746 | 0.23177 | 0.58814 |
| 190 | Veronica arvensis L. | Veronica arvensis | 0.02 | 1 | 0.053 | 6.7 | 5.3 | 4.5 | 5.3 | 5.8 | 0 | 0.75464 | 0.73051 | 1.36974 | 1.95508 | 0.45974 | 0.15567 | 0.6659 |
| 190 | Hypochaeris radicata L. | Hypochoeris radicata | 0.08 | 1 | 0.050 | 6.7 | 5.3 | 4.5 | 5.3 | 5.8 | 0 | 0.75464 | 0.73051 | 1.36974 | 1.95508 | 0.45974 | 0.15567 | 0.6659 |
| 190 | Bellis perennis L. | Bellis perennis | 0.1 | 1 | 0.040 | 7 | 5.6 | 4.4 | 5.9 | 5.4 | 0.1 | 0.64871 | 0.63945 | 1.70759 | 2.05577 | 0.56937 | 0.21649 | 0.54803 |
| 190 | Potentilla argentea L. | Potentilla argentea agg. | 0.02 | 1 | 0.044 | 7.3 | 6.1 | 3.8 | 6 | 5.1 | 0 | 0.62987 | 0.59281 | 1.27995 | 1.90491 | 0.6275 | 0.3036 | 0.42608 |
| 190 | Poa annua L. | Poa annua | 0.3 | 1 | 0.167 | NA | NA | NA | NA | NA | NA | NA | NA | NA | NA | NA | NA | NA |
| 190 | Bromus sterilis L. | Bromus sterilis | 0.12 | 1 | 0.044 | 6.7 | NA | NA | 6.5 | 7.7 | 0.1 | 0.70945 | 0.61373 | 0.80583 | 1.7717 | 0.43958 | 0.26577 | 0.51423 |
| 190 | Echium vulgare L. | Echium vulgare | 0.04 | 1 | 0.200 | NA | NA | NA | NA | NA | NA | NA | NA | NA | NA | NA | NA | NA |
| 191 | Medicago lupulina L. | Medicago lupulina | 0.12 | 1 | 0.050 | 6.7 | NA | NA | 6.5 | 7.7 | 0.1 | 0.70945 | 0.61373 | 0.80583 | 1.7717 | 0.43958 | 0.26577 | 0.51423 |
| 191 | Achillea millefolium L. | Achillea millefolium millefolium | 0.05 | 1 | 0.100 | NA | NA | NA | NA | NA | NA | NA | NA | NA | NA | NA | NA | NA |
| 191 | Erodium cicutarium (L.) | Erodium cicutarium cicutarium | 0.02 | 1 | 0.750 | NA | NA | NA | NA | NA | NA | NA | NA | NA | NA | NA | NA | NA |
| 191 | Taraxacum officinale F.H.Wigg. | Taraxacum officinale agg. | 0.05 | 1 | 0.409 | 7.4 | 5.3 | NA | 6 | 7.7 | 0.8 | 0.65201 | 0.63791 | 1.54898 | 2.03286 | 0.77966 | 0.37419 | 0.45721 |
| 191 | Plantago lanceolata L. | Plantago lanceolata | 0.08 | 1 | 0.380 | 7.4 | 5.3 | NA | 6 | 7.7 | 0.8 | 0.65201 | 0.63791 | 1.54898 | 2.03286 | 0.77966 | 0.37419 | 0.45721 |
| 191 | Hypochaeris radicata L. | Hypochoeris radicata | 0.1 | 1 | 0.050 | NA | NA | NA | NA | NA | NA | NA | NA | NA | NA | NA | NA | NA |
| 191 | Potentilla argentea L. | Potentilla argentea agg. | 0.05 | 1 | 0.430 | 7.4 | 5.3 | NA | 6 | 7.7 | 0.8 | 0.65201 | 0.63791 | 1.54898 | 2.03286 | 0.77966 | 0.37419 | 0.45721 |
| 191 | Poa annua L. | Poa annua | 0.3 | 1 | 0.408 | 7.4 | 5.3 | NA | 6 | 7.7 | 0.8 | 0.65201 | 0.63791 | 1.54898 | 2.03286 | 0.77966 | 0.37419 | 0.45721 |
| 191 | Bromus sterilis L. | Bromus sterilis | 0.12 | 1 | 0.200 | 7.4 | 5.3 | NA | 6 | 7.7 | 0.8 | 0.65201 | 0.63791 | 1.54898 | 2.03286 | 0.77966 | 0.37419 | 0.45721 |
| 191 | Echium vulgare L. | Echium vulgare | 0.11 | 1 | 0.306 | 7.4 | 5.3 | NA | 6 | 7.7 | 0.8 | 0.65201 | 0.63791 | 1.54898 | 2.03286 | 0.77966 | 0.37419 | 0.45721 |
| 192 | Medicago lupulina L. | Medicago lupulina | 0.05 | 0.96 | 0.380 | 7.4 | 5.3 | NA | 6 | 7.7 | 0.8 | 0.65201 | 0.63791 | 1.54898 | 2.03286 | 0.77966 | 0.37419 | 0.45721 |
| 192 | Achillea millefolium L. | Achillea millefolium millefolium | 0.08 | 0.96 | 0.150 | 7.4 | 5.3 | NA | 6 | 7.7 | 0.8 | 0.65201 | 0.63791 | 1.54898 | 2.03286 | 0.77966 | 0.37419 | 0.45721 |
| 192 | Hypochaeris radicata L. | Hypochoeris radicata | 0.2 | 0.96 | 0.350 | 7.4 | 5.3 | NA | 6 | 7.7 | 0.8 | 0.65201 | 0.63791 | 1.54898 | 2.03286 | 0.77966 | 0.37419 | 0.45721 |
| 192 | Taraxacum officinale F.H.Wigg. | Taraxacum officinale agg. | 0.05 | 0.96 | 0.450 | 7.4 | 5.3 | NA | 6 | 7.7 | 0.8 | 0.65201 | 0.63791 | 1.54898 | 2.03286 | 0.77966 | 0.37419 | 0.45721 |
| 192 | Bellis perennis L. | Bellis perennis | 0.05 | 0.96 | 0.200 | 7.4 | 5.3 | NA | 6 | 7.7 | 0.8 | 0.65201 | 0.63791 | 1.54898 | 2.03286 | 0.77966 | 0.37419 | 0.45721 |
| 192 | Jacobaea analoga (DC.) Veldkamp | Senecio jacobaea | 0.05 | 0.96 | 0.200 | 8.1 | NA | NA | 4.4 | 2.8 | 0.3 | 0.48868 | 0.46508 | 1.4426 | 2.04965 | 0.7278 | 0.33931 | 0.35628 |
| 192 | Poa annua L. | Poa annua | 0.4 | 0.96 | 0.550 | 7.4 | 5.3 | NA | 6 | 7.7 | 0.8 | 0.65201 | 0.63791 | 1.54898 | 2.03286 | 0.77966 | 0.37419 | 0.45721 |
| 192 | Bromus sterilis L. | Bromus sterilis | 0.08 | 0.96 | 0.350 | 7.4 | 5.3 | NA | 6 | 7.7 | 0.8 | 0.65201 | 0.63791 | 1.54898 | 2.03286 | 0.77966 | 0.37419 | 0.45721 |
| 193 | Medicago lupulina L. | Medicago lupulina | 0.02 | 1 | 0.170 | 6.8 | NA | 5.6 | 5.8 | NA | 0.1 | 0.52169 | 0.45097 | 0.93044 | 1.87015 | 0.67593 | 0.2975 | 0.27054 |
| 193 | Achillea millefolium L. | Achillea millefolium millefolium | 0.08 | 1 | 0.050 | NA | NA | NA | 7 | NA | 0 | 0.66957 | 0.20824 | 0.12967 | 1.34554 | 0.07292 | 0.20428 | 0.13248 |
| 193 | Erodium cicutarium (L.) | Erodium cicutarium cicutarium | 0.05 | 1 | 0.172 | NA | NA | NA | 7 | NA | 0 | 0.66957 | 0.20824 | 0.12967 | 1.34554 | 0.07292 | 0.20428 | 0.13248 |
| 193 | Plantago lanceolata L. | Plantago lanceolata | 0.1 | 1 | 0.211 | 6.8 | NA | 5.6 | 5.8 | NA | 0.1 | 0.52169 | 0.45097 | 0.93044 | 1.87015 | 0.67593 | 0.2975 | 0.27054 |
| 193 | Herniaria glabra L. | Herniaria glabra | 0.02 | 1 | 0.050 | NA | NA | NA | 7 | NA | 0 | 0.66957 | 0.20824 | 0.12967 | 1.34554 | 0.07292 | 0.20428 | 0.13248 |
| 193 | Potentilla argentea L. | Potentilla argentea agg. | 0.05 | 1 | 0.050 | NA | NA | NA | 7 | NA | 0 | 0.66957 | 0.20824 | 0.12967 | 1.34554 | 0.07292 | 0.20428 | 0.13248 |
| 193 | Taraxacum officinale F.H.Wigg. | Taraxacum officinale agg. | 0.08 | 1 | 0.090 | NA | NA | NA | 7 | NA | 0 | 0.66957 | 0.20824 | 0.12967 | 1.34554 | 0.07292 | 0.20428 | 0.13248 |
| 193 | Poa annua L. | Poa annua | 0.32 | 1 | 0.040 | 7.3 | NA | 3.9 | 6.1 | 3.9 | 0.1 | 0.55903 | 0.41538 | 0.65189 | 1.65181 | 0.43719 | 0.24279 | 0.21709 |
| 193 | Bromus sterilis L. | Bromus sterilis | 0.2 | 1 | 0.230 | 6.7 | NA | 4.9 | 6.8 | 5 | 0 | 0.66057 | 0.18432 | 0.11528 | 1.34075 | 0.06671 | 0.20032 | 0.12881 |
| 193 | Echium vulgare L. | Echium vulgare | 0.08 | 1 | 0.069 | NA | NA | NA | 7 | NA | 0 | 0.66957 | 0.20824 | 0.12967 | 1.34554 | 0.07292 | 0.20428 | 0.13248 |
| 194 | Medicago lupulina L. | Medicago lupulina | 0.08 | 1 | 0.062 | 7.3 | 5.4 | 4.1 | 7.6 | 4.2 | 0.4 | 0.58728 | 0.56055 | 1.34601 | 1.94534 | 0.82357 | 0.23743 | 0.34603 |
| 194 | Achillea millefolium L. | Achillea millefolium millefolium | 0.1 | 1 | 0.115 | 6.8 | NA | 5.6 | 5.8 | NA | 0.1 | 0.52169 | 0.45097 | 0.93044 | 1.87015 | 0.67593 | 0.2975 | 0.27054 |
| 194 | Erodium cicutarium (L.) | Erodium cicutarium cicutarium | 0.05 | 1 | 0.050 | 8.1 | NA | NA | 5.2 | NA | 0.1 | 0.51276 | 0.49581 | 1.51667 | 2.20989 | 0.98959 | 0.46997 | 0.3594 |
| 194 | Hypochaeris radicata L. | Hypochoeris radicata | 0.1 | 1 | 0.053 | 8.6 | NA | 2.7 | 4.4 | 1.6 | 0 | 0.53557 | 0.50485 | 1.29532 | 1.90211 | 0.72298 | 0.28495 | 0.2778 |
| 194 | Taraxacum officinale F.H.Wigg. | Taraxacum officinale agg. | 0.15 | 1 | 0.235 | 7.4 | 5.3 | NA | 6 | 7.7 | 0.8 | 0.65201 | 0.63791 | 1.54898 | 2.03286 | 0.77966 | 0.37419 | 0.45721 |
| 194 | Plantago lanceolata L. | Plantago lanceolata | 0.04 | 1 | 0.450 | 7.4 | 5.3 | NA | 6 | 7.7 | 0.8 | 0.65201 | 0.63791 | 1.54898 | 2.03286 | 0.77966 | 0.37419 | 0.45721 |
| 194 | Potentilla argentea L. | Potentilla argentea agg. | 0.02 | 1 | 0.100 | 8.6 | 5.9 | 3.5 | NA | 2.5 | 0 | 0.5503 | 0.53805 | 1.6668 | 2.00922 | 0.50104 | 0.3154 | 0.3619 |
| 194 | Poa annua L. | Poa annua | 0.26 | 1 | 0.200 | NA | NA | NA | NA | NA | NA | NA | NA | NA | NA | NA | NA | NA |
| 194 | Bromus sterilis L. | Bromus sterilis | 0.2 | 1 | 0.050 | 7.1 | NA | NA | NA | 4.6 | 0 | 0.69216 | 0.66495 | 1.32487 | 1.98453 | 0.49303 | 0.16764 | 0.56484 |
| 195 | Medicago lupulina L. | Medicago lupulina | 0.06 | 0.95 | 0.080 | 7.1 | 5.3 | 4.5 | 5.9 | 4.8 | 0.3 | 0.52518 | 0.5113 | 1.61045 | 2.05509 | 0.96495 | 0.27897 | 0.32911 |
| 195 | Achillea millefolium L. | Achillea millefolium millefolium | 0.06 | 0.95 | 0.057 | 8.6 | NA | 2.7 | 4.4 | 1.6 | 0 | 0.53557 | 0.50485 | 1.29532 | 1.90211 | 0.72298 | 0.28495 | 0.2778 |
| 195 | Echium vulgare L. | Echium vulgare | 0.23 | 0.95 | 0.100 | 7.1 | 5.3 | 4.5 | 5.9 | 4.8 | 0.3 | 0.52518 | 0.5113 | 1.61045 | 2.05509 | 0.96495 | 0.27897 | 0.32911 |
| 195 | Erodium cicutarium (L.) | Erodium cicutarium cicutarium | 0.04 | 0.95 | 0.156 | 7.4 | NA | NA | 6 | 6.1 | 0.8 | 0.58355 | 0.57243 | 1.60436 | 1.92909 | 0.71129 | 0.29823 | 0.3555 |
| 195 | Taraxacum officinale F.H.Wigg. | Taraxacum officinale agg. | 0.08 | 0.95 | 0.500 | 7.7 | NA | 3.7 | 5.8 | 6.4 | 0 | 0.70633 | 0.61633 | 0.84477 | 1.62276 | 0.49804 | 0.1651 | 0.39904 |
| 195 | Veronica arvensis L. | Veronica arvensis | 0.05 | 0.95 | 0.050 | 8.6 | NA | 2.7 | 4.4 | 1.6 | 0 | 0.53557 | 0.50485 | 1.29532 | 1.90211 | 0.72298 | 0.28495 | 0.2778 |
| 195 | Poa annua L. | Poa annua | 0.2 | 0.95 | 0.680 | 7.4 | 5.3 | NA | 6 | 7.7 | 0.8 | 0.65201 | 0.63791 | 1.54898 | 2.03286 | 0.77966 | 0.37419 | 0.45721 |
| 195 | Bromus sterilis L. | Bromus sterilis | 0.23 | 0.95 | 0.040 | 8.4 | NA | 3 | 6 | NA | 0 | 0.46249 | 0.43915 | 1.39189 | 1.83687 | 0.38437 | 0.30571 | 0.38883 |
| 196 | Medicago lupulina L. | Medicago lupulina | 0.05 | 0.95 | 0.185 | 8.6 | NA | 2.7 | 4.4 | 1.6 | 0 | 0.53557 | 0.50485 | 1.29532 | 1.90211 | 0.72298 | 0.28495 | 0.2778 |
| 196 | Achillea millefolium L. | Achillea millefolium millefolium | 0.08 | 0.95 | 0.118 | 7.1 | 5.3 | 4.5 | 5.9 | 4.8 | 0.3 | 0.52518 | 0.5113 | 1.61045 | 2.05509 | 0.96495 | 0.27897 | 0.32911 |
| 196 | Taraxacum officinale F.H.Wigg. | Taraxacum officinale agg. | 0.1 | 0.95 | 0.050 | 7.3 | 5.4 | 4.1 | 7.6 | 4.2 | 0.4 | 0.58728 | 0.56055 | 1.34601 | 1.94534 | 0.82357 | 0.23743 | 0.34603 |
| 196 | Potentilla argentea L. | Potentilla argentea agg. | 0.04 | 0.95 | 0.080 | 7.3 | NA | 3.9 | 6.1 | 3.9 | 0.1 | 0.55903 | 0.41538 | 0.65189 | 1.65181 | 0.43719 | 0.24279 | 0.21709 |
| 196 | Echium vulgare L. | Echium vulgare | 0.15 | 0.95 | 0.100 | 8.4 | 7.2 | 2.6 | 6.8 | 3.6 | 0.2 | 0.58203 | 0.49726 | 0.89718 | 1.5951 | 0.3257 | 0.17939 | 0.36267 |
| 196 | Poa annua L. | Poa annua | 0.33 | 0.95 | 0.132 | 7.3 | 5.4 | 4.1 | 7.6 | 4.2 | 0.4 | 0.58728 | 0.56055 | 1.34601 | 1.94534 | 0.82357 | 0.23743 | 0.34603 |
| 196 | Bromus sterilis L. | Bromus sterilis | 0.2 | 0.95 | 0.135 | 7.1 | 5.3 | 4.5 | 5.9 | 4.8 | 0.3 | 0.52518 | 0.5113 | 1.61045 | 2.05509 | 0.96495 | 0.27897 | 0.32911 |
| 197 | Medicago lupulina L. | Medicago lupulina | 0.1 | 0.98 | 0.090 | 8.1 | NA | NA | 4.4 | 2.8 | 0.3 | 0.48868 | 0.46508 | 1.4426 | 2.04965 | 0.7278 | 0.33931 | 0.35628 |
| 197 | Achillea millefolium L. | Achillea millefolium millefolium | 0.08 | 0.98 | 0.021 | 8.1 | NA | NA | 5.2 | NA | 0.1 | 0.51276 | 0.49581 | 1.51667 | 2.20989 | 0.98959 | 0.46997 | 0.3594 |
| 197 | Echium vulgare L. | Echium vulgare | 0.05 | 0.98 | 0.100 | 8.4 | 7.2 | 2.6 | 6.8 | 3.6 | 0.2 | 0.58203 | 0.49726 | 0.89718 | 1.5951 | 0.3257 | 0.17939 | 0.36267 |
| 197 | Plantago lanceolata L. | Plantago lanceolata | 0.05 | 0.98 | 0.167 | 8.1 | NA | NA | 4.4 | 2.8 | 0.3 | 0.48868 | 0.46508 | 1.4426 | 2.04965 | 0.7278 | 0.33931 | 0.35628 |
| 197 | Hypochaeris radicata L. | Hypochoeris radicata | 0.05 | 0.98 | 0.021 | 7.3 | NA | 3.9 | 6.1 | 3.9 | 0.1 | 0.55903 | 0.41538 | 0.65189 | 1.65181 | 0.43719 | 0.24279 | 0.21709 |
| 197 | Poa annua L. | Poa annua | 0.3 | 0.98 | 0.030 | 8.1 | NA | NA | 5.2 | NA | 0.1 | 0.51276 | 0.49581 | 1.51667 | 2.20989 | 0.98959 | 0.46997 | 0.3594 |

| | | | | | | | | | | | | | | | | | | |
|---|---|---|---|---|---|---|---|---|---|---|---|---|---|---|---|---|---|---|
| 197 | Bromus sterilis L. | Bromus sterilis | 0.35 | 0.98 | 0.110 | 7.6 | NA | NA | 5.8 | NA | 0.5 | 0.55139 | 0.54271 | 1.75426 | 2.13166 | 0.95232 | 0.27891 | 0.39876 |
| 198 | Medicago lupulina L. | Medicago lupulina | 0.05 | 1 | 0.040 | 7.3 | NA | 3.9 | 6.1 | 3.9 | 0.1 | 0.55903 | 0.41538 | 0.65189 | 1.65181 | 0.43719 | 0.24279 | 0.21709 |
| 198 | Achillea millefolium L. | Achillea millefolium millefolium | 0.12 | 1 | 0.050 | 8.6 | NA | 2.7 | 4.4 | 1.6 | 0 | 0.53557 | 0.50485 | 1.29532 | 1.90211 | 0.72298 | 0.28495 | 0.2778 |
| 198 | Senecio inaequidens DC. | Senecio inaequidens | 0.05 | 1 | 0.050 | 8.6 | NA | 2.7 | 4.4 | 1.6 | 0 | 0.53557 | 0.50485 | 1.29532 | 1.90211 | 0.72298 | 0.28495 | 0.2778 |
| 198 | Potentilla argentea L. | Potentilla argentea agg. | 0.02 | 1 | 0.040 | 8.6 | NA | 2.7 | 4.4 | 1.6 | 0 | 0.53557 | 0.50485 | 1.29532 | 1.90211 | 0.72298 | 0.28495 | 0.2778 |
| 198 | Bromus hordeaceus L. | Bromus hordeaceus (mollis) | 0.1 | 1 | 0.105 | NA | NA | NA | NA | NA | NA | NA | NA | NA | NA | NA | NA | NA |
| 198 | Poa annua L. | Poa annua | 0.44 | 1 | 0.020 | 8.6 | NA | 2.7 | 4.4 | 1.6 | 0 | 0.53557 | 0.50485 | 1.29532 | 1.90211 | 0.72298 | 0.28495 | 0.2778 |
| 198 | Bromus sterilis L. | Bromus sterilis | 0.11 | 1 | 0.100 | NA | NA | NA | NA | NA | NA | NA | NA | NA | NA | NA | NA | NA |
| 198 | Bellis perennis L. | Bellis perennis | 0.11 | 1 | 0.040 | 7.1 | 5.3 | 4.5 | 5.9 | 4.8 | 0.3 | 0.52518 | 0.5113 | 1.61045 | 2.05509 | 0.96495 | 0.27897 | 0.32911 |
| 199 | Medicago lupulina L. | Medicago lupulina | 0.05 | 1 | 0.050 | NA | NA | NA | NA | NA | NA | NA | NA | NA | NA | NA | NA | NA |
| 199 | Achillea millefolium L. | Achillea millefolium millefolium | 0.08 | 1 | 0.052 | NA | NA | NA | NA | NA | NA | NA | NA | NA | NA | NA | NA | NA |
| 199 | Hypochaeris radicata L. | Hypochoeris radicata | 0.1 | 1 | 0.100 | 7.1 | 5.3 | 4.5 | 5.9 | 4.8 | 0.3 | 0.52518 | 0.5113 | 1.61045 | 2.05509 | 0.96495 | 0.27897 | 0.32911 |
| 199 | Plantago lanceolata L. | Plantago lanceolata | 0.08 | 1 | 0.100 | 8.1 | NA | NA | 4.4 | 2.8 | 0.3 | 0.48868 | 0.46508 | 1.4426 | 2.04965 | 0.7278 | 0.33931 | 0.35628 |
| 199 | Taraxacum officinale F.H.Wigg. | Taraxacum officinale agg. | 0.05 | 1 | 0.242 | 8.5 | NA | 3.5 | 6.3 | 4.1 | 0.2 | 0.57633 | 0.55499 | 1.44489 | 1.81436 | 0.579 | 0.25616 | 0.27657 |
| 199 | Poa annua L. | Poa annua | 0.37 | 1 | 0.020 | 8.6 | NA | 2.7 | 4.4 | 1.6 | 0 | 0.53557 | 0.50485 | 1.29532 | 1.90211 | 0.72298 | 0.28495 | 0.2778 |
| 199 | Bromus sterilis L. | Bromus sterilis | 0.15 | 1 | 0.080 | 7.1 | 5.3 | 4.5 | 5.9 | 4.8 | 0.3 | 0.52518 | 0.5113 | 1.61045 | 2.05509 | 0.96495 | 0.27897 | 0.32911 |
| 199 | Myosotis arvensis Hill | Myosotis arvensis (intermedia) | 0.05 | 1 | 0.080 | 8.1 | NA | NA | 4.4 | 2.8 | 0.3 | 0.48868 | 0.46508 | 1.4426 | 2.04965 | 0.7278 | 0.33931 | 0.35628 |
| 199 | Bellis perennis L. | Bellis perennis | 0.05 | 1 | 0.050 | 7.1 | NA | NA | NA | 4.6 | 0 | 0.69216 | 0.66495 | 1.32487 | 1.98453 | 0.49303 | 0.16764 | 0.56484 |
| 199 | Potentilla argentea L. | Potentilla argentea agg. | 0.02 | 1 | 0.050 | 8.1 | NA | NA | 5.2 | NA | 0.1 | 0.51276 | 0.49581 | 1.51667 | 2.20989 | 0.98959 | 0.46997 | 0.3594 |
| 200 | Medicago lupulina L. | Medicago lupulina | 0.05 | 1 | 0.100 | 7.3 | 5.4 | 4.1 | 7.6 | 4.2 | 0.4 | 0.58728 | 0.56055 | 1.34601 | 1.94534 | 0.82357 | 0.23743 | 0.34603 |
| 200 | Achillea millefolium L. | Achillea millefolium millefolium | 0.1 | 1 | 0.100 | NA | NA | NA | NA | NA | NA | NA | NA | NA | NA | NA | NA | NA |
| 200 | Taraxacum officinale F.H.Wigg. | Taraxacum officinale agg. | 0.1 | 1 | 0.430 | 7.4 | 5.3 | NA | 6 | 7.7 | 0.8 | 0.65201 | 0.63791 | 1.54898 | 2.03286 | 0.77966 | 0.37419 | 0.45721 |
| 200 | Hypochaeris radicata L. | Hypochoeris radicata | 0.08 | 1 | 0.250 | 7.9 | 5.7 | 4.2 | 7 | 7.2 | 0.1 | 0.76998 | 0.7507 | 1.46055 | 1.90983 | 0.91556 | 0.1857 | 0.55131 |
| 200 | Echium vulgare L. | Echium vulgare | 0.14 | 1 | 0.010 | NA | NA | NA | NA | NA | NA | NA | NA | NA | NA | NA | NA | NA |
| 200 | Potentilla argentea L. | Potentilla argentea agg. | 0.01 | 1 | 0.051 | NA | NA | 6.4 | 6.8 | 6.8 | 0 | 0.66551 | 0.27982 | 0.17419 | 1.41859 | 0.08796 | 0.19647 | 0.14036 |
| 200 | Poa annua L. | Poa annua | 0.27 | 1 | 0.010 | 7.1 | 7.8 | 4.1 | 6.2 | 6 | 0.1 | 0.76637 | 0.65623 | 0.78647 | 1.82229 | 0.56568 | 0.18972 | 0.47454 |
| 200 | Bromus sterilis L. | Bromus sterilis | 0.1 | 1 | 0.050 | 7.4 | NA | NA | 6 | 6.1 | 0.8 | 0.58355 | 0.57243 | 1.60436 | 1.92909 | 0.71129 | 0.29823 | 0.3555 |
| 200 | Bromus hordeaceus L. | Bromus hordeaceus (mollis) | 0.15 | 1 | 0.010 | NA | NA | NA | NA | NA | NA | NA | NA | NA | NA | NA | NA | NA |
| 201 | Medicago lupulina L. | Medicago lupulina | 0.02 | 1 | 0.306 | 6.7 | NA | NA | 6.5 | 7.7 | 0.1 | 0.70945 | 0.61373 | 0.80583 | 1.7717 | 0.43958 | 0.26577 | 0.51423 |
| 201 | Achillea millefolium L. | Achillea millefolium millefolium | 0.1 | 1 | 0.091 | 7.2 | NA | NA | 6.6 | 6.8 | 0 | 0.81299 | 0.76483 | 1.09209 | 1.84162 | 0.41616 | 0.14633 | 0.71443 |
| 201 | Echium vulgare L. | Echium vulgare | 0.1 | 1 | 0.133 | NA | NA | NA | NA | NA | NA | NA | NA | NA | NA | NA | NA | NA |
| 201 | Vicia hirsuta (L.) Gray | Vicia hirsuta | 0.05 | 1 | 0.017 | 7.9 | NA | 4.1 | 5.8 | NA | 0.9 | 0.54139 | 0.51128 | 1.31363 | 1.93633 | 0.8112 | 0.25085 | 0.32953 |
| 201 | Myosotis arvensis Hill | Myosotis arvensis (intermedia) | 0.05 | 1 | 0.102 | 7.9 | NA | 4.1 | 5.8 | NA | 0.9 | 0.54139 | 0.51128 | 1.31363 | 1.93633 | 0.8112 | 0.25085 | 0.32953 |
| 201 | Poa annua L. | Poa annua | 0.33 | 1 | 0.029 | 7.9 | NA | 4.1 | 5.8 | NA | 0.9 | 0.54139 | 0.51128 | 1.31363 | 1.93633 | 0.8112 | 0.25085 | 0.32953 |
| 201 | Bromus sterilis L. | Bromus sterilis | 0.15 | 1 | 0.118 | 7.9 | NA | 4.1 | 5.8 | NA | 0.9 | 0.54139 | 0.51128 | 1.31363 | 1.93633 | 0.8112 | 0.25085 | 0.32953 |
| 201 | Bromus hordeaceus L. | Bromus hordeaceus (mollis) | 0.2 | 1 | 0.101 | 7.1 | NA | 4.4 | 7 | 5.9 | 0.1 | 0.82845 | 0.8207 | 1.75144 | 1.9998 | 0.52148 | 0.15007 | 0.73628 |
| 202 | Medicago lupulina L. | Medicago lupulina | 0.08 | 1 | 0.096 | 7.4 | 5.3 | NA | 6 | 7.7 | 0.8 | 0.65201 | 0.63791 | 1.54898 | 2.03286 | 0.77966 | 0.37419 | 0.45721 |
| 202 | Achillea millefolium L. | Achillea millefolium millefolium | 0.05 | 1 | 0.054 | NA | NA | NA | NA | NA | NA | NA | NA | NA | NA | NA | NA | NA |
| 202 | Taraxacum officinale F.H.Wigg. | Taraxacum officinale agg. | 0.13 | 1 | 0.100 | 7.6 | NA | NA | 6.5 | 6.3 | 0.7 | 0.75548 | 0.75048 | 1.8409 | 2.08419 | 0.63746 | 0.23177 | 0.58814 |
| 202 | Bellis perennis L. | Bellis perennis | 0.05 | 1 | 0.280 | 7.4 | 5 | NA | NA | NA | 0.1 | 0.52506 | 0.51076 | 1.56059 | 2.06148 | 1.07287 | 0.38107 | 0.34662 |
| 202 | Plantago lanceolata L. | Plantago lanceolata | 0.08 | 1 | 0.050 | NA | NA | 6.4 | 6.8 | 6.8 | 0 | 0.66551 | 0.27982 | 0.17419 | 1.41859 | 0.08796 | 0.19647 | 0.14036 |
| 202 | Myosotis arvensis Hill | Myosotis arvensis (intermedia) | 0.05 | 1 | 0.050 | 7.6 | NA | NA | 6.5 | 6.3 | 0.7 | 0.75548 | 0.75048 | 1.8409 | 2.08419 | 0.63746 | 0.23177 | 0.58814 |
| 202 | Veronica arvensis L. | Veronica arvensis | 0.05 | 1 | 0.111 | 6.7 | NA | NA | 6.5 | 7.7 | 0.1 | 0.70945 | 0.61373 | 0.80583 | 1.7717 | 0.43958 | 0.26577 | 0.51423 |
| 202 | Poa annua L. | Poa annua | 0.3 | 1 | 0.421 | 8.1 | NA | 3.9 | 6.9 | 6.4 | 0.4 | 0.80098 | 0.76713 | 1.29016 | 1.92909 | 0.75604 | 0.19458 | 0.52486 |
| 202 | Bromus sterilis L. | Bromus sterilis | 0.21 | 1 | 0.556 | NA | NA | 4.5 | 6.6 | 6.6 | 0 | 0.78059 | 0.60963 | 0.54418 | 1.63701 | 0.26733 | 0.16232 | 0.54354 |
| 203 | Achillea millefolium L. | Achillea millefolium millefolium | 0.05 | 1 | 0.500 | 7.7 | NA | NA | 5.6 | NA | 0.7 | 0.53223 | 0.52325 | 1.73719 | 2.14421 | 0.94278 | 0.39374 | 0.35054 |
| 203 | Potentilla argentea L. | Potentilla argentea agg. | 0.03 | 1 | 0.050 | 5.7 | NA | NA | 6.4 | 7.4 | 0 | 0.68859 | 0.26809 | 0.17316 | 1.34857 | 0.09545 | 0.19552 | 0.14349 |
| 203 | Hypochaeris radicata L. | Hypochoeris radicata | 0.13 | 1 | 0.056 | 5.7 | NA | NA | 6.4 | 7.4 | 0 | 0.68859 | 0.26809 | 0.17316 | 1.34857 | 0.09545 | 0.19552 | 0.14349 |
| 203 | Medicago lupulina L. | Medicago lupulina | 0.1 | 1 | 0.050 | 7.1 | 4.9 | NA | 6 | NA | 1 | 0.59517 | 0.59064 | 1.89772 | 2.12267 | 0.65091 | 0.41213 | 0.37185 |
| 203 | Plantago lanceolata L. | Plantago lanceolata | 0.13 | 1 | 0.080 | 6.3 | 6 | NA | 7.6 | 8 | 0 | 0.69438 | 0.37984 | 0.32264 | 1.37302 | 0.2083 | 0.19481 | 0.20797 |
| 203 | Poa annua L. | Poa annua | 0.14 | 1 | 0.031 | 6.7 | NA | NA | 6.5 | 7.7 | 0.1 | 0.70945 | 0.61373 | 0.80583 | 1.7717 | 0.43958 | 0.26577 | 0.51423 |
| 203 | Bromus sterilis L. | Bromus sterilis | 0.17 | 1 | 0.084 | 6.7 | NA | NA | 6.5 | 7.7 | 0.1 | 0.70945 | 0.61373 | 0.80583 | 1.7717 | 0.43958 | 0.26577 | 0.51423 |
| 203 | Bromus hordeaceus L. | Bromus hordeaceus (mollis) | 0.13 | 1 | 0.050 | 7.1 | 4.9 | NA | 6 | NA | 1 | 0.59517 | 0.59064 | 1.89772 | 2.12267 | 0.65091 | 0.41213 | 0.37185 |
| 203 | Myosotis arvensis Hill | Myosotis arvensis (intermedia) | 0.02 | 1 | 0.100 | NA | NA | NA | NA | NA | NA | NA | NA | NA | NA | NA | NA | NA |
| 203 | Vicia hirsuta (L.) Gray | Vicia hirsuta | 0.05 | 1 | 0.200 | 7.6 | NA | NA | 6.5 | 6.3 | 0.7 | 0.75548 | 0.75048 | 1.8409 | 2.08419 | 0.63746 | 0.23177 | 0.58814 |
| 203 | Echium vulgare L. | Echium vulgare | 0.05 | 1 | 0.105 | NA | NA | NA | NA | NA | NA | NA | NA | NA | NA | NA | NA | NA |
| 204 | Achillea millefolium L. | Achillea millefolium millefolium | 0.08 | 1 | 0.118 | 7.4 | 5.2 | 4.7 | 6.1 | NA | 0 | 0.77406 | 0.76726 | 1.79113 | 2.07279 | 0.64778 | 0.2119 | 0.65925 |
| 204 | Veronica arvensis L. | Veronica arvensis | 0.02 | 1 | 0.133 | 6.7 | NA | NA | 6.5 | 7.7 | 0.1 | 0.70945 | 0.61373 | 0.80583 | 1.7717 | 0.43958 | 0.26577 | 0.51423 |
| 204 | Hypochaeris radicata L. | Hypochoeris radicata | 0.1 | 1 | 0.200 | 6.3 | 6 | NA | 7.6 | 8 | 0 | 0.69438 | 0.37984 | 0.32264 | 1.37302 | 0.2083 | 0.19481 | 0.20797 |
| 204 | Taraxacum officinale F.H.Wigg. | Taraxacum officinale agg. | 0.1 | 1 | 0.100 | 6.3 | 5 | 7.3 | 5.8 | 6.5 | 0.7 | 0.53692 | 0.48641 | 1.02734 | 1.80642 | 0.6097 | 0.33182 | 0.26598 |
| 204 | Poa annua L. | Poa annua | 0.2 | 1 | 0.050 | 8.1 | NA | NA | 5.2 | NA | 0.1 | 0.51276 | 0.49581 | 1.51667 | 2.20989 | 0.98959 | 0.46997 | 0.3594 |
| 204 | Bromus sterilis L. | Bromus sterilis | 0.2 | 1 | 0.100 | NA | NA | NA | NA | NA | NA | NA | NA | NA | NA | NA | NA | NA |
| 204 | Myosotis arvensis Hill | Myosotis arvensis (intermedia) | 0.05 | 1 | 0.100 | NA | NA | 4.5 | 6.6 | 6.6 | 0 | 0.78059 | 0.60963 | 0.54418 | 1.63701 | 0.26733 | 0.16232 | 0.54354 |
| 204 | Bromus hordeaceus L. | Bromus hordeaceus (mollis) | 0.25 | 1 | 0.250 | 6.2 | 5.5 | NA | 6.9 | 8.5 | 0.1 | 0.60658 | 0.39025 | 0.44214 | 1.39281 | 0.24736 | 0.18526 | 0.18179 |
| 205 | Achillea millefolium L. | Achillea millefolium millefolium | 0.04 | 1 | 0.063 | 7.2 | 5.5 | 4.5 | 6 | 7.6 | 0.3 | 0.72905 | 0.70188 | 1.35981 | 1.84158 | 0.57912 | 0.20081 | 0.60725 |
| 205 | Potentilla argentea L. | Potentilla argentea agg. | 0.08 | 1 | 0.150 | NA | NA | NA | NA | NA | NA | NA | NA | NA | NA | NA | NA | NA |
| 205 | Hypochaeris radicata L. | Hypochoeris radicata | 0.15 | 1 | 0.080 | 7.4 | 5.3 | NA | 6 | 7.7 | 0.8 | 0.65201 | 0.63791 | 1.54898 | 2.03286 | 0.77966 | 0.37419 | 0.45721 |
| 205 | Poa annua L. | Poa annua | 0.43 | 1 | 0.200 | NA | NA | NA | NA | NA | NA | NA | NA | NA | NA | NA | NA | NA |
| 205 | Bromus sterilis L. | Bromus sterilis | 0.3 | 1 | 0.160 | 7.4 | 5.3 | NA | 6 | 7.7 | 0.8 | 0.65201 | 0.63791 | 1.54898 | 2.03286 | 0.77966 | 0.37419 | 0.45721 |
| 206 | Taraxacum officinale F.H.Wigg. | Taraxacum officinale agg. | 0.3 | 0.7 | 0.150 | 6.7 | NA | NA | 6.5 | 7.7 | 0.1 | 0.70945 | 0.61373 | 0.80583 | 1.7717 | 0.43958 | 0.26577 | 0.51423 |
| 206 | Poa annua L. | Poa annua | 0.3 | 0.7 | 0.250 | 7.4 | 5.3 | NA | 6 | 7.7 | 0.8 | 0.65201 | 0.63791 | 1.54898 | 2.03286 | 0.77966 | 0.37419 | 0.45721 |
| 206 | Stellaria media (L.) Vill. | Stellaria media agg. | 0.1 | 0.7 | 0.050 | NA | NA | NA | NA | NA | NA | NA | NA | NA | NA | NA | NA | NA |
| 207 | Stellaria media (L.) Vill. | Stellaria media agg. | 0.4 | 1 | 0.071 | 6.5 | NA | NA | 5.4 | NA | 0 | 0.59769 | 0.45571 | 0.64594 | 1.51584 | 0.28825 | 0.22901 | 0.32316 |
| 207 | Taraxacum officinale F.H.Wigg. | Taraxacum officinale agg. | 0.25 | 1 | 0.020 | 6.9 | 6.5 | NA | 7.3 | 6.3 | 0 | 0.67274 | 0.19896 | 0.13039 | 1.32576 | 0.04466 | 0.20186 | 0.11616 |
| 207 | Poa annua L. | Poa annua | 0.1 | 1 | 0.300 | 7.4 | 5.3 | NA | 6 | 7.7 | 0.8 | 0.65201 | 0.63791 | 1.54898 | 2.03286 | 0.77966 | 0.37419 | 0.45721 |
| 207 | Sisymbrium officinale (L.) Scop. | Sisymbrium officinale | 0.25 | 1 | 0.250 | 7.4 | 5.3 | NA | 6 | 7.7 | 0.8 | 0.65201 | 0.63791 | 1.54898 | 2.03286 | 0.77966 | 0.37419 | 0.45721 |
| 207 | NA | NA | 0 | 1 | 0.111 | NA | NA | 6.4 | 6.8 | 6.8 | 0 | 0.66551 | 0.27982 | 0.17419 | 1.41859 | 0.08796 | 0.19647 | 0.14036 |
| 208 | Trifolium repens L. | Trifolium repens | 0.5 | 1 | 0.050 | NA | NA | NA | NA | NA | NA | NA | NA | NA | NA | NA | NA | NA |
| 208 | Glechoma hederacea L. | Glechoma hederacea | 0.01 | 1 | 0.083 | NA | NA | 4.5 | 6.6 | 6.6 | 0 | 0.78059 | 0.60963 | 0.54418 | 1.63701 | 0.26733 | 0.16232 | 0.54354 |
| 208 | Poa annua L. | Poa annua | 0.4 | 1 | 0.153 | NA | NA | 4.5 | 6.6 | 6.6 | 0 | 0.78059 | 0.60963 | 0.54418 | 1.63701 | 0.26733 | 0.16232 | 0.54354 |
| 208 | Taraxacum officinale F.H.Wigg. | Taraxacum officinale agg. | 0.01 | 1 | 0.100 | 7.6 | NA | NA | 6.5 | 6.3 | 0.7 | 0.75548 | 0.75048 | 1.8409 | 2.08419 | 0.63746 | 0.23177 | 0.58814 |
| 208 | Dactylis glomerata L. | Dactylis glomerata | 0.04 | 1 | 0.170 | 5.7 | NA | NA | 7.3 | 5.4 | 0 | 0.6918 | 0.13881 | 0.03984 | 1.31841 | 0.01173 | 0.19643 | 0.10363 |
| 208 | Bellis perennis L. | Bellis perennis | 0.04 | 1 | 0.300 | NA | 5.4 | 5.7 | 6.8 | 7 | 0 | 0.67742 | 0.24559 | 0.14546 | 1.36309 | 0.06547 | 0.19208 | 0.12644 |
| 208 | NA | NA | 0 | 1 | 0.050 | 6.2 | 5.5 | NA | 6.9 | 8.5 | 0.1 | 0.60658 | 0.39025 | 0.44214 | 1.39281 | 0.24736 | 0.18526 | 0.18179 |
| 209 | Glechoma hederacea L. | Glechoma hederacea | 0.3 | 0.98 | 0.310 | 6.3 | 5 | 7.3 | 5.8 | 6.5 | 0.7 | 0.53692 | 0.48641 | 1.02734 | 1.80642 | 0.6097 | 0.33182 | 0.26598 |
| 209 | Geum urbanum L. | Geum urbanum | 0.01 | 0.98 | 0.050 | NA | NA | 6.4 | 6.8 | 6.8 | 0 | 0.66551 | 0.27982 | 0.17419 | 1.41859 | 0.08796 | 0.19647 | 0.14036 |
| 209 | Trifolium repens L. | Trifolium repens | 0.24 | 0.98 | 0.051 | 5.7 | NA | NA | 7.3 | 5.4 | 0 | 0.6918 | 0.13881 | 0.03984 | 1.31841 | 0.01173 | 0.19643 | 0.10363 |
| 209 | Ranunculus ficaria L. | Ranunculus ficaria (Ficaria verna) | 0.05 | 0.98 | 0.111 | 6.5 | NA | NA | 5.4 | NA | 0 | 0.59769 | 0.45571 | 0.64594 | 1.51584 | 0.28825 | 0.22901 | 0.32316 |
| 209 | Alliaria petiolata (M.Bieb.) | Alliaria petiolata | 0.01 | 0.98 | 0.300 | 7.3 | NA | NA | 7.6 | 7.7 | 0.4 | 0.75479 | 0.72524 | 1.28692 | 1.77374 | 0.61324 | 0.18283 | 0.56773 |
| 209 | Prunella vulgaris L. | Prunella vulgaris | 0.1 | 0.98 | 0.100 | 7.7 | NA | NA | 5.6 | NA | 0.7 | 0.53223 | 0.52325 | 1.73719 | 2.14421 | 0.94278 | 0.39374 | 0.35054 |
| 209 | Poa annua L. | Poa annua | 0.24 | 0.98 | 0.211 | NA | 5.4 | 5.7 | 6.8 | 7 | 0 | 0.67742 | 0.24559 | 0.14546 | 1.36309 | 0.06547 | 0.19208 | 0.12644 |
| 209 | Bromus sterilis L. | Bromus sterilis | 0.01 | 0.98 | 0.176 | 7.6 | NA | NA | 5.8 | NA | 0.5 | 0.55139 | 0.54271 | 1.75426 | 2.13166 | 0.95232 | 0.27891 | 0.39876 |
| 209 | Veronica hederifolia L. | Veronica hederifolia agg. | 0.01 | 0.98 | 0.200 | 8.4 | NA | 3 | 6 | NA | 0 | 0.46249 | 0.43915 | 1.39189 | 1.83687 | 0.38437 | 0.30571 | 0.38883 |
| 209 | Veronica chamaedrys L. | Veronica chamaedrys | 0.01 | 0.98 | 0.350 | NA | NA | NA | NA | NA | NA | NA | NA | NA | NA | NA | NA | NA |
| 209 | NA | NA | 0 | 0.98 | 0.500 | 7.4 | 5.3 | NA | 6 | 7.7 | 0.8 | 0.65201 | 0.63791 | 1.54898 | 2.03286 | 0.77966 | 0.37419 | 0.45721 |
| 210 | Taraxacum officinale F.H.Wigg. | Taraxacum officinale agg. | 0.1 | 1 | 0.400 | 7.4 | 5.3 | NA | 6 | 7.7 | 0.8 | 0.65201 | 0.63791 | 1.54898 | 2.03286 | 0.77966 | 0.37419 | 0.45721 |
| 210 | Stellaria media (L.) Vill. | Stellaria media agg. | 0.2 | 1 | 0.091 | 8.1 | NA | NA | 4.4 | 2.8 | 0.3 | 0.48868 | 0.46508 | 1.4426 | 2.04965 | 0.7278 | 0.33931 | 0.35628 |
| 210 | Poa annua L. | Poa annua | 0.6 | 1 | 0.091 | 7.7 | NA | NA | 5.6 | NA | 0.7 | 0.53223 | 0.52325 | 1.73719 | 2.14421 | 0.94278 | 0.39374 | 0.35054 |
| 210 | Oxalis corniculata L. | Oxalis corniculata | 0.01 | 1 | 0.050 | 7.3 | 6.1 | 3.8 | 6 | 5.1 | 0 | 0.62987 | 0.59281 | 1.27995 | 1.90491 | 0.6275 | 0.3036 | 0.42608 |
| 210 | Veronica arvensis L. | Veronica arvensis | 0.02 | 1 | 0.050 | NA | NA | NA | 7 | NA | 0 | 0.66957 | 0.20824 | 0.12967 | 1.34554 | 0.07292 | 0.20428 | 0.13248 |
| 210 | Polygonum aviculare L. | Polygonum aviculare agg. | 0.02 | 1 | 0.118 | NA | NA | NA | NA | NA | NA | NA | NA | NA | NA | NA | NA | NA |
| 210 | Plantago major L. | Plantago major ssp. major | 0.05 | 1 | 0.250 | NA | NA | NA | NA | NA | NA | NA | NA | NA | NA | NA | NA | NA |
| 210 | NA | NA | 0 | 1 | 0.050 | 7 | 5.6 | 4.4 | 5.9 | 5.4 | 0.1 | 0.64871 | 0.63945 | 1.70759 | 2.05577 | 0.56937 | 0.21649 | 0.54803 |
| 211 | Poa annua L. | Poa annua | 0.15 | 1 | 0.167 | NA | NA | NA | NA | NA | NA | NA | NA | NA | NA | NA | NA | NA |
| 211 | Alliaria petiolata (M.Bieb.) | Alliaria petiolata | 0.46 | 1 | 0.050 | NA | NA | NA | NA | NA | NA | NA | NA | NA | NA | NA | NA | NA |
| 211 | Geum urbanum L. | Geum urbanum | 0.15 | 1 | 0.050 | NA | NA | NA | NA | NA | NA | NA | NA | NA | NA | NA | NA | NA |

| | | | | | | | | | | | | | | | | |
|---|---|---|---|---|---|---|---|---|---|---|---|---|---|---|---|---|
| 211 | Rumex crispus L. | Rumex crispus | 0.05 | 1 | 0.250 | 7.7 | NA | NA | 5.6 | NA | 0.7 | 0.53223 | 0.52325 | 1.73719 | 2.14421 | 0.94278 | 0.39374 | 0.35054 |
| 211 | Chaerophyllum temulum L. | Chaerophyllum temulum | 0.05 | 1 | 0.050 | 6.3 | 5 | 7.3 | 5.8 | 6.5 | 0.7 | 0.53692 | 0.48641 | 1.02734 | 1.80642 | 0.6097 | 0.33182 | 0.26598 |
| 211 | Bromus sterilis L. | Bromus sterilis | 0.02 | 1 | 0.100 | 7.1 | 5.3 | 4.5 | 5.9 | 4.8 | 0.3 | 0.52518 | 0.5113 | 1.61045 | 2.05509 | 0.96495 | 0.27897 | 0.32911 |
| 211 | Galium aparine L. | Galium aparine aparine | 0.05 | 1 | 0.050 | 7.4 | 5.2 | 4.7 | 6.1 | NA | 0 | 0.77406 | 0.76726 | 1.79113 | 2.07279 | 0.64778 | 0.2119 | 0.65925 |
| 211 | Veronica hederifolia L. | Veronica hederifolia agg. | 0.01 | 1 | 0.053 | 7.8 | NA | 4.6 | 6.2 | 5.8 | 0.1 | 0.71321 | 0.68276 | 1.29546 | 1.8275 | 0.62188 | 0.18605 | 0.48742 |
| 211 | Taraxacum officinale F.H.Wigg. | Taraxacum officinale agg. | 0.01 | 1 | 0.056 | 7.1 | 4.9 | NA | 6 | NA | 1 | 0.59517 | 0.59064 | 1.89772 | 2.12267 | 0.65091 | 0.41213 | 0.37185 |
| 211 | ı galeobdolon subsp. argentatum (Smejkal) J. | NA | 0.05 | 1 | 0.059 | 7.1 | 4.9 | NA | 6 | NA | 1 | 0.59517 | 0.59064 | 1.89772 | 2.12267 | 0.65091 | 0.41213 | 0.37185 |
| 211 | NA | NA | 0 | 1 | 0.167 | 7.1 | 4.9 | NA | 6 | NA | 1 | 0.59517 | 0.59064 | 1.89772 | 2.12267 | 0.65091 | 0.41213 | 0.37185 |
| 212 | Alliaria petiolata (M.Bieb.) | Alliaria petiolata | 0.45 | 0.99 | 0.300 | 8 | NA | NA | 6.1 | 6.3 | 0.1 | 0.56333 | 0.55394 | 1.72195 | 2.16326 | 1.07172 | 0.43309 | 0.38415 |
| 212 | Geum urbanum L. | Geum urbanum | 0.2 | 0.99 | 0.110 | 8.1 | NA | NA | 4.4 | 2.8 | 0.3 | 0.48868 | 0.46508 | 1.4426 | 2.04965 | 0.7278 | 0.33931 | 0.35628 |
| 212 | Bromus sterilis L. | Bromus sterilis | 0.2 | 0.99 | 0.205 | 8.1 | NA | NA | 4.4 | 2.8 | 0.3 | 0.48868 | 0.46508 | 1.4426 | 2.04965 | 0.7278 | 0.33931 | 0.35628 |
| 212 | Taraxacum officinale F.H.Wigg. | Taraxacum officinale agg. | 0.01 | 0.99 | 0.118 | NA | NA | NA | NA | NA | NA | NA | NA | NA | NA | NA | NA | NA |
| 212 | Veronica hederifolia L. | Veronica hederifolia agg. | 0.01 | 0.99 | 0.161 | 7 | 5.6 | 4.4 | 5.9 | 5.4 | 0.1 | 0.64871 | 0.63945 | 1.70759 | 2.05577 | 0.56937 | 0.21649 | 0.54803 |
| 212 | Urtica dioica L. | Urtica dioica | 0.05 | 0.99 | 0.093 | NA | NA | NA | 7 | NA | 0 | 0.66957 | 0.20824 | 0.12967 | 1.34554 | 0.07292 | 0.20428 | 0.13248 |
| 212 | Poa annua L. | Poa annua | 0.01 | 0.99 | 0.200 | NA | NA | NA | 7 | NA | 0 | 0.66957 | 0.20824 | 0.12967 | 1.34554 | 0.07292 | 0.20428 | 0.13248 |
| 212 | Galium aparine L. | Galium aparine aparine | 0.01 | 0.99 | 0.020 | NA | NA | NA | NA | NA | NA | NA | NA | NA | NA | NA | NA | NA |
| 212 | Glechoma hederacea L. | Glechoma hederacea | 0.05 | 0.99 | 0.042 | 7.3 | 6.1 | 3.8 | 6 | 5.1 | 0 | 0.62987 | 0.59281 | 1.27995 | 1.90491 | 0.6275 | 0.3036 | 0.42608 |
| 212 | NA | NA | 0 | 0.99 | 0.200 | NA | NA | NA | NA | NA | NA | NA | NA | NA | NA | NA | NA | NA |
| 213 | Taraxacum officinale F.H.Wigg. | Taraxacum officinale agg. | 0.02 | 0.98 | 0.200 | 7.8 | NA | 4.6 | 6.2 | 5.8 | 0.1 | 0.71321 | 0.68276 | 1.29546 | 1.8275 | 0.62188 | 0.18605 | 0.48742 |
| 213 | Senecio vulgaris L. | Senecio vulgaris | 0.01 | 0.98 | 0.263 | NA | 5.4 | 5.7 | 6.8 | 7 | 0 | 0.67742 | 0.24559 | 0.14546 | 1.36309 | 0.06547 | 0.19208 | 0.12644 |
| 213 | Papaver rhoeas L. | Papaver rhoeas | 0.4 | 0.98 | 0.056 | 6.9 | 6.3 | 3.9 | 6.3 | 7.1 | 0.1 | 0.77916 | 0.67616 | 0.95204 | 1.78267 | 0.43197 | 0.25047 | 0.50688 |
| 213 | Stellaria media (L.) Vill. | Stellaria media agg. | 0.3 | 0.98 | 0.200 | 8.4 | NA | 4 | 6.3 | NA | 0.1 | 0.76525 | 0.73026 | 1.22636 | 1.78874 | 0.7034 | 0.16135 | 0.51807 |
| 213 | Herniaria glabra L. | Herniaria glabra | 0.2 | 0.98 | 0.040 | 7.1 | NA | NA | NA | 4.6 | 0 | 0.69216 | 0.66495 | 1.32487 | 1.98453 | 0.49303 | 0.16764 | 0.56484 |
| 213 | Sagina procumbens L. | Sagina procumbens | 0.02 | 0.98 | 0.082 | 7.3 | 6.2 | 7.3 | 5.3 | 7 | 0 | 0.64252 | 0.51787 | 0.68487 | 1.47944 | 0.40522 | 0.22979 | 0.19935 |
| 213 | Plantago major L. | Plantago major ssp. major | 0.02 | 0.98 | 0.085 | 8.1 | NA | NA | 4.4 | 2.8 | 0.3 | 0.48868 | 0.46508 | 1.4426 | 2.04965 | 0.7278 | 0.33931 | 0.35628 |
| 213 | Poa annua L. | Poa annua | 0.01 | 0.98 | 0.100 | NA | NA | NA | 7 | NA | 0 | 0.66957 | 0.20824 | 0.12967 | 1.34554 | 0.07292 | 0.20428 | 0.13248 |
| 214 | Taraxacum officinale F.H.Wigg. | Taraxacum officinale agg. | 0.9 | 0.9 | 0.090 | 7.6 | NA | NA | 6.5 | 6.3 | 0.7 | 0.75548 | 0.75048 | 1.8409 | 2.08419 | 0.63746 | 0.23177 | 0.58814 |
| 216 | Lactuca serriola L. | Lactuca serriola (scariola) | 0.21 | 0.33 | 0.550 | 7 | NA | NA | NA | NA | 0.3 | 0.56899 | 0.4288 | 0.64817 | 1.63031 | 0.49671 | 0.22558 | 0.25973 |
| 216 | Lamium purpureum L. | Lamium purpureum | 0.03 | 0.33 | 0.148 | NA | 5.4 | 5.7 | 6.8 | 7 | 0 | 0.67742 | 0.24559 | 0.14546 | 1.36309 | 0.06547 | 0.19208 | 0.12644 |
| 216 | Malva sylvestris L. | Malva sylvestris | 0.07 | 0.33 | 0.211 | 7.9 | NA | 4.1 | 5.8 | NA | 0.9 | 0.54139 | 0.51128 | 1.31363 | 1.93633 | 0.8112 | 0.25085 | 0.32953 |
| 216 | Calendula officinalis L. | NA | 0.02 | 0.33 | 0.170 | 7.3 | 6.1 | 3.8 | 6 | 5.1 | 0 | 0.62987 | 0.59281 | 1.27995 | 1.90491 | 0.6275 | 0.3036 | 0.42608 |
| 217 | Taraxacum officinale F.H.Wigg. | Taraxacum officinale agg. | 0.11 | 0.83 | 0.050 | 7.1 | 5.3 | 4.5 | 5.9 | 4.8 | 0.3 | 0.52518 | 0.5113 | 1.61045 | 2.05509 | 0.96495 | 0.27897 | 0.32911 |
| 217 | Achillea millefolium L. | Achillea millefolium millefolium | 0.13 | 0.83 | 0.050 | 7.3 | 5.4 | 4.1 | 7.6 | 4.2 | 0.4 | 0.58728 | 0.56055 | 1.34601 | 1.94534 | 0.82357 | 0.23743 | 0.34603 |
| 217 | Malva sylvestris L. | Malva sylvestris | 0.07 | 0.83 | 0.100 | 7.7 | NA | NA | 5.6 | NA | 0.7 | 0.53223 | 0.52325 | 1.73719 | 2.14421 | 0.94278 | 0.39374 | 0.35054 |
| 217 | Centaurea cyanus L. | Centaurea cyanus | 0.11 | 0.83 | 0.103 | 7.7 | NA | NA | 5.6 | NA | 0.7 | 0.53223 | 0.52325 | 1.73719 | 2.14421 | 0.94278 | 0.39374 | 0.35054 |
| 217 | Knautia arvensis (L.) Coult. | Knautia arvensis | 0.07 | 0.83 | 0.040 | 8.1 | NA | NA | 5.2 | NA | 0.1 | 0.51276 | 0.49581 | 1.51667 | 2.20989 | 0.98959 | 0.46997 | 0.3594 |
| 217 | Stachys sylvatica L. | Stachys sylvatica | 0.05 | 0.83 | 0.210 | 8.1 | NA | NA | 5.2 | NA | 0.1 | 0.51276 | 0.49581 | 1.51667 | 2.20989 | 0.98959 | 0.46997 | 0.3594 |
| 217 | Hypericum perforatum L. | Hypericum perforatum | 0.05 | 0.83 | 0.044 | 8.1 | NA | 3.5 | 6.1 | NA | 0.2 | 0.68175 | 0.6716 | 1.70824 | 1.99767 | 0.41848 | 0.25411 | 0.57318 |
| 217 | Cirsium vulgare (Savi) Ten. | Cirsium vulgare (lanceolatum) | 0.05 | 0.83 | 0.068 | 6.7 | NA | NA | 6.5 | 7.7 | 0.1 | 0.70945 | 0.61373 | 0.80583 | 1.7717 | 0.43958 | 0.26577 | 0.51423 |
| 217 | Artemisia vulgaris L. | Artemisia vulgaris | 0.05 | 0.83 | 0.455 | NA | NA | NA | 7 | NA | 0 | 0.66957 | 0.20824 | 0.12967 | 1.34554 | 0.07292 | 0.20428 | 0.13248 |
| 217 | Leucanthemum vulgare Lam. | Leucanthemum vulgare (leucanthem.) | 0.02 | 0.83 | 0.040 | 7.1 | 4.9 | NA | 6 | NA | 1 | 0.59517 | 0.59064 | 1.89772 | 2.12267 | 0.65091 | 0.41213 | 0.37185 |
| 217 | Cynosurus cristatus L. | Cynosurus cristatus | 0.02 | 0.83 | 0.320 | NA | NA | NA | NA | NA | NA | NA | NA | NA | NA | NA | NA | NA |
| 217 | Lepidium draba L. | Cardaria draba | 0.05 | 0.83 | 0.323 | NA | NA | NA | 7 | NA | 0 | 0.66957 | 0.20824 | 0.12967 | 1.34554 | 0.07292 | 0.20428 | 0.13248 |
| 217 | Lotus corniculatus L. | Lotus corniculatus | 0.05 | 0.83 | 0.167 | 6.5 | NA | NA | 5.4 | NA | 0 | 0.59769 | 0.45571 | 0.64594 | 1.51584 | 0.28825 | 0.22901 | 0.32316 |
| 218 | Taraxacum officinale F.H.Wigg. | Taraxacum officinale agg. | 0.05 | 0.6 | 0.200 | 7 | 5.6 | 4.4 | 5.9 | 5.4 | 0.1 | 0.64871 | 0.63945 | 1.70759 | 2.05577 | 0.56937 | 0.21649 | 0.54803 |
| 218 | Crepis biennis L. | Crepis biennis | 0.2 | 0.6 | 0.100 | 7 | 5.6 | 4.4 | 5.9 | 5.4 | 0.1 | 0.64871 | 0.63945 | 1.70759 | 2.05577 | 0.56937 | 0.21649 | 0.54803 |
| 218 | Achillea millefolium L. | Achillea millefolium millefolium | 0.01 | 0.6 | 0.050 | 7 | 5.6 | 4.4 | 5.9 | 5.4 | 0.1 | 0.64871 | 0.63945 | 1.70759 | 2.05577 | 0.56937 | 0.21649 | 0.54803 |
| 218 | Malva sylvestris L. | Malva sylvestris | 0.07 | 0.6 | 0.200 | 7.1 | 4.9 | NA | 6 | NA | 1 | 0.59517 | 0.59064 | 1.89772 | 2.12267 | 0.65091 | 0.41213 | 0.37185 |
| 218 | Hypochaeris radicata L. | Hypochoeris radicata | 0.05 | 0.6 | 0.200 | 7.1 | 4.9 | NA | 6 | NA | 1 | 0.59517 | 0.59064 | 1.89772 | 2.12267 | 0.65091 | 0.41213 | 0.37185 |
| 218 | Plantago lanceolata L. | Plantago lanceolata | 0.05 | 0.6 | 0.083 | 6.5 | NA | NA | 5.4 | NA | 0 | 0.59769 | 0.45571 | 0.64594 | 1.51584 | 0.28825 | 0.22901 | 0.32316 |
| 218 | Echium vulgare L. | Echium vulgare | 0.1 | 0.6 | 0.057 | 8.1 | NA | 3.5 | 7.5 | 5 | 0.3 | 0.75184 | 0.72654 | 1.3537 | 1.85024 | 0.717 | 0.18024 | 0.48387 |
| 218 | Myosotis arvensis Hill | Myosotis arvensis (intermedia) | 0.05 | 0.6 | 0.235 | 7.2 | NA | NA | 5.5 | 4.7 | 0.5 | 0.51067 | 0.4744 | 1.23473 | 1.90522 | 0.81709 | 0.28699 | 0.29055 |
| 218 | Poa annua L. | Poa annua | 0.02 | 0.6 | 0.250 | 7.9 | 6.2 | NA | 6.7 | 6.4 | 0.2 | 0.62749 | 0.58201 | 1.11218 | 1.53794 | 0.69579 | 0.21134 | 0.22838 |
| 219 | Taraxacum officinale F.H.Wigg. | Taraxacum officinale agg. | 0.1 | 0.49 | 0.050 | 6.9 | 5.9 | NA | 6.8 | 7 | 0 | 0.84157 | 0.83772 | 1.85474 | 2.04042 | 0.4856 | 0.14178 | 0.80173 |
| 219 | Crepis biennis L. | Crepis biennis | 0.1 | 0.49 | 0.154 | 7.3 | NA | NA | 6.5 | 2.7 | 0.3 | 0.50155 | 0.46101 | 1.19593 | 1.8416 | 0.70249 | 0.27259 | 0.24586 |
| 219 | Achillea millefolium L. | Achillea millefolium millefolium | 0.05 | 0.49 | 0.071 | 7.1 | NA | NA | 6.9 | 5.8 | 0.1 | 0.65728 | 0.63749 | 1.47109 | 1.96875 | 0.7311 | 0.22775 | 0.51018 |
| 219 | Myosotis arvensis Hill | Myosotis arvensis (intermedia) | 0.05 | 0.49 | 0.056 | 8.5 | NA | 2.5 | NA | NA | 0 | 0.506 | 0.44835 | 1.02121 | 1.67002 | 0.34316 | 0.25824 | 0.34742 |
| 219 | Ranunculus repens L. | Ranunculus repens | 0.02 | 0.49 | 0.050 | 7.8 | 5.3 | 3.8 | 6.6 | 5 | 0.2 | 0.49579 | 0.46867 | 1.36306 | 1.85033 | 0.62791 | 0.29201 | 0.3217 |
| 219 | Medicago lupulina L. | Medicago lupulina | 0.05 | 0.49 | 0.063 | NA | NA | NA | NA | NA | NA | NA | NA | NA | NA | NA | NA | NA |
| 219 | Leucanthemum vulgare Lam. | Leucanthemum vulgare (leucanthem.) | 0.1 | 0.49 | 0.111 | 6.8 | 6.2 | 4.4 | NA | NA | 0.3 | 0.7871 | 0.75023 | 1.37944 | 1.92919 | 0.40631 | 0.19607 | 0.63681 |
| 219 | Silene dioica (L.) Clairv. | Silene dioica (Melandrium rubrum) | 0.02 | 0.49 | 0.025 | 6.6 | 5.7 | NA | 6.3 | 7.6 | 0.1 | 0.67317 | 0.45023 | 0.43732 | 1.4267 | 0.22701 | 0.17304 | 0.27906 |
| 220 | Taraxacum officinale F.H.Wigg. | Taraxacum officinale agg. | 0.1 | 0.69 | 0.817 | 7.4 | 5.3 | NA | 6 | 7.7 | 0.8 | 0.65201 | 0.63791 | 1.54898 | 2.03286 | 0.77966 | 0.37419 | 0.45721 |
| 220 | Poa annua L. | Poa annua | 0.3 | 0.69 | 0.111 | 7.8 | 6.3 | NA | 7.3 | 5.3 | 0.2 | 0.5821 | 0.55876 | 1.40231 | 1.83895 | 0.97897 | 0.15853 | 0.34647 |
| 220 | Ranunculus repens L. | Ranunculus repens | 0.05 | 0.69 | 0.100 | 7.7 | NA | NA | 6.7 | 7.2 | 0.1 | 0.5681 | 0.51951 | 1.12401 | 1.73868 | 0.64408 | 0.31575 | 0.32626 |
| 220 | Achillea millefolium L. | Achillea millefolium millefolium | 0.02 | 0.69 | 0.368 | NA | NA | 4.5 | 6.6 | 6.6 | 0 | 0.78059 | 0.60963 | 0.54418 | 1.63701 | 0.26733 | 0.16232 | 0.54354 |
| 220 | Leucanthemum vulgare Lam. | Leucanthemum vulgare (leucanthem.) | 0.1 | 0.69 | 0.200 | 8.1 | NA | NA | 4.4 | 2.8 | 0.3 | 0.48868 | 0.46508 | 1.4426 | 2.04965 | 0.7278 | 0.33931 | 0.35628 |
| 220 | Crepis biennis L. | Crepis biennis | 0.1 | 0.69 | 0.100 | 7.3 | 5.4 | 4.1 | 7.6 | 4.2 | 0.4 | 0.58728 | 0.56055 | 1.34601 | 1.94534 | 0.82357 | 0.23743 | 0.34603 |
| 220 | Tragopogon pratensis L. | Tragopogon pratensis pratensis | 0.02 | 0.69 | 0.050 | 7.3 | 6.1 | 3.8 | 6 | 5.1 | 0 | 0.62987 | 0.59281 | 1.27995 | 1.90491 | 0.6275 | 0.3036 | 0.42608 |
| 221 | Crepis biennis L. | Crepis biennis | 0.56 | 0.85 | 0.100 | 7.3 | 5.4 | 4.1 | 7.6 | 4.2 | 0.4 | 0.58728 | 0.56055 | 1.34601 | 1.94534 | 0.82357 | 0.23743 | 0.34603 |
| 221 | Achillea millefolium L. | Achillea millefolium millefolium | 0.1 | 0.85 | 0.222 | 8.3 | 2.5 | 3.7 | 3.3 | 2 | 0 | 0.36542 | 0.32774 | 1.12918 | 1.79068 | 0.11557 | 0.28384 | 0.11777 |
| 221 | Poa annua L. | Poa annua | 0.07 | 0.85 | 0.054 | 8.2 | NA | 3.5 | 5.7 | 3.2 | 0 | 0.46142 | 0.41736 | 1.14878 | 1.78546 | 0.38405 | 0.27764 | 0.246 |
| 221 | Tragopogon pratensis L. | Tragopogon pratensis pratensis | 0.07 | 0.85 | 0.059 | 8.5 | NA | 2.5 | NA | NA | 0 | 0.506 | 0.44835 | 1.02121 | 1.67002 | 0.34316 | 0.25824 | 0.34742 |
| 221 | Lepidium draba L. | Cardaria draba | 0.05 | 0.85 | 0.022 | 8.5 | NA | 2.5 | NA | NA | 0 | 0.506 | 0.44835 | 1.02121 | 1.67002 | 0.34316 | 0.25824 | 0.34742 |
| 222 | Lepidium draba L. | Cardaria draba | 0.07 | 0.69 | 0.057 | 7.1 | 4.3 | NA | 5.1 | NA | 0.1 | 0.5318 | 0.50357 | 1.29074 | 2.04581 | 0.808 | 0.3645 | 0.36613 |
| 222 | Taraxacum officinale F.H.Wigg. | Taraxacum officinale agg. | 0.2 | 0.69 | 0.087 | 7.1 | 4.3 | NA | 5.1 | NA | 0.1 | 0.5318 | 0.50357 | 1.29074 | 2.04581 | 0.808 | 0.3645 | 0.36613 |
| 222 | Papaver rhoeas L. | Papaver rhoeas | 0.07 | 0.69 | 0.075 | 7.1 | 4.3 | NA | 5.1 | NA | 0.1 | 0.5318 | 0.50357 | 1.29074 | 2.04581 | 0.808 | 0.3645 | 0.36613 |
| 222 | Geranium pusillum L. | Geranium pusillum | 0.05 | 0.69 | 0.049 | 7.3 | 6.1 | 3.8 | 6 | 5.1 | 0 | 0.62987 | 0.59281 | 1.27995 | 1.90491 | 0.6275 | 0.3036 | 0.42608 |
| 222 | Sinapis arvensis L. | Sinapis arvensis | 0.3 | 0.69 | 0.250 | 6.7 | NA | NA | 5.2 | NA | 0.4 | 0.59706 | 0.49429 | 0.69366 | 1.8006 | 0.482 | 0.19483 | 0.19138 |
| 223 | Crepis biennis L. | Crepis biennis | 0.25 | 0.52 | 0.200 | NA | NA | NA | 6.6 | 7 | 0 | 0.72781 | 0.47089 | 0.36365 | 1.47514 | 0.21383 | 0.17215 | 0.37071 |
| 223 | Matricaria discoidea DC. | Matricaria discoidea | 0.02 | 0.52 | 0.111 | 7 | 5.6 | 4.4 | 5.9 | 5.4 | 0.1 | 0.64871 | 0.63945 | 1.70759 | 2.05577 | 0.56937 | 0.21649 | 0.54803 |
| 223 | Poa annua L. | Poa annua | 0.05 | 0.52 | 0.400 | 7.4 | 5.3 | NA | 6 | 7.7 | 0.8 | 0.65201 | 0.63791 | 1.54898 | 2.03286 | 0.77966 | 0.37419 | 0.45721 |
| 223 | Centaurea cyanus L. | Centaurea cyanus | 0.01 | 0.52 | 0.050 | 8.6 | 5.9 | 3.5 | NA | 2.5 | 0 | 0.5503 | 0.53805 | 1.6668 | 2.00922 | 0.50104 | 0.3154 | 0.3619 |
| 223 | Achillea millefolium L. | Achillea millefolium millefolium | 0.02 | 0.52 | 0.080 | 8.6 | NA | 2.7 | 4.4 | 1.6 | 0 | 0.53557 | 0.50485 | 1.29532 | 1.90211 | 0.72298 | 0.28495 | 0.2778 |
| 223 | Papaver rhoeas L. | Papaver rhoeas | 0.05 | 0.52 | 0.100 | 8.6 | NA | 2.7 | 4.4 | 1.6 | 0 | 0.53557 | 0.50485 | 1.29532 | 1.90211 | 0.72298 | 0.28495 | 0.2778 |
| 223 | Silene dioica (L.) Clairv. | Silene dioica (Melandrium rubrum) | 0.1 | 0.52 | 0.050 | 7.9 | NA | 4.1 | 5.8 | NA | 0.9 | 0.54139 | 0.51128 | 1.31363 | 1.93633 | 0.8112 | 0.25085 | 0.32953 |
| 223 | Taraxacum officinale F.H.Wigg. | Taraxacum officinale agg. | 0.02 | 0.52 | 0.080 | 6.7 | 5.3 | 4.5 | 5.3 | 5.8 | 0 | 0.75464 | 0.73051 | 1.36974 | 1.95508 | 0.45974 | 0.15567 | 0.6659 |
| 224 | Malva sylvestris L. | Malva sylvestris | 0.06 | 0.9 | 0.053 | 8.6 | NA | 2.7 | 4.4 | 1.6 | 0 | 0.53557 | 0.50485 | 1.29532 | 1.90211 | 0.72298 | 0.28495 | 0.2778 |
| 224 | Isatis tinctoria L. | Isatis tinctoria | 0.1 | 0.9 | 0.060 | 7.3 | 5.4 | 4.1 | 7.6 | 4.2 | 0.4 | 0.58728 | 0.56055 | 1.34601 | 1.94534 | 0.82357 | 0.23743 | 0.34603 |
| 224 | Crepis biennis L. | Crepis biennis | 0.3 | 0.9 | 0.050 | 8.1 | NA | 3.5 | 6.1 | NA | 0.2 | 0.68175 | 0.6716 | 1.70824 | 1.99767 | 0.41848 | 0.25411 | 0.57318 |
| 224 | Taraxacum officinale F.H.Wigg. | Taraxacum officinale agg. | 0.05 | 0.9 | 0.050 | 8.4 | NA | 3 | 6 | NA | 0 | 0.46249 | 0.43915 | 1.39189 | 1.83687 | 0.38437 | 0.30571 | 0.38883 |
| 224 | Sisymbrium officinale (L.) Scop. | Sisymbrium officinale | 0.15 | 0.9 | 0.100 | 8.1 | NA | NA | 4.4 | 2.8 | 0.3 | 0.48868 | 0.46508 | 1.4426 | 2.04965 | 0.7278 | 0.33931 | 0.35628 |
| 224 | Leucanthemum vulgare Lam. | Leucanthemum vulgare (leucanthem.) | 0.05 | 0.9 | 0.050 | 8.1 | NA | NA | 4.4 | 2.8 | 0.3 | 0.48868 | 0.46508 | 1.4426 | 2.04965 | 0.7278 | 0.33931 | 0.35628 |
| 224 | Phacelia tanacetifolia Benth. | NA | 0.01 | 0.9 | 0.080 | 8.1 | NA | 3.5 | 6.1 | NA | 0.2 | 0.68175 | 0.6716 | 1.70824 | 1.99767 | 0.41848 | 0.25411 | 0.57318 |
| 224 | Geranium pusillum L. | Geranium pusillum | 0.05 | 0.9 | 0.050 | NA | NA | NA | NA | NA | NA | NA | NA | NA | NA | NA | NA | NA |
| 224 | Echium vulgare L. | Echium vulgare | 0.05 | 0.9 | 0.100 | 8 | NA | 3.7 | NA | 2 | 0 | 0.55827 | 0.46888 | 0.89651 | 1.79355 | 0.39043 | 0.26528 | 0.38307 |
| 224 | Mercurialis annua L. | Mercurialis annua | 0.05 | 0.9 | 0.300 | 7.4 | 5.3 | NA | 6 | 7.7 | 0.8 | 0.65201 | 0.63791 | 1.54898 | 2.03286 | 0.77966 | 0.37419 | 0.45721 |
| 224 | Salvia pratensis L. | Salvia pratensis | 0.02 | 0.9 | 0.053 | NA | 5.4 | 5.7 | 6.8 | 7 | 0 | 0.67742 | 0.24559 | 0.14546 | 1.36309 | 0.06547 | 0.19208 | 0.12644 |
| 224 | Sinapis arvensis L. | Sinapis arvensis | 0.01 | 0.9 | 0.050 | 7.6 | NA | NA | 6.5 | 6.3 | 0.7 | 0.75548 | 0.75048 | 1.8409 | 2.08419 | 0.63746 | 0.23177 | 0.58814 |
| 225 | Crepis biennis L. | Crepis biennis | 0.25 | 0.93 | 0.540 | 7.4 | 5.3 | NA | 6 | 7.7 | 0.8 | 0.65201 | 0.63791 | 1.54898 | 2.03286 | 0.77966 | 0.37419 | 0.45721 |
| 225 | Alcea rosea L. | Alcea rosea | 0.14 | 0.93 | 0.100 | 7.9 | NA | 4.1 | 5.8 | NA | 0.9 | 0.54139 | 0.51128 | 1.31363 | 1.93633 | 0.8112 | 0.25085 | 0.32953 |
| 225 | Leucanthemum vulgare Lam. | Leucanthemum vulgare (leucanthem.) | 0.05 | 0.93 | 0.067 | 7.3 | 6.1 | 3.8 | 6 | 5.1 | 0 | 0.62987 | 0.59281 | 1.27995 | 1.90491 | 0.6275 | 0.3036 | 0.42608 |
| 225 | Taraxacum officinale F.H.Wigg. | Taraxacum officinale agg. | 0.05 | 0.93 | 0.111 | 8.4 | NA | 3 | 6 | NA | 0 | 0.46249 | 0.43915 | 1.39189 | 1.83687 | 0.38437 | 0.30571 | 0.38883 |
| 225 | Malva sylvestris L. | Malva sylvestris | 0.02 | 0.93 | 0.100 | 8.4 | NA | 3 | 6 | NA | 0 | 0.46249 | 0.43915 | 1.39189 | 1.83687 | 0.38437 | 0.30571 | 0.38883 |
| 225 | Achillea millefolium L. | Achillea millefolium millefolium | 0.02 | 0.93 | 0.431 | 7.4 | 5.3 | NA | 6 | 7.7 | 0.8 | 0.65201 | 0.63791 | 1.54898 | 2.03286 | 0.77966 | 0.37419 | 0.45721 |
| 225 | Senecio vulgaris L. | Senecio vulgaris | 0.1 | 0.93 | 0.100 | 8.4 | NA | 3 | 6 | NA | 0 | 0.46249 | 0.43915 | 1.39189 | 1.83687 | 0.38437 | 0.30571 | 0.38883 |
| 225 | Sinapis arvensis L. | Sinapis arvensis | 0.01 | 0.93 | 0.053 | 8.4 | 7.2 | 2.6 | 6.8 | 3.6 | 0.2 | 0.58203 | 0.49726 | 0.89718 | 1.5951 | 0.3257 | 0.17939 | 0.36267 |

| | | | | | | | | | | | | | | | | | | |
|---|---|---|---|---|---|---|---|---|---|---|---|---|---|---|---|---|---|---|
| 225 | Centaurea jacea L. | Centaurea jacea | 0.17 | 0.93 | 0.100 | 7.3 | 6.1 | 3.8 | 6 | 5.1 | 0 | 0.62987 | 0.59281 | 1.27995 | 1.90491 | 0.6275 | 0.3036 | 0.42608 |
| 225 | Isatis tinctoria L. | Isatis tinctoria | 0.01 | 0.93 | 0.450 | 7.4 | 5.3 | NA | 6 | 7.7 | 0.8 | 0.65201 | 0.63791 | 1.54898 | 2.03286 | 0.77966 | 0.37419 | 0.45721 |
| 225 | Silene dioica (L.) Clairv. | Silene dioica (Melandrium rubrum) | 0.02 | 0.93 | 0.020 | 7.3 | 5.4 | 4.1 | 7.6 | 4.2 | 0.4 | 0.58728 | 0.56055 | 1.34601 | 1.94534 | 0.82357 | 0.23743 | 0.34603 |
| 225 | Centaurea cyanus L. | Centaurea cyanus | 0.01 | 0.93 | 0.051 | 7.6 | NA | NA | 6.5 | 6.3 | 0.7 | 0.75548 | 0.75048 | 1.8409 | 2.08419 | 0.63746 | 0.23177 | 0.58814 |
| 225 | Rumex acetosa L. | Rumex acetosa | 0.01 | 0.93 | 0.150 | 7.6 | NA | NA | 6.5 | 6.3 | 0.7 | 0.75548 | 0.75048 | 1.8409 | 2.08419 | 0.63746 | 0.23177 | 0.58814 |
| 225 | Carduus nutant L. | Carduus nutans agg. | 0.05 | 0.93 | 0.051 | NA | NA | NA | NA | NA | NA | NA | NA | NA | NA | NA | NA | NA |
| 225 | Ranunculus repens L. | Ranunculus repens | 0.02 | 0.93 | 0.150 | NA | NA | NA | NA | NA | NA | NA | NA | NA | NA | NA | NA | NA |
| 226 | Phleum pratense L. | Phleum pratense pratense | 1 | 1 | 0.100 | NA | NA | NA | NA | NA | NA | NA | NA | NA | NA | NA | NA | NA |
| 227 | Phleum pratense L. | Phleum pratense pratense | 0.25 | 1 | 0.100 | NA | NA | NA | NA | NA | NA | NA | NA | NA | NA | NA | NA | NA |
| 227 | Veronica hederifolia L. | Veronica hederifolia agg. | 0.05 | 1 | 0.200 | 8.1 | NA | NA | 4.4 | 2.8 | 0.3 | 0.48868 | 0.46508 | 1.4426 | 2.04965 | 0.7278 | 0.33931 | 0.35628 |
| 227 | Poa annua L. | Poa annua | 0.25 | 1 | 0.050 | 7.3 | 6.1 | 3.8 | 6 | 5.1 | 0 | 0.62987 | 0.59281 | 1.27995 | 1.90491 | 0.6275 | 0.3036 | 0.42608 |
| 227 | Polygonum aviculare L. | Polygonum aviculare agg. | 0.1 | 1 | 0.056 | 8.6 | NA | 2.7 | 4.4 | 1.6 | 0 | 0.53557 | 0.50485 | 1.29532 | 1.90211 | 0.72298 | 0.28495 | 0.2778 |
| 227 | Erigeron canadensis L. | Conyza canadensis | 0.05 | 1 | 0.020 | 6.7 | NA | NA | 6.5 | 7.7 | 0.1 | 0.70945 | 0.61373 | 0.80583 | 1.7717 | 0.43958 | 0.26577 | 0.51423 |
| 227 | Ranunculus ficaria L. | Ranunculus ficaria (Ficaria verna) | 0.05 | 1 | 0.050 | 7.3 | NA | NA | 7.6 | 7.7 | 0.4 | 0.75479 | 0.72524 | 1.28692 | 1.77374 | 0.61324 | 0.18283 | 0.56773 |
| 227 | Malva L. | NA | 0.25 | 1 | 0.086 | 6.7 | NA | NA | 6.5 | 7.7 | 0.1 | 0.70945 | 0.61373 | 0.80583 | 1.7717 | 0.43958 | 0.26577 | 0.51423 |
| 228 | Stellaria media (L.) Vill. | Stellaria media agg. | 0.28 | 1 | 0.053 | NA | NA | NA | 7 | NA | 0 | 0.66957 | 0.20824 | 0.12967 | 1.34554 | 0.07292 | 0.20428 | 0.13248 |
| 228 | Veronica hederifolia L. | Veronica hederifolia agg. | 0.05 | 1 | 0.050 | 6.8 | NA | 5.6 | 5.8 | NA | 0.1 | 0.52169 | 0.45097 | 0.93044 | 1.87015 | 0.67593 | 0.2975 | 0.27054 |
| 228 | Poa annua L. | Poa annua | 0.05 | 1 | 0.040 | 6.8 | NA | 5.6 | 5.8 | NA | 0.1 | 0.52169 | 0.45097 | 0.93044 | 1.87015 | 0.67593 | 0.2975 | 0.27054 |
| 228 | Phleum pratense L. | Phleum pratense pratense | 0.28 | 1 | 0.090 | 6.8 | NA | 5.6 | 5.8 | NA | 0.1 | 0.52169 | 0.45097 | 0.93044 | 1.87015 | 0.67593 | 0.2975 | 0.27054 |
| 228 | Hedera helix L. | Hedera helix | 0.1 | 1 | 0.050 | 7.7 | NA | NA | 5.6 | NA | 0.7 | 0.53223 | 0.52325 | 1.73719 | 2.14421 | 0.94278 | 0.39374 | 0.35054 |
| 228 | Taraxacum officinale F.H.Wigg. | Taraxacum officinale agg. | 0.19 | 1 | 0.230 | 6.8 | NA | 5.6 | 5.8 | NA | 0.1 | 0.52169 | 0.45097 | 0.93044 | 1.87015 | 0.67593 | 0.2975 | 0.27054 |
| 228 | Polygonum aviculare L. | Polygonum aviculare agg. | 0.05 | 1 | 0.097 | 7.7 | NA | NA | 5.6 | NA | 0.7 | 0.53223 | 0.52325 | 1.73719 | 2.14421 | 0.94278 | 0.39374 | 0.35054 |
| 229 | Euphorbia peplus L. | Euphorbia peplus | 0.3 | 0.9 | 0.115 | NA | NA | NA | NA | NA | NA | NA | NA | NA | NA | NA | NA | NA |
| 229 | Tussilago farfara L. | Tussilago farfara | 0.1 | 0.9 | 0.020 | 7 | 5.6 | 4.4 | 5.9 | 5.4 | 0.1 | 0.64871 | 0.63945 | 1.70759 | 2.05577 | 0.56937 | 0.21649 | 0.54803 |
| 229 | Taraxacum officinale F.H.Wigg. | Taraxacum officinale agg. | 0.5 | 0.9 | 0.395 | 7.4 | 5.3 | NA | 6 | 7.7 | 0.8 | 0.65201 | 0.63791 | 1.54898 | 2.03286 | 0.77966 | 0.37419 | 0.45721 |
| 230 | Phleum pratense L. | Phleum pratense pratense | 0.64 | 1 | 0.118 | 8.6 | 5.9 | 3.5 | NA | 2.5 | 0 | 0.5503 | 0.53805 | 1.6668 | 2.00922 | 0.50104 | 0.3154 | 0.3619 |
| 230 | Sisymbrium officinale (L.) Scop. | Sisymbrium officinale | 0.24 | 1 | 0.100 | 7.9 | 5.1 | 5.1 | 6.7 | 7.7 | 0.2 | 0.75642 | 0.75467 | 2.08644 | 2.23861 | 0.83282 | 0.22413 | 0.56443 |
| 230 | Stellaria media (L.) Vill. | Stellaria media agg. | 0.05 | 1 | 0.100 | 7.1 | 4.9 | NA | 6 | NA | 1 | 0.59517 | 0.59064 | 1.89772 | 2.12267 | 0.65091 | 0.41213 | 0.37185 |
| 230 | Ranunculus ficaria L. | Ranunculus ficaria (Ficaria verna) | 0.05 | 1 | 0.050 | 8.4 | 7.2 | 2.6 | 6.8 | 3.6 | 0.2 | 0.58203 | 0.49726 | 0.89718 | 1.5951 | 0.3257 | 0.17939 | 0.36267 |
| 230 | Veronica hederifolia L. | Veronica hederifolia agg. | 0.02 | 1 | 0.600 | 7.4 | 5.3 | NA | 6 | 7.7 | 0.8 | 0.65201 | 0.63791 | 1.54898 | 2.03286 | 0.77966 | 0.37419 | 0.45721 |
| 231 | Taraxacum officinale F.H.Wigg. | Taraxacum officinale agg. | 0.65 | 1 | 0.045 | 7.3 | NA | 3.9 | 6.1 | 3.9 | 0.1 | 0.55903 | 0.41538 | 0.65189 | 1.65181 | 0.43719 | 0.24279 | 0.21709 |
| 231 | Poa annua L. | Poa annua | 0.15 | 1 | 0.050 | 8.1 | NA | NA | 5.2 | NA | 0.1 | 0.51276 | 0.49581 | 1.51667 | 2.20989 | 0.98959 | 0.46997 | 0.3594 |
| 231 | Chaerophyllum temulum L. | Chaerophyllum temulum | 0.05 | 1 | 0.052 | NA | NA | NA | NA | NA | NA | NA | NA | NA | NA | NA | NA | NA |
| 231 | Polygonum aviculare L. | Polygonum aviculare agg. | 0.05 | 1 | 0.320 | 7.4 | 5.3 | NA | 6 | 7.7 | 0.8 | 0.65201 | 0.63791 | 1.54898 | 2.03286 | 0.77966 | 0.37419 | 0.45721 |
| 231 | Veronica hederifolia L. | Veronica hederifolia agg. | 0.05 | 1 | 0.350 | 7.4 | 5.3 | NA | 6 | 7.7 | 0.8 | 0.65201 | 0.63791 | 1.54898 | 2.03286 | 0.77966 | 0.37419 | 0.45721 |
| 231 | Stellaria media (L.) Vill. | Stellaria media agg. | 0.05 | 1 | 0.050 | 7.3 | NA | 3.9 | 6.1 | 3.9 | 0.1 | 0.55903 | 0.41538 | 0.65189 | 1.65181 | 0.43719 | 0.24279 | 0.21709 |
| 232 | Taraxacum officinale F.H.Wigg. | Taraxacum officinale agg. | 0.4 | 0.9 | 0.050 | 6.6 | NA | NA | 6.9 | 5.2 | 0.4 | 0.54634 | 0.5169 | 1.30759 | 1.85566 | 0.79671 | 0.26499 | 0.28358 |
| 232 | Poa annua L. | Poa annua | 0.3 | 0.9 | 0.123 | 7.1 | 5.3 | 4.5 | 5.9 | 4.8 | 0.3 | 0.52518 | 0.5113 | 1.61045 | 2.05509 | 0.96495 | 0.27897 | 0.32911 |
| 232 | Stellaria media (L.) Vill. | Stellaria media agg. | 0.1 | 0.9 | 0.050 | 8.4 | 7.2 | 2.6 | 6.8 | 3.6 | 0.2 | 0.58203 | 0.49726 | 0.89718 | 1.5951 | 0.3257 | 0.17939 | 0.36267 |
| 232 | Hordeum murinum L. | Hordeum murinum agg. | 0.1 | 0.9 | 0.250 | 7.8 | 5.5 | 3.3 | 6.4 | 3.3 | 0.1 | 0.51939 | 0.38466 | 0.67536 | 1.6325 | 0.37226 | 0.27333 | 0.16989 |
| 233 | Poa annua L. | Poa annua | 0.3 | 0.95 | 0.550 | 7.4 | 5.3 | NA | 6 | 7.7 | 0.8 | 0.65201 | 0.63791 | 1.54898 | 2.03286 | 0.77966 | 0.37419 | 0.45721 |
| 233 | Taraxacum officinale F.H.Wigg. | Taraxacum officinale agg. | 0.15 | 0.95 | 0.053 | 8.1 | NA | NA | 4.4 | 2.8 | 0.3 | 0.48868 | 0.46508 | 1.4426 | 2.04965 | 0.7278 | 0.33931 | 0.35628 |
| 233 | Stellaria media (L.) Vill. | Stellaria media agg. | 0.1 | 0.95 | 0.162 | 6.6 | NA | NA | 6.9 | 5.2 | 0.4 | 0.54634 | 0.5169 | 1.30759 | 1.85566 | 0.79671 | 0.26499 | 0.28358 |
| 233 | Hordeum murinum L. | Hordeum murinum agg. | 0.4 | 0.95 | 0.110 | 7.3 | 5.4 | 4.1 | 7.6 | 4.2 | 0.4 | 0.58728 | 0.56055 | 1.34601 | 1.94534 | 0.82357 | 0.23743 | 0.34603 |
| 234 | Poa annua L. | Poa annua | 0.3 | 1 | 0.211 | 7 | NA | NA | NA | NA | 0.3 | 0.56899 | 0.4288 | 0.64817 | 1.63031 | 0.49671 | 0.22558 | 0.25973 |
| 234 | Hordeum murinum L. | Hordeum murinum agg. | 0.7 | 1 | 0.100 | 7.6 | NA | NA | 5.8 | NA | 0.5 | 0.55139 | 0.54271 | 1.75426 | 2.13166 | 0.95232 | 0.27891 | 0.39876 |
| 235 | Poa annua L. | Poa annua | 0.2 | 1 | 0.083 | 6.6 | NA | NA | 6.9 | 5.2 | 0.4 | 0.54634 | 0.5169 | 1.30759 | 1.85566 | 0.79671 | 0.26499 | 0.28358 |
| 235 | Hordeum murinum L. | Hordeum murinum agg. | 0.8 | 1 | 0.440 | 7.4 | 5.3 | NA | 6 | 7.7 | 0.8 | 0.65201 | 0.63791 | 1.54898 | 2.03286 | 0.77966 | 0.37419 | 0.45721 |
| 236 | Poa annua L. | Poa annua | 0.4 | 0.4 | 0.052 | NA | NA | NA | NA | NA | NA | NA | NA | NA | NA | NA | NA | NA |
| 237 | Poa annua L. | Poa annua | 0.1 | 0.45 | 0.121 | 7.3 | NA | 3.9 | 6.1 | 3.9 | 0.1 | 0.55903 | 0.41538 | 0.65189 | 1.65181 | 0.43719 | 0.24279 | 0.21709 |
| 237 | Stellaria media (L.) Vill. | Stellaria media agg. | 0.05 | 0.45 | 0.020 | 7.3 | NA | 3.9 | 6.1 | 3.9 | 0.1 | 0.55903 | 0.41538 | 0.65189 | 1.65181 | 0.43719 | 0.24279 | 0.21709 |
| 237 | Veronica hederifolia L. | Veronica hederifolia agg. | 0.25 | 0.45 | 0.020 | 7 | 5.6 | 4.4 | 5.9 | 5.4 | 0.1 | 0.64871 | 0.63945 | 1.70759 | 2.05577 | 0.56937 | 0.21649 | 0.54803 |
| 237 | Taraxacum officinale F.H.Wigg. | Taraxacum officinale agg. | 0.05 | 0.45 | 0.040 | 7.3 | NA | 3.9 | 6.1 | 3.9 | 0.1 | 0.55903 | 0.41538 | 0.65189 | 1.65181 | 0.43719 | 0.24279 | 0.21709 |
| 238 | Poa annua L. | Poa annua | 0.15 | 0.5 | 0.060 | 7.1 | 5.3 | 4.5 | 5.9 | 4.8 | 0.3 | 0.52518 | 0.5113 | 1.61045 | 2.05509 | 0.96495 | 0.27897 | 0.32911 |
| 238 | Stellaria media (L.) Vill. | Stellaria media agg. | 0.1 | 0.5 | 0.090 | 7.6 | NA | NA | 5.8 | NA | 0.5 | 0.55139 | 0.54271 | 1.75426 | 2.13166 | 0.95232 | 0.27891 | 0.39876 |
| 238 | Trifolium repens L. | Trifolium repens | 0.25 | 0.5 | 0.053 | 8.6 | NA | 2.7 | 4.4 | 1.6 | 0 | 0.53557 | 0.50485 | 1.29532 | 1.90211 | 0.72298 | 0.28495 | 0.2778 |
| 239 | Poa annua L. | Poa annua | 0.6 | 0.6 | 0.051 | 7.1 | 5.3 | 4.5 | 5.9 | 4.8 | 0.3 | 0.52518 | 0.5113 | 1.61045 | 2.05509 | 0.96495 | 0.27897 | 0.32911 |
| 240 | Poa annua L. | Poa annua | 0.5 | 1 | 0.020 | 8.6 | NA | 2.7 | 4.4 | 1.6 | 0 | 0.53557 | 0.50485 | 1.29532 | 1.90211 | 0.72298 | 0.28495 | 0.2778 |
| 240 | Hordeum murinum L. | Hordeum murinum agg. | 0.4 | 1 | 0.020 | 7 | 5.6 | 4.4 | 5.9 | 5.4 | 0.1 | 0.64871 | 0.63945 | 1.70759 | 2.05577 | 0.56937 | 0.21649 | 0.54803 |
| 240 | Taraxacum officinale F.H.Wigg. | Taraxacum officinale agg. | 0.1 | 1 | 0.080 | 7.1 | 5.3 | 4.5 | 5.9 | 4.8 | 0.3 | 0.52518 | 0.5113 | 1.61045 | 2.05509 | 0.96495 | 0.27897 | 0.32911 |
| 241 | Poa annua L. | Poa annua | 0.3 | 1 | 0.052 | 8.1 | NA | NA | 5.2 | NA | 0.1 | 0.51276 | 0.49581 | 1.51667 | 2.20989 | 0.98959 | 0.46997 | 0.3594 |
| 241 | Hordeum murinum L. | Hordeum murinum agg. | 0.3 | 1 | 0.020 | 8.6 | 5.9 | 3.5 | NA | 2.5 | 0 | 0.5503 | 0.53805 | 1.6668 | 2.00922 | 0.50104 | 0.3154 | 0.3619 |
| 241 | Taraxacum officinale F.H.Wigg. | Taraxacum officinale agg. | 0.2 | 1 | 0.150 | NA | NA | NA | NA | NA | NA | NA | NA | NA | NA | NA | NA | NA |
| 241 | Chaerophyllum temulum L. | Chaerophyllum temulum | 0.05 | 1 | 0.042 | 8.1 | NA | 3.5 | 6.1 | NA | 0.2 | 0.68175 | 0.6716 | 1.70824 | 1.99767 | 0.41848 | 0.25411 | 0.57318 |
| 241 | Veronica hederifolia L. | Veronica hederifolia agg. | 0.05 | 1 | 0.042 | 8.6 | NA | 2.7 | 4.4 | 1.6 | 0 | 0.53557 | 0.50485 | 1.29532 | 1.90211 | 0.72298 | 0.28495 | 0.2778 |
| 241 | Stellaria media (L.) Vill. | Stellaria media agg. | 0.1 | 1 | 0.051 | 7.1 | 5.3 | 4.5 | 5.9 | 4.8 | 0.3 | 0.52518 | 0.5113 | 1.61045 | 2.05509 | 0.96495 | 0.27897 | 0.32911 |
| 242 | Poa annua L. | Poa annua | 0.4 | 0.9 | 0.100 | 7.6 | NA | NA | 5.8 | NA | 0.5 | 0.55139 | 0.54271 | 1.75426 | 2.13166 | 0.95232 | 0.27891 | 0.39876 |
| 242 | Hordeum murinum L. | Hordeum murinum agg. | 0.1 | 0.9 | 0.050 | NA | NA | NA | NA | NA | NA | NA | NA | NA | NA | NA | NA | NA |
| 242 | Taraxacum officinale F.H.Wigg. | Taraxacum officinale agg. | 0.1 | 0.9 | 0.140 | 8.5 | NA | 3.5 | 6.3 | 4.1 | 0.2 | 0.57633 | 0.55499 | 1.44489 | 1.81436 | 0.579 | 0.25616 | 0.27657 |
| 242 | Chaerophyllum temulum L. | Chaerophyllum temulum | 0.05 | 0.9 | 0.050 | 6.7 | 5.3 | 4.5 | 5.3 | 5.8 | 0 | 0.75464 | 0.73051 | 1.36974 | 1.95508 | 0.45974 | 0.15567 | 0.6659 |
| 242 | Lamium purpureum L. | Lamium purpureum | 0.05 | 0.9 | 0.080 | 7.1 | 5.3 | 4.5 | 5.9 | 4.8 | 0.3 | 0.52518 | 0.5113 | 1.61045 | 2.05509 | 0.96495 | 0.27897 | 0.32911 |
| 242 | Veronica hederifolia L. | Veronica hederifolia agg. | 0.05 | 0.9 | 0.130 | 7.1 | 5.3 | 4.5 | 5.9 | 4.8 | 0.3 | 0.52518 | 0.5113 | 1.61045 | 2.05509 | 0.96495 | 0.27897 | 0.32911 |
| 242 | Stellaria media (L.) Vill. | Stellaria media agg. | 0.05 | 0.9 | 0.200 | 7.4 | 5.3 | NA | 6 | 7.7 | 0.8 | 0.65201 | 0.63791 | 1.54898 | 2.03286 | 0.77966 | 0.37419 | 0.45721 |
| 242 | Hedera helix L. | Hedera helix | 0.1 | 0.9 | 0.300 | NA | NA | NA | NA | NA | NA | NA | NA | NA | NA | NA | NA | NA |
| 243 | Poa annua L. | Poa annua | 0.5 | 1 | 0.040 | 7 | NA | NA | NA | NA | 0.3 | 0.56899 | 0.4288 | 0.64817 | 1.63031 | 0.49671 | 0.22558 | 0.25973 |
| 243 | Hordeum murinum L. | Hordeum murinum agg. | 0.4 | 1 | 0.010 | 5.2 | 5.9 | 5.4 | 7.3 | 8 | 0 | 0.67947 | 0.25214 | 0.1495 | 1.33661 | 0.07498 | 0.19025 | 0.12819 |
| 243 | Taraxacum officinale F.H.Wigg. | Taraxacum officinale agg. | 0.1 | 1 | 0.020 | 7 | 5.6 | 4.4 | 5.9 | 5.4 | 0.1 | 0.64871 | 0.63945 | 1.70759 | 2.05577 | 0.56937 | 0.21649 | 0.54803 |
| 248 | Poa annua L. | Poa annua | 0.6 | 1 | 0.050 | 5.7 | NA | NA | 6.4 | 7.4 | 0 | 0.68859 | 0.26809 | 0.17316 | 1.34857 | 0.09545 | 0.19552 | 0.14349 |
| 248 | Stellaria media (L.) Vill. | Stellaria media agg. | 0.4 | 1 | 0.010 | NA | NA | 4.5 | 6.6 | 6.6 | 0 | 0.78059 | 0.60963 | 0.54418 | 1.63701 | 0.26733 | 0.16232 | 0.54354 |
| 249 | Taraxacum officinale F.H.Wigg. | Taraxacum officinale agg. | 0.3 | 0.95 | 0.204 | 8.6 | 5.9 | 3.5 | NA | 2.5 | 0 | 0.5503 | 0.53805 | 1.6668 | 2.00922 | 0.50104 | 0.3154 | 0.3619 |
| 249 | Poa annua L. | Poa annua | 0.5 | 0.95 | 0.084 | 7.1 | 5.5 | 4.1 | 6.5 | 3.7 | 0 | 0.52269 | 0.47116 | 1.1242 | 1.93135 | 0.85876 | 0.19681 | 0.27698 |
| 249 | Cornus sanguinea L. | Cornus sanguinea | 0.15 | 0.95 | 0.117 | 8.1 | 6.2 | NA | 7.1 | 7.6 | 0.1 | 0.8111 | 0.79007 | 1.4109 | 1.89172 | 0.89447 | 0.17462 | 0.49967 |
| 250 | Taraxacum officinale F.H.Wigg. | Taraxacum officinale agg. | 0.25 | 1 | 0.102 | 6.7 | 5.3 | 4.5 | 5.3 | 5.8 | 0 | 0.75464 | 0.73051 | 1.36974 | 1.95508 | 0.45974 | 0.15567 | 0.6659 |
| 250 | Senecio vulgaris L. | Senecio vulgaris | 0.05 | 1 | 0.145 | 7.7 | NA | NA | 6.4 | NA | 0 | 0.51083 | 0.48089 | 1.29849 | 1.93695 | 0.96262 | 0.19429 | 0.30238 |
| 250 | Plantago major L. | Plantago major ssp. major | 0.1 | 1 | 0.082 | 7.4 | 5.3 | NA | 6 | 7.7 | 0.8 | 0.65201 | 0.63791 | 1.54898 | 2.03286 | 0.77966 | 0.37419 | 0.45721 |
| 250 | Sagina procumbens L. | Sagina procumbens | 0.05 | 1 | 0.072 | 7.4 | 5.9 | 3.8 | 6 | NA | 0 | 0.7907 | 0.77966 | 1.64404 | 2.03844 | 0.57393 | 0.17512 | 0.69169 |
| 250 | Stellaria media (L.) Vill. | Stellaria media agg. | 0.1 | 1 | 0.056 | NA | NA | NA | NA | NA | NA | NA | NA | NA | NA | NA | NA | NA |
| 250 | Poa annua L. | Poa annua | 0.1 | 1 | 0.050 | 7.8 | NA | 4.6 | 6.2 | 5.8 | 0.1 | 0.71321 | 0.68276 | 1.29546 | 1.8275 | 0.62188 | 0.18605 | 0.48742 |
| 250 | Oxalis corniculata L. | Oxalis corniculata | 0.05 | 1 | 0.100 | NA | NA | 5.3 | 6.2 | NA | 0.1 | 0.68682 | 0.14816 | 0.05813 | 1.31254 | 0.01433 | 0.20126 | 0.10505 |
| 250 | Erigeron canadensis L. | Conyza canadensis | 0.1 | 1 | 0.020 | NA | NA | 4.5 | 6.6 | 6.6 | 0 | 0.78059 | 0.60963 | 0.54418 | 1.63701 | 0.26733 | 0.16232 | 0.54354 |
| 250 | Hypochaeris radicata L. | Hypochoeris radicata | 0.2 | 1 | 0.050 | NA | NA | 4.5 | 6.6 | 6.6 | 0 | 0.78059 | 0.60963 | 0.54418 | 1.63701 | 0.26733 | 0.16232 | 0.54354 |
| 251 | Taraxacum officinale F.H.Wigg. | Taraxacum officinale agg. | 0.09 | 1 | 0.111 | 8.1 | NA | 3.9 | 6.9 | 6.4 | 0.4 | 0.80098 | 0.76713 | 1.29016 | 1.92909 | 0.75604 | 0.19458 | 0.52486 |
| 251 | Hypochaeris radicata L. | Hypochoeris radicata | 0.15 | 1 | 0.111 | NA | NA | NA | NA | NA | NA | NA | NA | NA | NA | NA | NA | NA |
| 251 | Oxalis corniculata L. | Oxalis corniculata | 0.2 | 1 | 0.050 | NA | NA | 4.5 | 6.6 | 6.6 | 0 | 0.78059 | 0.60963 | 0.54418 | 1.63701 | 0.26733 | 0.16232 | 0.54354 |
| 251 | Chelidonium majus L. | Chelidonium majus | 0.08 | 1 | 0.056 | 7.2 | NA | NA | 6.6 | 6.8 | 0 | 0.81299 | 0.76483 | 1.09209 | 1.84162 | 0.41616 | 0.14633 | 0.71443 |
| 251 | Geranium robertianum L. | Geranium robertianum | 0.1 | 1 | 0.100 | 6.7 | NA | NA | 6.5 | 7.7 | 0.1 | 0.70945 | 0.61373 | 0.80583 | 1.7717 | 0.43958 | 0.26577 | 0.51423 |
| 251 | Senecio vulgaris L. | Senecio vulgaris | 0.1 | 1 | 0.100 | NA | NA | 5.5 | 6.2 | 6.8 | 0 | 0.67445 | 0.21586 | 0.12312 | 1.32299 | 0.04699 | 0.18459 | 0.11909 |
| 251 | Euphorbia peplus L. | Euphorbia peplus | 0.08 | 1 | 0.051 | NA | NA | 4.5 | 6.6 | 6.6 | 0 | 0.78059 | 0.60963 | 0.54418 | 1.63701 | 0.26733 | 0.16232 | 0.54354 |
| 251 | Poa annua L. | Poa annua | 0.05 | 1 | 0.021 | 8.3 | 7.4 | 1.5 | 5 | NA | 0 | 0.82036 | 0.80731 | 1.56891 | 2.0257 | 0.90815 | 0.20703 | 0.54552 |
| 251 | Sagina procumbens L. | Sagina procumbens | 0.05 | 1 | 0.070 | 5.5 | 7 | 5.5 | 6 | 6.5 | 0 | 0.7044 | 0.27957 | 0.20257 | 1.35002 | 0.06865 | 0.19462 | 0.18523 |
| 251 | Erigeron canadensis L. | Conyza canadensis | 0.1 | 1 | 0.300 | 7.1 | 4.9 | NA | 6 | NA | 1 | 0.59517 | 0.59064 | 1.89772 | 2.12267 | 0.65091 | 0.41213 | 0.37185 |
| 252 | Poa annua L. | Poa annua | 0.8 | 1 | 0.053 | 7.1 | 4.9 | NA | 6 | NA | 1 | 0.59517 | 0.59064 | 1.89772 | 2.12267 | 0.65091 | 0.41213 | 0.37185 |
| 252 | Stellaria media (L.) Vill. | Stellaria media agg. | 0.2 | 1 | 0.200 | 7.4 | 5.3 | NA | 6 | 7.7 | 0.8 | 0.65201 | 0.63791 | 1.54898 | 2.03286 | 0.77966 | 0.37419 | 0.45721 |
| 253 | Taraxacum officinale F.H.Wigg. | Taraxacum officinale agg. | 0.8 | 1 | 0.050 | NA | NA | NA | NA | NA | NA | NA | NA | NA | NA | NA | NA | NA |
| 253 | Poa annua L. | Poa annua | 0.2 | 1 | 0.100 | 7.4 | 5.3 | NA | 6 | 7.7 | 0.8 | 0.65201 | 0.63791 | 1.54898 | 2.03286 | 0.77966 | 0.37419 | 0.45721 |
| 254 | Rumex obtusifolius L. | Rumex obtusifolius | 0.2 | 1 | 0.440 | NA | NA | NA | NA | NA | NA | NA | NA | NA | NA | NA | NA | NA |

| | | | | | | | | | | | | | | | | | |
|---|---|---|---|---|---|---|---|---|---|---|---|---|---|---|---|---|---|
| 254 | Cornus sanguinea L. | Cornus sanguinea | 0.2 | 1 | 0.100 | 7.2 | NA | NA | 6.5 | 8.4 | 0.1 | 0.5933 | 0.54722 | 1.04949 | 1.60983 | 0.64032 | 0.25162 | 0.31795 |
| 254 | Hedera helix L. | Hedera helix | 0.6 | 1 | 0.250 | NA | NA | NA | NA | NA | NA | NA | NA | NA | NA | NA | NA | NA |
| 255 | Taraxacum officinale F.H.Wigg. | Taraxacum officinale agg. | 0.33 | 0.98 | 0.100 | NA | 5.4 | 5.7 | 6.8 | 7 | 0 | 0.67742 | 0.24559 | 0.14546 | 1.36309 | 0.06547 | 0.19208 | 0.12644 |
| 255 | Poa annua L. | Poa annua | 0.5 | 0.98 | 0.150 | 7.9 | 5.7 | 4.2 | 7 | 7.2 | 0.1 | 0.76998 | 0.7507 | 1.46055 | 1.90983 | 0.91556 | 0.1857 | 0.55131 |
| 255 | Geranium molle L. | Geranium molle | 0.07 | 0.98 | 0.100 | 7.4 | 5.3 | NA | 6 | 7.7 | 0.8 | 0.65201 | 0.63791 | 1.54898 | 2.03286 | 0.77966 | 0.37419 | 0.45721 |
| 255 | Stellaria media (L.) Vill. | Stellaria media agg. | 0.03 | 0.98 | 0.150 | NA | NA | NA | NA | NA | NA | NA | NA | NA | NA | NA | NA | NA |
| 255 | Veronica hederifolia L. | Veronica hederifolia agg. | 0.05 | 0.98 | 0.200 | NA | NA | NA | NA | NA | NA | NA | NA | NA | NA | NA | NA | NA |
| 256 | Taraxacum officinale F.H.Wigg. | Taraxacum officinale agg. | 0.28 | 0.95 | 0.147 | NA | NA | NA | NA | NA | NA | NA | NA | NA | NA | NA | NA | NA |
| 256 | Poa annua L. | Poa annua | 0.4 | 0.95 | 0.050 | 7.4 | 5.3 | NA | 6 | 7.7 | 0.8 | 0.65201 | 0.63791 | 1.54898 | 2.03286 | 0.77966 | 0.37419 | 0.45721 |
| 256 | Geranium molle L. | Geranium molle | 0.1 | 0.95 | 0.071 | 7.4 | 5.3 | NA | 6 | 7.7 | 0.8 | 0.65201 | 0.63791 | 1.54898 | 2.03286 | 0.77966 | 0.37419 | 0.45721 |
| 256 | Stellaria media (L.) Vill. | Stellaria media agg. | 0.08 | 0.95 | 0.050 | 7 | 5.6 | 4.4 | 5.9 | 5.4 | 0.1 | 0.64871 | 0.63945 | 1.70759 | 2.05577 | 0.56937 | 0.21649 | 0.54803 |
| 256 | Capsella rubella Reut. | NA | 0.02 | 0.95 | 0.100 | NA | 5.4 | 5.7 | 6.8 | 7 | 0 | 0.67742 | 0.24559 | 0.14546 | 1.36309 | 0.06547 | 0.19208 | 0.12644 |
| 256 | Veronica arvensis L. | Veronica arvensis | 0.05 | 0.95 | 0.050 | 6.3 | 5 | 7.3 | 5.8 | 6.5 | 0.7 | 0.53692 | 0.48641 | 1.02734 | 1.80642 | 0.6097 | 0.33182 | 0.26598 |
| 256 | Plantago major L. | Plantago major ssp. major | 0.02 | 0.95 | 0.200 | 7.4 | 5.9 | 3.8 | 6 | NA | 0 | 0.7907 | 0.77966 | 1.64404 | 2.03844 | 0.57393 | 0.17512 | 0.69169 |
| 257 | Poa annua L. | Poa annua | 0.35 | 1 | 0.042 | 7.3 | 5.4 | 4.1 | 7.6 | 4.2 | 0.4 | 0.58728 | 0.56055 | 1.34601 | 1.94534 | 0.82357 | 0.23743 | 0.34603 |
| 257 | Stellaria media (L.) Vill. | Stellaria media agg. | 0.4 | 1 | 0.051 | 7 | 5.6 | 4.4 | 5.9 | 5.4 | 0.1 | 0.64871 | 0.63945 | 1.70759 | 2.05577 | 0.56937 | 0.21649 | 0.54803 |
| 257 | Cerastium fontanum Baumg. | Cerastium fontanum fontanum | 0.13 | 1 | 0.020 | 5.2 | 5.9 | 5.4 | 7.3 | 8 | 0 | 0.67947 | 0.25214 | 0.1495 | 1.33661 | 0.07498 | 0.19025 | 0.12819 |
| 257 | Sagina procumbens L. | Sagina procumbens | 0.05 | 1 | 0.110 | 6.3 | 5 | 7.3 | 5.8 | 6.5 | 0.7 | 0.53692 | 0.48641 | 1.02734 | 1.80642 | 0.6097 | 0.33182 | 0.26598 |
| 257 | Potentilla indica (Andrews) Th.Wolf | NA | 0.07 | 1 | 0.200 | 7.4 | 5.3 | NA | 6 | 7.7 | 0.8 | 0.65201 | 0.63791 | 1.54898 | 2.03286 | 0.77966 | 0.37419 | 0.45721 |
| 258 | Poa annua L. | Poa annua | 0.2 | 0.65 | 0.050 | 6.8 | NA | 5.6 | 5.8 | NA | 0.1 | 0.52169 | 0.45097 | 0.93044 | 1.87015 | 0.67593 | 0.2975 | 0.27054 |
| 258 | Stellaria media (L.) Vill. | Stellaria media agg. | 0.4 | 0.65 | 0.050 | NA | NA | 5.3 | 6.2 | NA | 0.1 | 0.68682 | 0.14816 | 0.05813 | 1.31254 | 0.01433 | 0.20126 | 0.10505 |
| 258 | Sagina procumbens L. | Sagina procumbens | 0.05 | 0.65 | 0.050 | NA | NA | NA | NA | NA | NA | NA | NA | NA | NA | NA | NA | NA |
| 259 | Oxalis corniculata L. | Oxalis corniculata | 0.25 | 1 | 0.333 | 6.8 | NA | 5.6 | 5.8 | NA | 0.1 | 0.52169 | 0.45097 | 0.93044 | 1.87015 | 0.67593 | 0.2975 | 0.27054 |
| 259 | Taraxacum officinale F.H.Wigg. | Taraxacum officinale agg. | 0.15 | 1 | 0.100 | 5.7 | NA | NA | 7.3 | 5.4 | 0 | 0.6918 | 0.13881 | 0.03984 | 1.31841 | 0.01173 | 0.19643 | 0.10363 |
| 259 | Poa annua L. | Poa annua | 0.2 | 1 | 0.050 | NA | NA | NA | NA | NA | NA | NA | NA | NA | NA | NA | NA | NA |
| 259 | NA | NA | 0.1 | 1 | 0.150 | 7.4 | 5.3 | NA | 6 | 7.7 | 0.8 | 0.65201 | 0.63791 | 1.54898 | 2.03286 | 0.77966 | 0.37419 | 0.45721 |
| 259 | Sagina procumbens L. | Sagina procumbens | 0.3 | 1 | 0.400 | 8.1 | NA | 3.9 | 6.9 | 6.4 | 0.4 | 0.80098 | 0.76713 | 1.29016 | 1.92909 | 0.75604 | 0.19458 | 0.52486 |
| 260 | Poa annua L. | Poa annua | 1 | 1 | 0.100 | 7 | 5.6 | 4.4 | 5.9 | 5.4 | 0.1 | 0.64871 | 0.63945 | 1.70759 | 2.05577 | 0.56937 | 0.21649 | 0.54803 |
| 261 | Taraxacum officinale F.H.Wigg. | Taraxacum officinale agg. | 0.1 | 1 | 0.050 | 7.1 | 5.3 | 4.5 | 5.9 | 4.8 | 0.3 | 0.52518 | 0.5113 | 1.61045 | 2.05509 | 0.96495 | 0.27897 | 0.32911 |
| 261 | Poa annua L. | Poa annua | 0.7 | 1 | 0.045 | 7.9 | NA | 4.1 | 5.8 | NA | 0.9 | 0.54139 | 0.51128 | 1.31363 | 1.93633 | 0.8112 | 0.25085 | 0.32953 |
| 261 | Sagina procumbens L. | Sagina procumbens | 0.2 | 1 | 0.045 | 7.9 | NA | 4.1 | 5.8 | NA | 0.9 | 0.54139 | 0.51128 | 1.31363 | 1.93633 | 0.8112 | 0.25085 | 0.32953 |
| 262 | Poa annua L. | Poa annua | 0.1 | 1 | 0.050 | 8.4 | NA | 3 | 6 | NA | 0 | 0.46249 | 0.43915 | 1.39189 | 1.83687 | 0.38437 | 0.30571 | 0.38883 |
| 262 | Capsella bursa-pastoris Medik. | Capsella bursa-pastoris | 0.3 | 1 | 0.100 | 6.7 | NA | NA | 6.5 | 7.7 | 0.1 | 0.70945 | 0.61373 | 0.80583 | 1.7717 | 0.43958 | 0.26577 | 0.51423 |
| 262 | Stellaria media (L.) Vill. | Stellaria media agg. | 0.4 | 1 | 0.050 | NA | NA | 5.3 | 6.2 | NA | 0.1 | 0.68682 | 0.14816 | 0.05813 | 1.31254 | 0.01433 | 0.20126 | 0.10505 |
| 262 | Polygonum aviculare L. | Polygonum aviculare agg. | 0.2 | 1 | 0.056 | 7 | 5.6 | 4.4 | 5.9 | 5.4 | 0.1 | 0.64871 | 0.63945 | 1.70759 | 2.05577 | 0.56937 | 0.21649 | 0.54803 |
| 263 | Poa annua L. | Poa annua | 0.5 | 0.95 | 0.200 | NA | NA | NA | 7 | NA | 0 | 0.66957 | 0.20824 | 0.12967 | 1.34554 | 0.07292 | 0.20428 | 0.13248 |
| 263 | Taraxacum officinale F.H.Wigg. | Taraxacum officinale agg. | 0.1 | 0.95 | 0.150 | 7.3 | 5.4 | 4.1 | 7.6 | 4.2 | 0.4 | 0.58728 | 0.56055 | 1.34601 | 1.94534 | 0.82357 | 0.23743 | 0.34603 |
| 263 | Medicago lupulina L. | Medicago lupulina | 0.2 | 0.95 | 0.050 | NA | NA | NA | NA | NA | NA | NA | NA | NA | NA | NA | NA | NA |
| 263 | Bromus sterilis L. | Bromus sterilis | 0.1 | 0.95 | 0.100 | 7.3 | 5.4 | 4.1 | 7.6 | 4.2 | 0.4 | 0.58728 | 0.56055 | 1.34601 | 1.94534 | 0.82357 | 0.23743 | 0.34603 |
| 263 | Sagina procumbens L. | Sagina procumbens | 0.05 | 0.95 | 0.263 | 7.3 | NA | NA | 7.6 | 7.7 | 0.4 | 0.75479 | 0.72524 | 1.28692 | 1.77374 | 0.61324 | 0.18283 | 0.56773 |
| 264 | Taraxacum officinale F.H.Wigg. | Taraxacum officinale agg. | 0.1 | 0.9 | 0.176 | 7.4 | 5.3 | NA | 6 | 7.7 | 0.8 | 0.65201 | 0.63791 | 1.54898 | 2.03286 | 0.77966 | 0.37419 | 0.45721 |
| 264 | Bromus sterilis L. | Bromus sterilis | 0.8 | 0.9 | 0.111 | 8.4 | NA | 4 | 6.3 | NA | 0.1 | 0.76525 | 0.73026 | 1.22636 | 1.78874 | 0.7034 | 0.16135 | 0.51807 |
| 265 | Sagina procumbens L. | Sagina procumbens | 0.1 | 0.9 | 0.040 | 7.4 | 5.9 | 3.8 | 6 | NA | 0 | 0.7907 | 0.77966 | 1.64404 | 2.03844 | 0.57393 | 0.17512 | 0.69169 |
| 265 | Taraxacum officinale F.H.Wigg. | Taraxacum officinale agg. | 0.1 | 0.9 | 0.080 | 7 | 5.6 | 4.4 | 5.9 | 5.4 | 0.1 | 0.64871 | 0.63945 | 1.70759 | 2.05577 | 0.56937 | 0.21649 | 0.54803 |
| 265 | Hordeum murinum L. | Hordeum murinum agg. | 0.7 | 0.9 | 0.013 | 8.4 | NA | 3 | 6 | NA | 0 | 0.46249 | 0.43915 | 1.39189 | 1.83687 | 0.38437 | 0.30571 | 0.38883 |
| 266 | Sagina procumbens L. | Sagina procumbens | 0.05 | 0.95 | 0.147 | 8.1 | NA | NA | 4.4 | 2.8 | 0.3 | 0.48868 | 0.46508 | 1.4426 | 2.04965 | 0.7278 | 0.33931 | 0.35628 |
| 266 | Stellaria media (L.) Vill. | Stellaria media agg. | 0.05 | 0.95 | 0.133 | 6.8 | NA | 5.6 | 5.8 | NA | 0.1 | 0.52169 | 0.45097 | 0.93044 | 1.87015 | 0.67593 | 0.2975 | 0.27054 |
| 266 | Hordeum murinum L. | Hordeum murinum agg. | 0.85 | 0.95 | 0.020 | 7.3 | 5.4 | 4.1 | 7.6 | 4.2 | 0.4 | 0.58728 | 0.56055 | 1.34601 | 1.94534 | 0.82357 | 0.23743 | 0.34603 |
| 267 | Hordeum murinum L. | Hordeum murinum agg. | 0.2 | 0.85 | 0.211 | 7.5 | NA | NA | 7 | NA | 0 | 0.62073 | 0.22498 | 0.24103 | 1.34507 | 0.09374 | 0.21377 | 0.12505 |
| 267 | Poa annua L. | Poa annua | 0.5 | 0.85 | 0.180 | 8.1 | NA | NA | 4.4 | 2.8 | 0.3 | 0.48868 | 0.46508 | 1.4426 | 2.04965 | 0.7278 | 0.33931 | 0.35628 |
| 267 | Stellaria media (L.) Vill. | Stellaria media agg. | 0.05 | 0.85 | 0.050 | 6.7 | NA | NA | 6.5 | 7.7 | 0.1 | 0.70945 | 0.61373 | 0.80583 | 1.7717 | 0.43958 | 0.26577 | 0.51423 |
| 267 | Capsella bursa-pastoris Medik. | Capsella bursa-pastoris | 0.1 | 0.85 | 0.102 | 8.1 | NA | NA | 4.4 | 2.8 | 0.3 | 0.48868 | 0.46508 | 1.4426 | 2.04965 | 0.7278 | 0.33931 | 0.35628 |
| 268 | Capsella bursa-pastoris Medik. | Capsella bursa-pastoris | 0.9 | 1 | 0.043 | 7.3 | NA | 8 | 7.3 | 7 | 0.3 | 0.46459 | 0.41665 | 0.98972 | 1.42624 | 0.46979 | 0.18153 | 0.14439 |
| 268 | Poa annua L. | Poa annua | 0.05 | 1 | 0.110 | 7.3 | 6.1 | 3.8 | 6 | 5.1 | 0 | 0.62987 | 0.59281 | 1.27995 | 1.90491 | 0.6275 | 0.3036 | 0.42608 |
| 268 | Stellaria media (L.) Vill. | Stellaria media agg. | 0.05 | 1 | 0.091 | NA | NA | NA | NA | NA | NA | NA | NA | NA | NA | NA | NA | NA |
| 268 | NA | NA | 0 | 1 | 0.080 | 7 | NA | NA | NA | NA | 0.3 | 0.56899 | 0.4288 | 0.64817 | 1.63031 | 0.49671 | 0.22558 | 0.25973 |
| 269 | Poa annua L. | Poa annua | 0.35 | 1 | 0.040 | 7.3 | 5.4 | 4.1 | 7.6 | 4.2 | 0.4 | 0.58728 | 0.56055 | 1.34601 | 1.94534 | 0.82357 | 0.23743 | 0.34603 |
| 269 | Taraxacum officinale F.H.Wigg. | Taraxacum officinale agg. | 0.35 | 1 | 0.200 | 8.1 | NA | 3.9 | 6.9 | 6.4 | 0.4 | 0.80098 | 0.76713 | 1.29016 | 1.92909 | 0.75604 | 0.19458 | 0.52486 |
| 269 | Stellaria media (L.) Vill. | Stellaria media agg. | 0.3 | 1 | 0.080 | 7.3 | 5.4 | 4.1 | 7.6 | 4.2 | 0.4 | 0.58728 | 0.56055 | 1.34601 | 1.94534 | 0.82357 | 0.23743 | 0.34603 |
| 270 | Poa annua L. | Poa annua | 0.9 | 1 | 0.134 | 6.6 | 5.6 | NA | 5.6 | 4.1 | 0.1 | 0.50969 | 0.50576 | 1.97957 | 2.23767 | 1.1694 | 0.28385 | 0.36995 |
| 270 | Jacobaea analoga (DC.) Veldkamp | Senecio jacobaea | 0.1 | 1 | 0.100 | 6.6 | 5.6 | NA | 5.6 | 4.1 | 0.1 | 0.50969 | 0.50576 | 1.97957 | 2.23767 | 1.1694 | 0.28385 | 0.36995 |
| 271 | Jacobaea analoga (DC.) Veldkamp | Senecio jacobaea | 0.4 | 1 | 0.040 | 7.3 | 5.4 | 4.1 | 7.6 | 4.2 | 0.4 | 0.58728 | 0.56055 | 1.34601 | 1.94534 | 0.82357 | 0.23743 | 0.34603 |
| 271 | Taraxacum officinale F.H.Wigg. | Taraxacum officinale agg. | 0.4 | 1 | 0.167 | NA | NA | NA | NA | NA | NA | NA | NA | NA | NA | NA | NA | NA |
| 271 | Poa annua L. | Poa annua | 0.2 | 1 | 0.081 | 7.2 | 5.5 | 4.5 | 6 | 7.6 | 0.3 | 0.72905 | 0.70188 | 1.35981 | 1.84158 | 0.57912 | 0.20081 | 0.60725 |
| 272 | Helianthus annuus L. | NA | 0.3 | 0.75 | 0.400 | 7.4 | 5.3 | NA | 6 | 7.7 | 0.8 | 0.65201 | 0.63791 | 1.54898 | 2.03286 | 0.77966 | 0.37419 | 0.45721 |
| 272 | Taraxacum officinale F.H.Wigg. | Taraxacum officinale agg. | 0.2 | 0.75 | 0.250 | NA | NA | NA | 7 | NA | 0 | 0.66957 | 0.20824 | 0.12967 | 1.34554 | 0.07292 | 0.20428 | 0.13248 |
| 272 | Stellaria media (L.) Vill. | Stellaria media agg. | 0.1 | 0.75 | 0.023 | 7.3 | 5.8 | 4.8 | 5.3 | 5.6 | 0 | 0.67083 | 0.63295 | 1.24583 | 1.99939 | 0.56795 | 0.28417 | 0.48569 |
| 272 | Poa annua L. | Poa annua | 0.15 | 0.75 | 0.353 | 8 | NA | NA | 6.1 | 6.3 | 0.1 | 0.56333 | 0.55394 | 1.72195 | 2.16326 | 1.07172 | 0.43309 | 0.38415 |
| 273 | Stellaria media (L.) Vill. | Stellaria media agg. | 0.6 | 0.8 | 0.050 | 7.6 | 5.3 | 3.9 | 7 | 5 | 0.2 | 0.63304 | 0.57145 | 1.00353 | 1.70929 | 0.62485 | 0.18325 | 0.34548 |
| 273 | Hedera helix L. | Hedera helix | 0.2 | 0.8 | 0.538 | 7.4 | 5.3 | NA | 6 | 7.7 | 0.8 | 0.65201 | 0.63791 | 1.54898 | 2.03286 | 0.77966 | 0.37419 | 0.45721 |
| 274 | Taraxacum officinale F.H.Wigg. | Taraxacum officinale agg. | 0.2 | 1 | 0.111 | NA | 5.4 | 5.7 | 6.8 | 7 | 0 | 0.67742 | 0.24559 | 0.14546 | 1.36309 | 0.06547 | 0.19208 | 0.12644 |
| 274 | Poa annua L. | Poa annua | 0.5 | 1 | 0.067 | 7.9 | 5.7 | 4.2 | 7 | 7.2 | 0.1 | 0.76998 | 0.7507 | 1.46055 | 1.90983 | 0.91556 | 0.1857 | 0.55131 |
| 274 | Veronica hederifolia L. | Veronica hederifolia agg. | 0.1 | 1 | 0.100 | 7.2 | 5.7 | 4.3 | 6.8 | NA | 0.1 | 0.54405 | 0.41742 | 0.70503 | 1.63482 | 0.49706 | 0.18636 | 0.24515 |
| 274 | Chelidonium majus L. | Chelidonium majus | 0.2 | 1 | 0.056 | 6.5 | 6.7 | 5.3 | 6.7 | 6.7 | 0 | 0.6884 | 0.39438 | 0.41506 | 1.39788 | 0.17506 | 0.19382 | 0.19382 |
| 275 | Trifolium repens L. | Trifolium repens | 0.6 | 1 | 0.053 | 6.5 | 6.7 | 5.3 | 6.7 | 6.7 | 0 | 0.6884 | 0.39438 | 0.41506 | 1.39788 | 0.17506 | 0.19382 | 0.19382 |
| 275 | Poa annua L. | Poa annua | 0.4 | 1 | 0.100 | 8.3 | 5 | NA | 7.2 | 7.8 | 0.1 | 0.67936 | 0.56623 | 0.73548 | 1.48508 | 0.50389 | 0.1867 | 0.27262 |
| 276 | Poa annua L. | Poa annua | 0.5 | 1 | 0.368 | 7.4 | 5.3 | NA | 6 | 7.7 | 0.8 | 0.65201 | 0.63791 | 1.54898 | 2.03286 | 0.77966 | 0.37419 | 0.45721 |
| 276 | Trifolium repens L. | Trifolium repens | 0.2 | 1 | 0.450 | 7.4 | 5.3 | NA | 6 | 7.7 | 0.8 | 0.65201 | 0.63791 | 1.54898 | 2.03286 | 0.77966 | 0.37419 | 0.45721 |
| 276 | Prunella vulgaris L. | Prunella vulgaris | 0.1 | 1 | 0.100 | 7.3 | 5.4 | 4.1 | 7.6 | 4.2 | 0.4 | 0.58728 | 0.56055 | 1.34601 | 1.94534 | 0.82357 | 0.23743 | 0.34603 |
| 276 | Ranunculus repens L. | Ranunculus repens | 0.1 | 1 | 0.050 | 7.9 | 6.2 | NA | 6.7 | 6.4 | 0.2 | 0.62749 | 0.58201 | 1.11218 | 1.53794 | 0.69579 | 0.21134 | 0.22838 |
| 276 | Taraxacum officinale F.H.Wigg. | Taraxacum officinale agg. | 0.05 | 1 | 0.083 | 8.2 | NA | 3.5 | 5.7 | 3.2 | 0 | 0.46142 | 0.41736 | 1.14878 | 1.78546 | 0.38405 | 0.27764 | 0.246 |
| 276 | Glechoma hederacea L. | Glechoma hederacea | 0.05 | 1 | 0.111 | 7.3 | 5.4 | 4.1 | 7.6 | 4.2 | 0.4 | 0.58728 | 0.56055 | 1.34601 | 1.94534 | 0.82357 | 0.23743 | 0.34603 |
| 277 | Glechoma hederacea L. | Glechoma hederacea | 0.1 | 1 | 0.215 | 7.3 | 5.4 | 4.1 | 7.6 | 4.2 | 0.4 | 0.58728 | 0.56055 | 1.34601 | 1.94534 | 0.82357 | 0.23743 | 0.34603 |
| 277 | Trifolium repens L. | Trifolium repens | 0.45 | 1 | 0.024 | 7 | 5.6 | 4.4 | 5.9 | 5.4 | 0.1 | 0.64871 | 0.63945 | 1.70759 | 2.05577 | 0.56937 | 0.21649 | 0.54803 |
| 277 | Poa annua L. | Poa annua | 0.4 | 1 | 0.054 | 7.7 | NA | NA | 5.6 | NA | 0.7 | 0.53223 | 0.52325 | 1.73719 | 2.14421 | 0.94278 | 0.39374 | 0.35054 |
| 277 | Bellis perennis L. | Bellis perennis | 0.05 | 1 | 0.115 | 7.3 | 5.4 | 4.1 | 7.6 | 4.2 | 0.4 | 0.58728 | 0.56055 | 1.34601 | 1.94534 | 0.82357 | 0.23743 | 0.34603 |
| 278 | Poa annua L. | Poa annua | 0.95 | 1 | 0.011 | 8.5 | NA | 2.5 | NA | NA | 0 | 0.506 | 0.44835 | 1.02121 | 1.67002 | 0.34316 | 0.25824 | 0.34742 |
| 278 | Trifolium repens L. | Trifolium repens | 0.05 | 1 | 0.060 | 7.3 | 5.4 | 4.1 | 7.6 | 4.2 | 0.4 | 0.58728 | 0.56055 | 1.34601 | 1.94534 | 0.82357 | 0.23743 | 0.34603 |
| 279 | Hordeum murinum L. | Hordeum murinum agg. | 0.95 | 1 | 0.020 | 7.3 | 5.4 | 4.1 | 7.6 | 4.2 | 0.4 | 0.58728 | 0.56055 | 1.34601 | 1.94534 | 0.82357 | 0.23743 | 0.34603 |
| 279 | Taraxacum officinale F.H.Wigg. | Taraxacum officinale agg. | 0.05 | 1 | 0.150 | 7.3 | 6.1 | 3.8 | 6 | 5.1 | 0 | 0.62987 | 0.59281 | 1.27995 | 1.90491 | 0.6275 | 0.3036 | 0.42608 |
| 281 | Poa annua L. | Poa annua | 1 | 1 | 0.050 | 8.8 | NA | 3.9 | 7.5 | 7.4 | 0.2 | 0.71714 | 0.69501 | 1.38943 | 1.78673 | 0.86446 | 0.18028 | 0.37675 |
| 282 | Taraxacum officinale F.H.Wigg. | Taraxacum officinale agg. | 0.85 | 0.9 | 0.111 | 6.3 | NA | 5.9 | 6.3 | 6.9 | 0.1 | 0.6037 | 0.3946 | 0.44368 | 1.50139 | 0.30164 | 0.21315 | 0.21494 |
| 282 | Poa annua L. | Poa annua | 0.05 | 0.9 | 0.071 | NA | NA | NA | NA | NA | NA | NA | NA | NA | NA | NA | NA | NA |
| 283 | Poa annua L. | Poa annua | 0.3 | 0.4 | 0.021 | 7.3 | 5.4 | 4.1 | 7.6 | 4.2 | 0.4 | 0.58728 | 0.56055 | 1.34601 | 1.94534 | 0.82357 | 0.23743 | 0.34603 |
| 283 | Stellaria media (L.) Vill. | Stellaria media agg. | 0.1 | 0.4 | 0.200 | 7.4 | 5.3 | NA | 6 | 7.7 | 0.8 | 0.65201 | 0.63791 | 1.54898 | 2.03286 | 0.77966 | 0.37419 | 0.45721 |
| 284 | Bromus sterilis L. | Bromus sterilis | 0.5 | 1 | 0.050 | 7.4 | 5.3 | NA | 6 | 7.7 | 0.8 | 0.65201 | 0.63791 | 1.54898 | 2.03286 | 0.77966 | 0.37419 | 0.45721 |
| 284 | Veronica hederifolia L. | Veronica hederifolia agg. | 0.05 | 1 | 0.100 | 7.4 | 5.3 | NA | 6 | 7.7 | 0.8 | 0.65201 | 0.63791 | 1.54898 | 2.03286 | 0.77966 | 0.37419 | 0.45721 |
| 284 | Sisymbrium officinale (L.) Scop. | Sisymbrium officinale | 0.2 | 1 | 0.050 | 8.6 | NA | 2.7 | 4.4 | 1.6 | 0 | 0.53557 | 0.50485 | 1.29532 | 1.90211 | 0.72298 | 0.28495 | 0.2778 |
| 284 | Taraxacum officinale F.H.Wigg. | Taraxacum officinale agg. | 0.1 | 1 | 0.050 | 8.4 | NA | 3 | 6 | NA | 0 | 0.46249 | 0.43915 | 1.39189 | 1.83687 | 0.38437 | 0.30571 | 0.38883 |
| 284 | Poa annua L. | Poa annua | 0.1 | 1 | 0.242 | 7.4 | 5.3 | NA | 6 | 7.7 | 0.8 | 0.65201 | 0.63791 | 1.54898 | 2.03286 | 0.77966 | 0.37419 | 0.45721 |
| 284 | Stellaria media (L.) Vill. | Stellaria media agg. | 0.05 | 1 | 0.020 | 6.7 | 5.3 | 4.5 | 5.3 | 5.8 | 0 | 0.75464 | 0.73051 | 1.36974 | 1.95508 | 0.45974 | 0.15567 | 0.6659 |
| 285 | Stellaria media (L.) Vill. | Stellaria media agg. | 0.05 | 1 | 0.050 | 8.6 | NA | 2.7 | 4.4 | 1.6 | 0 | 0.53557 | 0.50485 | 1.29532 | 1.90211 | 0.72298 | 0.28495 | 0.2778 |
| 285 | Taraxacum officinale F.H.Wigg. | Taraxacum officinale agg. | 0.1 | 1 | 0.330 | 7.4 | 5.3 | NA | 6 | 7.7 | 0.8 | 0.65201 | 0.63791 | 1.54898 | 2.03286 | 0.77966 | 0.37419 | 0.45721 |
| 285 | Trifolium repens L. | Trifolium repens | 0.2 | 1 | 0.100 | 8.4 | NA | 3 | 6 | NA | 0 | 0.46249 | 0.43915 | 1.39189 | 1.83687 | 0.38437 | 0.30571 | 0.38883 |
| 285 | Veronica hederifolia L. | Veronica hederifolia agg. | 0.1 | 1 | 0.500 | 7.4 | 5.3 | NA | 6 | 7.7 | 0.8 | 0.65201 | 0.63791 | 1.54898 | 2.03286 | 0.77966 | 0.37419 | 0.45721 |
| 285 | Bromus sterilis L. | Bromus sterilis | 0.44 | 1 | 0.020 | 5.2 | 5.9 | 5.4 | 7.3 | 8 | 0 | 0.67947 | 0.25214 | 0.1495 | 1.33661 | 0.07498 | 0.19025 | 0.12819 |

| | | | | | | | | | | | | | | | | | | |
|---|---|---|---|---|---|---|---|---|---|---|---|---|---|---|---|---|---|---|
| 285 | Medicago lupulina L. | Medicago lupulina | 0.05 | 1 | 0.340 | 7.4 | 5.3 | NA | 6 | 7.7 | 0.8 | 0.65201 | 0.63791 | 1.54898 | 2.03286 | 0.77966 | 0.37419 | 0.45721 |
| 285 | Veronica arvensis L. | Veronica arvensis | 0.01 | 1 | 0.050 | 6.7 | 5.3 | 4.5 | 5.3 | 5.8 | 0 | 0.75464 | 0.73051 | 1.36974 | 1.95508 | 0.45974 | 0.15567 | 0.6659 |
| 285 | Sisymbrium officinale (L.) Scop. | Sisymbrium officinale | 0.05 | 1 | 0.500 | NA | NA | NA | NA | NA | NA | NA | NA | NA | NA | NA | NA | NA |
| 286 | Ranunculus repens L. | Ranunculus repens | 0.3 | 1 | 0.053 | 5.2 | 5.9 | 5.4 | 7.3 | 8 | 0 | 0.67947 | 0.25214 | 0.1495 | 1.33661 | 0.07498 | 0.19025 | 0.12819 |
| 286 | Trifolium pratense L. | Trifolium pratense | 0.07 | 1 | 0.200 | 7.4 | 5.3 | NA | 6 | 7.7 | 0.8 | 0.65201 | 0.63791 | 1.54898 | 2.03286 | 0.77966 | 0.37419 | 0.45721 |
| 286 | Glechoma hederacea L. | Glechoma hederacea | 0.05 | 1 | 0.200 | NA | NA | NA | NA | NA | NA | NA | NA | NA | NA | NA | NA | NA |
| 286 | Urtica dioica L. | Urtica dioica | 0.25 | 1 | 0.400 | NA | NA | NA | NA | NA | NA | NA | NA | NA | NA | NA | NA | NA |
| 286 | Rumex obtusifolius L. | Rumex obtusifolius | 0.1 | 1 | 0.444 | NA | NA | NA | NA | NA | NA | NA | NA | NA | NA | NA | NA | NA |
| 286 | Poa annua L. | Poa annua | 0.05 | 1 | 0.133 | 6.7 | NA | NA | 6.5 | 7.7 | 0.1 | 0.70945 | 0.61373 | 0.80583 | 1.7717 | 0.43958 | 0.26577 | 0.51423 |
| 286 | Taraxacum officinale F.H.Wigg. | Taraxacum officinale agg. | 0.01 | 1 | 0.333 | 7.4 | 5.3 | NA | 6 | 7.7 | 0.8 | 0.65201 | 0.63791 | 1.54898 | 2.03286 | 0.77966 | 0.37419 | 0.45721 |
| 286 | Dactylis glomerata L. | Dactylis glomerata | 0.1 | 1 | 0.397 | 7.4 | 5.3 | NA | 6 | 7.7 | 0.8 | 0.65201 | 0.63791 | 1.54898 | 2.03286 | 0.77966 | 0.37419 | 0.45721 |
| 286 | Plantago major L. | Plantago major ssp. major | 0.07 | 1 | 0.040 | 8.4 | 7.2 | 2.6 | 6.8 | 3.6 | 0.2 | 0.58203 | 0.49726 | 0.89718 | 1.5951 | 0.3257 | 0.17939 | 0.36267 |
| 287 | Taraxacum officinale F.H.Wigg. | Taraxacum officinale agg. | 0.1 | 0.8 | 0.031 | 7.3 | 6.1 | 3.8 | 6 | 5.1 | 0 | 0.62987 | 0.59281 | 1.27995 | 1.90491 | 0.6275 | 0.3036 | 0.42608 |
| 287 | Medicago lupulina L. | Medicago lupulina | 0.1 | 0.8 | 0.020 | 7.3 | 5.4 | 4.1 | 7.6 | 4.2 | 0.4 | 0.58728 | 0.56055 | 1.34601 | 1.94534 | 0.82357 | 0.23743 | 0.34603 |
| 287 | Sisymbrium officinale (L.) Scop. | Sisymbrium officinale | 0.05 | 0.8 | 0.020 | 8.1 | NA | NA | 5.2 | NA | 0.1 | 0.51276 | 0.49581 | 1.51667 | 2.20989 | 0.98959 | 0.46997 | 0.3594 |
| 287 | Senecio vulgaris L. | Senecio vulgaris | 0.05 | 0.8 | 0.050 | 7.6 | NA | NA | 6.5 | 6.3 | 0.7 | 0.75548 | 0.75048 | 1.8409 | 2.08419 | 0.63746 | 0.23177 | 0.58814 |
| 287 | Bromus sterilis L. | Bromus sterilis | 0.2 | 0.8 | 0.200 | 7.4 | 5.3 | NA | 6 | 7.7 | 0.8 | 0.65201 | 0.63791 | 1.54898 | 2.03286 | 0.77966 | 0.37419 | 0.45721 |
| 287 | Stellaria media (L.) Vill. | Stellaria media agg. | 0.05 | 0.8 | 0.150 | 7 | NA | NA | NA | NA | 0.3 | 0.56899 | 0.4288 | 0.64817 | 1.63031 | 0.49671 | 0.22558 | 0.25973 |
| 287 | Poa annua L. | Poa annua | 0.05 | 0.8 | 0.100 | NA | NA | NA | NA | NA | NA | NA | NA | NA | NA | NA | NA | NA |
| 287 | Hordeum murinum L. | Hordeum murinum agg. | 0.2 | 0.8 | 0.138 | 6.8 | NA | 5.6 | 5.8 | NA | 0.1 | 0.52169 | 0.45097 | 0.93044 | 1.87015 | 0.67593 | 0.2975 | 0.27054 |
| 288 | Glechoma hederacea L. | Glechoma hederacea | 0.1 | 1 | 0.105 | NA | NA | NA | NA | NA | NA | NA | NA | NA | NA | NA | NA | NA |
| 288 | Veronica hederifolia L. | Veronica hederifolia agg. | 0.05 | 1 | 0.100 | NA | NA | NA | NA | NA | NA | NA | NA | NA | NA | NA | NA | NA |
| 288 | Sisymbrium officinale (L.) Scop. | Sisymbrium officinale | 0.15 | 1 | 0.080 | 7.3 | NA | 3.9 | 6.1 | 3.9 | 0.1 | 0.55903 | 0.41538 | 0.65189 | 1.65181 | 0.43719 | 0.24279 | 0.21709 |
| 288 | Taraxacum officinale F.H.Wigg. | Taraxacum officinale agg. | 0.15 | 1 | 0.150 | 7.7 | NA | NA | 5.6 | NA | 0.7 | 0.53223 | 0.52325 | 1.73719 | 2.14421 | 0.94278 | 0.39374 | 0.35054 |
| 288 | Geum urbanum L. | Geum urbanum | 0.1 | 1 | 0.040 | 5.7 | NA | NA | 6.4 | 7.4 | 0 | 0.68859 | 0.26809 | 0.17316 | 1.34857 | 0.09545 | 0.19552 | 0.14349 |
| 288 | Medicago lupulina L. | Medicago lupulina | 0.05 | 1 | 0.050 | 5.7 | NA | NA | 6.4 | 7.4 | 0 | 0.68859 | 0.26809 | 0.17316 | 1.34857 | 0.09545 | 0.19552 | 0.14349 |
| 288 | Bromus sterilis L. | Bromus sterilis | 0.3 | 1 | 0.097 | 6.7 | NA | NA | 5.2 | NA | 0.4 | 0.59706 | 0.49429 | 0.69366 | 1.8006 | 0.482 | 0.19483 | 0.19138 |
| 288 | Poa annua L. | Poa annua | 0.1 | 1 | 0.062 | 5.7 | NA | NA | 6.4 | 7.4 | 0 | 0.68859 | 0.26809 | 0.17316 | 1.34857 | 0.09545 | 0.19552 | 0.14349 |
| 289 | Veronica hederifolia L. | Veronica hederifolia agg. | 0.05 | 1 | 0.038 | 6.7 | NA | 4.9 | 6.8 | 5 | 0 | 0.66057 | 0.18432 | 0.11528 | 1.34075 | 0.06671 | 0.20032 | 0.12881 |
| 289 | Stellaria media (L.) Vill. | Stellaria media agg. | 0.05 | 1 | 0.100 | 6.8 | NA | 5.6 | 5.8 | NA | 0.1 | 0.52169 | 0.45097 | 0.93044 | 1.87015 | 0.67593 | 0.2975 | 0.27054 |
| 289 | Taraxacum officinale F.H.Wigg. | Taraxacum officinale agg. | 0.12 | 1 | 0.020 | 7.7 | NA | 4.6 | 4 | 3.9 | 0.4 | 0.73 | 0.72198 | 1.79063 | 2.10331 | 0.40223 | 0.22254 | 0.54071 |
| 289 | Poa annua L. | Poa annua | 0.08 | 1 | 0.610 | 7.4 | 5.3 | NA | 6 | 7.7 | 0.8 | 0.65201 | 0.63791 | 1.54898 | 2.03286 | 0.77966 | 0.37419 | 0.45721 |
| 289 | Sisymbrium officinale (L.) Scop. | Sisymbrium officinale | 0.15 | 1 | 0.020 | 7.1 | NA | NA | NA | 4.6 | 0 | 0.69216 | 0.66495 | 1.32487 | 1.98453 | 0.49303 | 0.16764 | 0.56484 |
| 289 | Bromus sterilis L. | Bromus sterilis | 0.25 | 1 | 0.023 | 7.1 | NA | NA | NA | 4.6 | 0 | 0.69216 | 0.66495 | 1.32487 | 1.98453 | 0.49303 | 0.16764 | 0.56484 |
| 289 | Hordeum murinum L. | Hordeum murinum agg. | 0.25 | 1 | 0.240 | 7.4 | 5.3 | NA | 6 | 7.7 | 0.8 | 0.65201 | 0.63791 | 1.54898 | 2.03286 | 0.77966 | 0.37419 | 0.45721 |
| 289 | Medicago lupulina L. | Medicago lupulina | 0.05 | 1 | 0.021 | 7 | 5.6 | 4.4 | 5.9 | 5.4 | 0.1 | 0.64871 | 0.63945 | 1.70759 | 2.05577 | 0.56937 | 0.21649 | 0.54803 |
| 290 | Geranium molle L. | Geranium molle | 0.05 | 1 | 0.350 | NA | NA | NA | NA | NA | NA | NA | NA | NA | NA | NA | NA | NA |
| 290 | Sisymbrium officinale (L.) Scop. | Sisymbrium officinale | 0.15 | 1 | 0.020 | 6.8 | 6.2 | 4.4 | NA | NA | 0.3 | 0.7871 | 0.75023 | 1.37944 | 1.92919 | 0.40631 | 0.19607 | 0.63681 |
| 290 | Stellaria media (L.) Vill. | Stellaria media agg. | 0.1 | 1 | 0.050 | 7.1 | 5.3 | 4.5 | 5.9 | 4.8 | 0.3 | 0.52518 | 0.5113 | 1.61045 | 2.05509 | 0.96495 | 0.27897 | 0.32911 |
| 290 | Taraxacum officinale F.H.Wigg. | Taraxacum officinale agg. | 0.2 | 1 | 0.062 | NA | NA | NA | NA | NA | NA | NA | NA | NA | NA | NA | NA | NA |
| 290 | Poa annua L. | Poa annua | 0.1 | 1 | 0.588 | 7.4 | 5.3 | NA | 6 | 7.7 | 0.8 | 0.65201 | 0.63791 | 1.54898 | 2.03286 | 0.77966 | 0.37419 | 0.45721 |
| 290 | Bromus sterilis L. | Bromus sterilis | 0.4 | 1 | 0.050 | 8.2 | NA | 2.1 | 6.6 | 1.7 | 0.4 | 0.4561 | 0.42853 | 1.29025 | 1.77233 | 0.32666 | 0.30641 | 0.32553 |
| 291 | Alliaria petiolata (M.Bieb.) | Alliaria petiolata | 0.02 | 0.75 | 0.390 | 7.4 | 5.3 | NA | 6 | 7.7 | 0.8 | 0.65201 | 0.63791 | 1.54898 | 2.03286 | 0.77966 | 0.37419 | 0.45721 |
| 291 | Veronica hederifolia L. | Veronica hederifolia agg. | 0.02 | 0.75 | 0.040 | 7.1 | 5.3 | 4.5 | 5.9 | 4.8 | 0.3 | 0.52518 | 0.5113 | 1.61045 | 2.05509 | 0.96495 | 0.27897 | 0.32911 |
| 291 | Bromus sterilis L. | Bromus sterilis | 0.15 | 0.75 | 0.053 | 7.3 | NA | 3.9 | 6.1 | 3.9 | 0.1 | 0.55903 | 0.41538 | 0.65189 | 1.65181 | 0.43719 | 0.24279 | 0.21709 |
| 291 | Poa annua L. | Poa annua | 0.12 | 0.75 | 0.108 | 7.3 | NA | 3.9 | 6.1 | 3.9 | 0.1 | 0.55903 | 0.41538 | 0.65189 | 1.65181 | 0.43719 | 0.24279 | 0.21709 |
| 291 | Stellaria media (L.) Vill. | Stellaria media agg. | 0.13 | 0.75 | 0.150 | 8 | NA | 3.7 | NA | 2 | 0 | 0.55827 | 0.46888 | 0.89651 | 1.79355 | 0.39043 | 0.26528 | 0.38307 |
| 291 | Urtica dioica L. | Urtica dioica | 0.05 | 0.75 | 0.453 | 7.4 | 5.3 | NA | 6 | 7.7 | 0.8 | 0.65201 | 0.63791 | 1.54898 | 2.03286 | 0.77966 | 0.37419 | 0.45721 |
| 291 | Sisymbrium officinale (L.) Scop. | Sisymbrium officinale | 0.02 | 0.75 | 0.400 | 7.4 | 5.3 | NA | 6 | 7.7 | 0.8 | 0.65201 | 0.63791 | 1.54898 | 2.03286 | 0.77966 | 0.37419 | 0.45721 |
| 291 | Trifolium repens L. | Trifolium repens | 0.07 | 0.75 | 0.033 | 7.3 | NA | 3.9 | 6.1 | 3.9 | 0.1 | 0.55903 | 0.41538 | 0.65189 | 1.65181 | 0.43719 | 0.24279 | 0.21709 |
| 291 | Taraxacum officinale F.H.Wigg. | Taraxacum officinale agg. | 0.1 | 0.75 | 0.022 | 8.1 | NA | NA | 5.2 | NA | 0.1 | 0.51276 | 0.49581 | 1.51667 | 2.20989 | 0.98959 | 0.46997 | 0.3594 |
| 291 | Plantago major L. | Plantago major ssp. major | 0.07 | 0.75 | 0.510 | 7.4 | 5.3 | NA | 6 | 7.7 | 0.8 | 0.65201 | 0.63791 | 1.54898 | 2.03286 | 0.77966 | 0.37419 | 0.45721 |
| 292 | Poa annua L. | Poa annua | 0.25 | 1 | 0.404 | 7.4 | 5.3 | NA | 6 | 7.7 | 0.8 | 0.65201 | 0.63791 | 1.54898 | 2.03286 | 0.77966 | 0.37419 | 0.45721 |
| 292 | Sisymbrium officinale (L.) Scop. | Sisymbrium officinale | 0.15 | 1 | 0.050 | 8.4 | NA | 3 | 6 | NA | 0 | 0.46249 | 0.43915 | 1.39189 | 1.83687 | 0.38437 | 0.30571 | 0.38883 |
| 292 | Veronica hederifolia L. | Veronica hederifolia agg. | 0.05 | 1 | 0.050 | 8.5 | NA | 3.5 | 6.3 | 4.1 | 0.2 | 0.57633 | 0.55499 | 1.44489 | 1.81436 | 0.579 | 0.25616 | 0.27657 |
| 292 | Stellaria media (L.) Vill. | Stellaria media agg. | 0.15 | 1 | 0.260 | 7.4 | 5.3 | NA | 6 | 7.7 | 0.8 | 0.65201 | 0.63791 | 1.54898 | 2.03286 | 0.77966 | 0.37419 | 0.45721 |
| 292 | Taraxacum officinale F.H.Wigg. | Taraxacum officinale agg. | 0.15 | 1 | 0.082 | 8.1 | NA | NA | 5.2 | NA | 0.1 | 0.51276 | 0.49581 | 1.51667 | 2.20989 | 0.98959 | 0.46997 | 0.3594 |
| 292 | Lolium perenne L. | Lolium perenne | 0.1 | 1 | 0.050 | 8.1 | NA | NA | 4.4 | 2.8 | 0.3 | 0.48868 | 0.46508 | 1.4426 | 2.04965 | 0.7278 | 0.33931 | 0.35628 |
| 292 | Bromus sterilis L. | Bromus sterilis | 0.15 | 1 | 0.080 | 8.1 | NA | NA | 4.4 | 2.8 | 0.3 | 0.48868 | 0.46508 | 1.4426 | 2.04965 | 0.7278 | 0.33931 | 0.35628 |
| 293 | Veronica hederifolia L. | Veronica hederifolia agg. | 0.05 | 1 | 0.100 | 8.1 | NA | NA | 4.4 | 2.8 | 0.3 | 0.48868 | 0.46508 | 1.4426 | 2.04965 | 0.7278 | 0.33931 | 0.35628 |
| 293 | Stellaria media (L.) Vill. | Stellaria media agg. | 0.1 | 1 | 0.052 | 7.8 | 5.3 | 3.8 | 6.6 | 5 | 0.2 | 0.49579 | 0.46867 | 1.36306 | 1.85033 | 0.62791 | 0.29201 | 0.3217 |
| 293 | Taraxacum officinale F.H.Wigg. | Taraxacum officinale agg. | 0.4 | 1 | 0.050 | 8.6 | NA | 2.7 | 4.4 | 1.6 | 0 | 0.53557 | 0.50485 | 1.29532 | 1.90211 | 0.72298 | 0.28495 | 0.2778 |
| 293 | Poa annua L. | Poa annua | 0.25 | 1 | 0.040 | 7.1 | 5.3 | 4.5 | 5.9 | 4.8 | 0.3 | 0.52518 | 0.5113 | 1.61045 | 2.05509 | 0.96495 | 0.27897 | 0.32911 |
| 293 | Bromus sterilis L. | Bromus sterilis | 0.2 | 1 | 0.084 | NA | NA | NA | NA | NA | NA | NA | NA | NA | NA | NA | NA | NA |
| 294 | Veronica hederifolia L. | Veronica hederifolia agg. | 0.1 | 0.68 | 0.158 | 8.5 | NA | 3.5 | 6.3 | 4.1 | 0.2 | 0.57633 | 0.55499 | 1.44489 | 1.81436 | 0.579 | 0.25616 | 0.27657 |
| 294 | Stellaria media (L.) Vill. | Stellaria media agg. | 0.23 | 0.68 | 0.051 | 8.1 | NA | NA | 4.4 | 2.8 | 0.3 | 0.48868 | 0.46508 | 1.4426 | 2.04965 | 0.7278 | 0.33931 | 0.35628 |
| 294 | Poa annua L. | Poa annua | 0.25 | 0.68 | 0.440 | 7.4 | 5.3 | NA | 6 | 7.7 | 0.8 | 0.65201 | 0.63791 | 1.54898 | 2.03286 | 0.77966 | 0.37419 | 0.45721 |
| 294 | Bromus sterilis L. | Bromus sterilis | 0.1 | 0.68 | 0.370 | 7.4 | 5.3 | NA | 6 | 7.7 | 0.8 | 0.65201 | 0.63791 | 1.54898 | 2.03286 | 0.77966 | 0.37419 | 0.45721 |
| 295 | Alliaria petiolata (M.Bieb.) | Alliaria petiolata | 0.19 | 1 | 0.010 | 8.6 | NA | 2.7 | 4.4 | 1.6 | 0 | 0.53557 | 0.50485 | 1.29532 | 1.90211 | 0.72298 | 0.28495 | 0.2778 |
| 295 | Veronica hederifolia L. | Veronica hederifolia agg. | 0.19 | 1 | 0.330 | 7.4 | 5.3 | NA | 6 | 7.7 | 0.8 | 0.65201 | 0.63791 | 1.54898 | 2.03286 | 0.77966 | 0.37419 | 0.45721 |
| 295 | Glechoma hederacea L. | Glechoma hederacea | 0.19 | 1 | 0.050 | 6.7 | 5.3 | 4.5 | 5.3 | 5.8 | 0 | 0.75464 | 0.73051 | 1.36974 | 1.95508 | 0.45974 | 0.15567 | 0.6659 |
| 295 | Taraxacum officinale F.H.Wigg. | Taraxacum officinale agg. | 0.05 | 1 | 0.140 | 7.4 | 5.3 | NA | 6 | 7.7 | 0.8 | 0.65201 | 0.63791 | 1.54898 | 2.03286 | 0.77966 | 0.37419 | 0.45721 |
| 295 | Poa annua L. | Poa annua | 0.05 | 1 | 0.200 | NA | NA | NA | NA | NA | NA | NA | NA | NA | NA | NA | NA | NA |
| 295 | Ranunculus repens L. | Ranunculus repens | 0.29 | 1 | 0.040 | 8.1 | NA | NA | 5.2 | NA | 0.1 | 0.51276 | 0.49581 | 1.51667 | 2.20989 | 0.98959 | 0.46997 | 0.3594 |
| 295 | Cerastium glomeratum Thuill. | Cerastium glomeratum | 0.02 | 1 | 0.102 | 6.8 | NA | 5.6 | 5.8 | NA | 0.1 | 0.52169 | 0.45097 | 0.93044 | 1.87015 | 0.67593 | 0.2975 | 0.27054 |
| 295 | Clematis vitalba L. | Clematis vitalba | 0.02 | 1 | 0.020 | 7.6 | NA | NA | 6.5 | 6.3 | 0.7 | 0.75548 | 0.75048 | 1.8409 | 2.08419 | 0.63746 | 0.23177 | 0.58814 |
| 296 | Alliaria petiolata (M.Bieb.) | Alliaria petiolata | 0.3 | 0.99 | 0.020 | NA | NA | NA | NA | NA | NA | NA | NA | NA | NA | NA | NA | NA |
| 296 | Veronica hederifolia L. | Veronica hederifolia agg. | 0.23 | 0.99 | 0.051 | 6.2 | 5.5 | NA | 6.9 | 8.5 | 0.1 | 0.60658 | 0.39025 | 0.44214 | 1.39281 | 0.24736 | 0.18526 | 0.18179 |
| 296 | Glechoma hederacea L. | Glechoma hederacea | 0.25 | 0.99 | 0.020 | 7.1 | 4.9 | NA | 6 | NA | 1 | 0.59517 | 0.59064 | 1.89772 | 2.12267 | 0.65091 | 0.41213 | 0.37185 |
| 296 | Cardamine hirsuta L. | Cardamine hirsuta | 0.07 | 0.99 | 0.060 | NA | NA | 7 | 7.1 | 7.1 | 0 | 0.67499 | 0.26365 | 0.14663 | 1.3586 | 0.0564 | 0.18297 | 0.11371 |
| 296 | Poa annua L. | Poa annua | 0.07 | 0.99 | 0.083 | 8.1 | NA | NA | 4.4 | 2.8 | 0.3 | 0.48868 | 0.46508 | 1.4426 | 2.04965 | 0.7278 | 0.33931 | 0.35628 |
| 296 | Betula pendula Roth | Betula pendula | 0.02 | 0.99 | 0.041 | 6.3 | 5 | 7.3 | 5.8 | 6.5 | 0.7 | 0.53692 | 0.48641 | 1.02734 | 1.80642 | 0.6097 | 0.33182 | 0.26598 |
| 296 | Veronica arvensis L. | Veronica arvensis | 0.01 | 0.99 | 0.145 | 7.2 | 5 | 5.1 | 6.1 | NA | 0.1 | 0.52316 | 0.5171 | 1.84298 | 2.10722 | 1.43318 | 0.13855 | 0.37874 |
| 296 | Cerastium glomeratum Thuill. | Cerastium glomeratum | 0.02 | 0.99 | 0.082 | 7.5 | NA | 4.2 | 6.8 | NA | 0 | 0.50996 | 0.50194 | 1.77695 | 2.09272 | 1.26941 | 0.17348 | 0.31239 |
| 296 | Polygonum aviculare L. | Polygonum aviculare agg. | 0.02 | 0.99 | 0.435 | 7.4 | 5.4 | 4.4 | 7.7 | 6.4 | 0.1 | 0.83003 | 0.82471 | 1.78482 | 1.97219 | 0.48138 | 0.13604 | 0.7566 |
| 297 | Alliaria petiolata (M.Bieb.) | Alliaria petiolata | 0.56 | 1 | 0.019 | NA | NA | NA | NA | NA | NA | NA | NA | NA | NA | NA | NA | NA |
| 297 | Veronica hederifolia L. | Veronica hederifolia agg. | 0.1 | 1 | 0.167 | 7.9 | 5.7 | 4.2 | 7 | 7.2 | 0.1 | 0.76998 | 0.7507 | 1.46055 | 1.90983 | 0.91556 | 0.1857 | 0.55131 |
| 297 | Glechoma hederacea L. | Glechoma hederacea | 0.1 | 1 | 0.022 | 8.1 | 6.2 | NA | 7.1 | 7.6 | 0.1 | 0.8111 | 0.79007 | 1.4109 | 1.89172 | 0.89447 | 0.17462 | 0.49967 |
| 297 | Clematis vitalba L. | Clematis vitalba | 0.02 | 1 | 0.050 | NA | NA | 6.4 | 6.8 | 6.8 | 0 | 0.66551 | 0.27982 | 0.17419 | 1.41859 | 0.08796 | 0.19647 | 0.14036 |
| 297 | Veronica arvensis L. | Veronica arvensis | 0.05 | 1 | 0.190 | NA | NA | NA | NA | NA | NA | NA | NA | NA | NA | NA | NA | NA |
| 297 | Poa annua L. | Poa annua | 0.05 | 1 | 0.050 | 6.7 | NA | NA | 6.5 | 7.7 | 0.1 | 0.70945 | 0.61373 | 0.80583 | 1.7717 | 0.43958 | 0.26577 | 0.51423 |
| 297 | NA | NA | 0.1 | 1 | 0.100 | 6.7 | NA | NA | 6.5 | 7.7 | 0.1 | 0.70945 | 0.61373 | 0.80583 | 1.7717 | 0.43958 | 0.26577 | 0.51423 |
| 297 | Galium aparine L. | Galium aparine aparine | 0.02 | 1 | 0.056 | NA | NA | 4.5 | 6.6 | 6.6 | 0 | 0.78059 | 0.60963 | 0.54418 | 1.63701 | 0.26733 | 0.16232 | 0.54354 |
| 298 | Alliaria petiolata (M.Bieb.) | Alliaria petiolata | 0.19 | 1 | 0.100 | 7.4 | 5.3 | NA | 6 | 7.7 | 0.8 | 0.65201 | 0.63791 | 1.54898 | 2.03286 | 0.77966 | 0.37419 | 0.45721 |
| 298 | Veronica hederifolia L. | Veronica hederifolia agg. | 0.05 | 1 | 0.100 | 7.2 | 5.5 | 4.5 | 6 | 7.6 | 0.3 | 0.72905 | 0.70188 | 1.35981 | 1.84158 | 0.57912 | 0.20081 | 0.60725 |
| 298 | Glechoma hederacea L. | Glechoma hederacea | 0.2 | 1 | 0.053 | 7 | 5.6 | 4.4 | 5.9 | 5.4 | 0.1 | 0.64871 | 0.63945 | 1.70759 | 2.05577 | 0.56937 | 0.21649 | 0.54803 |
| 298 | Poa annua L. | Poa annua | 0.3 | 1 | 0.050 | 6.3 | NA | 5.9 | 6.3 | 6.9 | 0.1 | 0.6037 | 0.3946 | 0.44368 | 1.50139 | 0.30164 | 0.21315 | 0.21494 |
| 298 | Geum urbanum L. | Geum urbanum | 0.1 | 1 | 0.050 | 6.7 | NA | NA | 6.5 | 7.7 | 0.1 | 0.70945 | 0.61373 | 0.80583 | 1.7717 | 0.43958 | 0.26577 | 0.51423 |
| 298 | Poa nemoralis L. | Poa nemoralis | 0.05 | 1 | 0.050 | 7.3 | 5.4 | 4.1 | 7.6 | 4.2 | 0.4 | 0.58728 | 0.56055 | 1.34601 | 1.94534 | 0.82357 | 0.23743 | 0.34603 |
| 298 | Acer campestre L. | Acer campestre | 0.1 | 1 | 0.050 | 7.4 | 5.3 | NA | 6 | 7.7 | 0.8 | 0.65201 | 0.63791 | 1.54898 | 2.03286 | 0.77966 | 0.37419 | 0.45721 |
| 298 | Trifolium repens L. | Trifolium repens | 0.01 | 1 | 0.063 | 6.7 | NA | NA | 6.5 | 7.7 | 0.1 | 0.70945 | 0.61373 | 0.80583 | 1.7717 | 0.43958 | 0.26577 | 0.51423 |
| 299 | Poa annua L. | Poa annua | 0.8 | 1 | 0.050 | 7.3 | 5.4 | 4.1 | 7.6 | 4.2 | 0.4 | 0.58728 | 0.56055 | 1.34601 | 1.94534 | 0.82357 | 0.23743 | 0.34603 |
| 299 | Ranunculus repens L. | Ranunculus repens | 0.05 | 1 | 0.250 | NA | NA | NA | NA | NA | NA | NA | NA | NA | NA | NA | NA | NA |
| 299 | Rumex obtusifolius L. | Rumex obtusifolius | 0.15 | 1 | 0.400 | NA | NA | NA | NA | NA | NA | NA | NA | NA | NA | NA | NA | NA |
| 300 | Alliaria petiolata (M.Bieb.) | Alliaria petiolata | 0.3 | 1 | 0.173 | 6.7 | NA | NA | 6.5 | 7.7 | 0.1 | 0.70945 | 0.61373 | 0.80583 | 1.7717 | 0.43958 | 0.26577 | 0.51423 |
| 300 | Glechoma hederacea L. | Glechoma hederacea | 0.4 | 1 | 0.100 | 8 | NA | NA | 6.1 | 6.3 | 0.1 | 0.56333 | 0.55394 | 1.72195 | 2.16326 | 1.07172 | 0.43309 | 0.38415 |

| | | | | | | | | | | | | | | | | | | |
|---|---|---|---|---|---|---|---|---|---|---|---|---|---|---|---|---|---|---|
| 300 | Poa annua L. | Poa annua | 0.3 | 1 | 0.290 | 6.3 | 5 | 7.3 | 5.8 | 6.5 | 0.7 | 0.53692 | 0.48641 | 1.02734 | 1.80642 | 0.6097 | 0.33182 | 0.26598 |
| 301 | Alliaria petiolata (M.Bieb.) | Alliaria petiolata | 0.1 | 1 | 0.020 | 7.2 | 5 | NA | 3.8 | 3.4 | 0.1 | 0.6826 | 0.18026 | 0.14111 | 1.34435 | 0.04142 | 0.20078 | 0.17144 |
| 301 | Glechoma hederacea L. | Glechoma hederacea | 0.4 | 1 | 0.050 | 7.4 | 5.3 | NA | 6 | 7.7 | 0.8 | 0.65201 | 0.63791 | 1.54898 | 2.03286 | 0.77966 | 0.37419 | 0.45721 |
| 301 | Medicago lupulina L. | Medicago lupulina | 0.1 | 1 | 0.050 | NA | NA | 5.4 | 6 | 4 | 0 | 0.68032 | 0.16727 | 0.07633 | 1.31114 | 0.02931 | 0.19319 | 0.11495 |
| 301 | Poa annua L. | Poa annua | 0.25 | 1 | 0.050 | 5.7 | NA | NA | 7.3 | 5.4 | 0 | 0.6918 | 0.13881 | 0.03984 | 1.31841 | 0.01173 | 0.19643 | 0.10363 |
| 301 | Ranunculus repens L. | Ranunculus repens | 0.05 | 1 | 0.040 | 6.9 | 5.9 | NA | 6.8 | 7 | 0 | 0.84157 | 0.83772 | 1.85474 | 2.04042 | 0.4856 | 0.14178 | 0.80173 |
| 301 | Acer campestre L. | Acer campestre | 0.05 | 1 | 0.051 | 7.1 | 4.9 | NA | 6 | NA | 1 | 0.59517 | 0.59064 | 1.89772 | 2.12267 | 0.65091 | 0.41213 | 0.37185 |
| 301 | Betula pendula Roth | Betula pendula | 0.05 | 1 | 0.160 | NA | NA | 4.5 | 6.6 | 6.6 | 0 | 0.78059 | 0.60963 | 0.54418 | 1.63701 | 0.26733 | 0.16232 | 0.54354 |
| 302 | Veronica arvensis L. | Veronica arvensis | 0.3 | 0.9 | 0.020 | 6.6 | 5.7 | NA | 6.3 | 7.6 | 0.1 | 0.67317 | 0.45023 | 0.43732 | 1.4267 | 0.22701 | 0.17304 | 0.27906 |
| 302 | Stellaria media (L.) Vill. | Stellaria media agg. | 0.3 | 0.9 | 0.050 | NA | 5.4 | 5.7 | 6.8 | 7 | 0 | 0.67742 | 0.24559 | 0.14546 | 1.36309 | 0.06547 | 0.19208 | 0.12644 |
| 302 | Acer campestre L. | Acer campestre | 0.2 | 0.9 | 0.020 | 7 | 5.6 | 4.4 | 5.9 | 5.4 | 0.1 | 0.64871 | 0.63945 | 1.70759 | 2.05577 | 0.56937 | 0.21649 | 0.54803 |
| 302 | Ranunculus ficaria L. | Ranunculus ficaria (Ficaria verna) | 0.1 | 0.9 | 0.100 | NA | NA | NA | NA | NA | NA | NA | NA | NA | NA | NA | NA | NA |
| 303 | Glechoma hederacea L. | Glechoma hederacea | 0.1 | 1 | 0.050 | 7.9 | 5.7 | 4.2 | 7 | 7.2 | 0.1 | 0.76998 | 0.7507 | 1.46055 | 1.90983 | 0.91556 | 0.1857 | 0.55131 |
| 303 | Trifolium repens L. | Trifolium repens | 0.05 | 1 | 0.200 | NA | NA | NA | 7 | NA | 0 | 0.66957 | 0.20824 | 0.12967 | 1.34554 | 0.07292 | 0.20428 | 0.13248 |
| 303 | Poa annua L. | Poa annua | 0.2 | 1 | 0.045 | 6.9 | 5.9 | NA | 6.8 | 7 | 0 | 0.84157 | 0.83772 | 1.85474 | 2.04042 | 0.4856 | 0.14178 | 0.80173 |
| 303 | Taraxacum officinale F.H.Wigg. | Taraxacum officinale agg. | 0.05 | 1 | 0.091 | 8.1 | NA | NA | 4.4 | 2.8 | 0.3 | 0.48868 | 0.46508 | 1.4426 | 2.04965 | 0.7278 | 0.33931 | 0.35628 |
| 303 | Geranium pusillum L. | Geranium pusillum | 0.2 | 1 | 0.300 | 7.4 | 5.3 | NA | 6 | 7.7 | 0.8 | 0.65201 | 0.63791 | 1.54898 | 2.03286 | 0.77966 | 0.37419 | 0.45721 |
| 303 | Veronica persica Poir. | Veronica persica (tournefortii) | 0.04 | 1 | 0.056 | NA | 5.4 | 5.7 | 6.8 | 7 | 0 | 0.67742 | 0.24559 | 0.14546 | 1.36309 | 0.06547 | 0.19208 | 0.12644 |
| 303 | Sonchus asper (L.) Hill | Sonchus asper | 0.08 | 1 | 0.100 | NA | 8.6 | 5.8 | 5 | NA | 0 | 0.68391 | 0.14543 | 0.0801 | 1.3317 | 0.00425 | 0.23391 | 0.10354 |
| 303 | Rubus pseudojaponicus Koidz. | Rubus caesius | 0.17 | 1 | 0.050 | 6.7 | NA | NA | 5.2 | NA | 0.4 | 0.59706 | 0.49429 | 0.69366 | 1.8006 | 0.482 | 0.19483 | 0.19138 |
| 303 | Veronica arvensis L. | Veronica arvensis | 0.1 | 1 | 0.050 | 6.7 | NA | NA | 5.2 | NA | 0.4 | 0.59706 | 0.49429 | 0.69366 | 1.8006 | 0.482 | 0.19483 | 0.19138 |
| 303 | Galium aparine L. | Galium aparine aparine | 0.01 | 1 | 0.118 | 8.4 | NA | 4 | 6.3 | NA | 0.1 | 0.76525 | 0.73026 | 1.22636 | 1.78874 | 0.7034 | 0.16135 | 0.51807 |
| 305 | Geranium pusillum L. | Geranium pusillum | 0.15 | 0.96 | 0.167 | 7.8 | NA | 4.6 | 6.2 | 5.8 | 0.1 | 0.71321 | 0.68276 | 1.29546 | 1.8275 | 0.62188 | 0.18605 | 0.48742 |
| 305 | Poa annua L. | Poa annua | 0.05 | 0.96 | 0.120 | 7.8 | NA | 4.6 | 6.2 | 5.8 | 0.1 | 0.71321 | 0.68276 | 1.29546 | 1.8275 | 0.62188 | 0.18605 | 0.48742 |
| 305 | Alliaria petiolata (M.Bieb.) | Alliaria petiolata | 0.1 | 0.96 | 0.059 | 7 | 5.6 | 4.4 | 5.9 | 5.4 | 0.1 | 0.64871 | 0.63945 | 1.70759 | 2.05577 | 0.56937 | 0.21649 | 0.54803 |
| 305 | Veronica hederifolia L. | Veronica hederifolia agg. | 0.08 | 0.96 | 0.179 | 8.4 | NA | 3 | 6 | NA | 0 | 0.46249 | 0.43915 | 1.39189 | 1.83687 | 0.38437 | 0.30571 | 0.38883 |
| 305 | Medicago lupulina L. | Medicago lupulina | 0.04 | 0.96 | 0.093 | 6.7 | NA | NA | 6.5 | 7.7 | 0.1 | 0.70945 | 0.61373 | 0.80583 | 1.7717 | 0.43958 | 0.26577 | 0.51423 |
| 305 | Acer campestre L. | Acer campestre | 0.1 | 0.96 | 0.083 | 7.7 | NA | NA | 5.6 | NA | 0.7 | 0.53223 | 0.52325 | 1.73719 | 2.14421 | 0.94278 | 0.39374 | 0.35054 |
| 305 | Clematis vitalba L. | Clematis vitalba | 0.05 | 0.96 | 0.051 | 7 | 5.6 | 4.4 | 5.9 | 5.4 | 0.1 | 0.64871 | 0.63945 | 1.70759 | 2.05577 | 0.56937 | 0.21649 | 0.54803 |
| 305 | Veronica arvensis L. | Veronica arvensis | 0.05 | 0.96 | 0.053 | NA | NA | NA | NA | NA | NA | NA | NA | NA | NA | NA | NA | NA |
| 305 | lene latifolia subsp. alba (Mill.) Greuter & Burd | Silene pratensis (alba) | 0.3 | 0.96 | 0.020 | 7.1 | 4.9 | NA | 6 | NA | 1 | 0.59517 | 0.59064 | 1.89772 | 2.12267 | 0.65091 | 0.41213 | 0.37185 |
| 305 | Urtica dioica L. | Urtica dioica | 0.02 | 0.96 | 0.220 | 7.4 | 5.3 | NA | 6 | 7.7 | 0.8 | 0.65201 | 0.63791 | 1.54898 | 2.03286 | 0.77966 | 0.37419 | 0.45721 |
| 305 | Galium aparine L. | Galium aparine aparine | 0.02 | 0.96 | 0.153 | 8.1 | NA | 3.9 | 6.9 | 6.4 | 0.4 | 0.80098 | 0.76713 | 1.29016 | 1.92909 | 0.75604 | 0.19458 | 0.52486 |
| 306 | Alliaria petiolata (M.Bieb.) | Alliaria petiolata | 1 | 1 | 0.128 | 8.1 | NA | 3.9 | 6.9 | 6.4 | 0.4 | 0.80098 | 0.76713 | 1.29016 | 1.92909 | 0.75604 | 0.19458 | 0.52486 |
| 307 | Glechoma hederacea L. | Glechoma hederacea | 0.15 | 0.98 | 0.080 | 7.1 | NA | NA | NA | 4.6 | 0 | 0.69216 | 0.66495 | 1.32487 | 1.98453 | 0.49303 | 0.16764 | 0.56484 |
| 307 | Viola odorata L. | Viola odorata | 0.3 | 0.98 | 0.050 | 8.1 | NA | 3.9 | 6.9 | 6.4 | 0.4 | 0.80098 | 0.76713 | 1.29016 | 1.92909 | 0.75604 | 0.19458 | 0.52486 |
| 307 | Poa annua L. | Poa annua | 0.2 | 0.98 | 0.210 | 8.1 | NA | 3.9 | 6.9 | 6.4 | 0.4 | 0.80098 | 0.76713 | 1.29016 | 1.92909 | 0.75604 | 0.19458 | 0.52486 |
| 307 | Veronica hederifolia L. | Veronica hederifolia agg. | 0.15 | 0.98 | 0.120 | 7.4 | 5.3 | NA | 6 | 7.7 | 0.8 | 0.65201 | 0.63791 | 1.54898 | 2.03286 | 0.77966 | 0.37419 | 0.45721 |
| 307 | Veronica arvensis L. | Veronica arvensis | 0.05 | 0.98 | 0.080 | 8.1 | NA | NA | 5.2 | NA | 0.1 | 0.51276 | 0.49581 | 1.51667 | 2.20989 | 0.98959 | 0.46997 | 0.3594 |
| 307 | Sagina procumbens L. | Sagina procumbens | 0.05 | 0.98 | 0.041 | 8.1 | NA | NA | 5.2 | NA | 0.1 | 0.51276 | 0.49581 | 1.51667 | 2.20989 | 0.98959 | 0.46997 | 0.3594 |
| 307 | Oxalis corniculata L. | Oxalis corniculata | 0.05 | 0.98 | 0.020 | 7 | 5.6 | 4.4 | 5.9 | 5.4 | 0.1 | 0.64871 | 0.63945 | 1.70759 | 2.05577 | 0.56937 | 0.21649 | 0.54803 |
| 307 | Clematis vitalba L. | Clematis vitalba | 0.03 | 0.98 | 0.040 | 7.3 | 6.1 | 3.8 | 6 | 5.1 | 0 | 0.62987 | 0.59281 | 1.27995 | 1.90491 | 0.6275 | 0.3036 | 0.42608 |
| 308 | Poa annua L. | Poa annua | 0.25 | 1 | 0.057 | NA | NA | NA | NA | NA | NA | NA | NA | NA | NA | NA | NA | NA |
| 308 | Veronica persica Poir. | Veronica persica (tournefortii) | 0.25 | 1 | 0.300 | 8 | NA | NA | 6.1 | 6.3 | 0.1 | 0.56333 | 0.55394 | 1.72195 | 2.16326 | 1.07172 | 0.43309 | 0.38415 |
| 308 | Arctium minus (Hill) Bernh. | Arctium minus | 0.38 | 1 | 0.056 | 7.4 | 5.8 | 4.2 | 6.7 | 5.3 | 0 | 0.62702 | 0.59749 | 1.35516 | 1.96644 | 0.57704 | 0.23154 | 0.49642 |
| 308 | Polygonum aviculare L. | Polygonum aviculare agg. | 0.1 | 1 | 0.167 | 6.8 | NA | 5.6 | 5.8 | NA | 0.1 | 0.52169 | 0.45097 | 0.93044 | 1.87015 | 0.67593 | 0.2975 | 0.27054 |
| 308 | Alliaria petiolata (M.Bieb.) | Alliaria petiolata | 0.02 | 1 | 0.150 | 6.8 | NA | 5.6 | 5.8 | NA | 0.1 | 0.52169 | 0.45097 | 0.93044 | 1.87015 | 0.67593 | 0.2975 | 0.27054 |
| 309 | Glechoma hederacea L. | Glechoma hederacea | 0.11 | 1 | 0.167 | 6.8 | NA | 5.6 | 5.8 | NA | 0.1 | 0.52169 | 0.45097 | 0.93044 | 1.87015 | 0.67593 | 0.2975 | 0.27054 |
| 309 | Viola odorata L. | Viola odorata | 0.17 | 1 | 0.188 | 7.7 | NA | 3.7 | 5.8 | 6.4 | 0 | 0.70633 | 0.61633 | 0.84477 | 1.62276 | 0.49804 | 0.1651 | 0.39904 |
| 309 | Poa annua L. | Poa annua | 0.17 | 1 | 0.033 | 6.6 | 5.7 | NA | 6.3 | 7.6 | 0.1 | 0.67317 | 0.45023 | 0.43732 | 1.4267 | 0.22701 | 0.17304 | 0.27906 |
| 309 | Acer campestre L. | Acer campestre | 0.17 | 1 | 0.050 | 8.1 | NA | 3.5 | 7.5 | 5 | 0.3 | 0.75184 | 0.72654 | 1.3537 | 1.85024 | 0.717 | 0.18024 | 0.48387 |
| 309 | Ranunculus repens L. | Ranunculus repens | 0.11 | 1 | 0.100 | 8.1 | NA | NA | 4.4 | 2.8 | 0.3 | 0.48868 | 0.46508 | 1.4426 | 2.04965 | 0.7278 | 0.33931 | 0.35628 |
| 309 | Veronica hederifolia L. | Veronica hederifolia agg. | 0.16 | 1 | 0.050 | 7.7 | NA | NA | 6.7 | 7.2 | 0.1 | 0.5681 | 0.51951 | 1.12401 | 1.73868 | 0.64408 | 0.31575 | 0.32626 |
| 309 | Clematis vitalba L. | Clematis vitalba | 0.02 | 1 | 0.150 | 8 | NA | NA | 6.1 | 6.3 | 0.1 | 0.56333 | 0.55394 | 1.72195 | 2.16326 | 1.07172 | 0.43309 | 0.38415 |
| 309 | Prunella vulgaris L. | Prunella vulgaris | 0.05 | 1 | 0.089 | 8.2 | NA | 3.5 | 5.7 | 3.2 | 0 | 0.46142 | 0.41736 | 1.14878 | 1.78546 | 0.38405 | 0.27764 | 0.246 |
| 309 | Galium aparine L. | Galium aparine aparine | 0.02 | 1 | 0.054 | 7.9 | NA | 4.1 | 5.8 | NA | 0.9 | 0.54139 | 0.51128 | 1.31363 | 1.93633 | 0.8112 | 0.25085 | 0.32953 |
| 309 | Alliaria petiolata (M.Bieb.) | Alliaria petiolata | 0.02 | 1 | 0.047 | 8.4 | NA | 3 | 6 | NA | 0 | 0.46249 | 0.43915 | 1.39189 | 1.83687 | 0.38437 | 0.30571 | 0.38883 |
| 310 | Alliaria petiolata (M.Bieb.) | Alliaria petiolata | 0.15 | 1 | 0.250 | 6.8 | NA | 5.6 | 5.8 | NA | 0.1 | 0.52169 | 0.45097 | 0.93044 | 1.87015 | 0.67593 | 0.2975 | 0.27054 |
| 310 | Glechoma hederacea L. | Glechoma hederacea | 0.3 | 1 | 0.023 | 7 | 5.6 | 4.4 | 5.9 | 5.4 | 0.1 | 0.64871 | 0.63945 | 1.70759 | 2.05577 | 0.56937 | 0.21649 | 0.54803 |
| 310 | Bellis perennis L. | Bellis perennis | 0.05 | 1 | 0.087 | 7.3 | 6.1 | 3.8 | 6 | 5.1 | 0 | 0.62987 | 0.59281 | 1.27995 | 1.90491 | 0.6275 | 0.3036 | 0.42608 |
| 310 | Geum urbanum L. | Geum urbanum | 0.3 | 1 | 0.039 | 7.2 | 6 | 4.1 | 6.8 | 4.9 | 0 | 0.72892 | 0.658 | 1.07912 | 1.91928 | 0.406 | 0.27624 | 0.47339 |
| 310 | Poa annua L. | Poa annua | 0.2 | 1 | 0.100 | NA | NA | NA | NA | NA | NA | NA | NA | NA | NA | NA | NA | NA |
| 311 | Glechoma hederacea L. | Glechoma hederacea | 0.1 | 1 | 0.029 | 7 | 5.6 | 4.4 | 5.9 | 5.4 | 0.1 | 0.64871 | 0.63945 | 1.70759 | 2.05577 | 0.56937 | 0.21649 | 0.54803 |
| 311 | Ranunculus repens L. | Ranunculus repens | 0.05 | 1 | 0.250 | NA | NA | NA | NA | NA | NA | NA | NA | NA | NA | NA | NA | NA |
| 311 | Veronica hederifolia L. | Veronica hederifolia agg. | 0.1 | 1 | 0.200 | NA | NA | NA | NA | NA | NA | NA | NA | NA | NA | NA | NA | NA |
| 311 | Urtica dioica L. | Urtica dioica | 0.05 | 1 | 0.510 | NA | NA | NA | NA | NA | NA | NA | NA | NA | NA | NA | NA | NA |
| 311 | Prunella vulgaris L. | Prunella vulgaris | 0.05 | 1 | 0.050 | 8.4 | NA | 3 | 6 | NA | 0 | 0.46249 | 0.43915 | 1.39189 | 1.83687 | 0.38437 | 0.30571 | 0.38883 |
| 311 | Galium aparine L. | Galium aparine aparine | 0.02 | 1 | 0.100 | 8 | 7.2 | 3.4 | 6.8 | 6.6 | 0 | 0.71596 | 0.67692 | 1.05842 | 1.65087 | 0.71218 | 0.18565 | 0.41952 |
| 311 | Geranium pusillum L. | Geranium pusillum | 0.05 | 1 | 0.158 | NA | NA | NA | NA | NA | NA | NA | NA | NA | NA | NA | NA | NA |
| 311 | Arctium minus (Hill) Bernh. | Arctium minus | 0.58 | 1 | 0.050 | 7.3 | NA | NA | 6.7 | 6.6 | 1 | 0.7393 | 0.71983 | 1.39902 | 1.79157 | 0.57605 | 0.19228 | 0.59493 |
| 312 | Glechoma hederacea L. | Glechoma hederacea | 0.21 | 1 | 0.370 | 7.4 | 5.3 | NA | 6 | 7.7 | 0.8 | 0.65201 | 0.63791 | 1.54898 | 2.03286 | 0.77966 | 0.37419 | 0.45721 |
| 312 | Veronica hederifolia L. | Veronica hederifolia agg. | 0.11 | 1 | 0.450 | NA | NA | NA | NA | NA | NA | NA | NA | NA | NA | NA | NA | NA |
| 312 | Urtica dioica L. | Urtica dioica | 0.05 | 1 | 0.350 | 7.4 | 5.3 | NA | 6 | 7.7 | 0.8 | 0.65201 | 0.63791 | 1.54898 | 2.03286 | 0.77966 | 0.37419 | 0.45721 |
| 312 | Ranunculus repens L. | Ranunculus repens | 0.31 | 1 | 0.150 | NA | NA | NA | NA | NA | NA | NA | NA | NA | NA | NA | NA | NA |
| 312 | Hedera helix L. | Hedera helix | 0.05 | 1 | 0.050 | 7.3 | 5.9 | 4.2 | 6 | 3.5 | 0 | 0.52833 | 0.51569 | 1.63653 | 2.07309 | 0.96127 | 0.27844 | 0.39266 |
| 312 | Geum urbanum L. | Geum urbanum | 0.05 | 1 | 0.270 | NA | NA | NA | NA | NA | NA | NA | NA | NA | NA | NA | NA | NA |
| 312 | Prunella vulgaris L. | Prunella vulgaris | 0.1 | 1 | 0.050 | 7.9 | 5.7 | 4.2 | 7 | 7.2 | 0.1 | 0.76998 | 0.7507 | 1.46055 | 1.90983 | 0.91556 | 0.1857 | 0.55131 |
| 312 | Veronica persica Poir. | Veronica persica (tournefortii) | 0.05 | 1 | 0.474 | 7.4 | 5.3 | NA | 6 | 7.7 | 0.8 | 0.65201 | 0.63791 | 1.54898 | 2.03286 | 0.77966 | 0.37419 | 0.45721 |
| 312 | Clematis vitalba L. | Clematis vitalba | 0.02 | 1 | 0.250 | NA | NA | NA | NA | NA | NA | NA | NA | NA | NA | NA | NA | NA |
| 312 | Ranunculus ficaria L. | Ranunculus ficaria (Ficaria verna) | 0.05 | 1 | 0.200 | NA | NA | NA | NA | NA | NA | NA | NA | NA | NA | NA | NA | NA |
| 313 | Rumex obtusifolius L. | Rumex obtusifolius | 0.62 | 1 | 0.077 | 6.7 | NA | NA | 6.5 | 7.7 | 0.1 | 0.70945 | 0.61373 | 0.80583 | 1.7717 | 0.43958 | 0.26577 | 0.51423 |
| 313 | Ranunculus repens L. | Ranunculus repens | 0.13 | 1 | 0.143 | 7.8 | NA | 4.6 | 6.2 | 5.8 | 0.1 | 0.71321 | 0.68276 | 1.29546 | 1.8275 | 0.62188 | 0.18605 | 0.48742 |
| 313 | Glechoma hederacea L. | Glechoma hederacea | 0.13 | 1 | 0.133 | 7.3 | 5.4 | 4.1 | 7.6 | 4.2 | 0.4 | 0.58728 | 0.56055 | 1.34601 | 1.94534 | 0.82357 | 0.23743 | 0.34603 |
| 313 | Ranunculus ficaria L. | Ranunculus ficaria (Ficaria verna) | 0.05 | 1 | 0.020 | 6.7 | NA | NA | 6.5 | 7.7 | 0.1 | 0.70945 | 0.61373 | 0.80583 | 1.7717 | 0.43958 | 0.26577 | 0.51423 |
| 313 | Taraxacum officinale F.H.Wigg. | Taraxacum officinale agg. | 0.05 | 1 | 0.350 | 7.1 | 5.3 | 4.5 | 5.9 | 4.8 | 0.3 | 0.52518 | 0.5113 | 1.61045 | 2.05509 | 0.96495 | 0.27897 | 0.32911 |
| 313 | Veronica arvensis L. | Veronica arvensis | 0.02 | 1 | 0.056 | NA | NA | NA | NA | NA | NA | NA | NA | NA | NA | NA | NA | NA |
| 314 | Poa annua L. | Poa annua | 0.28 | 0.98 | 0.110 | 7.4 | 5.3 | NA | 6 | 7.7 | 0.8 | 0.65201 | 0.63791 | 1.54898 | 2.03286 | 0.77966 | 0.37419 | 0.45721 |
| 314 | Cirsium vulgare (Savi) Ten. | Cirsium vulgare (lanceolatum) | 0.6 | 0.98 | 0.360 | 7.4 | 5.3 | NA | 6 | 7.7 | 0.8 | 0.65201 | 0.63791 | 1.54898 | 2.03286 | 0.77966 | 0.37419 | 0.45721 |
| 314 | Veronica arvensis L. | Veronica arvensis | 0.05 | 0.98 | 0.086 | NA | NA | NA | NA | NA | NA | NA | NA | NA | NA | NA | NA | NA |
| 314 | Acer campestre L. | Acer campestre | 0.05 | 0.98 | 0.126 | 7.4 | 5.3 | NA | 6 | 7.7 | 0.8 | 0.65201 | 0.63791 | 1.54898 | 2.03286 | 0.77966 | 0.37419 | 0.45721 |
| 315 | Veronica persica Poir. | Veronica persica (tournefortii) | 0.1 | 0.9 | 0.250 | 7.4 | 5.3 | NA | 6 | 7.7 | 0.8 | 0.65201 | 0.63791 | 1.54898 | 2.03286 | 0.77966 | 0.37419 | 0.45721 |
| 315 | Poa annua L. | Poa annua | 0.3 | 0.9 | 0.040 | 8.4 | NA | 3 | 6 | NA | 0 | 0.46249 | 0.43915 | 1.39189 | 1.83687 | 0.38437 | 0.30571 | 0.38883 |
| 315 | Veronica arvensis L. | Veronica arvensis | 0.1 | 0.9 | 0.040 | 8.4 | NA | 3 | 6 | NA | 0 | 0.46249 | 0.43915 | 1.39189 | 1.83687 | 0.38437 | 0.30571 | 0.38883 |
| 315 | Cardamine hirsuta L. | Cardamine hirsuta | 0.1 | 0.9 | 0.080 | 6.7 | NA | NA | 5.2 | NA | 0.4 | 0.59706 | 0.49429 | 0.69366 | 1.8006 | 0.482 | 0.19483 | 0.19138 |
| 315 | Prunella vulgaris L. | Prunella vulgaris | 0.3 | 0.9 | 0.050 | 7.7 | NA | NA | 5.6 | NA | 0.7 | 0.53223 | 0.52325 | 1.73719 | 2.14421 | 0.94278 | 0.39374 | 0.35054 |
| 316 | Poa annua L. | Poa annua | 0.3 | 1 | 0.077 | 6 | 5.8 | 5.3 | 6 | 6.3 | 0 | 0.7125 | 0.21953 | 0.14134 | 1.36241 | 0.05031 | 0.19015 | 0.17087 |
| 316 | Cardamine hirsuta L. | Cardamine hirsuta | 0.1 | 1 | 0.013 | 7 | 5.6 | 4.4 | 5.9 | 5.4 | 0.1 | 0.64871 | 0.63945 | 1.70759 | 2.05577 | 0.56937 | 0.21649 | 0.54803 |
| 316 | Veronica persica Poir. | Veronica persica (tournefortii) | 0.05 | 1 | 0.150 | 7.3 | 5.4 | 4.1 | 7.6 | 4.2 | 0.4 | 0.58728 | 0.56055 | 1.34601 | 1.94534 | 0.82357 | 0.23743 | 0.34603 |
| 316 | Sonchus oleraceus L. | Sonchus oleraceus | 0.3 | 1 | 0.300 | 7.4 | 5.3 | NA | 6 | 7.7 | 0.8 | 0.65201 | 0.63791 | 1.54898 | 2.03286 | 0.77966 | 0.37419 | 0.45721 |
| 316 | Acer campestre L. | Acer campestre | 0.1 | 1 | 0.020 | 6.8 | 6.2 | 4.4 | NA | NA | 0.3 | 0.7871 | 0.75023 | 1.37944 | 1.92919 | 0.40631 | 0.19607 | 0.63681 |
| 316 | Taraxacum officinale F.H.Wigg. | Taraxacum officinale agg. | 0.1 | 1 | 0.050 | 7.3 | NA | 3.9 | 6.1 | 3.9 | 0.1 | 0.55903 | 0.41538 | 0.65189 | 1.65181 | 0.43719 | 0.24279 | 0.21709 |
| 316 | Cynosurus cristatus L. | Cynosurus cristatus | 0.05 | 1 | 0.170 | 7.8 | 5.5 | 3.3 | 6.4 | 3.3 | 0.1 | 0.51939 | 0.38466 | 0.67536 | 1.6325 | 0.37226 | 0.27333 | 0.16989 |
| 317 | Glechoma hederacea L. | Glechoma hederacea | 0.2 | 1 | 0.300 | NA | NA | NA | NA | NA | NA | NA | NA | NA | NA | NA | NA | NA |
| 317 | Poa annua L. | Poa annua | 0.35 | 1 | 0.030 | 7 | 5.6 | 4.4 | 5.9 | 5.4 | 0.1 | 0.64871 | 0.63945 | 1.70759 | 2.05577 | 0.56937 | 0.21649 | 0.54803 |
| 317 | Ranunculus repens L. | Ranunculus repens | 0.3 | 1 | 0.100 | NA | NA | NA | NA | NA | NA | NA | NA | NA | NA | NA | NA | NA |
| 317 | Trifolium repens L. | Trifolium repens | 0.1 | 1 | 0.062 | 8.4 | NA | 3 | 6 | NA | 0 | 0.46249 | 0.43915 | 1.39189 | 1.83687 | 0.38437 | 0.30571 | 0.38883 |

| | | | | | | | | | | | | | | | | |
|---|---|---|---|---|---|---|---|---|---|---|---|---|---|---|---|---|
| 317 | Taraxacum officinale F.H.Wigg. | Taraxacum officinale agg. | 0.05 | 1 | 0.050 | 6.8 | 6.2 | 4.4 | NA | NA | 0.3 | 0.7871 | 0.75023 | 1.37944 | 1.92919 | 0.40631 | 0.19607 | 0.63681 |
| 318 | Glechoma hederacea L. | Glechoma hederacea | 0.2 | 0.95 | 0.100 | NA | NA | NA | NA | NA | NA | NA | NA | NA | NA | NA | NA | NA |
| 318 | Poa annua L. | Poa annua | 0.3 | 0.95 | 0.066 | 7.1 | 5.3 | 4.5 | 5.9 | 4.8 | 0.3 | 0.52518 | 0.5113 | 1.61045 | 2.05509 | 0.96495 | 0.27897 | 0.32911 |
| 318 | Ranunculus repens L. | Ranunculus repens | 0.25 | 0.95 | 0.108 | 7 | 5.6 | 4.4 | 5.9 | 5.4 | 0.1 | 0.64871 | 0.63945 | 1.70759 | 2.05577 | 0.56937 | 0.21649 | 0.54803 |
| 318 | Geum urbanum L. | Geum urbanum | 0.2 | 0.95 | 0.190 | 7.3 | NA | 3.9 | 6.1 | 3.9 | 0.1 | 0.55903 | 0.41538 | 0.65189 | 1.65181 | 0.43719 | 0.24279 | 0.21709 |
| 319 | Taraxacum officinale F.H.Wigg. | Taraxacum officinale agg. | 0.1 | 0.85 | 0.067 | 7.9 | NA | 4.1 | 5.8 | NA | 0.9 | 0.54139 | 0.51128 | 1.31363 | 1.93633 | 0.8112 | 0.25085 | 0.32953 |
| 319 | Cerastium semidecandrum L. | Cerastium semidecandrum | 0.4 | 0.85 | 0.101 | 7.6 | NA | NA | 5.8 | NA | 0.5 | 0.55139 | 0.54271 | 1.75426 | 2.13166 | 0.95232 | 0.27891 | 0.39876 |
| 319 | Bromus sterilis L. | Bromus sterilis | 0.2 | 0.85 | 0.300 | 7.8 | 5.5 | 3.3 | 6.4 | 3.3 | 0.1 | 0.51939 | 0.38466 | 0.67536 | 1.6325 | 0.37226 | 0.27333 | 0.16989 |
| 319 | Bromus hordeaceus L. | Bromus hordeaceus (mollis) | 0.15 | 0.85 | 0.540 | 7.4 | 5.3 | NA | 6 | 7.7 | 0.8 | 0.65201 | 0.63791 | 1.54898 | 2.03286 | 0.77966 | 0.37419 | 0.45721 |
| 320 | Taraxacum officinale F.H.Wigg. | Taraxacum officinale agg. | 0.2 | 1 | 0.230 | NA | NA | NA | NA | NA | NA | NA | NA | NA | NA | NA | NA | NA |
| 320 | Veronica arvensis L. | Veronica arvensis | 0.1 | 1 | 0.095 | 7.6 | NA | NA | 5.8 | NA | 0.5 | 0.55139 | 0.54271 | 1.75426 | 2.13166 | 0.95232 | 0.27891 | 0.39876 |
| 320 | Medicago lupulina L. | Medicago lupulina | 0.25 | 1 | 0.020 | 7 | 5.6 | 4.4 | 5.9 | 5.4 | 0.1 | 0.64871 | 0.63945 | 1.70759 | 2.05577 | 0.56937 | 0.21649 | 0.54803 |
| 320 | Cerastium semidecandrum L. | Cerastium semidecandrum | 0.2 | 1 | 0.050 | 8.4 | NA | 4 | 6.3 | NA | 0.1 | 0.76525 | 0.73026 | 1.22636 | 1.78874 | 0.7034 | 0.16135 | 0.51807 |
| 320 | Poa annua L. | Poa annua | 0.15 | 1 | 0.100 | 8.1 | NA | NA | 5.2 | NA | 0.1 | 0.51276 | 0.49581 | 1.51667 | 2.20989 | 0.98959 | 0.46997 | 0.3594 |
| 320 | Sisymbrium officinale (L.) Scop. | Sisymbrium officinale | 0.05 | 1 | 0.050 | 8.6 | NA | 2.7 | 4.4 | 1.6 | 0 | 0.53557 | 0.50485 | 1.29532 | 1.90211 | 0.72298 | 0.28495 | 0.2778 |
| 320 | Geranium molle L. | Geranium molle | 0.05 | 1 | 0.417 | 7.4 | 5.3 | NA | 6 | 7.7 | 0.8 | 0.65201 | 0.63791 | 1.54898 | 2.03286 | 0.77966 | 0.37419 | 0.45721 |
| 321 | Geranium molle L. | Geranium molle | 0.1 | 1 | 0.080 | NA | NA | NA | NA | NA | NA | NA | NA | NA | NA | NA | NA | NA |
| 321 | Veronica arvensis L. | Veronica arvensis | 0.1 | 1 | 0.020 | 8.6 | NA | 2.7 | 4.4 | 1.6 | 0 | 0.53557 | 0.50485 | 1.29532 | 1.90211 | 0.72298 | 0.28495 | 0.2778 |
| 321 | Sisymbrium officinale (L.) Scop. | Sisymbrium officinale | 0.05 | 1 | 0.053 | 7 | 5.6 | 4.4 | 5.9 | 5.4 | 0.1 | 0.64871 | 0.63945 | 1.70759 | 2.05577 | 0.56937 | 0.21649 | 0.54803 |
| 321 | Bromus sterilis L. | Bromus sterilis | 0.35 | 1 | 0.347 | 7.4 | 5.3 | NA | 6 | 7.7 | 0.8 | 0.65201 | 0.63791 | 1.54898 | 2.03286 | 0.77966 | 0.37419 | 0.45721 |
| 321 | Hordeum murinum L. | Hordeum murinum agg. | 0.4 | 1 | 0.306 | 7.4 | 5.3 | NA | 6 | 7.7 | 0.8 | 0.65201 | 0.63791 | 1.54898 | 2.03286 | 0.77966 | 0.37419 | 0.45721 |
| 322 | Cerastium semidecandrum L. | Cerastium semidecandrum | 0.2 | 1 | 0.110 | NA | NA | NA | NA | NA | NA | NA | NA | NA | NA | NA | NA | NA |
| 322 | Taraxacum officinale F.H.Wigg. | Taraxacum officinale agg. | 0.1 | 1 | 0.150 | NA | NA | NA | NA | NA | NA | NA | NA | NA | NA | NA | NA | NA |
| 322 | Sagina procumbens L. | Sagina procumbens | 0.1 | 1 | 0.270 | 7.4 | 5.3 | NA | 6 | 7.7 | 0.8 | 0.65201 | 0.63791 | 1.54898 | 2.03286 | 0.77966 | 0.37419 | 0.45721 |
| 322 | Poa annua L. | Poa annua | 0.5 | 1 | 0.150 | NA | NA | NA | NA | NA | NA | NA | NA | NA | NA | NA | NA | NA |
| 322 | Veronica arvensis L. | Veronica arvensis | 0.1 | 1 | 0.050 | 7 | 5.6 | 4.4 | 5.9 | 5.4 | 0.1 | 0.64871 | 0.63945 | 1.70759 | 2.05577 | 0.56937 | 0.21649 | 0.54803 |
| 323 | Trifolium repens L. | Trifolium repens | 0.05 | 1 | 0.170 | NA | NA | NA | NA | NA | NA | NA | NA | NA | NA | NA | NA | NA |
| 323 | Veronica arvensis L. | Veronica arvensis | 0.1 | 1 | 0.050 | 6.7 | 5.3 | 4.5 | 5.3 | 5.8 | 0 | 0.75464 | 0.73051 | 1.36974 | 1.95508 | 0.45974 | 0.15567 | 0.6659 |
| 323 | Achillea millefolium L. | Achillea millefolium millefolium | 0.05 | 1 | 0.245 | 7.4 | 5.3 | NA | 6 | 7.7 | 0.8 | 0.65201 | 0.63791 | 1.54898 | 2.03286 | 0.77966 | 0.37419 | 0.45721 |
| 323 | Poa annua L. | Poa annua | 0.4 | 1 | 0.050 | 7.3 | NA | NA | 5.8 | 5.9 | 0.5 | 0.63207 | 0.61738 | 1.52973 | 2.00883 | 0.70322 | 0.36696 | 0.39725 |
| 323 | Plantago lanceolata L. | Plantago lanceolata | 0.05 | 1 | 0.050 | 6.6 | 5.7 | NA | 6.3 | 7.6 | 0.1 | 0.67317 | 0.45023 | 0.43732 | 1.4267 | 0.22701 | 0.17304 | 0.27906 |
| 323 | Viola odorata L. | Viola odorata | 0.2 | 1 | 0.010 | 7.4 | 5.3 | NA | 6 | 7.7 | 0.8 | 0.65201 | 0.63791 | 1.54898 | 2.03286 | 0.77966 | 0.37419 | 0.45721 |
| 323 | Hypochaeris radicata L. | Hypochoeris radicata | 0.05 | 1 | 0.020 | 7.3 | NA | NA | 5.8 | 5.9 | 0.5 | 0.63207 | 0.61738 | 1.52973 | 2.00883 | 0.70322 | 0.36696 | 0.39725 |
| 323 | Potentilla reptans L. | Potentilla reptans | 0.05 | 1 | 0.060 | 7.3 | NA | 3.9 | 6.1 | 3.9 | 0.1 | 0.55903 | 0.41538 | 0.65189 | 1.65181 | 0.43719 | 0.24279 | 0.21709 |
| 323 | Medicago lupulina L. | Medicago lupulina | 0.05 | 1 | 0.083 | 7.1 | 5.3 | 4.5 | 5.9 | 4.8 | 0.3 | 0.52518 | 0.5113 | 1.61045 | 2.05509 | 0.96495 | 0.27897 | 0.32911 |
| 324 | Trifolium repens L. | Trifolium repens | 0.1 | 1.1 | 0.102 | 7.3 | 5.4 | 4.1 | 7.6 | 4.2 | 0.4 | 0.58728 | 0.56055 | 1.34601 | 1.94534 | 0.82357 | 0.23743 | 0.34603 |
| 324 | Bellis perennis L. | Bellis perennis | 0.1 | 1.1 | 0.059 | 8.1 | NA | 3.5 | 7.5 | 5 | 0.3 | 0.75184 | 0.72654 | 1.3537 | 1.85024 | 0.717 | 0.18024 | 0.48387 |
| 324 | Veronica arvensis L. | Veronica arvensis | 0.05 | 1.1 | 0.038 | 7.9 | NA | 4.1 | 5.8 | NA | 0.9 | 0.54139 | 0.51128 | 1.31363 | 1.93633 | 0.8112 | 0.25085 | 0.32953 |
| 324 | Hypochaeris radicata L. | Hypochoeris radicata | 0.1 | 1.1 | 0.056 | 7.7 | NA | NA | 6.4 | NA | 0 | 0.51083 | 0.48089 | 1.29849 | 1.93695 | 0.96262 | 0.19429 | 0.30238 |
| 324 | Achillea millefolium L. | Achillea millefolium millefolium | 0.05 | 1.1 | 0.022 | 7.9 | NA | 4.1 | 5.8 | NA | 0.9 | 0.54139 | 0.51128 | 1.31363 | 1.93633 | 0.8112 | 0.25085 | 0.32953 |
| 324 | Veronica persica Poir. | Veronica persica (tournefortii) | 0.05 | 1.1 | 0.250 | 8 | 9 | 4.8 | 7.3 | 8 | 3 | 0.69885 | 0.6622 | 1.07245 | 1.3127 | 0.43376 | 0.20235 | 0.40238 |
| 324 | Poa annua L. | Poa annua | 0.45 | 1.1 | 0.050 | 7.6 | NA | NA | 6.5 | 6.3 | 0.7 | 0.75548 | 0.75048 | 1.8409 | 2.08419 | 0.63746 | 0.23177 | 0.58814 |
| 324 | Trifolium dubium Sibth. | Trifolium dubium (minus) | 0.2 | 1.1 | 0.056 | 6.7 | NA | NA | 6.5 | 7.7 | 0.1 | 0.70945 | 0.61373 | 0.80583 | 1.7717 | 0.43958 | 0.26577 | 0.51423 |
| 325 | Bellis perennis L. | Bellis perennis | 0.15 | 1.1 | 0.050 | 7.1 | 7.8 | 4.1 | 6.2 | 6 | 0.1 | 0.76637 | 0.65623 | 0.78647 | 1.82229 | 0.56568 | 0.18972 | 0.47454 |
| 325 | Cerastium semidecandrum L. | Cerastium semidecandrum | 0.15 | 1.1 | 0.080 | 6.9 | 6.3 | 3.9 | 6.3 | 7.1 | 0.1 | 0.77916 | 0.67616 | 0.95204 | 1.78267 | 0.43197 | 0.25047 | 0.50688 |
| 325 | Medicago lupulina L. | Medicago lupulina | 0.1 | 1.1 | 0.021 | 7.3 | NA | NA | 5.8 | 5.9 | 0.5 | 0.63207 | 0.61738 | 1.52973 | 2.00883 | 0.70322 | 0.36696 | 0.39725 |
| 325 | Trifolium repens L. | Trifolium repens | 0.1 | 1.1 | 0.010 | 7 | 5.6 | 4.4 | 5.9 | 5.4 | 0.1 | 0.64871 | 0.63945 | 1.70759 | 2.05577 | 0.56937 | 0.21649 | 0.54803 |
| 325 | Achillea millefolium L. | Achillea millefolium millefolium | 0.05 | 1.1 | 0.010 | NA | NA | NA | NA | NA | NA | NA | NA | NA | NA | NA | NA | NA |
| 325 | Hypochaeris radicata L. | Hypochoeris radicata | 0.1 | 1.1 | 0.063 | 7.4 | 5.3 | NA | 6 | 7.7 | 0.8 | 0.65201 | 0.63791 | 1.54898 | 2.03286 | 0.77966 | 0.37419 | 0.45721 |
| 325 | Taraxacum officinale F.H.Wigg. | Taraxacum officinale agg. | 0.05 | 1.1 | 0.300 | NA | NA | NA | NA | NA | NA | NA | NA | NA | NA | NA | NA | NA |
| 325 | Poa annua L. | Poa annua | 0.25 | 1.1 | 0.250 | 8.1 | NA | 3.9 | 6.9 | 6.4 | 0.4 | 0.80098 | 0.76713 | 1.29016 | 1.92909 | 0.75604 | 0.19458 | 0.52486 |
| 325 | Stellaria media (L.) Vill. | Stellaria media agg. | 0.05 | 1.1 | 0.067 | 6.2 | 5.5 | NA | 6.9 | 8.5 | 0.1 | 0.60658 | 0.39025 | 0.44214 | 1.39281 | 0.24736 | 0.18526 | 0.18179 |
| 325 | Viola odorata L. | Viola odorata | 0.1 | 1.1 | 0.150 | NA | NA | NA | NA | NA | NA | NA | NA | NA | NA | NA | NA | NA |
| 326 | Bellis perennis L. | Bellis perennis | 0.05 | 1 | 0.020 | 7.3 | 5.8 | 4.8 | 5.3 | 5.6 | 0 | 0.67083 | 0.63295 | 1.24583 | 1.99939 | 0.56795 | 0.28417 | 0.48569 |
| 326 | Prunella vulgaris L. | Prunella vulgaris | 0.2 | 1 | 0.010 | 7 | 5.6 | 4.4 | 5.9 | 5.4 | 0.1 | 0.64871 | 0.63945 | 1.70759 | 2.05577 | 0.56937 | 0.21649 | 0.54803 |
| 326 | Medicago lupulina L. | Medicago lupulina | 0.1 | 1 | 0.100 | NA | NA | NA | NA | NA | NA | NA | NA | NA | NA | NA | NA | NA |
| 326 | Geranium molle L. | Geranium molle | 0.05 | 1 | 0.100 | 5.7 | NA | NA | 7.3 | 5.4 | 0 | 0.6918 | 0.13881 | 0.03984 | 1.31841 | 0.01173 | 0.19643 | 0.10363 |
| 326 | Cerastium semidecandrum L. | Cerastium semidecandrum | 0.05 | 1 | 0.050 | 7.2 | 5 | NA | 3.8 | 3.4 | 0.1 | 0.6826 | 0.18026 | 0.14111 | 1.34435 | 0.04142 | 0.20078 | 0.17144 |
| 326 | Poa annua L. | Poa annua | 0.3 | 1 | 0.080 | 7.3 | NA | NA | 6.7 | 6.6 | 1 | 0.7393 | 0.71983 | 1.39902 | 1.79157 | 0.57605 | 0.19228 | 0.59493 |
| 326 | Taraxacum officinale F.H.Wigg. | Taraxacum officinale agg. | 0.05 | 1 | 0.104 | 5.7 | NA | NA | 7.3 | 5.4 | 0 | 0.6918 | 0.13881 | 0.03984 | 1.31841 | 0.01173 | 0.19643 | 0.10363 |
| 326 | Viola odorata L. | Viola odorata | 0.2 | 1 | 0.051 | 7.1 | 7.8 | 4.1 | 6.2 | 6 | 0.1 | 0.76637 | 0.65623 | 0.78647 | 1.82229 | 0.56568 | 0.18972 | 0.47454 |
| 327 | Muscari armeniacum H.J.Veitch | NA | 0.65 | 1 | 0.020 | 6.9 | 6.5 | NA | 7.3 | 6.3 | 0 | 0.67274 | 0.19896 | 0.13039 | 1.32576 | 0.04466 | 0.20186 | 0.11616 |
| 327 | Medicago lupulina L. | Medicago lupulina | 0.1 | 1 | 0.050 | 7.4 | 5.9 | 3.8 | 6 | NA | 0 | 0.7907 | 0.77966 | 1.64404 | 2.03844 | 0.57393 | 0.17512 | 0.69169 |
| 327 | Poa annua L. | Poa annua | 0.2 | 1 | 0.100 | 6.8 | NA | 5.6 | 5.8 | NA | 0.1 | 0.52169 | 0.45097 | 0.93044 | 1.87015 | 0.67593 | 0.2975 | 0.27054 |
| 327 | Viola odorata L. | Viola odorata | 0.05 | 1 | 0.050 | 8 | 4.8 | NA | 5.3 | 4 | 0 | 0.50421 | 0.49749 | 1.83545 | 2.2368 | 1.2334 | 0.37538 | 0.34444 |
| 328 | Poa annua L. | Poa annua | 0.7 | 0.85 | 0.050 | 7.3 | 6.1 | 3.8 | 6 | 5.1 | 0 | 0.62987 | 0.59281 | 1.27995 | 1.90491 | 0.6275 | 0.3036 | 0.42608 |
| 328 | Cardamine hirsuta L. | Cardamine hirsuta | 0.05 | 0.85 | 0.050 | 8.1 | NA | NA | 4.4 | 2.8 | 0.3 | 0.48868 | 0.46508 | 1.4426 | 2.04965 | 0.7278 | 0.33931 | 0.35628 |
| 328 | Taraxacum officinale F.H.Wigg. | Taraxacum officinale agg. | 0.1 | 0.85 | 0.409 | 7.4 | 5.3 | NA | 6 | 7.7 | 0.8 | 0.65201 | 0.63791 | 1.54898 | 2.03286 | 0.77966 | 0.37419 | 0.45721 |
| 329 | Hedera helix L. | Hedera helix | 0.3 | 1 | 0.045 | NA | NA | NA | NA | NA | NA | NA | NA | NA | NA | NA | NA | NA |
| 329 | Veronica hederifolia L. | Veronica hederifolia agg. | 0.3 | 1 | 0.050 | NA | NA | NA | NA | NA | NA | NA | NA | NA | NA | NA | NA | NA |
| 329 | Poa annua L. | Poa annua | 0.4 | 1 | 0.111 | 6 | 5.8 | 5.3 | 6 | 6.3 | 0 | 0.7125 | 0.21953 | 0.14134 | 1.36241 | 0.05031 | 0.19015 | 0.17087 |
| 330 | Plantago major L. | Plantago major ssp. major | 0.2 | 1 | 0.050 | 7 | 5.6 | 4.4 | 5.9 | 5.4 | 0.1 | 0.64871 | 0.63945 | 1.70759 | 2.05577 | 0.56937 | 0.21649 | 0.54803 |
| 330 | Veronica arvensis L. | Veronica arvensis | 0.05 | 1 | 0.050 | 7.1 | 4.3 | NA | 5.1 | NA | 0.1 | 0.5318 | 0.50357 | 1.29074 | 2.04581 | 0.808 | 0.3645 | 0.36613 |
| 330 | Poa annua L. | Poa annua | 0.4 | 1 | 0.050 | 6.6 | 5.6 | NA | 5.6 | 4.1 | 0.1 | 0.50969 | 0.50576 | 1.97957 | 2.23767 | 1.1694 | 0.28385 | 0.36995 |
| 330 | Taraxacum officinale F.H.Wigg. | Taraxacum officinale agg. | 0.25 | 1 | 0.051 | 7.3 | 6.1 | 3.8 | 6 | 5.1 | 0 | 0.62987 | 0.59281 | 1.27995 | 1.90491 | 0.6275 | 0.3036 | 0.42608 |
| 330 | Stellaria media (L.) Vill. | Stellaria media agg. | 0.1 | 1 | 0.221 | 8.1 | NA | NA | 5.2 | NA | 0.1 | 0.51276 | 0.49581 | 1.51667 | 2.20989 | 0.98959 | 0.46997 | 0.3594 |
| 331 | Campanula persicifolia L. | Campanula persicifolia | 0.6 | 1 | 0.093 | 6.8 | NA | 5.6 | 5.8 | NA | 0.1 | 0.52169 | 0.45097 | 0.93044 | 1.87015 | 0.67593 | 0.2975 | 0.27054 |
| 331 | Taraxacum officinale F.H.Wigg. | Taraxacum officinale agg. | 0.2 | 1 | 0.067 | 7.1 | 4.3 | NA | 5.1 | NA | 0.1 | 0.5318 | 0.50357 | 1.29074 | 2.04581 | 0.808 | 0.3645 | 0.36613 |
| 331 | Poa annua L. | Poa annua | 0.1 | 1 | 0.102 | 7.3 | 6.1 | 3.8 | 6 | 5.1 | 0 | 0.62987 | 0.59281 | 1.27995 | 1.90491 | 0.6275 | 0.3036 | 0.42608 |
| 331 | Veronica arvensis L. | Veronica arvensis | 0.05 | 1 | 0.105 | 7.3 | 5.4 | 4.1 | 7.6 | 4.2 | 0.4 | 0.58728 | 0.56055 | 1.34601 | 1.94534 | 0.82357 | 0.23743 | 0.34603 |
| 331 | Hedera helix L. | Hedera helix | 0.05 | 1 | 0.050 | 7.8 | NA | 4.6 | 6.2 | 5.8 | 0.1 | 0.71321 | 0.68276 | 1.29546 | 1.8275 | 0.62188 | 0.18605 | 0.48742 |
| 332 | Trifolium repens L. | Trifolium repens | 0.25 | 0.9 | 0.050 | 7 | 5.6 | 4.4 | 5.9 | 5.4 | 0.1 | 0.64871 | 0.63945 | 1.70759 | 2.05577 | 0.56937 | 0.21649 | 0.54803 |
| 332 | Taraxacum officinale F.H.Wigg. | Taraxacum officinale agg. | 0.2 | 0.9 | 0.235 | 7.4 | 5.3 | NA | 6 | 7.7 | 0.8 | 0.65201 | 0.63791 | 1.54898 | 2.03286 | 0.77966 | 0.37419 | 0.45721 |
| 332 | Medicago lupulina L. | Medicago lupulina | 0.1 | 0.9 | 0.060 | NA | NA | NA | NA | NA | NA | NA | NA | NA | NA | NA | NA | NA |
| 332 | Bromus sterilis L. | Bromus sterilis | 0.15 | 0.9 | 0.057 | 5.7 | NA | NA | 6.4 | 7.4 | 0 | 0.68859 | 0.26809 | 0.17316 | 1.34857 | 0.09545 | 0.19552 | 0.14349 |
| 332 | Veronica arvensis L. | Veronica arvensis | 0.05 | 0.9 | 0.100 | 8 | NA | NA | 6.1 | 6.3 | 0.1 | 0.56333 | 0.55394 | 1.72195 | 2.16326 | 1.07172 | 0.43309 | 0.38415 |
| 332 | Geum urbanum L. | Geum urbanum | 0.05 | 0.9 | 0.110 | 7.4 | 5.3 | NA | 6 | 7.7 | 0.8 | 0.65201 | 0.63791 | 1.54898 | 2.03286 | 0.77966 | 0.37419 | 0.45721 |
| 332 | Symphoricarpos albus (L.) S.F.Blake | NA | 0.1 | 0.9 | 0.030 | 8 | NA | NA | 6.1 | 6.3 | 0.1 | 0.56333 | 0.55394 | 1.72195 | 2.16326 | 1.07172 | 0.43309 | 0.38415 |
| 333 | Taraxacum officinale F.H.Wigg. | Taraxacum officinale agg. | 0.45 | 1 | 0.050 | 7.9 | NA | 4.1 | 5.8 | NA | 0.9 | 0.54139 | 0.51128 | 1.31363 | 1.93633 | 0.8112 | 0.25085 | 0.32953 |
| 333 | Bromus sterilis L. | Bromus sterilis | 0.1 | 1 | 0.093 | 7.9 | NA | 4.1 | 5.8 | NA | 0.9 | 0.54139 | 0.51128 | 1.31363 | 1.93633 | 0.8112 | 0.25085 | 0.32953 |
| 333 | Sonchus oleraceus L. | Sonchus oleraceus | 0.1 | 1 | 0.020 | 6.7 | NA | NA | 6.5 | 7.7 | 0.1 | 0.70945 | 0.61373 | 0.80583 | 1.7717 | 0.43958 | 0.26577 | 0.51423 |
| 333 | Rosa fujisanensis Makino | NA | 0.05 | 1 | 0.040 | 7.3 | NA | NA | 5.8 | 5.9 | 0.5 | 0.63207 | 0.61738 | 1.52973 | 2.00883 | 0.70322 | 0.36696 | 0.39725 |
| 333 | Viola odorata L. | Viola odorata | 0.2 | 1 | 0.222 | 6.3 | 5 | 7.3 | 5.8 | 6.5 | 0.7 | 0.53692 | 0.48641 | 1.02734 | 1.80642 | 0.6097 | 0.33182 | 0.26598 |
| 333 | Laurus nobilis L. | NA | 0.1 | 1 | 0.023 | 8.3 | 5.2 | 3.5 | 6.7 | 4 | 0.1 | 0.59253 | 0.57367 | 1.48069 | 1.91606 | 0.44486 | 0.27236 | 0.42509 |
| 335 | Poa annua L. | Poa annua | 0.6 | 1 | 0.050 | 6.3 | 5 | 7.3 | 5.8 | 6.5 | 0.7 | 0.53692 | 0.48641 | 1.02734 | 1.80642 | 0.6097 | 0.33182 | 0.26598 |
| 335 | Veronica arvensis L. | Veronica arvensis | 0.05 | 1 | 0.020 | 7.4 | 5.9 | 3.8 | 6 | NA | 0 | 0.7907 | 0.77966 | 1.64404 | 2.03844 | 0.57393 | 0.17512 | 0.69169 |
| 335 | Bromus hordeaceus L. | Bromus hordeaceus (mollis) | 0.3 | 1 | 0.056 | 7.3 | NA | NA | 7.6 | 7.7 | 0.4 | 0.75479 | 0.72524 | 1.28692 | 1.77374 | 0.61324 | 0.18283 | 0.56773 |
| 335 | Taraxacum officinale F.H.Wigg. | Taraxacum officinale agg. | 0.05 | 1 | 0.063 | 7.3 | NA | NA | 7.6 | 7.7 | 0.4 | 0.75479 | 0.72524 | 1.28692 | 1.77374 | 0.61324 | 0.18283 | 0.56773 |
| 336 | Poa annua L. | Poa annua | 0.6 | 1 | 0.050 | 7.7 | NA | NA | 6.7 | 7.2 | 0.1 | 0.5681 | 0.51951 | 1.12401 | 1.73868 | 0.64408 | 0.31575 | 0.32626 |
| 336 | Veronica arvensis L. | Veronica arvensis | 0.05 | 1 | 0.083 | 7.3 | 6.1 | 3.8 | 6 | 5.1 | 0 | 0.62987 | 0.59281 | 1.27995 | 1.90491 | 0.6275 | 0.3036 | 0.42608 |
| 336 | Bromus hordeaceus L. | Bromus hordeaceus (mollis) | 0.1 | 1 | 0.022 | 7.4 | 5.9 | 3.8 | 6 | NA | 0 | 0.7907 | 0.77966 | 1.64404 | 2.03844 | 0.57393 | 0.17512 | 0.69169 |
| 336 | Trifolium repens L. | Trifolium repens | 0.25 | 1 | 0.059 | 7.3 | 6.1 | 3.8 | 6 | 5.1 | 0 | 0.62987 | 0.59281 | 1.27995 | 1.90491 | 0.6275 | 0.3036 | 0.42608 |
| 337 | Trifolium repens L. | Trifolium repens | 0.15 | 1 | 0.022 | 7 | 5.6 | 4.4 | 5.9 | 5.4 | 0.1 | 0.64871 | 0.63945 | 1.70759 | 2.05577 | 0.56937 | 0.21649 | 0.54803 |
| 337 | Poa annua L. | Poa annua | 0.45 | 1 | 0.046 | 6.7 | NA | 4.9 | 6.8 | 5 | 0 | 0.66057 | 0.18432 | 0.11528 | 1.34075 | 0.06671 | 0.20032 | 0.12881 |
| 337 | Glechoma hederacea L. | Glechoma hederacea | 0.1 | 1 | 0.054 | 7.3 | 5.4 | 4.1 | 7.6 | 4.2 | 0.4 | 0.58728 | 0.56055 | 1.34601 | 1.94534 | 0.82357 | 0.23743 | 0.34603 |
| 337 | Ranunculus repens L. | Ranunculus repens | 0.05 | 1 | 0.060 | 6.3 | 5 | 7.3 | 5.8 | 6.5 | 0.7 | 0.53692 | 0.48641 | 1.02734 | 1.80642 | 0.6097 | 0.33182 | 0.26598 |

| | | | | | | | | | | | | | | | | | | |
|---|---|---|---|---|---|---|---|---|---|---|---|---|---|---|---|---|---|---|
| 337 | Medicago lupulina L. | Medicago lupulina | 0.15 | 1 | 0.029 | 6.7 | 5.3 | 4.5 | 5.3 | 5.8 | 0 | 0.75464 | 0.73051 | 1.36974 | 1.95508 | 0.45974 | 0.15567 | 0.6659 |
| 337 | Cerastium fontanum Baumg. | Cerastium fontanum fontanum | 0.05 | 1 | 0.100 | 8.2 | NA | 2.1 | 6.6 | 1.7 | 0.4 | 0.4561 | 0.42853 | 1.29025 | 1.77233 | 0.32666 | 0.30641 | 0.32553 |
| 337 | Veronica arvensis L. | Veronica arvensis | 0.05 | 1 | 0.150 | 7.6 | NA | NA | 5.8 | NA | 0.5 | 0.55139 | 0.54271 | 1.75426 | 2.13166 | 0.95232 | 0.27891 | 0.39876 |
| 338 | Trifolium repens L. | Trifolium repens | 0.2 | 1 | 0.050 | 6.7 | 5.3 | 4.5 | 5.3 | 5.8 | 0 | 0.75464 | 0.73051 | 1.36974 | 1.95508 | 0.45974 | 0.15567 | 0.6659 |
| 338 | Poa annua L. | Poa annua | 0.5 | 1 | 0.200 | 7.4 | 5.3 | NA | 6 | 7.7 | 0.8 | 0.65201 | 0.63791 | 1.54898 | 2.03286 | 0.77966 | 0.37419 | 0.45721 |
| 338 | Glechoma hederacea L. | Glechoma hederacea | 0.05 | 1 | 0.105 | 8.4 | NA | 3 | 6 | NA | 0 | 0.46249 | 0.43915 | 1.39189 | 1.83687 | 0.38437 | 0.30571 | 0.38883 |
| 338 | Plantago lanceolata L. | Plantago lanceolata | 0.1 | 1 | 0.020 | 8.5 | NA | 3.5 | 6.3 | 4.1 | 0.2 | 0.57633 | 0.55499 | 1.44489 | 1.81436 | 0.579 | 0.25616 | 0.27657 |
| 338 | Taraxacum officinale F.H.Wigg. | Taraxacum officinale agg. | 0.05 | 1 | 0.210 | NA | NA | NA | NA | NA | NA | NA | NA | NA | NA | NA | NA | NA |
| 338 | Cerastium fontanum Baumg. | Cerastium fontanum fontanum | 0.05 | 1 | 0.190 | NA | NA | NA | NA | NA | NA | NA | NA | NA | NA | NA | NA | NA |
| 338 | Veronica serpyllifolia L. | Veronica serpyllifolia | 0.05 | 1 | 0.080 | NA | 5.4 | 5.7 | 6.8 | 7 | 0 | 0.67742 | 0.24559 | 0.14546 | 1.36309 | 0.06547 | 0.19208 | 0.12644 |
| 339 | Trifolium repens L. | Trifolium repens | 0.25 | 1 | 0.300 | 7.4 | 5.3 | NA | 6 | 7.7 | 0.8 | 0.65201 | 0.63791 | 1.54898 | 2.03286 | 0.77966 | 0.37419 | 0.45721 |
| 339 | Poa annua L. | Poa annua | 0.5 | 1 | 0.105 | NA | NA | NA | NA | NA | NA | NA | NA | NA | NA | NA | NA | NA |
| 339 | Plantago major L. | Plantago major ssp. major | 0.1 | 1 | 0.044 | 8.4 | NA | 3 | 6 | NA | 0 | 0.46249 | 0.43915 | 1.39189 | 1.83687 | 0.38437 | 0.30571 | 0.38883 |
| 339 | Capsella bursa-pastoris Medik. | Capsella bursa-pastoris | 0.05 | 1 | 0.154 | 7.6 | NA | NA | 5.8 | NA | 0.5 | 0.55139 | 0.54271 | 1.75426 | 2.13166 | 0.95232 | 0.27891 | 0.39876 |
| 339 | Medicago lupulina L. | Medicago lupulina | 0.1 | 1 | 0.050 | 7.1 | 5.3 | 4.5 | 5.9 | 4.8 | 0.3 | 0.52518 | 0.5113 | 1.61045 | 2.05509 | 0.96495 | 0.27897 | 0.32911 |
| 340 | Trifolium repens L. | Trifolium repens | 0.3 | 1 | 0.080 | 6.7 | NA | NA | 6.5 | 7.7 | 0.1 | 0.70945 | 0.61373 | 0.80583 | 1.7717 | 0.43958 | 0.26577 | 0.51423 |
| 340 | Poa annua L. | Poa annua | 0.7 | 1 | 0.172 | 7.4 | 5.3 | NA | 6 | 7.7 | 0.8 | 0.65201 | 0.63791 | 1.54898 | 2.03286 | 0.77966 | 0.37419 | 0.45721 |
| 341 | Poa annua L. | Poa annua | 0.1 | 1 | 0.084 | 6.3 | NA | 5.9 | 6.3 | 6.9 | 0.1 | 0.6037 | 0.3946 | 0.44368 | 1.50139 | 0.30164 | 0.21315 | 0.21494 |
| 341 | Polygonum aviculare L. | Polygonum aviculare agg. | 0.3 | 1 | 0.050 | NA | NA | NA | NA | NA | NA | NA | NA | NA | NA | NA | NA | NA |
| 341 | Hordeum murinum L. | Hordeum murinum agg. | 0.6 | 1 | 0.050 | 7.3 | NA | 3.9 | 6.1 | 3.9 | 0.1 | 0.55903 | 0.41538 | 0.65189 | 1.65181 | 0.43719 | 0.24279 | 0.21709 |
| 344 | Taraxacum officinale F.H.Wigg. | Taraxacum officinale agg. | 0.15 | 0.95 | 0.050 | 8 | NA | NA | 6.1 | 6.3 | 0.1 | 0.56333 | 0.55394 | 1.72195 | 2.16326 | 1.07172 | 0.43309 | 0.38415 |
| 344 | Bromus sterilis L. | Bromus sterilis | 0.4 | 0.95 | 0.120 | 8 | NA | NA | 6.1 | 6.3 | 0.1 | 0.56333 | 0.55394 | 1.72195 | 2.16326 | 1.07172 | 0.43309 | 0.38415 |
| 344 | Galium aparine L. | Galium aparine aparine | 0.1 | 0.95 | 0.056 | 7.3 | 5.4 | 4.1 | 7.6 | 4.2 | 0.4 | 0.58728 | 0.56055 | 1.34601 | 1.94534 | 0.82357 | 0.23743 | 0.34603 |
| 344 | Erigeron canadensis L. | Conyza canadensis | 0.05 | 0.95 | 0.154 | 6.8 | NA | 5.6 | 5.8 | NA | 0.1 | 0.52169 | 0.45097 | 0.93044 | 1.87015 | 0.67593 | 0.2975 | 0.27054 |
| 344 | Sonchus oleraceus L. | Sonchus oleraceus | 0.25 | 0.95 | 0.040 | NA | NA | NA | NA | NA | NA | NA | NA | NA | NA | NA | NA | NA |
| 345 | Taraxacum officinale F.H.Wigg. | Taraxacum officinale agg. | 0.4 | 0.9 | 0.130 | 7.1 | 4.9 | NA | 6 | NA | 1 | 0.59517 | 0.59064 | 1.89772 | 2.12267 | 0.65091 | 0.41213 | 0.37185 |
| 345 | Bromus sterilis L. | Bromus sterilis | 0.25 | 0.9 | 0.020 | 8.6 | NA | 2.7 | 4.4 | 1.6 | 0 | 0.53557 | 0.50485 | 1.29532 | 1.90211 | 0.72298 | 0.28495 | 0.2778 |
| 345 | Erigeron canadensis L. | Conyza canadensis | 0.2 | 0.9 | 0.050 | 7.3 | 5.4 | 4.1 | 7.6 | 4.2 | 0.4 | 0.58728 | 0.56055 | 1.34601 | 1.94534 | 0.82357 | 0.23743 | 0.34603 |
| 345 | Sagina procumbens L. | Sagina procumbens | 0.05 | 0.9 | 0.313 | 7.4 | 5.3 | NA | 6 | 7.7 | 0.8 | 0.65201 | 0.63791 | 1.54898 | 2.03286 | 0.77966 | 0.37419 | 0.45721 |
| 346 | Taraxacum officinale F.H.Wigg. | Taraxacum officinale agg. | 0.3 | 0.9 | 0.025 | 7 | 5.6 | 4.4 | 5.9 | 5.4 | 0.1 | 0.64871 | 0.63945 | 1.70759 | 2.05577 | 0.56937 | 0.21649 | 0.54803 |
| 346 | Bromus sterilis L. | Bromus sterilis | 0.25 | 0.9 | 0.197 | 7 | NA | NA | NA | NA | 0.3 | 0.56899 | 0.4288 | 0.64817 | 1.63031 | 0.49671 | 0.22558 | 0.25973 |
| 346 | Veronica arvensis L. | Veronica arvensis | 0.35 | 0.9 | 0.140 | 7.4 | 5.3 | NA | 6 | 7.7 | 0.8 | 0.65201 | 0.63791 | 1.54898 | 2.03286 | 0.77966 | 0.37419 | 0.45721 |
| 347 | Taraxacum officinale F.H.Wigg. | Taraxacum officinale agg. | 0.1 | 0.85 | 0.081 | 7 | NA | NA | NA | NA | 0.3 | 0.56899 | 0.4288 | 0.64817 | 1.63031 | 0.49671 | 0.22558 | 0.25973 |
| 347 | Medicago lupulina L. | Medicago lupulina | 0.2 | 0.85 | 0.290 | 7.4 | 5.3 | NA | 6 | 7.7 | 0.8 | 0.65201 | 0.63791 | 1.54898 | 2.03286 | 0.77966 | 0.37419 | 0.45721 |
| 347 | Veronica arvensis L. | Veronica arvensis | 0.25 | 0.85 | 0.316 | 7.4 | 5.3 | NA | 6 | 7.7 | 0.8 | 0.65201 | 0.63791 | 1.54898 | 2.03286 | 0.77966 | 0.37419 | 0.45721 |
| 347 | Sagina procumbens L. | Sagina procumbens | 0.05 | 0.85 | 0.092 | 8.1 | NA | NA | 4.4 | 2.8 | 0.3 | 0.48868 | 0.46508 | 1.4426 | 2.04965 | 0.7278 | 0.33931 | 0.35628 |
| 347 | Poa annua L. | Poa annua | 0.15 | 0.85 | 0.390 | 7.4 | 5.3 | NA | 6 | 7.7 | 0.8 | 0.65201 | 0.63791 | 1.54898 | 2.03286 | 0.77966 | 0.37419 | 0.45721 |
| 347 | Lactuca serriola L. | Lactuca serriola (scariola) | 0.1 | 0.85 | 0.020 | 8.6 | NA | 2.7 | 4.4 | 1.6 | 0 | 0.53557 | 0.50485 | 1.29532 | 1.90211 | 0.72298 | 0.28495 | 0.2778 |
| 348 | Taraxacum officinale F.H.Wigg. | Taraxacum officinale agg. | 0.05 | 0.9 | 0.300 | 7.4 | 5.3 | NA | 6 | 7.7 | 0.8 | 0.65201 | 0.63791 | 1.54898 | 2.03286 | 0.77966 | 0.37419 | 0.45721 |
| 348 | Medicago lupulina L. | Medicago lupulina | 0.2 | 0.9 | 0.083 | NA | NA | NA | NA | NA | NA | NA | NA | NA | NA | NA | NA | NA |
| 348 | Veronica arvensis L. | Veronica arvensis | 0.25 | 0.9 | 0.320 | 7.4 | 5.3 | NA | 6 | 7.7 | 0.8 | 0.65201 | 0.63791 | 1.54898 | 2.03286 | 0.77966 | 0.37419 | 0.45721 |
| 348 | Sagina procumbens L. | Sagina procumbens | 0.15 | 0.9 | 0.260 | 7.4 | 5.3 | NA | 6 | 7.7 | 0.8 | 0.65201 | 0.63791 | 1.54898 | 2.03286 | 0.77966 | 0.37419 | 0.45721 |
| 348 | Lactuca serriola L. | Lactuca serriola (scariola) | 0.1 | 0.9 | 0.211 | 7.4 | 5.3 | NA | 6 | 7.7 | 0.8 | 0.65201 | 0.63791 | 1.54898 | 2.03286 | 0.77966 | 0.37419 | 0.45721 |
| 348 | Erigeron canadensis L. | Conyza canadensis | 0.15 | 0.9 | 0.211 | NA | NA | NA | NA | NA | NA | NA | NA | NA | NA | NA | NA | NA |
| 350 | Poa annua L. | Poa annua | 0.6 | 1 | 0.357 | NA | NA | NA | NA | NA | NA | NA | NA | NA | NA | NA | NA | NA |
| 350 | Bellis perennis L. | Bellis perennis | 0.05 | 1 | 0.110 | 8.1 | NA | NA | 5.2 | NA | 0.1 | 0.51276 | 0.49581 | 1.51667 | 2.20989 | 0.98959 | 0.46997 | 0.3594 |
| 350 | Capsella bursa-pastoris Medik. | Capsella bursa-pastoris | 0.01 | 1 | 0.050 | 6.7 | 5.3 | 4.5 | 5.3 | 5.8 | 0 | 0.75464 | 0.73051 | 1.36974 | 1.95508 | 0.45974 | 0.15567 | 0.6659 |
| 350 | Lolium perenne L. | Lolium perenne | 0.3 | 1 | 0.100 | NA | NA | NA | NA | NA | NA | NA | NA | NA | NA | NA | NA | NA |
| 350 | Geranium pusillum L. | Geranium pusillum | 0.04 | 1 | 0.200 | 7.6 | NA | NA | 5.8 | NA | 0.5 | 0.55139 | 0.54271 | 1.75426 | 2.13166 | 0.95232 | 0.27891 | 0.39876 |
| 351 | Poa annua L. | Poa annua | 0.1 | 1 | 0.300 | 7.4 | 5.3 | NA | 6 | 7.7 | 0.8 | 0.65201 | 0.63791 | 1.54898 | 2.03286 | 0.77966 | 0.37419 | 0.45721 |
| 351 | Lolium perenne L. | Lolium perenne | 0.35 | 1 | 0.130 | 7.6 | NA | NA | 5.8 | NA | 0.5 | 0.55139 | 0.54271 | 1.75426 | 2.13166 | 0.95232 | 0.27891 | 0.39876 |
| 351 | Hordeum murinum L. | Hordeum murinum agg. | 0.55 | 1 | 0.250 | 7.6 | NA | NA | 5.8 | NA | 0.5 | 0.55139 | 0.54271 | 1.75426 | 2.13166 | 0.95232 | 0.27891 | 0.39876 |
| 352 | Poa annua L. | Poa annua | 0.3 | 1 | 0.010 | NA | NA | NA | NA | NA | NA | NA | NA | NA | NA | NA | NA | NA |
| 352 | Hordeum murinum L. | Hordeum murinum agg. | 0.7 | 1 | 0.010 | NA | NA | 4.5 | 6.6 | 6.6 | 0 | 0.78059 | 0.60963 | 0.54418 | 1.63701 | 0.26733 | 0.16232 | 0.54354 |
| 353 | Taraxacum officinale F.H.Wigg. | Taraxacum officinale agg. | 0.55 | 0.95 | 0.010 | 6.6 | 5.7 | NA | 6.3 | 7.6 | 0.1 | 0.67317 | 0.45023 | 0.43732 | 1.4267 | 0.22701 | 0.17304 | 0.27906 |
| 353 | Lamium album L. | Lamium album | 0.4 | 0.95 | 0.010 | 7.4 | 5.3 | NA | 6 | 7.7 | 0.8 | 0.65201 | 0.63791 | 1.54898 | 2.03286 | 0.77966 | 0.37419 | 0.45721 |
| 354 | Convolvulus arvensis L. | Convolvulus arvensis | 0.97 | 1 | 0.212 | 8.1 | 6.2 | NA | 7.1 | 7.6 | 0.1 | 0.8111 | 0.79007 | 1.4109 | 1.89172 | 0.89447 | 0.17462 | 0.49967 |
| 354 | Taraxacum officinale F.H.Wigg. | Taraxacum officinale agg. | 0.03 | 1 | 0.060 | 7.7 | NA | NA | 6.7 | 7.2 | 0.1 | 0.5681 | 0.51951 | 1.12401 | 1.73868 | 0.64408 | 0.31575 | 0.32626 |
| 355 | Taraxacum officinale F.H.Wigg. | Taraxacum officinale agg. | 0.98 | 0.98 | 0.167 | 8.5 | NA | 3.5 | 6.3 | 4.1 | 0.2 | 0.57633 | 0.55499 | 1.44489 | 1.81436 | 0.579 | 0.25616 | 0.27657 |
| 356 | Taraxacum officinale F.H.Wigg. | Taraxacum officinale agg. | 1 | 1 | 0.204 | 7.7 | NA | NA | 6.4 | NA | 0 | 0.51083 | 0.48089 | 1.29849 | 1.93695 | 0.96262 | 0.19429 | 0.30238 |
| 357 | Taraxacum officinale F.H.Wigg. | Taraxacum officinale agg. | 0.68 | 0.98 | 0.096 | 7.1 | NA | 4.4 | 7 | 5.9 | 0.1 | 0.82845 | 0.8207 | 1.75144 | 1.9998 | 0.52148 | 0.15007 | 0.73628 |
| 357 | Poa annua L. | Poa annua | 0.15 | 0.98 | 0.011 | NA | NA | NA | NA | NA | NA | NA | NA | NA | NA | NA | NA | NA |
| 357 | Sagina procumbens L. | Sagina procumbens | 0.15 | 0.98 | 0.108 | 7.2 | 5.5 | 4.5 | 6 | 7.6 | 0.3 | 0.72905 | 0.70188 | 1.35981 | 1.84158 | 0.57912 | 0.20081 | 0.60725 |
| 358 | Taraxacum officinale F.H.Wigg. | Taraxacum officinale agg. | 0.23 | 1 | 0.111 | NA | NA | 5.3 | 6.2 | NA | 0.1 | 0.68682 | 0.14816 | 0.05813 | 1.31254 | 0.01433 | 0.20126 | 0.10505 |
| 358 | Bellis perennis L. | Bellis perennis | 0.1 | 1 | 0.100 | 7.8 | NA | 4.6 | 6.2 | 5.8 | 0.1 | 0.71321 | 0.68276 | 1.29546 | 1.8275 | 0.62188 | 0.18605 | 0.48742 |
| 358 | Geum urbanum L. | Geum urbanum | 0.05 | 1 | 0.050 | 7.4 | 5.3 | NA | 6 | 7.7 | 0.8 | 0.65201 | 0.63791 | 1.54898 | 2.03286 | 0.77966 | 0.37419 | 0.45721 |
| 358 | Hypochaeris radicata L. | Hypochoeris radicata | 0.11 | 1 | 0.050 | 7.9 | 5.7 | 4.2 | 7 | 7.2 | 0.1 | 0.76998 | 0.7507 | 1.46055 | 1.90983 | 0.91556 | 0.1857 | 0.55131 |
| 358 | Veronica arvensis L. | Veronica arvensis | 0.08 | 1 | 0.100 | 7 | NA | NA | NA | NA | 0.3 | 0.56899 | 0.4288 | 0.64817 | 1.63031 | 0.49671 | 0.22558 | 0.25973 |
| 358 | Erigeron canadensis L. | Conyza canadensis | 0.12 | 1 | 0.250 | 8.1 | NA | 3.9 | 6.9 | 6.4 | 0.4 | 0.80098 | 0.76713 | 1.29016 | 1.92909 | 0.75604 | 0.19458 | 0.52486 |
| 358 | Trifolium dubium Sibth. | Trifolium dubium (minus) | 0.05 | 1 | 0.100 | 7.4 | 5.3 | NA | 6 | 7.7 | 0.8 | 0.65201 | 0.63791 | 1.54898 | 2.03286 | 0.77966 | 0.37419 | 0.45721 |
| 358 | Geranium molle L. | Geranium molle | 0.04 | 1 | 0.050 | 7.3 | 5.4 | 4.1 | 7.6 | 4.2 | 0.4 | 0.58728 | 0.56055 | 1.34601 | 1.94534 | 0.82357 | 0.23743 | 0.34603 |
| 358 | Cerastium semidecandrum L. | Cerastium semidecandrum | 0.03 | 1 | 0.027 | 7.9 | 5.7 | 4.2 | 7 | 7.2 | 0.1 | 0.76998 | 0.7507 | 1.46055 | 1.90983 | 0.91556 | 0.1857 | 0.55131 |
| 358 | Sagina procumbens L. | Sagina procumbens | 0.09 | 1 | 0.020 | 6.9 | 6.5 | NA | 7.3 | 6.3 | 0 | 0.67274 | 0.19896 | 0.13039 | 1.32576 | 0.04466 | 0.20186 | 0.11616 |
| 358 | Glechoma hederacea L. | Glechoma hederacea | 0.02 | 1 | 0.020 | 7.3 | 5.8 | 4.8 | 5.3 | 5.6 | 0 | 0.67083 | 0.63295 | 1.24583 | 1.99939 | 0.56795 | 0.28417 | 0.48569 |
| 358 | Stellaria media (L.) Vill. | Stellaria media agg. | 0.08 | 1 | 0.020 | 6.6 | 5.7 | NA | 6.3 | 7.6 | 0.1 | 0.67317 | 0.45023 | 0.43732 | 1.4267 | 0.22701 | 0.17304 | 0.27906 |
| 359 | Taraxacum officinale F.H.Wigg. | Taraxacum officinale agg. | 0.03 | 0.78 | 0.010 | 7.7 | NA | NA | 5.6 | NA | 0.7 | 0.53223 | 0.52325 | 1.73719 | 2.14421 | 0.94278 | 0.39374 | 0.35054 |
| 359 | Trifolium repens L. | Trifolium repens | 0.05 | 0.78 | 0.170 | 6.9 | 5.3 | 6.4 | 7.3 | 6.7 | 1 | 0.60499 | 0.3452 | 0.36505 | 1.40223 | 0.13192 | 0.19582 | 0.16678 |
| 359 | Ranunculus repens L. | Ranunculus repens | 0.18 | 0.78 | 0.052 | 6.9 | 6.5 | NA | 7.3 | 6.3 | 0 | 0.67274 | 0.19896 | 0.13039 | 1.32576 | 0.04466 | 0.20186 | 0.11616 |
| 359 | Hypochaeris radicata L. | Hypochoeris radicata | 0.16 | 0.78 | 0.031 | 6.9 | 6.5 | NA | 7.3 | 6.3 | 0 | 0.67274 | 0.19896 | 0.13039 | 1.32576 | 0.04466 | 0.20186 | 0.11616 |
| 359 | Cerastium semidecandrum L. | Cerastium semidecandrum | 0.01 | 0.78 | 0.050 | 6.8 | NA | 5.6 | 5.8 | NA | 0.1 | 0.52169 | 0.45097 | 0.93044 | 1.87015 | 0.67593 | 0.2975 | 0.27054 |
| 359 | Geranium molle L. | Geranium molle | 0.04 | 0.78 | 0.580 | 8.3 | 5 | NA | 7.2 | 7.8 | 0.1 | 0.67936 | 0.56623 | 0.73548 | 1.48508 | 0.50389 | 0.1867 | 0.27262 |
| 359 | Trifolium dubium Sibth. | Trifolium dubium (minus) | 0.09 | 0.78 | 0.050 | 6.9 | 5.9 | NA | 6.8 | 7 | 0 | 0.84157 | 0.83772 | 1.85474 | 2.04042 | 0.4856 | 0.14178 | 0.80173 |
| 359 | Prunella vulgaris L. | Prunella vulgaris | 0.18 | 0.78 | 0.050 | 6.6 | NA | NA | 6.9 | 5.2 | 0.4 | 0.54634 | 0.5169 | 1.30759 | 1.85566 | 0.79671 | 0.26499 | 0.28358 |
| 359 | Veronica arvensis L. | Veronica arvensis | 0.01 | 0.78 | 0.182 | 6.6 | 5.6 | NA | 5.6 | 4.1 | 0.1 | 0.50969 | 0.50576 | 1.97957 | 2.23767 | 1.1694 | 0.28385 | 0.36995 |
| 359 | Bellis perennis L. | Bellis perennis | 0.03 | 0.78 | 0.227 | 7.4 | 5.3 | NA | 6 | 7.7 | 0.8 | 0.65201 | 0.63791 | 1.54898 | 2.03286 | 0.77966 | 0.37419 | 0.45721 |
| 360 | Cerastium semidecandrum L. | Cerastium semidecandrum | 0.08 | 0.68 | 0.200 | NA | NA | NA | 7 | NA | 0 | 0.66957 | 0.20824 | 0.12967 | 1.34554 | 0.07292 | 0.20428 | 0.13248 |
| 360 | Hedera helix L. | Hedera helix | 0.05 | 0.68 | 0.040 | 7.3 | 6.1 | 3.8 | 6 | 5.1 | 0 | 0.62987 | 0.59281 | 1.27995 | 1.90491 | 0.6275 | 0.3036 | 0.42608 |
| 360 | Viola odorata L. | Viola odorata | 0.05 | 0.68 | 0.115 | 6.6 | 5.6 | NA | 5.6 | 4.1 | 0.1 | 0.50969 | 0.50576 | 1.97957 | 2.23767 | 1.1694 | 0.28385 | 0.36995 |
| 360 | Taraxacum officinale F.H.Wigg. | Taraxacum officinale agg. | 0.08 | 0.68 | 0.074 | 7.3 | 6.1 | 3.8 | 6 | 5.1 | 0 | 0.62987 | 0.59281 | 1.27995 | 1.90491 | 0.6275 | 0.3036 | 0.42608 |
| 360 | Hypochaeris radicata L. | Hypochoeris radicata | 0.1 | 0.68 | 0.071 | 7.1 | NA | NA | NA | 4.6 | 0 | 0.69216 | 0.66495 | 1.32487 | 1.98453 | 0.49303 | 0.16764 | 0.56484 |
| 360 | Veronica arvensis L. | Veronica arvensis | 0.04 | 0.68 | 0.047 | 7.3 | 6.1 | 3.8 | 6 | 5.1 | 0 | 0.62987 | 0.59281 | 1.27995 | 1.90491 | 0.6275 | 0.3036 | 0.42608 |
| 360 | Bellis perennis L. | Bellis perennis | 0.15 | 0.68 | 0.204 | 7.6 | NA | NA | 5.8 | NA | 0.5 | 0.55139 | 0.54271 | 1.75426 | 2.13166 | 0.95232 | 0.27891 | 0.39876 |
| 360 | Geranium molle L. | Geranium molle | 0.05 | 0.68 | 0.263 | 7.6 | NA | NA | 5.8 | NA | 0.5 | 0.55139 | 0.54271 | 1.75426 | 2.13166 | 0.95232 | 0.27891 | 0.39876 |
| 360 | Prunella vulgaris L. | Prunella vulgaris | 0.04 | 0.68 | 0.080 | 7.3 | NA | 3.9 | 6.1 | 3.9 | 0.1 | 0.55903 | 0.41538 | 0.65189 | 1.65181 | 0.43719 | 0.24279 | 0.21709 |
| 360 | Stellaria media (L.) Vill. | Stellaria media agg. | 0.02 | 0.68 | 0.150 | 8.5 | NA | 6.6 | 7.1 | 5.6 | 6.7 | 0.52046 | 0.51794 | 2.06848 | 2.16909 | 0.3404 | 0.42137 | 0.28121 |
| 360 | Erigeron canadensis L. | Conyza canadensis | 0.02 | 0.68 | 0.020 | 7.1 | 4.9 | NA | 6 | NA | 1 | 0.59517 | 0.59064 | 1.89772 | 2.12267 | 0.65091 | 0.41213 | 0.37185 |
| 361 | Bellis perennis L. | Bellis perennis | 0.15 | 0.56 | 0.034 | 7.7 | NA | NA | 5.6 | NA | 0.7 | 0.53223 | 0.52325 | 1.73719 | 2.14421 | 0.94278 | 0.39374 | 0.35054 |
| 361 | Geranium molle L. | Geranium molle | 0.08 | 0.56 | 0.250 | 7.4 | 5.3 | NA | 6 | 7.7 | 0.8 | 0.65201 | 0.63791 | 1.54898 | 2.03286 | 0.77966 | 0.37419 | 0.45721 |
| 361 | Taraxacum officinale F.H.Wigg. | Taraxacum officinale agg. | 0.04 | 0.56 | 0.220 | 7.3 | 6.1 | 3.8 | 6 | 5.1 | 0 | 0.62987 | 0.59281 | 1.27995 | 1.90491 | 0.6275 | 0.3036 | 0.42608 |
| 361 | Veronica arvensis L. | Veronica arvensis | 0.09 | 0.56 | 0.020 | 6.9 | 5.9 | NA | 6.8 | 7 | 0 | 0.84157 | 0.83772 | 1.85474 | 2.04042 | 0.4856 | 0.14178 | 0.80173 |
| 361 | Cerastium semidecandrum L. | Cerastium semidecandrum | 0.1 | 0.56 | 0.021 | 6.9 | 5.9 | NA | 6.8 | 7 | 0 | 0.84157 | 0.83772 | 1.85474 | 2.04042 | 0.4856 | 0.14178 | 0.80173 |
| 361 | Vicia hirsuta (L.) Gray | Vicia hirsuta | 0.04 | 0.56 | 0.020 | 7 | 5.6 | 4.4 | 5.9 | 5.4 | 0.1 | 0.64871 | 0.63945 | 1.70759 | 2.05577 | 0.56937 | 0.21649 | 0.54803 |
| 361 | Stellaria media (L.) Vill. | Stellaria media agg. | 0.04 | 0.56 | 0.030 | 6.3 | NA | 5.9 | 6.3 | 6.9 | 0.1 | 0.6037 | 0.3946 | 0.44368 | 1.50139 | 0.30164 | 0.21315 | 0.21494 |
| 361 | Draba verna L. | Erophila verna | 0.02 | 0.56 | 0.388 | 7.4 | 5.3 | NA | 6 | 7.7 | 0.8 | 0.65201 | 0.63791 | 1.54898 | 2.03286 | 0.77966 | 0.37419 | 0.45721 |
| 362 | Bellis perennis L. | Bellis perennis | 0.08 | 0.43 | 0.350 | 7.4 | 5.3 | NA | 6 | 7.7 | 0.8 | 0.65201 | 0.63791 | 1.54898 | 2.03286 | 0.77966 | 0.37419 | 0.45721 |
| 362 | Veronica arvensis L. | Veronica arvensis | 0.06 | 0.43 | 0.033 | 7.4 | 5.9 | 3.8 | 6 | NA | 0 | 0.7907 | 0.77966 | 1.64404 | 2.03844 | 0.57393 | 0.17512 | 0.69169 |

| | | | | | | | | | | | | | | | | | | |
|---|---|---|---|---|---|---|---|---|---|---|---|---|---|---|---|---|---|---|
| 362 | Cerastium semidecandrum L. | Cerastium semidecandrum | 0.15 | 0.43 | 0.022 | 7.4 | 5.2 | 4.7 | 6.1 | NA | 0 | 0.77406 | 0.76726 | 1.79113 | 2.07279 | 0.64778 | 0.2119 | 0.65925 |
| 362 | Viola odorata L. | Viola odorata | 0.04 | 0.43 | 0.024 | NA | NA | NA | 6.6 | NA | 0 | 0.65326 | 0.6245 | 1.32492 | 1.96985 | 0.60133 | 0.19748 | 0.49592 |
| 362 | Stellaria media (L.) Vill. | Stellaria media agg. | 0.04 | 0.43 | 0.046 | 6.7 | NA | NA | 5.2 | NA | 0.4 | 0.59706 | 0.49429 | 0.69366 | 1.8006 | 0.482 | 0.19483 | 0.19138 |
| 362 | Prunella vulgaris L. | Prunella vulgaris | 0.04 | 0.43 | 0.054 | 6.7 | NA | NA | 5.2 | NA | 0.4 | 0.59706 | 0.49429 | 0.69366 | 1.8006 | 0.482 | 0.19483 | 0.19138 |
| 362 | Geranium molle L. | Geranium molle | 0.02 | 0.43 | 0.030 | 8.4 | NA | 3 | 6 | NA | 0 | 0.46249 | 0.43915 | 1.39189 | 1.83687 | 0.38437 | 0.30571 | 0.38883 |
| 363 | Bellis perennis L. | Bellis perennis | 0.05 | 0.6 | 0.050 | 8.4 | NA | 3 | 6 | NA | 0 | 0.46249 | 0.43915 | 1.39189 | 1.83687 | 0.38437 | 0.30571 | 0.38883 |
| 363 | Taraxacum officinale F.H.Wigg. | Taraxacum officinale agg. | 0.08 | 0.6 | 0.100 | 8.1 | NA | NA | 5.2 | NA | 0.1 | 0.51276 | 0.49581 | 1.51667 | 2.20989 | 0.98959 | 0.46997 | 0.3594 |
| 363 | Geranium molle L. | Geranium molle | 0.04 | 0.6 | 0.200 | NA | NA | NA | NA | NA | NA | NA | NA | NA | NA | NA | NA | NA |
| 363 | Viola odorata L. | Viola odorata | 0.12 | 0.6 | 0.053 | 8 | NA | 3.7 | NA | 2 | 0 | 0.55827 | 0.46888 | 0.89651 | 1.79355 | 0.39043 | 0.26528 | 0.38307 |
| 363 | Prunella vulgaris L. | Prunella vulgaris | 0.08 | 0.6 | 0.050 | 8.1 | NA | NA | 6.3 | 3.9 | 3.1 | 0.54917 | 0.54189 | 1.63411 | 1.71985 | 0.26966 | 0.39041 | 0.32538 |
| 363 | Trifolium repens L. | Trifolium repens | 0.05 | 0.6 | 0.050 | 8.1 | NA | NA | 4.4 | 2.8 | 0.3 | 0.48868 | 0.46508 | 1.4426 | 2.04965 | 0.7278 | 0.33931 | 0.35628 |
| 363 | Veronica serpyllifolia L. | Veronica serpyllifolia | 0.04 | 0.6 | 0.050 | NA | NA | NA | NA | NA | NA | NA | NA | NA | NA | NA | NA | NA |
| 363 | Cerastium semidecandrum L. | Cerastium semidecandrum | 0.05 | 0.6 | 0.300 | 7.4 | 5.3 | NA | 6 | 7.7 | 0.8 | 0.65201 | 0.63791 | 1.54898 | 2.03286 | 0.77966 | 0.37419 | 0.45721 |
| 363 | Stellaria media (L.) Vill. | Stellaria media agg. | 0.04 | 0.6 | 0.220 | NA | NA | NA | NA | NA | NA | NA | NA | NA | NA | NA | NA | NA |
| 363 | Medicago lupulina L. | Medicago lupulina | 0.05 | 0.6 | 0.044 | 7 | 5.6 | 4.4 | 5.9 | 5.4 | 0.1 | 0.64871 | 0.63945 | 1.70759 | 2.05577 | 0.56937 | 0.21649 | 0.54803 |
| 364 | Myosotis ramosissima Rochel | Myosotis ramosissima (coll. Hispida) | 0.02 | 0.98 | 0.222 | 7.3 | 6.1 | 3.8 | 6 | 5.1 | 0 | 0.62987 | 0.59281 | 1.27995 | 1.90491 | 0.6275 | 0.3036 | 0.42608 |
| 364 | Cardamine hirsuta L. | Cardamine hirsuta | 0.02 | 0.98 | 0.220 | 8.4 | 7.2 | 2.6 | 6.8 | 3.6 | 0.2 | 0.58203 | 0.49726 | 0.89718 | 1.5951 | 0.3257 | 0.17939 | 0.36267 |
| 364 | Potentilla reptans L. | Potentilla reptans | 0.35 | 0.98 | 0.375 | 8.4 | 7.2 | 2.6 | 6.8 | 3.6 | 0.2 | 0.58203 | 0.49726 | 0.89718 | 1.5951 | 0.3257 | 0.17939 | 0.36267 |
| 364 | Taraxacum officinale F.H.Wigg. | Taraxacum officinale agg. | 0.02 | 0.98 | 0.190 | 7.4 | 5.3 | NA | 6 | 7.7 | 0.8 | 0.65201 | 0.63791 | 1.54898 | 2.03286 | 0.77966 | 0.37419 | 0.45721 |
| 364 | Medicago lupulina L. | Medicago lupulina | 0.02 | 0.98 | 0.040 | 6.7 | NA | NA | 5.2 | NA | 0.4 | 0.59706 | 0.49429 | 0.69366 | 1.8006 | 0.482 | 0.19483 | 0.19138 |
| 364 | Veronica arvensis L. | Veronica arvensis | 0.05 | 0.98 | 0.200 | 7.4 | 5.3 | NA | 6 | 7.7 | 0.8 | 0.65201 | 0.63791 | 1.54898 | 2.03286 | 0.77966 | 0.37419 | 0.45721 |
| 364 | Geranium molle L. | Geranium molle | 0.1 | 0.98 | 0.100 | 7.4 | 5.3 | NA | 6 | 7.7 | 0.8 | 0.65201 | 0.63791 | 1.54898 | 2.03286 | 0.77966 | 0.37419 | 0.45721 |
| 364 | Bromus hordeaceus L. | Bromus hordeaceus (mollis) | 0.2 | 0.98 | 0.097 | 6.8 | NA | 5.6 | 5.8 | NA | 0.1 | 0.52169 | 0.45097 | 0.93044 | 1.87015 | 0.67593 | 0.2975 | 0.27054 |
| 364 | Rosa canina | Rosa canina agg. | 0.2 | 0.98 | 0.062 | 6.7 | NA | NA | 5.2 | NA | 0.4 | 0.59706 | 0.49429 | 0.69366 | 1.8006 | 0.482 | 0.19483 | 0.19138 |
| 365 | Myosotis ramosissima Rochel | Myosotis ramosissima (coll. Hispida) | 0.04 | 0.95 | 0.115 | NA | NA | NA | NA | NA | NA | NA | NA | NA | NA | NA | NA | NA |
| 365 | NA | NA | 0.03 | 0.95 | 0.040 | 6.7 | NA | NA | 5.2 | NA | 0.4 | 0.59706 | 0.49429 | 0.69366 | 1.8006 | 0.482 | 0.19483 | 0.19138 |
| 365 | NA | NA | 0.04 | 0.95 | 0.040 | 8.5 | NA | 2.5 | NA | NA | 0 | 0.506 | 0.44835 | 1.02121 | 1.67002 | 0.34316 | 0.25824 | 0.34742 |
| 365 | Geranium molle L. | Geranium molle | 0.04 | 0.95 | 0.045 | 7.1 | 5.3 | 4.5 | 5.9 | 4.8 | 0.3 | 0.52518 | 0.5113 | 1.61045 | 2.05509 | 0.96495 | 0.27897 | 0.32911 |
| 365 | Rosa canina | Rosa canina agg. | 0.2 | 0.95 | 0.488 | 7.4 | 5.3 | NA | 6 | 7.7 | 0.8 | 0.65201 | 0.63791 | 1.54898 | 2.03286 | 0.77966 | 0.37419 | 0.45721 |
| 365 | Taraxacum officinale F.H.Wigg. | Taraxacum officinale agg. | 0.05 | 0.95 | 0.270 | 7.4 | 5.3 | NA | 6 | 7.7 | 0.8 | 0.65201 | 0.63791 | 1.54898 | 2.03286 | 0.77966 | 0.37419 | 0.45721 |
| 365 | Medicago lupulina L. | Medicago lupulina | 0.1 | 0.95 | 0.150 | 7.4 | 5.3 | NA | 6 | 7.7 | 0.8 | 0.65201 | 0.63791 | 1.54898 | 2.03286 | 0.77966 | 0.37419 | 0.45721 |
| 365 | Bromus hordeaceus L. | Bromus hordeaceus (mollis) | 0.25 | 0.95 | 0.040 | 8 | NA | NA | 6.1 | 6.3 | 0.1 | 0.56333 | 0.55394 | 1.72195 | 2.16326 | 1.07172 | 0.43309 | 0.38415 |
| 365 | Poa annua L. | Poa annua | 0.15 | 0.95 | 0.040 | NA | NA | NA | NA | NA | NA | NA | NA | NA | NA | NA | NA | NA |
| 365 | NA | NA | 0.05 | 0.95 | 0.263 | NA | NA | NA | NA | NA | NA | NA | NA | NA | NA | NA | NA | NA |
| 369 | Poa annua L. | Poa annua | 0.27 | 1 | 0.150 | NA | NA | NA | NA | NA | NA | NA | NA | NA | NA | NA | NA | NA |
| 369 | Cirsium vulgare (Savi) Ten. | Cirsium vulgare (lanceolatum) | 0.15 | 1 | 0.300 | 7.4 | 5.3 | NA | 6 | 7.7 | 0.8 | 0.65201 | 0.63791 | 1.54898 | 2.03286 | 0.77966 | 0.37419 | 0.45721 |
| 369 | Veronica serpyllifolia L. | Veronica serpyllifolia | 0.05 | 1 | 0.120 | NA | NA | NA | NA | NA | NA | NA | NA | NA | NA | NA | NA | NA |
| 369 | Taraxacum officinale F.H.Wigg. | Taraxacum officinale agg. | 0.2 | 1 | 0.200 | NA | NA | NA | NA | NA | NA | NA | NA | NA | NA | NA | NA | NA |
| 369 | Hypochaeris radicata L. | Hypochoeris radicata | 0.18 | 1 | 0.200 | NA | NA | NA | NA | NA | NA | NA | NA | NA | NA | NA | NA | NA |
| 369 | Sagina procumbens L. | Sagina procumbens | 0.02 | 1 | 0.242 | NA | NA | NA | NA | NA | NA | NA | NA | NA | NA | NA | NA | NA |
| 369 | Erigeron canadensis L. | Conyza canadensis | 0.05 | 1 | 0.050 | 8.1 | NA | NA | 5.2 | NA | 0.1 | 0.51276 | 0.49581 | 1.51667 | 2.20989 | 0.98959 | 0.46997 | 0.3594 |
| 369 | Hypericum perforatum L. | Hypericum perforatum | 0.08 | 1 | 0.150 | 7.6 | NA | NA | 5.8 | NA | 0.5 | 0.55139 | 0.54271 | 1.75426 | 2.13166 | 0.95232 | 0.27891 | 0.39876 |
| 370 | Cardamine hirsuta L. | Cardamine hirsuta | 0.05 | 1 | 0.210 | NA | NA | NA | NA | NA | NA | NA | NA | NA | NA | NA | NA | NA |
| 370 | Poa annua L. | Poa annua | 0.9 | 1 | 0.020 | 6.7 | 5.3 | 4.5 | 5.3 | 5.8 | 0 | 0.75464 | 0.73051 | 1.36974 | 1.95508 | 0.45974 | 0.15567 | 0.6659 |
| 370 | Parthenocissus quinquefolia Planch. | NA | 0.05 | 1 | 0.010 | NA | NA | 4.5 | 6.6 | 6.6 | 0 | 0.78059 | 0.60963 | 0.54418 | 1.63701 | 0.26733 | 0.16232 | 0.54354 |
| 371 | Cardamine hirsuta L. | Cardamine hirsuta | 0.05 | 1 | 0.010 | NA | NA | NA | NA | NA | NA | NA | NA | NA | NA | NA | NA | NA |
| 371 | Galium aparine L. | Galium aparine aparine | 0.2 | 1 | 0.051 | 6.3 | NA | 5.9 | 6.3 | 6.9 | 0.1 | 0.6037 | 0.3946 | 0.44368 | 1.50139 | 0.30164 | 0.21315 | 0.21494 |
| 371 | Sagina procumbens L. | Sagina procumbens | 0.08 | 1 | 0.061 | 7.8 | 6.8 | 4.6 | 6.2 | 6 | 0 | 0.79444 | 0.76111 | 1.13974 | 1.67291 | 0.96445 | 0.14 | 0.57297 |
| 371 | Erigeron canadensis L. | Conyza canadensis | 0.2 | 1 | 0.060 | 7.6 | 5.9 | NA | 6.4 | 7.3 | 0.1 | 0.70455 | 0.66978 | 1.21675 | 1.62781 | 0.62451 | 0.16011 | 0.39524 |
| 371 | Stellaria media (L.) Vill. | Stellaria media agg. | 0.05 | 1 | 0.083 | 6.7 | 5.3 | 4.5 | 5.3 | 5.8 | 0 | 0.75464 | 0.73051 | 1.36974 | 1.95508 | 0.45974 | 0.15567 | 0.6659 |
| 371 | Poa annua L. | Poa annua | 0.22 | 1 | 0.041 | NA | NA | NA | 6.8 | 7.5 | 0 | 0.62243 | 0.29646 | 0.26317 | 1.34606 | 0.11632 | 0.1748 | 0.13996 |
| 371 | Veronica arvensis L. | Veronica arvensis | 0.05 | 1 | 0.192 | NA | NA | NA | 6.8 | 7.5 | 0 | 0.62243 | 0.29646 | 0.26317 | 1.34606 | 0.11632 | 0.1748 | 0.13996 |
| 371 | Puccinellia distans (Jacq.) Parl. | Puccinellia distans (Atropis distans) | 0.15 | 1 | 0.056 | 7.4 | 5.9 | 3.8 | 6 | NA | 0 | 0.7907 | 0.77966 | 1.64404 | 2.03844 | 0.57393 | 0.17512 | 0.69169 |
| 372 | Cardamine hirsuta L. | Cardamine hirsuta | 0.05 | 0.95 | 0.011 | 7.4 | 5.4 | 4.4 | 7.7 | 6.4 | 0.1 | 0.83003 | 0.82471 | 1.78482 | 1.97219 | 0.48138 | 0.13604 | 0.7566 |
| 372 | Galium aparine L. | Galium aparine aparine | 0.4 | 0.95 | 0.200 | 8.1 | NA | NA | 4.4 | 2.8 | 0.3 | 0.48868 | 0.46508 | 1.4426 | 2.04965 | 0.7278 | 0.33931 | 0.35628 |
| 372 | Taraxacum officinale F.H.Wigg. | Taraxacum officinale agg. | 0.25 | 0.95 | 0.050 | 7.1 | 4.9 | NA | 6 | NA | 1 | 0.59517 | 0.59064 | 1.89772 | 2.12267 | 0.65091 | 0.41213 | 0.37185 |
| 372 | Geum urbanum L. | Geum urbanum | 0.25 | 0.95 | 0.070 | 7.3 | NA | NA | 5.8 | 5.9 | 0.5 | 0.63207 | 0.61738 | 1.52973 | 2.00883 | 0.70322 | 0.36696 | 0.39725 |
| 373 | Taraxacum officinale F.H.Wigg. | Taraxacum officinale agg. | 0.2 | 1 | 0.093 | 7.7 | NA | NA | 5.6 | NA | 0.7 | 0.53223 | 0.52325 | 1.73719 | 2.14421 | 0.94278 | 0.39374 | 0.35054 |
| 373 | Hordeum murinum L. | Hordeum murinum agg. | 0.8 | 1 | 0.020 | 7.6 | NA | NA | 6.5 | 6.3 | 0.7 | 0.75548 | 0.75048 | 1.8409 | 2.08419 | 0.63746 | 0.23177 | 0.58814 |
| 374 | Sagina procumbens L. | Sagina procumbens | 1 | 1 | 0.100 | 7 | 5.6 | 4.4 | 5.9 | 5.4 | 0.1 | 0.64871 | 0.63945 | 1.70759 | 2.05577 | 0.56937 | 0.21649 | 0.54803 |
| 375 | Taraxacum officinale F.H.Wigg. | Taraxacum officinale agg. | 0.5 | 0.9 | 0.052 | 7 | 5.6 | 4.4 | 5.9 | 5.4 | 0.1 | 0.64871 | 0.63945 | 1.70759 | 2.05577 | 0.56937 | 0.21649 | 0.54803 |
| 375 | Sagina procumbens L. | Sagina procumbens | 0.2 | 0.9 | 0.020 | 6.6 | 5.7 | NA | 6.3 | 7.6 | 0.1 | 0.67317 | 0.45023 | 0.43732 | 1.4267 | 0.22701 | 0.17304 | 0.27906 |
| 375 | Stellaria media (L.) Vill. | Stellaria media agg. | 0.15 | 0.9 | 0.020 | 6.9 | 6.5 | NA | 7.3 | 6.3 | 0 | 0.67274 | 0.19896 | 0.13039 | 1.32576 | 0.04466 | 0.20186 | 0.11616 |
| 375 | Euphorbia peplus L. | Euphorbia peplus | 0.05 | 0.9 | 0.050 | 7.3 | 5.4 | 4.1 | 7.6 | 4.2 | 0.4 | 0.58728 | 0.56055 | 1.34601 | 1.94534 | 0.82357 | 0.23743 | 0.34603 |
| 376 | Taraxacum officinale F.H.Wigg. | Taraxacum officinale agg. | 0.32 | 1 | 0.045 | 6.7 | NA | NA | 6.5 | 7.7 | 0.1 | 0.70945 | 0.61373 | 0.80583 | 1.7717 | 0.43958 | 0.26577 | 0.51423 |
| 376 | Hypochaeris radicata L. | Hypochoeris radicata | 0.33 | 1 | 0.030 | 8.4 | NA | 3 | 6 | NA | 0 | 0.46249 | 0.43915 | 1.39189 | 1.83687 | 0.38437 | 0.30571 | 0.38883 |
| 376 | Sagina procumbens L. | Sagina procumbens | 0.35 | 1 | 0.231 | 6.8 | NA | 5.6 | 5.8 | NA | 0.1 | 0.52169 | 0.45097 | 0.93044 | 1.87015 | 0.67593 | 0.2975 | 0.27054 |
| 377 | Taraxacum officinale F.H.Wigg. | Taraxacum officinale agg. | 0.55 | 0.9 | 0.059 | 6.8 | NA | 5.6 | 5.8 | NA | 0.1 | 0.52169 | 0.45097 | 0.93044 | 1.87015 | 0.67593 | 0.2975 | 0.27054 |
| 377 | Sagina procumbens L. | Sagina procumbens | 0.25 | 0.9 | 0.071 | 6.7 | NA | NA | 6.5 | 7.7 | 0.1 | 0.70945 | 0.61373 | 0.80583 | 1.7717 | 0.43958 | 0.26577 | 0.51423 |
| 377 | Lactuca serriola L. | Lactuca serriola (scariola) | 0.1 | 0.9 | 0.083 | 8.4 | NA | 3 | 6 | NA | 0 | 0.46249 | 0.43915 | 1.39189 | 1.83687 | 0.38437 | 0.30571 | 0.38883 |
| 378 | Taraxacum officinale F.H.Wigg. | Taraxacum officinale agg. | 0.6 | 1 | 0.204 | 7.5 | NA | NA | 7 | NA | 0 | 0.62073 | 0.22498 | 0.24103 | 1.34507 | 0.09374 | 0.21377 | 0.12505 |
| 378 | Stellaria media (L.) Vill. | Stellaria media agg. | 0.1 | 1 | 0.158 | 7.4 | 5.3 | NA | 6 | 7.7 | 0.8 | 0.65201 | 0.63791 | 1.54898 | 2.03286 | 0.77966 | 0.37419 | 0.45721 |
| 378 | Sagina procumbens L. | Sagina procumbens | 0.1 | 1 | 0.050 | NA | NA | NA | NA | NA | NA | NA | NA | NA | NA | NA | NA | NA |
| 378 | Lactuca serriola L. | Lactuca serriola (scariola) | 0.2 | 1 | 0.070 | 7.9 | NA | 4.1 | 5.8 | NA | 0.9 | 0.54139 | 0.51128 | 1.31363 | 1.93633 | 0.8112 | 0.25085 | 0.32953 |
| 379 | Taraxacum officinale F.H.Wigg. | Taraxacum officinale agg. | 0.5 | 1 | 0.250 | 7.4 | 5.3 | NA | 6 | 7.7 | 0.8 | 0.65201 | 0.63791 | 1.54898 | 2.03286 | 0.77966 | 0.37419 | 0.45721 |
| 379 | Sagina procumbens L. | Sagina procumbens | 0.05 | 1 | 0.155 | 7.4 | 5.3 | NA | 6 | 7.7 | 0.8 | 0.65201 | 0.63791 | 1.54898 | 2.03286 | 0.77966 | 0.37419 | 0.45721 |
| 379 | Poa compressa L. | Poa compressa | 0.45 | 1 | 0.057 | 8.1 | NA | 4.1 | 6.6 | NA | 0.7 | 0.57545 | 0.53032 | 1.14497 | 1.72102 | 0.70077 | 0.21269 | 0.32208 |
| 380 | Taraxacum officinale F.H.Wigg. | Taraxacum officinale agg. | 0.2 | 1 | 0.167 | 8.6 | 5.9 | 3.5 | NA | 2.5 | 0 | 0.5503 | 0.53805 | 1.6668 | 2.00922 | 0.50104 | 0.3154 | 0.3619 |
| 380 | Waldsteinia ternata Fritsch | NA | 0.74 | 1 | 0.150 | 7.4 | 5.3 | NA | 6 | 7.7 | 0.8 | 0.65201 | 0.63791 | 1.54898 | 2.03286 | 0.77966 | 0.37419 | 0.45721 |
| 380 | Medicago lupulina L. | Medicago lupulina | 0.02 | 1 | 0.050 | 6.7 | NA | 4.9 | 6.8 | 5 | 0 | 0.66057 | 0.18432 | 0.11528 | 1.34075 | 0.06671 | 0.20032 | 0.12881 |
| 380 | Vicia hirsuta (L.) Gray | Vicia hirsuta | 0.04 | 1 | 0.118 | 8.1 | NA | NA | 4.4 | 2.8 | 0.3 | 0.48868 | 0.46508 | 1.4426 | 2.04965 | 0.7278 | 0.33931 | 0.35628 |
| 381 | Taraxacum officinale F.H.Wigg. | Taraxacum officinale agg. | 0.2 | 0.98 | 0.043 | 6.3 | 5 | 7.3 | 5.8 | 6.5 | 0.7 | 0.53692 | 0.48641 | 1.02734 | 1.80642 | 0.6097 | 0.33182 | 0.26598 |
| 381 | Waldsteinia ternata Fritsch | NA | 0.15 | 0.98 | 0.011 | 7.9 | NA | 4.1 | 5.8 | NA | 0.9 | 0.54139 | 0.51128 | 1.31363 | 1.93633 | 0.8112 | 0.25085 | 0.32953 |
| 381 | Medicago lupulina L. | Medicago lupulina | 0.05 | 0.98 | 0.054 | 7.7 | NA | NA | 5.6 | NA | 0.7 | 0.53223 | 0.52325 | 1.73719 | 2.14421 | 0.94278 | 0.39374 | 0.35054 |
| 381 | Reynoutria japonica Houtt. | Reynoutria japonica (Polygon. cuspidat. | 0.08 | 0.98 | 0.030 | 7 | 5.6 | 4.4 | 5.9 | 5.4 | 0.1 | 0.64871 | 0.63945 | 1.70759 | 2.05577 | 0.56937 | 0.21649 | 0.54803 |
| 381 | Hypochaeris radicata L. | Hypochoeris radicata | 0.1 | 0.98 | 0.100 | 8.2 | NA | 2.1 | 6.6 | 1.7 | 0.4 | 0.4561 | 0.42853 | 1.29025 | 1.77233 | 0.32666 | 0.30641 | 0.32553 |
| 381 | Hordeum murinum L. | Hordeum murinum agg. | 0.15 | 0.98 | 0.240 | 7.4 | 5.3 | NA | 6 | 7.7 | 0.8 | 0.65201 | 0.63791 | 1.54898 | 2.03286 | 0.77966 | 0.37419 | 0.45721 |
| 381 | Poa annua L. | Poa annua | 0.23 | 0.98 | 0.020 | NA | NA | NA | 6.6 | 7 | 0 | 0.72781 | 0.47089 | 0.36365 | 1.47514 | 0.21383 | 0.17215 | 0.37071 |
| 381 | Sagina procumbens L. | Sagina procumbens | 0.02 | 0.98 | 0.021 | 6.7 | 5.3 | 4.5 | 5.3 | 5.8 | 0 | 0.75464 | 0.73051 | 1.36974 | 1.95508 | 0.45974 | 0.15567 | 0.6659 |
| 382 | Taraxacum officinale F.H.Wigg. | Taraxacum officinale agg. | 0.1 | 0.94 | 0.160 | NA | NA | NA | NA | NA | NA | NA | NA | NA | NA | NA | NA | NA |
| 382 | Waldsteinia ternata Fritsch | NA | 0.4 | 0.94 | 0.050 | 8 | NA | NA | 6.1 | 6.3 | 0.1 | 0.56333 | 0.55394 | 1.72195 | 2.16326 | 1.07172 | 0.43309 | 0.38415 |
| 382 | Lysimachia arvensis (L.) U.Manns & Anderb. | Anagallis arvensis | 0.2 | 0.94 | 0.130 | 6.8 | NA | 5.6 | 5.8 | NA | 0.1 | 0.52169 | 0.45097 | 0.93044 | 1.87015 | 0.67593 | 0.2975 | 0.27054 |
| 382 | Hypochaeris radicata L. | Hypochoeris radicata | 0.08 | 0.94 | 0.040 | 7.3 | NA | NA | 5.8 | 5.9 | 0.5 | 0.63207 | 0.61738 | 1.52973 | 2.00883 | 0.70322 | 0.36696 | 0.39725 |
| 382 | Epilobium hirsutum L. | Epilobium hirsutum | 0.04 | 0.94 | 0.181 | NA | NA | NA | NA | NA | NA | NA | NA | NA | NA | NA | NA | NA |
| 382 | Hordeum murinum L. | Hordeum murinum agg. | 0.12 | 0.94 | 0.062 | 6.3 | NA | 5.9 | 6.3 | 6.9 | 0.1 | 0.6037 | 0.3946 | 0.44368 | 1.50139 | 0.30164 | 0.21315 | 0.21494 |
| 384 | Poa annua L. | Poa annua | 0.3 | 1 | 0.321 | 7.4 | 5.3 | NA | 6 | 7.7 | 0.8 | 0.65201 | 0.63791 | 1.54898 | 2.03286 | 0.77966 | 0.37419 | 0.45721 |
| 384 | Taraxacum officinale F.H.Wigg. | Taraxacum officinale agg. | 0.3 | 1 | 0.050 | NA | NA | NA | NA | NA | NA | NA | NA | NA | NA | NA | NA | NA |
| 384 | Stellaria media (L.) Vill. | Stellaria media agg. | 0.3 | 1 | 0.020 | 7.3 | NA | 3.9 | 6.1 | 3.9 | 0.1 | 0.55903 | 0.41538 | 0.65189 | 1.65181 | 0.43719 | 0.24279 | 0.21709 |
| 384 | Viola odorata L. | Viola odorata | 0.1 | 1 | 0.227 | 7.4 | 5.3 | NA | 6 | 7.7 | 0.8 | 0.65201 | 0.63791 | 1.54898 | 2.03286 | 0.77966 | 0.37419 | 0.45721 |
| 385 | Dactylis glomerata L. | Dactylis glomerata | 0.22 | 1 | 0.123 | 7.3 | NA | 3.9 | 6.1 | 3.9 | 0.1 | 0.55903 | 0.41538 | 0.65189 | 1.65181 | 0.43719 | 0.24279 | 0.21709 |
| 385 | Bromus sterilis L. | Bromus sterilis | 0.22 | 1 | 0.053 | 6.8 | 6.2 | 4.4 | NA | NA | 0.3 | 0.7871 | 0.75023 | 1.37944 | 1.92919 | 0.40631 | 0.19607 | 0.63681 |
| 385 | Hordeum murinum L. | Hordeum murinum agg. | 0.22 | 1 | 0.053 | 8.5 | NA | 3.5 | 6.3 | 4.1 | 0.2 | 0.57633 | 0.55499 | 1.44489 | 1.81436 | 0.579 | 0.25616 | 0.27657 |
| 385 | Polygonum aviculare L. | Polygonum aviculare agg. | 0.09 | 1 | 0.306 | 7.4 | 5.3 | NA | 6 | 7.7 | 0.8 | 0.65201 | 0.63791 | 1.54898 | 2.03286 | 0.77966 | 0.37419 | 0.45721 |
| 385 | Geranium molle L. | Geranium molle | 0.11 | 1 | 0.100 | 8.5 | NA | 3.5 | 6.3 | 4.1 | 0.2 | 0.57633 | 0.55499 | 1.44489 | 1.81436 | 0.579 | 0.25616 | 0.27657 |
| 385 | Vicia hirsuta (L.) Gray | Vicia hirsuta | 0.08 | 1 | 0.120 | NA | NA | NA | NA | NA | NA | NA | NA | NA | NA | NA | NA | NA |
| 385 | Taraxacum officinale F.H.Wigg. | Taraxacum officinale agg. | 0.06 | 1 | 0.110 | 8.5 | NA | 3.5 | 6.3 | 4.1 | 0.2 | 0.57633 | 0.55499 | 1.44489 | 1.81436 | 0.579 | 0.25616 | 0.27657 |

| | | | | | | | | | | | | | | | | | | |
|---|---|---|---|---|---|---|---|---|---|---|---|---|---|---|---|---|---|---|
| 386 | Taraxacum officinale F.H.Wigg. | Taraxacum officinale agg. | 0.1 | 1 | 0.080 | 8.5 | NA | 3.5 | 6.3 | 4.1 | 0.2 | 0.57633 | 0.55499 | 1.44489 | 1.81436 | 0.579 | 0.25616 | 0.27657 |
| 386 | Trifolium repens L. | Trifolium repens | 0.3 | 1 | 0.020 | 8.6 | NA | 2.7 | 4.4 | 1.6 | 0 | 0.53557 | 0.50485 | 1.29532 | 1.90211 | 0.72298 | 0.28495 | 0.2778 |
| 386 | Polygonum aviculare L. | Polygonum aviculare agg. | 0.05 | 1 | 0.050 | 7.1 | NA | NA | NA | 4.6 | 0 | 0.69216 | 0.66495 | 1.32487 | 1.98453 | 0.49303 | 0.16764 | 0.56484 |
| 386 | Dactylis glomerata L. | Dactylis glomerata | 0.55 | 1 | 0.050 | NA | NA | 5.7 | 7 | 5.7 | 0 | 0.68851 | 0.1621 | 0.05682 | 1.30073 | 0.01259 | 0.1846 | 0.10299 |
| 387 | Dactylis glomerata L. | Dactylis glomerata | 0.17 | 0.88 | 0.024 | 7.7 | NA | NA | 6.4 | NA | 0 | 0.51083 | 0.48089 | 1.29849 | 1.93695 | 0.96262 | 0.19429 | 0.30238 |
| 387 | Lepidium draba L. | Cardaria draba | 0.17 | 0.88 | 0.029 | 7.5 | NA | 4.2 | 6.8 | NA | 0 | 0.50996 | 0.50194 | 1.77695 | 2.09272 | 1.26941 | 0.17348 | 0.31239 |
| 387 | Plantago lanceolata L. | Plantago lanceolata | 0.17 | 0.88 | 0.038 | NA | NA | NA | NA | NA | NA | NA | NA | NA | NA | NA | NA | NA |
| 387 | Geum urbanum L. | Geum urbanum | 0.13 | 0.88 | 0.056 | 8.5 | NA | 3.5 | 6.3 | 4.1 | 0.2 | 0.57633 | 0.55499 | 1.44489 | 1.81436 | 0.579 | 0.25616 | 0.27657 |
| 387 | Taraxacum officinale F.H.Wigg. | Taraxacum officinale agg. | 0.08 | 0.88 | 0.183 | 7.6 | NA | 5.3 | 5.9 | 3.3 | 0.7 | 0.51097 | 0.48199 | 1.33025 | 1.98017 | 1.04662 | 0.20459 | 0.27353 |
| 387 | Chaerophyllum temulum L. | Chaerophyllum temulum | 0.05 | 0.88 | 0.100 | 7.8 | NA | 4.6 | 6.2 | 5.8 | 0.1 | 0.71321 | 0.68276 | 1.29546 | 1.8275 | 0.62188 | 0.18605 | 0.48742 |
| 387 | Trifolium repens L. | Trifolium repens | 0.03 | 0.88 | 0.133 | NA | NA | NA | NA | NA | NA | NA | NA | NA | NA | NA | NA | NA |
| 387 | Bromus sterilis L. | Bromus sterilis | 0.08 | 0.88 | 0.010 | 6.6 | 5.7 | NA | 6.3 | 7.6 | 0.1 | 0.67317 | 0.45023 | 0.43732 | 1.4267 | 0.22701 | 0.17304 | 0.27906 |
| 388 | Bromus sterilis L. | Bromus sterilis | 0.7 | 1 | 0.313 | 7.7 | NA | 3.7 | 5.8 | 6.4 | 0 | 0.70633 | 0.61633 | 0.84477 | 1.62276 | 0.49804 | 0.1651 | 0.39904 |
| 388 | Poa annua L. | Poa annua | 0.2 | 1 | 0.020 | 5.2 | 5.9 | 5.4 | 7.3 | 8 | 0 | 0.67947 | 0.25214 | 0.1495 | 1.33661 | 0.07498 | 0.19025 | 0.12819 |
| 388 | Medicago lupulina L. | Medicago lupulina | 0.1 | 1 | 0.050 | NA | NA | 6.4 | 6.8 | 6.8 | 0 | 0.66551 | 0.27982 | 0.17419 | 1.41859 | 0.08796 | 0.19647 | 0.14036 |
| 389 | Bromus sterilis L. | Bromus sterilis | 0.65 | 0.95 | 0.091 | NA | NA | NA | 7 | NA | 0 | 0.66957 | 0.20824 | 0.12967 | 1.34554 | 0.07292 | 0.20428 | 0.13248 |
| 389 | Dactylis glomerata L. | Dactylis glomerata | 0.1 | 0.95 | 0.090 | 7.1 | 4.9 | NA | 6 | NA | 1 | 0.59517 | 0.59064 | 1.89772 | 2.12267 | 0.65091 | 0.41213 | 0.37185 |
| 389 | Achillea millefolium L. | Achillea millefolium millefolium | 0.2 | 0.95 | 0.013 | 7 | 5.6 | 4.4 | 5.9 | 5.4 | 0.1 | 0.64871 | 0.63945 | 1.70759 | 2.05577 | 0.56937 | 0.21649 | 0.54803 |
| 390 | Taraxacum officinale F.H.Wigg. | Taraxacum officinale agg. | 0.15 | 1 | 0.029 | 6.7 | NA | NA | 6.5 | 7.7 | 0.1 | 0.70945 | 0.61373 | 0.80583 | 1.7717 | 0.43958 | 0.26577 | 0.51423 |
| 390 | Poa annua L. | Poa annua | 0.7 | 1 | 0.036 | 8.1 | NA | 3.1 | 6.6 | NA | 0 | 0.56066 | 0.54465 | 1.58041 | 1.97627 | 0.33838 | 0.29416 | 0.45216 |
| 390 | Clematis vitalba L. | Clematis vitalba | 0.15 | 1 | 0.067 | 6.7 | NA | NA | 6.5 | 7.7 | 0.1 | 0.70945 | 0.61373 | 0.80583 | 1.7717 | 0.43958 | 0.26577 | 0.51423 |
| 398 | Taraxacum officinale F.H.Wigg. | Taraxacum officinale agg. | 0.1 | 1 | 0.020 | 7.3 | 5.8 | 4.8 | 5.3 | 5.6 | 0 | 0.67083 | 0.63295 | 1.24583 | 1.99939 | 0.56795 | 0.28417 | 0.48569 |
| 398 | Plantago lanceolata L. | Plantago lanceolata | 0.1 | 1 | 0.021 | 7.8 | NA | 4.6 | 6.2 | 5.8 | 0.1 | 0.71321 | 0.68276 | 1.29546 | 1.8275 | 0.62188 | 0.18605 | 0.48742 |
| 398 | Medicago lupulina L. | Medicago lupulina | 0.1 | 1 | 0.114 | 7.6 | NA | NA | 5.8 | NA | 0.5 | 0.55139 | 0.54271 | 1.75426 | 2.13166 | 0.95232 | 0.27891 | 0.39876 |
| 398 | Geranium molle L. | Geranium molle | 0.17 | 1 | 0.020 | 6.6 | NA | NA | 6.9 | 5.2 | 0.4 | 0.54634 | 0.5169 | 1.30759 | 1.85566 | 0.79671 | 0.26499 | 0.28358 |
| 398 | Dactylis glomerata L. | Dactylis glomerata | 0.08 | 1 | 0.030 | 6.6 | NA | NA | 6.9 | 5.2 | 0.4 | 0.54634 | 0.5169 | 1.30759 | 1.85566 | 0.79671 | 0.26499 | 0.28358 |
| 398 | Hordeum murinum L. | Hordeum murinum agg. | 0.05 | 1 | 0.054 | 6.7 | NA | 4.9 | 6.8 | 5 | 0 | 0.66057 | 0.18432 | 0.11528 | 1.34075 | 0.06671 | 0.20032 | 0.12881 |
| 398 | Lolium perenne L. | Lolium perenne | 0.1 | 1 | 0.046 | 8.4 | NA | 3 | 6 | NA | 0 | 0.46249 | 0.43915 | 1.39189 | 1.83687 | 0.38437 | 0.30571 | 0.38883 |
| 398 | Poa annua L. | Poa annua | 0.25 | 1 | 0.043 | 8.4 | NA | 3 | 6 | NA | 0 | 0.46249 | 0.43915 | 1.39189 | 1.83687 | 0.38437 | 0.30571 | 0.38883 |
| 398 | Bromus sterilis L. | Bromus sterilis | 0.05 | 1 | 0.060 | 7.3 | 6.1 | 3.8 | 6 | 5.1 | 0 | 0.62987 | 0.59281 | 1.27995 | 1.90491 | 0.6275 | 0.3036 | 0.42608 |
| 399 | Taraxacum officinale F.H.Wigg. | Taraxacum officinale agg. | 0.05 | 1 | 0.040 | 7 | 5.6 | 4.4 | 5.9 | 5.4 | 0.1 | 0.64871 | 0.63945 | 1.70759 | 2.05577 | 0.56937 | 0.21649 | 0.54803 |
| 399 | Veronica arvensis L. | Veronica arvensis | 0.04 | 1 | 0.130 | NA | NA | NA | NA | NA | NA | NA | NA | NA | NA | NA | NA | NA |
| 399 | Potentilla reptans L. | Potentilla reptans | 0.17 | 1 | 0.105 | 7.1 | 5.3 | 4.5 | 5.9 | 4.8 | 0.3 | 0.52518 | 0.5113 | 1.61045 | 2.05509 | 0.96495 | 0.27897 | 0.32911 |
| 399 | Plantago lanceolata L. | Plantago lanceolata | 0.05 | 1 | 0.100 | 6.7 | NA | NA | 6.5 | 7.7 | 0.1 | 0.70945 | 0.61373 | 0.80583 | 1.7717 | 0.43958 | 0.26577 | 0.51423 |
| 399 | Medicago lupulina L. | Medicago lupulina | 0.04 | 1 | 0.050 | 7.3 | NA | NA | 5.8 | 5.9 | 0.5 | 0.63207 | 0.61738 | 1.52973 | 2.00883 | 0.70322 | 0.36696 | 0.39725 |
| 399 | Hordeum murinum L. | Hordeum murinum agg. | 0.21 | 1 | 0.300 | 7.4 | 5.3 | NA | 6 | 7.7 | 0.8 | 0.65201 | 0.63791 | 1.54898 | 2.03286 | 0.77966 | 0.37419 | 0.45721 |
| 399 | Poa annua L. | Poa annua | 0.11 | 1 | 0.261 | 7.7 | NA | 3.7 | 5.8 | 6.4 | 0 | 0.70633 | 0.61633 | 0.84477 | 1.62276 | 0.49804 | 0.1651 | 0.39904 |
| 399 | Geranium molle L. | Geranium molle | 0.22 | 1 | 0.210 | 7.4 | 5.3 | NA | 6 | 7.7 | 0.8 | 0.65201 | 0.63791 | 1.54898 | 2.03286 | 0.77966 | 0.37419 | 0.45721 |
| 399 | Achillea millefolium L. | Achillea millefolium millefolium | 0.07 | 1 | 0.263 | 8.2 | NA | 2.1 | 6.6 | 1.7 | 0.4 | 0.4561 | 0.42853 | 1.29025 | 1.77233 | 0.32666 | 0.30641 | 0.32553 |
| 399 | Cerastium glomeratum Thuill. | Cerastium glomeratum | 0.02 | 1 | 0.133 | NA | NA | NA | NA | NA | NA | NA | NA | NA | NA | NA | NA | NA |
| 399 | Stellaria media (L.) Vill. | Stellaria media agg. | 0.02 | 1 | 0.040 | 8.5 | NA | 3.5 | 6.3 | 4.1 | 0.2 | 0.57633 | 0.55499 | 1.44489 | 1.81436 | 0.579 | 0.25616 | 0.27657 |
| 400 | Taraxacum officinale F.H.Wigg. | Taraxacum officinale agg. | 0.05 | 1 | 0.050 | 8.5 | NA | 3.5 | 6.3 | 4.1 | 0.2 | 0.57633 | 0.55499 | 1.44489 | 1.81436 | 0.579 | 0.25616 | 0.27657 |
| 400 | Plantago lanceolata L. | Plantago lanceolata | 0.3 | 1 | 0.010 | 6.6 | NA | 4.6 | 6.5 | 4.9 | 0 | 0.56883 | 0.37493 | 0.49917 | 1.61084 | 0.41638 | 0.18856 | 0.24929 |
| 400 | Geranium molle L. | Geranium molle | 0.25 | 1 | 0.024 | 8 | 4.8 | NA | 5.3 | 4 | 0 | 0.50421 | 0.49749 | 1.83545 | 2.2368 | 1.2334 | 0.37538 | 0.34444 |
| 400 | Medicago lupulina L. | Medicago lupulina | 0.05 | 1 | 0.033 | 7.4 | 5.3 | NA | 6 | 7.7 | 0.8 | 0.65201 | 0.63791 | 1.54898 | 2.03286 | 0.77966 | 0.37419 | 0.45721 |
| 400 | Hordeum murinum L. | Hordeum murinum agg. | 0.2 | 1 | 0.056 | 7.5 | NA | NA | 7.1 | NA | 0.1 | 0.81782 | 0.78043 | 1.27269 | 1.85713 | 0.65149 | 0.18293 | 0.6358 |
| 400 | Poa annua L. | Poa annua | 0.12 | 1 | 0.011 | 8.4 | NA | 3 | 7.7 | 3.3 | 0 | 0.62073 | 0.55773 | 1.05738 | 1.76031 | 0.36863 | 0.23711 | 0.33892 |
| 400 | Lolium perenne L. | Lolium perenne | 0.03 | 1 | 0.093 | 7.3 | NA | NA | 5.8 | 5.9 | 0.5 | 0.63207 | 0.61738 | 1.52973 | 2.00883 | 0.70322 | 0.36696 | 0.39725 |
| 401 | Taraxacum officinale F.H.Wigg. | Taraxacum officinale agg. | 0.06 | 1 | 0.021 | 6.2 | 5.5 | NA | 6.9 | 8.5 | 0.1 | 0.60658 | 0.39025 | 0.44214 | 1.39281 | 0.24736 | 0.18526 | 0.18179 |
| 401 | Plantago lanceolata L. | Plantago lanceolata | 0.21 | 1 | 0.020 | 6.3 | NA | 5.9 | 6.3 | 6.9 | 0.1 | 0.6037 | 0.3946 | 0.44368 | 1.50139 | 0.30164 | 0.21315 | 0.21494 |
| 401 | Potentilla reptans L. | Potentilla reptans | 0.15 | 1 | 0.038 | 8.1 | NA | NA | 5.2 | NA | 0.1 | 0.51276 | 0.49581 | 1.51667 | 2.20989 | 0.98959 | 0.46997 | 0.3594 |
| 401 | Trifolium repens L. | Trifolium repens | 0.1 | 1 | 0.029 | 7.8 | NA | 4.6 | 6.2 | 5.8 | 0.1 | 0.71321 | 0.68276 | 1.29546 | 1.8275 | 0.62188 | 0.18605 | 0.48742 |
| 401 | Medicago lupulina L. | Medicago lupulina | 0.08 | 1 | 0.053 | NA | NA | NA | NA | NA | NA | NA | NA | NA | NA | NA | NA | NA |
| 401 | Bellis perennis L. | Bellis perennis | 0.08 | 1 | 0.091 | NA | NA | NA | NA | NA | NA | NA | NA | NA | NA | NA | NA | NA |
| 401 | Achillea millefolium L. | Achillea millefolium millefolium | 0.05 | 1 | 0.020 | 6.7 | NA | NA | 6.5 | 7.7 | 0.1 | 0.70945 | 0.61373 | 0.80583 | 1.7717 | 0.43958 | 0.26577 | 0.51423 |
| 401 | Veronica persica Poir. | Veronica persica (tournefortii) | 0.02 | 1 | 0.054 | 8.1 | NA | NA | 4.4 | 2.8 | 0.3 | 0.48868 | 0.46508 | 1.4426 | 2.04965 | 0.7278 | 0.33931 | 0.35628 |
| 401 | Poa annua L. | Poa annua | 0.25 | 1 | 0.094 | 6.6 | 5.6 | NA | 5.6 | 4.1 | 0.1 | 0.50969 | 0.50576 | 1.97957 | 2.23767 | 1.1694 | 0.28385 | 0.36995 |
| 402 | Taraxacum officinale F.H.Wigg. | Taraxacum officinale agg. | 0.06 | 0.97 | 0.022 | 6.7 | 5.3 | 4.5 | 5.3 | 5.8 | 0 | 0.75464 | 0.73051 | 1.36974 | 1.95508 | 0.45974 | 0.15567 | 0.6659 |
| 402 | Plantago lanceolata L. | Plantago lanceolata | 0.18 | 0.97 | 0.100 | NA | NA | NA | NA | NA | NA | NA | NA | NA | NA | NA | NA | NA |
| 402 | Potentilla reptans L. | Potentilla reptans | 0.18 | 0.97 | 0.020 | 8.1 | NA | NA | 5.2 | NA | 0.1 | 0.51276 | 0.49581 | 1.51667 | 2.20989 | 0.98959 | 0.46997 | 0.3594 |
| 402 | Trifolium repens L. | Trifolium repens | 0.1 | 0.97 | 0.028 | 7.3 | NA | 3.9 | 6.1 | 3.9 | 0.1 | 0.55903 | 0.41538 | 0.65189 | 1.65181 | 0.43719 | 0.24279 | 0.21709 |
| 402 | Trifolium dubium Sibth. | Trifolium dubium (minus) | 0.13 | 0.97 | 0.062 | 7.3 | NA | 3.9 | 6.1 | 3.9 | 0.1 | 0.55903 | 0.41538 | 0.65189 | 1.65181 | 0.43719 | 0.24279 | 0.21709 |
| 402 | Bellis perennis L. | Bellis perennis | 0.04 | 0.97 | 0.023 | 6.8 | 6.2 | 4.4 | NA | NA | 0.3 | 0.7871 | 0.75023 | 1.37944 | 1.92919 | 0.40631 | 0.19607 | 0.63681 |
| 402 | Achillea millefolium L. | Achillea millefolium millefolium | 0.09 | 0.97 | 0.041 | 8.5 | NA | 3.5 | 6.3 | 4.1 | 0.2 | 0.57633 | 0.55499 | 1.44489 | 1.81436 | 0.579 | 0.25616 | 0.27657 |
| 402 | Veronica persica Poir. | Veronica persica (tournefortii) | 0.02 | 0.97 | 0.022 | 7.7 | NA | 3.4 | 7.8 | 3.7 | 0 | 0.50721 | 0.46976 | 1.21812 | 1.88354 | 0.85158 | 0.24893 | 0.19939 |
| 402 | Poa annua L. | Poa annua | 0.15 | 0.97 | 0.022 | NA | NA | NA | 6.8 | 7.5 | 0 | 0.62243 | 0.29646 | 0.26317 | 1.34606 | 0.11632 | 0.1748 | 0.13996 |
| 402 | Erigeron canadensis L. | Conyza canadensis | 0.02 | 0.97 | 0.021 | 6.6 | 5.7 | NA | 6.3 | 7.6 | 0.1 | 0.67317 | 0.45023 | 0.43732 | 1.4267 | 0.22701 | 0.17304 | 0.27906 |
| 403 | Taraxacum officinale F.H.Wigg. | Taraxacum officinale agg. | 0.25 | 1 | 0.080 | 6.7 | NA | NA | 6.5 | 7.7 | 0.1 | 0.70945 | 0.61373 | 0.80583 | 1.7717 | 0.43958 | 0.26577 | 0.51423 |
| 403 | Lolium perenne L. | Lolium perenne | 0.35 | 1 | 0.083 | 7.3 | 5.4 | 4.1 | 7.6 | 4.2 | 0.4 | 0.58728 | 0.56055 | 1.34601 | 1.94534 | 0.82357 | 0.23743 | 0.34603 |
| 403 | Poa annua L. | Poa annua | 0.22 | 1 | 0.106 | 6.8 | NA | 5.6 | 5.8 | NA | 0.1 | 0.52169 | 0.45097 | 0.93044 | 1.87015 | 0.67593 | 0.2975 | 0.27054 |
| 403 | Bellis perennis L. | Bellis perennis | 0.04 | 1 | 0.115 | 6.8 | NA | 5.6 | 5.8 | NA | 0.1 | 0.52169 | 0.45097 | 0.93044 | 1.87015 | 0.67593 | 0.2975 | 0.27054 |
| 403 | Trifolium dubium Sibth. | Trifolium dubium (minus) | 0.1 | 1 | 0.011 | 7.9 | NA | 4.1 | 5.8 | NA | 0.9 | 0.54139 | 0.51128 | 1.31363 | 1.93633 | 0.8112 | 0.25085 | 0.32953 |
| 403 | Veronica arvensis L. | Veronica arvensis | 0.02 | 1 | 0.119 | 8.1 | NA | NA | 4.4 | 2.8 | 0.3 | 0.48868 | 0.46508 | 1.4426 | 2.04965 | 0.7278 | 0.33931 | 0.35628 |
| 403 | Stellaria media (L.) Vill. | Stellaria media agg. | 0.02 | 1 | 0.150 | 7.4 | 5.3 | NA | 6 | 7.7 | 0.8 | 0.65201 | 0.63791 | 1.54898 | 2.03286 | 0.77966 | 0.37419 | 0.45721 |
| 404 | Taraxacum officinale F.H.Wigg. | Taraxacum officinale agg. | 0.2 | 1 | 0.181 | 7.4 | 5.3 | NA | 6 | 7.7 | 0.8 | 0.65201 | 0.63791 | 1.54898 | 2.03286 | 0.77966 | 0.37419 | 0.45721 |
| 404 | Lolium perenne L. | Lolium perenne | 0.25 | 1 | 0.077 | NA | NA | NA | NA | NA | NA | NA | NA | NA | NA | NA | NA | NA |
| 404 | Poa annua L. | Poa annua | 0.2 | 1 | 0.011 | 7.4 | 5.4 | 4.4 | 7.7 | 6.4 | 0.1 | 0.83003 | 0.82471 | 1.78482 | 1.97219 | 0.48138 | 0.13604 | 0.7566 |
| 404 | Bellis perennis L. | Bellis perennis | 0.21 | 1 | 0.011 | NA | NA | NA | NA | NA | NA | NA | NA | NA | NA | NA | NA | NA |
| 404 | Medicago lupulina L. | Medicago lupulina | 0.04 | 1 | 0.047 | 6.7 | NA | NA | 5.2 | NA | 0.4 | 0.59706 | 0.49429 | 0.69366 | 1.8006 | 0.482 | 0.19483 | 0.19138 |
| 404 | Geranium molle L. | Geranium molle | 0.04 | 1 | 0.115 | 8.1 | NA | NA | 4.4 | 2.8 | 0.3 | 0.48868 | 0.46508 | 1.4426 | 2.04965 | 0.7278 | 0.33931 | 0.35628 |
| 404 | Plantago major L. | Plantago major ssp. major | 0.04 | 1 | 0.043 | 6.7 | NA | 4.9 | 6.8 | 5 | 0 | 0.66057 | 0.18432 | 0.11528 | 1.34075 | 0.06671 | 0.20032 | 0.12881 |
| 404 | Veronica arvensis L. | Veronica arvensis | 0.02 | 1 | 0.075 | 7.4 | 5.3 | NA | 6 | 7.7 | 0.8 | 0.65201 | 0.63791 | 1.54898 | 2.03286 | 0.77966 | 0.37419 | 0.45721 |
| 405 | Taraxacum officinale F.H.Wigg. | Taraxacum officinale agg. | 0.8 | 1 | 0.169 | 7.4 | 5.3 | NA | 6 | 7.7 | 0.8 | 0.65201 | 0.63791 | 1.54898 | 2.03286 | 0.77966 | 0.37419 | 0.45721 |
| 405 | Solanum nigrum L. | Solanum nigrum | 0.2 | 1 | 0.011 | 7 | 5.6 | 4.4 | 5.9 | 5.4 | 0.1 | 0.64871 | 0.63945 | 1.70759 | 2.05577 | 0.56937 | 0.21649 | 0.54803 |
| 406 | Sonchus oleraceus L. | Sonchus oleraceus | 0.34 | 1 | 0.060 | 8.1 | NA | 3.5 | 7.5 | 5 | 0.3 | 0.75184 | 0.72654 | 1.3537 | 1.85024 | 0.717 | 0.18024 | 0.48387 |
| 406 | Clematis vitalba L. | Clematis vitalba | 0.33 | 1 | 0.011 | 7.8 | 5.1 | NA | NA | NA | 0.2 | 0.51255 | 0.49092 | 1.42792 | 1.96794 | 1.00937 | 0.22916 | 0.30884 |
| 406 | Solanum nigrum L. | Solanum nigrum | 0.33 | 1 | 0.087 | 7.4 | 5.3 | NA | 6 | 7.7 | 0.8 | 0.65201 | 0.63791 | 1.54898 | 2.03286 | 0.77966 | 0.37419 | 0.45721 |
| 407 | Taraxacum officinale F.H.Wigg. | Taraxacum officinale agg. | 0.25 | 0.75 | 0.060 | 7.3 | 5 | NA | 6.6 | 2.5 | 0.4 | 0.4995 | 0.45471 | 1.13703 | 1.80119 | 0.66561 | 0.27734 | 0.25548 |
| 407 | Oxalis corniculata L. | Oxalis corniculata | 0.5 | 0.75 | 0.054 | 8.8 | NA | 3.9 | 7.5 | 7.4 | 0.2 | 0.71714 | 0.69501 | 1.38943 | 1.78673 | 0.86446 | 0.18028 | 0.37675 |
| 409 | Taraxacum officinale F.H.Wigg. | Taraxacum officinale agg. | 0.5 | 0.9 | 0.087 | 6.8 | NA | 5.6 | 5.8 | NA | 0.1 | 0.52169 | 0.45097 | 0.93044 | 1.87015 | 0.67593 | 0.2975 | 0.27054 |
| 409 | Erigeron canadensis L. | Conyza canadensis | 0.3 | 0.9 | 0.022 | 6.3 | 5 | 7.3 | 5.8 | 6.5 | 0.7 | 0.53692 | 0.48641 | 1.02734 | 1.80642 | 0.6097 | 0.33182 | 0.26598 |
| 409 | Capsella bursa-pastoris Medik. | Capsella bursa-pastoris | 0.1 | 0.9 | 0.000 | NA | NA | NA | NA | NA | NA | NA | NA | NA | NA | NA | NA | NA |
| 411 | Taraxacum officinale F.H.Wigg. | Taraxacum officinale agg. | 0.18 | 0.9 | 0.000 | NA | NA | NA | NA | NA | NA | NA | NA | NA | NA | NA | NA | NA |
| 411 | Medicago lupulina L. | Medicago lupulina | 0.08 | 0.9 | 0.000 | NA | NA | NA | NA | NA | NA | NA | NA | NA | NA | NA | NA | NA |
| 411 | Poa annua L. | Poa annua | 0.25 | 0.9 | 0.000 | NA | NA | NA | NA | NA | NA | NA | NA | NA | NA | NA | NA | NA |
| 411 | Erodium cicutarium (L.) | Erodium cicutarium cicutarium | 0.04 | 0.9 | 0.000 | NA | NA | NA | NA | NA | NA | NA | NA | NA | NA | NA | NA | NA |
| 411 | Bromus sterilis L. | Bromus sterilis | 0.15 | 0.9 | 0.000 | NA | NA | NA | NA | NA | NA | NA | NA | NA | NA | NA | NA | NA |
| 411 | Ranunculus repens L. | Ranunculus repens | 0.2 | 0.9 | 0.000 | NA | NA | NA | NA | NA | NA | NA | NA | NA | NA | NA | NA | NA |
| 412 | Galium mollugo L. | Galium mollugo mollugo (elatum) | 0.21 | 0.76 | 0.276 | 7.2 | 5.7 | 4.3 | 6.8 | NA | 0.1 | 0.54405 | 0.41742 | 0.70503 | 1.63482 | 0.49706 | 0.18636 | 0.24515 |
| 412 | Pyracantha coccinea M.Roem. | Pyracantha coccinea | 0.15 | 0.76 | 1.000 | 7.3 | NA | 3.9 | 6.1 | 3.9 | 0.1 | 0.55903 | 0.41538 | 0.65189 | 1.65181 | 0.43719 | 0.24279 | 0.21709 |
| 412 | Taraxacum officinale F.H.Wigg. | Taraxacum officinale agg. | 0.11 | 0.76 | 0.769 | 7.2 | 5.7 | 4.3 | 6.8 | NA | 0.1 | 0.54405 | 0.41742 | 0.70503 | 1.63482 | 0.49706 | 0.18636 | 0.24515 |
| 412 | Bromus sterilis L. | Bromus sterilis | 0.15 | 0.76 | 0.143 | 4.8 | 5.1 | 6.3 | 6.6 | 6.9 | 0 | 0.67651 | 0.23653 | 0.12754 | 1.34711 | 0.06518 | 0.18881 | 0.12802 |
| 412 | Rumex acetosa L. | Rumex acetosa | 0.08 | 0.76 | 0.455 | 7.2 | 5.7 | 4.3 | 6.8 | NA | 0.1 | 0.54405 | 0.41742 | 0.70503 | 1.63482 | 0.49706 | 0.18636 | 0.24515 |
| 412 | Arenaria serpyllifolia L. | Arenaria serpyllifolia agg. | 0.02 | 0.76 | 1.000 | 8.4 | 7.2 | 2.6 | 6.8 | 3.6 | 0.2 | 0.58203 | 0.49726 | 0.89718 | 1.5951 | 0.3257 | 0.17939 | 0.36267 |
| 412 | Hypericum perforatum L. | Hypericum perforatum | 0.02 | 0.76 | 0.667 | 8.4 | 7.2 | 2.6 | 6.8 | 3.6 | 0.2 | 0.58203 | 0.49726 | 0.89718 | 1.5951 | 0.3257 | 0.17939 | 0.36267 |
| 412 | Achillea millefolium L. | Achillea millefolium millefolium | 0.02 | 0.76 | 1.000 | 8.4 | 7.2 | 2.6 | 6.8 | 3.6 | 0.2 | 0.58203 | 0.49726 | 0.89718 | 1.5951 | 0.3257 | 0.17939 | 0.36267 |

| | | | | | | | | | | | | | | | | | |
|---|---|---|---|---|---|---|---|---|---|---|---|---|---|---|---|---|---|
| 413 | Galium mollugo L. | Galium mollugo mollugo (elatum) | 0.2 | 0.62 | 0.167 | NA | NA | NA | 6.6 | 7 | 0 | 0.72781 | 0.47089 | 0.36365 | 1.47514 | 0.21383 | 0.17215 | 0.37071 |
| 413 | Brachypodium sylvaticum (Huds.) P.Beauv. | Brachypodium sylvaticum | 0.25 | 0.62 | 0.161 | 7.7 | NA | NA | 5.6 | NA | 0.7 | 0.53223 | 0.52325 | 1.73719 | 2.14421 | 0.94278 | 0.39374 | 0.35054 |
| 413 | Hedera helix L. | Hedera helix | 0.05 | 0.62 | 0.013 | 7.3 | 6.1 | 3.8 | 6 | 5.1 | 0 | 0.62987 | 0.59281 | 1.27995 | 1.90491 | 0.6275 | 0.3036 | 0.42608 |
| 413 | Acer platanoides L. | Acer platanoides | 0.12 | 0.62 | 0.031 | NA | NA | NA | NA | NA | NA | NA | NA | NA | NA | NA | NA | NA |
| 414 | Galium mollugo L. | Galium mollugo mollugo (elatum) | 0.25 | 0.51 | 0.055 | 8.1 | NA | NA | 5.2 | NA | 0.1 | 0.51276 | 0.49581 | 1.51667 | 2.20989 | 0.98959 | 0.46997 | 0.3594 |
| 414 | Medicago lupulina L. | Medicago lupulina | 0.12 | 0.51 | 0.471 | 7.4 | 5.3 | NA | 6 | 7.7 | 0.8 | 0.65201 | 0.63791 | 1.54898 | 2.03286 | 0.77966 | 0.37419 | 0.45721 |
| 414 | Origanum vulgare L. | Origanum vulgare | 0.1 | 0.51 | 0.167 | 7.8 | NA | 4.6 | 6.2 | 5.8 | 0.1 | 0.71321 | 0.68276 | 1.29546 | 1.8275 | 0.62188 | 0.18605 | 0.48742 |
| 414 | Hypericum perforatum L. | Hypericum perforatum | 0.02 | 0.51 | 0.455 | 7.8 | NA | 4.6 | 6.2 | 5.8 | 0.1 | 0.71321 | 0.68276 | 1.29546 | 1.8275 | 0.62188 | 0.18605 | 0.48742 |
| 414 | Taraxacum officinale F.H.Wigg. | Taraxacum officinale agg. | 0.02 | 0.51 | 0.857 | 7.8 | NA | 4.6 | 6.2 | 5.8 | 0.1 | 0.71321 | 0.68276 | 1.29546 | 1.8275 | 0.62188 | 0.18605 | 0.48742 |
| 417 | Hypericum perforatum L. | Hypericum perforatum | 0.07 | 0.07 | 0.750 | 7.8 | NA | 4.6 | 6.2 | 5.8 | 0.1 | 0.71321 | 0.68276 | 1.29546 | 1.8275 | 0.62188 | 0.18605 | 0.48742 |
| 418 | Galium mollugo L. | Galium mollugo mollugo (elatum) | 0.1 | 0.13 | 0.303 | 7.9 | 5.4 | 2.9 | 7.9 | 2.4 | 0 | 0.48324 | 0.22836 | 0.33987 | 1.12718 | 0.05497 | 0.20766 | 0.12487 |
| 418 | Hypericum perforatum L. | Hypericum perforatum | 0.03 | 0.13 | 0.462 | 7.1 | 4.9 | NA | 6 | NA | 1 | 0.59517 | 0.59064 | 1.89772 | 2.12267 | 0.65091 | 0.41213 | 0.37185 |
| 419 | Taraxacum officinale F.H.Wigg. | Taraxacum officinale agg. | 0.1 | 0.14 | 0.099 | 7.4 | 5.3 | NA | 6 | 7.7 | 0.8 | 0.65201 | 0.63791 | 1.54898 | 2.03286 | 0.77966 | 0.37419 | 0.45721 |
| 419 | Senecio inaequidens DC. | Senecio inaequidens | 0.02 | 0.14 | 0.188 | 6.7 | NA | NA | 6.5 | 7.7 | 0.1 | 0.70945 | 0.61373 | 0.80583 | 1.7717 | 0.43958 | 0.26577 | 0.51423 |
| 419 | Alliaria petiolata (M.Bieb.) | Alliaria petiolata | 0.02 | 0.14 | 0.500 | NA | NA | NA | NA | NA | NA | NA | NA | NA | NA | NA | NA | NA |
| 420 | Scrophularia nodosa L. | Scrophularia nodosa | 0.05 | 0.35 | 0.197 | 7.3 | 7.3 | 4.7 | 6 | 3.5 | 0 | 0.68706 | 0.13477 | 0.05158 | 1.3063 | 0.01001 | 0.21919 | 0.10465 |
| 420 | Hypericum perforatum L. | Hypericum perforatum | 0.07 | 0.35 | 0.323 | 7.2 | 5.7 | 4.3 | 6.8 | NA | 0.1 | 0.54405 | 0.41742 | 0.70503 | 1.63482 | 0.49706 | 0.18636 | 0.24515 |
| 420 | Cirsium arvense (L.) Scop. | Cirsium arvense | 0.05 | 0.35 | 0.490 | 7.2 | 5.7 | 4.3 | 6.8 | NA | 0.1 | 0.54405 | 0.41742 | 0.70503 | 1.63482 | 0.49706 | 0.18636 | 0.24515 |
| 420 | Lamium purpureum L. | Lamium purpureum | 0.05 | 0.35 | 0.231 | 7.3 | NA | 3.9 | 6.1 | 3.9 | 0.1 | 0.55903 | 0.41538 | 0.65189 | 1.65181 | 0.43719 | 0.24279 | 0.21709 |
| 420 | Rubus armeniacus Focke | NA | 0.02 | 0.35 | 0.714 | NA | NA | NA | NA | NA | NA | NA | NA | NA | NA | NA | NA | NA |
| 420 | Tanacetum vulgare L. | Tanacetum vulgare | 0.04 | 0.35 | 0.200 | 7.3 | NA | 3.9 | 6.1 | 3.9 | 0.1 | 0.55903 | 0.41538 | 0.65189 | 1.65181 | 0.43719 | 0.24279 | 0.21709 |
| 420 | Acer platanoides L. | Acer platanoides | 0.02 | 0.35 | 0.364 | NA | NA | NA | NA | NA | NA | NA | NA | NA | NA | NA | NA | NA |
| 420 | Cardamine hirsuta L. | Cardamine hirsuta | 0.05 | 0.35 | 0.156 | 6.7 | NA | 4.9 | 6.8 | 5 | 0 | 0.66057 | 0.18432 | 0.11528 | 1.34075 | 0.06671 | 0.20032 | 0.12881 |
| 421 | Galium mollugo L. | Galium mollugo mollugo (elatum) | 0.25 | 0.55 | 0.290 | NA | NA | 5.3 | 6.2 | NA | 0.1 | 0.68682 | 0.14816 | 0.05813 | 1.31254 | 0.01433 | 0.20126 | 0.10505 |
| 421 | Bromus sterilis L. | Bromus sterilis | 0.2 | 0.55 | 0.128 | NA | NA | NA | NA | NA | NA | NA | NA | NA | NA | NA | NA | NA |
| 421 | Taraxacum officinale F.H.Wigg. | Taraxacum officinale agg. | 0.05 | 0.55 | 0.079 | 6.5 | NA | NA | 5.4 | NA | 0 | 0.59769 | 0.45571 | 0.64594 | 1.51584 | 0.28825 | 0.22901 | 0.32316 |
| 421 | Achillea millefolium L. | Achillea millefolium millefolium | 0.05 | 0.55 | 0.556 | 7.2 | 5.7 | 4.3 | 6.8 | NA | 0.1 | 0.54405 | 0.41742 | 0.70503 | 1.63482 | 0.49706 | 0.18636 | 0.24515 |
| 422 | Fragaria vesca L. | Fragaria vesca | 0.05 | 0.32 | 0.286 | 7.2 | 5.7 | 4.3 | 6.8 | NA | 0.1 | 0.54405 | 0.41742 | 0.70503 | 1.63482 | 0.49706 | 0.18636 | 0.24515 |
| 422 | Galium mollugo L. | Galium mollugo mollugo (elatum) | 0.05 | 0.32 | 0.259 | NA | NA | NA | 6.6 | 7 | 0 | 0.72781 | 0.47089 | 0.36365 | 1.47514 | 0.21383 | 0.17215 | 0.37071 |
| 422 | Hypericum perforatum L. | Hypericum perforatum | 0.07 | 0.32 | 0.243 | NA | NA | NA | 6.6 | 7 | 0 | 0.72781 | 0.47089 | 0.36365 | 1.47514 | 0.21383 | 0.17215 | 0.37071 |
| 422 | Achillea millefolium L. | Achillea millefolium millefolium | 0.02 | 0.32 | 0.167 | NA | NA | NA | NA | NA | NA | NA | NA | NA | NA | NA | NA | NA |
| 422 | Plantago lanceolata L. | Plantago lanceolata | 0.05 | 0.32 | 0.209 | NA | NA | NA | 6.6 | 7 | 0 | 0.72781 | 0.47089 | 0.36365 | 1.47514 | 0.21383 | 0.17215 | 0.37071 |
| 422 | Geranium robertianum L. | Geranium robertianum | 0.06 | 0.32 | 0.397 | NA | NA | NA | NA | NA | NA | NA | NA | NA | NA | NA | NA | NA |
| 422 | Erigeron canadensis L. | Conyza canadensis | 0.02 | 0.32 | 0.446 | NA | NA | NA | NA | NA | NA | NA | NA | NA | NA | NA | NA | NA |
| 423 | Hedera helix L. | Hedera helix | 0.2 | 0.69 | 0.277 | NA | NA | NA | NA | NA | NA | NA | NA | NA | NA | NA | NA | NA |
| 423 | Erigeron canadensis L. | Conyza canadensis | 0.13 | 0.69 | 0.300 | NA | NA | NA | NA | NA | NA | NA | NA | NA | NA | NA | NA | NA |
| 423 | Hypericum perforatum L. | Hypericum perforatum | 0.05 | 0.69 | 0.013 | NA | NA | NA | NA | NA | NA | NA | NA | NA | NA | NA | NA | NA |
| 423 | Plantago lanceolata L. | Plantago lanceolata | 0.15 | 0.69 | 0.221 | NA | NA | NA | NA | NA | NA | NA | NA | NA | NA | NA | NA | NA |
| 423 | Galium mollugo L. | Galium mollugo mollugo (elatum) | 0.02 | 0.69 | 0.205 | NA | NA | NA | NA | NA | NA | NA | NA | NA | NA | NA | NA | NA |
| 423 | Arenaria serpyllifolia L. | Arenaria serpyllifolia agg. | 0.02 | 0.69 | 0.179 | 7.3 | NA | 3.9 | 6.1 | 3.9 | 0.1 | 0.55903 | 0.41538 | 0.65189 | 1.65181 | 0.43719 | 0.24279 | 0.21709 |
| 423 | Medicago lupulina L. | Medicago lupulina | 0.08 | 0.69 | 0.146 | NA | NA | NA | NA | NA | NA | NA | NA | NA | NA | NA | NA | NA |
| 423 | Bromus sterilis L. | Bromus sterilis | 0.04 | 0.69 | 0.446 | 7.3 | 5.4 | 4.1 | 7.6 | 4.2 | 0.4 | 0.58728 | 0.56055 | 1.34601 | 1.94534 | 0.82357 | 0.23743 | 0.34603 |
| 424 | Bromus sterilis L. | Bromus sterilis | 0.06 | 0.47 | 0.267 | 6.9 | 6.5 | NA | 7.3 | 6.3 | 0 | 0.67274 | 0.19896 | 0.13039 | 1.32576 | 0.04466 | 0.20186 | 0.11616 |
| 424 | Achillea millefolium L. | Achillea millefolium millefolium | 0.02 | 0.47 | 0.417 | 7.4 | 5.3 | NA | 6 | 7.7 | 0.8 | 0.65201 | 0.63791 | 1.54898 | 2.03286 | 0.77966 | 0.37419 | 0.45721 |
| 424 | Hypericum perforatum L. | Hypericum perforatum | 0.06 | 0.47 | 0.099 | 7.3 | 6.1 | 3.8 | 6 | 5.1 | 0 | 0.62987 | 0.59281 | 1.27995 | 1.90491 | 0.6275 | 0.3036 | 0.42608 |
| 424 | Medicago lupulina L. | Medicago lupulina | 0.05 | 0.47 | 0.123 | 7.3 | 6.1 | 3.8 | 6 | 5.1 | 0 | 0.62987 | 0.59281 | 1.27995 | 1.90491 | 0.6275 | 0.3036 | 0.42608 |
| 424 | Galium mollugo L. | Galium mollugo mollugo (elatum) | 0.02 | 0.47 | 0.338 | NA | NA | NA | NA | NA | NA | NA | NA | NA | NA | NA | NA | NA |
| 424 | Cerastium brachypetalum Pers. | Cerastium brachypetalum | 0.05 | 0.47 | 0.067 | 7.4 | 5.3 | NA | 6 | 7.7 | 0.8 | 0.65201 | 0.63791 | 1.54898 | 2.03286 | 0.77966 | 0.37419 | 0.45721 |
| 424 | Geranium robertianum L. | Geranium robertianum | 0.2 | 0.47 | 0.139 | NA | NA | NA | NA | NA | NA | NA | NA | NA | NA | NA | NA | NA |
| 424 | Arenaria serpyllifolia L. | Arenaria serpyllifolia agg. | 0.01 | 0.47 | 0.059 | 7.9 | 5.1 | 5.1 | 6.7 | 7.7 | 0.2 | 0.75642 | 0.75467 | 2.08644 | 2.23861 | 0.83282 | 0.22413 | 0.56443 |
| 425 | Cardamine hirsuta L. | Cardamine hirsuta | 0.05 | 0.63 | 0.147 | 7.4 | 5.3 | NA | 6 | 7.7 | 0.8 | 0.65201 | 0.63791 | 1.54898 | 2.03286 | 0.77966 | 0.37419 | 0.45721 |
| 425 | Poa pratensis L. | Poa pratensis aggr. | 0.38 | 0.63 | 0.137 | NA | NA | NA | NA | NA | NA | NA | NA | NA | NA | NA | NA | NA |
| 425 | Medicago lupulina L. | Medicago lupulina | 0.05 | 0.63 | 0.259 | NA | NA | NA | NA | NA | NA | NA | NA | NA | NA | NA | NA | NA |
| 425 | Hypericum perforatum L. | Hypericum perforatum | 0.05 | 0.63 | 0.250 | 8.4 | 7.2 | 2.6 | 6.8 | 3.6 | 0.2 | 0.58203 | 0.49726 | 0.89718 | 1.5951 | 0.3257 | 0.17939 | 0.36267 |
| 425 | Achillea millefolium L. | Achillea millefolium millefolium | 0.05 | 0.63 | 0.135 | 7.1 | 4.9 | NA | 6 | NA | 1 | 0.59517 | 0.59064 | 1.89772 | 2.12267 | 0.65091 | 0.41213 | 0.37185 |
| 425 | Plantago lanceolata L. | Plantago lanceolata | 0.05 | 0.63 | 0.254 | NA | NA | NA | NA | NA | NA | NA | NA | NA | NA | NA | NA | NA |
| 426 | Galium mollugo L. | Galium mollugo mollugo (elatum) | 0.2 | 0.36 | 0.233 | NA | NA | NA | NA | NA | NA | NA | NA | NA | NA | NA | NA | NA |
| 426 | Hypericum perforatum L. | Hypericum perforatum | 0.05 | 0.36 | 0.152 | 7 | 5.6 | 4.4 | 5.9 | 5.4 | 0.1 | 0.64871 | 0.63945 | 1.70759 | 2.05577 | 0.56937 | 0.21649 | 0.54803 |
| 426 | Achillea millefolium L. | Achillea millefolium millefolium | 0.02 | 0.36 | 0.169 | NA | NA | NA | NA | NA | NA | NA | NA | NA | NA | NA | NA | NA |
| 426 | Medicago lupulina L. | Medicago lupulina | 0.02 | 0.36 | 0.221 | NA | NA | NA | NA | NA | NA | NA | NA | NA | NA | NA | NA | NA |
| 426 | Cardamine hirsuta L. | Cardamine hirsuta | 0.02 | 0.36 | 0.500 | NA | NA | NA | NA | NA | NA | NA | NA | NA | NA | NA | NA | NA |
| 426 | Poa annua L. | Poa annua | 0.01 | 0.36 | 0.159 | NA | NA | NA | NA | NA | NA | NA | NA | NA | NA | NA | NA | NA |
| 426 | Bromus sterilis L. | Bromus sterilis | 0.04 | 0.36 | 0.322 | NA | NA | NA | NA | NA | NA | NA | NA | NA | NA | NA | NA | NA |
| 427 | Galium mollugo L. | Galium mollugo mollugo (elatum) | 0.18 | 0.63 | 0.273 | NA | NA | NA | NA | NA | NA | NA | NA | NA | NA | NA | NA | NA |
| 427 | Bromus sterilis L. | Bromus sterilis | 0.35 | 0.63 | 0.875 | 7.4 | 5.3 | NA | 6 | 7.7 | 0.8 | 0.65201 | 0.63791 | 1.54898 | 2.03286 | 0.77966 | 0.37419 | 0.45721 |
| 427 | Achillea millefolium L. | Achillea millefolium millefolium | 0.05 | 0.63 | 0.538 | 7.4 | 5.3 | NA | 6 | 7.7 | 0.8 | 0.65201 | 0.63791 | 1.54898 | 2.03286 | 0.77966 | 0.37419 | 0.45721 |
| 427 | Taraxacum officinale F.H.Wigg. | Taraxacum officinale agg. | 0.02 | 0.63 | 0.400 | 7.1 | 4.9 | NA | 6 | NA | 1 | 0.59517 | 0.59064 | 1.89772 | 2.12267 | 0.65091 | 0.41213 | 0.37185 |
| 427 | Hypericum perforatum L. | Hypericum perforatum | 0.02 | 0.63 | 0.429 | NA | NA | NA | NA | NA | NA | NA | NA | NA | NA | NA | NA | NA |
| 427 | Medicago lupulina L. | Medicago lupulina | 0.01 | 0.63 | 0.308 | 6.7 | NA | NA | 6.5 | 7.7 | 0.1 | 0.70945 | 0.61373 | 0.80583 | 1.7717 | 0.43958 | 0.26577 | 0.51423 |
| 428 | Senecio inaequidens DC. | Senecio inaequidens | 0.2 | 0.2 | 0.423 | NA | NA | NA | NA | NA | NA | NA | NA | NA | NA | NA | NA | NA |
| 429 | Senecio inaequidens DC. | Senecio inaequidens | 0.3 | 0.45 | 0.455 | NA | NA | NA | NA | NA | NA | NA | NA | NA | NA | NA | NA | NA |
| 429 | Taraxacum officinale F.H.Wigg. | Taraxacum officinale agg. | 0.15 | 0.45 | 0.500 | 7.4 | 5.3 | NA | 6 | 7.7 | 0.8 | 0.65201 | 0.63791 | 1.54898 | 2.03286 | 0.77966 | 0.37419 | 0.45721 |
| 431 | Senecio inaequidens DC. | Senecio inaequidens | 0.1 | 0.1 | 0.500 | 7.4 | 5.3 | NA | 6 | 7.7 | 0.8 | 0.65201 | 0.63791 | 1.54898 | 2.03286 | 0.77966 | 0.37419 | 0.45721 |
| 433 | Lapsana communis L. | Lapsana communis | 0.15 | 0.58 | 0.238 | 7.4 | 5.2 | 4.7 | 6.1 | NA | 0 | 0.77406 | 0.76726 | 1.79113 | 2.07279 | 0.64778 | 0.2119 | 0.65925 |
| 433 | Taraxacum officinale F.H.Wigg. | Taraxacum officinale agg. | 0.13 | 0.58 | 0.145 | NA | NA | NA | NA | NA | NA | NA | NA | NA | NA | NA | NA | NA |
| 433 | Veronica hederifolia L. | Veronica hederifolia agg. | 0.08 | 0.58 | 0.143 | 8 | 5 | NA | 6.1 | 6.6 | 0.4 | 0.62393 | 0.60267 | 1.39471 | 1.77442 | 0.6433 | 0.24014 | 0.43141 |
| 433 | Bromus sterilis L. | Bromus sterilis | 0.1 | 0.58 | 0.156 | 7.2 | 5.7 | 4.3 | 6.8 | NA | 0.1 | 0.54405 | 0.41742 | 0.70503 | 1.63482 | 0.49706 | 0.18636 | 0.24515 |
| 433 | Poa annua L. | Poa annua | 0.05 | 0.58 | 0.188 | 7.8 | NA | 4.6 | 6.2 | 5.8 | 0.1 | 0.71321 | 0.68276 | 1.29546 | 1.8275 | 0.62188 | 0.18605 | 0.48742 |
| 433 | Stellaria media (L.) Vill. | Stellaria media agg. | 0.05 | 0.58 | 0.556 | NA | NA | NA | NA | NA | NA | NA | NA | NA | NA | NA | NA | NA |
| 433 | Veronica arvensis L. | Veronica arvensis | 0.02 | 0.58 | 0.224 | NA | NA | NA | NA | NA | NA | NA | NA | NA | NA | NA | NA | NA |
| 434 | Lapsana communis L. | Lapsana communis | 0.17 | 0.7 | 0.186 | NA | NA | NA | NA | NA | NA | NA | NA | NA | NA | NA | NA | NA |
| 434 | Taraxacum officinale F.H.Wigg. | Taraxacum officinale agg. | 0.13 | 0.7 | 0.333 | NA | NA | 4.5 | 6.6 | 6.6 | 0 | 0.78059 | 0.60963 | 0.54418 | 1.63701 | 0.26733 | 0.16232 | 0.54354 |
| 434 | Sonchus oleraceus L. | Sonchus oleraceus | 0.05 | 0.7 | 0.233 | NA | NA | NA | NA | NA | NA | NA | NA | NA | NA | NA | NA | NA |
| 434 | Veronica arvensis L. | Veronica arvensis | 0.05 | 0.7 | 0.238 | NA | NA | NA | NA | NA | NA | NA | NA | NA | NA | NA | NA | NA |
| 434 | Stellaria media (L.) Vill. | Stellaria media agg. | 0.05 | 0.7 | 0.161 | 6.5 | NA | NA | 5.4 | NA | 0 | 0.59769 | 0.45571 | 0.64594 | 1.51584 | 0.28825 | 0.22901 | 0.32316 |
| 434 | Veronica hederifolia L. | Veronica hederifolia agg. | 0.08 | 0.7 | 0.426 | 6.5 | NA | NA | 5.4 | NA | 0 | 0.59769 | 0.45571 | 0.64594 | 1.51584 | 0.28825 | 0.22901 | 0.32316 |
| 434 | Poa annua L. | Poa annua | 0.08 | 0.7 | 0.300 | 6.2 | 5.5 | NA | 6.9 | 8.5 | 0.1 | 0.60658 | 0.39025 | 0.44214 | 1.39281 | 0.24736 | 0.18526 | 0.18179 |
| 434 | Tanacetum vulgare L. | Tanacetum vulgare | 0.09 | 0.7 | 0.339 | 7.4 | 5.3 | NA | 6 | 7.7 | 0.8 | 0.65201 | 0.63791 | 1.54898 | 2.03286 | 0.77966 | 0.37419 | 0.45721 |
| 435 | Lapsana communis L. | Lapsana communis | 0.1 | 0.6 | 0.015 | 6.5 | NA | NA | 5.4 | NA | 0 | 0.59769 | 0.45571 | 0.64594 | 1.51584 | 0.28825 | 0.22901 | 0.32316 |
| 435 | Taraxacum officinale F.H.Wigg. | Taraxacum officinale agg. | 0.1 | 0.6 | 0.033 | NA | NA | NA | NA | NA | NA | NA | NA | NA | NA | NA | NA | NA |
| 435 | Veronica hederifolia L. | Veronica hederifolia agg. | 0.2 | 0.6 | 0.179 | 4.7 | NA | NA | 5.8 | 6.6 | 0 | 0.68705 | 0.16985 | 0.0706 | 1.32057 | 0.02196 | 0.18337 | 0.11293 |
| 435 | Bromus sterilis L. | Bromus sterilis | 0.15 | 0.6 | 0.074 | 4.7 | 5.8 | 5.6 | 6.5 | 6.3 | 0 | 0.69843 | 0.15019 | 0.04615 | 1.3308 | 0.01233 | 0.18685 | 0.11235 |
| 435 | Hedera helix L. | Hedera helix | 0.05 | 0.6 | 0.411 | 8.1 | NA | 4.1 | 6.6 | NA | 0.7 | 0.57545 | 0.53032 | 1.14497 | 1.72102 | 0.70077 | 0.21269 | 0.32208 |
| 436 | Lapsana communis L. | Lapsana communis | 0.09 | 0.43 | 0.357 | 6.9 | 6.5 | NA | 7.3 | 6.3 | 0 | 0.67274 | 0.19896 | 0.13039 | 1.32576 | 0.04466 | 0.20186 | 0.11616 |
| 436 | Taraxacum officinale F.H.Wigg. | Taraxacum officinale agg. | 0.1 | 0.43 | 0.305 | 7.3 | NA | 3.9 | 6.1 | 3.9 | 0.1 | 0.55903 | 0.41538 | 0.65189 | 1.65181 | 0.43719 | 0.24279 | 0.21709 |
| 436 | Veronica hederifolia L. | Veronica hederifolia agg. | 0.05 | 0.43 | 0.149 | 7.3 | 6.1 | 3.8 | 6 | 5.1 | 0 | 0.62987 | 0.59281 | 1.27995 | 1.90491 | 0.6275 | 0.3036 | 0.42608 |
| 436 | Hedera helix L. | Hedera helix | 0.05 | 0.43 | 0.107 | NA | NA | NA | NA | NA | NA | NA | NA | NA | NA | NA | NA | NA |
| 436 | Cardamine hirsuta L. | Cardamine hirsuta | 0.09 | 0.43 | 0.417 | 7.1 | 4.9 | NA | 6 | NA | 1 | 0.59517 | 0.59064 | 1.89772 | 2.12267 | 0.65091 | 0.41213 | 0.37185 |
| 436 | Ranunculus ficaria L. | Ranunculus ficaria (Ficaria verna) | 0.05 | 0.43 | 0.099 | 8.1 | NA | NA | 4.4 | 2.8 | 0.3 | 0.48868 | 0.46508 | 1.4426 | 2.04965 | 0.7278 | 0.33931 | 0.35628 |
| 437 | Taraxacum officinale F.H.Wigg. | Taraxacum officinale agg. | 0.25 | 0.63 | 0.049 | NA | NA | NA | 6.6 | NA | 0 | 0.65326 | 0.6245 | 1.32492 | 1.96985 | 0.60133 | 0.19748 | 0.49592 |
| 437 | Bromus sterilis L. | Bromus sterilis | 0.15 | 0.63 | 0.185 | 7.1 | 4.9 | NA | 6 | NA | 1 | 0.59517 | 0.59064 | 1.89772 | 2.12267 | 0.65091 | 0.41213 | 0.37185 |
| 437 | Cardamine hirsuta L. | Cardamine hirsuta | 0.1 | 0.63 | 0.120 | NA | NA | NA | NA | NA | NA | NA | NA | NA | NA | NA | NA | NA |
| 437 | Hedera helix L. | Hedera helix | 0.05 | 0.63 | 0.063 | 7.3 | 6.1 | 3.8 | 6 | 5.1 | 0 | 0.62987 | 0.59281 | 1.27995 | 1.90491 | 0.6275 | 0.3036 | 0.42608 |
| 437 | Poa annua L. | Poa annua | 0.08 | 0.63 | 0.059 | 8.1 | NA | 3.1 | 6.6 | NA | 0 | 0.56066 | 0.54465 | 1.58041 | 1.97627 | 0.33838 | 0.29416 | 0.45216 |
| 438 | Taraxacum officinale F.H.Wigg. | Taraxacum officinale agg. | 0.25 | 0.56 | 0.221 | NA | NA | NA | NA | NA | NA | NA | NA | NA | NA | NA | NA | NA |
| 438 | Cardamine hirsuta L. | Cardamine hirsuta | 0.09 | 0.56 | 0.137 | 7 | 5.6 | 4.4 | 5.9 | 5.4 | 0.1 | 0.64871 | 0.63945 | 1.70759 | 2.05577 | 0.56937 | 0.21649 | 0.54803 |

| | | | | | | | | | | | | | | | | | | |
|---|---|---|---|---|---|---|---|---|---|---|---|---|---|---|---|---|---|---|
| 438 | Veronica hederifolia L. | Veronica hederifolia agg. | 0.05 | 0.56 | 0.172 | 7.4 | 5.3 | NA | 6 | 7.7 | 0.8 | 0.65201 | 0.63791 | 1.54898 | 2.03286 | 0.77966 | 0.37419 | 0.45721 |
| 438 | Hedera helix L. | Hedera helix | 0.09 | 0.56 | 0.167 | 7.4 | 5.3 | NA | 6 | 7.7 | 0.8 | 0.65201 | 0.63791 | 1.54898 | 2.03286 | 0.77966 | 0.37419 | 0.45721 |
| 438 | Stellaria media (L.) Vill. | Stellaria media agg. | 0.08 | 0.56 | 0.545 | 5.6 | 6.4 | 5.3 | 4 | 4 | 0 | 0.76452 | 0.18157 | 0.10031 | 1.31072 | 0.01535 | 0.20259 | 0.20926 |
| 439 | Taraxacum officinale F.H.Wigg. | Taraxacum officinale agg. | 0.13 | 0.47 | 0.243 | NA | NA | NA | NA | NA | NA | NA | NA | NA | NA | NA | NA | NA |
| 439 | Cardamine hirsuta L. | Cardamine hirsuta | 0.2 | 0.47 | 0.096 | 7 | 5.6 | 4.4 | 5.9 | 5.4 | 0.1 | 0.64871 | 0.63945 | 1.70759 | 2.05577 | 0.56937 | 0.21649 | 0.54803 |
| 439 | Veronica hederifolia L. | Veronica hederifolia agg. | 0.06 | 0.47 | 0.349 | 7.9 | 5.4 | 2.9 | 7.9 | 2.4 | 0 | 0.48324 | 0.22836 | 0.33987 | 1.12718 | 0.05497 | 0.20766 | 0.12487 |
| 439 | Stellaria media (L.) Vill. | Stellaria media agg. | 0.08 | 0.47 | 0.121 | 8.2 | NA | 2.2 | 7 | 2.1 | 0 | 0.50737 | 0.49072 | 1.5234 | 1.7929 | 0.17792 | 0.32947 | 0.3784 |
| 440 | Taraxacum officinale F.H.Wigg. | Taraxacum officinale agg. | 0.15 | 0.5 | 0.104 | 4.3 | 5.3 | 5.4 | 6.6 | 6.4 | 0 | 0.68604 | 0.17838 | 0.08272 | 1.32714 | 0.02309 | 0.19306 | 0.1228 |
| 440 | Urtica dioica L. | Urtica dioica | 0.15 | 0.5 | 0.074 | 7.9 | 5.4 | 2.9 | 7.9 | 2.4 | 0 | 0.48324 | 0.22836 | 0.33987 | 1.12718 | 0.05497 | 0.20766 | 0.12487 |
| 440 | Veronica hederifolia L. | Veronica hederifolia agg. | 0.1 | 0.5 | 0.270 | NA | 5.9 | 4.1 | NA | 2.9 | 0 | 0.61555 | 0.17954 | 0.13579 | 1.19563 | 0.03648 | 0.19058 | 0.11211 |
| 440 | Veronica arvensis L. | Veronica arvensis | 0.1 | 0.5 | 0.264 | 7 | 5.6 | 4.4 | 5.9 | 5.4 | 0.1 | 0.64871 | 0.63945 | 1.70759 | 2.05577 | 0.56937 | 0.21649 | 0.54803 |
| 445 | Trifolium repens L. | Trifolium repens | 0.1 | 0.62 | 0.364 | 7.4 | 5.3 | NA | 6 | 7.7 | 0.8 | 0.65201 | 0.63791 | 1.54898 | 2.03286 | 0.77966 | 0.37419 | 0.45721 |
| 445 | Poa annua L. | Poa annua | 0.21 | 0.62 | 0.125 | NA | NA | NA | NA | NA | NA | NA | NA | NA | NA | NA | NA | NA |
| 445 | Cerastium glomeratum Thuill. | Cerastium glomeratum | 0.05 | 0.62 | 0.154 | 7 | 5.6 | 4.4 | 5.9 | 5.4 | 0.1 | 0.64871 | 0.63945 | 1.70759 | 2.05577 | 0.56937 | 0.21649 | 0.54803 |
| 445 | Acer pseudoplatanus L. | Acer pseudoplatanus | 0.02 | 0.62 | 0.400 | 7.4 | 5.3 | NA | 6 | 7.7 | 0.8 | 0.65201 | 0.63791 | 1.54898 | 2.03286 | 0.77966 | 0.37419 | 0.45721 |
| 445 | Bromus sterilis L. | Bromus sterilis | 0.18 | 0.62 | 0.071 | 7.1 | 7.8 | 4.1 | 6.2 | 6 | 0.1 | 0.76637 | 0.65623 | 0.78647 | 1.82229 | 0.56568 | 0.18972 | 0.47454 |
| 445 | Taraxacum officinale F.H.Wigg. | Taraxacum officinale agg. | 0.03 | 0.62 | 0.154 | NA | NA | NA | NA | NA | NA | NA | NA | NA | NA | NA | NA | NA |
| 445 | Capsella bursa-pastoris Medik. | Capsella bursa-pastoris | 0.01 | 0.62 | 0.070 | 8.1 | NA | NA | 5.2 | NA | 0.1 | 0.51276 | 0.49581 | 1.51667 | 2.20989 | 0.98959 | 0.46997 | 0.3594 |
| 445 | Stellaria media (L.) Vill. | Stellaria media agg. | 0.02 | 0.62 | 0.182 | 7.3 | NA | NA | 5.8 | 5.9 | 0.5 | 0.63207 | 0.61738 | 1.52973 | 2.00883 | 0.70322 | 0.36696 | 0.39725 |
| 446 | Bromus sterilis L. | Bromus sterilis | 0.1 | 0.49 | 0.313 | 7.4 | 5.2 | 4.7 | 6.1 | NA | 0 | 0.77406 | 0.76726 | 1.79113 | 2.07279 | 0.64778 | 0.2119 | 0.65925 |
| 446 | Veronica arvensis L. | Veronica arvensis | 0.05 | 0.49 | 0.197 | NA | NA | NA | NA | NA | NA | NA | NA | NA | NA | NA | NA | NA |
| 446 | Stellaria media (L.) Vill. | Stellaria media agg. | 0.1 | 0.49 | 0.235 | 7.3 | 5.4 | 4.1 | 7.6 | 4.2 | 0.4 | 0.58728 | 0.56055 | 1.34601 | 1.94534 | 0.82357 | 0.23743 | 0.34603 |
| 446 | Geranium molle L. | Geranium molle | 0.02 | 0.49 | 0.219 | 7.3 | NA | 3.9 | 6.1 | 3.9 | 0.1 | 0.55903 | 0.41538 | 0.65189 | 1.65181 | 0.43719 | 0.24279 | 0.21709 |
| 446 | Taraxacum officinale F.H.Wigg. | Taraxacum officinale agg. | 0.05 | 0.49 | 0.072 | 7.3 | NA | 3.9 | 6.1 | 3.9 | 0.1 | 0.55903 | 0.41538 | 0.65189 | 1.65181 | 0.43719 | 0.24279 | 0.21709 |
| 446 | Acer pseudoplatanus L. | Acer pseudoplatanus | 0.05 | 0.49 | 0.043 | 7.9 | NA | 4.1 | 5.8 | NA | 0.9 | 0.54139 | 0.51128 | 1.31363 | 1.93633 | 0.8112 | 0.25085 | 0.32953 |
| 446 | Cerastium glomeratum Thuill. | Cerastium glomeratum | 0.05 | 0.49 | 0.603 | 7.1 | 5.1 | NA | NA | NA | 0.3 | 0.52904 | 0.48176 | 1.11642 | 1.87359 | 0.74801 | 0.282 | 0.30199 |
| 446 | Poa annua L. | Poa annua | 0.05 | 0.49 | 0.333 | NA | NA | NA | NA | NA | NA | NA | NA | NA | NA | NA | NA | NA |
| 446 | Medicago lupulina L. | Medicago lupulina | 0.01 | 0.49 | 0.138 | NA | NA | 4.5 | 6.6 | 6.6 | 0 | 0.78059 | 0.60963 | 0.54418 | 1.63701 | 0.26733 | 0.16232 | 0.54354 |
| 446 | NA | NA | 0.01 | 0.49 | 0.071 | 7.3 | NA | NA | 7.6 | 7.7 | 0.4 | 0.75479 | 0.72524 | 1.28692 | 1.77374 | 0.61324 | 0.18283 | 0.56773 |
| 447 | Geranium molle L. | Geranium molle | 0.01 | 0.79 | 0.250 | NA | NA | NA | NA | NA | NA | NA | NA | NA | NA | NA | NA | NA |
| 447 | Taraxacum officinale F.H.Wigg. | Taraxacum officinale agg. | 0.01 | 0.79 | 0.116 | NA | NA | 4.5 | 6.6 | 6.6 | 0 | 0.78059 | 0.60963 | 0.54418 | 1.63701 | 0.26733 | 0.16232 | 0.54354 |
| 447 | Bromus sterilis L. | Bromus sterilis | 0.2 | 0.79 | 0.159 | 6.5 | NA | NA | 5.4 | NA | 0 | 0.59769 | 0.45571 | 0.64594 | 1.51584 | 0.28825 | 0.22901 | 0.32316 |
| 447 | Stellaria media (L.) Vill. | Stellaria media agg. | 0.01 | 0.79 | 0.200 | NA | NA | 4.5 | 6.6 | 6.6 | 0 | 0.78059 | 0.60963 | 0.54418 | 1.63701 | 0.26733 | 0.16232 | 0.54354 |
| 447 | Acer pseudoplatanus L. | Acer pseudoplatanus | 0.5 | 0.79 | 0.517 | 6.7 | NA | NA | 6.5 | 7.7 | 0.1 | 0.70945 | 0.61373 | 0.80583 | 1.7717 | 0.43958 | 0.26577 | 0.51423 |
| 447 | Poa annua L. | Poa annua | 0.05 | 0.79 | 0.125 | NA | NA | NA | NA | NA | NA | NA | NA | NA | NA | NA | NA | NA |
| 447 | Capsella bursa-pastoris Medik. | Capsella bursa-pastoris | 0.01 | 0.79 | 0.089 | 8.1 | NA | 4.1 | 6.6 | NA | 0.7 | 0.57545 | 0.53032 | 1.14497 | 1.72102 | 0.70077 | 0.21269 | 0.32208 |
| 448 | Taraxacum officinale F.H.Wigg. | Taraxacum officinale agg. | 0.02 | 0.65 | 0.368 | 7.4 | 5.3 | NA | 6 | 7.7 | 0.8 | 0.65201 | 0.63791 | 1.54898 | 2.03286 | 0.77966 | 0.37419 | 0.45721 |
| 448 | Cardamine hirsuta L. | Cardamine hirsuta | 0.01 | 0.65 | 0.068 | 8.4 | NA | 4 | 6.3 | NA | 0.1 | 0.76525 | 0.73026 | 1.22636 | 1.78874 | 0.7034 | 0.16135 | 0.51807 |
| 448 | Poa annua L. | Poa annua | 0.1 | 0.65 | 0.179 | NA | NA | NA | NA | NA | NA | NA | NA | NA | NA | NA | NA | NA |
| 448 | Bromus sterilis L. | Bromus sterilis | 0.2 | 0.65 | 0.134 | 7.3 | 5.4 | 4.1 | 7.6 | 4.2 | 0.4 | 0.58728 | 0.56055 | 1.34601 | 1.94534 | 0.82357 | 0.23743 | 0.34603 |
| 448 | Sisymbrium officinale (L.) Scop. | Sisymbrium officinale | 0.02 | 0.65 | 0.133 | 7.1 | 5.3 | 4.5 | 5.9 | 4.8 | 0.3 | 0.52518 | 0.5113 | 1.61045 | 2.05509 | 0.96495 | 0.27897 | 0.32911 |
| 448 | Galium aparine L. | Galium aparine aparine | 0.15 | 0.65 | 0.167 | 7.8 | NA | 4.6 | 6.2 | 5.8 | 0.1 | 0.71321 | 0.68276 | 1.29546 | 1.8275 | 0.62188 | 0.18605 | 0.48742 |
| 448 | Cerastium glomeratum Thuill. | Cerastium glomeratum | 0.02 | 0.65 | 0.077 | NA | NA | NA | NA | NA | NA | NA | NA | NA | NA | NA | NA | NA |
| 448 | Stellaria media (L.) Vill. | Stellaria media agg. | 0.08 | 0.65 | 0.123 | NA | NA | NA | NA | NA | NA | NA | NA | NA | NA | NA | NA | NA |
| 448 | Acer campestre L. | Acer campestre | 0.02 | 0.65 | 0.123 | 6.8 | NA | 5.6 | 5.8 | NA | 0.1 | 0.52169 | 0.45097 | 0.93044 | 1.87015 | 0.67593 | 0.2975 | 0.27054 |
| 448 | Acer pseudoplatanus L. | Acer pseudoplatanus | 0.02 | 0.65 | 0.067 | 7.3 | NA | 3.9 | 6.1 | 3.9 | 0.1 | 0.55903 | 0.41538 | 0.65189 | 1.65181 | 0.43719 | 0.24279 | 0.21709 |
| 448 | NA | NA | 0.01 | 0.65 | 0.101 | 8.1 | NA | NA | 5.2 | NA | 0.1 | 0.51276 | 0.49581 | 1.51667 | 2.20989 | 0.98959 | 0.46997 | 0.3594 |
| 449 | Taraxacum officinale F.H.Wigg. | Taraxacum officinale agg. | 0.02 | 0.6 | 0.176 | 7.4 | 5.2 | 4.7 | 6.1 | NA | 0 | 0.77406 | 0.76726 | 1.79113 | 2.07279 | 0.64778 | 0.2119 | 0.65925 |
| 449 | Stellaria media (L.) Vill. | Stellaria media agg. | 0.31 | 0.6 | 0.147 | 7 | 5.6 | 4.4 | 5.9 | 5.4 | 0.1 | 0.64871 | 0.63945 | 1.70759 | 2.05577 | 0.56937 | 0.21649 | 0.54803 |
| 449 | Achillea millefolium L. | Achillea millefolium millefolium | 0.02 | 0.6 | 0.205 | 8.6 | 8 | 3.3 | 3 | 1.8 | 0.2 | 0.49702 | 0.46825 | 1.42549 | 1.99857 | 0.4756 | 0.32016 | 0.338 |
| 449 | Acer pseudoplatanus L. | Acer pseudoplatanus | 0.11 | 0.6 | 0.086 | 8.4 | NA | 3 | 6 | NA | 0 | 0.46249 | 0.43915 | 1.39189 | 1.83687 | 0.38437 | 0.30571 | 0.38883 |
| 449 | Silene latifolia Poir. | NA | 0.11 | 0.6 | 0.333 | 4.7 | NA | NA | 5.8 | 6.6 | 0 | 0.68705 | 0.16985 | 0.0706 | 1.32057 | 0.02196 | 0.18337 | 0.11293 |
| 449 | Acer campestre L. | Acer campestre | 0.01 | 0.6 | 0.143 | 6.5 | NA | NA | 5.4 | NA | 0 | 0.59769 | 0.45571 | 0.64594 | 1.51584 | 0.28825 | 0.22901 | 0.32316 |
| 449 | Cardamine hirsuta L. | Cardamine hirsuta | 0.02 | 0.6 | 0.250 | 7.1 | 4.9 | NA | 6 | NA | 1 | 0.59517 | 0.59064 | 1.89772 | 2.12267 | 0.65091 | 0.41213 | 0.37185 |
| 450 | Taraxacum officinale F.H.Wigg. | Taraxacum officinale agg. | 0.05 | 0.4 | 0.351 | 6.7 | NA | NA | 6.5 | 7.7 | 0.1 | 0.70945 | 0.61373 | 0.80583 | 1.7717 | 0.43958 | 0.26577 | 0.51423 |
| 450 | Stellaria media (L.) Vill. | Stellaria media agg. | 0.15 | 0.4 | 0.288 | 6.7 | NA | NA | 6.5 | 7.7 | 0.1 | 0.70945 | 0.61373 | 0.80583 | 1.7717 | 0.43958 | 0.26577 | 0.51423 |
| 450 | Erodium cicutarium (L.) | Erodium cicutarium cicutarium | 0.02 | 0.4 | 0.220 | 7.4 | 5.3 | NA | 6 | 7.7 | 0.8 | 0.65201 | 0.63791 | 1.54898 | 2.03286 | 0.77966 | 0.37419 | 0.45721 |
| 450 | Silene latifolia Poir. | NA | 0.04 | 0.4 | 0.152 | 7.9 | 5.4 | 2.9 | 7.9 | 2.4 | 0 | 0.48324 | 0.22836 | 0.33987 | 1.12718 | 0.05497 | 0.20766 | 0.12487 |
| 450 | Acer pseudoplatanus L. | Acer pseudoplatanus | 0.13 | 0.4 | 0.368 | 7.4 | 5.3 | NA | 6 | 7.7 | 0.8 | 0.65201 | 0.63791 | 1.54898 | 2.03286 | 0.77966 | 0.37419 | 0.45721 |
| 450 | Acer campestre L. | Acer campestre | 0.01 | 0.4 | 0.260 | 7 | 5.6 | 4.4 | 5.9 | 5.4 | 0.1 | 0.64871 | 0.63945 | 1.70759 | 2.05577 | 0.56937 | 0.21649 | 0.54803 |
| 451 | Acer pseudoplatanus L. | Acer pseudoplatanus | 0.1 | 0.56 | 0.095 | 7.9 | 5.4 | 2.9 | 7.9 | 2.4 | 0 | 0.48324 | 0.22836 | 0.33987 | 1.12718 | 0.05497 | 0.20766 | 0.12487 |
| 451 | Daucus carota L. | Daucus carota | 0.05 | 0.56 | 0.115 | 8.1 | NA | NA | 4.4 | 2.8 | 0.3 | 0.48868 | 0.46508 | 1.4426 | 2.04965 | 0.7278 | 0.33931 | 0.35628 |
| 451 | Sagina procumbens L. | Sagina procumbens | 0.09 | 0.56 | 0.182 | 6.7 | NA | NA | 6.5 | 7.7 | 0.1 | 0.70945 | 0.61373 | 0.80583 | 1.7717 | 0.43958 | 0.26577 | 0.51423 |
| 451 | Erigeron canadensis L. | Conyza canadensis | 0.05 | 0.56 | 0.308 | 7.8 | NA | 4.6 | 6.2 | 5.8 | 0.1 | 0.71321 | 0.68276 | 1.29546 | 1.8275 | 0.62188 | 0.18605 | 0.48742 |
| 451 | Poa annua L. | Poa annua | 0.08 | 0.56 | 0.200 | NA | NA | NA | NA | NA | NA | NA | NA | NA | NA | NA | NA | NA |
| 451 | Taraxacum officinale F.H.Wigg. | Taraxacum officinale agg. | 0.13 | 0.56 | 0.214 | 7.4 | 5.2 | 4.7 | 6.1 | NA | 0 | 0.77406 | 0.76726 | 1.79113 | 2.07279 | 0.64778 | 0.2119 | 0.65925 |
| 451 | Hypericum perforatum L. | Hypericum perforatum | 0.05 | 0.56 | 0.183 | NA | NA | 5.5 | 6.2 | 6.8 | 0 | 0.67445 | 0.21586 | 0.12312 | 1.32299 | 0.04699 | 0.18459 | 0.11909 |
| 451 | Stellaria media (L.) Vill. | Stellaria media agg. | 0.01 | 0.56 | 0.091 | 7.4 | 5.2 | 4.7 | 6.1 | NA | 0 | 0.77406 | 0.76726 | 1.79113 | 2.07279 | 0.64778 | 0.2119 | 0.65925 |
| 452 | Taraxacum officinale F.H.Wigg. | Taraxacum officinale agg. | 0.15 | 0.68 | 0.105 | 7.8 | 5.1 | NA | NA | NA | 0.2 | 0.51255 | 0.49092 | 1.42792 | 1.96794 | 1.00937 | 0.22916 | 0.30884 |
| 452 | Acer platanoides L. | Acer platanoides | 0.05 | 0.68 | 0.403 | NA | NA | 5.4 | 6.4 | 5.6 | 0 | 0.68187 | 0.18824 | 0.08946 | 1.3432 | 0.02985 | 0.19867 | 0.10919 |
| 452 | Poa annua L. | Poa annua | 0.25 | 0.68 | 0.143 | 7.2 | NA | NA | 6.6 | 6.8 | 0 | 0.81299 | 0.76483 | 1.09209 | 1.84162 | 0.41616 | 0.14633 | 0.71443 |
| 452 | Erigeron canadensis L. | Conyza canadensis | 0.07 | 0.68 | 0.091 | NA | NA | NA | NA | NA | NA | NA | NA | NA | NA | NA | NA | NA |
| 452 | Stellaria media (L.) Vill. | Stellaria media agg. | 0.08 | 0.68 | 0.063 | 7.9 | NA | 4.1 | 5.8 | NA | 0.9 | 0.54139 | 0.51128 | 1.31363 | 1.93633 | 0.8112 | 0.25085 | 0.32953 |
| 452 | Sagina procumbens L. | Sagina procumbens | 0.08 | 0.68 | 0.128 | 7.3 | NA | 3.9 | 6.1 | 3.9 | 0.1 | 0.55903 | 0.41538 | 0.65189 | 1.65181 | 0.43719 | 0.24279 | 0.21709 |
| 453 | Taraxacum officinale F.H.Wigg. | Taraxacum officinale agg. | 0.15 | 0.73 | 0.139 | 7.3 | NA | 3.9 | 6.1 | 3.9 | 0.1 | 0.55903 | 0.41538 | 0.65189 | 1.65181 | 0.43719 | 0.24279 | 0.21709 |
| 453 | Daucus carota L. | Daucus carota | 0.3 | 0.73 | 0.079 | 7.9 | NA | 4.1 | 5.8 | NA | 0.9 | 0.54139 | 0.51128 | 1.31363 | 1.93633 | 0.8112 | 0.25085 | 0.32953 |
| 453 | Lactuca serriola L. | Lactuca serriola (scariola) | 0.05 | 0.73 | 0.172 | NA | NA | NA | NA | NA | NA | NA | NA | NA | NA | NA | NA | NA |
| 453 | Erigeron canadensis L. | Conyza canadensis | 0.1 | 0.73 | 0.071 | 7 | 5.6 | 4.4 | 5.9 | 5.4 | 0.1 | 0.64871 | 0.63945 | 1.70759 | 2.05577 | 0.56937 | 0.21649 | 0.54803 |
| 453 | Sagina procumbens L. | Sagina procumbens | 0.05 | 0.73 | 0.083 | NA | NA | 5.3 | 6.2 | NA | 0.1 | 0.68682 | 0.14816 | 0.05813 | 1.31254 | 0.01433 | 0.20126 | 0.10505 |
| 453 | Poa annua L. | Poa annua | 0.08 | 0.73 | 0.116 | NA | NA | 5.3 | 6.2 | NA | 0.1 | 0.68682 | 0.14816 | 0.05813 | 1.31254 | 0.01433 | 0.20126 | 0.10505 |
| 454 | Hypericum perforatum L. | Hypericum perforatum | 0.15 | 0.84 | 0.079 | NA | NA | 5.3 | 6.2 | NA | 0.1 | 0.68682 | 0.14816 | 0.05813 | 1.31254 | 0.01433 | 0.20126 | 0.10505 |
| 454 | Clematis vitalba L. | Clematis vitalba | 0.3 | 0.84 | 0.089 | NA | NA | 4.5 | 6.6 | 6.6 | 0 | 0.78059 | 0.60963 | 0.54418 | 1.63701 | 0.26733 | 0.16232 | 0.54354 |
| 454 | Taraxacum officinale F.H.Wigg. | Taraxacum officinale agg. | 0.15 | 0.84 | 0.128 | NA | NA | 4.5 | 6.6 | 6.6 | 0 | 0.78059 | 0.60963 | 0.54418 | 1.63701 | 0.26733 | 0.16232 | 0.54354 |
| 454 | Erigeron canadensis L. | Conyza canadensis | 0.08 | 0.84 | 0.200 | 7 | 5.6 | 4.4 | 5.9 | 5.4 | 0.1 | 0.64871 | 0.63945 | 1.70759 | 2.05577 | 0.56937 | 0.21649 | 0.54803 |
| 454 | Sagina procumbens L. | Sagina procumbens | 0.09 | 0.84 | 0.033 | 7.9 | NA | 4.1 | 5.8 | NA | 0.9 | 0.54139 | 0.51128 | 1.31363 | 1.93633 | 0.8112 | 0.25085 | 0.32953 |
| 454 | Veronica arvensis L. | Veronica arvensis | 0.05 | 0.84 | 0.375 | 6.7 | NA | NA | 6.5 | 7.7 | 0.1 | 0.70945 | 0.61373 | 0.80583 | 1.7717 | 0.43958 | 0.26577 | 0.51423 |
| 454 | Poa annua L. | Poa annua | 0.02 | 0.84 | 0.103 | 7.8 | NA | 4.6 | 6.2 | 5.8 | 0.1 | 0.71321 | 0.68276 | 1.29546 | 1.8275 | 0.62188 | 0.18605 | 0.48742 |
| 455 | Taraxacum officinale F.H.Wigg. | Taraxacum officinale agg. | 0.12 | 0.82 | 0.137 | 7.8 | NA | 4.6 | 6.2 | 5.8 | 0.1 | 0.71321 | 0.68276 | 1.29546 | 1.8275 | 0.62188 | 0.18605 | 0.48742 |
| 455 | Hypericum perforatum L. | Hypericum perforatum | 0.25 | 0.82 | 0.095 | 7.8 | NA | 4.6 | 6.2 | 5.8 | 0.1 | 0.71321 | 0.68276 | 1.29546 | 1.8275 | 0.62188 | 0.18605 | 0.48742 |
| 455 | Medicago lupulina L. | Medicago lupulina | 0.11 | 0.82 | 0.122 | 8.6 | NA | 2.7 | 4.4 | 1.6 | 0 | 0.53557 | 0.50485 | 1.29532 | 1.90211 | 0.72298 | 0.28495 | 0.2778 |
| 455 | Potentilla argentea L. | Potentilla argentea agg. | 0.1 | 0.82 | 0.198 | 4.7 | 5.8 | 5.6 | 6.5 | 6.3 | 0 | 0.69843 | 0.15019 | 0.04615 | 1.3308 | 0.01233 | 0.18685 | 0.11235 |
| 455 | Carpinus betulus L. | Carpinus betulus | 0.15 | 0.82 | 0.067 | 7.9 | NA | 4.1 | 5.8 | NA | 0.9 | 0.54139 | 0.51128 | 1.31363 | 1.93633 | 0.8112 | 0.25085 | 0.32953 |
| 455 | Sagina procumbens L. | Sagina procumbens | 0.05 | 0.82 | 0.055 | 6.7 | 5.3 | 4.5 | 5.3 | 5.8 | 0 | 0.75464 | 0.73051 | 1.36974 | 1.95508 | 0.45974 | 0.15567 | 0.6659 |
| 455 | Veronica arvensis L. | Veronica arvensis | 0.02 | 0.82 | 0.037 | 6.6 | 5.6 | NA | 5.6 | 4.1 | 0.1 | 0.50969 | 0.50576 | 1.97957 | 2.23767 | 1.1694 | 0.28385 | 0.36995 |
| 455 | Poa annua L. | Poa annua | 0.02 | 0.82 | 0.154 | 8.4 | NA | 3 | 6 | NA | 0 | 0.46249 | 0.43915 | 1.39189 | 1.83687 | 0.38437 | 0.30571 | 0.38883 |
| 456 | Medicago lupulina L. | Medicago lupulina | 0.45 | 1.01 | 0.067 | 6.6 | 5.6 | NA | 5.6 | 4.1 | 0.1 | 0.50969 | 0.50576 | 1.97957 | 2.23767 | 1.1694 | 0.28385 | 0.36995 |
| 456 | Geranium molle L. | Geranium molle | 0.15 | 1.01 | 0.127 | 6.6 | 5.6 | NA | 5.6 | 4.1 | 0.1 | 0.50969 | 0.50576 | 1.97957 | 2.23767 | 1.1694 | 0.28385 | 0.36995 |
| 456 | Acer platanoides L. | Acer platanoides | 0.2 | 1.01 | 0.118 | 6.6 | 5.6 | NA | 5.6 | 4.1 | 0.1 | 0.50969 | 0.50576 | 1.97957 | 2.23767 | 1.1694 | 0.28385 | 0.36995 |
| 456 | Veronica arvensis L. | Veronica arvensis | 0.08 | 1.01 | 0.044 | 8.3 | 5.2 | 3.5 | 6.7 | 4 | 0.1 | 0.59253 | 0.57367 | 1.48069 | 1.91606 | 0.44486 | 0.27236 | 0.42509 |
| 456 | Sagina procumbens L. | Sagina procumbens | 0.08 | 1.01 | 0.068 | 7.4 | 5.3 | NA | 6 | 7.7 | 0.8 | 0.65201 | 0.63791 | 1.54898 | 2.03286 | 0.77966 | 0.37419 | 0.45721 |
| 456 | Erigeron canadensis L. | Conyza canadensis | 0.05 | 1.01 | 0.083 | 8.4 | NA | 4 | 6.3 | NA | 0.1 | 0.76525 | 0.73026 | 1.22636 | 1.78874 | 0.7034 | 0.16135 | 0.51807 |
| 457 | Clematis vitalba L. | Clematis vitalba | 0.2 | 0.75 | 0.152 | 7.1 | 4.9 | NA | 6 | NA | 1 | 0.59517 | 0.59064 | 1.89772 | 2.12267 | 0.65091 | 0.41213 | 0.37185 |
| 457 | Taraxacum officinale F.H.Wigg. | Taraxacum officinale agg. | 0.08 | 0.75 | 0.192 | 7.4 | 5.3 | NA | 6 | 7.7 | 0.8 | 0.65201 | 0.63791 | 1.54898 | 2.03286 | 0.77966 | 0.37419 | 0.45721 |
| 457 | Plantago lanceolata L. | Plantago lanceolata | 0.1 | 0.75 | 0.136 | 7.1 | 4.9 | NA | 6 | NA | 1 | 0.59517 | 0.59064 | 1.89772 | 2.12267 | 0.65091 | 0.41213 | 0.37185 |
| 457 | Achillea millefolium L. | Achillea millefolium millefolium | 0.05 | 0.75 | 0.233 | 6.5 | NA | NA | 5.4 | NA | 0 | 0.59769 | 0.45571 | 0.64594 | 1.51584 | 0.28825 | 0.22901 | 0.32316 |
| 457 | natherum elatius (L.) P.Beauv. ex J.Presl & C | Arrhenatherum elatius | 0.08 | 0.75 | 0.076 | 7.1 | 4.9 | NA | 6 | NA | 1 | 0.59517 | 0.59064 | 1.89772 | 2.12267 | 0.65091 | 0.41213 | 0.37185 |

| | | | | | | | | | | | | | | | | | | |
|---|---|---|---|---|---|---|---|---|---|---|---|---|---|---|---|---|---|---|
| 457 | Medicago lupulina L. | Medicago lupulina | 0.19 | 0.75 | 0.221 | 7.4 | 5.3 | NA | 6 | 7.7 | 0.8 | 0.65201 | 0.63791 | 1.54898 | 2.03286 | 0.77966 | 0.37419 | 0.45721 |
| 457 | Acer platanoides L. | Acer platanoides | 0.05 | 0.75 | 0.118 | 7 | 5.6 | 4.4 | 5.9 | 5.4 | 0.1 | 0.64871 | 0.63945 | 1.70759 | 2.05577 | 0.56937 | 0.21649 | 0.54803 |
| 458 | Poa annua L. | Poa annua | 0.25 | 0.6 | 0.200 | 7.4 | 5.3 | NA | 6 | 7.7 | 0.8 | 0.65201 | 0.63791 | 1.54898 | 2.03286 | 0.77966 | 0.37419 | 0.45721 |
| 458 | Sagina procumbens L. | Sagina procumbens | 0.25 | 0.6 | 0.032 | NA | NA | 5.5 | 6.2 | 6.8 | 0 | 0.67445 | 0.21586 | 0.12312 | 1.32299 | 0.04699 | 0.18459 | 0.11909 |
| 458 | Erigeron canadensis L. | Conyza canadensis | 0.1 | 0.6 | 0.207 | 7.4 | 5.3 | NA | 6 | 7.7 | 0.8 | 0.65201 | 0.63791 | 1.54898 | 2.03286 | 0.77966 | 0.37419 | 0.45721 |
| 459 | Bellis perennis L. | Bellis perennis | 0.05 | 0.91 | 0.182 | 7 | 5.6 | 4.4 | 5.9 | 5.4 | 0.1 | 0.64871 | 0.63945 | 1.70759 | 2.05577 | 0.56937 | 0.21649 | 0.54803 |
| 459 | Geranium molle L. | Geranium molle | 0.09 | 0.91 | 0.395 | NA | NA | NA | NA | NA | NA | NA | NA | NA | NA | NA | NA | NA |
| 459 | Hypochaeris radicata L. | Hypochoeris radicata | 0.09 | 0.91 | 0.143 | 7.1 | 4.9 | NA | 6 | NA | 1 | 0.59517 | 0.59064 | 1.89772 | 2.12267 | 0.65091 | 0.41213 | 0.37185 |
| 459 | Taraxacum officinale F.H.Wigg. | Taraxacum officinale agg. | 0.07 | 0.91 | 0.141 | NA | NA | NA | NA | NA | NA | NA | NA | NA | NA | NA | NA | NA |
| 459 | Myosotis arvensis Hill | Myosotis arvensis (intermedia) | 0.05 | 0.91 | 0.254 | 7.4 | 5.3 | NA | 6 | 7.7 | 0.8 | 0.65201 | 0.63791 | 1.54898 | 2.03286 | 0.77966 | 0.37419 | 0.45721 |
| 459 | Prunella vulgaris L. | Prunella vulgaris | 0.08 | 0.91 | 0.081 | NA | NA | 5.3 | 6.2 | NA | 0.1 | 0.68682 | 0.14816 | 0.05813 | 1.31254 | 0.01433 | 0.20126 | 0.10505 |
| 459 | Trifolium dubium Sibth. | Trifolium dubium (minus) | 0.05 | 0.91 | 0.217 | 7.1 | 5.3 | 4.5 | 5.9 | 4.8 | 0.3 | 0.52518 | 0.5113 | 1.61045 | 2.05509 | 0.96495 | 0.27897 | 0.32911 |
| 459 | Medicago lupulina L. | Medicago lupulina | 0.08 | 0.91 | 0.079 | 7.3 | 5.4 | 4.1 | 7.6 | 4.2 | 0.4 | 0.58728 | 0.56055 | 1.34601 | 1.94534 | 0.82357 | 0.23743 | 0.34603 |
| 459 | Oxalis corniculata L. | Oxalis corniculata | 0.05 | 0.91 | 0.086 | 7.4 | 5.3 | NA | 6 | 7.7 | 0.8 | 0.65201 | 0.63791 | 1.54898 | 2.03286 | 0.77966 | 0.37419 | 0.45721 |
| 459 | Poa annua L. | Poa annua | 0.05 | 0.91 | 0.071 | 6.7 | NA | NA | 6.5 | 7.7 | 0.1 | 0.70945 | 0.61373 | 0.80583 | 1.7717 | 0.43958 | 0.26577 | 0.51423 |
| 459 | Sedum sexangulare L. | Sedum sexangulare | 0.05 | 0.91 | 0.209 | 6.5 | NA | NA | 5.4 | NA | 0 | 0.59769 | 0.45571 | 0.64594 | 1.51584 | 0.28825 | 0.22901 | 0.32316 |
| 459 | Sagina procumbens L. | Sagina procumbens | 0.09 | 0.91 | 0.161 | NA | NA | 5.3 | 6.2 | NA | 0.1 | 0.68682 | 0.14816 | 0.05813 | 1.31254 | 0.01433 | 0.20126 | 0.10505 |
| 459 | Arenaria serpyllifolia L. | Arenaria serpyllifolia agg. | 0.01 | 0.91 | 0.081 | 7.3 | 5.8 | 4.8 | 5.3 | 5.6 | 0 | 0.67083 | 0.63295 | 1.24583 | 1.99939 | 0.56795 | 0.28417 | 0.48569 |
| 459 | NA | NA | 0.1 | 0.91 | 0.204 | NA | NA | NA | NA | NA | NA | NA | NA | NA | NA | NA | NA | NA |
| 460 | Geranium molle L. | Geranium molle | 0.1 | 0.81 | 0.161 | 7.1 | 4.9 | NA | 6 | NA | 1 | 0.59517 | 0.59064 | 1.89772 | 2.12267 | 0.65091 | 0.41213 | 0.37185 |
| 460 | Vicia sativa L. | Vicia sativa agg. | 0.04 | 0.81 | 0.118 | 6.7 | NA | NA | 6.5 | 7.7 | 0.1 | 0.70945 | 0.61373 | 0.80583 | 1.7717 | 0.43958 | 0.26577 | 0.51423 |
| 460 | Taraxacum officinale F.H.Wigg. | Taraxacum officinale agg. | 0.1 | 0.81 | 0.068 | 7.1 | 4.9 | NA | 6 | NA | 1 | 0.59517 | 0.59064 | 1.89772 | 2.12267 | 0.65091 | 0.41213 | 0.37185 |
| 460 | Trifolium dubium Sibth. | Trifolium dubium (minus) | 0.03 | 0.81 | 0.107 | 7.1 | 4.9 | NA | 6 | NA | 1 | 0.59517 | 0.59064 | 1.89772 | 2.12267 | 0.65091 | 0.41213 | 0.37185 |
| 460 | Oxalis corniculata L. | Oxalis corniculata | 0.02 | 0.81 | 0.183 | 4.4 | 5.9 | 5.4 | 5.5 | 5.3 | 0 | 0.695 | 0.13806 | 0.03343 | 1.32501 | 0.01288 | 0.19547 | 0.10889 |
| 460 | Bellis perennis L. | Bellis perennis | 0.08 | 0.81 | 0.079 | 7 | 5.6 | 4.4 | 5.9 | 5.4 | 0.1 | 0.64871 | 0.63945 | 1.70759 | 2.05577 | 0.56937 | 0.21649 | 0.54803 |
| 460 | Poa annua L. | Poa annua | 0.08 | 0.81 | 0.107 | 7.8 | 5.3 | NA | 6.6 | NA | 0.1 | 0.52944 | 0.48335 | 1.1288 | 1.79222 | 0.82465 | 0.16914 | 0.30679 |
| 460 | Myosotis arvensis Hill | Myosotis arvensis (intermedia) | 0.02 | 0.81 | 0.088 | 6.8 | NA | 5.6 | 5.8 | NA | 0.1 | 0.52169 | 0.45097 | 0.93044 | 1.87015 | 0.67593 | 0.2975 | 0.27054 |
| 460 | Cerastium semidecandrum L. | Cerastium semidecandrum | 0.03 | 0.81 | 0.025 | 7.1 | 7.8 | 4.1 | 6.2 | 6 | 0.1 | 0.76637 | 0.65623 | 0.78647 | 1.82229 | 0.56568 | 0.18972 | 0.47454 |
| 460 | Sedum sexangulare L. | Sedum sexangulare | 0.05 | 0.81 | 0.031 | 7.3 | 6.1 | 3.8 | 6 | 5.1 | 0 | 0.62987 | 0.59281 | 1.27995 | 1.90491 | 0.6275 | 0.3036 | 0.42608 |
| 460 | Hypochaeris radicata L. | Hypochoeris radicata | 0.1 | 0.81 | 0.200 | 7.3 | 5.4 | 4.1 | 7.6 | 4.2 | 0.4 | 0.58728 | 0.56055 | 1.34601 | 1.94534 | 0.82357 | 0.23743 | 0.34603 |
| 460 | Sagina procumbens L. | Sagina procumbens | 0.09 | 0.81 | 0.190 | 7.7 | NA | NA | 5.6 | NA | 0.7 | 0.53223 | 0.52325 | 1.73719 | 2.14421 | 0.94278 | 0.39374 | 0.35054 |
| 460 | Prunella vulgaris L. | Prunella vulgaris | 0.05 | 0.81 | 0.118 | NA | NA | NA | NA | NA | NA | NA | NA | NA | NA | NA | NA | NA |
| 460 | NA | NA | 0.02 | 0.81 | 0.221 | NA | NA | NA | NA | NA | NA | NA | NA | NA | NA | NA | NA | NA |
| 461 | Taraxacum officinale F.H.Wigg. | Taraxacum officinale agg. | 0.22 | 0.65 | 0.137 | 7.1 | 5.3 | 4.5 | 5.9 | 4.8 | 0.3 | 0.52518 | 0.5113 | 1.61045 | 2.05509 | 0.96495 | 0.27897 | 0.32911 |
| 461 | Sagina procumbens L. | Sagina procumbens | 0.12 | 0.65 | 0.172 | 7 | 5.6 | 4.4 | 5.9 | 5.4 | 0.1 | 0.64871 | 0.63945 | 1.70759 | 2.05577 | 0.56937 | 0.21649 | 0.54803 |
| 461 | Prunella vulgaris L. | Prunella vulgaris | 0.08 | 0.65 | 0.405 | 7.4 | 5.3 | NA | 6 | 7.7 | 0.8 | 0.65201 | 0.63791 | 1.54898 | 2.03286 | 0.77966 | 0.37419 | 0.45721 |
| 461 | Cerastium semidecandrum L. | Cerastium semidecandrum | 0.1 | 0.65 | 0.273 | NA | NA | NA | NA | NA | NA | NA | NA | NA | NA | NA | NA | NA |
| 461 | Geranium molle L. | Geranium molle | 0.02 | 0.65 | 0.085 | NA | NA | 5.3 | 6.2 | NA | 0.1 | 0.68682 | 0.14816 | 0.05813 | 1.31254 | 0.01433 | 0.20126 | 0.10505 |
| 461 | Oxalis corniculata L. | Oxalis corniculata | 0.02 | 0.65 | 0.121 | 7.3 | NA | NA | 5.8 | 5.9 | 0.5 | 0.63207 | 0.61738 | 1.52973 | 2.00883 | 0.70322 | 0.36696 | 0.39725 |
| 461 | Hypochaeris glabra L. | Hypochaeris glabra | 0.05 | 0.65 | 0.130 | 7 | 5.6 | 4.4 | 5.9 | 5.4 | 0.1 | 0.64871 | 0.63945 | 1.70759 | 2.05577 | 0.56937 | 0.21649 | 0.54803 |
| 461 | Arenaria serpyllifolia L. | Arenaria serpyllifolia agg. | 0.02 | 0.65 | 0.074 | 6.3 | NA | 5.9 | 6.3 | 6.9 | 0.1 | 0.6037 | 0.3946 | 0.44368 | 1.50139 | 0.30164 | 0.21315 | 0.21494 |
| 461 | Trifolium badium subsp. badium | Trifolium aureum | 0.02 | 0.65 | 0.040 | 7.1 | 4.9 | NA | 6 | NA | 1 | 0.59517 | 0.59064 | 1.89772 | 2.12267 | 0.65091 | 0.41213 | 0.37185 |
| 462 | Poa annua L. | Poa annua | 0.05 | 0.75 | 0.095 | 7.1 | 4.9 | NA | 6 | NA | 1 | 0.59517 | 0.59064 | 1.89772 | 2.12267 | 0.65091 | 0.41213 | 0.37185 |
| 462 | Taraxacum officinale F.H.Wigg. | Taraxacum officinale agg. | 0.09 | 0.75 | 0.092 | 7.8 | NA | 4.6 | 6.2 | 5.8 | 0.1 | 0.71321 | 0.68276 | 1.29546 | 1.8275 | 0.62188 | 0.18605 | 0.48742 |
| 462 | Hypericum perforatum L. | Hypericum perforatum | 0.05 | 0.75 | 0.211 | 7.1 | 4.9 | NA | 6 | NA | 1 | 0.59517 | 0.59064 | 1.89772 | 2.12267 | 0.65091 | 0.41213 | 0.37185 |
| 462 | Trifolium dubium Sibth. | Trifolium dubium (minus) | 0.05 | 0.75 | 0.143 | 7.4 | 5.3 | NA | 6 | 7.7 | 0.8 | 0.65201 | 0.63791 | 1.54898 | 2.03286 | 0.77966 | 0.37419 | 0.45721 |
| 462 | Medicago lupulina L. | Medicago lupulina | 0.15 | 0.75 | 0.273 | 7.4 | 5.3 | NA | 6 | 7.7 | 0.8 | 0.65201 | 0.63791 | 1.54898 | 2.03286 | 0.77966 | 0.37419 | 0.45721 |
| 462 | Trifolium repens L. | Trifolium repens | 0.04 | 0.75 | 0.317 | NA | NA | NA | NA | NA | NA | NA | NA | NA | NA | NA | NA | NA |
| 462 | Cerastium semidecandrum L. | Cerastium semidecandrum | 0.08 | 0.75 | 0.026 | 8.3 | 5.2 | 3.5 | 6.7 | 4 | 0.1 | 0.59253 | 0.57367 | 1.48069 | 1.91606 | 0.44486 | 0.27236 | 0.42509 |
| 462 | Hypochaeris radicata L. | Hypochoeris radicata | 0.08 | 0.75 | 0.196 | 7 | NA | 3.6 | 7.1 | 3.1 | 0 | 0.56346 | 0.34896 | 0.48122 | 1.51076 | 0.328 | 0.2423 | 0.13797 |
| 462 | Sedum sexangulare L. | Sedum sexangulare | 0.02 | 0.75 | 0.156 | 7.1 | 5.3 | 4.5 | 5.9 | 4.8 | 0.3 | 0.52518 | 0.5113 | 1.61045 | 2.05509 | 0.96495 | 0.27897 | 0.32911 |
| 462 | Vicia sativa L. | Vicia sativa agg. | 0.1 | 0.75 | 0.106 | 7.3 | 5.4 | 4.1 | 7.6 | 4.2 | 0.4 | 0.58728 | 0.56055 | 1.34601 | 1.94534 | 0.82357 | 0.23743 | 0.34603 |
| 462 | Senecio vulgaris L. | Senecio vulgaris | 0.01 | 0.75 | 0.056 | 7.9 | NA | 4.1 | 5.8 | NA | 0.9 | 0.54139 | 0.51128 | 1.31363 | 1.93633 | 0.8112 | 0.25085 | 0.32953 |
| 462 | Saxifraga tridactylites L. | Saxifraga tridactylites | 0.02 | 0.75 | 0.032 | NA | NA | NA | NA | NA | NA | NA | NA | NA | NA | NA | NA | NA |
| 462 | Myosotis arvensis Hill | Myosotis arvensis (intermedia) | 0.01 | 0.75 | 0.086 | 6.7 | NA | NA | 6.5 | 7.7 | 0.1 | 0.70945 | 0.61373 | 0.80583 | 1.7717 | 0.43958 | 0.26577 | 0.51423 |
| 463 | Taraxacum officinale F.H.Wigg. | Taraxacum officinale agg. | 0.11 | 0.79 | 0.114 | NA | NA | 4.5 | 6.6 | 6.6 | 0 | 0.78059 | 0.60963 | 0.54418 | 1.63701 | 0.26733 | 0.16232 | 0.54354 |
| 463 | Geranium molle L. | Geranium molle | 0.05 | 0.79 | 0.116 | NA | NA | 6.4 | 6.8 | 6.8 | 0 | 0.66551 | 0.27982 | 0.17419 | 1.41859 | 0.08796 | 0.19647 | 0.14036 |
| 463 | Bellis perennis L. | Bellis perennis | 0.08 | 0.79 | 0.143 | 6.7 | NA | NA | 6.5 | 7.7 | 0.1 | 0.70945 | 0.61373 | 0.80583 | 1.7717 | 0.43958 | 0.26577 | 0.51423 |
| 463 | Trifolium dubium Sibth. | Trifolium dubium (minus) | 0.1 | 0.79 | 0.170 | 6.7 | NA | NA | 6.5 | 7.7 | 0.1 | 0.70945 | 0.61373 | 0.80583 | 1.7717 | 0.43958 | 0.26577 | 0.51423 |
| 463 | Trifolium repens L. | Trifolium repens | 0.15 | 0.79 | 0.253 | NA | NA | NA | NA | NA | NA | NA | NA | NA | NA | NA | NA | NA |
| 463 | Hypericum perforatum L. | Hypericum perforatum | 0.05 | 0.79 | 0.154 | 7.4 | 5.3 | NA | 6 | 7.7 | 0.8 | 0.65201 | 0.63791 | 1.54898 | 2.03286 | 0.77966 | 0.37419 | 0.45721 |
| 463 | Myosotis arvensis Hill | Myosotis arvensis (intermedia) | 0.02 | 0.79 | 0.183 | 4.7 | NA | NA | 5.8 | 6.6 | 0 | 0.68705 | 0.16985 | 0.0706 | 1.32057 | 0.02196 | 0.18337 | 0.11293 |
| 463 | Sedum sexangulare L. | Sedum sexangulare | 0.01 | 0.79 | 0.050 | 8.1 | NA | 3.5 | 6.1 | NA | 0.2 | 0.68175 | 0.6716 | 1.70824 | 1.99767 | 0.41848 | 0.25411 | 0.57318 |
| 463 | Cerastium semidecandrum L. | Cerastium semidecandrum | 0.08 | 0.79 | 0.089 | 7.8 | NA | 4.6 | 6.2 | 5.8 | 0.1 | 0.71321 | 0.68276 | 1.29546 | 1.8275 | 0.62188 | 0.18605 | 0.48742 |
| 463 | Arenaria serpyllifolia L. | Arenaria serpyllifolia agg. | 0.01 | 0.79 | 0.118 | 7.1 | 4.9 | NA | 6 | NA | 1 | 0.59517 | 0.59064 | 1.89772 | 2.12267 | 0.65091 | 0.41213 | 0.37185 |
| 463 | Plantago major L. | Plantago major ssp. major | 0.04 | 0.79 | 0.060 | 7 | 5.6 | 4.4 | 5.9 | 5.4 | 0.1 | 0.64871 | 0.63945 | 1.70759 | 2.05577 | 0.56937 | 0.21649 | 0.54803 |
| 463 | Poa annua L. | Poa annua | 0.08 | 0.79 | 0.061 | 7.1 | 4.9 | NA | 6 | NA | 1 | 0.59517 | 0.59064 | 1.89772 | 2.12267 | 0.65091 | 0.41213 | 0.37185 |
| 463 | Senecio vulgaris L. | Senecio vulgaris | 0.01 | 0.79 | 0.079 | 7.1 | 4.9 | NA | 6 | NA | 1 | 0.59517 | 0.59064 | 1.89772 | 2.12267 | 0.65091 | 0.41213 | 0.37185 |
| 464 | Poa annua L. | Poa annua | 0.4 | 0.85 | 0.253 | 7.3 | 5.4 | 4.1 | 7.6 | 4.2 | 0.4 | 0.58728 | 0.56055 | 1.34601 | 1.94534 | 0.82357 | 0.23743 | 0.34603 |
| 464 | Matricaria discoidea DC. | Matricaria discoidea | 0.05 | 0.85 | 0.055 | 6.6 | 5.6 | NA | 5.6 | 4.1 | 0.1 | 0.50969 | 0.50576 | 1.97957 | 2.23767 | 1.1694 | 0.28385 | 0.36995 |
| 464 | Draba verna L. | Erophila verna | 0.05 | 0.85 | 0.099 | 8.1 | NA | NA | 5.2 | NA | 0.1 | 0.51276 | 0.49581 | 1.51667 | 2.20989 | 0.98959 | 0.46997 | 0.3594 |
| 464 | Capsella bursa-pastoris Medik. | Capsella bursa-pastoris | 0.15 | 0.85 | 0.031 | 7.1 | 7.8 | 4.1 | 6.2 | 6 | 0.1 | 0.76637 | 0.65623 | 0.78647 | 1.82229 | 0.56568 | 0.18972 | 0.47454 |
| 464 | Trifolium dubium Sibth. | Trifolium dubium (minus) | 0.1 | 0.85 | 0.053 | 7.7 | NA | NA | 5.6 | NA | 0.7 | 0.53223 | 0.52325 | 1.73719 | 2.14421 | 0.94278 | 0.39374 | 0.35054 |
| 464 | Taraxacum officinale F.H.Wigg. | Taraxacum officinale agg. | 0.1 | 0.85 | 0.063 | 7.3 | NA | 3.9 | 6.1 | 3.9 | 0.1 | 0.55903 | 0.41538 | 0.65189 | 1.65181 | 0.43719 | 0.24279 | 0.21709 |
| 465 | Poa annua L. | Poa annua | 0.1 | 0.68 | 0.178 | 6.6 | 5.6 | NA | 5.6 | 4.1 | 0.1 | 0.50969 | 0.50576 | 1.97957 | 2.23767 | 1.1694 | 0.28385 | 0.36995 |
| 465 | Bromus sterilis L. | Bromus sterilis | 0.15 | 0.68 | 0.172 | 7.1 | 4.9 | NA | 6 | NA | 1 | 0.59517 | 0.59064 | 1.89772 | 2.12267 | 0.65091 | 0.41213 | 0.37185 |
| 465 | Veronica arvensis L. | Veronica arvensis | 0.1 | 0.68 | 0.288 | NA | NA | NA | NA | NA | NA | NA | NA | NA | NA | NA | NA | NA |
| 465 | Arenaria serpyllifolia L. | Arenaria serpyllifolia agg. | 0.03 | 0.68 | 0.034 | 6.7 | NA | NA | 7.6 | 4.1 | 0 | 0.6581 | 0.15469 | 0.101 | 1.2995 | 0.02364 | 0.20663 | 0.10228 |
| 465 | Taraxacum officinale F.H.Wigg. | Taraxacum officinale agg. | 0.15 | 0.68 | 0.186 | 7.1 | 4.9 | NA | 6 | NA | 1 | 0.59517 | 0.59064 | 1.89772 | 2.12267 | 0.65091 | 0.41213 | 0.37185 |
| 465 | Cerastium semidecandrum L. | Cerastium semidecandrum | 0.15 | 0.68 | 0.076 | 8 | NA | 4.6 | 8 | 5.5 | 0.5 | 0.67376 | 0.64463 | 1.40205 | 1.95891 | 0.49218 | 0.31373 | 0.371 |
| 466 | Taraxacum officinale F.H.Wigg. | Taraxacum officinale agg. | 0.1 | 0.73 | 0.221 | 7.3 | NA | NA | 7.6 | 7.7 | 0.4 | 0.75479 | 0.72524 | 1.28692 | 1.77374 | 0.61324 | 0.18283 | 0.56773 |
| 466 | Veronica arvensis L. | Veronica arvensis | 0.1 | 0.73 | 0.074 | 7.1 | 4.9 | NA | 6 | NA | 1 | 0.59517 | 0.59064 | 1.89772 | 2.12267 | 0.65091 | 0.41213 | 0.37185 |
| 466 | Hypochaeris glabra L. | Hypochaeris glabra | 0.15 | 0.73 | 0.190 | 7.4 | 5.3 | NA | 6 | 7.7 | 0.8 | 0.65201 | 0.63791 | 1.54898 | 2.03286 | 0.77966 | 0.37419 | 0.45721 |
| 466 | Poa annua L. | Poa annua | 0.05 | 0.73 | 0.395 | 7.4 | 5.3 | NA | 6 | 7.7 | 0.8 | 0.65201 | 0.63791 | 1.54898 | 2.03286 | 0.77966 | 0.37419 | 0.45721 |
| 466 | Plantago lanceolata L. | Plantago lanceolata | 0.1 | 0.73 | 0.028 | 7.1 | 4.9 | NA | 6 | NA | 1 | 0.59517 | 0.59064 | 1.89772 | 2.12267 | 0.65091 | 0.41213 | 0.37185 |
| 466 | Trifolium dubium Sibth. | Trifolium dubium (minus) | 0.13 | 0.73 | 0.026 | 7.3 | NA | 3.9 | 6.1 | 3.9 | 0.1 | 0.55903 | 0.41538 | 0.65189 | 1.65181 | 0.43719 | 0.24279 | 0.21709 |
| 466 | Erigeron canadensis L. | Conyza canadensis | 0.05 | 0.73 | 0.194 | 4.7 | 5.8 | 5.6 | 6.5 | 6.3 | 0 | 0.69843 | 0.15019 | 0.04615 | 1.3308 | 0.01233 | 0.18685 | 0.11235 |
| 466 | Acer platanoides L. | Acer platanoides | 0.05 | 0.73 | 0.039 | 7.3 | NA | 3.9 | 6.1 | 3.9 | 0.1 | 0.55903 | 0.41538 | 0.65189 | 1.65181 | 0.43719 | 0.24279 | 0.21709 |
| 467 | Taraxacum officinale F.H.Wigg. | Taraxacum officinale agg. | 0.15 | 0.58 | 0.143 | 8.4 | 7.2 | 2.6 | 6.8 | 3.6 | 0.2 | 0.58203 | 0.49726 | 0.89718 | 1.5951 | 0.3257 | 0.17939 | 0.36267 |
| 467 | Poa annua L. | Poa annua | 0.1 | 0.58 | 0.057 | NA | NA | NA | NA | NA | NA | NA | NA | NA | NA | NA | NA | NA |
| 467 | Cerastium semidecandrum L. | Cerastium semidecandrum | 0.05 | 0.58 | 0.091 | 7.9 | NA | 4.1 | 5.8 | NA | 0.9 | 0.54139 | 0.51128 | 1.31363 | 1.93633 | 0.8112 | 0.25085 | 0.32953 |
| 467 | Veronica arvensis L. | Veronica arvensis | 0.1 | 0.58 | 0.188 | NA | NA | 5.5 | 6.2 | 6.8 | 0 | 0.67445 | 0.21586 | 0.12312 | 1.32299 | 0.04699 | 0.18459 | 0.11909 |
| 467 | Sagina procumbens L. | Sagina procumbens | 0.1 | 0.58 | 0.029 | 7.2 | 5.7 | 4.3 | 6.8 | NA | 0.1 | 0.54405 | 0.41742 | 0.70503 | 1.63482 | 0.49706 | 0.18636 | 0.24515 |
| 467 | Matricaria discoidea DC. | Matricaria discoidea | 0.08 | 0.58 | 0.079 | 7.3 | NA | 3.9 | 6.1 | 3.9 | 0.1 | 0.55903 | 0.41538 | 0.65189 | 1.65181 | 0.43719 | 0.24279 | 0.21709 |
| 468 | Erigeron canadensis L. | Conyza canadensis | 0.1 | 0.6 | 0.056 | 7.3 | 5.4 | 4.1 | 7.6 | 4.2 | 0.4 | 0.58728 | 0.56055 | 1.34601 | 1.94534 | 0.82357 | 0.23743 | 0.34603 |
| 468 | Senecio inaequidens DC. | Senecio inaequidens | 0.15 | 0.6 | 0.032 | 7.3 | NA | 3.9 | 6.1 | 3.9 | 0.1 | 0.55903 | 0.41538 | 0.65189 | 1.65181 | 0.43719 | 0.24279 | 0.21709 |
| 468 | Poa annua L. | Poa annua | 0.1 | 0.6 | 0.034 | 7 | 5.6 | 4.4 | 5.9 | 5.4 | 0.1 | 0.64871 | 0.63945 | 1.70759 | 2.05577 | 0.56937 | 0.21649 | 0.54803 |
| 468 | Acer pseudoplatanus L. | Acer pseudoplatanus | 0.2 | 0.6 | 0.114 | 7.4 | 5.3 | NA | 6 | 7.7 | 0.8 | 0.65201 | 0.63791 | 1.54898 | 2.03286 | 0.77966 | 0.37419 | 0.45721 |
| 468 | Lactuca serriola L. | Lactuca serriola (scariola) | 0.05 | 0.6 | 0.127 | 7.4 | 5.3 | NA | 6 | 7.7 | 0.8 | 0.65201 | 0.63791 | 1.54898 | 2.03286 | 0.77966 | 0.37419 | 0.45721 |
| 469 | Erigeron canadensis L. | Conyza canadensis | 0.25 | 0.55 | 0.032 | 4.7 | NA | NA | 5.8 | 6.6 | 0 | 0.68705 | 0.16985 | 0.0706 | 1.32057 | 0.02196 | 0.18337 | 0.11293 |
| 469 | Prunus virginiana L. | Prunus serotina | 0.3 | 0.55 | 0.102 | 7 | 5.6 | 4.4 | 5.9 | 5.4 | 0.1 | 0.64871 | 0.63945 | 1.70759 | 2.05577 | 0.56937 | 0.21649 | 0.54803 |
| 470 | Erigeron canadensis L. | Conyza canadensis | 0.3 | 0.35 | 0.308 | NA | NA | NA | NA | NA | NA | NA | NA | NA | NA | NA | NA | NA |
| 470 | Cardamine hirsuta L. | Cardamine hirsuta | 0.05 | 0.35 | 0.100 | 7.7 | NA | 3.7 | 5.8 | 6.4 | 0 | 0.70633 | 0.61633 | 0.84477 | 1.62276 | 0.49804 | 0.1651 | 0.39904 |
| 471 | Erigeron canadensis L. | Conyza canadensis | 0.15 | 0.2 | 0.143 | 7.4 | 5.3 | NA | 6 | 7.7 | 0.8 | 0.65201 | 0.63791 | 1.54898 | 2.03286 | 0.77966 | 0.37419 | 0.45721 |
| 471 | Sagina procumbens L. | Sagina procumbens | 0.05 | 0.2 | 0.110 | 7.4 | 5.3 | NA | 6 | 7.7 | 0.8 | 0.65201 | 0.63791 | 1.54898 | 2.03286 | 0.77966 | 0.37419 | 0.45721 |
| 472 | Taraxacum officinale F.H.Wigg. | Taraxacum officinale agg. | 0.09 | 0.37 | 0.024 | 7 | 5.6 | 4.4 | 5.9 | 5.4 | 0.1 | 0.64871 | 0.63945 | 1.70759 | 2.05577 | 0.56937 | 0.21649 | 0.54803 |

| | | | | | | | | | | | | | | | | | | |
|---|---|---|---|---|---|---|---|---|---|---|---|---|---|---|---|---|---|---|
| 472 | Stellaria media (L.) Vill. | Stellaria media agg. | 0.13 | 0.37 | 0.050 | 7.8 | NA | 4.6 | 6.2 | 5.8 | 0.1 | 0.71321 | 0.68276 | 1.29546 | 1.8275 | 0.62188 | 0.18605 | 0.48742 |
| 472 | Poa annua L. | Poa annua | 0.15 | 0.37 | 0.067 | 4.7 | 5.8 | 5.6 | 6.5 | 6.3 | 0 | 0.69843 | 0.15019 | 0.04615 | 1.3308 | 0.01233 | 0.18685 | 0.11235 |
| 474 | Asplenium ruta-muraria L. | Asplenium ruta-muraria | 0.1 | 0.33 | 0.088 | 7.3 | 5.4 | 4.1 | 7.6 | 4.2 | 0.4 | 0.58728 | 0.56055 | 1.34601 | 1.94534 | 0.82357 | 0.23743 | 0.34603 |
| 474 | Sagina procumbens L. | Sagina procumbens | 0.05 | 0.33 | 0.099 | 7.4 | 5.3 | NA | 6 | 7.7 | 0.8 | 0.65201 | 0.63791 | 1.54898 | 2.03286 | 0.77966 | 0.37419 | 0.45721 |
| 474 | Taraxacum officinale F.H.Wigg. | Taraxacum officinale agg. | 0.09 | 0.33 | 0.107 | 8.4 | NA | 3 | 6 | NA | 0 | 0.46249 | 0.43915 | 1.39189 | 1.83687 | 0.38437 | 0.30571 | 0.38883 |
| 474 | Poa annua L. | Poa annua | 0.09 | 0.33 | 0.025 | 6.7 | 5.3 | 4.5 | 5.3 | 5.8 | 0 | 0.75464 | 0.73051 | 1.36974 | 1.95508 | 0.45974 | 0.15567 | 0.6659 |
| 475 | Sagina procumbens L. | Sagina procumbens | 0.07 | 0.52 | 0.221 | 8.4 | NA | 3 | 6 | NA | 0 | 0.46249 | 0.43915 | 1.39189 | 1.83687 | 0.38437 | 0.30571 | 0.38883 |
| 475 | Veronica arvensis L. | Veronica arvensis | 0.05 | 0.52 | 0.068 | 7.8 | NA | 4.6 | 6.2 | 5.8 | 0.1 | 0.71321 | 0.68276 | 1.29546 | 1.8275 | 0.62188 | 0.18605 | 0.48742 |
| 475 | Stellaria media (L.) Vill. | Stellaria media agg. | 0.15 | 0.52 | 0.138 | 7.9 | 5.1 | 5.1 | 6.7 | 7.7 | 0.2 | 0.75642 | 0.75467 | 2.08644 | 2.23861 | 0.83282 | 0.22413 | 0.56443 |
| 475 | Poa annua L. | Poa annua | 0.1 | 0.52 | 0.273 | 7.4 | 5.3 | NA | 6 | 7.7 | 0.8 | 0.65201 | 0.63791 | 1.54898 | 2.03286 | 0.77966 | 0.37419 | 0.45721 |
| 475 | Taraxacum officinale F.H.Wigg. | Taraxacum officinale agg. | 0.15 | 0.52 | 0.102 | 6.7 | NA | NA | 6.5 | 7.7 | 0.1 | 0.70945 | 0.61373 | 0.80583 | 1.7717 | 0.43958 | 0.26577 | 0.51423 |
| 476 | Taraxacum officinale F.H.Wigg. | Taraxacum officinale agg. | 0.15 | 0.59 | 0.227 | 7.4 | 5.3 | NA | 6 | 7.7 | 0.8 | 0.65201 | 0.63791 | 1.54898 | 2.03286 | 0.77966 | 0.37419 | 0.45721 |
| 476 | Poa annua L. | Poa annua | 0.13 | 0.59 | 0.065 | 7.4 | 5.2 | 4.7 | 6.1 | NA | 0 | 0.77406 | 0.76726 | 1.79113 | 2.07279 | 0.64778 | 0.2119 | 0.65925 |
| 476 | Sagina procumbens L. | Sagina procumbens | 0.08 | 0.59 | 0.074 | NA | NA | 5.5 | 6.2 | 6.8 | 0 | 0.67445 | 0.21586 | 0.12312 | 1.32299 | 0.04699 | 0.18459 | 0.11909 |
| 476 | Hedera helix L. | Hedera helix | 0.05 | 0.59 | 0.026 | 7.9 | NA | 4.1 | 5.8 | NA | 0.9 | 0.54139 | 0.51128 | 1.31363 | 1.93633 | 0.8112 | 0.25085 | 0.32953 |
| 476 | Ligustrum vulgare L. | Ligustrum vulgare | 0.02 | 0.59 | 0.039 | NA | NA | NA | NA | NA | NA | NA | NA | NA | NA | NA | NA | NA |
| 476 | Stellaria media (L.) Vill. | Stellaria media agg. | 0.06 | 0.59 | 0.143 | 5.2 | 5.9 | 5.4 | 7.3 | 8 | 0 | 0.67947 | 0.25214 | 0.1495 | 1.33661 | 0.07498 | 0.19025 | 0.12819 |
| 476 | Veronica arvensis L. | Veronica arvensis | 0.05 | 0.59 | 0.114 | 7.8 | NA | NA | 7.1 | 5.7 | 0.2 | 0.62289 | 0.58174 | 1.15194 | 1.63402 | 0.69874 | 0.1726 | 0.30211 |
| 476 | NA | NA | 0.05 | 0.59 | 0.029 | 8.3 | 5.2 | 3.5 | 6.7 | 4 | 0.1 | 0.59253 | 0.57367 | 1.48069 | 1.91606 | 0.44486 | 0.27236 | 0.42509 |
| 477 | Taraxacum officinale F.H.Wigg. | Taraxacum officinale agg. | 0.1 | 0.43 | 0.043 | 7.2 | 5.7 | 4.3 | 6.8 | NA | 0.1 | 0.54405 | 0.41742 | 0.70503 | 1.63482 | 0.49706 | 0.18636 | 0.24515 |
| 477 | Asplenium ruta-muraria L. | Asplenium ruta-muraria | 0.15 | 0.43 | 0.079 | 7.9 | NA | 4.1 | 5.8 | NA | 0.9 | 0.54139 | 0.51128 | 1.31363 | 1.93633 | 0.8112 | 0.25085 | 0.32953 |
| 477 | Cardamine hirsuta L. | Cardamine hirsuta | 0.1 | 0.43 | 0.290 | NA | NA | NA | NA | NA | NA | NA | NA | NA | NA | NA | NA | NA |
| 477 | Sagina procumbens L. | Sagina procumbens | 0.08 | 0.43 | 0.204 | 6.7 | NA | NA | 6.5 | 7.7 | 0.1 | 0.70945 | 0.61373 | 0.80583 | 1.7717 | 0.43958 | 0.26577 | 0.51423 |
| 478 | Veronica arvensis L. | Veronica arvensis | 0.1 | 0.66 | 0.031 | 7.9 | 5.7 | 4.2 | 7 | 7.2 | 0.1 | 0.76998 | 0.7507 | 1.46055 | 1.90983 | 0.91556 | 0.1857 | 0.55131 |
| 478 | Saxifraga tridactylites L. | Saxifraga tridactylites | 0.08 | 0.66 | 0.183 | 7.7 | NA | 3.7 | 5.8 | 6.4 | 0 | 0.70633 | 0.61633 | 0.84477 | 1.62276 | 0.49804 | 0.1651 | 0.39904 |
| 478 | Asplenium ruta-muraria L. | Asplenium ruta-muraria | 0.1 | 0.66 | 0.325 | 4.7 | NA | NA | 5.8 | 6.6 | 0 | 0.68705 | 0.16985 | 0.0706 | 1.32057 | 0.02196 | 0.18337 | 0.11293 |
| 478 | Sagina procumbens L. | Sagina procumbens | 0.05 | 0.66 | 0.232 | NA | NA | NA | NA | NA | NA | NA | NA | NA | NA | NA | NA | NA |
| 478 | Plantago major L. | Plantago major ssp. major | 0.08 | 0.66 | 0.024 | 7.4 | 5.3 | NA | 6 | 7.7 | 0.8 | 0.65201 | 0.63791 | 1.54898 | 2.03286 | 0.77966 | 0.37419 | 0.45721 |
| 478 | Crepis sancta (L.) Babc. | Crepis sancta | 0.05 | 0.66 | 0.024 | 7.4 | 5.3 | NA | 6 | 7.7 | 0.8 | 0.65201 | 0.63791 | 1.54898 | 2.03286 | 0.77966 | 0.37419 | 0.45721 |
| 478 | Poa annua L. | Poa annua | 0.15 | 0.66 | 0.055 | 7.1 | 7.8 | 4.1 | 6.2 | 6 | 0.1 | 0.76637 | 0.65623 | 0.78647 | 1.82229 | 0.56568 | 0.18972 | 0.47454 |
| 478 | Oxalis corniculata L. | Oxalis corniculata | 0.05 | 0.66 | 0.077 | 8.6 | 8 | 3.3 | 3 | 1.8 | 0.2 | 0.49702 | 0.46825 | 1.42549 | 1.99857 | 0.4756 | 0.32016 | 0.338 |
| 479 | Taraxacum officinale F.H.Wigg. | Taraxacum officinale agg. | 0.13 | 0.77 | 0.107 | 8.1 | NA | NA | 4.4 | 2.8 | 0.3 | 0.48868 | 0.46508 | 1.4426 | 2.04965 | 0.7278 | 0.33931 | 0.35628 |
| 479 | Moehringia trinervia [Clairv.] | Moehringia trinervia | 0.08 | 0.77 | 0.013 | 7.3 | 5.3 | 2.7 | 6.3 | 1.5 | 0 | 0.47424 | 0.43924 | 1.22297 | 1.85433 | 0.44008 | 0.30378 | 0.24409 |
| 479 | Poa annua L. | Poa annua | 0.17 | 0.77 | 0.068 | 4.7 | 5.8 | 5.6 | 6.5 | 6.3 | 0 | 0.69843 | 0.15019 | 0.04615 | 1.3308 | 0.01233 | 0.18685 | 0.11235 |
| 479 | Veronica arvensis L. | Veronica arvensis | 0.1 | 0.77 | 0.085 | 7 | 5.6 | 4.4 | 5.9 | 5.4 | 0.1 | 0.64871 | 0.63945 | 1.70759 | 2.05577 | 0.56937 | 0.21649 | 0.54803 |
| 479 | Sonchus oleraceus L. | Sonchus oleraceus | 0.17 | 0.77 | 0.076 | 7.1 | 7.8 | 4.1 | 6.2 | 6 | 0.1 | 0.76637 | 0.65623 | 0.78647 | 1.82229 | 0.56568 | 0.18972 | 0.47454 |
| 479 | Capsella bursa-pastoris Medik. | Capsella bursa-pastoris | 0.05 | 0.77 | 0.065 | 7.1 | 4.9 | NA | 6 | NA | 1 | 0.59517 | 0.59064 | 1.89772 | 2.12267 | 0.65091 | 0.41213 | 0.37185 |
| 479 | Sagina procumbens L. | Sagina procumbens | 0.05 | 0.77 | 0.077 | 7.4 | 5.3 | NA | 6 | 7.7 | 0.8 | 0.65201 | 0.63791 | 1.54898 | 2.03286 | 0.77966 | 0.37419 | 0.45721 |
| 479 | Asplenium ruta-muraria L. | Asplenium ruta-muraria | 0.02 | 0.77 | 0.028 | 7.4 | 5.2 | 4.7 | 6.1 | NA | 0 | 0.77406 | 0.76726 | 1.79113 | 2.07279 | 0.64778 | 0.2119 | 0.65925 |
| 480 | Taraxacum officinale F.H.Wigg. | Taraxacum officinale agg. | 0.15 | 0.68 | 0.159 | 8.1 | NA | NA | 4.4 | 2.8 | 0.3 | 0.48868 | 0.46508 | 1.4426 | 2.04965 | 0.7278 | 0.33931 | 0.35628 |
| 480 | Asplenium ruta-muraria L. | Asplenium ruta-muraria | 0.05 | 0.68 | 0.057 | 4.7 | 5.8 | 5.6 | 6.5 | 6.3 | 0 | 0.69843 | 0.15019 | 0.04615 | 1.3308 | 0.01233 | 0.18685 | 0.11235 |
| 480 | Poa annua L. | Poa annua | 0.25 | 0.68 | 0.079 | 7.1 | 5.3 | 4.5 | 5.9 | 4.8 | 0.3 | 0.52518 | 0.5113 | 1.61045 | 2.05509 | 0.96495 | 0.27897 | 0.32911 |
| 480 | Veronica arvensis L. | Veronica arvensis | 0.08 | 0.68 | 0.048 | NA | NA | NA | NA | NA | NA | NA | NA | NA | NA | NA | NA | NA |
| 480 | Glechoma hederacea L. | Glechoma hederacea | 0.05 | 0.68 | 0.041 | 7.3 | 6.1 | 3.8 | 6 | 5.1 | 0 | 0.62987 | 0.59281 | 1.27995 | 1.90491 | 0.6275 | 0.3036 | 0.42608 |
| 480 | Sagina procumbens L. | Sagina procumbens | 0.05 | 0.68 | 0.013 | 6.7 | NA | NA | 6.5 | 7.7 | 0.1 | 0.70945 | 0.61373 | 0.80583 | 1.7717 | 0.43958 | 0.26577 | 0.51423 |
| 480 | Geranium robertianum L. | Geranium robertianum | 0.05 | 0.68 | 0.231 | 6.6 | 5.7 | NA | 6.3 | 7.6 | 0.1 | 0.67317 | 0.45023 | 0.43732 | 1.4267 | 0.22701 | 0.17304 | 0.27906 |
| 481 | Taraxacum officinale F.H.Wigg. | Taraxacum officinale agg. | 0.25 | 0.5 | 0.025 | 5.7 | NA | NA | 7.3 | 5.4 | 0 | 0.6918 | 0.13881 | 0.03984 | 1.31841 | 0.01173 | 0.19643 | 0.10363 |
| 481 | Veronica arvensis L. | Veronica arvensis | 0.13 | 0.5 | 0.089 | 7.3 | NA | 3.9 | 6.1 | 3.9 | 0.1 | 0.55903 | 0.41538 | 0.65189 | 1.65181 | 0.43719 | 0.24279 | 0.21709 |
| 481 | Poa annua L. | Poa annua | 0.1 | 0.5 | 0.025 | 6.7 | 5.3 | 4.5 | 5.3 | 5.8 | 0 | 0.75464 | 0.73051 | 1.36974 | 1.95508 | 0.45974 | 0.15567 | 0.6659 |
| 481 | Sagina procumbens L. | Sagina procumbens | 0.02 | 0.5 | 0.031 | 8.3 | 5.2 | 3.5 | 6.7 | 4 | 0.1 | 0.59253 | 0.57367 | 1.48069 | 1.91606 | 0.44486 | 0.27236 | 0.42509 |
| 482 | Taraxacum officinale F.H.Wigg. | Taraxacum officinale agg. | 0.1 | 0.63 | 0.027 | 7.3 | 5.3 | 2.7 | 6.3 | 1.5 | 0 | 0.47424 | 0.43924 | 1.22297 | 1.85433 | 0.44008 | 0.30378 | 0.24409 |
| 482 | Asplenium trichomanes L. | Asplenium trichomanes | 0.17 | 0.63 | 0.085 | NA | NA | NA | NA | NA | NA | NA | NA | NA | NA | NA | NA | NA |
| 482 | Asplenium ruta-muraria L. | Asplenium ruta-muraria | 0.06 | 0.63 | 0.079 | 6.5 | NA | NA | 5.4 | NA | 0 | 0.59769 | 0.45571 | 0.64594 | 1.51584 | 0.28825 | 0.22901 | 0.32316 |
| 482 | Geranium robertianum L. | Geranium robertianum | 0.02 | 0.63 | 0.028 | 6.7 | NA | NA | 6.5 | 7.7 | 0.1 | 0.70945 | 0.61373 | 0.80583 | 1.7717 | 0.43958 | 0.26577 | 0.51423 |
| 482 | Sagina procumbens L. | Sagina procumbens | 0.06 | 0.63 | 0.032 | 7.3 | NA | NA | 7.6 | 7.7 | 0.4 | 0.75479 | 0.72524 | 1.28692 | 1.77374 | 0.61324 | 0.18283 | 0.56773 |
| 482 | Poa annua L. | Poa annua | 0.12 | 0.63 | 0.063 | 7.8 | NA | 4.6 | 6.2 | 5.8 | 0.1 | 0.71321 | 0.68276 | 1.29546 | 1.8275 | 0.62188 | 0.18605 | 0.48742 |
| 482 | Cardamine hirsuta L. | Cardamine hirsuta | 0.05 | 0.63 | 0.056 | 6.5 | NA | NA | 5.4 | NA | 0 | 0.59769 | 0.45571 | 0.64594 | 1.51584 | 0.28825 | 0.22901 | 0.32316 |
| 482 | Veronica arvensis L. | Veronica arvensis | 0.05 | 0.63 | 0.016 | 7.3 | 5.4 | 4.1 | 7.6 | 4.2 | 0.4 | 0.58728 | 0.56055 | 1.34601 | 1.94534 | 0.82357 | 0.23743 | 0.34603 |
| 483 | Taraxacum officinale F.H.Wigg. | Taraxacum officinale agg. | 0.28 | 0.87 | 0.129 | 7.8 | NA | NA | 7.1 | 5.7 | 0.2 | 0.62289 | 0.58174 | 1.15194 | 1.63402 | 0.69874 | 0.1726 | 0.30211 |
| 483 | Veronica arvensis L. | Veronica arvensis | 0.23 | 0.87 | 0.016 | 7.4 | 5.2 | 4.7 | 6.1 | NA | 0 | 0.77406 | 0.76726 | 1.79113 | 2.07279 | 0.64778 | 0.2119 | 0.65925 |
| 483 | Hypochaeris radicata L. | Hypochoeris radicata | 0.1 | 0.87 | 0.633 | 4.7 | NA | NA | 5.8 | 6.6 | 0 | 0.68705 | 0.16985 | 0.0706 | 1.32057 | 0.02196 | 0.18337 | 0.11293 |
| 483 | Poa annua L. | Poa annua | 0.18 | 0.87 | 0.031 | 7.3 | 5.8 | 4.8 | 5.3 | 5.6 | 0 | 0.67083 | 0.63295 | 1.24583 | 1.99939 | 0.56795 | 0.28417 | 0.48569 |
| 483 | Erigeron canadensis L. | Conyza canadensis | 0.08 | 0.87 | 0.055 | 7.4 | 5.3 | NA | 6 | 7.7 | 0.8 | 0.65201 | 0.63791 | 1.54898 | 2.03286 | 0.77966 | 0.37419 | 0.45721 |
| 484 | Taraxacum officinale F.H.Wigg. | Taraxacum officinale agg. | 0.15 | 0.55 | 0.037 | 8.4 | NA | 3 | 6 | NA | 0 | 0.46249 | 0.43915 | 1.39189 | 1.83687 | 0.38437 | 0.30571 | 0.38883 |
| 484 | Poa annua L. | Poa annua | 0.2 | 0.55 | 0.101 | 8.4 | NA | 3 | 6 | NA | 0 | 0.46249 | 0.43915 | 1.39189 | 1.83687 | 0.38437 | 0.30571 | 0.38883 |
| 484 | Stellaria media (L.) Vill. | Stellaria media agg. | 0.1 | 0.55 | 0.026 | 7.9 | 5.4 | 2.9 | 7.9 | 2.4 | 0 | 0.48324 | 0.22836 | 0.33987 | 1.12718 | 0.05497 | 0.20766 | 0.12487 |
| 484 | Veronica arvensis L. | Veronica arvensis | 0.1 | 0.55 | 0.079 | 7 | 5.6 | 4.4 | 5.9 | 5.4 | 0.1 | 0.64871 | 0.63945 | 1.70759 | 2.05577 | 0.56937 | 0.21649 | 0.54803 |
| 485 | Poa annua L. | Poa annua | 0.35 | 0.4 | 0.106 | 8.8 | NA | 3 | NA | NA | 0 | 0.55609 | 0.48243 | 0.91574 | 1.84215 | 0.36932 | 0.29356 | 0.3049 |
| 485 | Taraxacum officinale F.H.Wigg. | Taraxacum officinale agg. | 0.05 | 0.4 | 0.028 | 7.4 | 5.3 | NA | 6 | 7.7 | 0.8 | 0.65201 | 0.63791 | 1.54898 | 2.03286 | 0.77966 | 0.37419 | 0.45721 |
| 486 | Poa annua L. | Poa annua | 0.35 | 0.65 | 0.123 | 6.7 | NA | NA | 6.5 | 7.7 | 0.1 | 0.70945 | 0.61373 | 0.80583 | 1.7717 | 0.43958 | 0.26577 | 0.51423 |
| 486 | Veronica arvensis L. | Veronica arvensis | 0.1 | 0.65 | 0.017 | 5.7 | NA | NA | 7.3 | 5.4 | 0 | 0.6918 | 0.13881 | 0.03984 | 1.31841 | 0.01173 | 0.19643 | 0.10363 |
| 486 | Erigeron canadensis L. | Conyza canadensis | 0.2 | 0.65 | 0.055 | 7.3 | 5.3 | 2.7 | 6.3 | 1.5 | 0 | 0.47424 | 0.43924 | 1.22297 | 1.85433 | 0.44008 | 0.30378 | 0.24409 |
| 487 | Taraxacum officinale F.H.Wigg. | Taraxacum officinale agg. | 0.15 | 0.38 | 0.062 | 7.3 | 5.3 | 2.7 | 6.3 | 1.5 | 0 | 0.47424 | 0.43924 | 1.22297 | 1.85433 | 0.44008 | 0.30378 | 0.24409 |
| 487 | Sagina procumbens L. | Sagina procumbens | 0.08 | 0.38 | 0.031 | 7.1 | 5.3 | 4 | 4.4 | 2.8 | 0 | 0.53846 | 0.48278 | 1.02807 | 1.87452 | 0.70983 | 0.24362 | 0.28858 |
| 487 | Poa annua L. | Poa annua | 0.15 | 0.38 | 0.133 | NA | NA | NA | 6.6 | NA | 0 | 0.65326 | 0.6245 | 1.32492 | 1.96985 | 0.60133 | 0.19748 | 0.49592 |
| 488 | Sagina procumbens L. | Sagina procumbens | 0.2 | 0.5 | 0.013 | 8.3 | 5.2 | 3.5 | 6.7 | 4 | 0.1 | 0.59253 | 0.57367 | 1.48069 | 1.91606 | 0.44486 | 0.27236 | 0.42509 |
| 488 | Poa annua L. | Poa annua | 0.2 | 0.5 | 0.143 | 6.5 | NA | NA | 5.4 | NA | 0 | 0.59769 | 0.45571 | 0.64594 | 1.51584 | 0.28825 | 0.22901 | 0.32316 |
| 488 | Taraxacum officinale F.H.Wigg. | Taraxacum officinale agg. | 0.1 | 0.5 | 0.116 | 7.3 | 5.4 | 4.1 | 7.6 | 4.2 | 0.4 | 0.58728 | 0.56055 | 1.34601 | 1.94534 | 0.82357 | 0.23743 | 0.34603 |
| 489 | Taraxacum officinale F.H.Wigg. | Taraxacum officinale agg. | 0.3 | 0.7 | 0.111 | NA | NA | NA | NA | NA | NA | NA | NA | NA | NA | NA | NA | NA |
| 489 | Oxalis corniculata L. | Oxalis corniculata | 0.05 | 0.7 | 0.102 | NA | NA | NA | NA | NA | NA | NA | NA | NA | NA | NA | NA | NA |
| 489 | Capsella bursa-pastoris Medik. | Capsella bursa-pastoris | 0.15 | 0.7 | 0.063 | 7.4 | 5.3 | NA | 6 | 7.7 | 0.8 | 0.65201 | 0.63791 | 1.54898 | 2.03286 | 0.77966 | 0.37419 | 0.45721 |
| 489 | Sagina procumbens L. | Sagina procumbens | 0.1 | 0.7 | 0.033 | 6.5 | NA | NA | 5.4 | NA | 0 | 0.59769 | 0.45571 | 0.64594 | 1.51584 | 0.28825 | 0.22901 | 0.32316 |
| 489 | Poa annua L. | Poa annua | 0.1 | 0.7 | 0.018 | 6.7 | NA | NA | 6.5 | 7.7 | 0.1 | 0.70945 | 0.61373 | 0.80583 | 1.7717 | 0.43958 | 0.26577 | 0.51423 |
| 490 | Sagina procumbens L. | Sagina procumbens | 0.3 | 0.65 | 0.123 | 8.1 | NA | NA | 4.4 | 2.8 | 0.3 | 0.48868 | 0.46508 | 1.4426 | 2.04965 | 0.7278 | 0.33931 | 0.35628 |
| 490 | Stellaria media (L.) Vill. | Stellaria media agg. | 0.2 | 0.65 | 0.013 | 7.2 | 5.5 | 4.5 | 6 | 7.6 | 0.3 | 0.72905 | 0.70188 | 1.35981 | 1.84158 | 0.57912 | 0.20081 | 0.60725 |
| 490 | Taraxacum officinale F.H.Wigg. | Taraxacum officinale agg. | 0.1 | 0.65 | 0.051 | 7.3 | NA | NA | 5.8 | 5.9 | 0.5 | 0.63207 | 0.61738 | 1.52973 | 2.00883 | 0.70322 | 0.36696 | 0.39725 |
| 490 | Poa annua L. | Poa annua | 0.05 | 0.65 | 0.058 | NA | NA | NA | NA | NA | NA | NA | NA | NA | NA | NA | NA | NA |
| 491 | Poa annua L. | Poa annua | 0.07 | 0.71 | 0.426 | NA | NA | 5.5 | 6.2 | 6.8 | 0 | 0.67445 | 0.21586 | 0.12312 | 1.32299 | 0.04699 | 0.18459 | 0.11909 |
| 491 | Bromus sterilis L. | Bromus sterilis | 0.3 | 0.71 | 0.102 | 4.7 | NA | NA | 5.8 | 6.6 | 0 | 0.68705 | 0.16985 | 0.0706 | 1.32057 | 0.02196 | 0.18337 | 0.11293 |
| 491 | Bellis perennis L. | Bellis perennis | 0.05 | 0.71 | 0.031 | 5.7 | NA | NA | 7.3 | 5.4 | 0 | 0.6918 | 0.13881 | 0.03984 | 1.31841 | 0.01173 | 0.19643 | 0.10363 |
| 491 | Geranium robertianum L. | Geranium robertianum | 0.13 | 0.71 | 0.099 | 7.1 | 4.9 | NA | 6 | NA | 1 | 0.59517 | 0.59064 | 1.89772 | 2.12267 | 0.65091 | 0.41213 | 0.37185 |
| 491 | Taraxacum officinale F.H.Wigg. | Taraxacum officinale agg. | 0.1 | 0.71 | 0.111 | 7.1 | 4.9 | NA | 6 | NA | 1 | 0.59517 | 0.59064 | 1.89772 | 2.12267 | 0.65091 | 0.41213 | 0.37185 |
| 491 | Sagina procumbens L. | Sagina procumbens | 0.02 | 0.71 | 0.027 | 8.2 | NA | 2.2 | 7 | 2.1 | 0 | 0.50737 | 0.49072 | 1.5234 | 1.7929 | 0.17792 | 0.32947 | 0.3784 |
| 491 | Capsella bursa-pastoris Medik. | Capsella bursa-pastoris | 0.02 | 0.71 | 0.101 | 7.4 | 5.3 | NA | 6 | 7.7 | 0.8 | 0.65201 | 0.63791 | 1.54898 | 2.03286 | 0.77966 | 0.37419 | 0.45721 |
| 491 | Stellaria media (L.) Vill. | Stellaria media agg. | 0.02 | 0.71 | 0.021 | 8.3 | 5.2 | 3.5 | 6.7 | 4 | 0.1 | 0.59253 | 0.57367 | 1.48069 | 1.91606 | 0.44486 | 0.27236 | 0.42509 |
| 492 | Taraxacum officinale F.H.Wigg. | Taraxacum officinale agg. | 0.25 | 0.55 | 0.032 | 6.7 | NA | NA | 6.5 | 7.7 | 0.1 | 0.70945 | 0.61373 | 0.80583 | 1.7717 | 0.43958 | 0.26577 | 0.51423 |
| 492 | Plantago major L. | Plantago major ssp. major | 0.1 | 0.55 | 0.102 | 7.3 | 5.8 | 4.8 | 5.3 | 5.6 | 0 | 0.67083 | 0.63295 | 1.24583 | 1.99939 | 0.56795 | 0.28417 | 0.48569 |
| 492 | Capsella bursa-pastoris Medik. | Capsella bursa-pastoris | 0.05 | 0.55 | 0.031 | 4.7 | NA | NA | 5.8 | 6.6 | 0 | 0.68705 | 0.16985 | 0.0706 | 1.32057 | 0.02196 | 0.18337 | 0.11293 |
| 492 | Poa annua L. | Poa annua | 0.15 | 0.55 | 0.062 | 6.8 | NA | 5.6 | 5.8 | NA | 0.1 | 0.52169 | 0.45097 | 0.93044 | 1.87015 | 0.67593 | 0.2975 | 0.27054 |
| 493 | Stellaria media (L.) Vill. | Stellaria media agg. | 0.15 | 0.8 | 0.013 | 6.7 | 5.3 | 4.5 | 5.3 | 5.8 | 0 | 0.75464 | 0.73051 | 1.36974 | 1.95508 | 0.45974 | 0.15567 | 0.6659 |
| 493 | Poa annua L. | Poa annua | 0.4 | 0.8 | 0.013 | 7.2 | 5.5 | 4.5 | 6 | 7.6 | 0.3 | 0.72905 | 0.70188 | 1.35981 | 1.84158 | 0.57912 | 0.20081 | 0.60725 |
| 493 | Capsella bursa-pastoris Medik. | Capsella bursa-pastoris | 0.25 | 0.8 | 0.102 | 7.4 | 5.3 | NA | 6 | 7.7 | 0.8 | 0.65201 | 0.63791 | 1.54898 | 2.03286 | 0.77966 | 0.37419 | 0.45721 |
| 494 | Taraxacum officinale F.H.Wigg. | Taraxacum officinale agg. | 0.3 | 0.6 | 0.013 | 7.4 | 5.2 | 4.7 | 6.1 | NA | 0 | 0.77406 | 0.76726 | 1.79113 | 2.07279 | 0.64778 | 0.2119 | 0.65925 |
| 494 | Poa annua L. | Poa annua | 0.3 | 0.6 | 0.020 | 7.3 | 5.4 | 4.1 | 7.6 | 4.2 | 0.4 | 0.58728 | 0.56055 | 1.34601 | 1.94534 | 0.82357 | 0.23743 | 0.34603 |
| 495 | Capsella bursa-pastoris Medik. | Capsella bursa-pastoris | 0.15 | 0.63 | 0.011 | 8.3 | 5.2 | 3.5 | 6.7 | 4 | 0.1 | 0.59253 | 0.57367 | 1.48069 | 1.91606 | 0.44486 | 0.27236 | 0.42509 |
| 495 | Poa annua L. | Poa annua | 0.16 | 0.63 | 0.020 | NA | NA | NA | NA | NA | NA | NA | NA | NA | NA | NA | NA | NA |

| | | | | | | | | | | | | | | | | | | |
|---|---|---|---|---|---|---|---|---|---|---|---|---|---|---|---|---|---|---|
| 495 | Taraxacum officinale F.H.Wigg. | Taraxacum officinale agg. | 0.2 | 0.63 | 0.015 | NA | NA | NA | NA | NA | NA | NA | NA | NA | NA | NA | NA | NA |
| 495 | Hypochaeris radicata L. | Hypochoeris radicata | 0.1 | 0.63 | 0.110 | NA | NA | NA | NA | NA | NA | NA | NA | NA | NA | NA | NA | NA |
| 495 | Sonchus oleraceus L. | Sonchus oleraceus | 0.02 | 0.63 | 0.025 | NA | NA | NA | NA | NA | NA | NA | NA | NA | NA | NA | NA | NA |

**TABLE S7: Soil variables measured from topsoil samples (0-4.5 cm) collected in April 2022.** Soil depth was measured separately in July 2023.

| Variable | Definition | Details |
|---|---|---|
| Core volume (Core_Vol) | Volume of sampling core (4.5 cm height, 5 cm diameter; $cm^3$); not used as environmental parameter | V = h * Pi * r² |
| Volume sampled | Volume of soil sampled at specific locations (hexagonal area, 6 sides; 3.5 × 6.5 × 4 cm; $cm^3$); not used as environmental parameter | V = 127.28 |
| Total fresh weight (Tot_FW) | Fresh weight of soil sample including skeleton fraction (g) | |
| Total fresh volume (Tot_Fsoil _Vol) | Total volume of fresh soil including skeleton fraction ($cm^3$) | nb_core * Core_Vol |
| Skeleton weight (SkW) | Mass of coarse fraction retained on 2 mm sieve (g) | |
| Skeleton volume (Sk_Vol) | Volume of soil skeletal fraction (particles > 2mm) estimated from quartz density (2.65 g $cm^{-3}$) | SkW / 2.65 |
| Fresh weight (FW) | Fresh weight of soil excluding skeleton fraction (g) | tot_FW – SkW |
| Fresh volume of soil (Fsoil_Vol) | Volume of fresh soil excluding skeleton fraction ($cm^3$) | tot_Fsoil_Vol – Sk_Vol |
| Total Dry weight_55 (DW_55_1/3) | Dry weight (55°C, 48 h) of subsample including skeleton fraction (g) | |
| Dry weight 55 (DW_55) | Dry weight (55°C, 48 h) of soil excluding skeleton fraction (g) | |
| Dry weight_105 (DW_105_1/3) | Dry weight (105°C, 48 h) of subsample excluding skeleton fraction (g) | |
| Dry weight_105 (DW_105) | Estimated oven-dry weight (105°C) of the total fresh soil sample (g) | DW_55 * (DW_105_1/3) / DW_55_1/3 |
| Soil water content (Soil_WC) | Gravimetric soil water content calculated using DW_105 | (FW – DW_105) / FW |
| Bulk density | Dry soil mass per unit fresh soil volume (g $cm^{-3}$); estimated from DW_105 and fresh soil volume excluding skeleton fraction | DW_105 / Fsoil_Vol |
| Effective soil volume | Soil volume available for root growth ($cm^3$); calculated differently for wall habitats due to discontinuous substrate availability | (Fsoil_Vol * Soil_depth) / Soil_surface; Walls: Soil_depth * Soil_surface |
| pH | pH value | Measured in water |
| C | Soil carbon content | High-temperature combustion (CNS analyser) |

| | | |
|---|---|---|
| N | Soil nitrogen content | High-temperature combustion (CNS analyser) |
| C/N | Ratio of soil carbon content and soil nitrogen content | |
| $Cl^-$ | Soluble chloride in soil (nmol $mg^{-1}$) | Ion chromatography |
| $F^-$ | Soluble fluoride in soil (nmol $mg^{-1}$) | Ion chromatography |
| $PO_4^{3-}$ | Soluble phosphate in soil (nmol $mg^{-1}$) | Ion chromatography |
| $NO_3^-$ | Nitrate in soil (nmol mg-1) | Ion chromatography |
| $SO_4^{2-}$ | Soluble sulfate in soil (nmol mg-1) | Ion chromatography |

**TABLE S8: Climatic variables measured at urban habitat patches. For temperature and humidity,** hygrochron data loggers (iButton, Whitewater, USA) were buried in soil (ca. 4 cm depth) at selected sites (n = 33)**.**

| Factors | Definition | Variables | Measured |
|---|---|---|---|
| Shading | Reduction in light availability due to tree canopy cover or building shadows (i.e. light not reaching the *A. thaliana* plants) | 1 | April 2022 |
| Light intensity | Light reaching the *A. thaliana* plants (inverse of shading) | 1 | April 2022 |
| Monthly maximum temperature | Mean of daily maximum temperatures per month | 13 | Sep 2021–Sep 2022 (monthly) |
| Monthly mean humidity | Mean relative humidity per month | 13 | Sep 2021–Sep 2022 (monthly) |
| Monthly mean temperature | Mean temperature per month | 13 | Sep 2021–Sep 2022 (monthly) |
| Monthly minimum temperature | Mean of daily minimum temperatures per month | 13 | Sep 2021–Sep 2022 (monthly) |
| Degree days | Thermal energy accumulation based on daily minimum and maximum temperatures; commonly used as a proxy for phenological development | 13 | Sep 2021–Sep 2022 (monthly) |
